# Macromolecular refinement of X-ray and cryo-electron microscopy structures with Phenix / OPLS3e for improved structure and ligand quality

**DOI:** 10.1101/2020.07.10.198093

**Authors:** Gydo C.P. van Zundert, Nigel W. Moriarty, Oleg V. Sobolev, Paul D. Adams, Kenneth W. Borrelli

**Affiliations:** Schrödinger, New York, New York, United States; Molecular Biosciences and Integrated Bioimaging, Lawrence Berkeley National Laboratory, Berkeley, CA 94720, USA; Department of Bioengineering, University of California at Berkeley, Berkeley, CA 94720, USA

## Abstract

Accurate macromolecular structure refinement is of paramount importance in structure based drug discovery as it provides a gateway to using ligand binding free energy calculations and ligand docking techniques. When dealing with high-resolution data, a simple restraint model may be preferred when the data is able to guide atom parameters to an unambiguous location. However, at lower resolution, the additional information contained in a complex force field may aid in refinement by avoiding implausible structures permitted by the simpler restraints. With the advent of the resolution revolution in cryo-electron microscopy, low resolution refinement is common, and likewise increases the need for a reliable force field. Here we report on the incorporation of the OPLS3e force field with the VSGB2.1 solvation model in the widely used structure determination package Phenix. The implementation is versatile and can be used in both reciprocal and real space refinement, alleviating the need for manually creating accurate ligand restraint dictionaries in the form of CIF files. Our results show significantly improved structure quality at lower resolution for X-ray refinement with reduced ligand strain, while showing only a slight increase in R_free_. For real space refinement of cryo-EM based structures, we find comparable quality structures, goodness-of-fit and reduced ligand strain. In addition, we explicitly show how structure quality is related to the map-model cross correlation as a function of data weight, and how it can be an insightful tool for detecting both over- and underfitting, especially when coupled with ligand energies. Further, we have compiled a user-friendly start-to-end script for refining structures with Phenix/OPLS3e, which is available starting with the Schrödinger 2020-3 distribution.

## Introduction

X-ray crystallography has been the main source of structural information accounting for about 90% of all entries in the PDB and has been the backbone of structural biology for the last decades. With the advent of the “resolution revolution” due to improved hardware and software, cryo-electron microscopy (cryo-EM) has become a viable approach for investigating larger molecular complexes and membrane proteins, potentially vastly expanding new drug targets amenable to structure based drug design (SBDD). Many SBDD methods require atomic accuracy for at least all of the heavy atoms. Unfortunately, the models fit to many X-ray crystallography datasets and virtually all cryo-EM datasets are hampered by low resolution, and contain some ambiguities at this scale.

Due to the high parameter to data ratio found in macromolecular refinement, more so at lower resolutions, knowledge based restraints are introduced to reduce overfitting, the most simple form being bond lengths and angles. At lower resolution, additional restraints can be included, such as secondary structure (Head et al. 2012), and deformable elastic network restraints (Schröder et al., 2014). The former is currently routinely used during cryo-EM structure refinement and low-resolution X-ray crystallography. Accurate restraints are thus paramount for obtaining physically realistic structures and ligands when data is relatively sparse. The problem is exacerbated in cryo-EM as no robust validation metric is generally agreed upon to detect overfitting, even though there is considerable concern and several methods have been proposed (DiMaio et al., 2013; Falkner and Schröder, 2013; Lagerstedt et al, 2020; Volkmann, 2009). This is in contrast to crystallographic refinement where the concept of R_free_ is universally used (Brunger, 1992), though the precise practicalities are not on a firm theoretical footing (Tickle et al., 1998).

Knowledge based structural restraints for the standard amino-acid residues are well established from the wealth of high-resolution data that is available in the Protein Data Bank (wwPDB consortium, 2019) and small molecule databases (Groom et al., 2016). The Engh and Huber equilibrium bond lengths and angles (Engh and Huber, 1991, 2001) were used for several decades, but have been superseded by conformation dependent restraint libraries (Moriarty et al., 2016; Tronrud et al., 2010) for the backbone atoms. Generating restraints for small molecules, however, is significantly harder, due to the vastly increased chemical space that they can occupy and limited available data; in conjunction with the observation that ligand densities are typically less resolved compared to their surroundings, either due to a superposition of states, partial occupancy, increased mobility, or any combination of these, this has led to a number of critical publications addressing the quality of ligands deposited in the PDB (Deller and Rupp, 2015; Liebeschuetz et al, 2012; Peach et al., 2017; Reynolds, 2014, Sitzmann et al., 2012). The wwPDB formed a working group to define proper ligand validation protocols to combat these issues, with several recommendations already being implemented (Adams et al., 2016).

Knowledge based restraints for ligands are usually provided through restraint dictionaries in CIF format, derived from high-resolution structure models extracted from small molecule crystallography databases, or quantum mechanical calculations, with several computer programs having been developed to automate the process (Steiner and Tucker, 2017). Large libraries containing restraints for monomers present in the PDB such as the REFMAC5 monomer library (Vagin et al., 2004) are extensively used in REFMAC (Kovalevskiy et al., 2018) and Phenix (Liebschner et. al., 2019). In addition, the latter uses the GeoStd library (http://sourceforge.net/projects/geostd) that is curated for accurate amino acid restraints. The restraints comprise the above mentioned bond length and angle restraints, but also include torsion angle, chirality, and planarity restraints, usually combined with a repulsive interaction energy term to prevent serious atomic clashes. However, for more complicated molecules such as macrocyclic peptides, generating an accurate restraint dictionary can still be a tedious and time consuming process. Furthermore, electrostatics and attractive Van der Waals forces are typically not taken into account, though it has been shown that electrostatics is important for modeling the hydration of DNA (Fenn et al., 2011), while including an implicit solvent improves stereochemistry for solvent accessible residues (Moulinier, 2003). Moreover, since cryo-EM results in Coulomb potential maps, the inclusion of electrostatics during refinement will ultimately be important to accurately represent the experimental data (Wang and Moore, 2017).

More sophisticated approaches introduce a physics-based force field or low-level quantum mechanics Hamiltonian to calculate energies and gradients during structure refinement. The use of a force field in refinement has already been introduced several decades ago in the well-known Xplor and CNS programs using the OPLS-AA parameters (Brunger et al., 1998). Also, the Schrodinger developed PrimeX refinement protocol uses the Prime force field, a combination of the OPLS3 and VSGB2.0 energy model, and we have shown that it results in significantly improved structures based on standard quality measures such as the number of clashes (Bell et al., 2012; Jianing et al., 2011). The Amber force field (Ponder and Case, 2003) is available in Phenix and a recent analysis likewise showed that structure quality is improved, more so at lower resolutions (Moriarty et al., 2020). The Q|R package applies QM to the entire protein (reciprocal or real space) improving geometries and hydrogen bonding networks (Wang et al. 2020; Zheng et al., 2017, 2020). A number of publications focused particularly on ligand geometries in reciprocal space refinement and are mainly effective in relieving ligand strain: Phenix-AFITT supplements the Phenix energy model with the MMFF-94 force field for ligands only (Janowski et al., 2009); Phenix-DIVCON takes a different approach by applying the AM1 quantum mechanical Hamiltonian (Borbulevych et al., 2014, Borbulevych 2016).

The recently released OPLS3e force field combines accurate partial charges from on-the-fly semi-empirical quantum mechanics calculations with accurate torsional profiles from a combination of a database covering a substantial portion of medicinal chemistry space and the ability to easily extend that database with high-level quantum mechanics calculations when encountering functional groups not already covered (Roos et al., 2019). Here we introduce the implementation of an interface between the principal refinement programs in the widely used macromolecular structure determination package Phenix and the OPLS3e force field and VSGB2.1 implicit solvation model (Li et al., 2011) through Schrödinger’s Prime software, which we refer to as Phenix/OPLS3e. The implementation is versatile supporting covalently bound ligands and multiconformer complexes, and can be used in reciprocal and real space refinement for either the whole or part of the structure, using `phenix.refine` (Afonine et al., 2012) and `phenix.real_space_refine` (Afonine et al., 2012; Afonine et al., 2018). It furthermore removes the need for accurate restraints files as ligands are automatically parameterized by the OPLS3e force field, either through its internal database or generated by the ForceFieldBuilder available in Maestro. We benchmarked our approach on 2284 cases using X-ray data and show that refinement with Phenix/OPLS3e results in structures with improved Molprobity scores and reduced whole structure and ligand strain energies. Larger improvements are seen at lower resolutions, with models showing a small increase in their R_free_ values. Interestingly, when we applied our method to 15 cryo-EM structural models, this resulted in models with similar MolProbity scores and real space cross correlation values but reduced ligand strain. We furthermore show that the inclusion of a high quality ligand force field provides an additional measure to reduce overfitting by constructing a ligand strain versus cross correlation curve by sampling over weight space through multiple refinements. Our analysis highlights some instances where structure quality and goodness-of-fit could be improved in current deposited PDB structures, though in all cases modelers have been wisely conservative in their refinement approaches.

The Phenix/OPLS3e implementation has already seen applications in cryo-EM enabled SBDD of macrocycles in ribosomes (Qi et al., 2019) and determining high-confidence ligand binding poses in ambiguous cryo-EM structures in the GemSpot pipeline (Robertson et al., 2020). We foresee our approach as an accessible and user-friendly implementation of a high quality force field to improve structure and ligand quality especially when using lower resolution data provided either by crystallography and cryo-EM, and as an additional measure for reducing overfitting in cryo-EM structure refinement and cryo-EM ligand fitting.

## Methods

### Overview

Macromolecular refinement with Phenix in general consists of three stages: an initial stage where all the input data is processed and prepared; the actual refinement stage which involves the concept of the macrocycle, an iterative sequence of distinct procedures that optimize different aspects of the model; and an end stage where the refinement is finalized and the structure and any other outputs are written to disk. An important procedure in a macrocycle is the optimization of the atom coordinates, which ultimately is a fine balancing act between the available experimental data and knowledge-based restraints. The target function *T* during coordinate optimization can be written as a simple linear function consisting of two terms

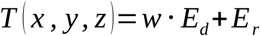

Where *E*_*r*_ and *E*_*d*_ are the (pseudo-)energies of the restraints and fit to the data respectively; *w* is a weight factor determining the impact of the restraints, and is typically determined automatically during refinement, using differing algorithms depending on the refinement engine, i.e. reciprocal space (`phenix.refine`) or real space refinement `(phenix.real_space_refine`). In the reciprocal space refinement engine, the total weight factor is the product of two terms, where the first term is determined by normalizing the restraints gradients against the data fit gradients, after performing a local minimization and a short molecular dynamics simulation (Adams et al., 1997); the second weight term is by default set to 0.5 or can be determined using a grid search that aims to optimize the metrics for model to data fit, while simultaneously maintaining overall geometric quality within a reasonable range of values. In real space refinement, when no NCS restraints are present, the weight factor is determined by refining small pieces of the structure against the density at different weights. The final weight is chosen such that a reasonable stereochemistry is maintained. In the case of NCS restraints the whole structure is refined instead of random pieces (Afonine et al., 2018).

The restraints energy *E*_*r*_ in turn can again be separated into a linear combination of terms given by

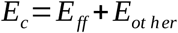

Where *E*_*ff*_ is the force field energy consisting of a linear combination of bond length, bond and torsion angle, planarity, chirality, parallelity, and nonbonded restraint terms. In the default Phenix force field the nonbonded restraints consist of solely a repulsive term to prevent atom clashes. The second *E*_*ot her*_ term is a collection of additional restraints such as non-crystallographic symmetry (NCS) restraints, deformable elastic network (DEN) restraints and others. Our aim is to use the energies and forces provided by the OPLS3e force field and VSGB2.1 energy model and exchange it for the default force field energy term *E*_*ff*_, while still allowing the use of restraints included in *E*_*ot her*_.

### Implementation details of the OPLS3e/VSGB force field in Phenix

Since the Schrödinger software stack is written primarily in Python3, while Phenix currently supports Python-2.7, this makes a direct connection between the software entities challenging. Thus, we decided to create an external program that runs in the background during a refinement using either the `phenix.refine` or `phenix.real_space_refine` engines. This acts as an energy server, running parallel to the refinement process. The external process is initialized using the starting structure in Maestro format and additional options that impact the energy calculation, such as including crystal symmetry or energy flags to include or exclude certain terms. During all subsequent energy calculations, the refinement protocol writes coordinates to disk that are then read by the external server. The external energy server internally updates its coordinates and calculates the energy and gradients of the system, which are then written to disk and read by the refinement program. In the case of a multiconformer model, the energy and gradients are calculated for each consistent conformer, after which the energies and gradients are summed and divided by the number of conformers. The concept of a consistent conformer simply entails that atoms carrying a certain alternate location ID interact only with atoms with no or the same alternate location ID.

Our implementation is versatile and allows for inclusion of the full range of Phenix energy terms, such as NCS and DEN restraints, alongside the OPLS3e energy terms. In addition, the force field can be used for only a subset of the structure, by providing a selection, such as a ligand and its surrounding residues. In this mode, atoms governed by the OPLS3e force field will be fully aware of its environment, while energies and gradients are calculated appropriately for the subset. A more restricted version applies the force field to a ligand that is decoupled from its environment, i.e. interaction energies with the macromolecule are not included, similar to the Phenix-AFFIT plugin. This is useful when higher resolution data is available for the macromolecule while the ligand densities remain ambiguous, or for larger complexes found typically in cryo-EM where refinement with the full force field can be lengthy. Crystal symmetry, which by default is not used, needs to be explicitly accounted for in the OPLS3e force field energy calculations when using crystallographic data by using the `schrodinger.use_symmetry=True` parameter. After obtaining energies and gradients, the default Phenix energy and gradient terms or the relevant selected atoms (default is all) are subtracted from the total energy and gradients, while the OPLS3e/VSGB force field energies and gradients are added with a weighting term (default is 10). In addition, the riding hydrogen model in the version of Phenix (1.16) used in this work was disabled for the force field selected part, as this increased the energy in the OPLS3e force field calculations. t should furthermore be noted that chemical components need to be consistent with the OPLS3e force field, meaning that the valency rules of atoms need to be adhered to, i.e. hydrogens need to be added to the system. Truncated residues are supported, though requires the addition of non-standard hydrogens.

### Performing a refinement with Phenix/OPLS3e

Initially the structure to be refined needs to be processed with Schrödinger’s `Protein Preparation Wizard`, which adds hydrogens and additional bonds between residues and ligands, among other things, so the structure is fully compliant with the OPLS3e force field specification. Although performing a refinement with the force field takes care of all structural restraints, restraint dictionaries describing the topology of ligands still need to be provided to Phenix for restrained atomic displacement (ADP) refinement and to prevent errors due to unknown atom types. Restraint file generation is provided by the `hetgrp_ffgen` utility, an internally used atom typing program, which generates restraints based on the OPLS2005 force field. After processing the input structure and generating the required restraint files, the structure can be refined through the command line. The OPLS3e force field can be requested by setting the parameters `schrodinger.use_schrodinger=True` and `schrodinger.maestro_file=<STRUCT_FN>` where `<STRUCT_FN>` is the processed input file in Maestro format.

The pipeline can be cumbersome to go through manually, as atom naming and residue naming conventions in Phenix are different compared to Schrödinger’s approach, thus there is a consolidated single entry point command line script within the Schrödinger distribution to provide a user-friendly experience. The minimal input solely consists of a PDB or Maestro structure file and a data file and in case of real space refinement the map resolution. The `phenix.refine` engine is used when a file containing reflections in either MTZ or mmCIF format is provided, and `phenix.real_space_refine` when a map in CCP4 or MRC format file is provided. Currently providing a real space map for reciprical space refinement is not supported, but is under developemnt for the next release. The calling signature is condensed to the following

~~~
$SCHRODINGER/run -FROM psp phenix.py <STRUCT_FN> <EXP_DATA_FN> [-resolution
<RESOLUTION>] [options]
~~~

The resolution option providing the cryo-EM map resolution is required solely for real space refinement. All Phenix refinement parameters can be provided through the command via key-value pairs, or by providing an input parameter file, which is directly passed on to the refinement engine. Before and after refining the structure, validation scripts are run including `phenix.molprobity` to calculate geometry statistics and R-factors, `phenix.real_space_correlation` for real space correlations on a per residue basis when dealing with reciprocal space refinement and `phenix.model_map_cc` for real space refinement plus total and per-atom based energies are calculated with Prime. All calculated energies and metrics are stored in the output Maestro structure file and are viewable in the Worktable in Maestro and are accessible as structure level and atom level properties. The user-friendly script is available in the Schrödinger 2020-3, and works ‘out of the box’ when both Phenix and the Schrödinger suite are installed and users have set the `SCHRODINGER` and `PHENIX` environment variables to the root directories of the Schrödinger and Phenix distributions.

### Benchmarking the impact of the OPLS3e force field on macromolecular refinement

The Phenix/OPLS3e protocol was benchmarked on a diverse dataset of 2284 protein-ligand cases. The 2284 cases were selected based on the following criteria: the macromolecule is only protein; the sequences contain less than 30% sequence identity among entries; and the entry should have at least 1 ligand according to Schrödinger’s LigandFinder (total of 4928 ligands). To ascertain the impact of the Schrödinger force field, the cases were divided in 6 different resolution bins with a width of 0.5 starting at 1.0Å and ending at 3.5Å resolution, where each bin contains 11, 863, 823, 433, 56 and 98 number of structures, respectively. Starting structures were downloaded from the PDB database with their associated reflection files in MTZ format. Each structure was prepared using Schrödinger’s `Protein Preparation Wizard (PPW)` tool to add and optimize hydrogen positions (heavy atoms positions were fixed), and to define bonds and linkages between residues, e.g. disulfide bridges. The output structure is further processed by renaming residues and atom names to be consistent with the naming conventions in Phenix. Waters near special positions were removed for Phenix/OPLS3e as these resulted in non-bonded overlaps in the current Schrödinger energy calculation routines. Input restraint files were generated using `hetgrp_ffgen`. Reciprocal space refinement was performed using `phenix.refine` using as input the PDB and Maestro file describing the structure, the PDB MTZ file, and generated restraint files as input files with the following refinement options:

~~~
main.number_of_macrocycles=5, optimize_xyz_weight=True,
optimize_adp_weight=True,
strategy=individual_sites+individual_adp+occupancies
weight_selection_criteria.bonds_rmsd=0.020
weight_selection_criteria.angles_rmsd=2.5
schrodinger.use_schrodinger=True schrodinger.use_symmetry=True
~~~

`schrodinger.maestro_file=<MAESTRO_FILE>,` where `<MAESTRO_FILE>` is the input structure in Maestro format. The macrocycle thus follows the iterative optimization scheme: 1) locally optimize atom positions; 2) optimize individual ADPs; and 3) optimize the atom occupancies atom positions for residues with alternate conformations or atoms with occupancies less than 1 but greater than 0. The default `individual_sites_real_space` strategy component was disabled, as this typically raised the energy dramatically, presumably due to rotamer idealization. The weight selection criteria for the automatic restraints weight determination were increased compared to default values for refinement at lower resolution, as shown in previous work (Bell et al., 2012). R_free_ flags were generated for the downloaded MTZ files, using `phenix.reflection_file_converter` using default values. For the reference Phenix workflow, which we will refer to as standard Phenix, the input structure and restraint files were prepared using `phenix.ready_set` with the option `add_h_to_water=True`. The output model, ligand restraint, and metal/link `edit` files were used as input to `phenix.refine` using the same options as described above, except for the Schrödinger specific options and the weight selection criteria were left at their default values. All re-refined output models were analyzed using `phenix.molprobity` and `phenix.real_space_correlation`, and energies were calculated using Prime. For `phenix.molprobity` we used the following options `keep_hydrogens=True.`Calculations were performed using Phenix-1.16.

### Re-refinement of high-resolution cryo-EM models

To determine the impact of the OPLS3e/VSGB force field in cryo-EM structure refinement, 15 protein-ligand complexes were retrieved from the PDB and EMDB containing pharmaceutically active compounds, a subset of cases we used in our recent GemSpot pipeline (Robertson et al., 2020). Structures were manually prepared using the Protein Preparation Wizard in Maestro. For all cases default options were used in the Preprocessing step, i.e. bond orders were assigned, hydrogens were added, metal zero-order bonds created and ligand protonation states determined. Since in some cases a substantial part of the structure was missing side chains, they were added using Prime for all cases. Protein protonation states and hydrogen bond networks were optimized and a restrained energy minimization of hydrogens only was performed. For 6NR2 (EMD-0487) and 6NR3 (EMD-0488) large stretches of unknown (UNK) residues were removed. The refinement pipeline is similar as described above for reciprocal space refinement, with the following differences: cross-correlations were calculated with `phenix.model_map_cc`; Ramachandran Z-scores were calculated with `phenix.rama_z;` and `phenix.real_space_refine` was used as refinement engine with different input parameters: the refinement strategy was set to only the global minimization and ADP optimization protocol, the weight factor of the experimental data in the refinement target function and the exclusion of crystallographic symmetry (*refinement.run=minimization_global+adp, weight=<WEIGHT>* and *schrodinger.use_symmetry=False*). Since cryo-EM modeling currently lacks a proper broadly accepted cross-validation term, determining an acceptable weight factor in the target function between the restraints and the experimental data is not straightforward. Default `phenix.real_space_refine` behavior is to perform a guided weight grid search, where pieces of the model are refined at different weights and monitoring the bond length and bond angle RMSD. Default values are set to 0.1Å and 1.0° angle deviation from ideality, which is too tight for the OPLS3e force field (Bell et al., 2012). Therefore, we performed an explicit weight scan by performing a full refinement at different weights. For comparison we also re-refined the structures using solely Phenix tools as described above and identical refinement options. Calculations were performed using Phenix-1.16, except for `phenix.rama_z` for which Phenix-1.18 was used.

### Limitations of current implementation

Our current implementation is lacking support for automatic water and ion placement and simulated annealing in torsion angle space. In addition, multiprocessor use for weight scanning within the refinement protocols is not available. Currently, only two conformers can be handled, as the Protein Preparation Wizard only takes into account the A and B conformer when adding hydrogens. Finally, as mentioned above, residues on special positions need to be removed to prevent high energies caused by clashes within the OPLS3e force field.

## Results and discussion

### Structure quality improvement increases with decreasing resolution

To measure the impact of the OPLS3e/VSGB force field on refinement, we applied Phenix/OPLS3e and standard Phenix reciprocal space refinement on 2284 protein ligand complexes at varying X-ray diffraction resolutions as described above. For each of the resolution bins we calculated the difference in R-factors and MolProbity validation metrics between the two refinement protocols by subtracting found values for the Phenix/OPLS3e refined structures from the standard Phenix refined structures for each case (Figure1a-b). The absolute values for all metrics for both protocols can be found in Figure S1 and Table S1.

**Figure 1.**
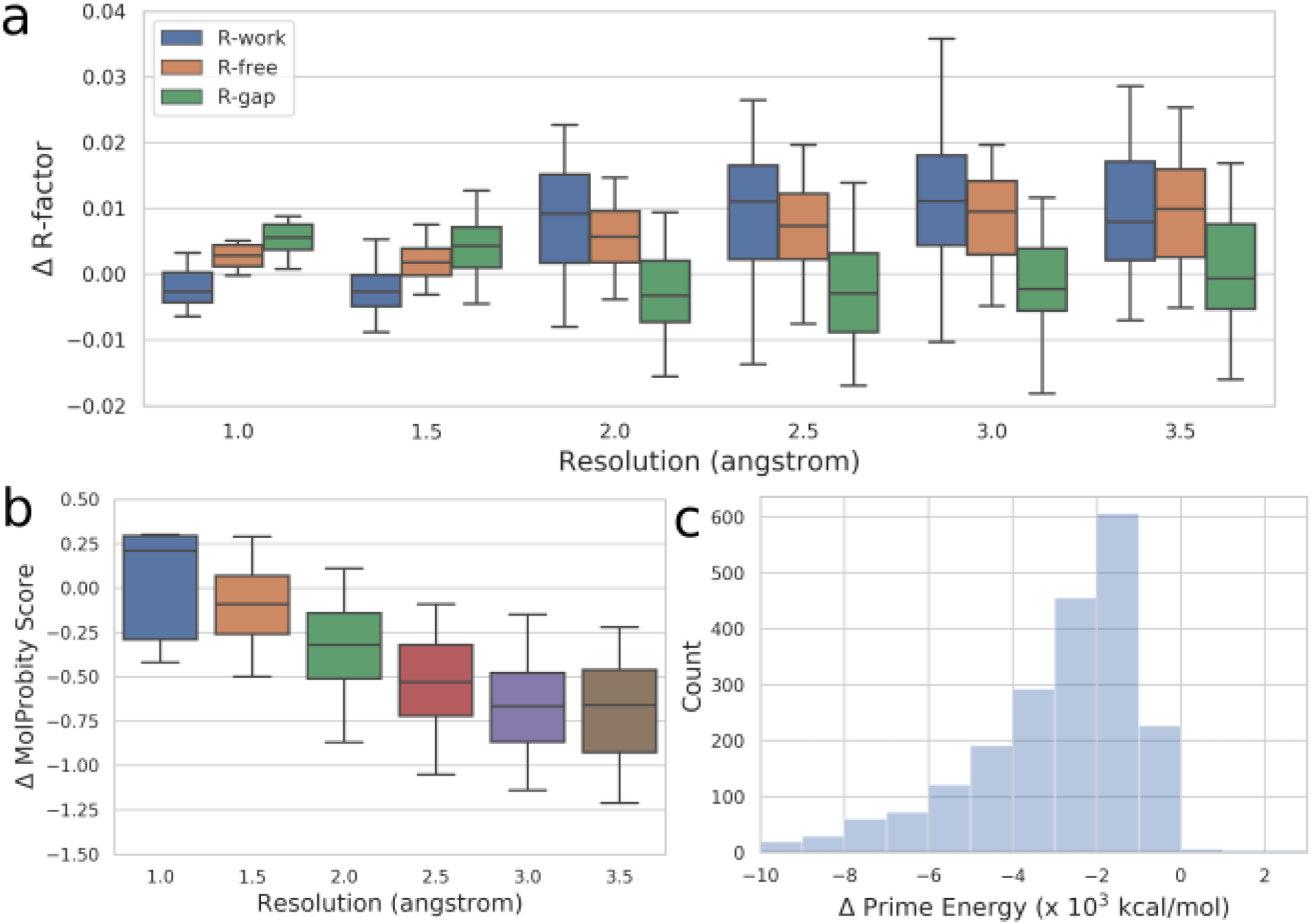
Global structure refinement metric differences. Shown are the difference in R_work_, R_free_ and R_gap_ (a) and MolProbity score (b) distributions between standard Phenix refinement and Phenix/OPLS3e refinement. Differences are calculated by subtracting Phenix refined structure metrics from Phenix/OPLS3e refined structure metrics. The whiskers in the box plots represent the 95th percentile, the box the 50th percentile and the line in the box represents the median. (c) Energy difference distribution between Phenix/OPLS3e refined structure and starting structure.

In general there is a slight increase in the median ΔRR_free_ value, ranging from 0.003 (0.3%) in the 1.0-1.5Å resolution bin to 0.010 at the 3.5-4.0Å range for models re-refined starting from the initial PDB structure. For ΔRR_gap_ we observe a different behavior, with increased values of 0.006 at high resolution and decreased values for lower resolution refinement starting at 2.0Å. The MolProbity score increases by 0.21 for high resolution refinements (1.0-1.5Å), but improves at lower resolutions with a small improvement of 0.09 at 1.5-2.0Å and 0.32-0.66 improvement at lower resolutions. As the MolProbity score is a combination of several factors, we also inspected other geometric metrics (see Figure S1). All terms except for C-beta deviations have improved median values for structures at resolutions lower than 1.5Å. Unsurprisingly, the structure energy is markedly reduced after refinement with inclusion of the OPLS3e energies indicating that the improvement in the structure quality is indeed due to the force field (Figure 1c), with the modus found at −1.5 × 10^3^ kcal/mol. As observed previously, the bond length and bond angle RMSD values increase by 0.09Å and 1.3-1.5°, with absolute RMSD values of 0.1Å and 2.3°, which is well in-line with observed values in high-resolution structures (Bell et al., 2012; Jaskolski et al. 2007). The only structure metric that suffers is the number of C-beta outliers, which is typically 0 or 1 in standard Phenix refinements, due to tight restraints. Phenix/OPLS3e typically results in an additional 1-3 C-beta outliers depending on resolution.

The improvement of structure quality in terms of the clash score has already been observed in our previous work describing PrimeX, with similar bond length and bond angle RMSDs found compared to ideality (Bell et al., 2012). Furthermore, during the development of this work and manuscript, Moriarty et al. (2020) reported on the inclusion of the Amber force field in Phenix reciprocal space refinement. The results show similar trends both in R-factors and MolProbity statistics showing increased R_work_ and R_free_ values, with a reduction in the MolProbity score and an increase in the C-beta outliers.

To properly compare the output structures of Phenix/OPLS3e and standard Phenix, it is important to take into account the weight optimization protocol used in `phenix.refine`. The weight optimization scheme is essentially a grid search where several weights are sampled, and the weight is chosen in such a way that certain structure based metrics fall within set cutoffs and within those bounds a specific metric is minimized. For example, for structures with a resolution better than 1.5Å, cutoffs are used for the bond length and angle RMSDs, and the weight resulting in the lowest R_free_ is chosen; this is in contrast to refinement at lower resolution such as between 2.5 and 3.5Å where besides the bond length and angle RMSDs also an acceptable R_gap_ and R_free_ range is compared to the lowest values found among all weights, and the structure containing the lowest clash score is chosen. The acceptable R_gap_ and R_free_ ranges are set to 6% and 1.5% points respectively. Taking these optimization parameters into account, we note that most Phenix/OPLS3e structures fall within the 1.5% R_free_ cutoff (Figure 1a), i.e. the median falls well below 1.5%, while nearly all exhibit an improved Clashscore (see Figure S1), where the 95% percentile whisker is well below zero. Thus, when the default optimization scheme would be provided with both the Phenix/OPLS3e re-refined structure and the standard Phenix structure, it would more often pick the Phenix/OPLS3e structure. The same arguments would hold for the other resolution bins starting from 2.0Å onward. In effect, this could also explain the behaviour between these resolution bins in Figure 1a where the R_work_ and R_free_ range is markedly increased compared to refinement at higher resolution. Obviously, changing the weight selection parameters might change the preferred force field used, as the standard Phenix restraints model seems better at finding lower R-factors, especially at higher resolution.

Next we inspected ligand quality by calculating the reduction in strain energy, i.e. energy before and after refinement, and their real space cross correlation for all 4928 ligand cases. Although the RMSDs between the starting and end conformation is typically smaller than 1Å (Figure 2a), and real space cross correlations remain about the same with a median difference of −0.001 (Figure 2b), in 99% of the cases the strain was markedly reduced (Figure 2c).

**Figure 2.**
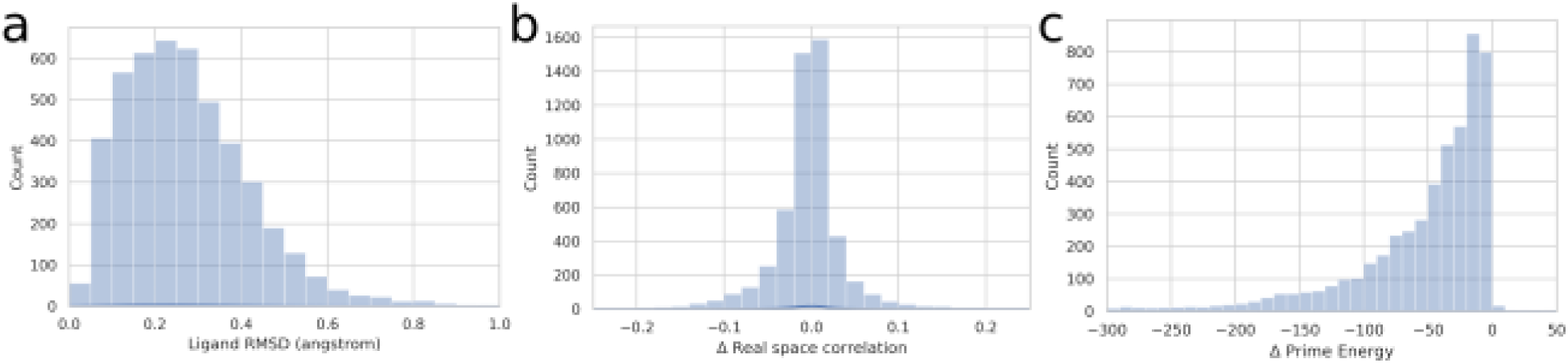
Ligand metrics before and after refinement with Phenix/OPLS3e. Distribution difference plots before and after Phenix/OPLS3e refinement are shown for ligand RMSD (a), real space cross correlation (b), and Prime Energy (c) when starting out with the PDB deposited structure.

In general, our results are in agreement with previous studies for both full structure refinement as well as ligand focused refinement protocols, showing that the impact of including the force field is mainly improving the detail of the models, inherent to the Phenix global minimization strategy employed during refinement (`refine.strategy=individual_sites+individual_adp+occupancies)` which applies a gradient based local minimization, a point that was adequately made recently by Moriarty et al. (2020).

### Re-refining cryo-EM structures with Phenix / OPLS3e

After validating our implementation in reciprocal space refinement using X-ray diffraction data, we applied our protocol on 15 deposited cryo-EM structures and maps, previously used in our GemSpot pipeline (Robertson et al., 2020). Since cryo-EM refinement lacks a robust cross-validation metric, we performed a grid scan over the map weight parameters for our cases, using both the Phenix standard restraints model and the OPLS3e force field. For each refinement, the MolProbity score, and cross correlation values were calculated and are shown in Figure 3 for Phenix/OPLS3e. Individual Clashscores and the Ramachandran z-scores are shown in Figure S2 and S3.

**Figure 3.**
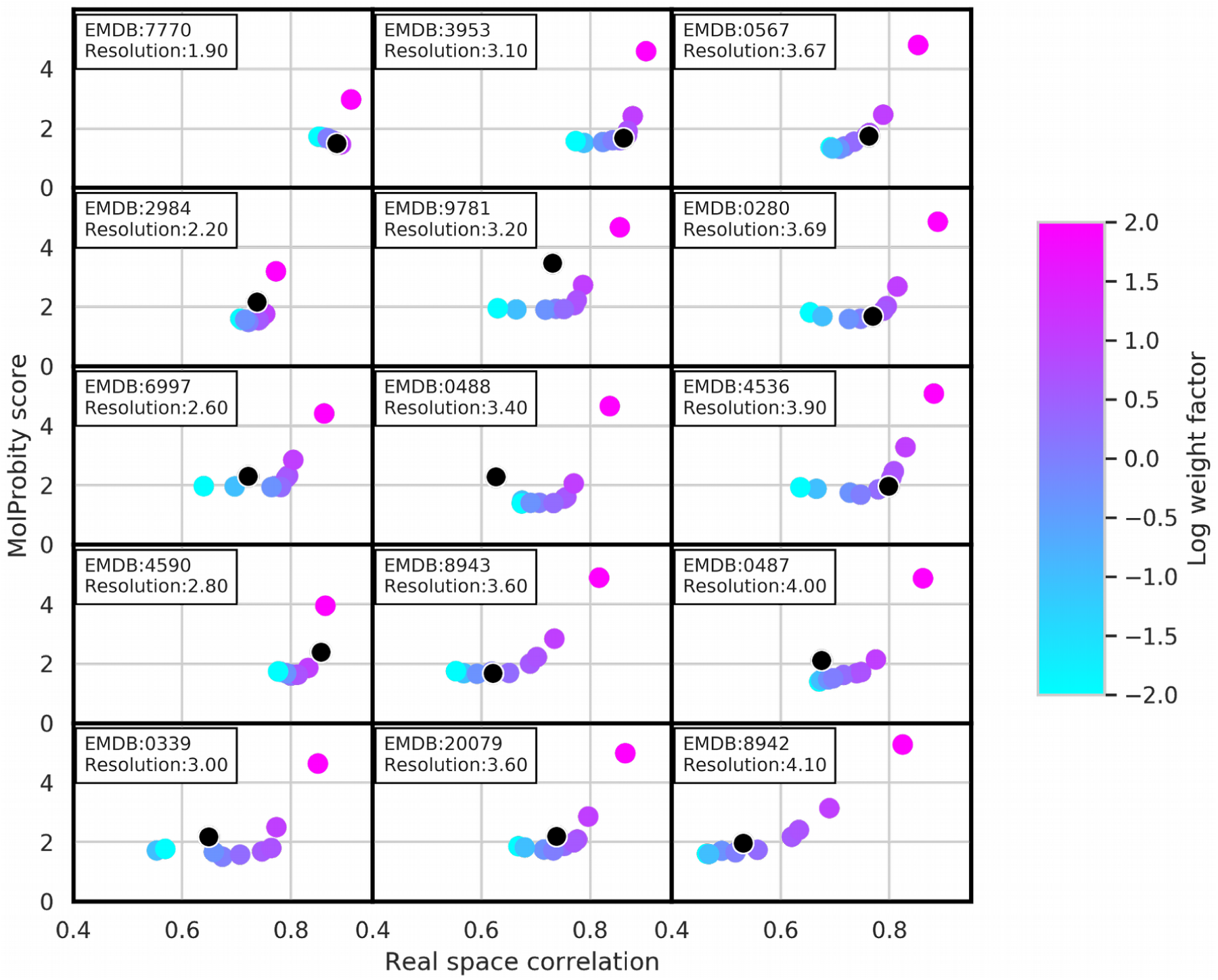
Phenix/OPLS3e real space refinement and the impact of data weight. The MolProbity score versus real space correlation is shown after refinement at different weight factors when using the OPLS3e force field. The black dot represents the deposited structure. The color of each dot represents the weight parameter for the impact of the Coulomb potential map used during refinement with blue indicating lower weights, and pink indicating higher weights.

It is well known that refinement is a balance between restraints and the available data, implying a trade-off between geometric quality and fit to the data. Interestingly, however, we find that although the cross-correlation value between the model and the map typically increases with increasing weight for the experimental data, structure quality proxied either by the MolProbity score remains relatively stable at lower weights, after which the score increases dramatically after a critical weight. Also, when inspecting other proxies, such as the Clashscore and the Ramachandran z-score, a similar behavior is observed (Figure S2 and S3). In cryo-EM modeling it is typically assumed that with equal structure quality the model with higher cross-correlation is preferred, while at equal cross-correlation the model with higher structure quality is preferred. Although there is no current metric that can formally decide how to balance cross-correlation and structure quality, and thus what weight to pick to generate the “most correct” structure, it is clear that there is a reasonable narrow range of acceptable weight values as can be seen in Figure 3, where a noticeable kink is observed in the quality-correlation plots. We therefore propose that providing a single cross correlation value and MolProbity score is not convincing to indicate that the best refined structure is obtained. Moreover, the shape of the quality-correlation curve indicates that the modeler is essentially free to choose the correlation value, reminiscent of earlier X-ray refinement before the advent of R_free_ where exceedingly low R-factors were obtained. However, cryo-EM lacking a cross-validation metric, a minimal check can be performed by staying within the stable structure quality regime. However, we note that in most cases a weight factor of 3 provides a reasonable estimate for the location of the break point.

To further narrow down the best weight, we also investigated the ligand energy against its local map-model correlation as a function of the weight factor (Figure 4). The shape of the curve follows a similar pattern as the MolProbity score with relatively low ligand energies, where the increase in energy is compared to the lowest energy conformer, at low to intermediate weights together with low real space correlations; while at high weights the correlation values and the energies increase. In addition to the knowledge based MolProbity metrics, the ligand energy provides an additional physics based check on overfitting as highly strained ligands are unlikely to be found. Since the resolution of cryo-EM maps is often non-uniform, the weight of the data during refinement could also be varied locally for optimal model to map fitting, even though this has not been investigated so far. Inspecting the ligand energy versus correlation at different weights provides one route to investigating local weight optimization, which is especially important for the binding site, as here accuracy is key. Based on our findings here coupled with the above observation for the global structure metrics, we advise an energy increase of the ligand to be no more than 1 log unit, i.e. 10 kcal / mol, within Phenix/OPLS3e refinement compared to the lowest energy conformer to reduce local overfitting.

**Figure 4.**
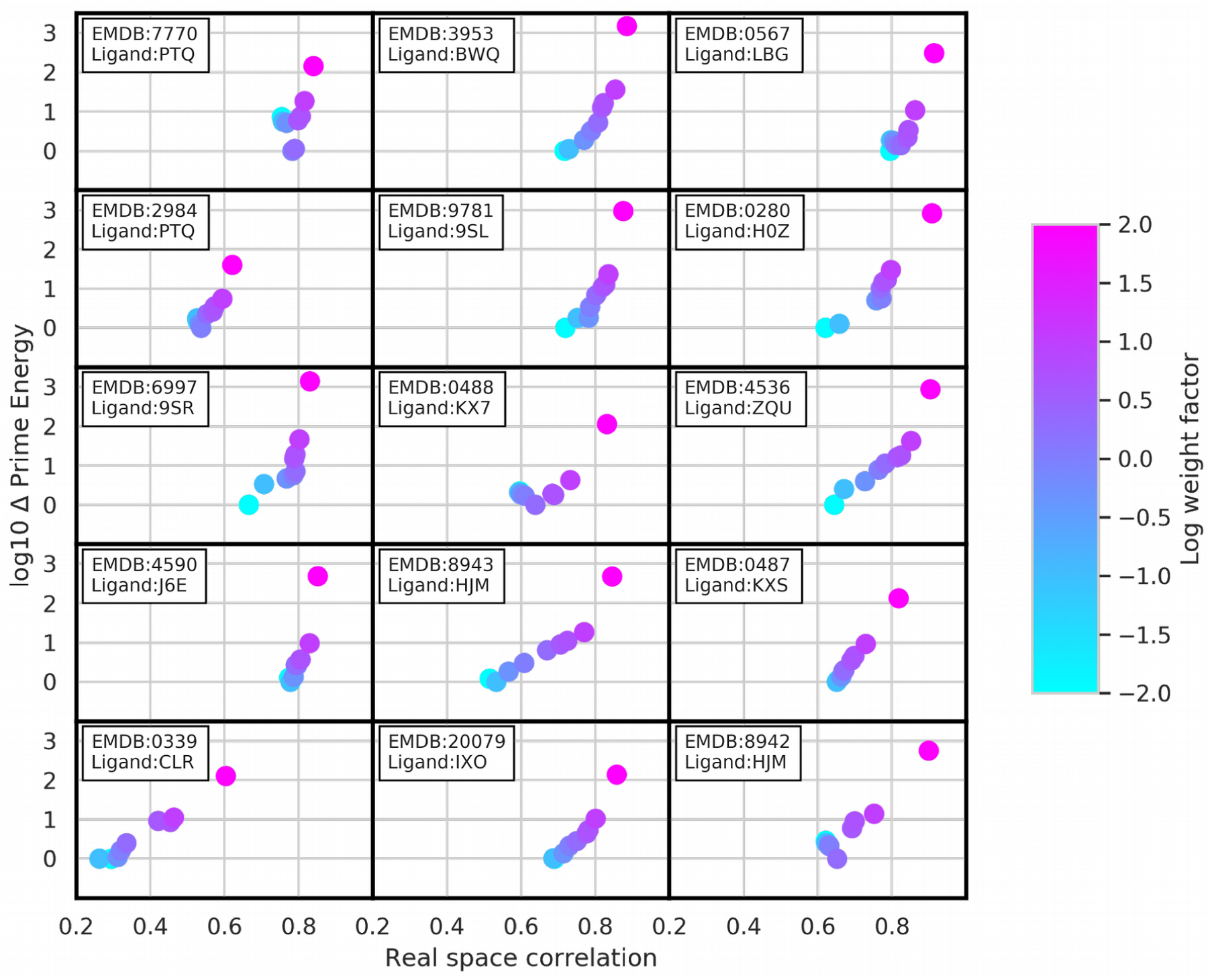
Ligand energies and correlations after Phenix/OPLS3e real space refinement. Energy differences against the lowest energy conformer for 5 ligands, one for each cryo-EM structure, against the cross correlation are shown. The energy of the lowest energy conformer was set to 1 and the energy axis is on a log scale. The color of the dot represents the impact of the Coulomb potential map, with blue tints indicating a low weight and pink tints indicating a high weight.

When comparing the standard Phenix force field and the OPLS3e force field (Figure S4), we find no significant difference in the structure quality-correlation curve between the two, owing to the additional restraints introduced by default in the real space refinement protocol including secondary structure, rotamer, NCS and Ramachandran restraints. Although refinements with Phenix/OPLS3e results in a curve starting at low correlations, refinement with the Phenix force field is more focused, i.e. even at very low weights the correlation remains rather stable as is the MolProbity score, and the curve is less smooth compared to Phenix/OPLS3e refinement but does still show a critical weight term after which structure quality deteriorates rapidly. This might be due to tighter restraints where even low weights on the geometric terms result in large forces at deviations from ideality, but we did not investigate this further.

Having established a narrow acceptable weight range, we can evaluate the current deposited ligand complexes (see black dots in Figure 3). When checking the location of the structure quality versus correlation locations of the deposited structures we find that most modelers have been conservative in their modeling and most structures are in the stable MolProbity --high-cross correlation region. Intriguing, several cases are found very close to the break point between structure quality and real space correlation, e.g. EMD-4536, EMD-0280, and EMD-3953. Other models can be further refined, i.e. both their real space correlation as well as their structure quality can be improved, such as EMD-6997, EMD-0339 and EMD-9781. For 2 cases, EMD-4590 and EMD-0567, the deposited model is found after the break point, and could indicate slight overfitting of the structure.

The procedure to vary the weight factor between data and model has been investigated earlier by Monroe et al. (2017) in the context of MDFF and Rosetta, where they found that most models could be “refined further”, although their dataset was mainly aimed at lower resolution cases down to 20Å. In addition, besides the difference in software platform used, they did not explicitly show how the weight factor relates to the cross correlation or how to decide on the optimal weight factor at high resolution cryo-EM modeling.

Finally, to help with determining a near-optimal weight, we implemented an automatic weight scanning tool that runs several refinements at different weights and produces a plot similar to Figures 3 and 4 showing MolProbity score versus real space correlation as the global metrics and ligand energy versus correlation for local ligand metrics.

## Conclusion

Here we have described our implementation of Phenix/OPLS3e, which incorporates the OPLS3e force field and VSGB2.1 solvation model into the popular refinement package Phenix, for both reciprocal and real space refinement. Our approach alleviates the need to handcraft an accurate restraints dictionary for ligands as gradients are provided by the force field and includes additional physics based forces, most notably electrostatics and a GBSA implicit solvent model. More specifically, for reciprocal space refinement a clear improvement in structure quality is observed starting at resolutions worse than 1.5Å, while R_free_ is generally slightly increased compared to the standard Phenix restraints model, results that are in line with previous observations (Bell et al, 2012; Moriarty et al., 2020). For real space refinement of cryo-EM structures, no significant improvement is noticed in refinement when comparing the structure quality-correlation plots between the two restraints models by scanning across weights, though global and ligand energies are significantly reduced when using Phenix/OPLS3e. We furthermore have shown that, although cryo-EM currently has no universally used cross validation metric, and thus no formal target to determine the weight ratio for data versus restraints, there is actually only a relatively narrow weight range where structure quality is acceptable while correlation is maximized. Insights into local overfitting can be obtained by inspecting ligand strain compared to the low energy conformer. The nature of the quality-correlation curve implies that a weight scan should preferably be performed to find the optimal point that balances structure quality and data fit, i.e. MolProbity score and cross correlation, although we find that a weight value of 3 often provides a reasonable estimate. We’ve also shown that most structures deposited are well situated on the quality-correlation curve, with no clear overfitting observed though several structures could be further refined resulting in higher real space correlation and structure quality. Intriguingly, in other instances we found the deposited structure was exactly situated near the break point between structure quality and cross correlation. Finally, the use of the OPLS3e/VSGB force field within Phenix is made user-friendly by a tool that includes preparing the input structure, performing the refinement and calculating several validation metrics, and which is available in the 2020-3 release of the Schrödinger software. This release of the software is compatible with Phenix version 1.16 and later.

## Acknowledgments

This work was supported by the US National Institutes of Health (NIH) (grant P01GM063210) and the Phenix Industrial Consortium. This work was supported in part by the US Department of Energy under Contract No. DE-AC02-05CH1123.

## Supplemental information

**Figure S1.**
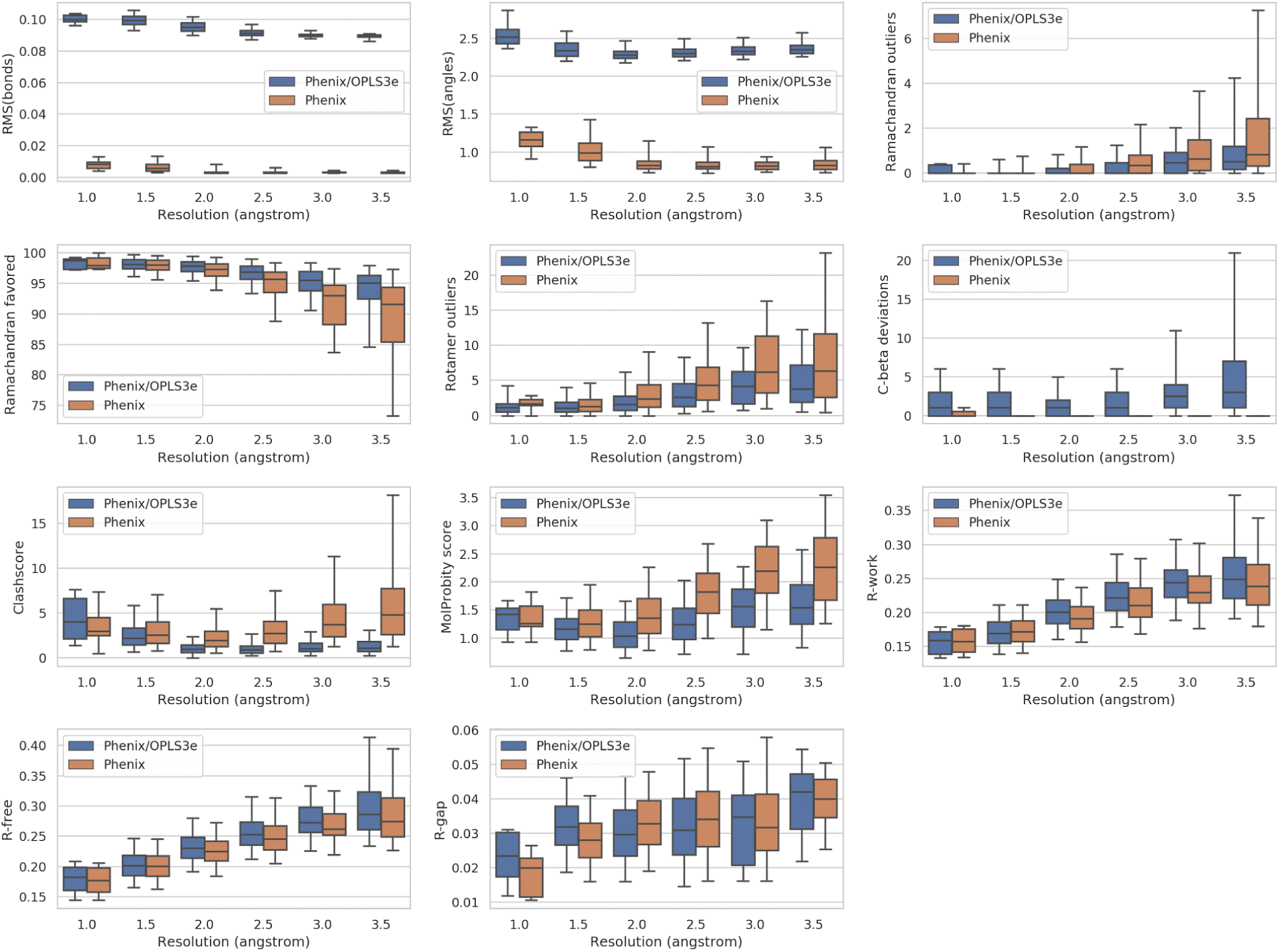
Reciprocal space refinement metrics after refinement with standard Phenix and Phenix/OPLS3e. Refinement metric distributions are shown for all individual MolProbity components and R-factors after refinement with Phenix/OPLS3e (blue) and Phenix (orange). The whiskers encompass the 95th percentile, the box the 50^th^ percentile, and the line in the box the median.

**Figure S2.**
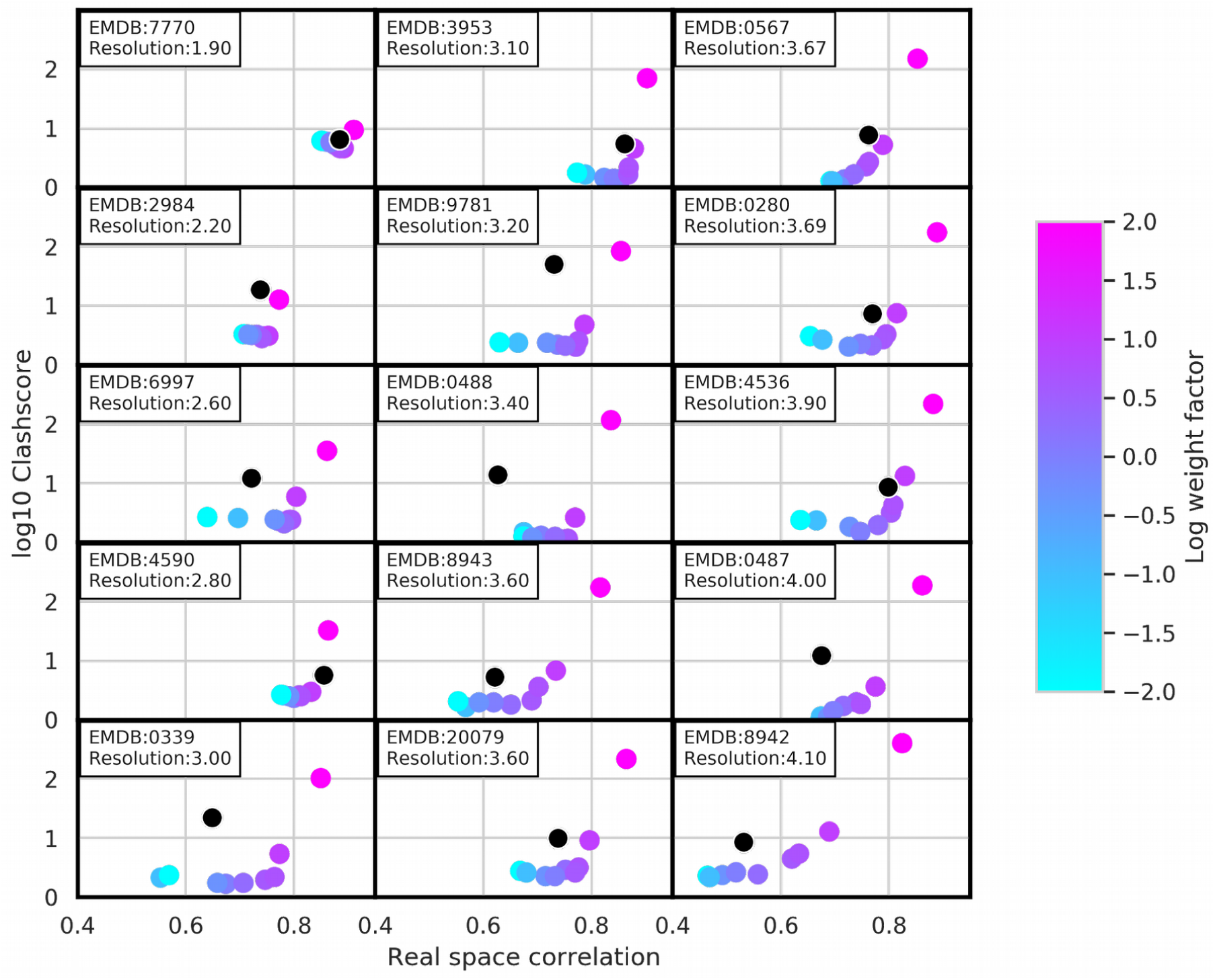
Phenix/OPLS3e real space refinement and the impact of data weight on the MolProbity Clashscore. The MolProbity score versus real space correlation is shown after refinement at different weight factors when using the OPLS3e force field. The black dot represents the deposited structure. The color of each dot represents the weight parameter for the impact of the Coulomb potential map used during refinement with blue indicating lower weights, and pink indicating higher weights.

**Figure S3.**
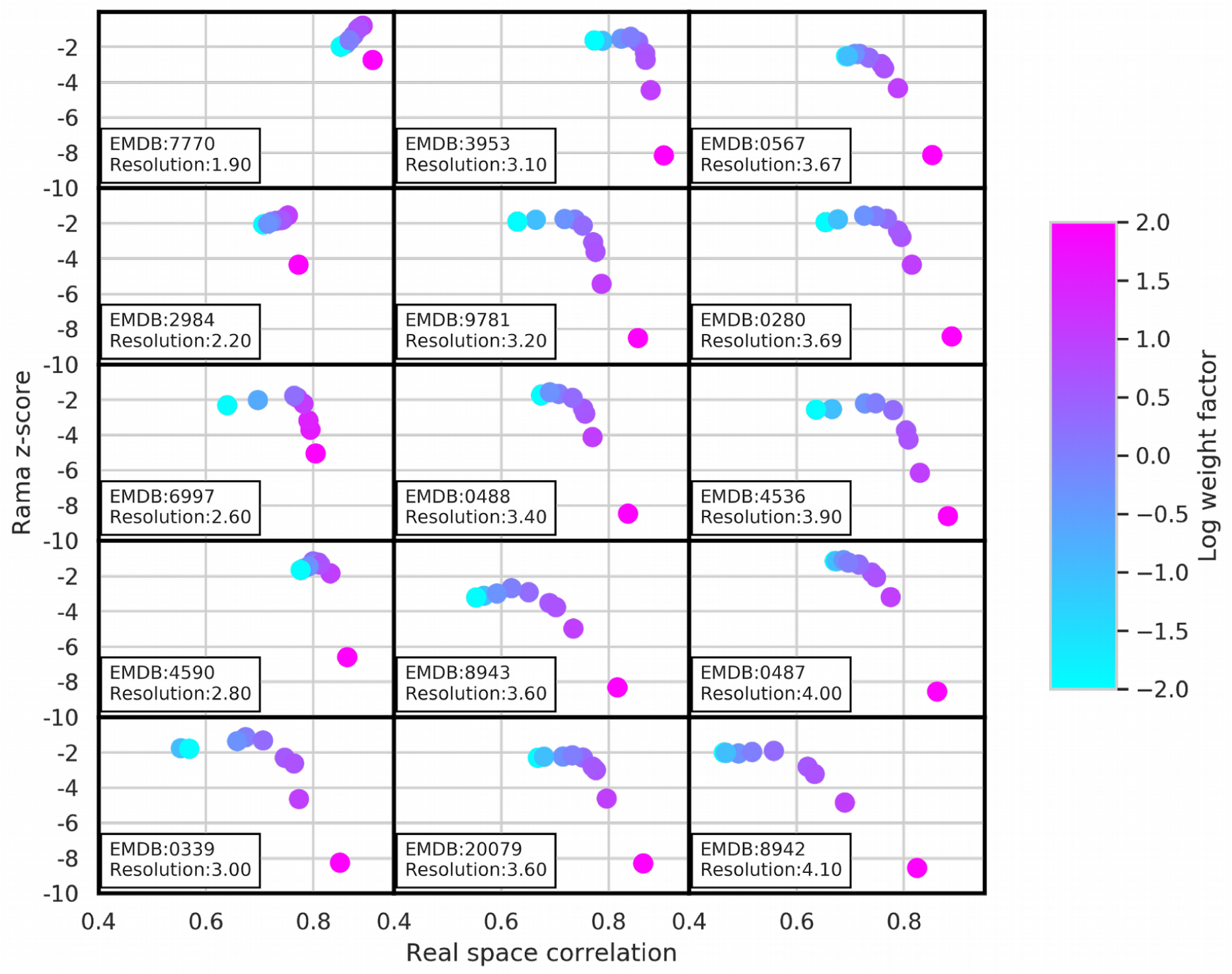
Phenix/OPLS3e real space refinement and the impact of data weight on the Ramachandran z-score. The Ramachandran z-score versus real space correlation is shown after refinement at different weight factors when using the OPLS3e force field. The color of each dot represents the weight parameter for the impact of the Coulomb potential map used during refinement with blue indicating lower weights, and pink indicating higher weights.

**Figure S4.**
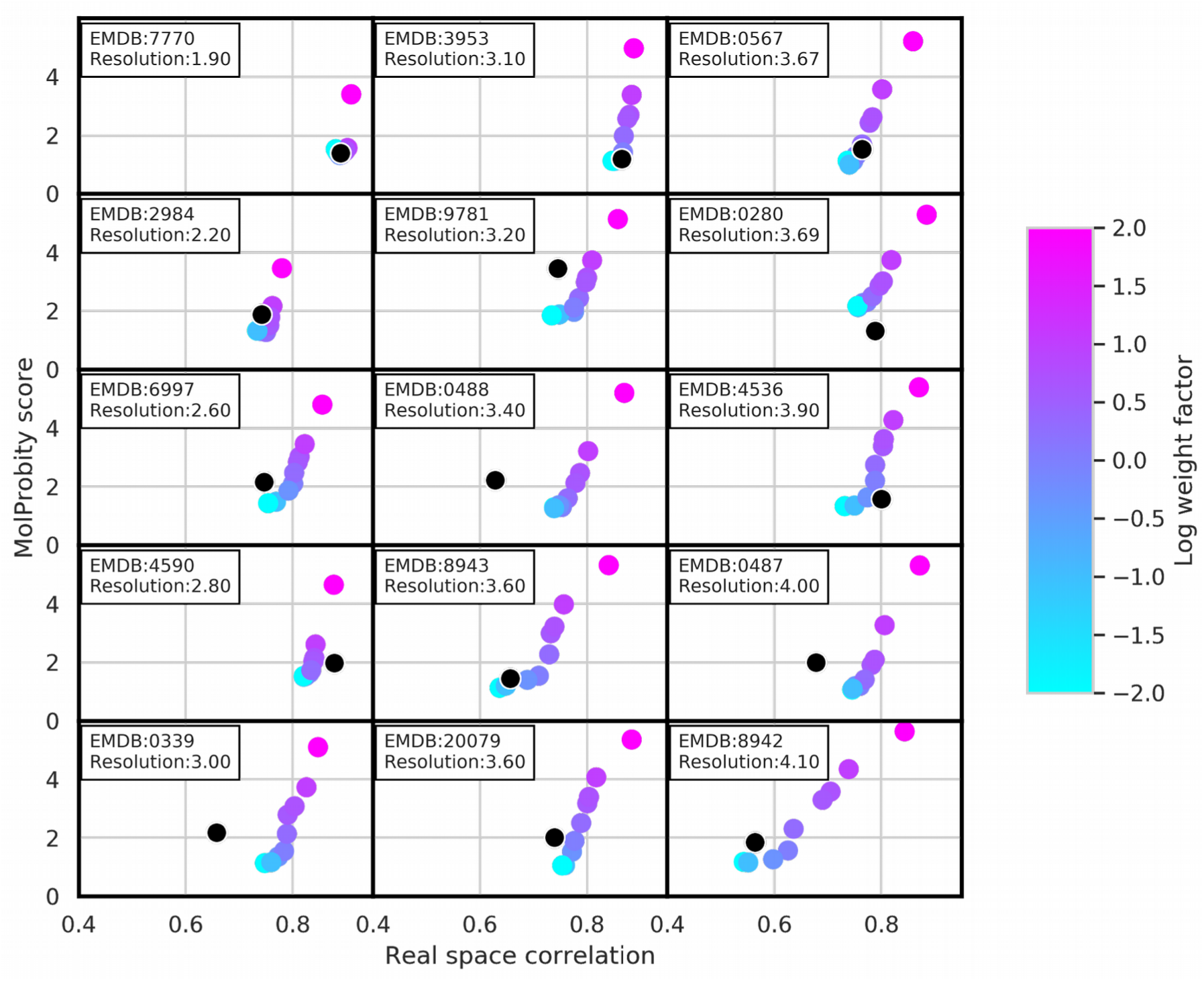
Standard Phenix real space refinement and the impact of data weight on the MolProbity score. Refinement metrics for 15 cryo-EM cases showing their MolProbity score and real space correlation after refinement with standard Phenix at different weights. The black dot represents the deposited structure. The color of the dot indicates the weight of the cryo-EM map with blue tints indicating low weights and pink tints indicating how weights.

## Notes

### Competing Interest Statement

The authors have declared no competing interest.

## References

Adams, P D and Pannu, N S and Read, R J and Brünger, A T Cross-validated maximum likelihood enhances crystallographic simulated annealing refinement. Proceedings of the National Academy of Sciences of the United States of America, 1997, Vol. 94, pp. 5018–5023

Adams, P.D., Afonine, P.V., Bunkóczi, G., Chen, V.B., Davis, I.W., Echols, N., Headd, J.J., Hung, L.-W., Kapral, G.J., Grosse-Kunstleve, R.W., McCoy, A.J., Moriarty, N.W., Oeffner, R., Read, R. J., Richardson, D.C., Richardson, J.S., Terwilliger, T.C. and Zwart, P.H. PHENIX: a comprehensive Python-based system for macromolecular structure solution. Acta crystallographica. Section D, Biological crystallography, 2010, Vol. 66, pp. 213–221.

Adams, P. D., Aertgeerts, K., Bauer, C., Bell, J. A., Berman, H. M., Bhat, T. N., Blaney, J. M., Bolton, E., Bricogne, G., Brown, D., Burley, S. K., Case, D. A., Clark, K. L., Darden, T., Emsley, P., Feher, V. A., Feng, Z., Groom, C. R., Harris, S. F., Hendle, J., Holder, T., Joachimiak, A., Kleywegt, G. J., Krojer, T., Marcotrigiano, J., Mark, A. E., Markley, J. L., Miller, M., Minor, W., Montelione, G. T., Murshudov, G., Nakagawa, A., Nakamura, H., Nicholls, A., Nicklaus, M., Nolte, R. T., Padyana, A. K., Peishoff, C. E., Pieniazek, S., Read, R. J., Shao, C., Sheriff, S., Smart, O., Soisson, S., Spurlino, J., Stouch, T., Svobodova, R., Tempel, W., Terwilliger, T. C., Tronrud, D., Velankar, S., Ward, S. C., Warren, G. L., Westbrook, J. D., Williams, P., Yang, H. and Young, J. Outcome of the First wwPDB/CCDC/D3R Ligand Validation Workshop. Structure, 2016, Vol. 24, pp. 502–508

Afonine, P. V., Grosse-Kunstleve, R. W., Echols, N., Headd, J. J., Moriarty, N. W., Mustyakimov, M., Terwilliger, T. C., Urzhumtsev, A., Zwart, P. H. and Adams, P. D. Towards automated crystallographic structure refinement with phenix.refine. Acta crystallographica. Section D, Biological crystallography, 2012, Vol. 68, pp. 352–367

Afonine, P.V., Poon, B.K., Read, R.J., Sobolev, O.V., Terwilliger, T.C., Urzhumtsev, A. and Adams, P.D. Real-space refinement in PHENIX for cryo-EM and crystallography. Acta crystallographica. Section D, Structural biology, 2018, Vol. 74, pp. 531–544

Bell, J. A., Ho, K. L. and Farid, R. Significant reduction in errors associated with nonbonded contacts in protein crystal structures: automated all-atom refinement with PrimeX. Acta crystallographica. Section D, Biological crystallography, 2012, Vol. 68, pp. 935–952

Beshnova, D. A., Pereira, J. and Lamzin, V. S. Estimation of the protein-ligand interaction energy for model building and validation. Acta crystallographica. Section D, Structural biology, 2017, Vol. 73, pp. 195–202

Borbulevych, O. Y., Plumley, J. A., Martin, R. I., Merz, K. M. and Westerhoff, L. M. Accurate macromolecular crystallographic refinement: incorporation of the linear scaling, semiempirical quantum-mechanics program DivCon into the PHENIX refinement package. Acta crystallographica. Section D, Biological crystallography, 2014, Vol. 70, pp. 1233–1247

Brunger, Axel T Free R value: a novel statistical quantity for assessing the accuracy of crystal structures Nature, 1992, Vol. 355, pp. 472.

Brunger, A.T., Adams, P.D., Clore, G.M., DeLano, W.L., Gros, P., Grosse-Kunstleve, R.W., Jiang, J-S, Kuszewski, J., Nilges, M., Pannu, N.S., Read, R.J., Rice, L.M., Simonson, T. and Warren, G.L. Crystallography & NMR system: A new software suite for macromolecular structure determination Acta Crystallographica Section D: Biological Crystallography, 1998, Vol. 54, pp. 905–921

Deller, M. C. and Rupp, B. Models of protein-ligand crystal structures: trust, but verify. Journal of computer-aided molecular design, 2015, Vol. 29, pp. 817–836

DiMaio, F., Zhang, J., Chiu, W., and Baker, D. Cryo-em model validation using independent map reconstructions Protein Science, 2013, Vol. 22, pp. 865–868.

Engh, R. A and Huber, R. Accurate bond and angle parameters for X-ray protein structure refinement Acta Crystallographica Section A: Foundations of Crystallography, 1991, Vol. 47, pp. 392–400

Engh, R.A. and Huber, R. Structure quality and target parameters International Tables for Crystallography, Vol. F, edited by M. G. Rossmann & E. Arnold, pp. 382–392. Dordrecht: Kluwer Academic Publishers

Groom, C.R, Bruno, I.J., Lightfoot, M.P. and Ward, S.C. The Cambridge Structural Database Acta Cryst. 2016, Vol. B72, pp. 171–179

Falkner, B. and Schröder, G.F. Cross-validation in cryo-EM-based structural modeling Proceedings of the National Academy of Sciences, 2013, Vol. 110, pp.8930–8935.

Fenn, T. D., Schnieders, M. J., Mustyakimov, M., Wu, C., Langan, P., Pande, V. S. and Brunger, A. T. Reintroducing electrostatics into macromolecular crystallographic refinement: application to neutron crystallography and DNA hydration. Structure (London, England : 1993), 2011, Vol. 19, pp. 523–533

Headd, J. J., Echols, N., Afonine, P. V., Grosse-Kunstleve, R. W., Chen, V. B., Moriarty, N. W., Richardson, D. C., Richardson, J. S. and Adams, P. D. Use of knowledge-based restraints in phenix.refine to improve macromolecular refinement at low resolution. Acta crystallographica. Section D, Biological crystallography, 2012, Vol. 68, pp. 381–390

Jaskolski, M., Gilski, M., Dauter, Z. and Wlodawer, A. Stereochemical restraints revisited: how accurate are refinement targets and how much should protein structures be allowed to deviate from them? Acta crystallographica. Section D, Biological crystallography, 2007, Vol. 63, pp. 611–620

Janowski, P. A., Moriarty, N. W., Kelley, B. P., Case, D. A., York, D. M., Adams, P. D. and Warren, G. L. Improved ligand geometries in crystallographic refinement using AFITT in PHENIX. Acta crystallographica. Section D, Structural biology, 2016, Vol. 72, pp. 1062–1072

Joosten, R. P., Long, F., Murshudov, G. N. and Perrakis, A. The PDB_REDO server for macromolecular structure model optimization. IUCrJ, 2014, Vol. 1, pp. 213–220

Li, J., Abel, R., Zhu, K., Cao, Y., Zhao, S. and Friesner, R. A. The VSGB 2.0 model: a next generation energy model for high resolution protein structure modeling Proteins: Structure, Function, and Bioinformatics, 2011, Vol. 79, pp. 2794–2812

Korostelev, A., Fenley, M.O., and Chapman, M.S. Impact of a Poisson-Boltzmann electrostatic restraint on protein structures refined at medium resolution Acta Crystallographica Section D: Biological Crystallography, 2004, Vol. 60, pp. 1786–1794

Kovalevskiy, Oleg and Nicholls, Robert A and Long, Fei and Carlon, Azzurra and Murshudov, Garib N Overview of refinement procedures within REFMAC5: utilizing data from different sources. Acta crystallographica. Section D, Structural biology, 2018, Vol. 74, pp. 215–227.

Lagerstedt, J.A., Jakobi, A., Burnley, T., Patwardhan, A., Topf, M. and Winn, M. Comparing cryo-EM reconstructions and validating atomic model fit using difference maps. Journal of Chemical Information and Modeling, 2020, Vol. 60, pp. 2552–2560

Li, Q., Pellegrino, J., Lee, D.J., Tran, A., Wang, R., Park, J., Ji, K., Chow, D., Zhang, N., Brilot, A., Biel, J., van Zundert, G.C.P., Borelli, K., Shinabarger, D., Wolfe, C., Murray, B., Jacobson, M.P., Fraser, J.S., Seiple, I.B. Synthesis and Mechanism of Action of Group a Streptogramin Antibiotics That Overcome Resistance ChemRxiv, 2019, doi: 10.26434/chemrxiv.8346107.v2.

Liebeschuetz, J., Hennemann, J., Olsson, T. and Groom, C. R. The good, the bad and the twisted: a survey of ligand geometry in protein crystal structures. Journal of computer-aided molecular design, 2012, Vol. 26, pp. 169–183

Liebschner, D., Afonine, P. V., Baker, M. L., Bunkoczi, G., Chen, V. B., Croll, T. I., Hintze, B., Hung, L.-W., Jain, S., McCoy, A. J., Moriarty, N. W., Oeffner, R. D., Poon, B. K., Prisant, M. G., Read, R. J., Richardson, J. S., Richardson, D. C., Sammito, M. D., Sobolev, O. V., Stockwell, D. H., Terwilliger, T. C., Urzhumtsev, A. G., Videau, L. L., Williams, C. J. & Adams, P. D. (2019). Macromolecular structure determination using X-rays, neutrons and electrons: recent developments in Phenix. Acta crystallographica. Section D, 2019, Vol. 75, pp. 861–877.

Monroe, L. and Terashi, G. and Kihara, D. Variability of protein structure models from electron microscopy Structure, 2017, Vol. 25, pp.592–602

Moriarty, N. W., Tronrud, D. E., Adams, P. D. and Karplus, P. A. A new default restraint library for the protein backbone in Phenix: a conformation-dependent geometry goes mainstream. Acta crystallographica. Section D, Structural biology, 2016, Vol. 72, pp. 176–179

Moriarty, N. W., Janowski, P. A., Swails, J. M., Nguyen, H., Richardson, J. S., Case, D. A. and Adams, P. D. Improved chemistry restraints for crystallographic refinement by integrating the Amber force field into Phenix Acta Cryst. D, 2020, Vol. 76, pp. 51–62.

Moulinier, L. and Case, D. A and Simonson, T. Reintroducing electrostatics into protein X-ray structure refinement: bulk solvent treated as a dielectric continuum Acta Crystallographica Section D: Biological Crystallography, 2003, pp. 2094–2103

Peach, M. L., Cachau, R. E. and Nicklaus, M. C. Conformational energy range of ligands in protein crystal structures: The difficult quest for accurate understanding Journal of Molecular Recognition, Wiley Online Library, 2017, Vol. 30, pp. E2618

Ponder, J.W. and Case, D.A. Force fields for protein simulations. Adv. Prot. Chem., 2003, Vol. 66, pp. 27–85.

Reynolds, C. H. Protein-ligand cocrystal structures: we can do better. ACS medicinal chemistry letters, 2014, Vol. 5, pp. 727–729

Robertson, M.l J., van Zundert, G. C. P., Borrelli, K., and Skiniotis, G. GemSpot: A Pipeline for Robust Modeling of Ligands into CryoEM Maps Structure, 2020, Vol. 28, Issue 6, pp. 707-716.e3

Roos, K., Wu, C., Damm, W., Reboul, M., Stevenson, J.M., Lu, C., Dahlgren, M.K., Mondal, S., Chen, W., Wang, L. Abel, R., Friesner, R.A. and Harder, E.D. OPLS3e: Extending force field coverage for drug-like small molecules Journal of chemical theory and computation, 2019, Vol. 15, pp.1863–1874

Schröder, G.F. and Levitt, M. and Brunger, A.T. Deformable elastic network refinement for low-resolution macromolecular crystallography. Acta crystallographica. Section D, Biological crystallography, 2014, Vol. 70, pp. 2241–2255

Sitzmann, M., Weidlich, I. E., Filippov, I. V., Liao, C., Peach, M. L., Ihlenfeldt, W.-D., Karki, R. G., Borodina, Y. V., Cachau, R. E. and Nicklaus, M. C. PDB ligand conformational energies calculated quantum-mechanically. Journal of chemical information and modeling, 2012, Vol. 52, pp. 739–756

Smart, O. S., Horský, V., Gore, S., Svobodová Vařeková, R., Bendová, V., Kleywegt, G. J. and Velankar, S. Validation of ligands in macromolecular structures determined by X-ray crystallography. Acta crystallographica. Section D, Structural biology, 2018, Vol. 74, pp. 228–236

Steiner, R. A. and Tucker, J. A. Keep it together: restraints in crystallographic refinement of macromolecule-ligand complexes. Acta crystallographica. Section D, Structural biology, 2017, Vol. 73, pp. 93–102

Tickle, I.J., Laskowski, R.A., and Moss, D.S. R_free_ and the R_free_ Ratio. I. Derivation of Expected Values of Cross-Validation Residuals Used in Macromolecular Least-Squares Refinement. Acta Crystallographica, Section D, Structural Biology, 1998, Vol. 54, pp. 547–557.

Tronrud, Dale E and Berkholz, Donald S and Karplus, P Andrew Using a conformation-dependent stereochemical library improves crystallographic refinement of proteins. Acta crystallographica. Section D, Biological crystallography, 2010, Vol. 66, pp. 834–843

Vagin, Alexei A and Steiner, Roberto A and Lebedev, Andrey A and Potterton, Liz and McNicholas, Stuart and Long, Fei and Murshudov, Garib N REFMAC5 dictionary: organization of prior chemical knowledge and guidelines for its use. Acta crystallographica. Section D, Biological crystallography, 2004, Vol. 60, pp. 2184–2195

Volkmann, N. Confidence intervals for fitting of atomic models into low-resolution densities Acta Crystallographica Section D: Biological Crystallography, 2009, Vol. 65, pp. 679–689.

Wang, J. and Moore, P.B. On the interpretation of electron microscopic maps of biological macromolecules Protein Science, 2017, Vol. 26, pp. 122–129.

Wang, L., Kruse, H., Sogolev, O.V., Moriarty, N.W., Waller, M.P., Afonine, P.V. and Biczysko, M. Real-space quantum-based refinement for cryo-EM: Q|R#3 Biorxiv, doi: https://doi.org/10.1101/2020.05.25.115386

Williams, C. J., Headd, J. J., Moriarty, N. W., Prisant, M. G., Videau, L. L., Deis, L. N., Verma, V., Keedy, D. A., Hintze, B. J., Chen, V. B., Jain, S., Lewis, S. M., Arendall, W. B., Snoeyink, J., Adams, P. D., Lovell, S. C., Richardson, J. S. and Richardson, D. C. MolProbity: More and better reference data for improved all-atom structure validation. Protein science : a publication of the Protein Society, 2018, Vol. 27, pp. 293–315

wwPDB consortium Protein Data Bank: the single global archive for 3D macromolecular structure data Nucleic Acids Research, 2019, Vol. 47, pp. D520–D528

Zheng M., Moriarty N.W., Xu Y.T., Reimers J.R., Afonine P.V. and Waller M.P. Solving the scalability issue in quantum-based refinement: Q|R#1 Acta Crystallographica Section D: Biological Crystallography, 2017, Vol. 73, pp. 1020–1028,

Zheng M, Biczysko M, Xu Y, Moriarty NW, Kruse H, Urzhumtsev A, Waller MP, Afonine PV. Including crystallographic symmetry in quantum-based refinement: Q|R#2. Acta Crystallographica Section D: Biological Crystallography, 2020, Vol., 76, pp. 41–50

