## Supplementary material for "Macromolecular refinement of X-ray and cryo-electron microscopy structures with Phenix / OPLS3e for improved structure and ligand quality": Table S1

| pdbid | protocol | Resolution | RMS(bonds) | RMS(angles) | Ramachandran_outliers | Ramachandran_favored | Rotamer_outliers | C-beta_deviations | Clashscore | MolProbity_score | R-work | R-free | R-gap |
|---|---|---|---|---|---|---|---|---|---|---|---|---|---|
| 108m | opls | 2.67 | 0.0957 | 2.31 | 0 | 97.35 | 3.15 | 1 | 4.2 | 1.71 | 0.136 | 0.1805 | 0.0445 |
| 108m | noschrodinger | 2.67 | 0.0035 | 0.75 | 0 | 96.71 | 3.91 | 0 | 7.26 | 2.06 | 0.1226 | 0.1712 | 0.0486 |
| 1a0r | opls | 2.8 | 0.0907 | 2.33 | 0.17 | 96.56 | 3.93 | 2 | 1.08 | 1.49 | 0.2301 | 0.2674 | 0.0373 |
| 1a0r | noschrodinger | 2.8 | 0.0027 | 0.76 | 0.51 | 95.21 | 3.52 | 0 | 3.47 | 1.89 | 0.2191 | 0.2581 | 0.039 |
| 1a27 | opls | 1.9 | 0.0968 | 2.49 | 0.36 | 97.51 | 5.96 | 6 | 1.79 | 1.63 | 0.1924 | 0.2372 | 0.0448 |
| 1a27 | noschrodinger | 1.9 | 0.0038 | 0.98 | 0.35 | 97.17 | 8.51 | 0 | 3.62 | 2.01 | 0.1915 | 0.2308 | 0.0393 |
| 1a53 | opls | 2.02 | 0.1017 | 2.33 | 0.41 | 98.36 | 8.52 | 1 | 1.47 | 1.59 | 0.1384 | 0.1667 | 0.0283 |
| 1a53 | noschrodinger | 2.02 | 0.0027 | 0.82 | 0.41 | 97.96 | 8.48 | 0 | 1.23 | 1.56 | 0.1437 | 0.1686 | 0.0249 |
| 1ahn | opls | 2.4 | 0.0926 | 2.32 | 0 | 96.97 | 7.97 | 4 | 0.38 | 1.5 | 0.2336 | 0.2663 | 0.0327 |
| 1ahn | noschrodinger | 2.4 | 0.0028 | 0.81 | 0.6 | 91.02 | 7.97 | 0 | 3.8 | 2.37 | 0.2179 | 0.2645 | 0.0466 |
| 1ai7 | opls | 2.5 | 0.1004 | 2.2 | 0 | 98.15 | 2.36 | 2 | 0.5 | 0.96 | 0.1347 | 0.1754 | 0.0407 |
| 1ai7 | noschrodinger | 2.5 | 0.0032 | 0.84 | 0.13 | 97.89 | 5.64 | 0 | 1.76 | 1.53 | 0.1214 | 0.1666 | 0.0452 |
| 1aod | opls | 2.64 | 0.0947 | 2.21 | 0.37 | 97.76 | 2.85 | 0 | 0.9 | 1.17 | 0.1471 | 0.197 | 0.0499 |
| 1aod | noschrodinger | 2.64 | 0.0095 | 1.16 | 0.37 | 97.43 | 4.45 | 0 | 3.37 | 1.73 | 0.1441 | 0.2007 | 0.0566 |
| 1aur | opls | 2.51 | 0.0959 | 2.28 | 0 | 98.6 | 5.43 | 1 | 1.04 | 1.36 | 0.171 | 0.2167 | 0.0457 |
| 1aur | noschrodinger | 2.51 | 0.0029 | 0.8 | 0.23 | 97.69 | 8.24 | 0 | 2.09 | 1.74 | 0.1586 | 0.208 | 0.0494 |
| 1awb | opls | 2.5 | 0.0979 | 2.25 | 0.37 | 97.77 | 1.32 | 1 | 1.07 | 0.96 | 0.1491 | 0.1905 | 0.0414 |
| 1awb | noschrodinger | 2.5 | 0.0035 | 0.93 | 0 | 97.78 | 1.98 | 0 | 2.15 | 1.26 | 0.141 | 0.1806 | 0.0396 |
| 1azs | opls | 2.32 | 0.0915 | 2.29 | 0.14 | 97.16 | 0.64 | 2 | 0.26 | 0.75 | 0.2086 | 0.2455 | 0.0369 |
| 1azs | noschrodinger | 2.32 | 0.0025 | 0.75 | 0.14 | 96.47 | 1.59 | 0 | 1.39 | 1.26 | 0.2078 | 0.2407 | 0.0329 |
| 1b1c | opls | 1.93 | 0.094 | 2.48 | 0 | 98.16 | 2.17 | 2 | 1.92 | 1.21 | 0.152 | 0.1955 | 0.0435 |
| 1b1c | noschrodinger | 1.93 | 0.0061 | 0.93 | 0 | 99.39 | 2.17 | 0 | 1.54 | 1.15 | 0.1552 | 0.1894 | 0.0342 |
| 1b5e | opls | 1.6 | 0.0939 | 2.43 | 0 | 98.1 | 3.76 | 6 | 2.43 | 1.46 | 0.1548 | 0.1768 | 0.022 |
| 1b5e | noschrodinger | 1.6 | 0.0076 | 0.97 | 0 | 97.7 | 3.49 | 0 | 1.66 | 1.39 | 0.158 | 0.1754 | 0.0174 |
| 1bqs | opls | 2.2 | 0.0947 | 2.58 | 7.77 | 85.44 | 10.59 | 9 | 4.8 | 2.68 | 0.2372 | 0.2635 | 0.0263 |
| 1bqs | noschrodinger | 2.2 | 0.0038 | 1.03 | 8.7 | 84.06 | 23.98 | 0 | 10.88 | 3.28 | 0.2293 | 0.268 | 0.0387 |
| 1btn | opls | 2.03 | 0.0987 | 2.36 | 0 | 99.03 | 5.49 | 1 | 0 | 1.06 | 0.1686 | 0.2212 | 0.0526 |
| 1btn | noschrodinger | 2.03 | 0.0025 | 0.72 | 0 | 98.08 | 5.43 | 0 | 4.04 | 1.75 | 0.1811 | 0.224 | 0.0429 |
| 1byh | opls | 2.57 | 0.0936 | 2.27 | 0.47 | 95.26 | 9.14 | 3 | 2.42 | 2.08 | 0.1782 | 0.2363 | 0.0581 |
| 1byh | noschrodinger | 2.57 | 0.0047 | 1.02 | 1.42 | 93.87 | 12.5 | 0 | 5.13 | 2.51 | 0.1738 | 0.2231 | 0.0493 |
| 1c3q | opls | 2 | 0.0939 | 2.32 | 0.47 | 96.44 | 1.4 | 3 | 1.04 | 1.15 | 0.1689 | 0.2137 | 0.0448 |
| 1c3q | noschrodinger | 2 | 0.006 | 0.95 | 0.59 | 95.74 | 1.23 | 0 | 2.34 | 1.38 | 0.1698 | 0.2115 | 0.0417 |
| 1c9k | opls | 2.1 | 0.0967 | 2.59 | 0.47 | 98.11 | 9.09 | 1 | 0.89 | 1.5 | 0.2292 | 0.2631 | 0.0339 |
| 1c9k | noschrodinger | 2.1 | 0.0033 | 0.8 | 0.4 | 96.79 | 14.51 | 0 | 7.46 | 2.49 | 0.2209 | 0.2571 | 0.0362 |
| 1cf5 | opls | 2.43 | 0.0916 | 2.36 | 1.02 | 95.33 | 13.59 | 9 | 0.97 | 1.98 | 0.2089 | 0.2545 | 0.0456 |
| 1cf5 | noschrodinger | 2.43 | 0.0077 | 1.28 | 2.23 | 91.5 | 19.95 | 0 | 7.87 | 2.92 | 0.1882 | 0.2387 | 0.0505 |
| 1cr1 | opls | 2.1 | 0.0912 | 2.39 | 0.47 | 96.74 | 4.62 | 1 | 2.08 | 1.69 | 0.2461 | 0.2736 | 0.0275 |
| 1cr1 | noschrodinger | 2.1 | 0.0029 | 0.8 | 0 | 93.67 | 12 | 0 | 5.5 | 2.54 | 0.2345 | 0.2735 | 0.039 |
| 1cr5 | opls | 2.3 | 0.0963 | 2.18 | 0 | 97.85 | 2.39 | 0 | 1.18 | 1.16 | 0.2215 | 0.2541 | 0.0326 |
| 1cr5 | noschrodinger | 2.3 | 0.0029 | 0.77 | 0.19 | 96.52 | 2.81 | 0 | 4.63 | 1.8 | 0.1962 | 0.2426 | 0.0464 |
| 1ctt | opls | 2.04 | 0.0926 | 2.31 | 0 | 97.94 | 8.41 | 1 | 2.48 | 1.75 | 0.182 | 0.1992 | 0.0172 |
| 1ctt | noschrodinger | 2.04 | 0.003 | 0.83 | 0 | 96.92 | 9.69 | 0 | 2.25 | 1.94 | 0.1775 | 0.1914 | 0.0139 |
| 1cv8 | opls | 1.75 | 0.0987 | 2.29 | 0 | 98.83 | 0 | 0 | 2.17 | 0.99 | 0.147 | 0.1714 | 0.0244 |
| 1cv8 | noschrodinger | 1.75 | 0.0096 | 1.17 | 0 | 98.83 | 1.37 | 0 | 0.72 | 0.84 | 0.1492 | 0.1719 | 0.0227 |
| 1d1g | opls | 2.1 | 0.0953 | 2.55 | 0 | 99.38 | 2.41 | 3 | 2.51 | 1.33 | 0.1635 | 0.2074 | 0.0439 |
| 1d1g | noschrodinger | 2.1 | 0.0045 | 0.95 | 0 | 99.07 | 2.4 | 2 | 2.9 | 1.37 | 0.1581 | 0.2034 | 0.0453 |

| | | | | | | | | | | | | | |
|---|---|---|---|---|---|---|---|---|---|---|---|---|---|
| 1d2n | opls | 1.75 | 0.1023 | 2.83 | 0 | 97.94 | 0.9 | 3 | 2.46 | 1.04 | 0.1677 | 0.2082 | 0.0405 |
| 1d2n | noschrodinger | 1.75 | 0.0057 | 0.95 | 0 | 97.13 | 0 | 0 | 4.67 | 1.4 | 0.1753 | 0.2127 | 0.0374 |
| 1d2s | opls | 1.55 | 0.0977 | 2.31 | 0 | 96.93 | 1.31 | 0 | 1.78 | 1.21 | 0.1895 | 0.2247 | 0.0352 |
| 1d2s | noschrodinger | 1.55 | 0.0068 | 1.11 | 0 | 96.99 | 3.23 | 0 | 1.79 | 1.5 | 0.1893 | 0.2233 | 0.034 |
| 1dek | opls | 2.02 | 0.0976 | 2.41 | 0 | 96.5 | 2.93 | 0 | 1.44 | 1.46 | 0.1645 | 0.2015 | 0.037 |
| 1dek | noschrodinger | 2.02 | 0.003 | 0.78 | 0.22 | 94.13 | 4.85 | 0 | 1.96 | 1.88 | 0.1705 | 0.2051 | 0.0346 |
| 1dnl | opls | 1.8 | 0.0976 | 2.72 | 0.54 | 97.28 | 4.62 | 4 | 2.67 | 1.69 | 0.1807 | 0.202 | 0.0213 |
| 1dnl | noschrodinger | 1.8 | 0.0115 | 1.17 | 0.51 | 97.46 | 5.03 | 0 | 7.7 | 2.06 | 0.1771 | 0.2005 | 0.0234 |
| 1drj | opls | 1.96 | 0.0974 | 2.24 | 0.37 | 98.13 | 0.95 | 1 | 0 | 0.5 | 0.1567 | 0.2011 | 0.0444 |
| 1drj | noschrodinger | 1.96 | 0.0057 | 0.86 | 0.37 | 96.65 | 1.42 | 0 | 1.22 | 1.17 | 0.1527 | 0.2012 | 0.0485 |
| 1drk | opls | 1.8 | 0.0993 | 2.29 | 0.37 | 98.51 | 1.44 | 2 | 0.74 | 0.85 | 0.1545 | 0.2007 | 0.0462 |
| 1drk | noschrodinger | 1.8 | 0.0039 | 0.9 | 0.37 | 98.51 | 1.43 | 0 | 2.21 | 1.11 | 0.1587 | 0.2013 | 0.0426 |
| 1ds6 | opls | 2.3 | 0.0947 | 2.29 | 1.43 | 96.56 | 0.95 | 2 | 0.52 | 0.9 | 0.2265 | 0.2545 | 0.028 |
| 1ds6 | noschrodinger | 2.3 | 0.003 | 0.86 | 1.97 | 94.38 | 1.9 | 0 | 1.05 | 1.4 | 0.2091 | 0.25 | 0.0409 |
| 1dtp | opls | 2.51 | 0.0896 | 2.37 | 0.55 | 95.05 | 8.44 | 1 | 0.69 | 1.77 | 0.2637 | 0.3051 | 0.0414 |
| 1dtp | noschrodinger | 2.51 | 0.0036 | 0.97 | 2.13 | 85.11 | 16.23 | 0 | 5.5 | 2.87 | 0.2215 | 0.2687 | 0.0472 |
| 1ei9 | opls | 2.25 | 0.0923 | 2.33 | 0 | 96.01 | 3.33 | 0 | 0.22 | 1.26 | 0.2453 | 0.2779 | 0.0326 |
| 1ei9 | noschrodinger | 2.25 | 0.0053 | 0.9 | 0.36 | 94.58 | 4.98 | 0 | 2.66 | 1.95 | 0.2269 | 0.2759 | 0.049 |
| 1eom | opls | 2.1 | 0.101 | 2.22 | 0.36 | 97.15 | 0.86 | 0 | 0.22 | 0.74 | 0.1794 | 0.2084 | 0.029 |
| 1eom | noschrodinger | 2.1 | 0.0058 | 0.98 | 0.71 | 96.8 | 1.29 | 0 | 6.6 | 1.65 | 0.1606 | 0.1966 | 0.036 |
| 1ep3 | opls | 2.1 | 0.0998 | 2.3 | 0 | 96.83 | 3.22 | 2 | 1.59 | 1.49 | 0.1914 | 0.2231 | 0.0317 |
| 1ep3 | noschrodinger | 2.1 | 0.0033 | 0.85 | 0.53 | 95.95 | 3.85 | 0 | 3.24 | 1.84 | 0.1727 | 0.2123 | 0.0396 |
| 1ex9 | opls | 2.54 | 0.0952 | 2.5 | 0 | 97.16 | 1.3 | 3 | 1.16 | 1.07 | 0.1806 | 0.2226 | 0.042 |
| 1ex9 | noschrodinger | 2.54 | 0.0029 | 0.78 | 0 | 96.47 | 2.16 | 0 | 1.17 | 1.32 | 0.1904 | 0.2299 | 0.0395 |
| 1exb | opls | 2.1 | 0.098 | 2.36 | 0.25 | 98.75 | 2.56 | 0 | 1.04 | 1.11 | 0.1926 | 0.205 | 0.0124 |
| 1exb | noschrodinger | 2.1 | 0.0063 | 1.02 | 0.24 | 98.31 | 2.27 | 0 | 2.99 | 1.36 | 0.172 | 0.1957 | 0.0237 |
| 1exc | opls | 2.71 | 0.0938 | 2.34 | 0.28 | 97.79 | 5.1 | 1 | 1.17 | 1.41 | 0.2005 | 0.2457 | 0.0452 |
| 1exc | noschrodinger | 2.71 | 0.0028 | 0.79 | 0.55 | 93.99 | 9.12 | 0 | 5.32 | 2.42 | 0.1859 | 0.2282 | 0.0423 |
| 1f5v | opls | 1.55 | 0.0989 | 2.48 | 0.42 | 97.05 | 3.23 | 6 | 2.22 | 1.55 | 0.1463 | 0.1802 | 0.0339 |
| 1f5v | noschrodinger | 1.55 | 0.0068 | 1.05 | 0.21 | 97.06 | 2.48 | 0 | 3.39 | 1.6 | 0.1515 | 0.176 | 0.0245 |
| 1f7s | opls | 2 | 0.0981 | 2.28 | 0 | 98.35 | 1.85 | 1 | 1.45 | 1.08 | 0.1944 | 0.2266 | 0.0322 |
| 1f7s | noschrodinger | 2 | 0.003 | 0.75 | 0 | 97.54 | 1.83 | 0 | 1.93 | 1.25 | 0.1871 | 0.2183 | 0.0312 |
| 1fq1 | opls | 2.8 | 0.0905 | 2.44 | 1.71 | 89.55 | 5.97 | 6 | 1.81 | 2.09 | 0.2817 | 0.3029 | 0.0212 |
| 1fq1 | noschrodinger | 2.8 | 0.0032 | 0.9 | 5.51 | 78.81 | 9.72 | 0 | 11.39 | 3.07 | 0.267 | 0.2879 | 0.0209 |
| 1fsl | opls | 2.31 | 0.0923 | 2.21 | 0 | 97.86 | 2.22 | 2 | 1.31 | 1.15 | 0.1438 | 0.1897 | 0.0459 |
| 1fsl | noschrodinger | 2.31 | 0.0041 | 0.75 | 0.71 | 96.81 | 2.64 | 0 | 3.73 | 1.68 | 0.1299 | 0.1833 | 0.0534 |
| 1g0t | opls | 2.56 | 0.0917 | 2.21 | 0.24 | 97.17 | 0.28 | 0 | 0.76 | 0.89 | 0.2156 | 0.2478 | 0.0322 |
| 1g0t | noschrodinger | 2.56 | 0.003 | 0.76 | 0.47 | 95.78 | 0.56 | 0 | 1.07 | 1.1 | 0.1988 | 0.2434 | 0.0446 |
| 1g5t | opls | 1.9 | 0.0994 | 2.31 | 0 | 98.05 | 3.17 | 1 | 1.6 | 1.29 | 0.1914 | 0.238 | 0.0466 |
| 1g5t | noschrodinger | 1.9 | 0.0047 | 1.49 | 0 | 99.35 | 4.76 | 0 | 4.4 | 1.73 | 0.1898 | 0.2298 | 0.04 |
| 1g81 | opls | 1.7 | 0.1004 | 2.23 | 0 | 98.76 | 2.08 | 0 | 2.23 | 1.24 | 0.184 | 0.2173 | 0.0333 |
| 1g81 | noschrodinger | 1.7 | 0.0032 | 0.87 | 0 | 98.77 | 2.07 | 0 | 0.37 | 0.87 | 0.1829 | 0.2198 | 0.0369 |
| 1gc6 | opls | 2.9 | 0.0906 | 2.28 | 1.37 | 95.21 | 5.24 | 0 | 1.4 | 1.75 | 0.2087 | 0.2441 | 0.0354 |
| 1gc6 | noschrodinger | 2.9 | 0.0032 | 0.81 | 2.03 | 93.22 | 8.96 | 0 | 3 | 2.25 | 0.2105 | 0.2442 | 0.0337 |
| 1gfm | opls | 3.3 | 0.0941 | 2.18 | 0 | 96.44 | 3.82 | 0 | 0.38 | 1.31 | 0.164 | 0.2058 | 0.0418 |
| 1gfm | noschrodinger | 3.3 | 0.0028 | 0.77 | 0 | 94.67 | 3.44 | 0 | 2.5 | 1.81 | 0.1422 | 0.195 | 0.0528 |
| 1gka | opls | 3 | 0.0896 | 2.32 | 0.57 | 95.4 | 5.21 | 3 | 1.03 | 1.67 | 0.2209 | 0.2415 | 0.0206 |
| 1gka | noschrodinger | 3 | 0.0043 | 0.98 | 0.86 | 89.71 | 11.33 | 0 | 5.86 | 2.68 | 0.2271 | 0.2459 | 0.0188 |
| 1gky | opls | 2.01 | 0.1011 | 2.26 | 0 | 98.37 | 1.9 | 0 | 0 | 0.71 | 0.1841 | 0.2188 | 0.0347 |

| | | | | | | | | | | | | | |
|---|---|---|---|---|---|---|---|---|---|---|---|---|---|
| 1gky | noschrodinger | 2.01 | 0.0028 | 0.79 | 0 | 98.37 | 3.8 | 0 | 1.71 | 1.36 | 0.1782 | 0.2073 | 0.0291 |
| 1gny | opls | 1.63 | 0.1057 | 2.22 | 0 | 98 | 2.52 | 1 | 1.31 | 1.16 | 0.145 | 0.1777 | 0.0327 |
| 1gny | noschrodinger | 1.63 | 0.005 | 1.33 | 0 | 98.01 | 3.36 | 0 | 5.24 | 1.68 | 0.1496 | 0.1825 | 0.0329 |
| 1gqc | opls | 2.6 | 0.0971 | 2.29 | 0.21 | 97.26 | 2.77 | 0 | 1.05 | 1.28 | 0.1743 | 0.2092 | 0.0349 |
| 1gqc | noschrodinger | 2.6 | 0.0028 | 0.78 | 0 | 96.45 | 3.75 | 0 | 1.05 | 1.48 | 0.1589 | 0.208 | 0.0491 |
| 1gs5 | opls | 1.44 | 0.0996 | 2.37 | 0.39 | 97.24 | 1.44 | 1 | 2.47 | 1.29 | 0.1842 | 0.2025 | 0.0183 |
| 1gs5 | noschrodinger | 1.44 | 0.0051 | 1.07 | 0 | 97.27 | 2.37 | 1 | 7.4 | 1.83 | 0.1803 | 0.2028 | 0.0225 |
| 1gu3 | opls | 2.3 | 0.0952 | 2.55 | 0 | 99.28 | 6.54 | 0 | 2.39 | 1.64 | 0.231 | 0.2631 | 0.0321 |
| 1gu3 | noschrodinger | 2.3 | 0.0032 | 0.98 | 0 | 99.29 | 7.41 | 0 | 5.25 | 1.94 | 0.2171 | 0.2638 | 0.0467 |
| 1gui | opls | 1.9 | 0.0954 | 2.26 | 0 | 98.03 | 0 | 0 | 1.53 | 0.9 | 0.1685 | 0.1935 | 0.025 |
| 1gui | noschrodinger | 1.9 | 0.0143 | 1.55 | 0 | 97.39 | 1.49 | 0 | 7.68 | 1.67 | 0.1751 | 0.1936 | 0.0185 |
| 1gwv | opls | 2.5 | 0.094 | 2.22 | 0 | 97.02 | 1.74 | 1 | 1.35 | 1.22 | 0.1947 | 0.2228 | 0.0281 |
| 1gwv | noschrodinger | 2.5 | 0.003 | 0.81 | 0 | 96.49 | 2.52 | 0 | 2.28 | 1.54 | 0.1859 | 0.2205 | 0.0346 |
| 1h0a | opls | 1.7 | 0.1024 | 2.47 | 0 | 99.35 | 3.25 | 4 | 3.54 | 1.53 | 0.1819 | 0.2069 | 0.025 |
| 1h0a | noschrodinger | 1.7 | 0.0079 | 0.98 | 0 | 98.72 | 4.49 | 0 | 3.53 | 1.64 | 0.1856 | 0.2076 | 0.022 |
| 1h2h | opls | 2.49 | 0.0916 | 2.25 | 1.37 | 95.43 | 2.99 | 1 | 1.32 | 1.54 | 0.2256 | 0.268 | 0.0424 |
| 1h2h | noschrodinger | 2.49 | 0.0028 | 0.77 | 2.18 | 93.45 | 2.43 | 0 | 3.44 | 1.86 | 0.2118 | 0.27 | 0.0582 |
| 1h65 | opls | 2 | 0.1001 | 2.27 | 0 | 98.91 | 0.77 | 5 | 1.66 | 0.92 | 0.162 | 0.2099 | 0.0479 |
| 1h65 | noschrodinger | 2 | 0.0031 | 0.83 | 0 | 98.43 | 0.91 | 0 | 1.74 | 0.93 | 0.1685 | 0.2114 | 0.0429 |
| 1h6h | opls | 1.7 | 0.1037 | 2.44 | 0 | 98.57 | 0 | 0 | 0.42 | 0.65 | 0.1727 | 0.201 | 0.0283 |
| 1h6h | noschrodinger | 1.7 | 0.0127 | 1.19 | 0.71 | 97.16 | 1.6 | 0 | 2.11 | 1.29 | 0.1633 | 0.2009 | 0.0376 |
| 1h7g | opls | 2.13 | 0.1008 | 2.28 | 0 | 98.12 | 2.01 | 0 | 0.92 | 1.01 | 0.1845 | 0.2066 | 0.0221 |
| 1h7g | noschrodinger | 2.13 | 0.0027 | 0.8 | 0 | 97.93 | 2.49 | 0 | 1.45 | 1.2 | 0.1653 | 0.2013 | 0.036 |
| 1h8p | opls | 1.82 | 0.1018 | 2.17 | 0 | 96.51 | 0 | 2 | 1.35 | 1.09 | 0.1705 | 0.197 | 0.0265 |
| 1h8p | noschrodinger | 1.82 | 0.0064 | 0.95 | 0 | 95.93 | 0 | 0 | 2.03 | 1.25 | 0.1687 | 0.1992 | 0.0305 |
| 1hd2 | opls | 1.5 | 0.1056 | 2.62 | 0 | 96.2 | 1.56 | 0 | 2.46 | 1.43 | 0.1529 | 0.1621 | 0.0092 |
| 1hd2 | noschrodinger | 1.5 | 0.0061 | 1.03 | 0 | 95.6 | 1.56 | 0 | 2.45 | 1.48 | 0.1557 | 0.1649 | 0.0092 |
| 1hg5 | opls | 2 | 0.097 | 2.38 | 0.38 | 96.54 | 0.43 | 0 | 0.23 | 0.81 | 0.188 | 0.2004 | 0.0124 |
| 1hg5 | noschrodinger | 2 | 0.0065 | 1.31 | 0.77 | 93.87 | 2.13 | 0 | 3.49 | 1.8 | 0.1808 | 0.2042 | 0.0234 |
| 1hkq | opls | 2.6 | 0.0928 | 2.39 | 1.22 | 94.31 | 3.18 | 2 | 1.21 | 1.61 | 0.2094 | 0.2621 | 0.0527 |
| 1hkq | noschrodinger | 2.6 | 0.0028 | 0.75 | 3.25 | 91.46 | 5.45 | 0 | 4.59 | 2.3 | 0.2314 | 0.2666 | 0.0352 |
| 1htw | opls | 1.74 | 0.0996 | 2.43 | 0.86 | 96.11 | 6.2 | 1 | 3.26 | 1.98 | 0.1811 | 0.2302 | 0.0491 |
| 1htw | noschrodinger | 1.74 | 0.0082 | 1.13 | 0.85 | 94.44 | 6.37 | 0 | 4.18 | 2.19 | 0.188 | 0.2299 | 0.0419 |
| 1hx9 | opls | 3.31 | 0.0919 | 2.32 | 0.19 | 97.88 | 1.07 | 1 | 0.47 | 0.72 | 0.1971 | 0.2333 | 0.0362 |
| 1hx9 | noschrodinger | 3.31 | 0.0027 | 0.74 | 0.77 | 93.83 | 1.92 | 0 | 2.24 | 1.63 | 0.1768 | 0.2181 | 0.0413 |
| 1i36 | opls | 2 | 0.0994 | 2.51 | 0.41 | 96.69 | 2.22 | 4 | 1.36 | 1.34 | 0.1808 | 0.2193 | 0.0385 |
| 1i36 | noschrodinger | 2 | 0.0086 | 1.1 | 1.18 | 93.7 | 2.93 | 0 | 4.85 | 2.02 | 0.1781 | 0.2162 | 0.0381 |
| 1i5d | opls | 2.7 | 0.0918 | 2.62 | 0.61 | 96.95 | 9.55 | 1 | 1.65 | 1.84 | 0.3068 | 0.3421 | 0.0353 |
| 1i5d | noschrodinger | 2.7 | 0.0033 | 0.89 | 2.66 | 81.91 | 17.09 | 0 | 10.34 | 3.18 | 0.2894 | 0.3166 | 0.0272 |
| 1i69 | opls | 2.7 | 0.0902 | 2.31 | 1.01 | 94.97 | 3.82 | 1 | 1.69 | 1.71 | 0.2273 | 0.2599 | 0.0326 |
| 1i69 | noschrodinger | 2.7 | 0.0031 | 0.87 | 1.73 | 88.37 | 6.36 | 0 | 5.22 | 2.48 | 0.2117 | 0.2454 | 0.0337 |
| 1i72 | opls | 1.98 | 0.0988 | 2.26 | 0.34 | 96.98 | 0.36 | 2 | 0.98 | 0.97 | 0.1661 | 0.2025 | 0.0364 |
| 1i72 | noschrodinger | 1.98 | 0.0091 | 1 | 0.99 | 96.03 | 0.35 | 0 | 2.37 | 1.29 | 0.1651 | 0.2044 | 0.0393 |
| 1ia9 | opls | 2 | 0.096 | 2.3 | 0.37 | 97.44 | 6.02 | 1 | 1.55 | 1.6 | 0.2344 | 0.26 | 0.0256 |
| 1ia9 | noschrodinger | 2 | 0.0031 | 0.81 | 0.73 | 95.45 | 8.57 | 0 | 6.31 | 2.37 | 0.2238 | 0.2456 | 0.0218 |
| 1if2 | opls | 2 | 0.101 | 2.22 | 0 | 97.98 | 0.5 | 0 | 1.04 | 0.81 | 0.119 | 0.1477 | 0.0287 |
| 1if2 | noschrodinger | 2 | 0.0029 | 0.81 | 0 | 97.57 | 0.5 | 0 | 0.26 | 0.69 | 0.1189 | 0.1384 | 0.0195 |
| 1iim | opls | 2.01 | 0.0986 | 2.45 | 0.18 | 99.11 | 2.71 | 4 | 3.26 | 1.45 | 0.1778 | 0.2194 | 0.0416 |
| 1iim | noschrodinger | 2.01 | 0.0032 | 0.9 | 0.35 | 98.78 | 2.89 | 0 | 2.83 | 1.42 | 0.1877 | 0.2178 | 0.0301 |

| | | | | | | | | | | | | |
|---|---|---|---|---|---|---|---|---|---|---|---|---|
| 1ikt | opls | 1.75 | 0.1018 | 2.28 | 0 | 98.23 | 0 | 0 | 3.69 | 1.16 | 0.1775 | 0.2261 | 0.0486 |
| 1ikt | noschrodinger | 1.75 | 0.0029 | 0.84 | 0.88 | 96.46 | 0 | 0 | 2.65 | 1.28 | 0.1824 | 0.2199 | 0.0375 |
| 1ioo | opls | 1.55 | 0.1071 | 2.35 | 0 | 97.41 | 0.56 | 0 | 3 | 1.21 | 0.1517 | 0.1864 | 0.0347 |
| 1ioo | noschrodinger | 1.55 | 0.007 | 1.17 | 0.26 | 97.42 | 0.28 | 0 | 7.05 | 1.5 | 0.1597 | 0.1845 | 0.0248 |
| 1ipc | opls | 1.9 | 0.0996 | 2.36 | 0 | 96.69 | 1.82 | 1 | 0.97 | 1.2 | 0.1615 | 0.1931 | 0.0316 |
| 1ipc | noschrodinger | 1.9 | 0.0081 | 1.09 | 0 | 97.25 | 3.01 | 0 | 2.27 | 1.51 | 0.1623 | 0.1968 | 0.0345 |
| 1iwn | opls | 2.2 | 0.0964 | 2.26 | 0 | 98.29 | 1.31 | 1 | 1.74 | 1.02 | 0.2111 | 0.2514 | 0.0403 |
| 1iwn | noschrodinger | 2.2 | 0.0034 | 0.83 | 0 | 97.14 | 4.58 | 0 | 1.74 | 1.59 | 0.198 | 0.2435 | 0.0455 |
| 1ixr | opls | 3.19 | 0.0912 | 2.41 | 2.56 | 87.36 | 8.47 | 4 | 2.48 | 2.35 | 0.3076 | 0.3329 | 0.0253 |
| 1ixr | noschrodinger | 3.19 | 0.0035 | 0.93 | 7.96 | 70.06 | 14.83 | 0 | 17.89 | 3.48 | 0.302 | 0.319 | 0.017 |
| 1jay | opls | 1.65 | 0.098 | 2.42 | 0 | 98.09 | 1.14 | 4 | 1 | 0.84 | 0.1568 | 0.1868 | 0.03 |
| 1jay | noschrodinger | 1.65 | 0.0081 | 1.08 | 0 | 98.33 | 1.14 | 0 | 1 | 0.84 | 0.1576 | 0.1835 | 0.0259 |
| 1jd3 | opls | 2.03 | 0.096 | 2.22 | 0 | 98.14 | 2.08 | 1 | 0 | 0.74 | 0.1831 | 0.2067 | 0.0236 |
| 1jd3 | noschrodinger | 2.03 | 0.0082 | 1.2 | 0 | 97.53 | 6.21 | 0 | 1.11 | 1.52 | 0.1589 | 0.197 | 0.0381 |
| 1jjv | opls | 2.02 | 0.1019 | 2.33 | 0 | 97.83 | 1.2 | 1 | 0.63 | 0.81 | 0.1951 | 0.2212 | 0.0261 |
| 1jjv | noschrodinger | 2.02 | 0.0028 | 0.82 | 0.53 | 96.81 | 2.35 | 0 | 0.95 | 1.26 | 0.1715 | 0.2147 | 0.0432 |
| 1jnw | opls | 2.07 | 0.1028 | 2.26 | 0.5 | 96.5 | 0.54 | 1 | 1.38 | 1.1 | 0.1678 | 0.2203 | 0.0525 |
| 1jnw | noschrodinger | 2.07 | 0.0028 | 0.77 | 0.47 | 96.23 | 1.06 | 0 | 1.66 | 1.19 | 0.1689 | 0.2268 | 0.0579 |
| 1jr8 | opls | 1.5 | 0.1022 | 2.39 | 0 | 98.53 | 1.09 | 0 | 2.83 | 1.1 | 0.166 | 0.1949 | 0.0289 |
| 1jr8 | noschrodinger | 1.5 | 0.0097 | 1.29 | 0 | 98.06 | 0 | 0 | 5.66 | 1.31 | 0.1625 | 0.1964 | 0.0339 |
| 1jtk | opls | 2.04 | 0.0981 | 2.42 | 0 | 99.21 | 1.9 | 0 | 1 | 1.01 | 0.1798 | 0.2183 | 0.0385 |
| 1jtk | noschrodinger | 2.04 | 0.0034 | 0.79 | 0 | 99.22 | 2.8 | 0 | 2.5 | 1.37 | 0.1687 | 0.2132 | 0.0445 |
| 1juv | opls | 1.7 | 0.0968 | 2.5 | 0 | 99.47 | 2.98 | 0 | 5.1 | 1.63 | 0.207 | 0.2291 | 0.0221 |
| 1juv | noschrodinger | 1.7 | 0.013 | 1.5 | 0 | 99.48 | 2.37 | 0 | 4.42 | 1.5 | 0.2044 | 0.2319 | 0.0275 |
| 1jwa | opls | 2.93 | 0.09 | 2.39 | 0 | 94.44 | 4.33 | 0 | 1.1 | 1.68 | 0.1916 | 0.2494 | 0.0578 |
| 1jwa | noschrodinger | 2.93 | 0.003 | 0.83 | 0.68 | 92.12 | 10.68 | 0 | 2.21 | 2.26 | 0.1937 | 0.2298 | 0.0361 |
| 1jwb | opls | 2.1 | 0.0959 | 2.34 | 0 | 95.19 | 1.98 | 1 | 0.62 | 1.26 | 0.2023 | 0.2334 | 0.0311 |
| 1jwb | noschrodinger | 2.1 | 0.0031 | 0.8 | 0.32 | 93.95 | 3.16 | 0 | 3.52 | 1.93 | 0.1894 | 0.2267 | 0.0373 |
| 1jx0 | opls | 2.8 | 0.0937 | 3.12 | 0.49 | 92.68 | 6.53 | 38 | 7.12 | 2.47 | 0.2841 | 0.3238 | 0.0397 |
| 1jx0 | noschrodinger | 2.8 | 0.0031 | 0.81 | 0.48 | 89.42 | 7.08 | 0 | 5.42 | 2.5 | 0.3153 | 0.3291 | 0.0138 |
| 1jxi | opls | 2.5 | 0.0919 | 2.35 | 0 | 96.62 | 5.96 | 3 | 2.86 | 1.88 | 0.2244 | 0.251 | 0.0266 |
| 1jxi | noschrodinger | 2.5 | 0.003 | 0.78 | 1.01 | 93.95 | 8.76 | 0 | 3.27 | 2.24 | 0.2078 | 0.2475 | 0.0397 |
| 1jy8 | opls | 2.5 | 0.0954 | 2.44 | 0 | 98.01 | 0.88 | 0 | 0 | 0.5 | 0.1983 | 0.2144 | 0.0161 |
| 1jy8 | noschrodinger | 2.5 | 0.003 | 0.99 | 0 | 98.04 | 2.59 | 0 | 1.86 | 1.26 | 0.1804 | 0.2028 | 0.0224 |
| 1k12 | opls | 1.9 | 0.0997 | 2.27 | 0.65 | 94.81 | 1.56 | 1 | 3.87 | 1.68 | 0.1694 | 0.1998 | 0.0304 |
| 1k12 | noschrodinger | 1.9 | 0.0071 | 1 | 0.64 | 97.44 | 3.1 | 0 | 3.44 | 1.62 | 0.1716 | 0.2016 | 0.03 |
| 1kak | opls | 2.5 | 0.0964 | 2.3 | 0.68 | 94.58 | 2.22 | 2 | 1.03 | 1.44 | 0.1976 | 0.2337 | 0.0361 |
| 1kak | noschrodinger | 2.5 | 0.0028 | 0.78 | 1.69 | 91.86 | 2.59 | 0 | 5.17 | 2.08 | 0.1786 | 0.2228 | 0.0442 |
| 1kw8 | opls | 2 | 0.0976 | 2.23 | 0 | 95.77 | 0 | 0 | 0.23 | 0.88 | 0.1668 | 0.1853 | 0.0185 |
| 1kw8 | noschrodinger | 2 | 0.0031 | 0.81 | 0 | 96.5 | 0 | 0 | 1.81 | 1.17 | 0.1562 | 0.1763 | 0.0201 |
| 1kws | opls | 2.1 | 0.0979 | 2.27 | 0 | 97.75 | 0 | 1 | 0.65 | 0.77 | 0.1694 | 0.2032 | 0.0338 |
| 1kws | noschrodinger | 2.1 | 0.003 | 0.83 | 0 | 97.52 | 0.25 | 0 | 1.69 | 1.02 | 0.1616 | 0.1985 | 0.0369 |
| 1kyq | opls | 1.9 | 0.099 | 2.4 | 1.23 | 96.05 | 3.39 | 10 | 2.69 | 1.73 | 0.1812 | 0.2165 | 0.0353 |
| 1kyq | noschrodinger | 1.9 | 0.0085 | 1.02 | 1.65 | 93.39 | 5.19 | 0 | 6.45 | 2.33 | 0.1778 | 0.2117 | 0.0339 |
| 1l8j | opls | 2 | 0.0938 | 2.3 | 0.61 | 95.73 | 2.67 | 0 | 2.66 | 1.67 | 0.2014 | 0.2304 | 0.029 |
| 1l8j | noschrodinger | 2 | 0.0036 | 0.91 | 0.6 | 93.45 | 1.32 | 0 | 4.73 | 1.77 | 0.1981 | 0.2282 | 0.0301 |
| 1lc3 | opls | 1.5 | 0.1001 | 2.41 | 0 | 99.31 | 1.59 | 1 | 1.86 | 1.1 | 0.1698 | 0.2082 | 0.0384 |
| 1lc3 | noschrodinger | 1.5 | 0.004 | 0.98 | 0 | 98.62 | 0.79 | 0 | 1.86 | 0.95 | 0.1749 | 0.2103 | 0.0354 |
| 1lcs | opls | 2.5 | 0.0936 | 2.3 | 0 | 97.28 | 1.74 | 0 | 0.31 | 0.93 | 0.2386 | 0.2574 | 0.0188 |

| | | | | | | | | | | | | | |
|---|---|---|---|---|---|---|---|---|---|---|---|---|---|
| 1lcs | noschrodinger | 2.5 | 0.0032 | 0.86 | 0.25 | 95.81 | 3.18 | 0 | 4.45 | 1.89 | 0.1946 | 0.2399 | 0.0453 |
| 1lho | opls | 2 | 0.096 | 2.23 | 0 | 97.69 | 0.65 | 0 | 0.7 | 0.79 | 0.1858 | 0.2176 | 0.0318 |
| 1lho | noschrodinger | 2 | 0.0029 | 0.86 | 0.57 | 97.7 | 0.65 | 0 | 1.05 | 0.87 | 0.1716 | 0.2111 | 0.0395 |
| 1lkd | opls | 1.7 | 0.1024 | 2.36 | 0 | 97.53 | 0 | 1 | 1.35 | 0.96 | 0.171 | 0.1979 | 0.0269 |
| 1lkd | noschrodinger | 1.7 | 0.0078 | 1.06 | 0 | 97.19 | 0 | 0 | 3.16 | 1.25 | 0.1733 | 0.1984 | 0.0251 |
| 1lo7 | opls | 1.5 | 0.1025 | 2.42 | 0 | 95.65 | 1.57 | 3 | 2.53 | 1.49 | 0.1428 | 0.1684 | 0.0256 |
| 1lo7 | noschrodinger | 1.5 | 0.0103 | 1.38 | 0 | 95.65 | 1.57 | 0 | 2.95 | 1.54 | 0.1425 | 0.1655 | 0.023 |
| 1lo9 | opls | 2.8 | 0.0913 | 2.46 | 0 | 96.38 | 4.1 | 2 | 0.86 | 1.47 | 0.1801 | 0.233 | 0.0529 |
| 1lo9 | noschrodinger | 2.8 | 0.0029 | 0.82 | 0 | 93.48 | 9.84 | 0 | 5.19 | 2.46 | 0.1599 | 0.2135 | 0.0536 |
| 1lsl | opls | 1.9 | 0.1042 | 2.34 | 1.82 | 93.64 | 5.32 | 4 | 5.41 | 2.26 | 0.2408 | 0.2785 | 0.0377 |
| 1lsl | noschrodinger | 1.9 | 0.0031 | 0.92 | 0.9 | 95.5 | 8.42 | 0 | 8.43 | 2.47 | 0.2469 | 0.2846 | 0.0377 |
| 1lua | opls | 1.91 | 0.1002 | 2.2 | 0 | 97.77 | 0.8 | 1 | 0.31 | 0.67 | 0.1694 | 0.1962 | 0.0268 |
| 1lua | noschrodinger | 1.91 | 0.0031 | 0.81 | 0 | 97.78 | 0.96 | 0 | 0.77 | 0.79 | 0.1747 | 0.1959 | 0.0212 |
| 1m49 | opls | 2 | 0.0938 | 2.87 | 0 | 99.1 | 0.9 | 0 | 1.21 | 0.84 | 0.2175 | 0.2578 | 0.0403 |
| 1m49 | noschrodinger | 2 | 0.0031 | 0.77 | 0 | 99.12 | 2.19 | 0 | 1.21 | 1.1 | 0.2152 | 0.2594 | 0.0442 |
| 1m4i | opls | 1.5 | 0.1009 | 2.49 | 0 | 99.15 | 1.79 | 5 | 2.4 | 1.21 | 0.1469 | 0.1704 | 0.0235 |
| 1m4i | noschrodinger | 1.5 | 0.0139 | 1.5 | 0 | 98.87 | 2.14 | 0 | 2.4 | 1.27 | 0.1481 | 0.173 | 0.0249 |
| 1mg9 | opls | 1.85 | 0.0916 | 2.36 | 0.46 | 97.71 | 8.21 | 4 | 1.64 | 1.67 | 0.2488 | 0.2774 | 0.0286 |
| 1mg9 | noschrodinger | 1.85 | 0.0069 | 1.13 | 1.81 | 97.29 | 8.12 | 0 | 5.22 | 2.1 | 0.2457 | 0.2791 | 0.0334 |
| 1mjh | opls | 1.8 | 0.1006 | 2.25 | 0 | 98.55 | 0.38 | 2 | 1.24 | 0.84 | 0.1756 | 0.2076 | 0.032 |
| 1mjh | noschrodinger | 1.8 | 0.0083 | 1.06 | 0 | 97.85 | 0.38 | 0 | 2.7 | 1.09 | 0.1767 | 0.2024 | 0.0257 |
| 1moy | opls | 1.55 | 0.0969 | 2.53 | 0 | 99.17 | 0 | 1 | 8.76 | 1.47 | 0.1522 | 0.1781 | 0.0259 |
| 1moy | noschrodinger | 1.55 | 0.0099 | 1.21 | 0 | 99.17 | 0 | 0 | 3.61 | 1.15 | 0.154 | 0.1721 | 0.0181 |
| 1mvl | opls | 2 | 0.1012 | 2.48 | 0 | 100 | 2.7 | 3 | 2.15 | 1.32 | 0.1446 | 0.1874 | 0.0428 |
| 1mvl | noschrodinger | 2 | 0.0032 | 0.85 | 0 | 98.82 | 4 | 0 | 1.79 | 1.39 | 0.1567 | 0.1871 | 0.0304 |
| 1mzu | opls | 2 | 0.0947 | 2.32 | 0 | 95.64 | 1.52 | 2 | 1.56 | 1.34 | 0.2526 | 0.287 | 0.0344 |
| 1mzu | noschrodinger | 2 | 0.0027 | 0.79 | 0 | 95.3 | 2.63 | 0 | 2.16 | 1.64 | 0.2395 | 0.279 | 0.0395 |
| 1n06 | opls | 2 | 0.0961 | 2.28 | 0.33 | 97.36 | 1.85 | 0 | 0.59 | 1.02 | 0.2184 | 0.2547 | 0.0363 |
| 1n06 | noschrodinger | 2 | 0.0038 | 0.88 | 0.66 | 96.72 | 2.56 | 0 | 1.97 | 1.48 | 0.2039 | 0.252 | 0.0481 |
| 1n08 | opls | 1.6 | 0.0998 | 2.22 | 0 | 97.03 | 1.85 | 1 | 0.59 | 1.07 | 0.1866 | 0.2024 | 0.0158 |
| 1n08 | noschrodinger | 1.6 | 0.003 | 0.88 | 0.33 | 97.7 | 1.83 | 0 | 0.98 | 1.06 | 0.1867 | 0.2014 | 0.0147 |
| 1n0s | opls | 2 | 0.0932 | 2.46 | 0 | 96.75 | 5.78 | 7 | 3.1 | 1.88 | 0.1737 | 0.215 | 0.0413 |
| 1n0s | noschrodinger | 2 | 0.0133 | 1.26 | 0 | 96.49 | 8.72 | 2 | 5.1 | 2.22 | 0.1778 | 0.2132 | 0.0354 |
| 1n52 | opls | 2 | 0.0957 | 3.1 | 0.69 | 96.66 | 2.27 | 3 | 1.63 | 1.39 | 0.2361 | 0.2683 | 0.0322 |
| 1n52 | noschrodinger | 2 | 0.0028 | 0.75 | 0.11 | 94.04 | 3.39 | 0 | 4.11 | 2 | 0.2438 | 0.2765 | 0.0327 |
| 1n5q | opls | 1.74 | 0.0991 | 2.48 | 0 | 99.54 | 2.25 | 1 | 4.61 | 1.5 | 0.1677 | 0.1994 | 0.0317 |
| 1n5q | noschrodinger | 1.74 | 0.0062 | 1.08 | 0 | 98.64 | 3.93 | 0 | 9.53 | 1.95 | 0.1657 | 0.1959 | 0.0302 |
| 1n5v | opls | 2.24 | 0.0937 | 2.38 | 0 | 97.71 | 2.82 | 0 | 0.89 | 1.18 | 0.2072 | 0.2401 | 0.0329 |
| 1n5v | noschrodinger | 2.24 | 0.0031 | 0.88 | 0 | 98.18 | 5.08 | 0 | 3.85 | 1.71 | 0.1894 | 0.2319 | 0.0425 |
| 1nae | opls | 2.05 | 0.1054 | 2.14 | 0 | 96.83 | 0 | 0 | 1.56 | 1.1 | 0.1721 | 0.2109 | 0.0388 |
| 1nae | noschrodinger | 2.05 | 0.0028 | 0.87 | 0.79 | 96.85 | 1.94 | 0 | 3.13 | 1.51 | 0.1409 | 0.1977 | 0.0568 |
| 1ne8 | opls | 2.1 | 0.0979 | 2.26 | 0 | 97.17 | 0 | 0 | 0.52 | 0.83 | 0.174 | 0.197 | 0.023 |
| 1ne8 | noschrodinger | 2.1 | 0.0032 | 0.8 | 0 | 97.37 | 0 | 0 | 0.53 | 0.8 | 0.1564 | 0.182 | 0.0256 |
| 1nni | opls | 2.43 | 0.0983 | 2.28 | 0.65 | 98.04 | 2.24 | 0 | 0.37 | 0.9 | 0.2105 | 0.2318 | 0.0213 |
| 1nni | noschrodinger | 2.43 | 0.0032 | 0.8 | 1.18 | 97.04 | 2.13 | 0 | 2.95 | 1.5 | 0.2252 | 0.2428 | 0.0176 |
| 1nox | opls | 1.6 | 0.0965 | 2.36 | 0 | 99.49 | 0.64 | 1 | 0.62 | 0.7 | 0.1484 | 0.1777 | 0.0293 |
| 1nox | noschrodinger | 1.6 | 0.0142 | 1.43 | 0 | 99.49 | 0.64 | 0 | 1.23 | 0.84 | 0.1495 | 0.178 | 0.0285 |
| 1nxj | opls | 1.9 | 0.0984 | 2.51 | 0 | 99.35 | 0 | 4 | 2.63 | 1.05 | 0.1568 | 0.1834 | 0.0266 |
| 1nxj | noschrodinger | 1.9 | 0.0063 | 1.21 | 0 | 100 | 0.56 | 0 | 1.91 | 0.95 | 0.1626 | 0.1878 | 0.0252 |

| | | | | | | | | | | | | | |
|---|---|---|---|---|---|---|---|---|---|---|---|---|---|
| 1o91 | opls | 1.9 | 0.098 | 2.18 | 0.78 | 94.57 | 0.31 | 0 | 0.81 | 1.13 | 0.148 | 0.1726 | 0.0246 |
| 1o91 | noschrodinger | 1.9 | 0.0062 | 1.14 | 0.78 | 95.09 | 0 | 0 | 2.48 | 1.37 | 0.149 | 0.1726 | 0.0236 |
| 1oc3 | opls | 2 | 0.0927 | 2.28 | 0.21 | 94.56 | 2.11 | 0 | 0.69 | 1.34 | 0.2346 | 0.2479 | 0.0133 |
| 1oc3 | noschrodinger | 2 | 0.003 | 0.85 | 0.42 | 94.35 | 4.75 | 0 | 1.37 | 1.77 | 0.2265 | 0.2458 | 0.0193 |
| 1od6 | opls | 1.5 | 0.1018 | 3.14 | 0 | 97.97 | 1.55 | 2 | 1.56 | 1.05 | 0.1716 | 0.2023 | 0.0307 |
| 1od6 | noschrodinger | 1.5 | 0.0138 | 1.38 | 0.66 | 97.35 | 3.82 | 0 | 3.12 | 1.67 | 0.1776 | 0.2039 | 0.0263 |
| 1og4 | opls | 2.6 | 0.0977 | 2.31 | 0.45 | 95.93 | 1.03 | 3 | 1.92 | 1.25 | 0.1971 | 0.2265 | 0.0294 |
| 1og4 | noschrodinger | 2.6 | 0.0028 | 0.84 | 0.9 | 93.67 | 1.03 | 0 | 4.39 | 1.65 | 0.1758 | 0.2088 | 0.033 |
| 1ogx | opls | 2.02 | 0.0883 | 2.43 | 0.51 | 95.92 | 4.95 | 2 | 1.28 | 1.66 | 0.1552 | 0.1887 | 0.0335 |
| 1ogx | noschrodinger | 2.02 | 0.0096 | 1.16 | 1.21 | 94.35 | 9 | 0 | 2.57 | 2.15 | 0.1561 | 0.1874 | 0.0313 |
| 1okc | opls | 2.2 | 0.0888 | 2.39 | 0.36 | 98.56 | 4.04 | 4 | 1.88 | 1.41 | 0.2002 | 0.2349 | 0.0347 |
| 1okc | noschrodinger | 2.2 | 0.0037 | 0.89 | 1.38 | 97.59 | 4.04 | 0 | 4.25 | 1.75 | 0.1983 | 0.2332 | 0.0349 |
| 1okk | opls | 2.05 | 0.1016 | 2.25 | 0.18 | 99.45 | 1.36 | 0 | 0.67 | 0.82 | 0.1524 | 0.1843 | 0.0319 |
| 1okk | noschrodinger | 2.05 | 0.0032 | 0.8 | 0.18 | 98.72 | 0.9 | 0 | 1.23 | 0.84 | 0.1467 | 0.1763 | 0.0296 |
| 1omz | opls | 2.1 | 0.0956 | 2.21 | 0.42 | 95.76 | 1.6 | 1 | 0.74 | 1.18 | 0.2006 | 0.2252 | 0.0246 |
| 1omz | noschrodinger | 2.1 | 0.0056 | 0.95 | 0.2 | 96.2 | 2.28 | 0 | 1.98 | 1.49 | 0.1782 | 0.2146 | 0.0364 |
| 1on8 | opls | 2.7 | 0.0918 | 2.27 | 0.81 | 95.73 | 3.74 | 1 | 0.71 | 1.46 | 0.1987 | 0.2285 | 0.0298 |
| 1on8 | noschrodinger | 2.7 | 0.0033 | 0.83 | 0.77 | 94.63 | 5.47 | 0 | 1.42 | 1.81 | 0.177 | 0.2223 | 0.0453 |
| 1oo5 | opls | 2.5 | 0.0918 | 2.3 | 0.23 | 97.9 | 7.58 | 4 | 0.88 | 1.46 | 0.2242 | 0.2616 | 0.0374 |
| 1oo5 | noschrodinger | 2.5 | 0.0028 | 0.76 | 0.23 | 96.98 | 11.73 | 0 | 2.35 | 2 | 0.2122 | 0.2473 | 0.0351 |
| 1otw | opls | 2.3 | 0.0941 | 2.31 | 0.2 | 97.42 | 1.65 | 1 | 1.73 | 1.21 | 0.2196 | 0.245 | 0.0254 |
| 1otw | noschrodinger | 2.3 | 0.0029 | 0.78 | 0.2 | 96.23 | 5.63 | 0 | 1.88 | 1.78 | 0.2088 | 0.2426 | 0.0338 |
| 1ow3 | opls | 1.8 | 0.1015 | 2.4 | 0 | 98.1 | 1.51 | 4 | 3.47 | 1.27 | 0.1458 | 0.1852 | 0.0394 |
| 1ow3 | noschrodinger | 1.8 | 0.004 | 0.98 | 0 | 98.65 | 2.1 | 0 | 4.47 | 1.47 | 0.157 | 0.1809 | 0.0239 |
| 1ow4 | opls | 1.7 | 0.1013 | 3.92 | 0 | 99.57 | 4.25 | 1 | 12.9 | 2.1 | 0.1709 | 0.2083 | 0.0374 |
| 1ow4 | noschrodinger | 1.7 | 0.0098 | 1.49 | 0 | 99.57 | 4.25 | 0 | 7.53 | 1.89 | 0.1719 | 0.2027 | 0.0308 |
| 1p3y | opls | 2.54 | 0.0942 | 2.34 | 0 | 98.18 | 4.08 | 0 | 1.07 | 1.27 | 0.2401 | 0.2705 | 0.0304 |
| 1p3y | noschrodinger | 2.54 | 0.0027 | 0.75 | 0 | 95.81 | 4.03 | 0 | 3.25 | 1.87 | 0.2121 | 0.2439 | 0.0318 |
| 1p4t | opls | 2.55 | 0.0874 | 2.16 | 0 | 97.26 | 3.36 | 0 | 1.18 | 1.37 | 0.2363 | 0.258 | 0.0217 |
| 1p4t | noschrodinger | 2.55 | 0.0027 | 0.78 | 1.31 | 96.08 | 5.88 | 0 | 0.4 | 1.5 | 0.2156 | 0.2419 | 0.0263 |
| 1p5z | opls | 1.6 | 0.0985 | 2.29 | 0 | 97.67 | 0.48 | 0 | 0.78 | 0.82 | 0.1639 | 0.2025 | 0.0386 |
| 1p5z | noschrodinger | 1.6 | 0.0073 | 1.02 | 0 | 96.89 | 2.4 | 0 | 1.05 | 1.28 | 0.1697 | 0.1995 | 0.0298 |
| 1pa9 | opls | 2 | 0.1005 | 2.3 | 0.36 | 95.62 | 3.81 | 4 | 1.62 | 1.66 | 0.186 | 0.225 | 0.039 |
| 1pa9 | noschrodinger | 2 | 0.0042 | 0.86 | 0.36 | 96 | 3.81 | 0 | 3.48 | 1.85 | 0.1773 | 0.2189 | 0.0416 |
| 1pge | opls | 3.5 | 0.0897 | 2.29 | 0.09 | 94.98 | 7.22 | 3 | 1.59 | 1.91 | 0.1726 | 0.2113 | 0.0387 |
| 1pge | noschrodinger | 3.5 | 0.0039 | 0.91 | 0.46 | 92.44 | 11.42 | 0 | 5.21 | 2.55 | 0.1509 | 0.1947 | 0.0438 |
| 1pl3 | opls | 1.9 | 0.0989 | 2.24 | 0 | 98.36 | 0 | 4 | 1.91 | 0.96 | 0.1734 | 0.2061 | 0.0327 |
| 1pl3 | noschrodinger | 1.9 | 0.0057 | 0.99 | 0 | 97.54 | 0.67 | 0 | 3.65 | 1.25 | 0.175 | 0.2067 | 0.0317 |
| 1pno | opls | 2.1 | 0.0974 | 2.35 | 0 | 98.3 | 3.2 | 0 | 0.73 | 1.12 | 0.193 | 0.2143 | 0.0213 |
| 1pno | noschrodinger | 2.1 | 0.0041 | 0.92 | 0 | 97.17 | 4.61 | 0 | 0.91 | 1.43 | 0.1864 | 0.209 | 0.0226 |
| 1pub | opls | 2.51 | 0.0928 | 2.27 | 0.63 | 94.97 | 2.76 | 2 | 0.39 | 1.32 | 0.2433 | 0.2627 | 0.0194 |
| 1pub | noschrodinger | 2.51 | 0.0048 | 0.95 | 2.5 | 89.38 | 6.85 | 0 | 5.95 | 2.53 | 0.2416 | 0.2564 | 0.0148 |
| 1puc | opls | 1.92 | 0.0971 | 2.31 | 0 | 97.94 | 1.1 | 1 | 0 | 0.55 | 0.1766 | 0.2061 | 0.0295 |
| 1puc | noschrodinger | 1.92 | 0.0109 | 1.64 | 0 | 97.98 | 1.08 | 0 | 6.29 | 1.38 | 0.1643 | 0.2005 | 0.0362 |
| 1puj | opls | 2 | 0.098 | 2.29 | 0.78 | 96.47 | 0.89 | 0 | 2.09 | 1.21 | 0.1905 | 0.22 | 0.0295 |
| 1puj | noschrodinger | 2 | 0.0032 | 0.93 | 0.78 | 94.94 | 1.77 | 0 | 2.56 | 1.58 | 0.1884 | 0.2181 | 0.0297 |
| 1pvs | opls | 2.39 | 0.0959 | 2.27 | 0 | 98.75 | 2.49 | 3 | 0.68 | 1.02 | 0.2141 | 0.2411 | 0.027 |
| 1pvs | noschrodinger | 2.39 | 0.0033 | 0.79 | 0.18 | 97.68 | 2.93 | 0 | 0.9 | 1.2 | 0.2004 | 0.2383 | 0.0379 |
| 1pwq | opls | 3.52 | 0.0907 | 2.32 | 0.82 | 93.06 | 5.66 | 9 | 1.24 | 1.86 | 0.1911 | 0.2402 | 0.0491 |

| | | | | | | | | | | | |
|---|---|---|---|---|---|---|---|---|---|---|---|
| 1pwq | noschrodinger | 3.52 | 0.003 | 0.8 | 3.5 | 80.73 | 9.3 | 0 | 6.94 | 2.84 | 0.1852 | 0.2286 | 0.0434 |
| 1q0z | opls | 1.97 | 0.0995 | 2.56 | 0 | 97.96 | 2.61 | 1 | 2.41 | 1.35 | 0.1432 | 0.186 | 0.0428 |
| 1q0z | noschrodinger | 1.97 | 0.0036 | 1.12 | 0 | 97.63 | 1.74 | 0 | 1.54 | 1.16 | 0.1475 | 0.182 | 0.0345 |
| 1q1a | opls | 1.5 | 0.1011 | 2.46 | 0 | 98.58 | 1.18 | 2 | 2.93 | 1.14 | 0.148 | 0.1759 | 0.0279 |
| 1q1a | noschrodinger | 1.5 | 0.0113 | 1.26 | 0 | 97.55 | 0.78 | 0 | 5.88 | 1.41 | 0.1475 | 0.177 | 0.0295 |
| 1q2o | opls | 1.74 | 0.0979 | 2.3 | 0 | 98.25 | 1.02 | 2 | 3.34 | 1.13 | 0.157 | 0.1797 | 0.0227 |
| 1q2o | noschrodinger | 1.74 | 0.0068 | 1.15 | 0 | 97.88 | 1.16 | 0 | 4.19 | 1.28 | 0.1581 | 0.1811 | 0.023 |
| 1q33 | opls | 1.8 | 0.0984 | 2.31 | 0 | 96.35 | 1.63 | 2 | 1.06 | 1.21 | 0.1751 | 0.2052 | 0.0301 |
| 1q33 | noschrodinger | 1.8 | 0.0122 | 1.2 | 0.34 | 95.17 | 1.59 | 0 | 2.34 | 1.5 | 0.1734 | 0.2036 | 0.0302 |
| 1q4u | opls | 1.6 | 0.1008 | 2.51 | 0 | 99.27 | 3.54 | 0 | 2.45 | 1.45 | 0.1595 | 0.187 | 0.0275 |
| 1q4u | noschrodinger | 1.6 | 0.0144 | 1.62 | 0 | 99.27 | 3.07 | 0 | 1.56 | 1.27 | 0.1614 | 0.1857 | 0.0243 |
| 1q6i | opls | 2.25 | 0.0956 | 2.19 | 0 | 98.76 | 1.47 | 1 | 0.59 | 0.83 | 0.2059 | 0.242 | 0.0361 |
| 1q6i | noschrodinger | 2.25 | 0.0029 | 0.82 | 0 | 98.08 | 3.48 | 0 | 1.77 | 1.35 | 0.1941 | 0.2326 | 0.0385 |
| 1qb7 | opls | 1.41 | 0.101 | 2.87 | 0.86 | 98.71 | 4.31 | 1 | 4 | 1.67 | 0.1661 | 0.1959 | 0.0298 |
| 1qb7 | noschrodinger | 1.41 | 0.0064 | 1.11 | 0.43 | 98.72 | 2.86 | 0 | 2.95 | 1.43 | 0.1708 | 0.193 | 0.0222 |
| 1qb8 | opls | 2 | 0.0993 | 2.29 | 0 | 98.28 | 1.98 | 1 | 1.62 | 1.14 | 0.1886 | 0.2133 | 0.0247 |
| 1qb8 | noschrodinger | 2 | 0.0028 | 0.84 | 0.43 | 97.44 | 1.97 | 0 | 1.91 | 1.29 | 0.1845 | 0.2134 | 0.0289 |
| 1qfo | opls | 1.85 | 0.0958 | 2.23 | 0 | 97.39 | 1.37 | 1 | 0.18 | 0.79 | 0.1745 | 0.2095 | 0.035 |
| 1qfo | noschrodinger | 1.85 | 0.013 | 1.28 | 0 | 97.26 | 0.99 | 0 | 3.05 | 1.23 | 0.1701 | 0.2028 | 0.0327 |
| 1qh6 | opls | 2 | 0.1047 | 2.32 | 0 | 98.04 | 2.23 | 1 | 0.62 | 0.97 | 0.1515 | 0.177 | 0.0255 |
| 1qh6 | noschrodinger | 2 | 0.003 | 0.85 | 0 | 97.3 | 1.11 | 0 | 1.85 | 1.11 | 0.1339 | 0.1628 | 0.0289 |
| 1qjp | opls | 1.65 | 0.0926 | 3.03 | 0 | 99.2 | 9.17 | 0 | 2.42 | 1.76 | 0.2113 | 0.238 | 0.0267 |
| 1qjp | noschrodinger | 1.65 | 0.0138 | 1.46 | 0 | 99.22 | 5.36 | 0 | 11.41 | 2.13 | 0.2028 | 0.2336 | 0.0308 |
| 1r6n | opls | 2.42 | 0.0902 | 2.25 | 0 | 94.82 | 1.13 | 0 | 0.91 | 1.17 | 0.2005 | 0.2233 | 0.0228 |
| 1r6n | noschrodinger | 2.42 | 0.0026 | 0.8 | 0.52 | 94.33 | 3.37 | 0 | 1.53 | 1.68 | 0.1875 | 0.2136 | 0.0261 |
| 1rbc | opls | 1.77 | 0.0955 | 2.33 | 0 | 97.32 | 2.73 | 3 | 5.84 | 1.78 | 0.1693 | 0.2065 | 0.0372 |
| 1rbc | noschrodinger | 1.77 | 0.0037 | 0.84 | 0 | 94.78 | 6.19 | 0 | 2.66 | 2.01 | 0.1769 | 0.2078 | 0.0309 |
| 1rgl | opls | 1.92 | 0.0983 | 2.16 | 0 | 98.98 | 3.57 | 0 | 0 | 0.92 | 0.1342 | 0.1849 | 0.0507 |
| 1rgl | noschrodinger | 1.92 | 0.0031 | 0.9 | 0 | 99.02 | 4.71 | 0 | 1.34 | 1.37 | 0.134 | 0.1833 | 0.0493 |
| 1rhu | opls | 2.51 | 0.0916 | 2.29 | 0 | 98.96 | 1.03 | 1 | 2.37 | 1.03 | 0.168 | 0.211 | 0.043 |
| 1rhu | noschrodinger | 2.51 | 0.0029 | 0.91 | 0 | 98.24 | 2.9 | 0 | 2.11 | 1.33 | 0.1515 | 0.2026 | 0.0511 |
| 1rki | opls | 1.6 | 0.0944 | 2.89 | 0 | 97.89 | 3.63 | 1 | 1.17 | 1.28 | 0.1812 | 0.2124 | 0.0312 |
| 1rki | noschrodinger | 1.6 | 0.0087 | 0.99 | 0 | 98.45 | 5.58 | 0 | 4.98 | 1.83 | 0.1858 | 0.2092 | 0.0234 |
| 1rlj | opls | 2 | 0.1008 | 2.37 | 0 | 96.18 | 2.54 | 2 | 0.91 | 1.34 | 0.1721 | 0.1905 | 0.0184 |
| 1rlj | noschrodinger | 2 | 0.0026 | 0.77 | 0 | 97.74 | 4.17 | 0 | 0.91 | 1.3 | 0.1717 | 0.1896 | 0.0179 |
| 1rwn | opls | 2 | 0.102 | 2.29 | 0 | 97.65 | 1.72 | 1 | 1.91 | 1.21 | 0.19 | 0.2296 | 0.0396 |
| 1rwn | noschrodinger | 2 | 0.0029 | 0.8 | 0.39 | 96.11 | 7.73 | 0 | 1.19 | 1.77 | 0.1777 | 0.2213 | 0.0436 |
| 1s3z | opls | 2 | 0.097 | 2.41 | 0 | 99.65 | 3.31 | 1 | 2.67 | 1.45 | 0.1535 | 0.1989 | 0.0454 |
| 1s3z | noschrodinger | 2 | 0.0034 | 0.94 | 0 | 99.65 | 4.13 | 0 | 1.64 | 1.38 | 0.1574 | 0.1937 | 0.0363 |
| 1s4m | opls | 2 | 0.0955 | 2.25 | 0.38 | 98.11 | 0.63 | 4 | 0.99 | 0.79 | 0.2095 | 0.2427 | 0.0332 |
| 1s4m | noschrodinger | 2 | 0.0034 | 0.79 | 0.93 | 96.28 | 3.7 | 0 | 4.32 | 1.89 | 0.1942 | 0.2366 | 0.0424 |
| 1s68 | opls | 1.89 | 0.1015 | 2.37 | 0 | 96.51 | 3.48 | 4 | 5.11 | 1.91 | 0.1511 | 0.1949 | 0.0438 |
| 1s68 | noschrodinger | 1.89 | 0.0063 | 0.99 | 0.43 | 95.67 | 2.96 | 0 | 7.53 | 2.07 | 0.154 | 0.196 | 0.042 |
| 1s7f | opls | 1.99 | 0.0978 | 2.3 | 0 | 100 | 3.95 | 0 | 0.69 | 1.18 | 0.1844 | 0.227 | 0.0426 |
| 1s7f | noschrodinger | 1.99 | 0.0032 | 0.85 | 0 | 99.44 | 7.14 | 0 | 3.44 | 1.78 | 0.1865 | 0.2271 | 0.0406 |
| 1sbr | opls | 2.3 | 0.0902 | 2.3 | 0.33 | 98.7 | 1.78 | 0 | 1.78 | 1.13 | 0.2105 | 0.2421 | 0.0316 |
| 1sbr | noschrodinger | 2.3 | 0.0033 | 0.79 | 0.28 | 95.22 | 7.72 | 0 | 3.05 | 2.1 | 0.2167 | 0.2413 | 0.0246 |
| 1sfj | opls | 2.4 | 0.0971 | 2.31 | 0.45 | 97.29 | 6.6 | 0 | 1.35 | 1.62 | 0.223 | 0.2596 | 0.0366 |
| 1sfj | noschrodinger | 2.4 | 0.0029 | 0.75 | 1.34 | 94.2 | 6.76 | 0 | 4.88 | 2.28 | 0.2057 | 0.2474 | 0.0417 |

| | | | | | | | | | | | | | |
|---|---|---|---|---|---|---|---|---|---|---|---|---|---|
| 1sm4 | opls | 2 | 0.0986 | 2.44 | 0.34 | 96.59 | 6.75 | 3 | 1.89 | 1.8 | 0.227 | 0.2397 | 0.0127 |
| 1sm4 | noschrodinger | 2 | 0.0029 | 0.83 | 0.85 | 97.11 | 5.14 | 0 | 3.46 | 1.84 | 0.2243 | 0.2438 | 0.0195 |
| 1t27 | opls | 2.1 | 0.0956 | 2.25 | 0 | 96.62 | 3.32 | 0 | 1.09 | 1.43 | 0.1918 | 0.2373 | 0.0455 |
| 1t27 | noschrodinger | 2.1 | 0.0026 | 0.76 | 0 | 94.38 | 3.31 | 0 | 2.63 | 1.83 | 0.1778 | 0.2261 | 0.0483 |
| 1t41 | opls | 1 | 0.0995 | 3.26 | 0.36 | 98.93 | 4.23 | 8 | 10.3 | 2.01 | 0.1113 | 0.1232 | 0.0119 |
| 1t41 | noschrodinger | 1 | 0.0079 | 1.24 | 0 | 99.36 | 3.41 | 0 | 5.55 | 1.71 | 0.108 | 0.1143 | 0.0063 |
| 1t44 | opls | 2 | 0.1002 | 2.22 | 0 | 98.38 | 4.02 | 3 | 0.89 | 1.23 | 0.1667 | 0.1927 | 0.026 |
| 1t44 | noschrodinger | 2 | 0.0029 | 0.86 | 0 | 98.99 | 5.87 | 0 | 1.01 | 1.38 | 0.1614 | 0.1865 | 0.0251 |
| 1t57 | opls | 2.3 | 0.0915 | 2.44 | 0.4 | 95.56 | 7.8 | 0 | 2.33 | 2 | 0.2672 | 0.3001 | 0.0329 |
| 1t57 | noschrodinger | 2.3 | 0.0032 | 0.82 | 1.09 | 88.95 | 10.81 | 0 | 6.52 | 2.72 | 0.2701 | 0.3046 | 0.0345 |
| 1t6y | opls | 2.8 | 0.0905 | 2.35 | 0 | 97.49 | 3.41 | 5 | 0.77 | 1.25 | 0.2362 | 0.283 | 0.0468 |
| 1t6y | noschrodinger | 2.8 | 0.003 | 0.83 | 0.76 | 91.79 | 7.56 | 0 | 6.19 | 2.5 | 0.2456 | 0.2868 | 0.0412 |
| 1tc2 | opls | 1.81 | 0.0961 | 2.3 | 0 | 97.96 | 1.85 | 2 | 1.29 | 1.07 | 0.1634 | 0.1955 | 0.0321 |
| 1tc2 | noschrodinger | 1.81 | 0.0036 | 0.9 | 0 | 98.15 | 2.77 | 0 | 1.62 | 1.25 | 0.1662 | 0.197 | 0.0308 |
| 1te2 | opls | 1.76 | 0.1047 | 2.33 | 0 | 99.74 | 1.39 | 0 | 3.75 | 1.27 | 0.1752 | 0.2109 | 0.0357 |
| 1te2 | noschrodinger | 1.76 | 0.0049 | 0.84 | 0 | 99.07 | 1.6 | 0 | 3.6 | 1.3 | 0.1742 | 0.2081 | 0.0339 |
| 1tgj | opls | 2 | 0.0978 | 2.4 | 0 | 96.33 | 1.9 | 0 | 0 | 0.96 | 0.1623 | 0.2155 | 0.0532 |
| 1tgj | noschrodinger | 2 | 0.0028 | 0.8 | 0 | 97.27 | 0.96 | 0 | 0 | 0.64 | 0.1599 | 0.2038 | 0.0439 |
| 1th8 | opls | 2.3 | 0.0942 | 2.34 | 0 | 98.32 | 1.91 | 0 | 0.77 | 0.96 | 0.1992 | 0.2399 | 0.0407 |
| 1th8 | noschrodinger | 2.3 | 0.0026 | 0.79 | 0 | 98.76 | 2.36 | 0 | 0.78 | 1.03 | 0.1922 | 0.2273 | 0.0351 |
| 1thq | opls | 1.9 | 0.0888 | 2.19 | 0 | 100 | 0 | 0 | 1.14 | 0.82 | 0.1971 | 0.2428 | 0.0457 |
| 1thq | noschrodinger | 1.9 | 0.0094 | 1 | 0 | 99.3 | 1.67 | 0 | 1.92 | 1.12 | 0.2004 | 0.2349 | 0.0345 |
| 1tlw | opls | 3 | 0.0893 | 2.27 | 0.62 | 94.87 | 4.08 | 3 | 2.38 | 1.84 | 0.2589 | 0.2703 | 0.0114 |
| 1tlw | noschrodinger | 3 | 0.0055 | 1 | 1.63 | 90.63 | 4.75 | 0 | 10.68 | 2.59 | 0.2538 | 0.27 | 0.0162 |
| 1tu1 | opls | 1.95 | 0.1009 | 2.43 | 0 | 98.93 | 3.49 | 3 | 5.24 | 1.69 | 0.1903 | 0.2313 | 0.041 |
| 1tu1 | noschrodinger | 1.95 | 0.0051 | 0.91 | 0 | 98.59 | 2.69 | 0 | 4.03 | 1.51 | 0.1941 | 0.2293 | 0.0352 |
| 1tuv | opls | 1.7 | 0.1076 | 2.58 | 0 | 97.98 | 3.53 | 3 | 4.32 | 1.63 | 0.2063 | 0.2411 | 0.0348 |
| 1tuv | noschrodinger | 1.7 | 0.0121 | 1.34 | 0.99 | 97.03 | 6.98 | 0 | 6.19 | 2.15 | 0.206 | 0.2375 | 0.0315 |
| 1tvp | opls | 1.6 | 0.1035 | 2.2 | 0 | 94.45 | 0 | 4 | 3.05 | 1.47 | 0.1303 | 0.1667 | 0.0364 |
| 1tvp | noschrodinger | 1.6 | 0.0042 | 0.89 | 0 | 94.31 | 0 | 0 | 1.92 | 1.34 | 0.1359 | 0.1596 | 0.0237 |
| 1txz | opls | 2.05 | 0.0939 | 2.31 | 0.42 | 96.62 | 1.41 | 0 | 0.25 | 0.92 | 0.181 | 0.2132 | 0.0322 |
| 1txz | noschrodinger | 2.05 | 0.0028 | 0.82 | 0.41 | 96.27 | 1.85 | 0 | 1.49 | 1.34 | 0.1747 | 0.2063 | 0.0316 |
| 1tzf | opls | 2.1 | 0.0972 | 2.26 | 0 | 95.92 | 1.91 | 0 | 2.23 | 1.5 | 0.196 | 0.2228 | 0.0268 |
| 1tzf | noschrodinger | 2.1 | 0.003 | 0.8 | 0.4 | 96.36 | 4.76 | 0 | 1.73 | 1.69 | 0.1966 | 0.2173 | 0.0207 |
| 1u3g | opls | 2.5 | 0.0958 | 2.44 | 1.23 | 96.3 | 4.64 | 1 | 3.19 | 1.86 | 0.2192 | 0.2591 | 0.0399 |
| 1u3g | noschrodinger | 2.5 | 0.0027 | 0.83 | 1.85 | 93.83 | 5.96 | 1 | 3.56 | 2.15 | 0.1936 | 0.2392 | 0.0456 |
| 1u7z | opls | 2.3 | 0.0947 | 2.3 | 0.16 | 96.69 | 2.8 | 1 | 1.74 | 1.48 | 0.203 | 0.2432 | 0.0402 |
| 1u7z | noschrodinger | 2.3 | 0.0031 | 0.83 | 0.31 | 97.04 | 4.43 | 0 | 4.26 | 1.87 | 0.1891 | 0.231 | 0.0419 |
| 1uaq | opls | 1.6 | 0.1071 | 2.3 | 0 | 98.99 | 0.79 | 5 | 1.49 | 0.89 | 0.1569 | 0.1931 | 0.0362 |
| 1uaq | noschrodinger | 1.6 | 0.0126 | 1.16 | 0 | 98.99 | 0 | 0 | 1.71 | 0.92 | 0.1574 | 0.1924 | 0.035 |
| 1uei | opls | 2.6 | 0.0921 | 2.28 | 0 | 99.45 | 4.02 | 0 | 0.3 | 1.07 | 0.204 | 0.2232 | 0.0192 |
| 1uei | noschrodinger | 2.6 | 0.0027 | 0.74 | 0 | 98.77 | 6.27 | 0 | 1.68 | 1.52 | 0.1916 | 0.2173 | 0.0257 |
| 1uj2 | opls | 1.8 | 0.1013 | 2.29 | 0 | 98.99 | 1.6 | 1 | 1.86 | 1.1 | 0.1656 | 0.1919 | 0.0263 |
| 1uj2 | noschrodinger | 1.8 | 0.0038 | 0.87 | 0 | 99.76 | 1.33 | 0 | 3.02 | 1.19 | 0.1693 | 0.1903 | 0.021 |
| 1uqx | opls | 1.7 | 0.1065 | 2.29 | 0 | 96.36 | 1.04 | 0 | 8.01 | 1.69 | 0.1176 | 0.1559 | 0.0383 |
| 1uqx | noschrodinger | 1.7 | 0.0051 | 1.04 | 0 | 96.4 | 0 | 0 | 1.14 | 1.06 | 0.1188 | 0.1496 | 0.0308 |
| 1uz0 | opls | 1.9 | 0.0999 | 2.11 | 0 | 100 | 1.02 | 0 | 0.5 | 0.68 | 0.1556 | 0.1864 | 0.0308 |
| 1uz0 | noschrodinger | 1.9 | 0.0055 | 1.36 | 0 | 100 | 1.01 | 0 | 5.52 | 1.3 | 0.1551 | 0.1859 | 0.0308 |
| 1v0c | opls | 2.2 | 0.0989 | 2.31 | 0 | 99.42 | 0.67 | 1 | 1.36 | 0.87 | 0.1835 | 0.2135 | 0.03 |

| | | | | | | | | | | | | | |
|---|---|---|---|---|---|---|---|---|---|---|---|---|---|
| 1v0c | noschrodinger | 2.2 | 0.0032 | 0.91 | 0 | 99.42 | 0 | 0 | 1.7 | 0.92 | 0.1674 | 0.2022 | 0.0348 |
| 1v2b | opls | 1.6 | 0.1046 | 2.75 | 0 | 97.67 | 0.4 | 1 | 1.07 | 0.88 | 0.1568 | 0.1863 | 0.0295 |
| 1v2b | noschrodinger | 1.6 | 0.003 | 1.12 | 0 | 97.91 | 0 | 0 | 1.07 | 0.83 | 0.1596 | 0.1832 | 0.0236 |
| 1v2g | opls | 2 | 0.0969 | 2.42 | 0 | 96.49 | 1.36 | 2 | 1.05 | 1.14 | 0.2107 | 0.2598 | 0.0491 |
| 1v2g | noschrodinger | 2 | 0.0041 | 0.84 | 0 | 96.59 | 1.34 | 0 | 1.4 | 1.19 | 0.2134 | 0.2539 | 0.0405 |
| 1v84 | opls | 1.82 | 0.0988 | 2.37 | 1.05 | 96.64 | 3.04 | 4 | 1.6 | 1.49 | 0.1657 | 0.1931 | 0.0274 |
| 1v84 | noschrodinger | 1.82 | 0.0045 | 1.01 | 1.04 | 96.67 | 1.63 | 0 | 2.95 | 1.46 | 0.17 | 0.1949 | 0.0249 |
| 1v8t | opls | 2.6 | 0.0917 | 2.21 | 0 | 95.89 | 2.44 | 0 | 0 | 1.08 | 0.2052 | 0.236 | 0.0308 |
| 1v8t | noschrodinger | 2.6 | 0.0035 | 0.88 | 0 | 94.59 | 3.23 | 0 | 1.63 | 1.67 | 0.1921 | 0.2402 | 0.0481 |
| 1v9h | opls | 2 | 0.0975 | 2.2 | 0 | 95.81 | 0 | 0 | 0.66 | 1.01 | 0.1839 | 0.211 | 0.0271 |
| 1v9h | noschrodinger | 2 | 0.0027 | 0.77 | 0 | 94.76 | 0.61 | 0 | 1.32 | 1.22 | 0.1851 | 0.216 | 0.0309 |
| 1vc8 | opls | 2 | 0.1005 | 2.36 | 0 | 98.78 | 2.5 | 1 | 0.49 | 0.97 | 0.1687 | 0.2092 | 0.0405 |
| 1vc8 | noschrodinger | 2 | 0.0027 | 0.81 | 0 | 98.39 | 2.48 | 0 | 1.71 | 1.22 | 0.1717 | 0.2079 | 0.0362 |
| 1vht | opls | 1.59 | 0.0999 | 2.31 | 0 | 99.61 | 1.53 | 2 | 1.17 | 0.97 | 0.1802 | 0.2115 | 0.0313 |
| 1vht | noschrodinger | 1.59 | 0.0032 | 0.85 | 0 | 99.16 | 1.3 | 0 | 2.15 | 1.07 | 0.1841 | 0.2101 | 0.026 |
| 1vi0 | opls | 1.63 | 0.0965 | 2.85 | 0 | 97.52 | 5.06 | 6 | 4.68 | 1.87 | 0.1961 | 0.2374 | 0.0413 |
| 1vi0 | noschrodinger | 1.63 | 0.0054 | 0.94 | 0 | 98.39 | 5.45 | 0 | 5.33 | 1.85 | 0.2013 | 0.2309 | 0.0296 |
| 1vio | opls | 1.59 | 0.1024 | 2.4 | 0 | 97.69 | 1.49 | 1 | 2.72 | 1.26 | 0.1609 | 0.1874 | 0.0265 |
| 1vio | noschrodinger | 1.59 | 0.0042 | 0.88 | 0 | 97.35 | 0.74 | 0 | 2.18 | 1.12 | 0.1645 | 0.1874 | 0.0229 |
| 1vkj | opls | 2.5 | 0.0905 | 2.39 | 0 | 97.48 | 0.66 | 12 | 1.74 | 1.03 | 0.23 | 0.2576 | 0.0276 |
| 1vkj | noschrodinger | 2.5 | 0.0029 | 0.82 | 0.13 | 95.89 | 0.66 | 0 | 1.45 | 1.17 | 0.2437 | 0.2549 | 0.0112 |
| 1vp6 | opls | 1.7 | 0.1019 | 2.41 | 0 | 100 | 1.43 | 0 | 1.94 | 1.08 | 0.1628 | 0.1998 | 0.037 |
| 1vp6 | noschrodinger | 1.7 | 0.0072 | 0.99 | 0 | 100 | 1.42 | 0 | 8 | 1.55 | 0.1654 | 0.1974 | 0.032 |
| 1vpa | opls | 2.67 | 0.091 | 2.26 | 0 | 97.41 | 0.29 | 0 | 0.71 | 0.85 | 0.2089 | 0.2396 | 0.0307 |
| 1vpa | noschrodinger | 2.67 | 0.0028 | 0.85 | 0.23 | 96.34 | 2.03 | 0 | 2.16 | 1.47 | 0.1949 | 0.2315 | 0.0366 |
| 1vpm | opls | 1.66 | 0.0952 | 2.33 | 0 | 98.89 | 1.87 | 4 | 1.65 | 1.12 | 0.1796 | 0.2042 | 0.0246 |
| 1vpm | noschrodinger | 1.66 | 0.0039 | 0.91 | 0 | 98.02 | 2.33 | 0 | 3.19 | 1.39 | 0.1792 | 0.2052 | 0.026 |
| 1vq2 | opls | 1.92 | 0.0928 | 2.61 | 0.62 | 97.5 | 3.62 | 2 | 3.41 | 1.66 | 0.1942 | 0.2193 | 0.0251 |
| 1vq2 | noschrodinger | 1.92 | 0.0059 | 1.07 | 0 | 97.63 | 2.16 | 0 | 3.81 | 1.5 | 0.2046 | 0.2195 | 0.0149 |
| 1vr0 | opls | 2.49 | 0.0908 | 2.26 | 0 | 96.24 | 1.55 | 0 | 0.89 | 1.17 | 0.1941 | 0.2431 | 0.049 |
| 1vr0 | noschrodinger | 2.49 | 0.0028 | 0.75 | 0 | 95.87 | 2.66 | 0 | 1.26 | 1.46 | 0.1887 | 0.2371 | 0.0484 |
| 1w0h | opls | 1.59 | 0.1074 | 2.36 | 0 | 97.8 | 0 | 1 | 2.09 | 1.03 | 0.1574 | 0.1827 | 0.0253 |
| 1w0h | noschrodinger | 1.59 | 0.0062 | 0.93 | 0 | 97.47 | 1.06 | 0 | 2.4 | 1.15 | 0.1592 | 0.1784 | 0.0192 |
| 1w0y | opls | 2.5 | 0.0969 | 2.22 | 0 | 97.6 | 1.4 | 0 | 1.1 | 1.01 | 0.2115 | 0.2371 | 0.0256 |
| 1w0y | noschrodinger | 2.5 | 0.0028 | 0.89 | 0 | 96.16 | 1.58 | 0 | 1.9 | 1.37 | 0.185 | 0.2259 | 0.0409 |
| 1w2y | opls | 1.65 | 0.1005 | 2.39 | 0 | 99.52 | 1.01 | 5 | 2.09 | 0.98 | 0.1925 | 0.2216 | 0.0291 |
| 1w2y | noschrodinger | 1.65 | 0.0081 | 0.95 | 0.47 | 98.58 | 1.75 | 0 | 2 | 1.15 | 0.1883 | 0.2181 | 0.0298 |
| 1w8g | opls | 2 | 0.1037 | 2.26 | 0 | 98.21 | 1.05 | 0 | 3.8 | 1.18 | 0.1709 | 0.2036 | 0.0327 |
| 1w8g | noschrodinger | 2 | 0.003 | 0.79 | 0 | 98.21 | 1.57 | 0 | 3.8 | 1.32 | 0.156 | 0.1894 | 0.0334 |
| 1wlj | opls | 1.9 | 0.0958 | 2.69 | 0 | 99.34 | 0.71 | 0 | 2.21 | 1 | 0.1748 | 0.2048 | 0.03 |
| 1wlj | noschrodinger | 1.9 | 0.0048 | 0.89 | 0 | 99.39 | 0 | 0 | 1.86 | 0.95 | 0.1819 | 0.2038 | 0.0219 |
| 1wn6 | opls | 1.8 | 0.0977 | 2.38 | 0 | 98.78 | 1.55 | 0 | 2.37 | 1.16 | 0.183 | 0.1983 | 0.0153 |
| 1wn6 | noschrodinger | 1.8 | 0.0074 | 1.23 | 0 | 98.79 | 1.53 | 0 | 5.83 | 1.46 | 0.1806 | 0.2026 | 0.022 |
| 1wol | opls | 1.62 | 0.0981 | 2.56 | 1.68 | 93.28 | 5.83 | 4 | 3.51 | 2.16 | 0.1695 | 0.2006 | 0.0311 |
| 1wol | noschrodinger | 1.62 | 0.0141 | 1.51 | 1.67 | 92.5 | 5.77 | 0 | 8.53 | 2.51 | 0.1714 | 0.1997 | 0.0283 |
| 1wom | opls | 2.5 | 0.0929 | 2.31 | 0.19 | 97.49 | 4.87 | 0 | 1.42 | 1.5 | 0.2147 | 0.2361 | 0.0214 |
| 1wom | noschrodinger | 2.5 | 0.0027 | 0.79 | 0.56 | 96.65 | 6.64 | 0 | 2.96 | 1.92 | 0.2001 | 0.2243 | 0.0242 |
| 1wpx | opls | 2.7 | 0.0904 | 2.3 | 0.16 | 95.11 | 5.19 | 7 | 1.34 | 1.74 | 0.1584 | 0.189 | 0.0306 |
| 1wpx | noschrodinger | 2.7 | 0.0082 | 1.13 | 0.49 | 93.19 | 5.54 | 0 | 3.19 | 2.11 | 0.1539 | 0.1872 | 0.0333 |

| | | | | | | | | | | | | | |
|---|---|---|---|---|---|---|---|---|---|---|---|---|---|
| 1wv4 | opls | 2.2 | 0.0934 | 2.39 | 0.98 | 94.44 | 5.13 | 2 | 1.5 | 1.81 | 0.2712 | 0.2978 | 0.0266 |
| 1wv4 | noschrodinger | 2.2 | 0.0027 | 0.79 | 2.27 | 90.26 | 8.36 | 0 | 6.01 | 2.57 | 0.2676 | 0.2934 | 0.0258 |
| 1wvc | opls | 2.5 | 0.0942 | 2.32 | 0 | 95.45 | 1.46 | 0 | 1.25 | 1.29 | 0.2265 | 0.257 | 0.0305 |
| 1wvc | noschrodinger | 2.5 | 0.0031 | 0.86 | 0.4 | 91.13 | 9.66 | 0 | 5.77 | 2.58 | 0.2142 | 0.2602 | 0.046 |
| 1wwz | opls | 1.75 | 0.1026 | 2.39 | 0 | 97.4 | 1.15 | 0 | 0.75 | 0.9 | 0.164 | 0.2002 | 0.0362 |
| 1wwz | noschrodinger | 1.75 | 0.0035 | 0.83 | 0 | 96.77 | 0.38 | 0 | 1.31 | 1.06 | 0.1707 | 0.1986 | 0.0279 |
| 1wzu | opls | 2 | 0.0972 | 2.34 | 0 | 98.45 | 1.3 | 0 | 2.48 | 1.12 | 0.1828 | 0.2259 | 0.0431 |
| 1wzu | noschrodinger | 2 | 0.0112 | 1.14 | 0 | 98.48 | 1.28 | 0 | 4.3 | 1.29 | 0.183 | 0.2304 | 0.0474 |
| 1x6u | opls | 2.71 | 0.0912 | 2.42 | 0.75 | 95.09 | 6.28 | 1 | 0.94 | 1.73 | 0.2844 | 0.3149 | 0.0305 |
| 1x6u | noschrodinger | 2.71 | 0.0029 | 0.79 | 1.12 | 91.42 | 6.19 | 0 | 4.02 | 2.3 | 0.2335 | 0.2807 | 0.0472 |
| 1x77 | opls | 2.7 | 0.096 | 2.32 | 0.88 | 96.2 | 1.84 | 3 | 0.92 | 1.24 | 0.2238 | 0.2556 | 0.0318 |
| 1x77 | noschrodinger | 2.7 | 0.0029 | 0.81 | 1.16 | 95.64 | 2.55 | 0 | 3.69 | 1.77 | 0.205 | 0.2547 | 0.0497 |
| 1x82 | opls | 1.5 | 0.1026 | 2.24 | 0 | 98.86 | 3.27 | 0 | 1.97 | 1.35 | 0.1531 | 0.1833 | 0.0302 |
| 1x82 | noschrodinger | 1.5 | 0.0094 | 1.17 | 0.53 | 98.94 | 3.14 | 0 | 1.64 | 1.29 | 0.1523 | 0.181 | 0.0287 |
| 1x92 | opls | 2.28 | 0.0968 | 2.29 | 0 | 95.57 | 1.85 | 0 | 0.5 | 1.18 | 0.1705 | 0.1942 | 0.0237 |
| 1x92 | noschrodinger | 2.28 | 0.0029 | 0.81 | 0 | 95.61 | 2.76 | 0 | 1.99 | 1.61 | 0.1629 | 0.1916 | 0.0287 |
| 1xa9 | opls | 2.5 | 0.0963 | 2.32 | 0.46 | 97.26 | 6.19 | 0 | 1.41 | 1.61 | 0.2351 | 0.2653 | 0.0302 |
| 1xa9 | noschrodinger | 2.5 | 0.003 | 0.86 | 0.45 | 96.36 | 9.28 | 0 | 5.73 | 2.29 | 0.2172 | 0.2586 | 0.0414 |
| 1xdd | opls | 2.2 | 0.0974 | 2.15 | 0 | 96.37 | 0.62 | 0 | 1.14 | 1.07 | 0.1552 | 0.1895 | 0.0343 |
| 1xdd | noschrodinger | 2.2 | 0.0107 | 1.26 | 0 | 95.28 | 0.62 | 0 | 2.28 | 1.34 | 0.147 | 0.1842 | 0.0372 |
| 1xe7 | opls | 1.7 | 0.095 | 2.21 | 0 | 99.63 | 1.84 | 2 | 1.32 | 1.06 | 0.1894 | 0.2179 | 0.0285 |
| 1xe7 | noschrodinger | 1.7 | 0.0034 | 0.84 | 0.18 | 99.27 | 2.02 | 0 | 1.98 | 1.2 | 0.1956 | 0.2181 | 0.0225 |
| 1xio | opls | 2 | 0.0881 | 2.29 | 0.48 | 98.1 | 0 | 3 | 2.56 | 1.04 | 0.2029 | 0.2348 | 0.0319 |
| 1xio | noschrodinger | 2 | 0.0035 | 1.2 | 0.47 | 98.59 | 0.52 | 0 | 4.44 | 1.22 | 0.208 | 0.235 | 0.027 |
| 1xk5 | opls | 2.4 | 0.0942 | 2.5 | 1 | 96.52 | 8.84 | 2 | 1.82 | 1.89 | 0.237 | 0.2819 | 0.0449 |
| 1xk5 | noschrodinger | 2.4 | 0.0028 | 0.84 | 1.49 | 95.05 | 9.89 | 0 | 9.41 | 2.6 | 0.2158 | 0.2653 | 0.0495 |
| 1xqa | opls | 1.8 | 0.1028 | 2.28 | 0 | 98.45 | 1.61 | 0 | 4.07 | 1.35 | 0.1619 | 0.2098 | 0.0479 |
| 1xqa | noschrodinger | 1.8 | 0.0048 | 0.94 | 0 | 99.07 | 2.06 | 0 | 5.96 | 1.57 | 0.1585 | 0.1986 | 0.0401 |
| 1xvp | opls | 2.6 | 0.0914 | 2.45 | 0.21 | 94.22 | 2.98 | 4 | 2.13 | 1.74 | 0.3187 | 0.3253 | 0.0066 |
| 1xvp | noschrodinger | 2.6 | 0.0041 | 0.89 | 1.92 | 85.45 | 4.53 | 0 | 9.26 | 2.64 | 0.2902 | 0.3138 | 0.0236 |
| 1xw4 | opls | 2 | 0.0982 | 2.26 | 0 | 100 | 2.17 | 0 | 0 | 0.76 | 0.2199 | 0.2454 | 0.0255 |
| 1xw4 | noschrodinger | 2 | 0.0031 | 0.88 | 0 | 100 | 1.09 | 0 | 1.75 | 0.96 | 0.2099 | 0.2418 | 0.0319 |
| 1xwv | opls | 1.83 | 0.099 | 2.26 | 0.4 | 97.62 | 0.9 | 0 | 1.67 | 1 | 0.2047 | 0.249 | 0.0443 |
| 1xwv | noschrodinger | 1.83 | 0.0033 | 0.89 | 0.79 | 97.64 | 2.67 | 0 | 5.68 | 1.71 | 0.2046 | 0.2456 | 0.041 |
| 1xz8 | opls | 2.8 | 0.0926 | 2.42 | 0.64 | 95.5 | 3.69 | 1 | 0.76 | 1.48 | 0.2129 | 0.2592 | 0.0463 |
| 1xz8 | noschrodinger | 2.8 | 0.0034 | 0.85 | 0.95 | 92.06 | 3.27 | 0 | 4.17 | 2.08 | 0.1924 | 0.2471 | 0.0547 |
| 1y30 | opls | 2.2 | 0.1 | 2.29 | 0 | 97.52 | 0 | 0 | 0.67 | 0.82 | 0.1976 | 0.2228 | 0.0252 |
| 1y30 | noschrodinger | 2.2 | 0.0029 | 0.82 | 0 | 96.81 | 1.26 | 0 | 2.22 | 1.27 | 0.1793 | 0.2249 | 0.0456 |
| 1y7p | opls | 1.9 | 0.0953 | 2.31 | 0.32 | 98.06 | 1.35 | 1 | 3.07 | 1.2 | 0.1879 | 0.2109 | 0.023 |
| 1y7p | noschrodinger | 1.9 | 0.01 | 1.03 | 0.64 | 96.15 | 2.5 | 0 | 7.15 | 1.96 | 0.1857 | 0.209 | 0.0233 |
| 1y7u | opls | 2.8 | 0.0932 | 2.61 | 0 | 98.01 | 12.19 | 4 | 4.93 | 2.08 | 0.1812 | 0.24 | 0.0588 |
| 1y7u | noschrodinger | 2.8 | 0.0028 | 0.8 | 0 | 96.5 | 11.65 | 0 | 4.45 | 2.26 | 0.1892 | 0.238 | 0.0488 |
| 1y89 | opls | 2 | 0.1031 | 2.27 | 0 | 98.65 | 0.54 | 0 | 1.08 | 0.81 | 0.1771 | 0.1979 | 0.0208 |
| 1y89 | noschrodinger | 2 | 0.0032 | 0.8 | 0 | 98.73 | 1.04 | 0 | 1.63 | 0.93 | 0.1647 | 0.1864 | 0.0217 |
| 1y9l | opls | 1.5 | 0.0994 | 2.46 | 0 | 100 | 1.87 | 2 | 5.25 | 1.49 | 0.21 | 0.2523 | 0.0423 |
| 1y9l | noschrodinger | 1.5 | 0.0053 | 0.99 | 0 | 98.15 | 0.93 | 0 | 4.18 | 1.2 | 0.2156 | 0.2434 | 0.0278 |
| 1yat | opls | 2.5 | 0.092 | 2.4 | 0.92 | 95.41 | 1.09 | 0 | 0 | 0.85 | 0.1709 | 0.1954 | 0.0245 |
| 1yat | noschrodinger | 2.5 | 0.0032 | 0.98 | 0.9 | 94.59 | 4.26 | 0 | 2.21 | 1.85 | 0.1598 | 0.1857 | 0.0259 |
| 1ycd | opls | 1.7 | 0.1098 | 2.18 | 0 | 98.72 | 0.72 | 1 | 1.32 | 0.86 | 0.1464 | 0.1763 | 0.0299 |

| | | | | | | | | | | | | | |
|---|---|---|---|---|---|---|---|---|---|---|---|---|---|
| 1ycd | noschrodinger | 1.7 | 0.0039 | 0.89 | 0 | 98.94 | 1.43 | 0 | 1.71 | 1.04 | 0.1453 | 0.1763 | 0.031 |
| 1ylr | opls | 1.7 | 0.105 | 2.28 | 0 | 99.53 | 0.56 | 6 | 1.61 | 0.91 | 0.167 | 0.2016 | 0.0346 |
| 1ylr | noschrodinger | 1.7 | 0.0032 | 0.79 | 0 | 99.07 | 0.56 | 0 | 1.91 | 0.95 | 0.1764 | 0.2009 | 0.0245 |
| 1ylu | opls | 2 | 0.1035 | 2.25 | 0 | 99.3 | 2.25 | 1 | 2.35 | 1.28 | 0.1569 | 0.1978 | 0.0409 |
| 1ylu | noschrodinger | 2 | 0.0027 | 0.76 | 0 | 99.3 | 2.25 | 0 | 1.18 | 1.1 | 0.1423 | 0.1835 | 0.0412 |
| 1yoa | opls | 1.9 | 0.0935 | 2.29 | 0 | 97.44 | 3.08 | 1 | 0.38 | 1.12 | 0.1863 | 0.2097 | 0.0234 |
| 1yoa | noschrodinger | 1.9 | 0.0089 | 1.1 | 0 | 97.45 | 3.82 | 0 | 1.9 | 1.5 | 0.1831 | 0.2089 | 0.0258 |
| 1yr9 | opls | 2.8 | 0.0912 | 2.39 | 2.06 | 92.18 | 8.14 | 1 | 1.99 | 2.14 | 0.244 | 0.2914 | 0.0474 |
| 1yr9 | noschrodinger | 2.8 | 0.0036 | 0.82 | 4.1 | 80.74 | 13.06 | 0 | 7.7 | 3 | 0.2225 | 0.2653 | 0.0428 |
| 1yrb | opls | 1.75 | 0.0982 | 2.46 | 0.39 | 99.02 | 5.34 | 7 | 3.35 | 1.68 | 0.1805 | 0.2135 | 0.033 |
| 1yrb | noschrodinger | 1.75 | 0.0105 | 1.16 | 0.39 | 98.83 | 4.48 | 0 | 7.84 | 1.92 | 0.1802 | 0.2096 | 0.0294 |
| 1ytz | opls | 3 | 0.0906 | 2.42 | 2.09 | 93.72 | 7.99 | 5 | 2.91 | 2.18 | 0.2546 | 0.2927 | 0.0381 |
| 1ytz | noschrodinger | 3 | 0.0031 | 0.73 | 3.65 | 84.38 | 11.76 | 0 | 5.67 | 2.79 | 0.2431 | 0.2848 | 0.0417 |
| 1yuk | opls | 1.8 | 0.1007 | 2.52 | 0.5 | 97.52 | 4.35 | 3 | 6.33 | 1.93 | 0.1934 | 0.2348 | 0.0414 |
| 1yuk | noschrodinger | 1.8 | 0.0089 | 1.27 | 0.96 | 96.63 | 4.26 | 0 | 9.39 | 2.19 | 0.1989 | 0.2394 | 0.0405 |
| 1yun | opls | 1.88 | 0.0977 | 2.56 | 0.48 | 97.83 | 5.44 | 5 | 3.98 | 1.78 | 0.2045 | 0.2427 | 0.0382 |
| 1yun | noschrodinger | 1.88 | 0.0079 | 1.42 | 0.24 | 97.6 | 9.4 | 0 | 7.37 | 2.23 | 0.211 | 0.2475 | 0.0365 |
| 1ywm | opls | 1.87 | 0.1033 | 2.25 | 0 | 98.87 | 0.63 | 0 | 2.1 | 0.98 | 0.1808 | 0.2188 | 0.038 |
| 1ywm | noschrodinger | 1.87 | 0.0087 | 1.12 | 0 | 98.88 | 0.63 | 0 | 2.44 | 1.03 | 0.1819 | 0.2244 | 0.0425 |
| 1yzx | opls | 1.86 | 0.0956 | 2.34 | 0 | 99.07 | 0.53 | 1 | 0.98 | 0.79 | 0.1448 | 0.1777 | 0.0329 |
| 1yzx | noschrodinger | 1.86 | 0.009 | 0.98 | 0 | 98.15 | 1.06 | 0 | 1.12 | 0.84 | 0.1508 | 0.1794 | 0.0286 |
| 1z4r | opls | 1.68 | 0.099 | 2.32 | 0 | 98.75 | 2.11 | 1 | 2.53 | 1.28 | 0.1684 | 0.1975 | 0.0291 |
| 1z4r | noschrodinger | 1.68 | 0.0071 | 0.98 | 0 | 98.76 | 1.4 | 0 | 1.45 | 0.99 | 0.1631 | 0.1971 | 0.034 |
| 1za4 | opls | 1.9 | 0.0949 | 2.4 | 0 | 96.43 | 2.92 | 0 | 2.15 | 1.58 | 0.2281 | 0.2777 | 0.0496 |
| 1za4 | noschrodinger | 1.9 | 0.0134 | 1.28 | 0.49 | 96.55 | 2.3 | 0 | 4.61 | 1.73 | 0.2195 | 0.2639 | 0.0444 |
| 1zak | opls | 3.2 | 0.0904 | 2.37 | 1.15 | 94.04 | 5.5 | 2 | 0.7 | 1.69 | 0.2032 | 0.257 | 0.0538 |
| 1zak | noschrodinger | 3.2 | 0.0029 | 0.81 | 1.61 | 91.28 | 7.33 | 0 | 4.76 | 2.41 | 0.2066 | 0.2658 | 0.0592 |
| 1zar | opls | 1.75 | 0.103 | 2.21 | 0.39 | 98.83 | 0.43 | 0 | 1.57 | 0.9 | 0.1622 | 0.1839 | 0.0217 |
| 1zar | noschrodinger | 1.75 | 0.0064 | 0.96 | 0.38 | 98.85 | 0.43 | 0 | 3.39 | 1.13 | 0.1598 | 0.1848 | 0.025 |
| 1zct | opls | 2.6 | 0.0916 | 2.26 | 0 | 96.63 | 2.43 | 0 | 0.61 | 1.21 | 0.2444 | 0.2626 | 0.0182 |
| 1zct | noschrodinger | 2.6 | 0.0028 | 0.8 | 0.39 | 95.47 | 3.07 | 0 | 1.84 | 1.63 | 0.2222 | 0.2506 | 0.0284 |
| 1zrh | opls | 2.1 | 0.0982 | 2.23 | 0 | 98.85 | 1.25 | 1 | 0.91 | 0.85 | 0.19 | 0.2184 | 0.0284 |
| 1zrh | noschrodinger | 2.1 | 0.0031 | 0.83 | 0 | 98.47 | 1.66 | 0 | 0.91 | 0.94 | 0.1788 | 0.2076 | 0.0288 |
| 1zsk | opls | 1.9 | 0.1015 | 2.67 | 0 | 97.02 | 2.93 | 4 | 4.18 | 1.73 | 0.144 | 0.1793 | 0.0353 |
| 1zsk | noschrodinger | 1.9 | 0.0096 | 1.12 | 0 | 97.46 | 2.91 | 0 | 3.39 | 1.59 | 0.1479 | 0.1864 | 0.0385 |
| 1ztf | opls | 2 | 0.0997 | 2.34 | 0 | 98.1 | 2.44 | 0 | 1.45 | 1.18 | 0.1849 | 0.231 | 0.0461 |
| 1ztf | noschrodinger | 2 | 0.0098 | 0.93 | 0.8 | 96.02 | 6.98 | 0 | 3.4 | 2.05 | 0.1724 | 0.2215 | 0.0491 |
| 1zx5 | opls | 2.3 | 0.0985 | 2.19 | 0 | 97.9 | 0.79 | 1 | 1.22 | 0.86 | 0.1594 | 0.1823 | 0.0229 |
| 1zx5 | noschrodinger | 2.3 | 0.0028 | 0.78 | 0 | 96.98 | 1.55 | 0 | 1.02 | 1.12 | 0.1634 | 0.179 | 0.0156 |
| 1zzm | opls | 1.79 | 0.1056 | 2.33 | 0.39 | 98.44 | 1.4 | 2 | 2.42 | 1.13 | 0.1499 | 0.1985 | 0.0486 |
| 1zzm | noschrodinger | 1.79 | 0.0046 | 0.96 | 0.78 | 97.28 | 1.86 | 0 | 2.9 | 1.42 | 0.1574 | 0.198 | 0.0406 |
| 1zzq | opls | 1.9 | 0.1208 | 2.73 | 0.12 | 96.89 | 1.77 | 6 | 5.05 | 1.64 | 0.1848 | 0.2224 | 0.0376 |
| 1zzq | noschrodinger | 1.9 | 0.004 | 0.95 | 0.12 | 97.16 | 1.21 | 1 | 3.95 | 1.4 | 0.191 | 0.2179 | 0.0269 |
| 227l | opls | 1.86 | 0.0993 | 2.25 | 0 | 98.1 | 2.27 | 2 | 0.76 | 1.01 | 0.1464 | 0.1889 | 0.0425 |
| 227l | noschrodinger | 1.86 | 0.0078 | 0.91 | 0 | 98.75 | 4.48 | 0 | 2.28 | 1.5 | 0.1499 | 0.1917 | 0.0418 |
| 2a15 | opls | 1.68 | 0.1056 | 2.71 | 0 | 98.46 | 0.83 | 5 | 4.27 | 1.21 | 0.1615 | 0.1911 | 0.0296 |
| 2a15 | noschrodinger | 1.68 | 0.0076 | 1.07 | 0 | 98.47 | 0.82 | 0 | 4.25 | 1.21 | 0.1676 | 0.1894 | 0.0218 |
| 2a3h | opls | 2.01 | 0.1041 | 2.2 | 0 | 96.63 | 0 | 0 | 1.29 | 1.07 | 0.1263 | 0.1647 | 0.0384 |
| 2a3h | noschrodinger | 2.01 | 0.0036 | 0.88 | 0 | 96.64 | 0 | 0 | 0.86 | 0.98 | 0.1139 | 0.1586 | 0.0447 |

| | | | | | | | | | | | | | |
|---|---|---|---|---|---|---|---|---|---|---|---|---|---|
| 2a41 | opls | 2.6 | 0.0939 | 2.29 | 0 | 97.7 | 2.99 | 3 | 0.67 | 1.14 | 0.1842 | 0.2183 | 0.0341 |
| 2a41 | noschrodinger | 2.6 | 0.0032 | 0.83 | 0.3 | 97.56 | 6.29 | 0 | 2.77 | 1.76 | 0.1738 | 0.2178 | 0.044 |
| 2a42 | opls | 1.85 | 0.0988 | 2.33 | 0 | 98.01 | 1.62 | 3 | 1.69 | 1.08 | 0.1794 | 0.2188 | 0.0394 |
| 2a42 | noschrodinger | 1.85 | 0.0029 | 0.82 | 0.33 | 98.03 | 1.8 | 0 | 1.98 | 1.16 | 0.192 | 0.2109 | 0.0189 |
| 2a84 | opls | 1.55 | 0.103 | 2.59 | 0 | 99.26 | 0.48 | 2 | 2.8 | 1.07 | 0.1438 | 0.176 | 0.0322 |
| 2a84 | noschrodinger | 1.55 | 0.0065 | 1.08 | 0 | 99.27 | 0.47 | 0 | 0.94 | 0.78 | 0.1505 | 0.1759 | 0.0254 |
| 2a8e | opls | 2 | 0.0951 | 2.23 | 0 | 97.4 | 3.85 | 0 | 1.73 | 1.49 | 0.2121 | 0.2278 | 0.0157 |
| 2a8e | noschrodinger | 2 | 0.0026 | 0.76 | 0 | 96.63 | 2.66 | 0 | 1.16 | 1.37 | 0.1998 | 0.2221 | 0.0223 |
| 2aco | opls | 1.8 | 0.1048 | 2.23 | 0 | 99.39 | 0.36 | 2 | 3.74 | 1.16 | 0.1614 | 0.1928 | 0.0314 |
| 2aco | noschrodinger | 1.8 | 0.0042 | 0.92 | 0 | 99.39 | 0.72 | 0 | 3.19 | 1.11 | 0.1648 | 0.1917 | 0.0269 |
| 2ag8 | opls | 2.09 | 0.1 | 2.35 | 0 | 99.15 | 4.69 | 0 | 2.22 | 1.51 | 0.1994 | 0.2454 | 0.046 |
| 2ag8 | noschrodinger | 2.09 | 0.003 | 0.79 | 0.38 | 98.08 | 5.97 | 0 | 3.21 | 1.7 | 0.1832 | 0.2304 | 0.0472 |
| 2aho | opls | 3 | 0.0916 | 2.37 | 2.03 | 92.51 | 7.72 | 7 | 2.44 | 2.17 | 0.2371 | 0.2826 | 0.0455 |
| 2aho | noschrodinger | 3 | 0.0034 | 0.9 | 3.23 | 83.69 | 10.96 | 0 | 11.11 | 3.03 | 0.2291 | 0.2629 | 0.0338 |
| 2ans | opls | 2.56 | 0.0941 | 2.22 | 0 | 98.05 | 4.91 | 2 | 0.96 | 1.31 | 0.1962 | 0.22 | 0.0238 |
| 2ans | noschrodinger | 2.56 | 0.0025 | 0.73 | 0 | 95.35 | 4.87 | 0 | 0.96 | 1.63 | 0.1826 | 0.2173 | 0.0347 |
| 2aqx | opls | 2.5 | 0.0926 | 2.33 | 0 | 96.49 | 5.15 | 3 | 1.59 | 1.68 | 0.2059 | 0.2284 | 0.0225 |
| 2aqx | noschrodinger | 2.5 | 0.0036 | 0.86 | 0.52 | 95.45 | 8.09 | 1 | 2.76 | 2.07 | 0.1904 | 0.2248 | 0.0344 |
| 2arp | opls | 2 | 0.0973 | 2.53 | 0 | 97.52 | 3.12 | 1 | 1.52 | 1.37 | 0.2138 | 0.2358 | 0.022 |
| 2arp | noschrodinger | 2 | 0.003 | 0.8 | 0.41 | 93.06 | 5.33 | 0 | 5.33 | 2.28 | 0.2104 | 0.2392 | 0.0288 |
| 2aru | opls | 2.51 | 0.0937 | 2.32 | 0.42 | 96.67 | 2.39 | 0 | 0.5 | 1.17 | 0.1962 | 0.2304 | 0.0342 |
| 2aru | noschrodinger | 2.51 | 0.0025 | 0.79 | 0.41 | 95.47 | 2.84 | 0 | 2.78 | 1.73 | 0.1951 | 0.2267 | 0.0316 |
| 2atq | opls | 3.2 | 0.0904 | 2.34 | 2.2 | 89.39 | 5.86 | 11 | 1.23 | 1.99 | 0.2684 | 0.2868 | 0.0184 |
| 2atq | noschrodinger | 3.2 | 0.0034 | 0.87 | 5.62 | 76.93 | 8.08 | 0 | 13.56 | 3.1 | 0.2614 | 0.276 | 0.0146 |
| 2atz | opls | 2 | 0.0961 | 2.28 | 0.62 | 98.15 | 3.33 | 0 | 1.01 | 1.19 | 0.239 | 0.2712 | 0.0322 |
| 2atz | noschrodinger | 2 | 0.0027 | 0.77 | 1.15 | 98.28 | 4.52 | 0 | 3.7 | 1.66 | 0.2286 | 0.262 | 0.0334 |
| 2b0c | opls | 2 | 0.0961 | 2.27 | 0 | 98.19 | 1.88 | 1 | 2.19 | 1.2 | 0.1892 | 0.2186 | 0.0294 |
| 2b0c | noschrodinger | 2 | 0.0058 | 0.88 | 0 | 98.48 | 2.34 | 0 | 1.88 | 1.23 | 0.1843 | 0.2139 | 0.0296 |
| 2b1r | opls | 2.2 | 0.1007 | 2.35 | 0 | 94.83 | 1.47 | 3 | 1.04 | 1.29 | 0.1475 | 0.1931 | 0.0456 |
| 2b1r | noschrodinger | 2.2 | 0.0046 | 0.91 | 0 | 95.04 | 1.46 | 0 | 1.56 | 1.37 | 0.155 | 0.1984 | 0.0434 |
| 2b3d | opls | 2.1 | 0.1013 | 2.26 | 2.13 | 94.68 | 1.24 | 3 | 3.54 | 1.58 | 0.1931 | 0.2328 | 0.0397 |
| 2b3d | noschrodinger | 2.1 | 0.0027 | 0.79 | 2.38 | 93.65 | 2.16 | 0 | 3.06 | 1.77 | 0.1837 | 0.2297 | 0.046 |
| 2bbh | opls | 1.85 | 0.0979 | 2.99 | 0 | 99.02 | 2.36 | 0 | 3.87 | 1.46 | 0.1862 | 0.2179 | 0.0317 |
| 2bbh | noschrodinger | 1.85 | 0.0078 | 1.71 | 0 | 98.16 | 4.59 | 0 | 6.29 | 1.85 | 0.1882 | 0.2232 | 0.035 |
| 2bfr | opls | 2.49 | 0.0927 | 2.18 | 0 | 98.41 | 0 | 0 | 0.66 | 0.72 | 0.1818 | 0.2181 | 0.0363 |
| 2bfr | noschrodinger | 2.49 | 0.0028 | 0.81 | 0 | 97.88 | 1.33 | 0 | 0.67 | 0.84 | 0.1733 | 0.2191 | 0.0458 |
| 2bkj | opls | 2.08 | 0.0899 | 2.31 | 0 | 99.14 | 1.16 | 0 | 0.29 | 0.66 | 0.1677 | 0.1958 | 0.0281 |
| 2bkj | noschrodinger | 2.08 | 0.0032 | 0.79 | 0 | 98.01 | 1.44 | 0 | 0.87 | 0.89 | 0.1591 | 0.1927 | 0.0336 |
| 2bkv | opls | 1.5 | 0.0998 | 3.13 | 0 | 97.68 | 4.07 | 0 | 6.04 | 1.86 | 0.1707 | 0.2029 | 0.0322 |
| 2bkv | noschrodinger | 1.5 | 0.0141 | 1.42 | 0 | 97.7 | 4.03 | 0 | 4.23 | 1.73 | 0.1713 | 0.1987 | 0.0274 |
| 2bl9 | opls | 1.73 | 0.103 | 2.27 | 0 | 98.58 | 3 | 0 | 1.64 | 1.28 | 0.1841 | 0.2158 | 0.0317 |
| 2bl9 | noschrodinger | 1.73 | 0.008 | 0.95 | 0 | 98.58 | 3 | 0 | 3.28 | 1.48 | 0.1888 | 0.2174 | 0.0286 |
| 2blc | opls | 2.25 | 0.1002 | 2.27 | 0 | 98.04 | 2.02 | 1 | 0.27 | 0.83 | 0.2075 | 0.2416 | 0.0341 |
| 2blc | noschrodinger | 2.25 | 0.0027 | 0.75 | 0.47 | 95.26 | 3.02 | 0 | 3.82 | 1.86 | 0.1999 | 0.243 | 0.0431 |
| 2bml | opls | 2.6 | 0.0904 | 2.32 | 0.41 | 94.72 | 4.33 | 3 | 0.7 | 1.57 | 0.2405 | 0.2594 | 0.0189 |
| 2bml | noschrodinger | 2.6 | 0.0025 | 0.89 | 1.21 | 93.12 | 6.7 | 0 | 4.7 | 2.31 | 0.2072 | 0.251 | 0.0438 |
| 2bne | opls | 2.28 | 0.0928 | 2.32 | 0 | 97.83 | 3.51 | 2 | 0.68 | 1.17 | 0.2018 | 0.2378 | 0.036 |
| 2bne | noschrodinger | 2.28 | 0.0027 | 0.75 | 0.43 | 94.83 | 4.03 | 0 | 4.47 | 2.04 | 0.1792 | 0.2246 | 0.0454 |
| 2bt1 | opls | 2.7 | 0.0935 | 2.3 | 1.59 | 97.21 | 4.93 | 1 | 0.98 | 1.46 | 0.1912 | 0.2285 | 0.0373 |

| | | | | | | | | | | | | | |
|---|---|---|---|---|---|---|---|---|---|---|---|---|---|
| 2bt1 | noschrodinger | 2.7 | 0.0036 | 0.94 | 0.79 | 94.84 | 11.61 | 0 | 3.18 | 2.28 | 0.1918 | 0.2241 | 0.0323 |
| 2bue | opls | 1.7 | 0.1031 | 2.39 | 0 | 98.87 | 0 | 0 | 4.65 | 1.24 | 0.1511 | 0.1804 | 0.0293 |
| 2bue | noschrodinger | 1.7 | 0.0066 | 1.03 | 0 | 98.87 | 1.28 | 0 | 2.99 | 1.17 | 0.1544 | 0.1804 | 0.026 |
| 2bur | opls | 1.79 | 0.0981 | 2.27 | 0 | 97.25 | 0.81 | 0 | 1.86 | 1.09 | 0.1358 | 0.1619 | 0.0261 |
| 2bur | noschrodinger | 1.79 | 0.0123 | 1.2 | 0 | 96.79 | 0.81 | 0 | 1.43 | 1.08 | 0.1355 | 0.1657 | 0.0302 |
| 2byc | opls | 1.9 | 0.0946 | 2.4 | 0 | 97.32 | 4.76 | 1 | 2.24 | 1.65 | 0.2303 | 0.2725 | 0.0422 |
| 2byc | noschrodinger | 1.9 | 0.004 | 0.89 | 0.38 | 96.6 | 5.98 | 0 | 4.49 | 2.03 | 0.2275 | 0.2727 | 0.0452 |
| 2byo | opls | 2.1 | 0.0971 | 2.46 | 0 | 96.61 | 0.67 | 0 | 0.35 | 0.84 | 0.2038 | 0.2393 | 0.0355 |
| 2byo | noschrodinger | 2.1 | 0.0028 | 0.84 | 0 | 97.21 | 2.63 | 0 | 1.04 | 1.27 | 0.196 | 0.238 | 0.042 |
| 2bzg | opls | 1.58 | 0.1019 | 2.2 | 0 | 98.62 | 1.5 | 0 | 1.32 | 0.99 | 0.192 | 0.2167 | 0.0247 |
| 2bzg | noschrodinger | 1.58 | 0.0074 | 1.11 | 0 | 99.12 | 1.96 | 0 | 1.58 | 1.13 | 0.1899 | 0.2187 | 0.0288 |
| 2c3x | opls | 2.7 | 0.0963 | 2.24 | 0 | 97.81 | 5.19 | 0 | 1.69 | 1.51 | 0.2293 | 0.2645 | 0.0352 |
| 2c3x | noschrodinger | 2.7 | 0.0035 | 0.9 | 0 | 96.17 | 10.39 | 0 | 1.7 | 1.96 | 0.2157 | 0.2504 | 0.0347 |
| 2c4i | opls | 1.95 | 0.0956 | 2.25 | 0.43 | 97.83 | 1.43 | 0 | 2.35 | 1.17 | 0.1835 | 0.2252 | 0.0417 |
| 2c4i | noschrodinger | 1.95 | 0.005 | 0.96 | 0.43 | 97.01 | 2.34 | 0 | 2.88 | 1.53 | 0.1852 | 0.2249 | 0.0397 |
| 2c9a | opls | 2.69 | 0.091 | 2.29 | 0.41 | 96.34 | 2.28 | 1 | 0.48 | 1.18 | 0.2455 | 0.2735 | 0.028 |
| 2c9a | noschrodinger | 2.69 | 0.0044 | 0.89 | 0.78 | 92.22 | 6.7 | 0 | 6.43 | 2.46 | 0.2294 | 0.2606 | 0.0312 |
| 2cbg | opls | 2.49 | 0.0946 | 2.24 | 0.49 | 97.06 | 4.71 | 2 | 1.51 | 1.57 | 0.1775 | 0.2091 | 0.0316 |
| 2cbg | noschrodinger | 2.49 | 0.003 | 0.8 | 1.46 | 94.66 | 8.72 | 0 | 2.12 | 2.07 | 0.1576 | 0.2064 | 0.0488 |
| 2cfa | opls | 2.3 | 0.0944 | 2.55 | 0.31 | 99.08 | 9.03 | 3 | 2.49 | 1.76 | 0.1783 | 0.2134 | 0.0351 |
| 2cfa | noschrodinger | 2.3 | 0.0094 | 0.93 | 0.29 | 98.53 | 10.71 | 0 | 3.34 | 1.91 | 0.1816 | 0.2126 | 0.031 |
| 2cis | opls | 1.62 | 0.1038 | 2.26 | 0 | 96.82 | 0.78 | 1 | 1.72 | 1.12 | 0.1612 | 0.1913 | 0.0301 |
| 2cis | noschrodinger | 1.62 | 0.0052 | 1.05 | 0 | 97.17 | 0.78 | 0 | 1.5 | 1.04 | 0.1626 | 0.189 | 0.0264 |
| 2cm9 | opls | 2.3 | 0.0926 | 2.24 | 0 | 96.53 | 1.57 | 2 | 0.44 | 1.03 | 0.1899 | 0.2141 | 0.0242 |
| 2cm9 | noschrodinger | 2.3 | 0.0026 | 0.72 | 0.69 | 97.92 | 3.15 | 0 | 2.18 | 1.39 | 0.1793 | 0.2097 | 0.0304 |
| 2cns | opls | 2.5 | 0.0932 | 2.44 | 0.23 | 96.6 | 2.44 | 3 | 0.8 | 1.26 | 0.1943 | 0.2345 | 0.0402 |
| 2cns | noschrodinger | 2.5 | 0.0027 | 0.77 | 0.67 | 95.97 | 3.47 | 0 | 2.28 | 1.69 | 0.2036 | 0.2306 | 0.027 |
| 2cul | opls | 1.65 | 0.0997 | 2.19 | 0 | 98.17 | 1.13 | 0 | 1.1 | 0.86 | 0.1476 | 0.17 | 0.0224 |
| 2cul | noschrodinger | 1.65 | 0.0039 | 0.9 | 0 | 97.74 | 1.12 | 0 | 0.82 | 0.85 | 0.148 | 0.168 | 0.02 |
| 2cvj | opls | 2 | 0.0984 | 2.29 | 0 | 95.4 | 0.74 | 0 | 1.43 | 1.2 | 0.1733 | 0.2032 | 0.0299 |
| 2cvj | noschrodinger | 2 | 0.0033 | 0.85 | 0 | 95.51 | 0.73 | 0 | 1.43 | 1.19 | 0.1752 | 0.2034 | 0.0282 |
| 2cy2 | opls | 2 | 0.0997 | 2.31 | 0 | 98.81 | 5.43 | 1 | 1.41 | 1.43 | 0.1861 | 0.2185 | 0.0324 |
| 2cy2 | noschrodinger | 2 | 0.0123 | 1.15 | 0 | 99.42 | 5.34 | 0 | 0.71 | 1.28 | 0.1893 | 0.221 | 0.0317 |
| 2cz6 | opls | 1.5 | 0.1088 | 2.35 | 0 | 97.73 | 1.43 | 3 | 4.14 | 1.37 | 0.1247 | 0.1468 | 0.0221 |
| 2cz6 | noschrodinger | 1.5 | 0.0056 | 1.04 | 0 | 98 | 1.42 | 1 | 2.3 | 1.13 | 0.126 | 0.1455 | 0.0195 |
| 2czl | opls | 1.55 | 0.1032 | 2.4 | 0 | 98.49 | 0.47 | 0 | 1.16 | 0.83 | 0.1608 | 0.1839 | 0.0231 |
| 2czl | noschrodinger | 1.55 | 0.0058 | 1.16 | 0 | 98.11 | 0.47 | 0 | 2.78 | 1.07 | 0.1653 | 0.1863 | 0.021 |
| 2d1e | opls | 1.51 | 0.1048 | 2.5 | 0 | 97.45 | 2.16 | 1 | 7.24 | 1.76 | 0.1525 | 0.1842 | 0.0317 |
| 2d1e | noschrodinger | 1.51 | 0.0104 | 1.25 | 0 | 96.68 | 2.59 | 0 | 8.19 | 1.97 | 0.1509 | 0.1775 | 0.0266 |
| 2d3y | opls | 1.55 | 0.1014 | 2.33 | 0 | 97.69 | 0.58 | 1 | 1.72 | 0.99 | 0.148 | 0.1805 | 0.0325 |
| 2d3y | noschrodinger | 1.55 | 0.0134 | 2.15 | 0 | 97.7 | 0.58 | 0 | 2.58 | 1.11 | 0.2161 | 0.2488 | 0.0327 |
| 2d5a | opls | 1.7 | 0.0982 | 2.44 | 0 | 99.28 | 1.6 | 0 | 4.55 | 1.38 | 0.1834 | 0.2113 | 0.0279 |
| 2d5a | noschrodinger | 1.7 | 0.0118 | 1.3 | 0 | 99.29 | 2.38 | 0 | 5.79 | 1.6 | 0.1773 | 0.2056 | 0.0283 |
| 2dcl | opls | 2.29 | 0.0945 | 2.32 | 0 | 99.28 | 3.14 | 0 | 0.61 | 1.08 | 0.2329 | 0.2652 | 0.0323 |
| 2dcl | noschrodinger | 2.29 | 0.0033 | 0.84 | 0.35 | 96.45 | 7.34 | 0 | 3.44 | 2.03 | 0.2184 | 0.2551 | 0.0367 |
| 2ddo | opls | 2.6 | 0.0944 | 2.31 | 0 | 96.25 | 3.64 | 0 | 1.08 | 1.49 | 0.2017 | 0.2576 | 0.0559 |
| 2ddo | noschrodinger | 2.6 | 0.0027 | 0.79 | 0.58 | 95.59 | 5.18 | 0 | 3.14 | 1.95 | 0.2012 | 0.2594 | 0.0582 |
| 2djh | opls | 1.75 | 0.101 | 2.25 | 0 | 97.85 | 1.2 | 1 | 0.65 | 0.81 | 0.177 | 0.2092 | 0.0322 |
| 2djh | noschrodinger | 1.75 | 0.0078 | 1.34 | 0 | 96.81 | 1.2 | 0 | 5.83 | 1.58 | 0.1771 | 0.2098 | 0.0327 |

| | | | | | | | | | | | | | |
|---|---|---|---|---|---|---|---|---|---|---|---|---|---|
| 2dkg | opls | 2 | 0.1002 | 2.27 | 0 | 96.34 | 1.23 | 0 | 1.17 | 1.14 | 0.189 | 0.2161 | 0.0271 |
| 2dkg | noschrodinger | 2 | 0.0029 | 0.78 | 0 | 95.92 | 1.22 | 0 | 3.51 | 1.49 | 0.1692 | 0.2085 | 0.0393 |
| 2doh | opls | 2 | 0.1057 | 2.57 | 0 | 96 | 3.47 | 2 | 1 | 1.48 | 0.2431 | 0.2805 | 0.0374 |
| 2doh | noschrodinger | 2 | 0.0027 | 0.82 | 0 | 95.65 | 9.04 | 0 | 7.69 | 2.45 | 0.2312 | 0.2708 | 0.0396 |
| 2dqa | opls | 1.6 | 0.1026 | 2.51 | 0 | 98.35 | 1.74 | 4 | 6.02 | 1.51 | 0.142 | 0.18 | 0.038 |
| 2dqa | noschrodinger | 1.6 | 0.0136 | 1.4 | 0 | 98.35 | 0.87 | 0 | 12.9 | 1.62 | 0.1443 | 0.1814 | 0.0371 |
| 2drz | opls | 2.19 | 0.097 | 2.18 | 0 | 96.46 | 1.83 | 0 | 1.2 | 1.27 | 0.2021 | 0.2421 | 0.04 |
| 2drz | noschrodinger | 2.19 | 0.0041 | 0.83 | 0.39 | 96.48 | 1.82 | 0 | 2.65 | 1.48 | 0.194 | 0.2376 | 0.0436 |
| 2dsc | opls | 2 | 0.0986 | 2.29 | 0.26 | 97.4 | 2.42 | 1 | 1.46 | 1.29 | 0.1842 | 0.2147 | 0.0305 |
| 2dsc | noschrodinger | 2 | 0.0035 | 0.81 | 0 | 97.15 | 3.61 | 0 | 4.54 | 1.81 | 0.1791 | 0.2201 | 0.041 |
| 2dtn | opls | 2.5 | 0.0948 | 2.27 | 0 | 98.51 | 2.67 | 0 | 1.01 | 1.12 | 0.2164 | 0.2423 | 0.0259 |
| 2dtn | noschrodinger | 2.5 | 0.0025 | 0.7 | 0 | 97.54 | 2.65 | 0 | 2.74 | 1.48 | 0.1963 | 0.2376 | 0.0413 |
| 2dvn | opls | 1.6 | 0.1022 | 2.38 | 0 | 98.91 | 0.96 | 0 | 1.62 | 0.91 | 0.1452 | 0.1656 | 0.0204 |
| 2dvn | noschrodinger | 1.6 | 0.0099 | 1.14 | 0 | 98.91 | 0.96 | 0 | 2.92 | 1.08 | 0.1443 | 0.1624 | 0.0181 |
| 2dy5 | opls | 2.71 | 0.0902 | 2.33 | 0 | 97.16 | 3.3 | 0 | 0.28 | 1.15 | 0.1989 | 0.2338 | 0.0349 |
| 2dy5 | noschrodinger | 2.71 | 0.0051 | 0.95 | 0.47 | 97.17 | 6.56 | 0 | 1.12 | 1.59 | 0.1685 | 0.213 | 0.0445 |
| 2dy9 | opls | 2.01 | 0.0987 | 2.27 | 0.66 | 94.43 | 1.14 | 1 | 0.39 | 1.06 | 0.2528 | 0.2771 | 0.0243 |
| 2dy9 | noschrodinger | 2.01 | 0.0028 | 0.78 | 1.31 | 95.1 | 1.13 | 0 | 2.17 | 1.37 | 0.2334 | 0.2788 | 0.0454 |
| 2e1m | opls | 2.59 | 0.09 | 2.32 | 0.34 | 94.69 | 1.64 | 4 | 0.85 | 1.29 | 0.2413 | 0.2648 | 0.0235 |
| 2e1m | noschrodinger | 2.59 | 0.0037 | 0.88 | 2.55 | 90.49 | 4.29 | 0 | 5.85 | 2.34 | 0.2235 | 0.2611 | 0.0376 |
| 2e2o | opls | 1.65 | 0.1045 | 2.4 | 0 | 98.99 | 4.68 | 4 | 2.18 | 1.5 | 0.1613 | 0.2029 | 0.0416 |
| 2e2o | noschrodinger | 1.65 | 0.0041 | 0.91 | 0 | 99.66 | 2.54 | 0 | 2.61 | 1.36 | 0.1646 | 0.1963 | 0.0317 |
| 2e2p | opls | 2 | 0.0996 | 2.28 | 0.34 | 98.47 | 2.8 | 3 | 1.3 | 1.19 | 0.1555 | 0.1988 | 0.0433 |
| 2e2p | noschrodinger | 2 | 0.0029 | 0.8 | 0.34 | 98.82 | 2.77 | 0 | 2.17 | 1.33 | 0.1635 | 0.1958 | 0.0323 |
| 2e3q | opls | 2.08 | 0.0972 | 2.3 | 0.43 | 98.71 | 1.9 | 0 | 1.02 | 1.01 | 0.1759 | 0.2158 | 0.0399 |
| 2e3q | noschrodinger | 2.08 | 0.0026 | 0.79 | 0.86 | 99.14 | 2.36 | 0 | 1.02 | 1.08 | 0.1853 | 0.216 | 0.0307 |
| 2e4p | opls | 2 | 0.0968 | 2.24 | 0 | 97.57 | 1.06 | 1 | 0.59 | 0.8 | 0.2142 | 0.2446 | 0.0304 |
| 2e4p | noschrodinger | 2 | 0.0052 | 1.04 | 0 | 98.11 | 4.23 | 0 | 2.05 | 1.45 | 0.1866 | 0.2392 | 0.0526 |
| 2e5x | opls | 2 | 0.1015 | 2.24 | 0 | 99.45 | 0 | 0 | 1.31 | 0.86 | 0.1895 | 0.2182 | 0.0287 |
| 2e5x | noschrodinger | 2 | 0.0029 | 0.77 | 0 | 98.91 | 0.64 | 0 | 2.3 | 1.01 | 0.1762 | 0.2131 | 0.0369 |
| 2e5y | opls | 1.79 | 0.0992 | 2.41 | 0.38 | 96.15 | 1.87 | 2 | 1.17 | 1.3 | 0.186 | 0.235 | 0.049 |
| 2e5y | noschrodinger | 1.79 | 0.0046 | 0.98 | 0.76 | 95.8 | 3.24 | 0 | 4.45 | 1.9 | 0.187 | 0.2293 | 0.0423 |
| 2.00E+83 | opls | 1.52 | 0.1071 | 2.45 | 0 | 98.74 | 0 | 1 | 6 | 1.33 | 0.1293 | 0.1596 | 0.0303 |
| 2.00E+83 | noschrodinger | 1.52 | 0.0075 | 1.1 | 0 | 98.33 | 0 | 0 | 4.43 | 1.22 | 0.1339 | 0.1576 | 0.0237 |
| 2ea1 | opls | 1.8 | 0.0974 | 2.56 | 1.65 | 94.63 | 5.08 | 3 | 4.18 | 2.11 | 0.1855 | 0.2263 | 0.0408 |
| 2ea1 | noschrodinger | 1.8 | 0.0124 | 1.23 | 1.65 | 95.47 | 2.02 | 0 | 6.53 | 1.91 | 0.1872 | 0.2235 | 0.0363 |
| 2ei5 | opls | 1.88 | 0.0977 | 2.22 | 0 | 99.01 | 0 | 2 | 0.3 | 0.61 | 0.1759 | 0.206 | 0.0301 |
| 2ei5 | noschrodinger | 1.88 | 0.0083 | 0.95 | 0 | 99.04 | 0 | 0 | 2.08 | 0.98 | 0.1774 | 0.2018 | 0.0244 |
| 2eis | opls | 2.1 | 0.0997 | 2.35 | 0.47 | 98.11 | 5.43 | 1 | 2.29 | 1.57 | 0.1993 | 0.2201 | 0.0208 |
| 2eis | noschrodinger | 2.1 | 0.0031 | 0.84 | 0.84 | 97.9 | 6.19 | 0 | 4.83 | 1.88 | 0.1995 | 0.2175 | 0.018 |
| 2eix | opls | 1.56 | 0.1003 | 2.3 | 0 | 99.38 | 1.17 | 1 | 1.13 | 0.87 | 0.1743 | 0.1931 | 0.0188 |
| 2eix | noschrodinger | 1.56 | 0.0066 | 1 | 0 | 99.59 | 1.64 | 0 | 2 | 1.13 | 0.1792 | 0.1946 | 0.0154 |
| 2erb | opls | 1.5 | 0.1053 | 2.31 | 0 | 99.17 | 0.88 | 1 | 2.41 | 1.02 | 0.1452 | 0.1698 | 0.0246 |
| 2erb | noschrodinger | 1.5 | 0.0123 | 1.28 | 0 | 99.17 | 1.75 | 0 | 4.34 | 1.4 | 0.1433 | 0.1712 | 0.0279 |
| 2erv | opls | 2 | 0.0926 | 2.17 | 0 | 97.56 | 1.35 | 1 | 0.64 | 0.9 | 0.2052 | 0.2273 | 0.0221 |
| 2erv | noschrodinger | 2 | 0.0026 | 0.79 | 0 | 97.59 | 1.35 | 0 | 1.55 | 1.08 | 0.1938 | 0.2131 | 0.0193 |
| 2esr | opls | 1.8 | 0.0974 | 2.35 | 0 | 97.2 | 1.19 | 4 | 3.55 | 1.35 | 0.1882 | 0.2195 | 0.0313 |
| 2esr | noschrodinger | 1.8 | 0.0046 | 1.25 | 0 | 96.2 | 0.76 | 0 | 2.77 | 1.32 | 0.1966 | 0.2207 | 0.0241 |
| 2et1 | opls | 1.61 | 0.1004 | 2.3 | 0 | 97.98 | 0 | 1 | 1.01 | 0.8 | 0.142 | 0.1744 | 0.0324 |

| | | | | | | | | | | | | | |
|---|---|---|---|---|---|---|---|---|---|---|---|---|---|
| 2et1 | noschrodinger | 1.61 | 0.0063 | 1.12 | 0 | 97.49 | 0.6 | 0 | 3.35 | 1.23 | 0.1471 | 0.1739 | 0.0268 |
| 2ev1 | opls | 1.6 | 0.1007 | 2.54 | 0 | 98.32 | 1.32 | 3 | 4.9 | 1.35 | 0.1871 | 0.2163 | 0.0292 |
| 2ev1 | noschrodinger | 1.6 | 0.009 | 1.06 | 0 | 98.07 | 0.99 | 0 | 5.43 | 1.29 | 0.1862 | 0.2165 | 0.0303 |
| 2ev3 | opls | 2.68 | 0.0909 | 2.35 | 0.28 | 96.14 | 1.71 | 0 | 0.35 | 1.07 | 0.2535 | 0.2845 | 0.031 |
| 2ev3 | noschrodinger | 2.68 | 0.0032 | 0.77 | 0.55 | 95.07 | 3.06 | 0 | 1.73 | 1.64 | 0.2497 | 0.277 | 0.0273 |
| 2ev4 | opls | 2.28 | 0.0939 | 2.35 | 0.55 | 97.52 | 2.01 | 1 | 2.56 | 1.37 | 0.2422 | 0.2638 | 0.0216 |
| 2ev4 | noschrodinger | 2.28 | 0.0033 | 0.83 | 0.82 | 95.62 | 3.32 | 0 | 5.8 | 2.02 | 0.2168 | 0.2511 | 0.0343 |
| 2evd | opls | 2 | 0.0978 | 2.31 | 0 | 97.47 | 1.69 | 0 | 0.57 | 0.97 | 0.2011 | 0.2281 | 0.027 |
| 2evd | noschrodinger | 2 | 0.0031 | 0.79 | 0 | 98.5 | 1.68 | 0 | 2.86 | 1.25 | 0.1763 | 0.2193 | 0.043 |
| 2eve | opls | 1.55 | 0.1042 | 2.52 | 0 | 99.27 | 0.83 | 2 | 3.73 | 1.16 | 0.1588 | 0.1911 | 0.0323 |
| 2eve | noschrodinger | 1.55 | 0.0118 | 1.17 | 0 | 99.32 | 0 | 0 | 1.25 | 0.84 | 0.1635 | 0.191 | 0.0275 |
| 2ews | opls | 2.05 | 0.0974 | 2.24 | 0.79 | 97.43 | 2.42 | 1 | 0.75 | 1.14 | 0.2139 | 0.2489 | 0.035 |
| 2ews | noschrodinger | 2.05 | 0.0028 | 0.78 | 0.59 | 95.69 | 2.16 | 0 | 2.25 | 1.56 | 0.2053 | 0.2351 | 0.0298 |
| 2f07 | opls | 2.33 | 0.0928 | 2.29 | 0.52 | 97.66 | 4.37 | 2 | 0.47 | 1.22 | 0.2026 | 0.2333 | 0.0307 |
| 2f07 | noschrodinger | 2.33 | 0.0097 | 1.16 | 0.77 | 95.63 | 6.36 | 0 | 4.5 | 2.14 | 0.1942 | 0.2328 | 0.0386 |
| 2f17 | opls | 2.5 | 0.093 | 2.35 | 0.61 | 95.34 | 2.28 | 1 | 0.88 | 1.36 | 0.1979 | 0.2397 | 0.0418 |
| 2f17 | noschrodinger | 2.5 | 0.0027 | 0.79 | 1.21 | 93.52 | 2.97 | 0 | 4.01 | 1.97 | 0.2131 | 0.2495 | 0.0364 |
| 2f1t | opls | 2.44 | 0.0875 | 2.51 | 1.88 | 92.84 | 6.25 | 1 | 1.37 | 1.93 | 0.3445 | 0.3659 | 0.0214 |
| 2f1t | noschrodinger | 2.44 | 0.003 | 0.88 | 4.1 | 88.08 | 9.66 | 0 | 10.65 | 2.89 | 0.3296 | 0.3493 | 0.0197 |
| 2f2c | opls | 2.79 | 0.0884 | 2.33 | 0.7 | 94.85 | 5.81 | 5 | 1.22 | 1.78 | 0.3035 | 0.3222 | 0.0187 |
| 2f2c | noschrodinger | 2.79 | 0.0028 | 0.81 | 1.97 | 89.57 | 7.62 | 0 | 2.97 | 2.32 | 0.2934 | 0.3156 | 0.0222 |
| 2f2e | opls | 1.85 | 0.1037 | 2.43 | 0.76 | 96.97 | 1.32 | 3 | 1.57 | 1.17 | 0.1703 | 0.2194 | 0.0491 |
| 2f2e | noschrodinger | 1.85 | 0.0114 | 1.15 | 0.72 | 96.4 | 2.15 | 0 | 5.68 | 1.8 | 0.1656 | 0.2151 | 0.0495 |
| 2f2l | opls | 2.06 | 0.0962 | 2.24 | 0 | 99.07 | 1.42 | 4 | 0.9 | 0.89 | 0.19 | 0.223 | 0.033 |
| 2f2l | noschrodinger | 2.06 | 0.0065 | 0.96 | 0 | 98.47 | 2.46 | 0 | 2.72 | 1.36 | 0.1753 | 0.2155 | 0.0402 |
| 2f3x | opls | 3.1 | 0.0897 | 2.46 | 0 | 97.74 | 7.5 | 3 | 1.3 | 1.58 | 0.2377 | 0.2519 | 0.0142 |
| 2f3x | noschrodinger | 3.1 | 0.0032 | 0.82 | 1.77 | 93.26 | 13.64 | 0 | 3.07 | 2.4 | 0.2071 | 0.243 | 0.0359 |
| 2f4o | opls | 2.26 | 0.0913 | 2.35 | 0.59 | 91.79 | 4.62 | 1 | 0.68 | 1.72 | 0.2798 | 0.3053 | 0.0255 |
| 2f4o | noschrodinger | 2.26 | 0.0027 | 0.77 | 1.16 | 92.2 | 8.26 | 0 | 1.89 | 2.13 | 0.2464 | 0.2867 | 0.0403 |
| 2f62 | opls | 1.5 | 0.0991 | 2.29 | 0 | 98.51 | 0.74 | 3 | 2.29 | 1.01 | 0.1504 | 0.1799 | 0.0295 |
| 2f62 | noschrodinger | 1.5 | 0.0078 | 0.96 | 0 | 98.72 | 1.74 | 0 | 1.14 | 1.01 | 0.1529 | 0.1769 | 0.024 |
| 2f8d | opls | 2.7 | 0.1017 | 2.26 | 0 | 98.15 | 2.59 | 2 | 0.28 | 0.92 | 0.1776 | 0.2088 | 0.0312 |
| 2f8d | noschrodinger | 2.7 | 0.0029 | 0.79 | 0 | 97.7 | 4.65 | 0 | 1.38 | 1.44 | 0.1626 | 0.2043 | 0.0417 |
| 2fa1 | opls | 1.6 | 0.1021 | 2.45 | 0.64 | 97.76 | 3.1 | 2 | 4.19 | 1.63 | 0.193 | 0.2191 | 0.0261 |
| 2fa1 | noschrodinger | 1.6 | 0.0056 | 0.98 | 0.96 | 96.18 | 5.17 | 0 | 4.94 | 2.06 | 0.1869 | 0.2174 | 0.0305 |
| 2fde | opls | 2.71 | 0.0943 | 2.32 | 0 | 97.94 | 3.01 | 1 | 2.17 | 1.37 | 0.2012 | 0.2466 | 0.0454 |
| 2fde | noschrodinger | 2.71 | 0.0029 | 0.78 | 0 | 97.42 | 7.23 | 0 | 4.35 | 1.98 | 0.1867 | 0.2332 | 0.0465 |
| 2feo | opls | 2.7 | 0.093 | 2.49 | 3.3 | 90.57 | 7.47 | 1 | 2.32 | 2.21 | 0.2851 | 0.3293 | 0.0442 |
| 2feo | noschrodinger | 2.7 | 0.0148 | 1.82 | 12.79 | 66.67 | 15.43 | 1 | 46.94 | 3.92 | 0.2798 | 0.337 | 0.0572 |
| 2ff3 | opls | 2 | 0.0987 | 2.27 | 0 | 98.32 | 4.18 | 5 | 1.44 | 1.35 | 0.1838 | 0.211 | 0.0272 |
| 2ff3 | noschrodinger | 2 | 0.0089 | 1.02 | 0 | 98.12 | 4.9 | 0 | 3.68 | 1.68 | 0.1818 | 0.2111 | 0.0293 |
| 2fh5 | opls | 2.45 | 0.0947 | 2.32 | 0.34 | 96.98 | 3.65 | 2 | 1.18 | 1.43 | 0.1883 | 0.2254 | 0.0371 |
| 2fh5 | noschrodinger | 2.45 | 0.0027 | 0.79 | 0.66 | 96.36 | 4.32 | 0 | 1.58 | 1.63 | 0.1933 | 0.2289 | 0.0356 |
| 2fiw | opls | 2.35 | 0.0912 | 2.39 | 0 | 99.22 | 4.27 | 2 | 1.94 | 1.44 | 0.2155 | 0.2471 | 0.0316 |
| 2fiw | noschrodinger | 2.35 | 0.0029 | 0.84 | 0 | 98.1 | 9.3 | 0 | 2.72 | 1.8 | 0.2162 | 0.2371 | 0.0209 |
| 2fj1 | opls | 2.2 | 0.0963 | 2.37 | 0 | 96.08 | 6.25 | 1 | 0.59 | 1.57 | 0.1741 | 0.209 | 0.0349 |
| 2fj1 | noschrodinger | 2.2 | 0.0046 | 1.07 | 0.98 | 95.61 | 6.78 | 0 | 2.64 | 1.99 | 0.1617 | 0.2035 | 0.0418 |
| 2fre | opls | 1.9 | 0.1001 | 2.31 | 0 | 99.18 | 0.94 | 1 | 2.38 | 1.02 | 0.1746 | 0.2128 | 0.0382 |
| 2fre | noschrodinger | 1.9 | 0.0041 | 0.87 | 0 | 99.24 | 2.42 | 0 | 2.39 | 1.31 | 0.1831 | 0.2134 | 0.0303 |

| | | | | | | | | | | | | | |
|---|---|---|---|---|---|---|---|---|---|---|---|---|---|
| 2ft0 | opls | 1.66 | 0.1012 | 2.38 | 0 | 96.11 | 1.79 | 4 | 1.89 | 1.41 | 0.1793 | 0.2127 | 0.0334 |
| 2ft0 | noschrodinger | 1.66 | 0.0083 | 1.06 | 0 | 96.14 | 2.09 | 0 | 2.76 | 1.57 | 0.1835 | 0.211 | 0.0275 |
| 2fue | opls | 1.75 | 0.101 | 2.34 | 0 | 97.01 | 0.47 | 0 | 2.53 | 1.21 | 0.1991 | 0.2355 | 0.0364 |
| 2fue | noschrodinger | 1.75 | 0.0049 | 0.81 | 0 | 97.54 | 0.92 | 0 | 0.77 | 0.84 | 0.2037 | 0.2345 | 0.0308 |
| 2fxv | opls | 2.05 | 0.0953 | 2.28 | 0.27 | 96.24 | 1.9 | 0 | 0.5 | 1.14 | 0.2041 | 0.2284 | 0.0243 |
| 2fxv | noschrodinger | 2.05 | 0.0028 | 0.8 | 0.8 | 95.74 | 2.52 | 0 | 1.51 | 1.49 | 0.1932 | 0.2224 | 0.0292 |
| 2g09 | opls | 2.1 | 0.1049 | 2.27 | 0 | 98.69 | 0.79 | 2 | 1.38 | 0.87 | 0.1867 | 0.2135 | 0.0268 |
| 2g09 | noschrodinger | 2.1 | 0.0027 | 0.76 | 0 | 97.92 | 1.73 | 0 | 1.38 | 1.07 | 0.1731 | 0.2048 | 0.0317 |
| 2g1a | opls | 2 | 0.1064 | 2.25 | 0 | 98.3 | 0.57 | 4 | 0.61 | 0.7 | 0.1655 | 0.2072 | 0.0417 |
| 2g1a | noschrodinger | 2 | 0.003 | 0.83 | 0 | 98.79 | 1.98 | 0 | 2.3 | 1.23 | 0.1563 | 0.2027 | 0.0464 |
| 2g7c | opls | 2 | 0.0984 | 2.21 | 0.61 | 97.15 | 1.53 | 5 | 1.98 | 1.26 | 0.1694 | 0.2051 | 0.0357 |
| 2g7c | noschrodinger | 2 | 0.0058 | 0.96 | 1.01 | 96.97 | 1.77 | 0 | 5.81 | 1.68 | 0.1744 | 0.2059 | 0.0315 |
| 2g7z | opls | 2.05 | 0.0989 | 2.24 | 0 | 97.8 | 0.68 | 0 | 1.15 | 0.87 | 0.183 | 0.2008 | 0.0178 |
| 2g7z | noschrodinger | 2.05 | 0.0036 | 0.92 | 0 | 97.42 | 1.76 | 0 | 3.23 | 1.42 | 0.1715 | 0.1949 | 0.0234 |
| 2g8j | opls | 1.61 | 0.0952 | 2.35 | 0.36 | 95.29 | 2.55 | 7 | 3.2 | 1.75 | 0.1617 | 0.1908 | 0.0291 |
| 2g8j | noschrodinger | 1.61 | 0.0041 | 0.91 | 0.94 | 95 | 3.09 | 0 | 1.88 | 1.67 | 0.1657 | 0.1932 | 0.0275 |
| 2gc0 | opls | 2 | 0.0997 | 2.16 | 0 | 98.1 | 2.26 | 0 | 1.5 | 1.16 | 0.179 | 0.2111 | 0.0321 |
| 2gc0 | noschrodinger | 2 | 0.0056 | 0.85 | 0 | 98.39 | 2.55 | 0 | 1.33 | 1.17 | 0.1643 | 0.1999 | 0.0356 |
| 2gdj | opls | 2.39 | 0.0904 | 2.29 | 0 | 97.82 | 3.52 | 1 | 0.8 | 1.21 | 0.2088 | 0.2571 | 0.0483 |
| 2gdj | noschrodinger | 2.39 | 0.0031 | 0.79 | 0.43 | 95.24 | 4.98 | 0 | 2.94 | 1.95 | 0.1907 | 0.246 | 0.0553 |
| 2ggs | opls | 1.7 | 0.101 | 2.42 | 1.11 | 95.36 | 2.97 | 12 | 4.33 | 1.9 | 0.1844 | 0.2251 | 0.0407 |
| 2ggs | noschrodinger | 1.7 | 0.0089 | 1.16 | 1.66 | 94.1 | 4.64 | 0 | 5.89 | 2.23 | 0.1881 | 0.2209 | 0.0328 |
| 2gja | opls | 1.85 | 0.0971 | 2.56 | 0 | 98.14 | 3.82 | 3 | 2.14 | 1.43 | 0.1761 | 0.2119 | 0.0358 |
| 2gja | noschrodinger | 1.85 | 0.0077 | 1.12 | 0 | 98.15 | 3.79 | 0 | 2.92 | 1.52 | 0.1821 | 0.211 | 0.0289 |
| 2gkj | opls | 1.7 | 0.1019 | 2.29 | 0 | 97.79 | 1.69 | 2 | 2.32 | 1.23 | 0.1394 | 0.1704 | 0.031 |
| 2gkj | noschrodinger | 1.7 | 0.0045 | 0.91 | 0 | 97.79 | 1.26 | 0 | 0.46 | 0.79 | 0.1437 | 0.1703 | 0.0266 |
| 2gpj | opls | 2.2 | 0.0923 | 2.62 | 0 | 97.24 | 1.02 | 0 | 1.48 | 1.03 | 0.203 | 0.2362 | 0.0332 |
| 2gpj | noschrodinger | 2.2 | 0.0028 | 0.79 | 0 | 96.28 | 1.49 | 0 | 2.97 | 1.47 | 0.192 | 0.2257 | 0.0337 |
| 2gqu | opls | 1.6 | 0.1056 | 2.48 | 0 | 98.11 | 1.42 | 1 | 7.16 | 1.51 | 0.1576 | 0.2007 | 0.0431 |
| 2gqu | noschrodinger | 1.6 | 0.0056 | 0.95 | 0.38 | 97.73 | 2.37 | 0 | 7.16 | 1.74 | 0.1611 | 0.1984 | 0.0373 |
| 2h55 | opls | 1.99 | 0.1052 | 2.38 | 0 | 97.07 | 0 | 1 | 2.3 | 1.17 | 0.1386 | 0.1768 | 0.0382 |
| 2h55 | noschrodinger | 1.99 | 0.0039 | 0.89 | 0 | 97.58 | 0.53 | 0 | 1.72 | 1.01 | 0.1454 | 0.1784 | 0.033 |
| 2h8g | opls | 1.5 | 0.1059 | 2.59 | 0.21 | 96.91 | 2.22 | 3 | 3.59 | 1.6 | 0.1562 | 0.1949 | 0.0387 |
| 2h8g | noschrodinger | 1.5 | 0.0066 | 1.13 | 0 | 97.12 | 1.23 | 0 | 5.06 | 1.49 | 0.1611 | 0.1899 | 0.0288 |
| 2hj3 | opls | 2.5 | 0.0951 | 2.61 | 0.51 | 98.46 | 2.79 | 0 | 2.29 | 1.35 | 0.23 | 0.278 | 0.048 |
| 2hj3 | noschrodinger | 2.5 | 0.0031 | 0.77 | 1.02 | 95.43 | 3.33 | 0 | 4.01 | 1.9 | 0.2186 | 0.2641 | 0.0455 |
| 2hmv | opls | 2.2 | 0.0959 | 2.41 | 0 | 97.76 | 7.08 | 5 | 1.36 | 1.57 | 0.2322 | 0.2581 | 0.0259 |
| 2hmv | noschrodinger | 2.2 | 0.0027 | 0.83 | 0 | 97.04 | 7.49 | 0 | 4.79 | 2.08 | 0.2198 | 0.2564 | 0.0366 |
| 2hq9 | opls | 1.95 | 0.0953 | 2.39 | 0 | 98.7 | 0.91 | 2 | 3.53 | 1.14 | 0.185 | 0.2125 | 0.0275 |
| 2hq9 | noschrodinger | 1.95 | 0.0069 | 0.87 | 0 | 98.49 | 0.88 | 0 | 3.32 | 1.12 | 0.188 | 0.2109 | 0.0229 |
| 2htw | opls | 3.5 | 0.0898 | 2.3 | 1.04 | 89.06 | 3.04 | 1 | 1.7 | 1.86 | 0.2326 | 0.2614 | 0.0288 |
| 2htw | noschrodinger | 3.5 | 0.0031 | 0.82 | 2.07 | 80.83 | 5.44 | 0 | 8.52 | 2.74 | 0.202 | 0.239 | 0.037 |
| 2hvw | opls | 1.67 | 0.1069 | 3.69 | 0 | 98.14 | 2.01 | 1 | 7.32 | 1.63 | 0.124 | 0.153 | 0.029 |
| 2hvw | noschrodinger | 1.67 | 0.0076 | 1.63 | 0 | 98.38 | 1.75 | 1 | 7.28 | 1.58 | 0.1268 | 0.151 | 0.0242 |
| 2hw1 | opls | 2.1 | 0.1001 | 2.3 | 0 | 98.28 | 1.63 | 0 | 0.65 | 0.87 | 0.183 | 0.2124 | 0.0294 |
| 2hw1 | noschrodinger | 2.1 | 0.0029 | 0.84 | 0 | 98.64 | 2.04 | 0 | 1.51 | 1.13 | 0.1807 | 0.2109 | 0.0302 |
| 2hx5 | opls | 1.5 | 0.1003 | 2.43 | 0 | 97.71 | 1.59 | 0 | 4.09 | 1.41 | 0.1529 | 0.1765 | 0.0236 |
| 2hx5 | noschrodinger | 1.5 | 0.0123 | 1.22 | 0 | 98.59 | 1.55 | 0 | 2.45 | 1.17 | 0.1546 | 0.1733 | 0.0187 |
| 2hza | opls | 2 | 0.0879 | 2.37 | 0 | 98.97 | 2.03 | 0 | 0.51 | 0.91 | 0.2402 | 0.2881 | 0.0479 |

| | | | | | | | | | | | | | |
|---|---|---|---|---|---|---|---|---|---|---|---|---|---|
| 2hza | noschrodinger | 2 | 0.0029 | 0.75 | 0.8 | 96.02 | 3.47 | 0 | 2.61 | 1.73 | 0.2374 | 0.2814 | 0.044 |
| 2i55 | opls | 2.9 | 0.0908 | 2.27 | 0 | 96.09 | 5.13 | 3 | 0.34 | 1.43 | 0.2432 | 0.2741 | 0.0309 |
| 2i55 | noschrodinger | 2.9 | 0.0033 | 0.79 | 0.56 | 92.36 | 6.7 | 0 | 1.9 | 2.05 | 0.2341 | 0.2696 | 0.0355 |
| 2i5e | opls | 2.1 | 0.0938 | 2.37 | 0 | 96.09 | 7.14 | 6 | 1.48 | 1.8 | 0.2105 | 0.2422 | 0.0317 |
| 2i5e | noschrodinger | 2.1 | 0.0032 | 1.05 | 0 | 94.71 | 11.88 | 0 | 4.9 | 2.44 | 0.2028 | 0.2326 | 0.0298 |
| 2i6k | opls | 1.99 | 0.0939 | 2.33 | 0 | 96.71 | 2.01 | 3 | 3.12 | 1.54 | 0.208 | 0.241 | 0.033 |
| 2i6k | noschrodinger | 1.99 | 0.0072 | 0.98 | 0 | 95.68 | 3.2 | 1 | 4.22 | 1.89 | 0.2045 | 0.2359 | 0.0314 |
| 2i6p | opls | 2.28 | 0.0924 | 2.34 | 0.65 | 92.26 | 7.97 | 2 | 1.92 | 2.12 | 0.2158 | 0.2729 | 0.0571 |
| 2i6p | noschrodinger | 2.28 | 0.0027 | 0.76 | 3.82 | 84.71 | 5 | 0 | 6.16 | 2.53 | 0.2263 | 0.2683 | 0.042 |
| 2i76 | opls | 2.6 | 0.0899 | 2.5 | 1.89 | 91.96 | 6.55 | 2 | 3.9 | 2.28 | 0.2608 | 0.2925 | 0.0317 |
| 2i76 | noschrodinger | 2.6 | 0.0031 | 0.81 | 3.82 | 83.23 | 9.25 | 0 | 6.96 | 2.81 | 0.2352 | 0.2846 | 0.0494 |
| 2ibn | opls | 1.5 | 0.1003 | 2.43 | 0 | 96.77 | 1.23 | 3 | 2.8 | 1.34 | 0.1868 | 0.2123 | 0.0255 |
| 2ibn | noschrodinger | 1.5 | 0.0139 | 1.3 | 0 | 97.41 | 1.61 | 0 | 2.93 | 1.36 | 0.1881 | 0.2126 | 0.0245 |
| 2ifr | opls | 1.95 | 0.3632 | 7.54 | 2 | 91.5 | 4.02 | 21 | 59.67 | 3.21 | 0.187 | 0.2174 | 0.0304 |
| 2ifr | noschrodinger | 1.95 | 0.0077 | 1.36 | 0.99 | 94.55 | 2.27 | 0 | 5.69 | 1.95 | 0.1838 | 0.2084 | 0.0246 |
| 2ihb | opls | 2.71 | 0.0886 | 2.23 | 0 | 98.54 | 0.85 | 1 | 0.15 | 0.56 | 0.2162 | 0.2412 | 0.025 |
| 2ihb | noschrodinger | 2.71 | 0.003 | 0.75 | 0 | 97.87 | 1.98 | 0 | 0.6 | 0.96 | 0.2 | 0.2347 | 0.0347 |
| 2iho | opls | 2.41 | 0.0959 | 2.22 | 0.69 | 96.19 | 1.67 | 1 | 1.73 | 1.35 | 0.1875 | 0.215 | 0.0275 |
| 2iho | noschrodinger | 2.41 | 0.003 | 0.8 | 1.03 | 95.52 | 1.66 | 0 | 4.53 | 1.71 | 0.1852 | 0.2182 | 0.033 |
| 2il4 | opls | 2.05 | 0.0959 | 2.42 | 0 | 97.7 | 0 | 0 | 0 | 0.57 | 0.1709 | 0.2163 | 0.0454 |
| 2il4 | noschrodinger | 2.05 | 0.008 | 0.99 | 0 | 97.73 | 2.44 | 0 | 0.65 | 1.07 | 0.1657 | 0.2136 | 0.0479 |
| 2img | opls | 1.93 | 0.0997 | 2.33 | 0 | 95.83 | 2.5 | 1 | 1.28 | 1.44 | 0.1629 | 0.2017 | 0.0388 |
| 2img | noschrodinger | 1.93 | 0.0039 | 0.88 | 0 | 95.92 | 2.48 | 0 | 5.57 | 1.88 | 0.1681 | 0.204 | 0.0359 |
| 2ite | opls | 1.6 | 0.1064 | 2.2 | 0 | 100 | 1.39 | 0 | 3.3 | 1.23 | 0.1494 | 0.1893 | 0.0399 |
| 2ite | noschrodinger | 1.6 | 0.0044 | 0.94 | 0 | 99.16 | 2.27 | 0 | 2.8 | 1.34 | 0.1534 | 0.1864 | 0.033 |
| 2ivf | opls | 1.88 | 0.0997 | 2.78 | 0 | 96.9 | 1.73 | 10 | 2.54 | 1.4 | 0.1554 | 0.1784 | 0.023 |
| 2ivf | noschrodinger | 1.88 | 0.0133 | 1.4 | 0 | 96.77 | 1.81 | 0 | 4.56 | 1.63 | 0.1569 | 0.1843 | 0.0274 |
| 2iw5 | opls | 2.57 | 0.0919 | 2.25 | 0 | 98.11 | 0.88 | 1 | 0.63 | 0.71 | 0.2328 | 0.2477 | 0.0149 |
| 2iw5 | noschrodinger | 2.57 | 0.0028 | 0.75 | 0.13 | 97.74 | 2.2 | 0 | 0.78 | 1.06 | 0.2363 | 0.2494 | 0.0131 |
| 2ixh | opls | 2 | 0.1067 | 2.25 | 0.28 | 98.34 | 1.32 | 1 | 0.67 | 0.81 | 0.2252 | 0.2517 | 0.0265 |
| 2ixh | noschrodinger | 2 | 0.0038 | 0.82 | 0.55 | 98.08 | 2.61 | 0 | 1.51 | 1.21 | 0.2003 | 0.2437 | 0.0434 |
| 2j0b | opls | 2.1 | 0.0942 | 2.33 | 0.43 | 97.86 | 4.72 | 2 | 1.78 | 1.48 | 0.1916 | 0.2122 | 0.0206 |
| 2j0b | noschrodinger | 2.1 | 0.0031 | 0.87 | 0.42 | 97.47 | 5.12 | 0 | 2.56 | 1.69 | 0.1781 | 0.1978 | 0.0197 |
| 2j34 | opls | 2.01 | 0.0985 | 2.15 | 0 | 97.48 | 0 | 1 | 0.23 | 0.69 | 0.1818 | 0.22 | 0.0382 |
| 2j34 | noschrodinger | 2.01 | 0.0034 | 0.86 | 0 | 97.87 | 0.83 | 0 | 0.68 | 0.75 | 0.1746 | 0.2143 | 0.0397 |
| 2j44 | opls | 2.1 | 0.0979 | 2.36 | 2.35 | 88.26 | 9.42 | 4 | 3.54 | 2.48 | 0.2456 | 0.2819 | 0.0363 |
| 2j44 | noschrodinger | 2.1 | 0.0045 | 1.13 | 2.79 | 87.44 | 12.44 | 0 | 13.1 | 3.07 | 0.2321 | 0.2712 | 0.0391 |
| 2j7l | opls | 2.6 | 0.0926 | 2.18 | 0 | 99.01 | 1.62 | 0 | 1.18 | 0.99 | 0.2186 | 0.2598 | 0.0412 |
| 2j7l | noschrodinger | 2.6 | 0.0032 | 1.21 | 0 | 94.17 | 2.14 | 0 | 4.45 | 1.87 | 0.217 | 0.2622 | 0.0452 |
| 2j7m | opls | 2.3 | 0.1053 | 2.2 | 0 | 99.29 | 0.83 | 1 | 0.45 | 0.66 | 0.1911 | 0.2273 | 0.0362 |
| 2j7m | noschrodinger | 2.3 | 0.0083 | 0.96 | 0 | 97.87 | 0.83 | 0 | 3.16 | 1.14 | 0.1764 | 0.2153 | 0.0389 |
| 2j83 | opls | 2 | 0.0996 | 2.21 | 0 | 97.23 | 0.45 | 4 | 0.37 | 0.78 | 0.161 | 0.1839 | 0.0229 |
| 2j83 | noschrodinger | 2 | 0.0028 | 0.86 | 0.2 | 97.24 | 0.67 | 0 | 1.37 | 1.01 | 0.1604 | 0.1765 | 0.0161 |
| 2j9r | opls | 2.7 | 0.0917 | 2.34 | 0 | 98.26 | 3.36 | 1 | 0.71 | 1.13 | 0.2084 | 0.2304 | 0.022 |
| 2j9r | noschrodinger | 2.7 | 0.003 | 0.79 | 0 | 95.98 | 5.3 | 0 | 2.5 | 1.86 | 0.2019 | 0.2333 | 0.0314 |
| 2jao | opls | 2 | 0.1001 | 2.47 | 0 | 98.45 | 1.1 | 1 | 1.81 | 0.97 | 0.1675 | 0.186 | 0.0185 |
| 2jao | noschrodinger | 2 | 0.0036 | 1.31 | 0 | 97.94 | 1.64 | 0 | 2.11 | 1.16 | 0.1578 | 0.1819 | 0.0241 |
| 2jaq | opls | 2.3 | 0.0928 | 2.26 | 0 | 97.75 | 2.03 | 3 | 0.62 | 1 | 0.2268 | 0.2552 | 0.0284 |
| 2jaq | noschrodinger | 2.3 | 0.0028 | 0.74 | 0.27 | 96.7 | 3.98 | 0 | 1.87 | 1.61 | 0.2156 | 0.2557 | 0.0401 |

| | | | | | | | | | | | | | |
|---|---|---|---|---|---|---|---|---|---|---|---|---|---|
| 2jat | opls | 2.6 | 0.09 | 2.26 | 0 | 98.63 | 2.86 | 2 | 0.31 | 0.96 | 0.2438 | 0.2582 | 0.0144 |
| 2jat | noschrodinger | 2.6 | 0.0028 | 0.74 | 0 | 95.97 | 5.04 | 0 | 2.46 | 1.84 | 0.244 | 0.2677 | 0.0237 |
| 2jcb | opls | 1.6 | 0.0995 | 2.32 | 0.26 | 97.35 | 3.45 | 3 | 1.87 | 1.48 | 0.1898 | 0.2294 | 0.0396 |
| 2jcb | noschrodinger | 1.6 | 0.0029 | 0.84 | 0.79 | 96.58 | 3.71 | 0 | 2.8 | 1.72 | 0.1971 | 0.2296 | 0.0325 |
| 2jdc | opls | 1.6 | 0.0995 | 2.29 | 0 | 99.3 | 0.83 | 0 | 2.08 | 0.98 | 0.1529 | 0.1893 | 0.0364 |
| 2jdc | noschrodinger | 1.6 | 0.0059 | 1.01 | 0 | 99.3 | 0 | 0 | 0.42 | 0.65 | 0.1534 | 0.1867 | 0.0333 |
| 2jf2 | opls | 1.8 | 0.1071 | 2.91 | 0 | 98.47 | 4.11 | 3 | 6.37 | 1.82 | 0.1491 | 0.1895 | 0.0404 |
| 2jf2 | noschrodinger | 1.8 | 0.0057 | 1.03 | 0 | 98.47 | 1.37 | 0 | 7.07 | 1.49 | 0.1532 | 0.1855 | 0.0323 |
| 2jf3 | opls | 3 | 0.0916 | 2.36 | 0 | 98.07 | 1.46 | 3 | 0.25 | 0.72 | 0.1903 | 0.226 | 0.0357 |
| 2jf3 | noschrodinger | 3 | 0.003 | 0.78 | 0 | 97.31 | 0.48 | 0 | 0.75 | 0.87 | 0.1863 | 0.2115 | 0.0252 |
| 2jiv | opls | 3.5 | 0.0906 | 2.46 | 1.23 | 91.56 | 13.57 | 5 | 2.25 | 2.36 | 0.2506 | 0.2873 | 0.0367 |
| 2jiv | noschrodinger | 3.5 | 0.0034 | 0.85 | 5.66 | 74.15 | 28.91 | 0 | 15.58 | 3.61 | 0.2308 | 0.2709 | 0.0401 |
| 2no5 | opls | 2.6 | 0.0927 | 2.3 | 0 | 97.49 | 3.64 | 4 | 1.37 | 1.4 | 0.2166 | 0.238 | 0.0214 |
| 2no5 | noschrodinger | 2.6 | 0.0027 | 0.78 | 0 | 96.83 | 7.49 | 0 | 1.23 | 1.7 | 0.2006 | 0.2371 | 0.0365 |
| 2nr4 | opls | 1.85 | 0.0969 | 2.26 | 0 | 97.18 | 1.6 | 0 | 2.32 | 1.32 | 0.2113 | 0.2358 | 0.0245 |
| 2nr4 | noschrodinger | 1.85 | 0.0051 | 0.87 | 0 | 96.95 | 2.51 | 0 | 3.5 | 1.62 | 0.2117 | 0.2338 | 0.0221 |
| 2nsl | opls | 2 | 0.0991 | 2.29 | 0 | 98.11 | 0 | 1 | 0.41 | 0.65 | 0.1518 | 0.1922 | 0.0404 |
| 2nsl | noschrodinger | 2 | 0.003 | 0.82 | 0 | 98.12 | 0 | 0 | 2.47 | 1.03 | 0.1588 | 0.1933 | 0.0345 |
| 2nuo | opls | 1.5 | 0.1023 | 3.33 | 0 | 97.48 | 4.65 | 0 | 3.9 | 1.79 | 0.1543 | 0.1863 | 0.032 |
| 2nuo | noschrodinger | 1.5 | 0.0043 | 1.11 | 0 | 97.48 | 1.86 | 0 | 4.63 | 1.55 | 0.1552 | 0.1859 | 0.0307 |
| 2nuy | opls | 2.5 | 0.0944 | 2.21 | 0 | 97.02 | 0.58 | 1 | 0.21 | 0.75 | 0.1887 | 0.2034 | 0.0147 |
| 2nuy | noschrodinger | 2.5 | 0.0028 | 0.79 | 0 | 96.85 | 0.58 | 0 | 0.75 | 0.93 | 0.1736 | 0.1998 | 0.0262 |
| 2nyk | opls | 1.95 | 0.0948 | 2.5 | 0 | 98.58 | 2.37 | 1 | 3.36 | 1.41 | 0.1967 | 0.2418 | 0.0451 |
| 2nyk | noschrodinger | 1.95 | 0.0038 | 0.82 | 0 | 98.24 | 1.86 | 0 | 2.6 | 1.25 | 0.2078 | 0.2365 | 0.0287 |
| 2nzu | opls | 2.09 | 0.0925 | 2.31 | 0.28 | 96.34 | 4.89 | 2 | 0.89 | 1.54 | 0.2464 | 0.2808 | 0.0344 |
| 2nzu | noschrodinger | 2.09 | 0.0027 | 0.77 | 0.56 | 95.21 | 5.54 | 0 | 4.09 | 2.09 | 0.2428 | 0.2701 | 0.0273 |
| 2o28 | opls | 1.8 | 0.0994 | 2.31 | 0 | 98.88 | 2.15 | 1 | 1.34 | 1.11 | 0.1655 | 0.1903 | 0.0248 |
| 2o28 | noschrodinger | 1.8 | 0.0058 | 1.46 | 0 | 98.04 | 1.83 | 0 | 3.19 | 1.31 | 0.164 | 0.1908 | 0.0268 |
| 2o4t | opls | 1.95 | 0.0947 | 2.25 | 0 | 100 | 1.39 | 0 | 0 | 0.61 | 0.2019 | 0.2437 | 0.0418 |
| 2o4t | noschrodinger | 1.95 | 0.0027 | 0.69 | 0 | 98.86 | 1.37 | 0 | 1.42 | 0.98 | 0.2107 | 0.2397 | 0.029 |
| 2ocf | opls | 2.95 | 0.0905 | 2.41 | 0 | 97.43 | 12.32 | 4 | 0.57 | 1.63 | 0.2343 | 0.2707 | 0.0364 |
| 2ocf | noschrodinger | 2.95 | 0.0059 | 0.99 | 0.64 | 94.59 | 16.72 | 0 | 3.43 | 2.44 | 0.2387 | 0.2678 | 0.0291 |
| 2odh | opls | 1.6 | 0.1006 | 2.53 | 0.88 | 96.48 | 2.34 | 4 | 3.56 | 1.66 | 0.2055 | 0.2465 | 0.041 |
| 2odh | noschrodinger | 1.6 | 0.005 | 0.86 | 0.87 | 96.51 | 3.24 | 0 | 3.61 | 1.77 | 0.215 | 0.2487 | 0.0337 |
| 2oh6 | opls | 2.1 | 0.0966 | 2.35 | 0 | 97.55 | 0.46 | 0 | 0.49 | 0.76 | 0.1434 | 0.1895 | 0.0461 |
| 2oh6 | noschrodinger | 2.1 | 0.0027 | 0.84 | 0 | 96.73 | 0 | 0 | 1.49 | 1.09 | 0.1317 | 0.176 | 0.0443 |
| 2oi2 | opls | 2.5 | 0.0925 | 2.31 | 0 | 97.41 | 2.7 | 1 | 0.47 | 1.11 | 0.2122 | 0.2304 | 0.0182 |
| 2oi2 | noschrodinger | 2.5 | 0.003 | 0.77 | 0.36 | 94.66 | 5.86 | 0 | 3.52 | 2.09 | 0.2034 | 0.2261 | 0.0227 |
| 2okg | opls | 1.65 | 0.0999 | 2.34 | 0 | 99.1 | 1 | 5 | 3.92 | 1.18 | 0.1877 | 0.2288 | 0.0411 |
| 2okg | noschrodinger | 1.65 | 0.0061 | 0.93 | 0 | 99.19 | 0.71 | 0 | 2.78 | 1.07 | 0.1904 | 0.2267 | 0.0363 |
| 2on6 | opls | 2.5 | 0.0935 | 2.4 | 0.36 | 98.18 | 5.17 | 2 | 0.68 | 1.26 | 0.1998 | 0.2284 | 0.0286 |
| 2on6 | noschrodinger | 2.5 | 0.0035 | 0.86 | 0.72 | 96.76 | 8.09 | 0 | 2.03 | 1.86 | 0.206 | 0.2336 | 0.0276 |
| 2oo5 | opls | 2.6 | 0.0904 | 2.31 | 0.11 | 96.42 | 2.21 | 1 | 0.49 | 1.17 | 0.2138 | 0.2535 | 0.0397 |
| 2oo5 | noschrodinger | 2.6 | 0.0027 | 0.8 | 0.32 | 96.01 | 2.88 | 0 | 1.12 | 1.44 | 0.2103 | 0.2478 | 0.0375 |
| 2oqo | opls | 2 | 0.0956 | 2.29 | 1.14 | 97.16 | 1.99 | 0 | 1.29 | 1.23 | 0.1983 | 0.2331 | 0.0348 |
| 2oqo | noschrodinger | 2 | 0.0078 | 1.11 | 0.56 | 96.05 | 1.97 | 0 | 5.16 | 1.77 | 0.187 | 0.2256 | 0.0386 |
| 2orv | opls | 2.3 | 0.0964 | 2.41 | 0 | 98.66 | 7.06 | 0 | 1.95 | 1.61 | 0.1941 | 0.2218 | 0.0277 |
| 2orv | noschrodinger | 2.3 | 0.0039 | 0.89 | 0 | 96.83 | 8.53 | 0 | 2.55 | 1.94 | 0.1972 | 0.2187 | 0.0215 |
| 2orw | opls | 1.5 | 0.1012 | 2.23 | 0 | 98.47 | 0.34 | 0 | 1.78 | 0.94 | 0.1627 | 0.184 | 0.0213 |

| | | | | | | | | | | | | | |
|---|---|---|---|---|---|---|---|---|---|---|---|---|---|
| 2orw | noschrodinger | 1.5 | 0.0041 | 0.95 | 0.3 | 98.19 | 1 | 0 | 0.89 | 0.77 | 0.1606 | 0.1848 | 0.0242 |
| 2ou5 | opls | 1.6 | 0.1012 | 2.24 | 0 | 98.09 | 0.35 | 1 | 2.27 | 1.01 | 0.1804 | 0.2117 | 0.0313 |
| 2ou5 | noschrodinger | 1.6 | 0.0041 | 0.91 | 0 | 98.27 | 0.69 | 0 | 2.1 | 0.98 | 0.1802 | 0.2097 | 0.0295 |
| 2ovd | opls | 1.75 | 0.1034 | 2.23 | 0.64 | 96.79 | 1.57 | 0 | 3.96 | 1.53 | 0.18 | 0.2179 | 0.0379 |
| 2ovd | noschrodinger | 1.75 | 0.0044 | 0.86 | 0 | 94.94 | 0.78 | 0 | 1.98 | 1.32 | 0.1842 | 0.218 | 0.0338 |
| 2oys | opls | 1.97 | 0.1001 | 2.23 | 0.23 | 97.91 | 1.71 | 4 | 1.06 | 1.01 | 0.195 | 0.242 | 0.047 |
| 2oys | noschrodinger | 1.97 | 0.0054 | 0.76 | 0.22 | 98.46 | 1.67 | 0 | 1.19 | 1 | 0.1872 | 0.2344 | 0.0472 |
| 2oz5 | opls | 1.72 | 0.0978 | 2.32 | 0 | 96.79 | 1.45 | 2 | 1.81 | 1.26 | 0.1781 | 0.2101 | 0.032 |
| 2oz5 | noschrodinger | 1.72 | 0.0054 | 1 | 0 | 97.48 | 1.93 | 0 | 1.84 | 1.27 | 0.1778 | 0.2108 | 0.033 |
| 2p0d | opls | 1.8 | 0.0971 | 2.5 | 0 | 97.83 | 2.27 | 4 | 1.7 | 1.23 | 0.1748 | 0.2204 | 0.0456 |
| 2p0d | noschrodinger | 1.8 | 0.0035 | 0.92 | 0 | 96.12 | 2.22 | 0 | 2.29 | 1.54 | 0.1869 | 0.2141 | 0.0272 |
| 2p0t | opls | 2.19 | 0.0923 | 2.62 | 0 | 98.59 | 11.11 | 3 | 0.8 | 1.55 | 0.2452 | 0.2951 | 0.0499 |
| 2p0t | noschrodinger | 2.19 | 0.0029 | 0.81 | 0.68 | 96.58 | 15.62 | 0 | 1.2 | 1.96 | 0.2409 | 0.2922 | 0.0513 |
| 2p1m | opls | 1.8 | 0.1041 | 2.37 | 0 | 98.44 | 3.64 | 5 | 1.15 | 1.25 | 0.185 | 0.2179 | 0.0329 |
| 2p1m | noschrodinger | 1.8 | 0.0096 | 1.38 | 0 | 98.61 | 4.13 | 0 | 3.06 | 1.56 | 0.1873 | 0.2198 | 0.0325 |
| 2p1p | opls | 2.21 | 0.0986 | 2.35 | 0 | 98.16 | 3.77 | 2 | 0.66 | 1.15 | 0.1987 | 0.2211 | 0.0224 |
| 2p1p | noschrodinger | 2.21 | 0.0037 | 0.97 | 0 | 97.71 | 5.97 | 0 | 1.9 | 1.61 | 0.1927 | 0.2112 | 0.0185 |
| 2p2v | opls | 1.85 | 0.0971 | 2.29 | 0 | 97.39 | 1.2 | 4 | 1.69 | 1.1 | 0.1815 | 0.2049 | 0.0234 |
| 2p2v | noschrodinger | 1.85 | 0.0061 | 0.85 | 0 | 97.79 | 1.19 | 0 | 2.33 | 1.12 | 0.1878 | 0.2042 | 0.0164 |
| 2p39 | opls | 1.5 | 0.0966 | 2.34 | 0 | 98.31 | 0 | 0 | 0.89 | 0.77 | 0.1668 | 0.1883 | 0.0215 |
| 2p39 | noschrodinger | 1.5 | 0.0184 | 1.88 | 0 | 97.14 | 0.87 | 1 | 0.9 | 0.93 | 0.1633 | 0.1894 | 0.0261 |
| 2p71 | opls | 2.01 | 0.0915 | 2.34 | 2.33 | 96.12 | 5.36 | 1 | 0.48 | 1.49 | 0.2374 | 0.2841 | 0.0467 |
| 2p71 | noschrodinger | 2.01 | 0.0034 | 0.83 | 2.31 | 93.08 | 11.5 | 0 | 1.96 | 2.21 | 0.2178 | 0.2702 | 0.0524 |
| 2p9h | opls | 2 | 0.1057 | 2.34 | 0.57 | 94.53 | 3.9 | 6 | 1.1 | 1.64 | 0.1986 | 0.2284 | 0.0298 |
| 2p9h | noschrodinger | 2 | 0.003 | 0.85 | 0.94 | 94.94 | 6.82 | 0 | 5.12 | 2.26 | 0.1835 | 0.2176 | 0.0341 |
| 2pa7 | opls | 1.5 | 0.104 | 2.76 | 0 | 96.58 | 1.21 | 7 | 3.29 | 1.4 | 0.176 | 0.2079 | 0.0319 |
| 2pa7 | noschrodinger | 1.5 | 0.0057 | 1 | 0 | 96.6 | 4.4 | 0 | 4.38 | 1.92 | 0.1829 | 0.2047 | 0.0218 |
| 2pae | opls | 2.5 | 0.0968 | 2.33 | 0 | 96.97 | 3.31 | 1 | 0.45 | 1.23 | 0.2276 | 0.2594 | 0.0318 |
| 2pae | noschrodinger | 2.5 | 0.0036 | 0.77 | 0 | 93.23 | 7.38 | 0 | 4.97 | 2.36 | 0.2097 | 0.2627 | 0.053 |
| 2paf | opls | 3.5 | 0.0887 | 2.5 | 1.21 | 92.91 | 7.53 | 8 | 1.22 | 1.96 | 0.199 | 0.2559 | 0.0569 |
| 2paf | noschrodinger | 3.5 | 0.0061 | 0.96 | 3.18 | 81.65 | 13.64 | 0 | 10.35 | 3.11 | 0.1888 | 0.2336 | 0.0448 |
| 2par | opls | 2.1 | 0.0896 | 2.29 | 0 | 98.63 | 1.1 | 0 | 0.54 | 0.71 | 0.2147 | 0.24 | 0.0253 |
| 2par | noschrodinger | 2.1 | 0.0032 | 0.83 | 0.29 | 98.26 | 1.81 | 0 | 2 | 1.16 | 0.1976 | 0.2319 | 0.0343 |
| 2pke | opls | 1.81 | 0.0949 | 2.32 | 0 | 98.16 | 1.45 | 4 | 1.26 | 0.97 | 0.187 | 0.2216 | 0.0346 |
| 2pke | noschrodinger | 1.81 | 0.0031 | 0.84 | 0.89 | 96.66 | 2.53 | 0 | 3.54 | 1.66 | 0.1948 | 0.222 | 0.0272 |
| 2pkp | opls | 2.1 | 0.1007 | 2.25 | 0.61 | 96.93 | 0 | 0 | 0.38 | 0.82 | 0.1951 | 0.2317 | 0.0366 |
| 2pkp | noschrodinger | 2.1 | 0.0028 | 0.77 | 0 | 98.79 | 0 | 0 | 2.65 | 1.05 | 0.1816 | 0.2287 | 0.0471 |
| 2pnn | opls | 2.5 | 0.0937 | 2.43 | 0.42 | 95.76 | 8.7 | 2 | 1.27 | 1.86 | 0.2478 | 0.2759 | 0.0281 |
| 2pnn | noschrodinger | 2.5 | 0.0035 | 0.91 | 1.22 | 93.9 | 16.59 | 0 | 3.8 | 2.5 | 0.2411 | 0.2691 | 0.028 |
| 2pnz | opls | 2.14 | 0.0956 | 2.34 | 0.2 | 97.59 | 6.92 | 2 | 1.71 | 1.65 | 0.2239 | 0.2432 | 0.0193 |
| 2pnz | noschrodinger | 2.14 | 0.0034 | 0.9 | 0.8 | 96.39 | 9.48 | 0 | 4.65 | 2.22 | 0.2088 | 0.2344 | 0.0256 |
| 2pr6 | opls | 1.94 | 0.101 | 2.27 | 0.43 | 97.83 | 2.22 | 2 | 2.65 | 1.36 | 0.1797 | 0.2162 | 0.0365 |
| 2pr6 | noschrodinger | 1.94 | 0.0032 | 0.84 | 0.4 | 97.59 | 1.72 | 0 | 2.66 | 1.32 | 0.1823 | 0.2103 | 0.028 |
| 2psw | opls | 2.1 | 0.0922 | 2.37 | 0 | 98.7 | 1.82 | 2 | 0.26 | 0.8 | 0.2132 | 0.237 | 0.0238 |
| 2psw | noschrodinger | 2.1 | 0.0028 | 0.82 | 0.22 | 98.23 | 3.06 | 0 | 0.79 | 1.12 | 0.2062 | 0.2304 | 0.0242 |
| 2ptf | opls | 2.35 | 0.094 | 2.31 | 0 | 99.22 | 1.76 | 1 | 1.23 | 1.03 | 0.1891 | 0.2344 | 0.0453 |
| 2ptf | noschrodinger | 2.35 | 0.0026 | 0.74 | 0 | 97.69 | 2.91 | 0 | 1.84 | 1.36 | 0.2016 | 0.231 | 0.0294 |
| 2px3 | opls | 3.2 | 0.0903 | 2.33 | 1.59 | 92.83 | 5.78 | 1 | 0.73 | 1.77 | 0.2308 | 0.2649 | 0.0341 |
| 2px3 | noschrodinger | 3.2 | 0.0031 | 0.84 | 1.98 | 85.32 | 11.5 | 0 | 7.04 | 2.85 | 0.2181 | 0.2568 | 0.0387 |

| | | | | | | | | | | | | | |
|---|---|---|---|---|---|---|---|---|---|---|---|---|---|
| 2q1a | opls | 2.5 | 0.0955 | 2.27 | 0 | 94.43 | 1.17 | 1 | 0.43 | 1.08 | 0.2133 | 0.2278 | 0.0145 |
| 2q1a | noschrodinger | 2.5 | 0.0027 | 0.79 | 0.69 | 93.77 | 2.71 | 0 | 1.28 | 1.59 | 0.2077 | 0.2206 | 0.0129 |
| 2q32 | opls | 2.4 | 0.0939 | 2.23 | 0 | 98.35 | 1.6 | 0 | 0.68 | 0.88 | 0.2088 | 0.2416 | 0.0328 |
| 2q32 | noschrodinger | 2.4 | 0.0029 | 0.76 | 0 | 98.13 | 3.2 | 0 | 1.64 | 1.3 | 0.196 | 0.2287 | 0.0327 |
| 2q37 | opls | 2.5 | 0.0903 | 2.45 | 0.83 | 96.69 | 7.96 | 0 | 1.34 | 1.76 | 0.2345 | 0.2666 | 0.0321 |
| 2q37 | noschrodinger | 2.5 | 0.0028 | 0.77 | 2.17 | 92.75 | 8.33 | 0 | 0.45 | 1.82 | 0.221 | 0.2455 | 0.0245 |
| 2q9p | opls | 1.65 | 0.1011 | 2.45 | 0 | 99.22 | 0.85 | 0 | 2.61 | 1.05 | 0.1871 | 0.2048 | 0.0177 |
| 2q9p | noschrodinger | 1.65 | 0.0087 | 2.97 | 0 | 99.23 | 0 | 0 | 3.5 | 1.14 | 0.178 | 0.2022 | 0.0242 |
| 2qcx | opls | 2.2 | 0.0929 | 2.23 | 0 | 98.87 | 1.87 | 0 | 1.1 | 1.02 | 0.2054 | 0.2284 | 0.023 |
| 2qcx | noschrodinger | 2.2 | 0.0037 | 0.74 | 0.22 | 98.65 | 2.39 | 0 | 1.93 | 1.25 | 0.1871 | 0.2226 | 0.0355 |
| 2qdt | opls | 2 | 0.1009 | 2.25 | 0 | 96.2 | 2.96 | 0 | 0.5 | 1.29 | 0.1862 | 0.201 | 0.0148 |
| 2qdt | noschrodinger | 2 | 0.0027 | 0.81 | 0 | 95.83 | 3.94 | 0 | 1.24 | 1.58 | 0.1755 | 0.1937 | 0.0182 |
| 2qg6 | opls | 1.5 | 0.1064 | 2.46 | 0 | 96.88 | 0 | 0 | 1.65 | 1.1 | 0.1656 | 0.1944 | 0.0288 |
| 2qg6 | noschrodinger | 1.5 | 0.0094 | 1.08 | 0 | 97.75 | 0.59 | 0 | 0.99 | 0.85 | 0.168 | 0.1969 | 0.0289 |
| 2qgu | opls | 1.5 | 0.1084 | 2.32 | 0 | 98.84 | 0.67 | 0 | 4.98 | 1.26 | 0.155 | 0.1833 | 0.0283 |
| 2qgu | noschrodinger | 1.5 | 0.0035 | 0.85 | 0 | 98.87 | 1.32 | 0 | 1.66 | 1.01 | 0.1605 | 0.1813 | 0.0208 |
| 2qhs | opls | 1.5 | 0.1086 | 2.26 | 0 | 96.14 | 0.6 | 0 | 1.81 | 1.2 | 0.1574 | 0.1772 | 0.0198 |
| 2qhs | noschrodinger | 1.5 | 0.004 | 0.91 | 0 | 96.15 | 0.6 | 0 | 1.51 | 1.15 | 0.1612 | 0.1757 | 0.0145 |
| 2qhv | opls | 1.6 | 0.1012 | 2.23 | 0 | 96.6 | 0.6 | 0 | 1.2 | 1.06 | 0.1815 | 0.2002 | 0.0187 |
| 2qhv | noschrodinger | 1.6 | 0.0043 | 0.94 | 0 | 96.63 | 0.6 | 0 | 0.9 | 0.99 | 0.1858 | 0.1996 | 0.0138 |
| 2qib | opls | 1.7 | 0.1043 | 2.28 | 0 | 100 | 1.19 | 0 | 2.47 | 1.09 | 0.1694 | 0.2027 | 0.0333 |
| 2qib | noschrodinger | 1.7 | 0.0069 | 0.97 | 0 | 99.76 | 1.71 | 0 | 3.64 | 1.33 | 0.1638 | 0.2016 | 0.0378 |
| 2qkh | opls | 1.9 | 0.0974 | 2.18 | 0 | 98.32 | 1 | 0 | 3.73 | 1.16 | 0.1658 | 0.1932 | 0.0274 |
| 2qkh | noschrodinger | 1.9 | 0.0125 | 1.21 | 0 | 98.36 | 0.97 | 0 | 3.74 | 1.16 | 0.166 | 0.1966 | 0.0306 |
| 2qmy | opls | 2.5 | 0.0971 | 2.24 | 0 | 96.7 | 1.8 | 4 | 1.21 | 1.24 | 0.1916 | 0.225 | 0.0334 |
| 2qmy | noschrodinger | 2.5 | 0.0044 | 0.87 | 0 | 95.39 | 2.58 | 0 | 3.24 | 1.75 | 0.167 | 0.2214 | 0.0544 |
| 2qu8 | opls | 2.01 | 0.0938 | 2.51 | 0 | 99.42 | 6.51 | 3 | 1.28 | 1.47 | 0.1875 | 0.2384 | 0.0509 |
| 2qu8 | noschrodinger | 2.01 | 0.0027 | 0.82 | 0 | 98.92 | 6.36 | 0 | 2.25 | 1.61 | 0.1968 | 0.2343 | 0.0375 |
| 2qws | opls | 2.52 | 0.0789 | 2.08 | 0 | 95.04 | 0.99 | 0 | 0.54 | 1.03 | 0.3157 | 0.3666 | 0.0509 |
| 2qws | noschrodinger | 2.52 | 0.0023 | 0.71 | 0.81 | 93.5 | 0.97 | 0 | 0.52 | 1.1 | 0.308 | 0.3688 | 0.0608 |
| 2qxx | opls | 1.99 | 0.0952 | 2.28 | 0 | 98.11 | 3.37 | 2 | 1.34 | 1.26 | 0.1609 | 0.1969 | 0.036 |
| 2qxx | noschrodinger | 1.99 | 0.004 | 0.91 | 0 | 98.93 | 4.55 | 0 | 1.34 | 1.36 | 0.1557 | 0.1962 | 0.0405 |
| 2r4o | opls | 3.58 | 0.0897 | 2.34 | 2.4 | 92.57 | 5.79 | 7 | 1.57 | 1.95 | 0.2446 | 0.2834 | 0.0388 |
| 2r4o | noschrodinger | 3.58 | 0.0035 | 0.86 | 4.89 | 79.36 | 9.42 | 0 | 11.72 | 3.07 | 0.2319 | 0.2719 | 0.04 |
| 2r96 | opls | 2.6 | 0.0921 | 2.34 | 0.52 | 97.65 | 4.78 | 5 | 1.83 | 1.53 | 0.1984 | 0.2214 | 0.023 |
| 2r96 | noschrodinger | 2.6 | 0.0031 | 0.91 | 1.03 | 97.18 | 8.19 | 0 | 5.17 | 2.12 | 0.1931 | 0.2138 | 0.0207 |
| 2rde | opls | 1.89 | 0.1033 | 2.32 | 0 | 97.29 | 1.01 | 4 | 2.06 | 1.11 | 0.163 | 0.2021 | 0.0391 |
| 2rde | noschrodinger | 1.89 | 0.0084 | 1.08 | 0 | 97.08 | 1.01 | 0 | 4.12 | 1.36 | 0.1694 | 0.2027 | 0.0333 |
| 2rdg | opls | 1.6 | 0.1069 | 2.31 | 0 | 100 | 0 | 0 | 2.47 | 1.03 | 0.1498 | 0.1789 | 0.0291 |
| 2rdg | noschrodinger | 1.6 | 0.0069 | 1.13 | 0 | 98.42 | 0.56 | 0 | 5.26 | 1.28 | 0.1529 | 0.1778 | 0.0249 |
| 2rds | opls | 1.63 | 0.1062 | 2.52 | 0 | 98.84 | 1.29 | 1 | 7.07 | 1.47 | 0.1514 | 0.1926 | 0.0412 |
| 2rds | noschrodinger | 1.63 | 0.0073 | 1.12 | 0 | 98.46 | 1.28 | 0 | 9.19 | 1.57 | 0.1555 | 0.1873 | 0.0318 |
| 2ref | opls | 2.75 | 0.0893 | 2.43 | 0.5 | 96.25 | 3.17 | 4 | 0.72 | 1.36 | 0.2152 | 0.2505 | 0.0353 |
| 2ref | noschrodinger | 2.75 | 0.0028 | 0.89 | 0.25 | 95.77 | 5.51 | 0 | 3.16 | 1.96 | 0.1967 | 0.2341 | 0.0374 |
| 2rhq | opls | 2.2 | 0.0947 | 2.28 | 0.29 | 96.29 | 2.18 | 4 | 1.02 | 1.31 | 0.2167 | 0.2523 | 0.0356 |
| 2rhq | noschrodinger | 2.2 | 0.0029 | 0.79 | 1.23 | 94.12 | 5.75 | 0 | 3.6 | 2.12 | 0.2019 | 0.2374 | 0.0355 |
| 2rin | opls | 1.8 | 0.0943 | 2.7 | 1.23 | 93.16 | 10.65 | 13 | 12.68 | 2.84 | 0.1544 | 0.1926 | 0.0382 |
| 2rin | noschrodinger | 1.8 | 0.0143 | 1.65 | 0.87 | 89.69 | 15.8 | 0 | 23.78 | 3.34 | 0.1443 | 0.1846 | 0.0403 |
| 2uxi | opls | 2.5 | 0.0941 | 2.39 | 0.27 | 97.28 | 6.44 | 2 | 2.15 | 1.74 | 0.2721 | 0.2944 | 0.0223 |

| | | | | | | | | | | | | | |
|---|---|---|---|---|---|---|---|---|---|---|---|---|---|
| 2uxi | noschrodinger | 2.5 | 0.0028 | 0.76 | 0 | 93.61 | 9.51 | 0 | 6.66 | 2.53 | 0.245 | 0.2853 | 0.0403 |
| 2v1b | opls | 1.55 | 0.095 | 2.42 | 0 | 98.15 | 2.36 | 5 | 2.36 | 1.3 | 0.1545 | 0.1946 | 0.0401 |
| 2v1b | noschrodinger | 1.55 | 0.0076 | 1.14 | 0 | 99.3 | 3.6 | 0 | 2.36 | 1.44 | 0.1598 | 0.1928 | 0.033 |
| 2v2b | opls | 1.5 | 0.1 | 2.48 | 0.37 | 97.4 | 2.21 | 3 | 2.13 | 1.37 | 0.2095 | 0.2333 | 0.0238 |
| 2v2b | noschrodinger | 1.5 | 0.0102 | 1.06 | 0 | 98.15 | 2.21 | 0 | 2.37 | 1.28 | 0.2082 | 0.231 | 0.0228 |
| 2v30 | opls | 2 | 0.1026 | 2.26 | 0 | 98.05 | 0.89 | 2 | 3.03 | 1.09 | 0.1579 | 0.1777 | 0.0198 |
| 2v30 | noschrodinger | 2 | 0.0035 | 0.84 | 0 | 98.25 | 0.89 | 0 | 3.95 | 1.18 | 0.1441 | 0.1677 | 0.0236 |
| 2v5e | opls | 2.35 | 0.0984 | 2.25 | 0 | 96.88 | 1.14 | 0 | 0.63 | 0.94 | 0.1863 | 0.2246 | 0.0383 |
| 2v5e | noschrodinger | 2.35 | 0.0029 | 0.77 | 0.34 | 96.97 | 1.87 | 0 | 2.52 | 1.42 | 0.1761 | 0.2141 | 0.038 |
| 2v6t | opls | 3.1 | 0.0905 | 2.46 | 0.52 | 96.89 | 9.7 | 3 | 1.24 | 1.78 | 0.2187 | 0.239 | 0.0203 |
| 2v6t | noschrodinger | 3.1 | 0.0033 | 0.86 | 1.54 | 91.79 | 13.77 | 0 | 3.72 | 2.52 | 0.2135 | 0.2384 | 0.0249 |
| 2v73 | opls | 2 | 0.0975 | 2.41 | 3.39 | 92.37 | 3.59 | 6 | 1.93 | 1.85 | 0.2303 | 0.2796 | 0.0493 |
| 2v73 | noschrodinger | 2 | 0.006 | 1.02 | 3.6 | 89.47 | 4.9 | 1 | 9.35 | 2.58 | 0.2357 | 0.2745 | 0.0388 |
| 2v8l | opls | 1.8 | 0.0998 | 2.16 | 0 | 93.27 | 0 | 0 | 2.29 | 1.44 | 0.1865 | 0.2333 | 0.0468 |
| 2v8l | noschrodinger | 1.8 | 0.0089 | 0.98 | 0 | 94.23 | 0 | 0 | 2.3 | 1.4 | 0.1891 | 0.234 | 0.0449 |
| 2v8q | opls | 2.1 | 0.1003 | 2.36 | 1.29 | 94.4 | 8.26 | 9 | 2.13 | 2.06 | 0.2062 | 0.2234 | 0.0172 |
| 2v8q | noschrodinger | 2.1 | 0.0029 | 0.85 | 1.71 | 93.18 | 9.09 | 0 | 2.89 | 2.25 | 0.2079 | 0.223 | 0.0151 |
| 2vbt | opls | 2.7 | 0.0933 | 2.28 | 0 | 97.74 | 2.48 | 0 | 0 | 0.86 | 0.215 | 0.2399 | 0.0249 |
| 2vbt | noschrodinger | 2.7 | 0.003 | 0.75 | 0 | 94.78 | 4.13 | 0 | 0.87 | 1.59 | 0.2105 | 0.2515 | 0.041 |
| 2vbv | opls | 2.4 | 0.0926 | 2.27 | 0.4 | 99.2 | 5.13 | 1 | 0.87 | 1.31 | 0.222 | 0.2515 | 0.0295 |
| 2vbv | noschrodinger | 2.4 | 0.0029 | 0.82 | 1.13 | 94.74 | 11.49 | 0 | 2.62 | 2.22 | 0.2275 | 0.2486 | 0.0211 |
| 2vfk | opls | 1.5 | 0.1018 | 2.36 | 0 | 98.48 | 0.57 | 2 | 1.81 | 0.94 | 0.1583 | 0.1856 | 0.0273 |
| 2vfk | noschrodinger | 1.5 | 0.0107 | 1.14 | 0 | 98.52 | 0.56 | 0 | 2.11 | 0.98 | 0.1599 | 0.1878 | 0.0279 |
| 2vfl | opls | 2.25 | 0.0922 | 2.27 | 0 | 97.97 | 0 | 0 | 1.2 | 0.84 | 0.2057 | 0.2391 | 0.0334 |
| 2vfl | noschrodinger | 2.25 | 0.0033 | 0.79 | 0 | 98.03 | 0 | 0 | 0.9 | 0.77 | 0.2067 | 0.2335 | 0.0268 |
| 2vh9 | opls | 2 | 0.0996 | 2.45 | 0 | 98.11 | 1.29 | 0 | 1.96 | 1.05 | 0.175 | 0.1948 | 0.0198 |
| 2vh9 | noschrodinger | 2 | 0.0029 | 0.94 | 0 | 96.97 | 1.51 | 0 | 3.69 | 1.47 | 0.165 | 0.1841 | 0.0191 |
| 2vi0 | opls | 1.51 | 0.1038 | 2.39 | 0 | 99.26 | 1.23 | 4 | 3.86 | 1.24 | 0.1488 | 0.1871 | 0.0383 |
| 2vi0 | noschrodinger | 1.51 | 0.0046 | 0.94 | 0 | 98.53 | 1.23 | 0 | 2.99 | 1.16 | 0.1529 | 0.1847 | 0.0318 |
| 2vii | opls | 2.85 | 0.0955 | 2.36 | 0 | 98.28 | 1.92 | 1 | 1.01 | 1.01 | 0.181 | 0.2227 | 0.0417 |
| 2vii | noschrodinger | 2.85 | 0.0031 | 0.82 | 0 | 98.35 | 2.39 | 0 | 4.07 | 1.48 | 0.1674 | 0.207 | 0.0396 |
| 2vj2 | opls | 2.5 | 0.0977 | 2.24 | 0.68 | 94.58 | 1.58 | 1 | 1.16 | 1.35 | 0.2028 | 0.2508 | 0.048 |
| 2vj2 | noschrodinger | 2.5 | 0.0035 | 0.82 | 0 | 93.22 | 1.19 | 0 | 2.56 | 1.54 | 0.2046 | 0.2445 | 0.0399 |
| 2vlp | opls | 2 | 0.1007 | 2.39 | 0 | 99.03 | 2.67 | 1 | 0.59 | 1.02 | 0.1777 | 0.208 | 0.0303 |
| 2vlp | noschrodinger | 2 | 0.0028 | 0.8 | 0 | 100 | 3.17 | 0 | 2.07 | 1.36 | 0.1652 | 0.1981 | 0.0329 |
| 2vp2 | opls | 2.5 | 0.0938 | 2.29 | 0.26 | 96.66 | 3.03 | 2 | 0.3 | 1.19 | 0.201 | 0.2358 | 0.0348 |
| 2vp2 | noschrodinger | 2.5 | 0.0028 | 0.81 | 0.51 | 95.14 | 4.66 | 0 | 0.9 | 1.62 | 0.1884 | 0.22 | 0.0316 |
| 2vpp | opls | 2.2 | 0.0976 | 2.28 | 0.28 | 95.79 | 0.58 | 1 | 0.64 | 1 | 0.2151 | 0.2396 | 0.0245 |
| 2vpp | noschrodinger | 2.2 | 0.0028 | 0.8 | 0 | 96.4 | 3.17 | 0 | 1.75 | 1.55 | 0.1966 | 0.2295 | 0.0329 |
| 2vq5 | opls | 2.09 | 0.0942 | 2.25 | 0.34 | 97.97 | 4.38 | 1 | 2.27 | 1.5 | 0.2215 | 0.2395 | 0.018 |
| 2vq5 | noschrodinger | 2.09 | 0.0028 | 0.83 | 0.93 | 97.52 | 5.56 | 0 | 3.4 | 1.8 | 0.2151 | 0.2299 | 0.0148 |
| 2vrb | opls | 2 | 0.0962 | 2.25 | 0 | 98.58 | 2.16 | 1 | 0.9 | 1.03 | 0.2345 | 0.256 | 0.0215 |
| 2vrb | noschrodinger | 2 | 0.0031 | 0.83 | 0 | 99.29 | 6.44 | 0 | 2.03 | 1.59 | 0.2107 | 0.2478 | 0.0371 |
| 2vzq | opls | 1.7 | 0.1091 | 2.28 | 0.4 | 97.98 | 2.55 | 3 | 2.95 | 1.4 | 0.1573 | 0.2018 | 0.0445 |
| 2vzq | noschrodinger | 1.7 | 0.0032 | 0.85 | 0 | 95.97 | 2.55 | 0 | 3.75 | 1.75 | 0.1622 | 0.194 | 0.0318 |
| 2w1q | opls | 1.6 | 0.1074 | 2.27 | 0 | 97.8 | 0.83 | 2 | 3.4 | 1.18 | 0.1476 | 0.1855 | 0.0379 |
| 2w1q | noschrodinger | 1.6 | 0.0039 | 0.91 | 0 | 97.82 | 1.65 | 0 | 2.49 | 1.24 | 0.1528 | 0.1833 | 0.0305 |
| 2w4y | opls | 2.76 | 0.0904 | 2.33 | 1.12 | 94.68 | 3.88 | 1 | 1.9 | 1.77 | 0.4132 | 0.4243 | 0.0111 |
| 2w4y | noschrodinger | 2.76 | 0.0045 | 0.87 | 1.39 | 88.33 | 4.85 | 0 | 12.3 | 2.72 | 0.4073 | 0.4171 | 0.0098 |

| | | | | | | | | | | | | | |
|---|---|---|---|---|---|---|---|---|---|---|---|---|---|
| 2w4z | opls | 3.5 | 0.0897 | 2.39 | 1.12 | 93.28 | 5.83 | 2 | 1.04 | 1.82 | 0.512 | 0.5193 | 0.0073 |
| 2w4z | noschrodinger | 3.5 | 0.0034 | 0.77 | 1.39 | 93.89 | 7.77 | 0 | 6.23 | 2.43 | 0.515 | 0.5244 | 0.0094 |
| 2w52 | opls | 1.56 | 0.1029 | 2.24 | 0 | 98.31 | 0 | 0 | 2.77 | 1.07 | 0.1316 | 0.1579 | 0.0263 |
| 2w52 | noschrodinger | 1.56 | 0.0051 | 1.16 | 0 | 98.31 | 0.41 | 0 | 6.6 | 1.36 | 0.1332 | 0.1542 | 0.021 |
| 2w5p | opls | 1.93 | 0.0999 | 2.33 | 0.49 | 97.04 | 0.29 | 10 | 2.67 | 1.22 | 0.1839 | 0.23 | 0.0461 |
| 2w5p | noschrodinger | 1.93 | 0.0034 | 0.91 | 0.49 | 96.34 | 4.03 | 0 | 4.71 | 1.95 | 0.1908 | 0.2295 | 0.0387 |
| 2wal | opls | 2.4 | 0.0925 | 2.45 | 3.3 | 92.31 | 10.04 | 3 | 1.17 | 2.07 | 0.2681 | 0.2892 | 0.0211 |
| 2wal | noschrodinger | 2.4 | 0.0028 | 0.81 | 3.97 | 92.42 | 12.93 | 0 | 3.05 | 2.41 | 0.2624 | 0.2818 | 0.0194 |
| 2wcm | opls | 1.5 | 0.0981 | 2.22 | 0 | 98.54 | 0 | 0 | 1.72 | 0.93 | 0.19 | 0.229 | 0.039 |
| 2wcm | noschrodinger | 1.5 | 0.0049 | 0.84 | 0 | 98.56 | 0 | 0 | 1.72 | 0.93 | 0.1882 | 0.2203 | 0.0321 |
| 2wcu | opls | 1.9 | 0.0933 | 2.52 | 0 | 97.6 | 5.24 | 6 | 1.68 | 1.55 | 0.1442 | 0.1885 | 0.0443 |
| 2wcu | noschrodinger | 1.9 | 0.0056 | 0.91 | 0 | 97.96 | 3.6 | 0 | 2.31 | 1.44 | 0.155 | 0.1887 | 0.0337 |
| 2we0 | opls | 2.01 | 0.0988 | 2.28 | 0 | 98.73 | 1.4 | 0 | 0.5 | 0.78 | 0.1831 | 0.2191 | 0.036 |
| 2we0 | noschrodinger | 2.01 | 0.0031 | 0.84 | 0 | 97.94 | 1.84 | 0 | 2.26 | 1.22 | 0.1772 | 0.2107 | 0.0335 |
| 2wex | opls | 1.91 | 0.0945 | 2.39 | 0.67 | 96 | 4.41 | 1 | 3.15 | 1.87 | 0.2272 | 0.2582 | 0.031 |
| 2wex | noschrodinger | 1.91 | 0.0076 | 1.02 | 0.65 | 94.77 | 6.47 | 0 | 4.34 | 2.19 | 0.2353 | 0.2609 | 0.0256 |
| 2wjg | opls | 2.2 | 0.095 | 2.3 | 0.3 | 96.39 | 6.74 | 1 | 1.47 | 1.75 | 0.2546 | 0.2719 | 0.0173 |
| 2wjg | noschrodinger | 2.2 | 0.003 | 0.84 | 1.19 | 91.99 | 10.21 | 0 | 2.21 | 2.25 | 0.2383 | 0.2605 | 0.0222 |
| 2wjh | opls | 2.1 | 0.0964 | 2.27 | 0 | 98.46 | 6.81 | 1 | 0.56 | 1.32 | 0.223 | 0.2523 | 0.0293 |
| 2wjh | noschrodinger | 2.1 | 0.0028 | 0.81 | 0.61 | 96.34 | 9.54 | 0 | 2.44 | 2.02 | 0.2034 | 0.2422 | 0.0388 |
| 2wjq | opls | 2 | 0.0921 | 2.06 | 0 | 98.34 | 1.15 | 0 | 0.27 | 0.65 | 0.197 | 0.2186 | 0.0216 |
| 2wjq | noschrodinger | 2 | 0.0026 | 0.74 | 0 | 97.01 | 1.14 | 0 | 1.08 | 1.03 | 0.1891 | 0.2108 | 0.0217 |
| 2wm3 | opls | 1.85 | 0.0989 | 2.56 | 0.75 | 97.75 | 0.41 | 4 | 4.76 | 1.3 | 0.1735 | 0.2151 | 0.0416 |
| 2wm3 | noschrodinger | 1.85 | 0.0135 | 1.29 | 0 | 97.62 | 0.41 | 0 | 6.04 | 1.41 | 0.1758 | 0.2116 | 0.0358 |
| 2wmd | opls | 2 | 0.0992 | 2.25 | 0 | 97.75 | 0.42 | 0 | 0.64 | 0.77 | 0.2385 | 0.2776 | 0.0391 |
| 2wmd | noschrodinger | 2 | 0.0027 | 0.76 | 0 | 96.59 | 1.67 | 0 | 1.94 | 1.35 | 0.2327 | 0.2751 | 0.0424 |
| 2wmm | opls | 2.3 | 0.0956 | 2.31 | 0 | 96.86 | 5.04 | 0 | 0.79 | 1.47 | 0.2101 | 0.2453 | 0.0352 |
| 2wmm | noschrodinger | 2.3 | 0.0039 | 0.77 | 0.31 | 95.31 | 6.79 | 0 | 2.57 | 2 | 0.2015 | 0.2406 | 0.0391 |
| 2wnb | opls | 1.55 | 0.1028 | 2.32 | 0 | 98.45 | 0 | 2 | 3.83 | 1.17 | 0.1501 | 0.1852 | 0.0351 |
| 2wnb | noschrodinger | 1.55 | 0.0058 | 1.13 | 0 | 97.77 | 1.17 | 0 | 5.32 | 1.39 | 0.1563 | 0.1806 | 0.0243 |
| 2wne | opls | 2.12 | 0.098 | 2.3 | 0 | 97.29 | 0.42 | 0 | 0.65 | 0.85 | 0.1687 | 0.209 | 0.0403 |
| 2wne | noschrodinger | 2.12 | 0.0043 | 0.97 | 0 | 97.3 | 1.26 | 0 | 5.81 | 1.53 | 0.163 | 0.1904 | 0.0274 |
| 2wns | opls | 1.9 | 0.0978 | 2.46 | 0 | 97.93 | 0.87 | 3 | 1.74 | 0.95 | 0.1695 | 0.2159 | 0.0464 |
| 2wns | noschrodinger | 1.9 | 0.0035 | 0.8 | 0 | 97.7 | 0.86 | 0 | 2.05 | 1.04 | 0.1816 | 0.212 | 0.0304 |
| 2wqq | opls | 2.25 | 0.0932 | 2.27 | 0 | 96.61 | 0.92 | 0 | 0.97 | 1.01 | 0.2133 | 0.2501 | 0.0368 |
| 2wqq | noschrodinger | 2.25 | 0.0026 | 0.74 | 0 | 96.65 | 4.07 | 0 | 1.94 | 1.64 | 0.1957 | 0.2352 | 0.0395 |
| 2wqw | opls | 2.24 | 0.0934 | 2.24 | 0 | 96.11 | 1.14 | 2 | 0.54 | 0.99 | 0.2072 | 0.2339 | 0.0267 |
| 2wqw | noschrodinger | 2.24 | 0.0027 | 0.81 | 0 | 95.42 | 3.21 | 0 | 1.18 | 1.54 | 0.2071 | 0.2241 | 0.017 |
| 2wsx | opls | 3.41 | 0.0906 | 2.32 | 0.07 | 97.74 | 3.99 | 3 | 0.76 | 1.26 | 0.2574 | 0.2747 | 0.0173 |
| 2wsx | noschrodinger | 3.41 | 0.0035 | 0.81 | 0.74 | 88.73 | 9.84 | 0 | 8.92 | 2.81 | 0.2252 | 0.2569 | 0.0317 |
| 2wtn | opls | 2.1 | 0.0942 | 2.34 | 0 | 96.61 | 1.2 | 3 | 1.26 | 1.13 | 0.1804 | 0.2061 | 0.0257 |
| 2wtn | noschrodinger | 2.1 | 0.0064 | 1.05 | 0.61 | 96.16 | 1.68 | 0 | 2.9 | 1.51 | 0.1706 | 0.2011 | 0.0305 |
| 2wu1 | opls | 2.2 | 0.0961 | 2.3 | 0 | 96.01 | 3.54 | 1 | 0.71 | 1.42 | 0.1691 | 0.1981 | 0.029 |
| 2wu1 | noschrodinger | 2.2 | 0.0031 | 0.83 | 0.38 | 95.64 | 6.39 | 0 | 2 | 1.88 | 0.1577 | 0.188 | 0.0303 |
| 2wvq | opls | 2 | 0.0962 | 2.37 | 0 | 98.51 | 8.76 | 5 | 1.63 | 1.63 | 0.2274 | 0.2589 | 0.0315 |
| 2wvq | noschrodinger | 2 | 0.0029 | 0.84 | 0 | 98.06 | 13.24 | 0 | 1.62 | 1.76 | 0.2301 | 0.2536 | 0.0235 |
| 2wzv | opls | 1.75 | 0.0985 | 2.38 | 0 | 98.97 | 0.89 | 1 | 0.99 | 0.79 | 0.1445 | 0.1668 | 0.0223 |
| 2wzv | noschrodinger | 1.75 | 0.0042 | 0.84 | 0 | 99.09 | 0.85 | 0 | 0.71 | 0.73 | 0.148 | 0.1635 | 0.0155 |
| 2x0j | opls | 2.79 | 0.0915 | 2.33 | 1.03 | 95.88 | 2.16 | 4 | 0.22 | 1.12 | 0.2082 | 0.243 | 0.0348 |

| | | | | | | | | | | | | | |
|---|---|---|---|---|---|---|---|---|---|---|---|---|---|
| 2x0j | noschrodinger | 2.79 | 0.0035 | 0.85 | 1.37 | 92.12 | 4.31 | 0 | 5.65 | 2.27 | 0.2121 | 0.248 | 0.0359 |
| 2x27 | opls | 2.4 | 0.0925 | 2.21 | 0.5 | 97.01 | 0 | 1 | 0.86 | 0.94 | 0.1901 | 0.2488 | 0.0587 |
| 2x27 | noschrodinger | 2.4 | 0.0034 | 0.81 | 0.49 | 97.07 | 0.59 | 0 | 1.46 | 1.05 | 0.1878 | 0.2324 | 0.0446 |
| 2x32 | opls | 1.55 | 0.1045 | 2.35 | 0 | 98.54 | 1.88 | 6 | 4.05 | 1.4 | 0.1573 | 0.1873 | 0.03 |
| 2x32 | noschrodinger | 1.55 | 0.0038 | 0.92 | 0 | 98.55 | 2.48 | 0 | 3.7 | 1.46 | 0.1623 | 0.1836 | 0.0213 |
| 2x61 | opls | 1.95 | 0.0953 | 2.29 | 0 | 98.51 | 0.68 | 1 | 1.43 | 0.88 | 0.156 | 0.1903 | 0.0343 |
| 2x61 | noschrodinger | 1.95 | 0.0093 | 1.1 | 0 | 98.32 | 1.13 | 1 | 3.58 | 1.19 | 0.1614 | 0.1959 | 0.0345 |
| 2x63 | opls | 2 | 0.0983 | 2.24 | 0 | 98.43 | 0.86 | 0 | 0.93 | 0.78 | 0.1909 | 0.2021 | 0.0112 |
| 2x63 | noschrodinger | 2 | 0.0025 | 0.74 | 0 | 98.05 | 2.13 | 0 | 2.1 | 1.23 | 0.1782 | 0.1961 | 0.0179 |
| 2x6j | opls | 3.5 | 0.0908 | 2.42 | 1.21 | 95.06 | 9.03 | 15 | 1.72 | 2 | 0.1967 | 0.2425 | 0.0458 |
| 2x6j | noschrodinger | 3.5 | 0.0035 | 0.87 | 1.48 | 93.61 | 18.57 | 0 | 10.51 | 2.93 | 0.1789 | 0.2273 | 0.0484 |
| 2x6u | opls | 1.9 | 0.0996 | 2.18 | 0 | 100 | 1.88 | 0 | 1.68 | 1.13 | 0.1974 | 0.2331 | 0.0357 |
| 2x6u | noschrodinger | 1.9 | 0.0025 | 0.71 | 0 | 99.42 | 1.23 | 0 | 0.34 | 0.69 | 0.2029 | 0.2313 | 0.0284 |
| 2x9f | opls | 1.75 | 0.099 | 2.23 | 0 | 99.07 | 0.47 | 0 | 1.19 | 0.83 | 0.161 | 0.195 | 0.034 |
| 2x9f | noschrodinger | 1.75 | 0.0084 | 0.98 | 0 | 97.68 | 0.46 | 0 | 0.97 | 0.86 | 0.1591 | 0.196 | 0.0369 |
| 2x9o | opls | 1.55 | 0.1036 | 2.57 | 0 | 99.1 | 0 | 1 | 3.66 | 1.16 | 0.1606 | 0.1943 | 0.0337 |
| 2x9o | noschrodinger | 1.55 | 0.0038 | 0.95 | 0 | 98.69 | 1.43 | 0 | 2.87 | 1.19 | 0.171 | 0.1934 | 0.0224 |
| 2xb1 | opls | 1.9 | 0.1002 | 2.36 | 0 | 98.89 | 3.11 | 1 | 4.35 | 1.59 | 0.1642 | 0.2087 | 0.0445 |
| 2xb1 | noschrodinger | 1.9 | 0.0081 | 0.98 | 0 | 98.92 | 2.42 | 0 | 4.36 | 1.51 | 0.1655 | 0.2093 | 0.0438 |
| 2xb5 | opls | 2.5 | 0.0928 | 2.38 | 0.98 | 96.08 | 9.88 | 1 | 0.6 | 1.72 | 0.1693 | 0.2173 | 0.048 |
| 2xb5 | noschrodinger | 2.5 | 0.003 | 0.87 | 1.46 | 95.12 | 11.05 | 0 | 1.2 | 1.97 | 0.1612 | 0.2127 | 0.0515 |
| 2xga | opls | 2.3 | 0.0936 | 2.31 | 0 | 97.94 | 0.86 | 2 | 0 | 0.51 | 0.2283 | 0.2493 | 0.021 |
| 2xga | noschrodinger | 2.3 | 0.0029 | 0.87 | 0 | 98.38 | 0.84 | 0 | 3.37 | 1.13 | 0.1988 | 0.2387 | 0.0399 |
| 2xgw | opls | 2.1 | 0.1018 | 2.19 | 0 | 99.36 | 2.1 | 1 | 0.75 | 0.98 | 0.1432 | 0.1696 | 0.0264 |
| 2xgw | noschrodinger | 2.1 | 0.003 | 0.75 | 0 | 99.38 | 2.1 | 0 | 2.64 | 1.29 | 0.1382 | 0.1628 | 0.0246 |
| 2xj9 | opls | 2.8 | 0.0923 | 2.41 | 0.78 | 95.9 | 8.69 | 1 | 1.19 | 1.83 | 0.2023 | 0.2559 | 0.0536 |
| 2xj9 | noschrodinger | 2.8 | 0.0034 | 0.85 | 1.15 | 92.31 | 15.78 | 0 | 5.47 | 2.68 | 0.1858 | 0.227 | 0.0412 |
| 2xmy | opls | 1.9 | 0.0994 | 2.42 | 0 | 96.53 | 9.13 | 2 | 2.61 | 2 | 0.1909 | 0.2414 | 0.0505 |
| 2xmy | noschrodinger | 1.9 | 0.0038 | 0.88 | 0 | 96.22 | 12.03 | 0 | 8.83 | 2.55 | 0.1925 | 0.2325 | 0.04 |
| 2xnb | opls | 1.8 | 0.0995 | 2.49 | 0.34 | 95.86 | 12.36 | 2 | 4.46 | 2.34 | 0.2127 | 0.2474 | 0.0347 |
| 2xnb | noschrodinger | 1.8 | 0.0031 | 0.89 | 0.34 | 95.55 | 14.13 | 0 | 8.13 | 2.63 | 0.2257 | 0.2467 | 0.021 |
| 2xoc | opls | 1.89 | 0.1072 | 2.54 | 0 | 98.04 | 1.32 | 2 | 2.34 | 1.11 | 0.1491 | 0.1794 | 0.0303 |
| 2xoc | noschrodinger | 1.89 | 0.008 | 0.98 | 0 | 98.08 | 1.58 | 0 | 1.91 | 1.11 | 0.1554 | 0.1776 | 0.0222 |
| 2xov | opls | 1.65 | 0.0899 | 2.41 | 0 | 98.86 | 0 | 0 | 2.31 | 1.01 | 0.1755 | 0.2001 | 0.0246 |
| 2xov | noschrodinger | 1.65 | 0.0129 | 1.39 | 0 | 97.77 | 0 | 0 | 4.55 | 1.28 | 0.1786 | 0.1996 | 0.021 |
| 2xoy | opls | 2.6 | 0.0927 | 2.34 | 0 | 96.74 | 0 | 1 | 0.6 | 0.9 | 0.206 | 0.2378 | 0.0318 |
| 2xoy | noschrodinger | 2.6 | 0.005 | 0.82 | 0 | 96.41 | 0.8 | 0 | 0.75 | 0.98 | 0.1978 | 0.2236 | 0.0258 |
| 2xoz | opls | 2.37 | 0.0964 | 2.3 | 0 | 96.57 | 0.8 | 0 | 0.74 | 0.96 | 0.2003 | 0.2256 | 0.0253 |
| 2xoz | noschrodinger | 2.37 | 0.0043 | 0.86 | 0 | 97.6 | 2.9 | 0 | 0.89 | 1.21 | 0.1788 | 0.2183 | 0.0395 |
| 2xp3 | opls | 2 | 0.0972 | 2.53 | 0 | 100 | 0 | 0 | 1.27 | 0.85 | 0.2384 | 0.2697 | 0.0313 |
| 2xp3 | noschrodinger | 2 | 0.0027 | 0.75 | 0 | 99.29 | 3.97 | 0 | 2.54 | 1.49 | 0.2211 | 0.2628 | 0.0417 |
| 2xrh | opls | 1.5 | 0.1008 | 2.31 | 1.03 | 97.94 | 2.2 | 1 | 1.75 | 1.21 | 0.1944 | 0.2272 | 0.0328 |
| 2xrh | noschrodinger | 1.5 | 0.0056 | 0.93 | 1.02 | 97.96 | 2.2 | 0 | 5.25 | 1.55 | 0.1882 | 0.2156 | 0.0274 |
| 2xtm | opls | 1.7 | 0.0964 | 2.38 | 0 | 99.38 | 1.2 | 2 | 1.4 | 0.93 | 0.1786 | 0.2024 | 0.0238 |
| 2xtm | noschrodinger | 1.7 | 0.0073 | 1 | 0 | 98.75 | 0.89 | 0 | 0.63 | 0.71 | 0.1828 | 0.201 | 0.0182 |
| 2xto | opls | 2.8 | 0.0895 | 2.31 | 0 | 97.85 | 2.14 | 1 | 1 | 1.08 | 0.2186 | 0.268 | 0.0494 |
| 2xto | noschrodinger | 2.8 | 0.0026 | 0.76 | 0 | 96.8 | 4.76 | 0 | 0.86 | 1.48 | 0.208 | 0.2613 | 0.0533 |
| 2xup | opls | 2.7 | 0.0899 | 2.33 | 0.57 | 95.94 | 1.48 | 4 | 1.11 | 1.23 | 0.2152 | 0.2411 | 0.0259 |
| 2xup | noschrodinger | 2.7 | 0.003 | 0.88 | 0.75 | 94.94 | 3.51 | 0 | 2.99 | 1.85 | 0.2219 | 0.2394 | 0.0175 |

| | | | | | | | | | | | | | |
|---|---|---|---|---|---|---|---|---|---|---|---|---|---|
| 2xw7 | opls | 2 | 0.0943 | 2.36 | 0 | 97.37 | 1.69 | 0 | 1.78 | 1.23 | 0.1965 | 0.2138 | 0.0173 |
| 2xw7 | noschrodinger | 2 | 0.0036 | 0.9 | 0.58 | 96.52 | 3.34 | 0 | 2.84 | 1.7 | 0.1896 | 0.2133 | 0.0237 |
| 2y1b | opls | 2 | 0.0966 | 2.25 | 0 | 97.59 | 1.41 | 0 | 0 | 0.7 | 0.2197 | 0.233 | 0.0133 |
| 2y1b | noschrodinger | 2 | 0.01 | 1.14 | 0 | 96.43 | 5.56 | 0 | 3.87 | 1.98 | 0.1974 | 0.2197 | 0.0223 |
| 2y33 | opls | 2 | 0.0977 | 2.25 | 0.5 | 95.98 | 0.57 | 0 | 0.3 | 0.89 | 0.1943 | 0.2138 | 0.0195 |
| 2y33 | noschrodinger | 2 | 0.0034 | 0.86 | 0 | 97.63 | 1.71 | 0 | 0.9 | 1.03 | 0.1783 | 0.2075 | 0.0292 |
| 2y7p | opls | 1.85 | 0.0988 | 2.4 | 0 | 99.01 | 3.26 | 1 | 1.69 | 1.31 | 0.165 | 0.2023 | 0.0373 |
| 2y7p | noschrodinger | 1.85 | 0.0116 | 1.2 | 0.47 | 98.12 | 3.76 | 0 | 1.41 | 1.31 | 0.1672 | 0.2023 | 0.0351 |
| 2yb1 | opls | 1.9 | 0.1015 | 2.3 | 0 | 98.2 | 0 | 1 | 1.41 | 0.87 | 0.1421 | 0.1812 | 0.0391 |
| 2yb1 | noschrodinger | 1.9 | 0.003 | 0.83 | 0 | 97.86 | 0.47 | 0 | 1.41 | 0.91 | 0.1484 | 0.1797 | 0.0313 |
| 2ybq | opls | 2 | 0.0962 | 2.36 | 0 | 98.28 | 0.54 | 0 | 1.82 | 0.94 | 0.2465 | 0.276 | 0.0295 |
| 2ybq | noschrodinger | 2 | 0.0028 | 0.83 | 0 | 98.31 | 1.6 | 0 | 1.56 | 1.06 | 0.2374 | 0.2749 | 0.0375 |
| 2ycz | opls | 3.65 | 0.0914 | 2.3 | 0.17 | 96.35 | 1 | 0 | 0 | 0.74 | 0.2432 | 0.2873 | 0.0441 |
| 2ycz | noschrodinger | 3.65 | 0.0027 | 0.72 | 0 | 93.25 | 1.59 | 0 | 1.46 | 1.47 | 0.2373 | 0.2759 | 0.0386 |
| 2yd4 | opls | 1.65 | 0.1039 | 2.36 | 0 | 99 | 0.53 | 2 | 2.35 | 1.02 | 0.1597 | 0.1848 | 0.0251 |
| 2yd4 | noschrodinger | 1.65 | 0.0056 | 0.98 | 0 | 98.51 | 0.53 | 0 | 1.76 | 0.93 | 0.1649 | 0.1853 | 0.0204 |
| 2yd8 | opls | 2.05 | 0.0967 | 2.23 | 0 | 98.97 | 0 | 0 | 0 | 0.5 | 0.2306 | 0.2556 | 0.025 |
| 2yd8 | noschrodinger | 2.05 | 0.007 | 1 | 0 | 97.96 | 0 | 0 | 0.99 | 0.8 | 0.201 | 0.2384 | 0.0374 |
| 2yfa | opls | 1.8 | 0.1035 | 2.34 | 0.22 | 99.14 | 2.38 | 3 | 1.08 | 1.1 | 0.1976 | 0.2387 | 0.0411 |
| 2yfa | noschrodinger | 1.8 | 0.008 | 0.99 | 0 | 98.72 | 2.63 | 0 | 3.25 | 1.44 | 0.1947 | 0.233 | 0.0383 |
| 2ygo | opls | 1.84 | 0.0958 | 2.33 | 1.26 | 96.23 | 0.69 | 1 | 1.02 | 1.05 | 0.1812 | 0.2086 | 0.0274 |
| 2ygo | noschrodinger | 1.84 | 0.0117 | 0.93 | 0.62 | 98.12 | 2.08 | 0 | 1.37 | 1.11 | 0.1803 | 0.2078 | 0.0275 |
| 2ygp | opls | 2.22 | 0.0927 | 2.52 | 0.63 | 93.08 | 0.7 | 0 | 1.01 | 1.24 | 0.1929 | 0.2342 | 0.0413 |
| 2ygp | noschrodinger | 2.22 | 0.0112 | 0.89 | 0.62 | 96.88 | 0.69 | 0 | 2.7 | 1.25 | 0.1853 | 0.2251 | 0.0398 |
| 2ygq | opls | 3.95 | 0.0875 | 2.42 | 0 | 95.38 | 0.51 | 0 | 0.78 | 1.07 | 0.2501 | 0.2961 | 0.046 |
| 2ygq | noschrodinger | 3.95 | 0.0029 | 0.77 | 0 | 92.89 | 1.51 | 0 | 1.82 | 1.53 | 0.2498 | 0.2957 | 0.0459 |
| 2yio | opls | 2.43 | 0.0929 | 2.31 | 0.79 | 96.44 | 0.46 | 2 | 0.77 | 0.98 | 0.1677 | 0.2008 | 0.0331 |
| 2yio | noschrodinger | 2.43 | 0.0029 | 0.8 | 0.39 | 96.47 | 1.37 | 0 | 1.29 | 1.19 | 0.1728 | 0.1981 | 0.0253 |
| 2yjm | opls | 1.84 | 0.0965 | 2.24 | 0 | 98.76 | 0.69 | 1 | 0.72 | 0.73 | 0.1926 | 0.226 | 0.0334 |
| 2yjm | noschrodinger | 1.84 | 0.0029 | 0.73 | 0 | 98.77 | 0.69 | 0 | 0.73 | 0.73 | 0.1895 | 0.2187 | 0.0292 |
| 2ykt | opls | 2.11 | 0.0932 | 2.23 | 0 | 98.68 | 1.44 | 1 | 1.02 | 0.92 | 0.2385 | 0.2692 | 0.0307 |
| 2ykt | noschrodinger | 2.11 | 0.0029 | 0.71 | 0 | 98.7 | 5.26 | 0 | 1.03 | 1.35 | 0.2268 | 0.2595 | 0.0327 |
| 2yn4 | opls | 1.74 | 0.102 | 2.59 | 0 | 98.65 | 1.63 | 5 | 10.21 | 1.69 | 0.1491 | 0.1906 | 0.0415 |
| 2yn4 | noschrodinger | 1.74 | 0.008 | 1.04 | 0 | 98.43 | 2.1 | 0 | 7.8 | 1.67 | 0.1528 | 0.1855 | 0.0327 |
| 2ypj | opls | 2.35 | 0.0862 | 2.14 | 0 | 99.08 | 0 | 0 | 0 | 0.5 | 0.236 | 0.281 | 0.045 |
| 2ypj | noschrodinger | 2.35 | 0.0029 | 0.89 | 0 | 95.83 | 3.03 | 0 | 4 | 1.84 | 0.2334 | 0.2775 | 0.0441 |
| 2yut | opls | 2.2 | 0.0967 | 2.26 | 0 | 98.94 | 0 | 0 | 1.28 | 0.85 | 0.1724 | 0.2038 | 0.0314 |
| 2yut | noschrodinger | 2.2 | 0.0027 | 0.79 | 0 | 99.49 | 0 | 0 | 2.57 | 1.04 | 0.1748 | 0.1992 | 0.0244 |
| 2yvo | opls | 1.67 | 0.099 | 2.34 | 0.56 | 98.88 | 0.65 | 0 | 1.02 | 0.8 | 0.1407 | 0.1691 | 0.0284 |
| 2yvo | noschrodinger | 1.67 | 0.0077 | 1.16 | 0.56 | 98.89 | 1.29 | 0 | 1.7 | 1.01 | 0.1449 | 0.1691 | 0.0242 |
| 2ywl | opls | 1.6 | 0.0991 | 2.23 | 0 | 97.7 | 1.11 | 2 | 1.96 | 1.06 | 0.1545 | 0.1856 | 0.0311 |
| 2ywl | noschrodinger | 1.6 | 0.0048 | 1.04 | 0 | 97.47 | 1.09 | 0 | 2.32 | 1.15 | 0.1563 | 0.1832 | 0.0269 |
| 2yww | opls | 2 | 0.0965 | 2.71 | 0 | 98.56 | 0.76 | 1 | 1.44 | 0.88 | 0.1943 | 0.225 | 0.0307 |
| 2yww | noschrodinger | 2 | 0.0039 | 1.03 | 0 | 97.13 | 0.76 | 0 | 2.47 | 1.19 | 0.1916 | 0.2224 | 0.0308 |
| 2yy8 | opls | 2.48 | 0.0936 | 2.41 | 0.59 | 95.25 | 10.03 | 5 | 2.61 | 2.14 | 0.1936 | 0.2379 | 0.0443 |
| 2yy8 | noschrodinger | 2.48 | 0.0028 | 0.79 | 0.58 | 93.27 | 12.24 | 0 | 3.83 | 2.43 | 0.1886 | 0.2353 | 0.0467 |
| 2yzj | opls | 1.66 | 0.0975 | 2.22 | 0 | 95.56 | 2.51 | 0 | 2.33 | 1.62 | 0.1637 | 0.1833 | 0.0196 |
| 2yzj | noschrodinger | 1.66 | 0.0111 | 1.31 | 0 | 95.51 | 2.96 | 0 | 2.33 | 1.68 | 0.1585 | 0.187 | 0.0285 |
| 2yzo | opls | 1.85 | 0.0935 | 2.27 | 0 | 97.73 | 1.85 | 1 | 1.28 | 1.11 | 0.1817 | 0.2128 | 0.0311 |

| | | | | | | | | | | | | | |
|---|---|---|---|---|---|---|---|---|---|---|---|---|---|
| 2yzo | noschrodinger | 1.85 | 0.0106 | 1.19 | 0.68 | 97.74 | 1.31 | 1 | 4 | 1.33 | 0.1798 | 0.2162 | 0.0364 |
| 2z4i | opls | 2.6 | 0.0861 | 2.23 | 0.75 | 95.11 | 3.7 | 2 | 1.6 | 1.68 | 0.2266 | 0.2545 | 0.0279 |
| 2z4i | noschrodinger | 2.6 | 0.0028 | 0.79 | 2.12 | 93.12 | 6.6 | 0 | 3.25 | 2.18 | 0.2173 | 0.2449 | 0.0276 |
| 2z64 | opls | 2.84 | 0.0915 | 2.39 | 0.82 | 91.35 | 5.75 | 4 | 1.96 | 2.05 | 0.2629 | 0.2879 | 0.025 |
| 2z64 | noschrodinger | 2.84 | 0.0035 | 0.88 | 1.92 | 85.62 | 8.82 | 0 | 5.74 | 2.68 | 0.251 | 0.2807 | 0.0297 |
| 2z6d | opls | 2 | 0.0971 | 2.27 | 0 | 100 | 4.57 | 1 | 1.66 | 1.42 | 0.1741 | 0.2181 | 0.044 |
| 2z6d | noschrodinger | 2 | 0.0057 | 0.88 | 0 | 99.55 | 3.02 | 0 | 1.39 | 1.23 | 0.1756 | 0.2142 | 0.0386 |
| 2za0 | opls | 1.7 | 0.1018 | 2.33 | 0.57 | 97.72 | 0.96 | 2 | 2.28 | 1.07 | 0.1693 | 0.2001 | 0.0308 |
| 2za0 | noschrodinger | 1.7 | 0.0059 | 1.02 | 0.28 | 97.73 | 1.27 | 0 | 0.88 | 0.91 | 0.1672 | 0.1983 | 0.0311 |
| 2ze7 | opls | 2.1 | 0.0921 | 2.42 | 0 | 98.65 | 3.68 | 0 | 0.54 | 1.12 | 0.2554 | 0.2774 | 0.022 |
| 2ze7 | noschrodinger | 2.1 | 0.0031 | 0.82 | 0.89 | 95.98 | 9.38 | 0 | 1.63 | 1.93 | 0.2404 | 0.2813 | 0.0409 |
| 2zfk | opls | 3.61 | 0.0879 | 2.68 | 5.42 | 82.08 | 18.9 | 7 | 3.86 | 2.85 | 0.3737 | 0.4201 | 0.0464 |
| 2zfk | noschrodinger | 3.61 | 0.0035 | 1.03 | 13.18 | 65.59 | 24.81 | 0 | 24.5 | 3.82 | 0.3301 | 0.3947 | 0.0646 |
| 2zg1 | opls | 2.7 | 0.0921 | 2.3 | 1.01 | 91.46 | 6.95 | 0 | 2.11 | 2.13 | 0.2556 | 0.2913 | 0.0357 |
| 2zg1 | noschrodinger | 2.7 | 0.0027 | 0.8 | 4.46 | 85.15 | 5.79 | 0 | 5.44 | 2.53 | 0.2727 | 0.3057 | 0.033 |
| 2zgn | opls | 2.5 | 0.0942 | 2.24 | 0 | 96.45 | 4.96 | 2 | 1.89 | 1.71 | 0.198 | 0.2381 | 0.0401 |
| 2zgn | noschrodinger | 2.5 | 0.0028 | 0.79 | 0 | 94.87 | 5.34 | 0 | 1.47 | 1.79 | 0.2091 | 0.2361 | 0.027 |
| 2zgw | opls | 1.5 | 0.1037 | 2.37 | 0 | 97.61 | 0.25 | 0 | 2.9 | 1.16 | 0.1492 | 0.1762 | 0.027 |
| 2zgw | noschrodinger | 1.5 | 0.0061 | 1.06 | 0 | 96.98 | 0.49 | 0 | 3.69 | 1.33 | 0.1543 | 0.1785 | 0.0242 |
| 2ziy | opls | 3.72 | 0.09 | 2.34 | 1.09 | 92.39 | 5.79 | 1 | 0.67 | 1.77 | 0.2783 | 0.3306 | 0.0523 |
| 2ziy | noschrodinger | 3.72 | 0.0032 | 1.06 | 2.45 | 80.71 | 9 | 0 | 6.39 | 2.8 | 0.2832 | 0.3138 | 0.0306 |
| 2zmf | opls | 2.1 | 0.095 | 2.25 | 0.32 | 97.45 | 4.44 | 1 | 1.95 | 1.56 | 0.2146 | 0.2495 | 0.0349 |
| 2zmf | noschrodinger | 2.1 | 0.0029 | 0.8 | 0.29 | 96.52 | 6.21 | 0 | 1.77 | 1.76 | 0.2112 | 0.2426 | 0.0314 |
| 2zqo | opls | 1.8 | 0.1023 | 2.32 | 0 | 97.64 | 2.23 | 0 | 1.17 | 1.17 | 0.1807 | 0.2183 | 0.0376 |
| 2zqo | noschrodinger | 1.8 | 0.0039 | 0.85 | 0 | 97.66 | 2.21 | 0 | 2.34 | 1.35 | 0.1846 | 0.2179 | 0.0333 |
| 2zrg | opls | 3.5 | 0.0866 | 2.38 | 0 | 96.33 | 4.46 | 2 | 0.23 | 1.33 | 0.1831 | 0.2313 | 0.0482 |
| 2zrg | noschrodinger | 3.5 | 0.0033 | 0.87 | 2.31 | 86.47 | 6.44 | 0 | 7.52 | 2.66 | 0.1791 | 0.2303 | 0.0512 |
| 2zsi | opls | 1.8 | 0.0983 | 2.26 | 0 | 98.45 | 0.29 | 0 | 1.11 | 0.82 | 0.1779 | 0.2024 | 0.0245 |
| 2zsi | noschrodinger | 1.8 | 0.0092 | 1.06 | 0 | 97.46 | 0.58 | 0 | 3.18 | 1.22 | 0.1715 | 0.2044 | 0.0329 |
| 2zx2 | opls | 1.8 | 0.1042 | 2.21 | 0 | 98.18 | 0 | 0 | 0.66 | 0.72 | 0.1602 | 0.1865 | 0.0263 |
| 2zx2 | noschrodinger | 1.8 | 0.0059 | 0.95 | 0 | 98.45 | 0 | 0 | 0.82 | 0.76 | 0.1603 | 0.1869 | 0.0266 |
| 2zx3 | opls | 2.11 | 0.0999 | 2.21 | 0 | 97.66 | 0.28 | 0 | 0.65 | 0.79 | 0.1893 | 0.2152 | 0.0259 |
| 2zx3 | noschrodinger | 2.11 | 0.0048 | 0.87 | 0.26 | 97.15 | 0.28 | 0 | 1.94 | 1.11 | 0.1673 | 0.2057 | 0.0384 |
| 2zx4 | opls | 2.69 | 0.09 | 2.23 | 0 | 97.92 | 0.29 | 0 | 0.65 | 0.73 | 0.1996 | 0.23 | 0.0304 |
| 2zx4 | noschrodinger | 2.69 | 0.0029 | 0.83 | 0 | 95.6 | 2.31 | 0 | 2.78 | 1.65 | 0.1827 | 0.2182 | 0.0355 |
| 2zyt | opls | 1.55 | 0.0984 | 2.28 | 0 | 98.61 | 1.06 | 1 | 1.77 | 0.95 | 0.1632 | 0.1936 | 0.0304 |
| 2zyt | noschrodinger | 1.55 | 0.0072 | 1.05 | 0 | 98.96 | 1.76 | 0 | 1.77 | 1.12 | 0.1676 | 0.1949 | 0.0273 |
| 3a1s | opls | 1.5 | 0.1005 | 2.34 | 0.22 | 98.47 | 3.22 | 5 | 4.5 | 1.61 | 0.1659 | 0.1839 | 0.018 |
| 3a1s | noschrodinger | 1.5 | 0.0046 | 1 | 0.2 | 97.62 | 3.37 | 0 | 5.82 | 1.8 | 0.1684 | 0.1793 | 0.0109 |
| 3a2y | opls | 1.95 | 0.0976 | 2.26 | 0.55 | 97.25 | 0.65 | 3 | 1.01 | 0.94 | 0.1837 | 0.2314 | 0.0477 |
| 3a2y | noschrodinger | 1.95 | 0.0025 | 0.74 | 1.09 | 96.72 | 0.64 | 0 | 2.37 | 1.22 | 0.1934 | 0.2242 | 0.0308 |
| 3a3g | opls | 2 | 0.0961 | 2.32 | 0.58 | 96.5 | 5.14 | 3 | 0.71 | 1.5 | 0.205 | 0.2211 | 0.0161 |
| 3a3g | noschrodinger | 2 | 0.0029 | 0.84 | 0.29 | 96.28 | 7.57 | 0 | 3.37 | 2.05 | 0.1857 | 0.2096 | 0.0239 |
| 3a4m | opls | 1.79 | 0.0963 | 2.31 | 0 | 100 | 4.5 | 3 | 2.64 | 1.55 | 0.2102 | 0.2404 | 0.0302 |
| 3a4m | noschrodinger | 1.79 | 0.0035 | 0.83 | 0 | 99.13 | 6.67 | 0 | 5.78 | 1.94 | 0.206 | 0.2378 | 0.0318 |
| 3a6u | opls | 2.55 | 0.0981 | 2.28 | 0 | 97.58 | 0.93 | 0 | 1.93 | 1.05 | 0.1941 | 0.2465 | 0.0524 |
| 3a6u | noschrodinger | 2.55 | 0.0057 | 0.87 | 0 | 96.77 | 1.87 | 0 | 6.29 | 1.75 | 0.1864 | 0.2421 | 0.0557 |
| 3ab8 | opls | 1.7 | 0.1004 | 2.51 | 0 | 99.17 | 0.27 | 1 | 0.88 | 0.77 | 0.1468 | 0.1946 | 0.0478 |
| 3ab8 | noschrodinger | 1.7 | 0.0048 | 1.08 | 0 | 99.42 | 0 | 0 | 1.39 | 0.87 | 0.1551 | 0.1953 | 0.0402 |

| | | | | | | | | | | | | | |
|---|---|---|---|---|---|---|---|---|---|---|---|---|---|
| 3acb | opls | 2.06 | 0.0994 | 2.21 | 0 | 95.62 | 0 | 1 | 0.75 | 1.04 | 0.188 | 0.2166 | 0.0286 |
| 3acb | noschrodinger | 2.06 | 0.0028 | 0.81 | 0 | 96.91 | 0 | 0 | 0.75 | 0.92 | 0.1792 | 0.2119 | 0.0327 |
| 3acg | opls | 1.5 | 0.1109 | 2.31 | 0 | 97.38 | 1.24 | 1 | 2.96 | 1.28 | 0.1392 | 0.1707 | 0.0315 |
| 3acg | noschrodinger | 1.5 | 0.0083 | 1.13 | 0 | 97.38 | 1.24 | 0 | 1.98 | 1.16 | 0.1394 | 0.1665 | 0.0271 |
| 3acw | opls | 1.63 | 0.1014 | 2.25 | 0 | 99.29 | 0.78 | 1 | 0.63 | 0.71 | 0.1901 | 0.2155 | 0.0254 |
| 3acw | noschrodinger | 1.63 | 0.0039 | 0.97 | 0 | 99.29 | 1.17 | 0 | 1.05 | 0.86 | 0.1963 | 0.2186 | 0.0223 |
| 3afk | opls | 1.9 | 0.0999 | 2.21 | 0 | 96.93 | 2.17 | 0 | 2.32 | 1.45 | 0.16 | 0.2049 | 0.0449 |
| 3afk | noschrodinger | 1.9 | 0.0066 | 1.07 | 0 | 96.66 | 1.42 | 0 | 5.83 | 1.65 | 0.1634 | 0.2089 | 0.0455 |
| 3agc | opls | 2 | 0.0971 | 2.33 | 0.6 | 97.22 | 4.1 | 5 | 1.51 | 1.5 | 0.1906 | 0.2272 | 0.0366 |
| 3agc | noschrodinger | 2 | 0.0032 | 0.86 | 0.76 | 96.94 | 5.28 | 0 | 1.63 | 1.64 | 0.1934 | 0.2183 | 0.0249 |
| 3ah2 | opls | 1.7 | 0.1052 | 2.35 | 0 | 97.33 | 0.91 | 4 | 3.3 | 1.25 | 0.1542 | 0.1802 | 0.026 |
| 3ah2 | noschrodinger | 1.7 | 0.0061 | 0.98 | 0 | 97.51 | 0.18 | 0 | 1.7 | 1.02 | 0.1557 | 0.1792 | 0.0235 |
| 3ai4 | opls | 1.6 | 0.1025 | 2.31 | 0 | 99.19 | 1.34 | 0 | 3.67 | 1.25 | 0.1747 | 0.2194 | 0.0447 |
| 3ai4 | noschrodinger | 1.6 | 0.0029 | 0.84 | 0 | 98.8 | 3.1 | 0 | 1.96 | 1.34 | 0.1805 | 0.2145 | 0.034 |
| 3ai9 | opls | 1.55 | 0.0989 | 2.39 | 0 | 97.4 | 1.63 | 2 | 1.73 | 1.21 | 0.1912 | 0.2279 | 0.0367 |
| 3ai9 | noschrodinger | 1.55 | 0.0111 | 1.1 | 0.48 | 97.13 | 2.09 | 0 | 2.89 | 1.48 | 0.1941 | 0.225 | 0.0309 |
| 3ajh | opls | 2.25 | 0.0933 | 2.38 | 0.84 | 95.37 | 3.12 | 4 | 1.03 | 1.5 | 0.2653 | 0.309 | 0.0437 |
| 3ajh | noschrodinger | 2.25 | 0.003 | 0.92 | 2.31 | 92.24 | 7.64 | 0 | 7.72 | 2.57 | 0.24 | 0.29 | 0.05 |
| 3ajm | opls | 2.3 | 0.094 | 2.31 | 0 | 98.17 | 4.55 | 2 | 0.77 | 1.24 | 0.2327 | 0.257 | 0.0243 |
| 3ajm | noschrodinger | 2.3 | 0.0028 | 0.71 | 0 | 96.88 | 5.63 | 0 | 2.92 | 1.84 | 0.2208 | 0.2523 | 0.0315 |
| 3ake | opls | 1.5 | 0.0987 | 2.56 | 0 | 98 | 1.91 | 0 | 1.56 | 1.11 | 0.1799 | 0.2007 | 0.0208 |
| 3ake | noschrodinger | 1.5 | 0.0102 | 1.17 | 0 | 99.01 | 1.89 | 0 | 2.81 | 1.28 | 0.1836 | 0.1991 | 0.0155 |
| 3amp | opls | 1.78 | 0.1004 | 2.19 | 0 | 96.44 | 0.45 | 0 | 1.67 | 1.15 | 0.1526 | 0.188 | 0.0354 |
| 3amp | noschrodinger | 1.78 | 0.007 | 0.97 | 0 | 96.44 | 0.89 | 0 | 1.55 | 1.13 | 0.1563 | 0.1866 | 0.0303 |
| 3aot | opls | 2.2 | 0.0905 | 2.47 | 1 | 97.5 | 2.82 | 0 | 2.08 | 1.42 | 0.2021 | 0.2295 | 0.0274 |
| 3aot | noschrodinger | 2.2 | 0.0027 | 0.77 | 1.49 | 96.02 | 3.37 | 0 | 1.49 | 1.56 | 0.215 | 0.2354 | 0.0204 |
| 3aqt | opls | 2.5 | 0.0947 | 2.28 | 0 | 98.83 | 2.05 | 2 | 0.82 | 0.99 | 0.2132 | 0.2357 | 0.0225 |
| 3aqt | noschrodinger | 2.5 | 0.0027 | 0.72 | 0 | 96.34 | 2.92 | 0 | 2.48 | 1.63 | 0.2006 | 0.2284 | 0.0278 |
| 3aqx | opls | 2.05 | 0.0956 | 2.13 | 0 | 98.46 | 0.61 | 0 | 0.54 | 0.68 | 0.1873 | 0.2204 | 0.0331 |
| 3aqx | noschrodinger | 2.05 | 0.0026 | 0.85 | 0 | 100 | 1.21 | 0 | 4.56 | 1.29 | 0.1794 | 0.2106 | 0.0312 |
| 3asz | opls | 2.25 | 0.0967 | 2.27 | 0.25 | 98.52 | 1.18 | 1 | 0.3 | 0.67 | 0.2008 | 0.2263 | 0.0255 |
| 3asz | noschrodinger | 2.25 | 0.003 | 0.8 | 0.25 | 98.28 | 1.76 | 0 | 1.34 | 1.05 | 0.1811 | 0.2126 | 0.0315 |
| 3b18 | opls | 2.75 | 0.0925 | 2.42 | 1.27 | 93.04 | 5.76 | 1 | 1.49 | 1.91 | 0.2376 | 0.2801 | 0.0425 |
| 3b18 | noschrodinger | 2.75 | 0.003 | 0.86 | 2.45 | 88.96 | 12.14 | 0 | 7.11 | 2.79 | 0.2219 | 0.2595 | 0.0376 |
| 3b6c | opls | 2.3 | 0.0945 | 2.43 | 0.26 | 97.12 | 5.31 | 0 | 1.43 | 1.59 | 0.2485 | 0.2944 | 0.0459 |
| 3b6c | noschrodinger | 2.3 | 0.0031 | 0.79 | 1.01 | 94.46 | 10.87 | 0 | 6.55 | 2.53 | 0.2493 | 0.2963 | 0.047 |
| 3b6j | opls | 2.05 | 0.0986 | 2.37 | 0 | 99.74 | 1.94 | 1 | 2.33 | 1.23 | 0.1751 | 0.1954 | 0.0203 |
| 3b6j | noschrodinger | 2.05 | 0.0028 | 0.81 | 0 | 99.74 | 2.58 | 0 | 2.48 | 1.34 | 0.1678 | 0.1889 | 0.0211 |
| 3b89 | opls | 2.6 | 0.0919 | 2.37 | 0.45 | 95.93 | 1.6 | 1 | 1.07 | 1.25 | 0.213 | 0.2549 | 0.0419 |
| 3b89 | noschrodinger | 2.6 | 0.0028 | 0.82 | 0 | 94.02 | 3.14 | 0 | 2.95 | 1.86 | 0.2043 | 0.2602 | 0.0559 |
| 3bb4 | opls | 2.85 | 0.0918 | 2.4 | 0.85 | 96.58 | 6.28 | 5 | 1.29 | 1.68 | 0.1974 | 0.2492 | 0.0518 |
| 3bb4 | noschrodinger | 2.85 | 0.0033 | 1.03 | 1.69 | 92.37 | 13.88 | 0 | 3.37 | 2.47 | 0.201 | 0.2383 | 0.0373 |
| 3beg | opls | 2.9 | 0.0887 | 2.42 | 0.55 | 92.58 | 7.61 | 8 | 1.17 | 1.96 | 0.2797 | 0.3243 | 0.0446 |
| 3beg | noschrodinger | 2.9 | 0.0034 | 0.87 | 3.05 | 84.04 | 16.25 | 0 | 5.91 | 2.92 | 0.2806 | 0.3046 | 0.024 |
| 3bem | opls | 1.65 | 0.0962 | 2.42 | 0 | 99.71 | 0.3 | 2 | 3.15 | 1.11 | 0.1391 | 0.1767 | 0.0376 |
| 3bem | noschrodinger | 1.65 | 0.014 | 1.36 | 0 | 99.27 | 1.38 | 0 | 3.16 | 1.21 | 0.1416 | 0.1796 | 0.038 |
| 3bf8 | opls | 1.68 | 0.1073 | 2.35 | 0 | 97.21 | 2.15 | 6 | 3 | 1.49 | 0.1908 | 0.2305 | 0.0397 |
| 3bf8 | noschrodinger | 1.68 | 0.0039 | 0.83 | 0 | 97.43 | 1.9 | 0 | 2.25 | 1.33 | 0.1958 | 0.2206 | 0.0248 |
| 3bfv | opls | 1.8 | 0.1002 | 2.33 | 0.21 | 98.53 | 1.18 | 4 | 1.72 | 0.98 | 0.1451 | 0.1877 | 0.0426 |

| | | | | | | | | | | | |
|---|---|---|---|---|---|---|---|---|---|---|---|
| 3bfv | noschrodinger | 1.8 | 0.0037 | 0.94 | 0.42 | 98.33 | 1.41 | 0 | 1.98 | 1.08 | 0.1511 | 0.1842 | 0.0331 |
| 3bgd | opls | 2 | 0.0976 | 2.28 | 0 | 97.82 | 2 | 1 | 1.7 | 1.19 | 0.2295 | 0.2488 | 0.0193 |
| 3bgd | noschrodinger | 2 | 0.0026 | 0.79 | 0.22 | 97.17 | 3.23 | 0 | 2.36 | 1.55 | 0.2092 | 0.2423 | 0.0331 |
| 3bm1 | opls | 2 | 0.1068 | 2.3 | 0.29 | 97.65 | 3.64 | 1 | 1.45 | 1.38 | 0.2062 | 0.2475 | 0.0413 |
| 3bm1 | noschrodinger | 2 | 0.0029 | 0.8 | 0.58 | 96.24 | 3.96 | 0 | 4.53 | 1.94 | 0.1765 | 0.2285 | 0.052 |
| 3bnk | opls | 2.05 | 0.0936 | 2.25 | 0 | 97 | 4.15 | 0 | 2.72 | 1.7 | 0.2009 | 0.2425 | 0.0416 |
| 3bnk | noschrodinger | 2.05 | 0.0031 | 0.85 | 0 | 97.29 | 4.44 | 0 | 4.43 | 1.85 | 0.1917 | 0.2265 | 0.0348 |
| 3bpb | opls | 2.81 | 0.092 | 2.31 | 0 | 95.95 | 3.55 | 2 | 0.88 | 1.46 | 0.2285 | 0.2797 | 0.0512 |
| 3bpb | noschrodinger | 2.81 | 0.0031 | 0.8 | 0.2 | 91.87 | 4.24 | 0 | 2.26 | 1.97 | 0.214 | 0.2721 | 0.0581 |
| 3bpl | opls | 2.93 | 0.0926 | 2.26 | 0.2 | 95.47 | 3.14 | 1 | 1.16 | 1.52 | 0.2308 | 0.271 | 0.0402 |
| 3bpl | noschrodinger | 2.93 | 0.0028 | 0.76 | 0.39 | 92.64 | 4.36 | 0 | 3.14 | 2.05 | 0.2136 | 0.2562 | 0.0426 |
| 3bpx | opls | 1.63 | 0.1001 | 2.42 | 0 | 99.29 | 1.82 | 1 | 4.77 | 1.44 | 0.2415 | 0.2965 | 0.055 |
| 3bpx | noschrodinger | 1.63 | 0.0043 | 0.78 | 0 | 99.29 | 1.09 | 0 | 6.47 | 1.39 | 0.2376 | 0.2862 | 0.0486 |
| 3br3 | opls | 2.8 | 0.0887 | 2.15 | 0 | 97.37 | 1.64 | 1 | 0 | 0.79 | 0.2427 | 0.2764 | 0.0337 |
| 3br3 | noschrodinger | 2.8 | 0.007 | 0.84 | 0.83 | 94.75 | 3.25 | 0 | 1.7 | 1.67 | 0.2355 | 0.277 | 0.0415 |
| 3brq | opls | 2 | 0.097 | 2.33 | 0.41 | 95.85 | 2.55 | 6 | 1.41 | 1.47 | 0.1972 | 0.2326 | 0.0354 |
| 3brq | noschrodinger | 2 | 0.003 | 0.85 | 0.56 | 96.3 | 5.96 | 0 | 2.47 | 1.87 | 0.1846 | 0.2238 | 0.0392 |
| 3bsf | opls | 2.91 | 0.0913 | 2.3 | 1.03 | 94.21 | 4.31 | 0 | 0.53 | 1.55 | 0.2377 | 0.2724 | 0.0347 |
| 3bsf | noschrodinger | 2.91 | 0.0033 | 0.82 | 1.24 | 89.46 | 5.5 | 0 | 4.65 | 2.36 | 0.2482 | 0.2681 | 0.0199 |
| 3bwu | opls | 1.76 | 0.1051 | 2.56 | 0 | 99.29 | 1.54 | 5 | 6.81 | 1.52 | 0.1581 | 0.1927 | 0.0346 |
| 3bwu | noschrodinger | 1.76 | 0.0072 | 1 | 0 | 99.32 | 1.02 | 0 | 4.18 | 1.21 | 0.1611 | 0.1937 | 0.0326 |
| 3by9 | opls | 1.7 | 0.1044 | 2.25 | 0 | 98.2 | 1.06 | 2 | 3 | 1.11 | 0.1453 | 0.181 | 0.0357 |
| 3by9 | noschrodinger | 1.7 | 0.0063 | 0.94 | 0 | 98.44 | 1.05 | 0 | 1.96 | 0.98 | 0.1474 | 0.1771 | 0.0297 |
| 3c03 | opls | 1.9 | 0.094 | 2.44 | 0 | 97.3 | 4.07 | 5 | 0.62 | 1.3 | 0.1888 | 0.2323 | 0.0435 |
| 3c03 | noschrodinger | 1.9 | 0.0064 | 1.05 | 0 | 96.81 | 4.02 | 0 | 4.34 | 1.87 | 0.1897 | 0.2295 | 0.0398 |
| 3c1q | opls | 1.7 | 0.104 | 2.33 | 0 | 100 | 0 | 1 | 3.19 | 1.11 | 0.1521 | 0.1934 | 0.0413 |
| 3c1q | noschrodinger | 1.7 | 0.0051 | 0.96 | 0 | 100 | 0.49 | 0 | 2.67 | 1.05 | 0.1552 | 0.1914 | 0.0362 |
| 3c37 | opls | 1.7 | 0.0962 | 2.21 | 0 | 99.24 | 1.92 | 1 | 2.37 | 1.23 | 0.1997 | 0.2307 | 0.031 |
| 3c37 | noschrodinger | 1.7 | 0.0051 | 0.85 | 0 | 99.08 | 2.64 | 0 | 2.93 | 1.4 | 0.1954 | 0.2254 | 0.03 |
| 3c51 | opls | 3.55 | 0.0907 | 2.31 | 0.66 | 94.32 | 1.78 | 4 | 0.98 | 1.37 | 0.2546 | 0.2947 | 0.0401 |
| 3c51 | noschrodinger | 3.55 | 0.0027 | 0.77 | 0.76 | 89.63 | 5.91 | 0 | 1.5 | 2.04 | 0.2453 | 0.2906 | 0.0453 |
| 3c5x | opls | 2.21 | 0.0912 | 2.33 | 0.22 | 96.05 | 2.45 | 2 | 1.33 | 1.43 | 0.2411 | 0.2632 | 0.0221 |
| 3c5x | noschrodinger | 2.21 | 0.0188 | 1.06 | 0.43 | 95.46 | 4.35 | 0 | 5.85 | 2.12 | 0.227 | 0.2535 | 0.0265 |
| 3c6e | opls | 2.59 | 0.09 | 2.44 | 0.44 | 94.47 | 5.88 | 8 | 0.8 | 1.71 | 0.249 | 0.2805 | 0.0315 |
| 3c6e | noschrodinger | 2.59 | 0.0035 | 0.95 | 0.43 | 91.54 | 6.28 | 0 | 6.67 | 2.48 | 0.2613 | 0.2817 | 0.0204 |
| 3ccf | opls | 1.9 | 0.099 | 2.2 | 0 | 98.1 | 1.34 | 0 | 0.67 | 0.81 | 0.1823 | 0.2128 | 0.0305 |
| 3ccf | noschrodinger | 1.9 | 0.0032 | 0.83 | 0 | 97.48 | 1.83 | 0 | 1.61 | 1.21 | 0.1839 | 0.2094 | 0.0255 |
| 3cgy | opls | 2.6 | 0.0888 | 2.45 | 0 | 96.1 | 1.91 | 1 | 0.34 | 1.1 | 0.2241 | 0.2628 | 0.0387 |
| 3cgy | noschrodinger | 2.6 | 0.0067 | 1.26 | 1.37 | 93.72 | 5.61 | 0 | 6.44 | 2.34 | 0.2152 | 0.2617 | 0.0465 |
| 3cj8 | opls | 1.95 | 0.0991 | 2.38 | 0 | 98.36 | 2.87 | 2 | 1.92 | 1.3 | 0.1962 | 0.2399 | 0.0437 |
| 3cj8 | noschrodinger | 1.95 | 0.0035 | 0.87 | 0 | 98.15 | 3.95 | 0 | 2.02 | 1.42 | 0.2028 | 0.2337 | 0.0309 |
| 3cjd | opls | 1.79 | 0.0966 | 2.36 | 0 | 99.68 | 1.44 | 4 | 1.03 | 0.92 | 0.2032 | 0.2344 | 0.0312 |
| 3cjd | noschrodinger | 1.79 | 0.0055 | 0.93 | 0 | 100 | 1.04 | 0 | 1.56 | 0.91 | 0.2012 | 0.2327 | 0.0315 |
| 3ckl | opls | 2 | 0.1016 | 2.2 | 0 | 97.6 | 0.75 | 0 | 1.43 | 0.96 | 0.1844 | 0.2158 | 0.0314 |
| 3ckl | noschrodinger | 2 | 0.0028 | 0.8 | 0 | 97.92 | 1.31 | 0 | 1.95 | 1.07 | 0.1736 | 0.207 | 0.0334 |
| 3clp | opls | 2 | 0.0968 | 2.26 | 0 | 99.18 | 1.95 | 0 | 1.25 | 1.07 | 0.1841 | 0.2179 | 0.0338 |
| 3clp | noschrodinger | 2 | 0.0081 | 1.01 | 0 | 100 | 2.93 | 0 | 0.75 | 1.09 | 0.1767 | 0.2208 | 0.0441 |
| 3cnm | opls | 1.65 | 0.103 | 2.5 | 0 | 98.39 | 1.39 | 3 | 4.47 | 1.33 | 0.1661 | 0.1987 | 0.0326 |
| 3cnm | noschrodinger | 1.65 | 0.0093 | 1.1 | 0 | 98.71 | 2.78 | 0 | 4.09 | 1.53 | 0.1692 | 0.1984 | 0.0292 |

| | | | | | | | | | | | | | |
|---|---|---|---|---|---|---|---|---|---|---|---|---|---|
| 3cql | opls | 1.5 | 0.099 | 2.31 | 0 | 97.71 | 0.52 | 2 | 2.12 | 1.05 | 0.1377 | 0.1605 | 0.0228 |
| 3cql | noschrodinger | 1.5 | 0.0051 | 0.98 | 0 | 98.13 | 0.52 | 0 | 1.72 | 0.93 | 0.1423 | 0.158 | 0.0157 |
| 3cqo | opls | 2.32 | 0.0961 | 2.23 | 0.12 | 94.46 | 1.76 | 1 | 0.81 | 1.32 | 0.187 | 0.2165 | 0.0295 |
| 3cqo | noschrodinger | 2.32 | 0.0037 | 0.86 | 0.34 | 93.81 | 2.55 | 0 | 3.4 | 1.85 | 0.1703 | 0.2108 | 0.0405 |
| 3crc | opls | 2.97 | 0.0905 | 2.61 | 2.58 | 93.44 | 15.78 | 14 | 2.63 | 2.39 | 0.2518 | 0.2981 | 0.0463 |
| 3crc | noschrodinger | 2.97 | 0.0051 | 1.33 | 6 | 83.14 | 23.95 | 1 | 19.46 | 3.52 | 0.2368 | 0.2864 | 0.0496 |
| 3ctr | opls | 2.1 | 0.099 | 2.29 | 0 | 100 | 1.52 | 1 | 0 | 0.64 | 0.2153 | 0.229 | 0.0137 |
| 3ctr | noschrodinger | 2.1 | 0.0028 | 0.76 | 0 | 98.63 | 1.52 | 0 | 3.28 | 1.26 | 0.1939 | 0.2255 | 0.0316 |
| 3cu2 | opls | 1.91 | 0.0991 | 2.23 | 0 | 98.02 | 0.25 | 1 | 2.09 | 0.98 | 0.1606 | 0.1966 | 0.036 |
| 3cu2 | noschrodinger | 1.91 | 0.0027 | 0.8 | 0.21 | 96.36 | 0.49 | 0 | 1.44 | 1.12 | 0.1654 | 0.1919 | 0.0265 |
| 3cu3 | opls | 2 | 0.0993 | 2.24 | 0 | 99.3 | 0.76 | 1 | 1.53 | 0.9 | 0.1723 | 0.2087 | 0.0364 |
| 3cu3 | noschrodinger | 2 | 0.0083 | 0.96 | 0 | 99.38 | 0.74 | 0 | 1.54 | 0.9 | 0.1557 | 0.203 | 0.0473 |
| 3cwq | opls | 2.47 | 0.0912 | 2.39 | 0 | 95.34 | 9.01 | 2 | 1.27 | 1.9 | 0.2522 | 0.2659 | 0.0137 |
| 3cwq | noschrodinger | 2.47 | 0.0035 | 0.89 | 0.25 | 93.95 | 15.81 | 0 | 3.64 | 2.47 | 0.226 | 0.2542 | 0.0282 |
| 3cx6 | opls | 2.5 | 0.0941 | 2.38 | 0.21 | 97.95 | 4.68 | 1 | 1.08 | 1.33 | 0.2235 | 0.264 | 0.0405 |
| 3cx6 | noschrodinger | 2.5 | 0.0027 | 0.77 | 0.61 | 94.93 | 6.59 | 0 | 1.93 | 1.93 | 0.2086 | 0.2591 | 0.0505 |
| 3cx8 | opls | 2 | 0.0976 | 2.3 | 0.2 | 97.6 | 2.62 | 3 | 0.95 | 1.19 | 0.2113 | 0.2325 | 0.0212 |
| 3cx8 | noschrodinger | 2 | 0.0026 | 0.79 | 0.79 | 96.83 | 3.04 | 0 | 2.27 | 1.57 | 0.2031 | 0.2267 | 0.0236 |
| 3cz1 | opls | 1.5 | 0.1016 | 2.3 | 0 | 99.56 | 2.2 | 3 | 4.92 | 1.52 | 0.156 | 0.191 | 0.035 |
| 3cz1 | noschrodinger | 1.5 | 0.0053 | 0.96 | 0 | 99.12 | 2.2 | 0 | 5.44 | 1.55 | 0.1584 | 0.1871 | 0.0287 |
| 3d1l | opls | 2.19 | 0.0946 | 2.25 | 0 | 97.36 | 1.85 | 0 | 1.19 | 1.16 | 0.2194 | 0.24 | 0.0206 |
| 3d1l | noschrodinger | 2.19 | 0.0028 | 0.78 | 0.2 | 96.67 | 4.46 | 0 | 2.02 | 1.68 | 0.203 | 0.2323 | 0.0293 |
| 3d36 | opls | 2.03 | 0.099 | 2.27 | 0.22 | 98.27 | 1.24 | 1 | 0.38 | 0.71 | 0.2224 | 0.2461 | 0.0237 |
| 3d36 | noschrodinger | 2.03 | 0.0027 | 0.8 | 0.21 | 98.72 | 2.68 | 0 | 1.4 | 1.2 | 0.2128 | 0.2433 | 0.0305 |
| 3d40 | opls | 1.54 | 0.1012 | 2.52 | 0 | 98.73 | 1.47 | 3 | 5.08 | 1.4 | 0.152 | 0.1731 | 0.0211 |
| 3d40 | noschrodinger | 1.54 | 0.0111 | 1.34 | 0 | 98.36 | 1.94 | 0 | 4.85 | 1.47 | 0.1527 | 0.1725 | 0.0198 |
| 3d41 | opls | 2.2 | 0.0928 | 2.37 | 0 | 96.84 | 2.86 | 2 | 0.73 | 1.27 | 0.2193 | 0.2531 | 0.0338 |
| 3d41 | noschrodinger | 2.2 | 0.0032 | 0.89 | 0.38 | 96.21 | 3.32 | 0 | 2.2 | 1.65 | 0.2056 | 0.2417 | 0.0361 |
| 3d72 | opls | 1.52 | 0.1052 | 2.42 | 0.68 | 97.26 | 2.29 | 1 | 2.85 | 1.49 | 0.1962 | 0.2291 | 0.0329 |
| 3d72 | noschrodinger | 1.52 | 0.0123 | 1.29 | 0.34 | 96.94 | 3.79 | 0 | 3.47 | 1.76 | 0.1977 | 0.2367 | 0.039 |
| 3daw | opls | 2.55 | 0.0933 | 2.32 | 0.2 | 96.42 | 4.52 | 3 | 0.86 | 1.5 | 0.222 | 0.25 | 0.028 |
| 3daw | noschrodinger | 2.55 | 0.003 | 0.85 | 0.39 | 94.29 | 8.58 | 0 | 2.96 | 2.18 | 0.208 | 0.2414 | 0.0334 |
| 3dba | opls | 2.57 | 0.0925 | 2.33 | 0.6 | 93.45 | 4.55 | 2 | 1.59 | 1.83 | 0.2897 | 0.3107 | 0.021 |
| 3dba | noschrodinger | 2.57 | 0.0029 | 0.81 | 1.48 | 90.53 | 6.13 | 0 | 3.88 | 2.31 | 0.2736 | 0.3077 | 0.0341 |
| 3ddc | opls | 1.8 | 0.095 | 2.33 | 0 | 97.92 | 3.04 | 3 | 2.26 | 1.39 | 0.173 | 0.2042 | 0.0312 |
| 3ddc | noschrodinger | 1.8 | 0.0036 | 0.98 | 0 | 96.93 | 3.37 | 0 | 2.06 | 1.56 | 0.1771 | 0.2006 | 0.0235 |
| 3ddj | opls | 1.8 | 0.1 | 2.28 | 0 | 98.7 | 0 | 0 | 2.14 | 0.99 | 0.1586 | 0.2046 | 0.046 |
| 3ddj | noschrodinger | 1.8 | 0.0034 | 0.83 | 0 | 98.91 | 0 | 0 | 0.22 | 0.58 | 0.1634 | 0.2012 | 0.0378 |
| 3ddy | opls | 2.5 | 0.1033 | 2.32 | 0 | 95.83 | 5.52 | 1 | 1.86 | 1.8 | 0.2034 | 0.2299 | 0.0265 |
| 3ddy | noschrodinger | 2.5 | 0.0033 | 0.84 | 0 | 95.91 | 6.12 | 0 | 4.48 | 2.1 | 0.1841 | 0.2256 | 0.0415 |
| 3deu | opls | 2.3 | 0.0906 | 2.32 | 0 | 99.6 | 2.15 | 3 | 0 | 0.75 | 0.2538 | 0.2709 | 0.0171 |
| 3deu | noschrodinger | 2.3 | 0.003 | 0.75 | 0.39 | 98.03 | 6.75 | 0 | 0.9 | 1.4 | 0.2314 | 0.2665 | 0.0351 |
| 3dm8 | opls | 1.8 | 0.1032 | 2.34 | 0 | 99.2 | 3.51 | 0 | 1.35 | 1.28 | 0.1784 | 0.2018 | 0.0234 |
| 3dm8 | noschrodinger | 1.8 | 0.004 | 0.88 | 0 | 100 | 4.31 | 0 | 1.36 | 1.35 | 0.1832 | 0.2003 | 0.0171 |
| 3doe | opls | 2.25 | 0.0945 | 2.36 | 0 | 97.96 | 7.09 | 5 | 1.81 | 1.6 | 0.2092 | 0.2437 | 0.0345 |
| 3doe | noschrodinger | 2.25 | 0.0029 | 0.84 | 0.34 | 94.95 | 11.85 | 0 | 3.83 | 2.34 | 0.1953 | 0.2349 | 0.0396 |
| 3dof | opls | 3.3 | 0.0909 | 2.42 | 1.65 | 92.74 | 5.84 | 3 | 1.78 | 1.98 | 0.2679 | 0.3139 | 0.046 |
| 3dof | noschrodinger | 3.3 | 0.0034 | 0.85 | 1.31 | 86.56 | 9.78 | 0 | 4.76 | 2.63 | 0.2512 | 0.2907 | 0.0395 |
| 3dt6 | opls | 2.1 | 0.0932 | 2.25 | 0 | 97.04 | 1.81 | 0 | 0 | 0.86 | 0.1901 | 0.2176 | 0.0275 |

| | | | | | | | | | | | | | |
|---|---|---|---|---|---|---|---|---|---|---|---|---|---|
| 3dt6 | noschrodinger | 2.1 | 0.0027 | 0.79 | 0 | 98.1 | 1.8 | 0 | 1.9 | 1.15 | 0.1914 | 0.2209 | 0.0295 |
| 3dtt | opls | 1.7 | 0.1026 | 2.25 | 0 | 97.87 | 0.32 | 1 | 0.9 | 0.8 | 0.1541 | 0.1904 | 0.0363 |
| 3dtt | noschrodinger | 1.7 | 0.008 | 1.05 | 0 | 98.16 | 0.31 | 0 | 0.3 | 0.61 | 0.151 | 0.1881 | 0.0371 |
| 3dx5 | opls | 2.11 | 0.0973 | 2.25 | 0 | 99.21 | 2.65 | 2 | 3.46 | 1.46 | 0.1718 | 0.1852 | 0.0134 |
| 3dx5 | noschrodinger | 2.11 | 0.0037 | 0.93 | 0 | 98.9 | 2.93 | 0 | 3.85 | 1.53 | 0.1551 | 0.1762 | 0.0211 |
| 3dxo | opls | 2.7 | 0.0906 | 2.21 | 0 | 97.64 | 0.56 | 0 | 0.54 | 0.76 | 0.2294 | 0.2563 | 0.0269 |
| 3dxo | noschrodinger | 2.7 | 0.0027 | 0.75 | 0 | 96.97 | 2.15 | 0 | 2.17 | 1.42 | 0.2178 | 0.2616 | 0.0438 |
| 3e3u | opls | 1.56 | 0.1031 | 2.31 | 0.52 | 97.93 | 0 | 0 | 4.6 | 1.25 | 0.1359 | 0.1615 | 0.0256 |
| 3e3u | noschrodinger | 1.56 | 0.0061 | 1.09 | 0 | 97.94 | 0 | 0 | 2.3 | 1.02 | 0.1394 | 0.163 | 0.0236 |
| 3e3z | opls | 1.7 | 0.1035 | 2.26 | 0 | 97.09 | 1.89 | 1 | 1.68 | 1.29 | 0.152 | 0.1819 | 0.0299 |
| 3e3z | noschrodinger | 1.7 | 0.0096 | 1.14 | 0 | 97.7 | 1.23 | 0 | 4.37 | 1.35 | 0.1455 | 0.1794 | 0.0339 |
| 3e5s | opls | 2 | 0.097 | 2.29 | 0 | 96.03 | 2.25 | 1 | 2.05 | 1.52 | 0.1685 | 0.2036 | 0.0351 |
| 3e5s | noschrodinger | 2 | 0.0042 | 0.92 | 0.39 | 94.88 | 2.68 | 0 | 3.19 | 1.79 | 0.1715 | 0.199 | 0.0275 |
| 3e6p | opls | 2.1 | 0.0958 | 2.3 | 1.02 | 92.58 | 3.81 | 5 | 1.63 | 1.82 | 0.2164 | 0.2284 | 0.012 |
| 3e6p | noschrodinger | 2.1 | 0.0031 | 0.84 | 0.51 | 90.89 | 7.85 | 0 | 2.72 | 2.26 | 0.2075 | 0.2282 | 0.0207 |
| 3e8w | opls | 2.5 | 0.0941 | 2.31 | 0 | 96.76 | 1.07 | 0 | 0.28 | 0.83 | 0.181 | 0.2204 | 0.0394 |
| 3e8w | noschrodinger | 2.5 | 0.0029 | 0.88 | 0.46 | 95.85 | 2.66 | 0 | 1.39 | 1.48 | 0.1801 | 0.2361 | 0.056 |
| 3e8x | opls | 2.09 | 0.097 | 2.41 | 0 | 98.48 | 1.6 | 0 | 1.12 | 0.97 | 0.1922 | 0.2124 | 0.0202 |
| 3e8x | noschrodinger | 2.09 | 0.0031 | 0.81 | 0 | 97.64 | 3.08 | 0 | 2.5 | 1.48 | 0.1809 | 0.2015 | 0.0206 |
| 3ea0 | opls | 2.2 | 0.0925 | 2.35 | 0 | 97.8 | 4.51 | 3 | 1.03 | 1.35 | 0.2164 | 0.2549 | 0.0385 |
| 3ea0 | noschrodinger | 2.2 | 0.0029 | 0.83 | 0 | 97.48 | 7.66 | 0 | 2.45 | 1.8 | 0.1993 | 0.2403 | 0.041 |
| 3ebw | opls | 2.7 | 0.0869 | 2.31 | 0.7 | 91.55 | 5.84 | 6 | 1.22 | 1.93 | 0.2765 | 0.2892 | 0.0127 |
| 3ebw | noschrodinger | 2.7 | 0.0027 | 0.79 | 4.25 | 87.58 | 10.21 | 0 | 3.7 | 2.53 | 0.2611 | 0.2792 | 0.0181 |
| 3ec2 | opls | 2.25 | 0.0931 | 2.44 | 0.61 | 97.56 | 2.58 | 2 | 1.69 | 1.33 | 0.2469 | 0.2696 | 0.0227 |
| 3ec2 | noschrodinger | 2.25 | 0.0046 | 0.97 | 0 | 96.53 | 8.18 | 0 | 8.16 | 2.36 | 0.2298 | 0.257 | 0.0272 |
| 3ec6 | opls | 1.6 | 0.0947 | 2.54 | 0 | 98.28 | 0.79 | 3 | 5.84 | 1.32 | 0.1661 | 0.2024 | 0.0363 |
| 3ec6 | noschrodinger | 1.6 | 0.0084 | 1.06 | 0 | 99.19 | 3.03 | 0 | 3.33 | 1.49 | 0.1683 | 0.1972 | 0.0289 |
| 3eea | opls | 1.78 | 0.0947 | 2.28 | 0 | 99.32 | 1.49 | 0 | 1.77 | 1.07 | 0.1962 | 0.2183 | 0.0221 |
| 3eea | noschrodinger | 1.78 | 0.0032 | 0.85 | 0 | 98.99 | 1.48 | 0 | 3.36 | 1.26 | 0.2 | 0.2161 | 0.0161 |
| 3eeb | opls | 2.1 | 0.097 | 2.3 | 0.25 | 95.99 | 1.19 | 1 | 0.8 | 1.08 | 0.204 | 0.2359 | 0.0319 |
| 3eeb | noschrodinger | 2.1 | 0.005 | 1.57 | 0.25 | 95.77 | 2.07 | 0 | 1.77 | 1.47 | 0.1928 | 0.2333 | 0.0405 |
| 3eeh | opls | 1.95 | 0.1003 | 2.44 | 0.88 | 97.35 | 0 | 1 | 0.54 | 0.81 | 0.1848 | 0.2144 | 0.0296 |
| 3eeh | noschrodinger | 1.95 | 0.0143 | 1.25 | 0.88 | 97.37 | 0 | 1 | 2.69 | 1.18 | 0.1885 | 0.2147 | 0.0262 |
| 3ef5 | opls | 2.4 | 0.0915 | 2.33 | 0.4 | 96.03 | 5.14 | 1 | 0.69 | 1.54 | 0.2413 | 0.2708 | 0.0295 |
| 3ef5 | noschrodinger | 2.4 | 0.0026 | 0.81 | 0.78 | 95.29 | 7.37 | 0 | 3.23 | 2.1 | 0.2405 | 0.2784 | 0.0379 |
| 3efr | opls | 2.55 | 0.0925 | 2.28 | 0 | 94.47 | 4.94 | 2 | 2.19 | 1.9 | 0.2224 | 0.2621 | 0.0397 |
| 3efr | noschrodinger | 2.55 | 0.0033 | 0.84 | 0.43 | 92.64 | 5.87 | 0 | 3.22 | 2.16 | 0.214 | 0.2627 | 0.0487 |
| 3egq | opls | 2.53 | 0.0807 | 2.22 | 0 | 99.3 | 1.96 | 1 | 0.77 | 0.97 | 0.2485 | 0.2821 | 0.0336 |
| 3egq | noschrodinger | 2.53 | 0.0037 | 0.78 | 0.31 | 96.92 | 2.19 | 0 | 1.79 | 1.38 | 0.2682 | 0.3018 | 0.0336 |
| 3egv | opls | 1.75 | 0.1034 | 2.45 | 0 | 97.54 | 1.22 | 3 | 3.08 | 1.26 | 0.1827 | 0.2062 | 0.0235 |
| 3egv | noschrodinger | 1.75 | 0.0076 | 1.1 | 0.31 | 97.86 | 2.43 | 0 | 1.93 | 1.28 | 0.1847 | 0.2071 | 0.0224 |
| 3egw | opls | 1.9 | 0.0992 | 2.25 | 0.15 | 97.05 | 1.75 | 5 | 2.16 | 1.34 | 0.1448 | 0.1757 | 0.0309 |
| 3egw | noschrodinger | 1.9 | 0.0091 | 1.43 | 0.36 | 96.8 | 1.75 | 0 | 2.42 | 1.4 | 0.1505 | 0.182 | 0.0315 |
| 3ehg | opls | 1.74 | 0.1041 | 2.35 | 0 | 98.31 | 0 | 0 | 1.88 | 0.95 | 0.1654 | 0.2013 | 0.0359 |
| 3ehg | noschrodinger | 1.74 | 0.0043 | 0.98 | 0 | 98.37 | 0.86 | 0 | 6.59 | 1.36 | 0.1647 | 0.2001 | 0.0354 |
| 3en8 | opls | 1.85 | 0.0976 | 2.32 | 0 | 100 | 0.97 | 1 | 4.19 | 1.2 | 0.178 | 0.2129 | 0.0349 |
| 3en8 | noschrodinger | 1.85 | 0.004 | 0.89 | 0 | 100 | 0.93 | 0 | 2.8 | 1.07 | 0.1849 | 0.2166 | 0.0317 |
| 3es6 | opls | 3.23 | 0.0887 | 2.43 | 0.77 | 94.33 | 8.6 | 11 | 1.21 | 1.93 | 0.1777 | 0.2228 | 0.0451 |
| 3es6 | noschrodinger | 3.23 | 0.0037 | 0.91 | 0.77 | 92.05 | 12 | 0 | 5.9 | 2.63 | 0.1732 | 0.2198 | 0.0466 |

| | | | | | | | | | | | | | |
|---|---|---|---|---|---|---|---|---|---|---|---|---|---|
| 3esl | opls | 1.74 | 0.0971 | 2.27 | 0 | 98.97 | 1.04 | 2 | 1.3 | 0.87 | 0.1689 | 0.2001 | 0.0312 |
| 3esl | noschrodinger | 1.74 | 0.0056 | 0.87 | 0 | 98.98 | 1.29 | 0 | 1.01 | 0.88 | 0.1712 | 0.1982 | 0.027 |
| 3esm | opls | 1.65 | 0.0988 | 2.23 | 0 | 97.71 | 0 | 1 | 2.43 | 1.09 | 0.212 | 0.253 | 0.041 |
| 3esm | noschrodinger | 1.65 | 0.0064 | 0.96 | 0.76 | 96.21 | 0.88 | 0 | 0.97 | 1.05 | 0.2166 | 0.2492 | 0.0326 |
| 3euj | opls | 3.1 | 0.0919 | 2.3 | 1.17 | 94.55 | 4.47 | 1 | 2.13 | 1.85 | 0.2439 | 0.2667 | 0.0228 |
| 3euj | noschrodinger | 3.1 | 0.0034 | 0.79 | 1.35 | 85.3 | 6.67 | 0 | 8.55 | 2.74 | 0.2378 | 0.262 | 0.0242 |
| 3evi | opls | 2.7 | 0.0915 | 2.29 | 0 | 98.68 | 7.28 | 2 | 0.52 | 1.33 | 0.2365 | 0.2769 | 0.0404 |
| 3evi | noschrodinger | 2.7 | 0.0031 | 0.81 | 0 | 97.84 | 14.76 | 0 | 2.08 | 1.9 | 0.2395 | 0.2642 | 0.0247 |
| 3evo | opls | 1.5 | 0.1059 | 2.34 | 0 | 98.46 | 1.71 | 2 | 3.01 | 1.27 | 0.1386 | 0.1629 | 0.0243 |
| 3evo | noschrodinger | 1.5 | 0.0076 | 1.13 | 0 | 98.47 | 2.12 | 0 | 1.16 | 1.08 | 0.1416 | 0.1622 | 0.0206 |
| 3ewk | opls | 2.34 | 0.0935 | 2.97 | 4.93 | 90.13 | 11.96 | 10 | 4.39 | 2.58 | 0.2432 | 0.2627 | 0.0195 |
| 3ewk | noschrodinger | 2.34 | 0.009 | 1.61 | 6.22 | 86.67 | 15.59 | 0 | 20.66 | 3.34 | 0.2323 | 0.2722 | 0.0399 |
| 3exm | opls | 1.65 | 0.1059 | 2.45 | 0 | 98.55 | 2.22 | 3 | 4.27 | 1.47 | 0.1514 | 0.1877 | 0.0363 |
| 3exm | noschrodinger | 1.65 | 0.0061 | 1.04 | 0 | 98.08 | 2.78 | 1 | 9.11 | 1.82 | 0.1586 | 0.1874 | 0.0288 |
| 3eyk | opls | 2.5 | 0.0989 | 2.22 | 0.21 | 94.42 | 0.47 | 3 | 0.66 | 1.1 | 0.216 | 0.2365 | 0.0205 |
| 3eyk | noschrodinger | 2.5 | 0.003 | 0.79 | 0.62 | 95.07 | 1.17 | 0 | 2.37 | 1.41 | 0.2004 | 0.2279 | 0.0275 |
| 3f2v | opls | 2 | 0.0971 | 2.3 | 0 | 97.02 | 1.41 | 0 | 1.42 | 1.16 | 0.1841 | 0.224 | 0.0399 |
| 3f2v | noschrodinger | 2 | 0.0077 | 0.9 | 0 | 97.06 | 1.39 | 0 | 3.56 | 1.42 | 0.1834 | 0.2265 | 0.0431 |
| 3f4f | opls | 2 | 0.1024 | 2.35 | 0 | 98.76 | 1.2 | 2 | 1.54 | 0.96 | 0.1693 | 0.1932 | 0.0239 |
| 3f4f | noschrodinger | 2 | 0.0034 | 0.83 | 0 | 97.79 | 1.5 | 0 | 2.61 | 1.23 | 0.1576 | 0.1797 | 0.0221 |
| 3f5a | opls | 2 | 0.0972 | 2.29 | 0 | 96.62 | 1.59 | 2 | 0.55 | 1.06 | 0.1722 | 0.1924 | 0.0202 |
| 3f5a | noschrodinger | 2 | 0.0031 | 0.83 | 0 | 95.54 | 0.52 | 0 | 4.98 | 1.57 | 0.1678 | 0.1912 | 0.0234 |
| 3f5b | opls | 2 | 0.0943 | 2.35 | 0 | 96.15 | 3.27 | 1 | 2.39 | 1.67 | 0.1923 | 0.2284 | 0.0361 |
| 3f5b | noschrodinger | 2 | 0.0084 | 1.01 | 1.19 | 97.02 | 5.62 | 0 | 2.05 | 1.72 | 0.1913 | 0.2291 | 0.0378 |
| 3f7q | opls | 1.75 | 0.1023 | 2.27 | 0.24 | 97.39 | 1.33 | 2 | 1.72 | 1.14 | 0.1522 | 0.1836 | 0.0314 |
| 3f7q | noschrodinger | 1.75 | 0.0037 | 0.9 | 0.24 | 98.35 | 1.32 | 0 | 2.29 | 1.1 | 0.158 | 0.1815 | 0.0235 |
| 3f7r | opls | 2.04 | 0.0979 | 2.2 | 0 | 96.21 | 1.08 | 1 | 0.88 | 1.05 | 0.1877 | 0.2108 | 0.0231 |
| 3f7r | noschrodinger | 2.04 | 0.008 | 1.01 | 0.47 | 96.7 | 1.6 | 0 | 2.34 | 1.38 | 0.1718 | 0.2054 | 0.0336 |
| 3f8z | opls | 2.01 | 0.1044 | 2.26 | 0 | 97.83 | 4.76 | 2 | 2.86 | 1.63 | 0.1699 | 0.2013 | 0.0314 |
| 3f8z | noschrodinger | 2.01 | 0.0035 | 1.02 | 0 | 97.83 | 4.76 | 0 | 2.22 | 1.55 | 0.1617 | 0.1947 | 0.033 |
| 3fbr | opls | 3.5 | 0.0904 | 2.38 | 0.92 | 94.46 | 5.95 | 1 | 1.84 | 1.91 | 0.2835 | 0.3082 | 0.0247 |
| 3fbr | noschrodinger | 3.5 | 0.0029 | 0.79 | 2.76 | 85.25 | 9.43 | 0 | 4.98 | 2.66 | 0.2685 | 0.2955 | 0.027 |
| 3fbs | opls | 2.15 | 0.0973 | 2.3 | 0.34 | 95.35 | 0.92 | 2 | 0.55 | 1.01 | 0.1954 | 0.22 | 0.0246 |
| 3fbs | noschrodinger | 2.15 | 0.0028 | 0.8 | 0.17 | 95.38 | 2.5 | 0 | 0.99 | 1.42 | 0.1802 | 0.2106 | 0.0304 |
| 3fbu | opls | 1.8 | 0.0957 | 2.24 | 0 | 99.06 | 0 | 1 | 0.89 | 0.77 | 0.1764 | 0.2053 | 0.0289 |
| 3fbu | noschrodinger | 1.8 | 0.0038 | 0.87 | 0 | 98.15 | 0.68 | 0 | 0.54 | 0.68 | 0.1833 | 0.2026 | 0.0193 |
| 3fdx | opls | 1.58 | 0.0996 | 2.7 | 0.87 | 98.27 | 2.46 | 4 | 7.78 | 1.72 | 0.1956 | 0.2283 | 0.0327 |
| 3fdx | noschrodinger | 1.58 | 0.0116 | 1.36 | 0.82 | 97.14 | 1.95 | 0 | 8.84 | 1.85 | 0.1938 | 0.2286 | 0.0348 |
| 3fe2 | opls | 2.6 | 0.0924 | 2.54 | 0 | 98.48 | 5.03 | 5 | 1.46 | 1.42 | 0.1859 | 0.2426 | 0.0567 |
| 3fe2 | noschrodinger | 2.6 | 0.0039 | 0.86 | 0.43 | 97.63 | 7.04 | 0 | 1.33 | 1.58 | 0.1992 | 0.2401 | 0.0409 |
| 3ff4 | opls | 2.1 | 0.0961 | 2.27 | 0 | 98.23 | 0.92 | 0 | 1.97 | 0.96 | 0.1877 | 0.2129 | 0.0252 |
| 3ff4 | noschrodinger | 2.1 | 0.0037 | 0.82 | 0 | 99.16 | 1.8 | 0 | 0.98 | 0.99 | 0.1813 | 0.2116 | 0.0303 |
| 3fid | opls | 1.9 | 0.0974 | 2.37 | 0 | 99.32 | 1.69 | 2 | 2.53 | 1.21 | 0.1722 | 0.2164 | 0.0442 |
| 3fid | noschrodinger | 1.9 | 0.0128 | 1.27 | 0 | 98.81 | 2.11 | 0 | 2.22 | 1.24 | 0.1732 | 0.2126 | 0.0394 |
| 3fmo | opls | 2.51 | 0.091 | 2.31 | 0.15 | 96.45 | 4.84 | 4 | 2.02 | 1.73 | 0.1893 | 0.2128 | 0.0235 |
| 3fmo | noschrodinger | 2.51 | 0.0054 | 1.01 | 0.62 | 94.3 | 6.74 | 0 | 4.53 | 2.24 | 0.1938 | 0.2117 | 0.0179 |
| 3fmz | opls | 2.9 | 0.0902 | 2.23 | 0.58 | 94.74 | 5.67 | 2 | 1.26 | 1.78 | 0.2285 | 0.2541 | 0.0256 |
| 3fmz | noschrodinger | 2.9 | 0.003 | 0.79 | 0.87 | 91.86 | 9.27 | 0 | 3.23 | 2.34 | 0.2051 | 0.2502 | 0.0451 |
| 3fna | opls | 2.03 | 0.0896 | 2.36 | 0 | 98.31 | 2.59 | 0 | 0.49 | 0.99 | 0.2331 | 0.2647 | 0.0316 |

| | | | | | | | | | | | | | |
|---|---|---|---|---|---|---|---|---|---|---|---|---|---|
| 3fna | noschrodinger | 2.03 | 0.0031 | 0.85 | 0 | 97.87 | 4.17 | 0 | 1.49 | 1.39 | 0.235 | 0.2731 | 0.0381 |
| 3fnn | opls | 2.3 | 0.0935 | 2.32 | 0 | 98.17 | 3.72 | 0 | 0.83 | 1.19 | 0.2137 | 0.239 | 0.0253 |
| 3fnn | noschrodinger | 2.3 | 0.0028 | 0.76 | 0 | 95.93 | 5.24 | 0 | 1.11 | 1.64 | 0.2088 | 0.2335 | 0.0247 |
| 3fo5 | opls | 2 | 0.0937 | 2.28 | 0 | 98.45 | 1.43 | 2 | 1.56 | 1.02 | 0.2055 | 0.2372 | 0.0317 |
| 3fo5 | noschrodinger | 2 | 0.0028 | 0.78 | 0 | 97.85 | 2.36 | 0 | 1.56 | 1.22 | 0.2078 | 0.233 | 0.0252 |
| 3fpf | opls | 1.66 | 0.1018 | 2.53 | 0 | 98.66 | 3.51 | 7 | 5.23 | 1.69 | 0.1685 | 0.1984 | 0.0299 |
| 3fpf | noschrodinger | 1.66 | 0.0074 | 1.13 | 0 | 98.47 | 3.71 | 2 | 5.7 | 1.74 | 0.1721 | 0.1959 | 0.0238 |
| 3fpg | opls | 2 | 0.102 | 2.35 | 0 | 98.08 | 3.07 | 2 | 2.36 | 1.39 | 0.2105 | 0.2518 | 0.0413 |
| 3fpg | noschrodinger | 2 | 0.0032 | 0.8 | 0 | 97.9 | 4.59 | 0 | 4.6 | 1.76 | 0.1982 | 0.2409 | 0.0427 |
| 3fpt | opls | 2.7 | 0.0893 | 3.32 | 2.55 | 93.19 | 9.76 | 26 | 4.16 | 2.39 | 0.2158 | 0.2493 | 0.0335 |
| 3fpt | noschrodinger | 2.7 | 0.0046 | 1.07 | 2.06 | 91.77 | 14.9 | 0 | 8.91 | 2.86 | 0.2392 | 0.2527 | 0.0135 |
| 3fqq | opls | 2.2 | 0.0976 | 2.28 | 0 | 98.72 | 4.12 | 1 | 1.22 | 1.31 | 0.1962 | 0.2448 | 0.0486 |
| 3fqq | noschrodinger | 2.2 | 0.0031 | 0.84 | 0 | 97.14 | 4.09 | 0 | 1.43 | 1.5 | 0.2035 | 0.246 | 0.0425 |
| 3frl | opls | 2.25 | 0.0958 | 2.24 | 0.24 | 97.61 | 2.62 | 1 | 1.37 | 1.27 | 0.211 | 0.2324 | 0.0214 |
| 3frl | noschrodinger | 2.25 | 0.0032 | 0.84 | 0.22 | 96.47 | 3.27 | 0 | 1.51 | 1.52 | 0.209 | 0.2358 | 0.0268 |
| 3fs8 | opls | 1.7 | 0.1003 | 2.54 | 0 | 96.86 | 1.1 | 3 | 2.37 | 1.24 | 0.1734 | 0.2152 | 0.0418 |
| 3fs8 | noschrodinger | 1.7 | 0.0057 | 1.03 | 0 | 97.27 | 3.49 | 0 | 2.6 | 1.6 | 0.1737 | 0.213 | 0.0393 |
| 3fus | opls | 4 | 0.0804 | 2.82 | 7.07 | 72.05 | 14.87 | 5 | 6.02 | 3.04 | 0.3712 | 0.4113 | 0.0401 |
| 3fus | noschrodinger | 4 | 0.0064 | 1.19 | 12.67 | 65 | 21.69 | 0 | 27.68 | 3.83 | 0.3384 | 0.3837 | 0.0453 |
| 3fws | opls | 2.03 | 0.0941 | 2.47 | 0.35 | 97.19 | 4.15 | 1 | 1.66 | 1.53 | 0.2102 | 0.2447 | 0.0345 |
| 3fws | noschrodinger | 2.03 | 0.0033 | 0.94 | 0.35 | 96.52 | 8.99 | 0 | 4.16 | 2.15 | 0.1896 | 0.226 | 0.0364 |
| 3fwz | opls | 1.79 | 0.0974 | 2.29 | 0 | 99.17 | 0.46 | 0 | 2.02 | 0.97 | 0.1588 | 0.1879 | 0.0291 |
| 3fwz | noschrodinger | 1.79 | 0.0036 | 0.87 | 0 | 99.26 | 0.45 | 0 | 2.25 | 1 | 0.1636 | 0.1877 | 0.0241 |
| 3fyb | opls | 1.8 | 0.0978 | 2.54 | 0 | 98.66 | 0.71 | 0 | 3.97 | 1.18 | 0.1921 | 0.2225 | 0.0304 |
| 3fyb | noschrodinger | 1.8 | 0.0131 | 1.02 | 0 | 98.35 | 1.32 | 0 | 11.26 | 1.66 | 0.1811 | 0.2239 | 0.0428 |
| 3fyk | opls | 3.5 | 0.09 | 2.39 | 1.1 | 88.24 | 7.54 | 2 | 2.15 | 2.25 | 0.2554 | 0.2838 | 0.0284 |
| 3fyk | noschrodinger | 3.5 | 0.0033 | 0.94 | 7.25 | 72.46 | 11.33 | 0 | 10.11 | 3.15 | 0.2374 | 0.265 | 0.0276 |
| 3g0m | opls | 1.76 | 0.0958 | 2.34 | 0 | 100 | 0 | 0 | 0.88 | 0.77 | 0.1972 | 0.2464 | 0.0492 |
| 3g0m | noschrodinger | 1.76 | 0.0043 | 0.78 | 0 | 100 | 0 | 0 | 1.76 | 0.93 | 0.2004 | 0.2473 | 0.0469 |
| 3g2p | opls | 2.95 | 0.0897 | 2.37 | 1.35 | 96.64 | 4.72 | 4 | 0.53 | 1.41 | 0.2278 | 0.2532 | 0.0254 |
| 3g2p | noschrodinger | 2.95 | 0.0034 | 0.87 | 1.91 | 92.34 | 4.91 | 0 | 2.29 | 2.01 | 0.2174 | 0.246 | 0.0286 |
| 3g3r | opls | 2 | 0.0974 | 2.24 | 0 | 99.06 | 0.59 | 1 | 1.26 | 0.85 | 0.208 | 0.2475 | 0.0395 |
| 3g3r | noschrodinger | 2 | 0.0027 | 0.76 | 0 | 97.52 | 1.76 | 0 | 2.32 | 1.29 | 0.1992 | 0.2438 | 0.0446 |
| 3g7p | opls | 2 | 0.095 | 2.39 | 0.89 | 96.43 | 0.88 | 0 | 1.24 | 1.08 | 0.1935 | 0.2307 | 0.0372 |
| 3g7p | noschrodinger | 2 | 0.0114 | 1.16 | 0 | 95.8 | 1.65 | 0 | 4.15 | 1.65 | 0.1976 | 0.2272 | 0.0296 |
| 3gan | opls | 2 | 0.0906 | 2.58 | 0 | 96.74 | 0 | 0 | 0.95 | 0.99 | 0.1868 | 0.2127 | 0.0259 |
| 3gan | noschrodinger | 2 | 0.004 | 1.18 | 0 | 98.17 | 0 | 0 | 1.45 | 0.88 | 0.1914 | 0.2194 | 0.028 |
| 3gb5 | opls | 2 | 0.0976 | 2.32 | 0.57 | 98.86 | 2.35 | 2 | 2.21 | 1.28 | 0.1609 | 0.177 | 0.0161 |
| 3gb5 | noschrodinger | 2 | 0.0046 | 0.89 | 0 | 98.32 | 3.47 | 0 | 2.22 | 1.41 | 0.1647 | 0.1787 | 0.014 |
| 3gg7 | opls | 1.5 | 0.0975 | 2.32 | 0 | 98.27 | 0.98 | 0 | 1.3 | 0.85 | 0.1699 | 0.192 | 0.0221 |
| 3gg7 | noschrodinger | 1.5 | 0.0063 | 1.05 | 0 | 98.34 | 0.98 | 0 | 0.52 | 0.68 | 0.1725 | 0.1937 | 0.0212 |
| 3ggl | opls | 3 | 0.0898 | 2.28 | 0 | 94.81 | 6.2 | 0 | 2.02 | 1.93 | 0.2445 | 0.2683 | 0.0238 |
| 3ggl | noschrodinger | 3 | 0.0029 | 0.82 | 1.26 | 87.42 | 11.36 | 0 | 4.04 | 2.6 | 0.23 | 0.2564 | 0.0264 |
| 3gk6 | opls | 1.8 | 0.0962 | 2.27 | 0 | 94.85 | 2.29 | 1 | 1.2 | 1.47 | 0.2075 | 0.245 | 0.0375 |
| 3gk6 | noschrodinger | 1.8 | 0.0052 | 0.87 | 0 | 96.64 | 0.73 | 0 | 0.8 | 0.97 | 0.2112 | 0.2415 | 0.0303 |
| 3gkj | opls | 1.6 | 0.0948 | 2.54 | 0 | 97.3 | 3.5 | 7 | 3.67 | 1.7 | 0.1632 | 0.2041 | 0.0409 |
| 3gkj | noschrodinger | 1.6 | 0.0049 | 1.03 | 0 | 96.86 | 2.49 | 0 | 3.67 | 1.65 | 0.1718 | 0.2032 | 0.0314 |
| 3glv | opls | 1.99 | 0.0988 | 2.55 | 0.42 | 97.03 | 1.42 | 2 | 2.25 | 1.29 | 0.1785 | 0.2241 | 0.0456 |
| 3glv | noschrodinger | 1.99 | 0.0044 | 0.78 | 0.42 | 97.08 | 2.34 | 0 | 2.26 | 1.45 | 0.1913 | 0.2219 | 0.0306 |

| | | | | | | | | | | | | | |
|---|---|---|---|---|---|---|---|---|---|---|---|---|---|
| 3goc | opls | 1.5 | 0.1082 | 2.5 | 0 | 98 | 0.29 | 4 | 3.39 | 1.13 | 0.152 | 0.1822 | 0.0302 |
| 3goc | noschrodinger | 1.5 | 0.0044 | 0.99 | 0 | 98.01 | 0 | 0 | 1.77 | 0.93 | 0.1581 | 0.1811 | 0.023 |
| 3gqk | opls | 2.5 | 0.0901 | 2.29 | 0.62 | 96.88 | 5.22 | 0 | 1.93 | 1.69 | 0.2881 | 0.3125 | 0.0244 |
| 3gqk | noschrodinger | 2.5 | 0.0028 | 0.85 | 1.24 | 88.82 | 5.19 | 0 | 3.1 | 2.22 | 0.2892 | 0.3077 | 0.0185 |
| 3gud | opls | 2.2 | 0.0938 | 2.41 | 0.43 | 96.55 | 8.37 | 4 | 1.55 | 1.82 | 0.2572 | 0.2824 | 0.0252 |
| 3gud | noschrodinger | 2.2 | 0.0032 | 0.81 | 0.85 | 91.91 | 13.11 | 0 | 6.46 | 2.69 | 0.2535 | 0.2969 | 0.0434 |
| 3guz | opls | 1.67 | 0.0988 | 2.28 | 0 | 99.04 | 1.75 | 1 | 1.11 | 1 | 0.1839 | 0.2099 | 0.026 |
| 3guz | noschrodinger | 1.67 | 0.0053 | 0.95 | 0 | 99.07 | 2.76 | 0 | 2.78 | 1.4 | 0.1862 | 0.2047 | 0.0185 |
| 3gv1 | opls | 2 | 0.094 | 2.23 | 0 | 99.19 | 1.47 | 1 | 0.91 | 0.9 | 0.1864 | 0.2083 | 0.0219 |
| 3gv1 | noschrodinger | 2 | 0.0031 | 0.82 | 0.24 | 98.31 | 2.51 | 0 | 1.67 | 1.22 | 0.1709 | 0.1958 | 0.0249 |
| 3gwl | opls | 2.1 | 0.098 | 2.52 | 0 | 98.51 | 0.52 | 1 | 0.82 | 0.75 | 0.2003 | 0.2459 | 0.0456 |
| 3gwl | noschrodinger | 2.1 | 0.0027 | 0.72 | 0 | 99.02 | 1.03 | 0 | 1.91 | 0.97 | 0.1983 | 0.248 | 0.0497 |
| 3gwn | opls | 1.78 | 0.0992 | 2.45 | 0 | 98.62 | 0 | 0 | 1.24 | 0.84 | 0.1385 | 0.1806 | 0.0421 |
| 3gwn | noschrodinger | 1.78 | 0.0037 | 0.88 | 0 | 98.65 | 0.47 | 0 | 2.97 | 1.09 | 0.1449 | 0.1827 | 0.0378 |
| 3gyd | opls | 1.79 | 0.0962 | 2.3 | 0 | 99.32 | 0.34 | 3 | 2.19 | 0.99 | 0.1654 | 0.2063 | 0.0409 |
| 3gyd | noschrodinger | 1.79 | 0.0033 | 0.86 | 0.28 | 99.15 | 1.61 | 0 | 1.18 | 0.99 | 0.1744 | 0.2041 | 0.0297 |
| 3h2b | opls | 2 | 0.0978 | 2.36 | 0 | 99.17 | 2.98 | 4 | 0.84 | 1.12 | 0.1591 | 0.2005 | 0.0414 |
| 3h2b | noschrodinger | 2 | 0.0032 | 0.9 | 0 | 98.45 | 2.88 | 0 | 1.18 | 1.18 | 0.1678 | 0.2015 | 0.0337 |
| 3h4s | opls | 2.4 | 0.0942 | 2.35 | 0.46 | 97.49 | 5.53 | 1 | 0.96 | 1.45 | 0.2165 | 0.256 | 0.0395 |
| 3h4s | noschrodinger | 2.4 | 0.0032 | 0.84 | 0.68 | 95.72 | 8.17 | 0 | 1.92 | 1.95 | 0.2054 | 0.2525 | 0.0471 |
| 3h7j | opls | 1.87 | 0.0946 | 2.24 | 0.23 | 97.96 | 3.55 | 1 | 1.4 | 1.3 | 0.1942 | 0.2229 | 0.0287 |
| 3h7j | noschrodinger | 1.87 | 0.0039 | 0.94 | 0.23 | 98.2 | 6.06 | 0 | 3.37 | 1.72 | 0.1932 | 0.2202 | 0.027 |
| 3h8v | opls | 2 | 0.0938 | 2.24 | 0.25 | 97.74 | 0.55 | 1 | 0.87 | 0.83 | 0.2489 | 0.2674 | 0.0185 |
| 3h8v | noschrodinger | 2 | 0.0029 | 0.79 | 0 | 97.28 | 3.01 | 0 | 1.18 | 1.33 | 0.2414 | 0.262 | 0.0206 |
| 3h9a | opls | 2.04 | 0.0924 | 2.44 | 0 | 96.84 | 2.42 | 1 | 0.98 | 1.27 | 0.2098 | 0.2318 | 0.022 |
| 3h9a | noschrodinger | 2.04 | 0.0029 | 0.81 | 0 | 97.09 | 3.77 | 0 | 2.1 | 1.58 | 0.2006 | 0.2308 | 0.0302 |
| 3ha2 | opls | 1.8 | 0.0943 | 2.26 | 0 | 96.69 | 0 | 1 | 0.93 | 0.99 | 0.1807 | 0.2186 | 0.0379 |
| 3ha2 | noschrodinger | 1.8 | 0.0031 | 0.81 | 0 | 96.69 | 0.7 | 0 | 1.48 | 1.1 | 0.1872 | 0.2196 | 0.0324 |
| 3ham | opls | 2.3 | 0.0945 | 2.32 | 0.68 | 95.93 | 6.16 | 11 | 1.29 | 1.73 | 0.2263 | 0.2676 | 0.0413 |
| 3ham | noschrodinger | 2.3 | 0.0062 | 1.06 | 1.01 | 93.43 | 9.53 | 0 | 4.46 | 2.4 | 0.2202 | 0.2684 | 0.0482 |
| 3hav | opls | 2.4 | 0.0921 | 2.4 | 1.41 | 93.76 | 8.32 | 18 | 2.22 | 2.11 | 0.2494 | 0.2668 | 0.0174 |
| 3hav | noschrodinger | 2.4 | 0.0038 | 1.17 | 2.78 | 88.99 | 9.46 | 0 | 5.08 | 2.59 | 0.2362 | 0.2567 | 0.0205 |
| 3hbn | opls | 1.85 | 0.099 | 2.35 | 0.37 | 98.17 | 0 | 0 | 1.27 | 0.85 | 0.1536 | 0.1933 | 0.0397 |
| 3hbn | noschrodinger | 1.85 | 0.0037 | 0.88 | 0.36 | 97.86 | 0.78 | 0 | 0.64 | 0.74 | 0.1596 | 0.1922 | 0.0326 |
| 3hch | opls | 2.1 | 0.0942 | 2.25 | 0 | 97.14 | 1.98 | 5 | 0.4 | 1.02 | 0.1886 | 0.2087 | 0.0201 |
| 3hch | noschrodinger | 2.1 | 0.0034 | 0.9 | 0 | 96.44 | 3.15 | 4 | 3.61 | 1.76 | 0.1847 | 0.2005 | 0.0158 |
| 3hee | opls | 2 | 0.0971 | 2.34 | 0 | 99.65 | 1.33 | 3 | 1.32 | 0.95 | 0.1653 | 0.1936 | 0.0283 |
| 3hee | noschrodinger | 2 | 0.0062 | 0.86 | 0 | 98.97 | 1.3 | 0 | 1.76 | 1.02 | 0.1666 | 0.1908 | 0.0242 |
| 3hf6 | opls | 1.8 | 0.0976 | 2.44 | 0.42 | 98.31 | 3.86 | 4 | 3.54 | 1.59 | 0.1526 | 0.1919 | 0.0393 |
| 3hf6 | noschrodinger | 1.8 | 0.0035 | 0.87 | 0 | 98.91 | 2.81 | 0 | 5.38 | 1.63 | 0.1581 | 0.1866 | 0.0285 |
| 3hf7 | opls | 2.75 | 0.0876 | 2.38 | 0 | 96.08 | 6.8 | 0 | 0 | 1.4 | 0.2448 | 0.272 | 0.0272 |
| 3hf7 | noschrodinger | 2.75 | 0.0029 | 0.81 | 0 | 94.4 | 13.59 | 0 | 2.58 | 2.29 | 0.2313 | 0.2614 | 0.0301 |
| 3hhx | opls | 1.89 | 0.0996 | 2.29 | 0 | 98.42 | 1.41 | 2 | 1.48 | 1 | 0.2228 | 0.2661 | 0.0433 |
| 3hhx | noschrodinger | 1.89 | 0.0029 | 0.83 | 0.39 | 98.03 | 2.35 | 0 | 2.24 | 1.28 | 0.224 | 0.2605 | 0.0365 |
| 3hhy | opls | 1.55 | 0.1008 | 2.62 | 0 | 98.42 | 4.63 | 7 | 4.14 | 1.7 | 0.1831 | 0.2187 | 0.0356 |
| 3hhy | noschrodinger | 1.55 | 0.0122 | 1.3 | 0 | 98.43 | 5.56 | 1 | 7.35 | 1.97 | 0.1848 | 0.2179 | 0.0331 |
| 3hjk | opls | 1.98 | 0.1009 | 2.29 | 0 | 97.89 | 2.7 | 3 | 2.08 | 1.33 | 0.211 | 0.2604 | 0.0494 |
| 3hjk | noschrodinger | 1.98 | 0.0043 | 0.86 | 0 | 97.2 | 4.21 | 0 | 2.5 | 1.66 | 0.2044 | 0.2521 | 0.0477 |
| 3hkv | opls | 2.1 | 0.0971 | 2.33 | 0 | 98.94 | 3.06 | 3 | 1.45 | 1.25 | 0.1992 | 0.2237 | 0.0245 |

| | | | | | | | | | | | | | |
|---|---|---|---|---|---|---|---|---|---|---|---|---|---|
| 3hkv | noschrodinger | 2.1 | 0.0057 | 0.94 | 0 | 98.68 | 4.56 | 0 | 3.23 | 1.62 | 0.1844 | 0.2121 | 0.0277 |
| 3hl4 | opls | 2.2 | 0.0976 | 2.27 | 0 | 97.69 | 0 | 0 | 1.58 | 0.97 | 0.2272 | 0.2489 | 0.0217 |
| 3hl4 | noschrodinger | 2.2 | 0.0028 | 0.8 | 0 | 97.99 | 1.62 | 0 | 2.46 | 1.19 | 0.2225 | 0.2505 | 0.028 |
| 3hnh | opls | 1.8 | 0.0946 | 2.36 | 0.44 | 98.25 | 1 | 1 | 1.03 | 0.8 | 0.2053 | 0.2442 | 0.0389 |
| 3hnh | noschrodinger | 1.8 | 0.005 | 0.89 | 0.43 | 97.84 | 3.45 | 0 | 3.36 | 1.57 | 0.2054 | 0.24 | 0.0346 |
| 3ho6 | opls | 1.6 | 0.104 | 2.26 | 0.21 | 98.12 | 0.86 | 1 | 1.97 | 0.96 | 0.1817 | 0.2158 | 0.0341 |
| 3ho6 | noschrodinger | 1.6 | 0.0044 | 1.12 | 0.21 | 98.35 | 0.85 | 0 | 2.59 | 1.04 | 0.1792 | 0.2127 | 0.0335 |
| 3hpe | opls | 2.1 | 0.0927 | 2.25 | 0.31 | 95.65 | 2.82 | 1 | 1.47 | 1.53 | 0.2493 | 0.2761 | 0.0268 |
| 3hpe | noschrodinger | 2.1 | 0.0029 | 0.83 | 0 | 93.83 | 5.94 | 0 | 4.24 | 2.2 | 0.2283 | 0.2659 | 0.0376 |
| 3hy8 | opls | 2.3 | 0.0927 | 2.41 | 1.94 | 93.69 | 3.78 | 0 | 1.43 | 1.73 | 0.2375 | 0.2935 | 0.056 |
| 3hy8 | noschrodinger | 2.3 | 0.0042 | 0.88 | 2.9 | 90.34 | 3.23 | 0 | 6.88 | 2.31 | 0.2368 | 0.2969 | 0.0601 |
| 3hyo | opls | 1.85 | 0.1006 | 2.27 | 0 | 96.51 | 0 | 1 | 1.21 | 1.07 | 0.1651 | 0.2019 | 0.0368 |
| 3hyo | noschrodinger | 1.85 | 0.0037 | 0.94 | 0 | 96.31 | 0.93 | 0 | 1.94 | 1.21 | 0.1657 | 0.2004 | 0.0347 |
| 3hys | opls | 2.3 | 0.0993 | 2.28 | 0 | 98.11 | 1.43 | 2 | 0.72 | 0.85 | 0.1853 | 0.2028 | 0.0175 |
| 3hys | noschrodinger | 2.3 | 0.0033 | 0.88 | 0 | 98.5 | 3.32 | 0 | 2.05 | 1.37 | 0.1686 | 0.192 | 0.0234 |
| 3hzo | opls | 2.6 | 0.097 | 2.29 | 0 | 97.74 | 1.91 | 0 | 0.61 | 0.98 | 0.1752 | 0.2129 | 0.0377 |
| 3hzo | noschrodinger | 2.6 | 0.0032 | 0.86 | 0 | 98.31 | 5.25 | 0 | 1.47 | 1.43 | 0.1687 | 0.1984 | 0.0297 |
| 3hzs | opls | 2.1 | 0.0955 | 2.27 | 0.49 | 97.57 | 1.04 | 1 | 2.22 | 1.1 | 0.1951 | 0.2308 | 0.0357 |
| 3hzs | noschrodinger | 2.1 | 0.0101 | 1.33 | 0.48 | 97.58 | 2.58 | 0 | 1.66 | 1.32 | 0.1921 | 0.2306 | 0.0385 |
| 3i3x | opls | 1.96 | 0.0945 | 2.34 | 0 | 97.25 | 1.09 | 3 | 0.9 | 0.94 | 0.2085 | 0.2325 | 0.024 |
| 3i3x | noschrodinger | 1.96 | 0.0033 | 0.87 | 0 | 96.76 | 1.24 | 0 | 1.81 | 1.21 | 0.203 | 0.2289 | 0.0259 |
| 3i5c | opls | 1.94 | 0.0975 | 2.44 | 0 | 99.21 | 3.4 | 3 | 1.71 | 1.33 | 0.1997 | 0.2354 | 0.0357 |
| 3i5c | noschrodinger | 1.94 | 0.0138 | 1.44 | 0 | 97.95 | 3.68 | 0 | 4.85 | 1.7 | 0.2012 | 0.242 | 0.0408 |
| 3i7d | opls | 2.3 | 0.0954 | 2.19 | 0 | 99.26 | 0.42 | 0 | 0.42 | 0.65 | 0.2141 | 0.2473 | 0.0332 |
| 3i7d | noschrodinger | 2.3 | 0.0027 | 0.77 | 0 | 99.04 | 1.58 | 0 | 0.64 | 0.86 | 0.198 | 0.2391 | 0.0411 |
| 3i95 | opls | 1.4 | 0.104 | 2.66 | 0 | 97.92 | 1.65 | 2 | 6.35 | 1.53 | 0.1462 | 0.1678 | 0.0216 |
| 3i95 | noschrodinger | 1.4 | 0.0105 | 1.22 | 0 | 97.93 | 1.64 | 0 | 2.35 | 1.2 | 0.1497 | 0.1639 | 0.0142 |
| 3ibq | opls | 2 | 0.0983 | 2.31 | 0 | 96.5 | 0.48 | 0 | 1.46 | 1.11 | 0.1684 | 0.1966 | 0.0282 |
| 3ibq | noschrodinger | 2 | 0.0031 | 0.86 | 0 | 97.04 | 1.41 | 0 | 1.95 | 1.24 | 0.1708 | 0.1963 | 0.0255 |
| 3igd | opls | 2.4 | 0.0941 | 2.33 | 0 | 98.52 | 3.6 | 0 | 0.92 | 1.2 | 0.2965 | 0.3206 | 0.0241 |
| 3igd | noschrodinger | 2.4 | 0.0027 | 0.74 | 1.47 | 91.18 | 4.46 | 0 | 3.66 | 2.16 | 0.2718 | 0.3157 | 0.0439 |
| 3igs | opls | 1.5 | 0.109 | 2.62 | 0 | 97.2 | 1.1 | 5 | 8.25 | 1.63 | 0.1308 | 0.1554 | 0.0246 |
| 3igs | noschrodinger | 1.5 | 0.0059 | 1.08 | 0 | 97.16 | 2.13 | 4 | 5.49 | 1.7 | 0.1364 | 0.152 | 0.0156 |
| 3iif | opls | 2.1 | 0.0937 | 2.32 | 0 | 96.77 | 5.48 | 4 | 0.81 | 1.51 | 0.1995 | 0.2217 | 0.0222 |
| 3iif | noschrodinger | 2.1 | 0.0038 | 0.99 | 0.55 | 96.12 | 7.44 | 0 | 2.44 | 1.95 | 0.1851 | 0.2152 | 0.0301 |
| 3iij | opls | 1.76 | 0.1012 | 2.71 | 0 | 96.53 | 1.2 | 1 | 0.68 | 1.01 | 0.166 | 0.1903 | 0.0243 |
| 3iij | noschrodinger | 1.76 | 0.0132 | 1.29 | 0 | 94.8 | 1.81 | 0 | 1.02 | 1.35 | 0.1659 | 0.1915 | 0.0256 |
| 3iil | opls | 2 | 0.1018 | 2.54 | 0 | 98.27 | 2.44 | 1 | 1.37 | 1.16 | 0.1714 | 0.1936 | 0.0222 |
| 3iil | noschrodinger | 2 | 0.0072 | 0.99 | 0 | 97.11 | 2.44 | 0 | 0.69 | 1.18 | 0.1697 | 0.1973 | 0.0276 |
| 3im0 | opls | 1.66 | 0.1083 | 2.31 | 0.42 | 97.08 | 2.35 | 2 | 7.02 | 1.83 | 0.1318 | 0.1647 | 0.0329 |
| 3im0 | noschrodinger | 1.66 | 0.0069 | 1.04 | 0.41 | 96.27 | 3.74 | 0 | 7.01 | 2.07 | 0.1336 | 0.1628 | 0.0292 |
| 3ivl | opls | 2.2 | 0.0897 | 2.28 | 0 | 98.88 | 8.85 | 4 | 1.62 | 1.63 | 0.2242 | 0.2473 | 0.0231 |
| 3ivl | noschrodinger | 2.2 | 0.003 | 0.83 | 0 | 99.18 | 13.24 | 0 | 2.55 | 1.89 | 0.2299 | 0.2494 | 0.0195 |
| 3jqy | opls | 1.7 | 0.1058 | 2.3 | 0 | 98.48 | 1.2 | 2 | 2.17 | 1.05 | 0.1576 | 0.185 | 0.0274 |
| 3jqy | noschrodinger | 1.7 | 0.0037 | 0.82 | 0 | 98.49 | 1.19 | 0 | 1.98 | 1.02 | 0.1577 | 0.1829 | 0.0252 |
| 3jr7 | opls | 2 | 0.1004 | 2.28 | 0 | 96.63 | 1.53 | 1 | 1.38 | 1.22 | 0.1678 | 0.1942 | 0.0264 |
| 3jr7 | noschrodinger | 2 | 0.0029 | 0.75 | 0 | 97.45 | 1.26 | 0 | 1.15 | 1.01 | 0.1715 | 0.1903 | 0.0188 |
| 3jt7 | opls | 2.1 | 0.0967 | 2.27 | 0 | 97.65 | 0.27 | 5 | 1.77 | 1.01 | 0.2053 | 0.2292 | 0.0239 |
| 3jt7 | noschrodinger | 2.1 | 0.0031 | 0.83 | 0 | 98.02 | 1.77 | 0 | 2.89 | 1.27 | 0.1844 | 0.2217 | 0.0373 |

| | | | | | | | | | | | |
|---|---|---|---|---|---|---|---|---|---|---|---|
| 3jt8 | opls | 1.95 | 0.0952 | 2.48 | 0.12 | 96.53 | 0.96 | 11 | 5.31 | 1.51 | 0.1988 | 0.2371 | 0.0383 |
| 3jt8 | noschrodinger | 1.95 | 0.0079 | 0.97 | 0 | 97.9 | 1.5 | 0 | 4.06 | 1.35 | 0.2078 | 0.2353 | 0.0275 |
| 3jtf | opls | 2 | 0.0962 | 2.37 | 0 | 98.12 | 1.82 | 0 | 1.85 | 1.14 | 0.1948 | 0.2232 | 0.0284 |
| 3jtf | noschrodinger | 2 | 0.0029 | 0.83 | 0 | 97.94 | 1.73 | 0 | 2.55 | 1.24 | 0.1767 | 0.2203 | 0.0436 |
| 3juo | opls | 2.2 | 0.0933 | 2.22 | 0 | 97.97 | 1.5 | 0 | 0.78 | 0.89 | 0.2161 | 0.2523 | 0.0362 |
| 3juo | noschrodinger | 2.2 | 0.0026 | 0.75 | 0 | 98.38 | 1.87 | 0 | 1.37 | 1.07 | 0.2043 | 0.2393 | 0.035 |
| 3jz0 | opls | 2 | 0.1035 | 2.22 | 0 | 97.04 | 1.22 | 5 | 1.55 | 1.13 | 0.1555 | 0.2003 | 0.0448 |
| 3jz0 | noschrodinger | 2 | 0.0052 | 0.82 | 0 | 97.92 | 2.23 | 0 | 0.78 | 1.03 | 0.1596 | 0.1939 | 0.0343 |
| 3k0t | opls | 2.1 | 0.0969 | 2.44 | 0 | 99.08 | 1.09 | 0 | 1.6 | 0.94 | 0.1757 | 0.1973 | 0.0216 |
| 3k0t | noschrodinger | 2.1 | 0.0029 | 0.84 | 0 | 99.45 | 1.04 | 0 | 1.24 | 0.86 | 0.1603 | 0.1849 | 0.0246 |
| 3k0z | opls | 1.91 | 0.1002 | 2.2 | 0 | 97.91 | 0 | 0 | 0.82 | 0.78 | 0.1705 | 0.204 | 0.0335 |
| 3k0z | noschrodinger | 1.91 | 0.0038 | 0.92 | 0 | 98.3 | 0 | 0 | 1.45 | 0.88 | 0.1661 | 0.1993 | 0.0332 |
| 3k2p | opls | 2.04 | 0.0979 | 2.27 | 0.39 | 98.44 | 8.8 | 0 | 1.2 | 1.55 | 0.2284 | 0.2619 | 0.0335 |
| 3k2p | noschrodinger | 2.04 | 0.0037 | 0.9 | 0 | 98.45 | 13.3 | 0 | 2.41 | 1.88 | 0.2091 | 0.246 | 0.0369 |
| 3k2v | opls | 1.95 | 0.099 | 2.66 | 0 | 99.52 | 3.29 | 2 | 6.07 | 1.73 | 0.1558 | 0.1845 | 0.0287 |
| 3k2v | noschrodinger | 1.95 | 0.007 | 1 | 0 | 99.61 | 3.42 | 0 | 3.14 | 1.51 | 0.1623 | 0.1862 | 0.0239 |
| 3k4y | opls | 2.54 | 0.0933 | 2.32 | 0.2 | 96.4 | 2.25 | 1 | 0.83 | 1.26 | 0.2064 | 0.2341 | 0.0277 |
| 3k4y | noschrodinger | 2.54 | 0.003 | 0.77 | 1 | 92.83 | 2.91 | 0 | 5.47 | 2.1 | 0.1938 | 0.2312 | 0.0374 |
| 3k65 | opls | 1.85 | 0.0976 | 2.54 | 0 | 96.39 | 2.69 | 8 | 2.95 | 1.65 | 0.1773 | 0.2175 | 0.0402 |
| 3k65 | noschrodinger | 1.85 | 0.013 | 1.21 | 0 | 95.06 | 2.65 | 0 | 3.68 | 1.82 | 0.1822 | 0.2194 | 0.0372 |
| 3k93 | opls | 2.15 | 0.094 | 2.2 | 0 | 98.34 | 0 | 0 | 1.92 | 0.96 | 0.184 | 0.2134 | 0.0294 |
| 3k93 | noschrodinger | 2.15 | 0.0028 | 0.73 | 0 | 98.64 | 1.04 | 0 | 0.28 | 0.62 | 0.1799 | 0.2077 | 0.0278 |
| 3kac | opls | 2 | 0.1017 | 2.36 | 0 | 99.55 | 1.03 | 1 | 2.23 | 1.01 | 0.1604 | 0.2167 | 0.0563 |
| 3kac | noschrodinger | 2 | 0.0037 | 0.84 | 0 | 99.55 | 1.03 | 0 | 1.13 | 0.83 | 0.1641 | 0.2151 | 0.051 |
| 3kas | opls | 2.39 | 0.0933 | 2.32 | 0.26 | 95.76 | 2.36 | 4 | 1.02 | 1.38 | 0.198 | 0.2312 | 0.0332 |
| 3kas | noschrodinger | 2.39 | 0.0031 | 0.77 | 0.13 | 95.4 | 3.07 | 0 | 3.6 | 1.84 | 0.1902 | 0.2228 | 0.0326 |
| 3kb2 | opls | 2.2 | 0.096 | 2.4 | 0.62 | 95.98 | 2 | 2 | 2.12 | 1.49 | 0.1895 | 0.2353 | 0.0458 |
| 3kb2 | noschrodinger | 2.2 | 0.0025 | 0.72 | 0.3 | 95.18 | 2.62 | 0 | 1.06 | 1.46 | 0.2028 | 0.2393 | 0.0365 |
| 3keo | opls | 1.5 | 0.1041 | 2.58 | 0 | 98.76 | 0.84 | 4 | 6.01 | 1.33 | 0.1874 | 0.2148 | 0.0274 |
| 3keo | noschrodinger | 1.5 | 0.0133 | 1.3 | 0 | 98.28 | 0.83 | 0 | 3.82 | 1.17 | 0.1879 | 0.2119 | 0.024 |
| 3kgy | opls | 1.5 | 0.1007 | 2.33 | 0 | 99.22 | 0.27 | 0 | 4.5 | 1.23 | 0.1437 | 0.1692 | 0.0255 |
| 3kgy | noschrodinger | 1.5 | 0.0075 | 1.2 | 0 | 99.31 | 0.26 | 0 | 4.87 | 1.25 | 0.1449 | 0.171 | 0.0261 |
| 3kh5 | opls | 2.1 | 0.096 | 2.39 | 0.36 | 97.83 | 3.64 | 2 | 1.49 | 1.36 | 0.2109 | 0.2365 | 0.0256 |
| 3kh5 | noschrodinger | 2.1 | 0.0042 | 0.95 | 0.36 | 96.38 | 6.88 | 0 | 4.89 | 2.13 | 0.1965 | 0.2267 | 0.0302 |
| 3ki6 | opls | 1.54 | 0.1047 | 2.53 | 0 | 97.91 | 1.26 | 1 | 3.51 | 1.24 | 0.164 | 0.1936 | 0.0296 |
| 3ki6 | noschrodinger | 1.54 | 0.0079 | 1.04 | 0 | 97.94 | 1.24 | 0 | 0.96 | 0.87 | 0.1693 | 0.1917 | 0.0224 |
| 3kjf | opls | 2 | 0.0971 | 2.28 | 0 | 97.84 | 1.89 | 0 | 0.76 | 0.99 | 0.1654 | 0.1967 | 0.0313 |
| 3kjf | noschrodinger | 2 | 0.0042 | 0.9 | 0 | 97.06 | 1.86 | 0 | 1.27 | 1.22 | 0.1706 | 0.1931 | 0.0225 |
| 3kp2 | opls | 2.55 | 0.0946 | 2.41 | 0.38 | 98.11 | 4.84 | 0 | 1.94 | 1.48 | 0.2392 | 0.2753 | 0.0361 |
| 3kp2 | noschrodinger | 2.55 | 0.0029 | 0.94 | 2.26 | 95.86 | 8.4 | 0 | 3.03 | 2.08 | 0.2253 | 0.2749 | 0.0496 |
| 3kp6 | opls | 2.45 | 0.0941 | 2.3 | 0 | 96.99 | 3.2 | 1 | 1.7 | 1.48 | 0.2396 | 0.2868 | 0.0472 |
| 3kp6 | noschrodinger | 2.45 | 0.0027 | 0.67 | 0 | 95.15 | 6.75 | 0 | 4.47 | 2.19 | 0.2228 | 0.2627 | 0.0399 |
| 3kp9 | opls | 3.6 | 0.0903 | 2.4 | 0.4 | 95.2 | 6.93 | 1 | 1.24 | 1.82 | 0.2717 | 0.307 | 0.0353 |
| 3kp9 | noschrodinger | 3.6 | 0.0032 | 0.87 | 1.98 | 85.38 | 15.76 | 0 | 8.7 | 3.03 | 0.2767 | 0.309 | 0.0323 |
| 3kpc | opls | 1.79 | 0.1031 | 2.44 | 0 | 100 | 2.7 | 1 | 6.72 | 1.7 | 0.1628 | 0.1988 | 0.036 |
| 3kpc | noschrodinger | 1.79 | 0.0043 | 1.07 | 0 | 100 | 2.68 | 0 | 6.73 | 1.7 | 0.1677 | 0.1964 | 0.0287 |
| 3kqs | opls | 2 | 0.0988 | 2.28 | 0 | 98.07 | 0.23 | 0 | 0.97 | 0.79 | 0.2015 | 0.2255 | 0.024 |
| 3kqs | noschrodinger | 2 | 0.0027 | 0.83 | 0 | 98.46 | 0.7 | 0 | 1.58 | 0.9 | 0.1951 | 0.2259 | 0.0308 |
| 3ksm | opls | 1.9 | 0.0978 | 2.28 | 0 | 97.29 | 0.47 | 1 | 1.53 | 1.03 | 0.144 | 0.176 | 0.032 |

| | | | | | | | | | | | | | |
|---|---|---|---|---|---|---|---|---|---|---|---|---|---|
| 3ksm | noschrodinger | 1.9 | 0.0047 | 0.96 | 0 | 97.06 | 1.14 | 0 | 2.7 | 1.27 | 0.1467 | 0.1746 | 0.0279 |
| 3ksp | opls | 2.6 | 0.0878 | 2.23 | 0 | 98.17 | 1.79 | 0 | 1.4 | 1.06 | 0.2508 | 0.2677 | 0.0169 |
| 3ksp | noschrodinger | 2.6 | 0.0028 | 0.79 | 0 | 95.28 | 3.39 | 0 | 1.41 | 1.61 | 0.2413 | 0.2642 | 0.0229 |
| 3ku3 | opls | 1.5 | 0.1024 | 2.54 | 0.2 | 97.75 | 2.06 | 4 | 3.78 | 1.46 | 0.191 | 0.2262 | 0.0352 |
| 3ku3 | noschrodinger | 1.5 | 0.0097 | 1.09 | 0 | 97.36 | 2.28 | 0 | 2.39 | 1.42 | 0.1956 | 0.2252 | 0.0296 |
| 3kw2 | opls | 2 | 0.0931 | 2.25 | 0 | 95.91 | 2.02 | 1 | 0.78 | 1.26 | 0.2043 | 0.2377 | 0.0334 |
| 3kw2 | noschrodinger | 2 | 0.0028 | 0.77 | 0 | 96.38 | 4.18 | 0 | 1.43 | 1.59 | 0.1996 | 0.2219 | 0.0223 |
| 3kwt | opls | 1.89 | 0.0946 | 2.41 | 0.96 | 97.12 | 1.96 | 1 | 2.1 | 1.36 | 0.206 | 0.2425 | 0.0365 |
| 3kwt | noschrodinger | 1.89 | 0.0112 | 1.12 | 1.85 | 96.3 | 4.72 | 0 | 2.64 | 1.81 | 0.2105 | 0.2401 | 0.0296 |
| 3kyf | opls | 2.09 | 0.0981 | 2.32 | 0 | 99.56 | 1.01 | 1 | 0.52 | 0.68 | 0.2007 | 0.2439 | 0.0432 |
| 3kyf | noschrodinger | 2.09 | 0.0025 | 0.72 | 0.44 | 98.25 | 2 | 0 | 2.08 | 1.21 | 0.1937 | 0.241 | 0.0473 |
| 3kyg | opls | 2.1 | 0.097 | 2.28 | 0.45 | 95.98 | 2.5 | 0 | 0.66 | 1.3 | 0.2091 | 0.2323 | 0.0232 |
| 3kyg | noschrodinger | 2.1 | 0.0041 | 0.81 | 0.22 | 96.89 | 2.99 | 0 | 3.06 | 1.64 | 0.194 | 0.2225 | 0.0285 |
| 3kyq | opls | 2.44 | 0.0951 | 2.25 | 0 | 98.41 | 4.71 | 0 | 0 | 1.01 | 0.2015 | 0.2316 | 0.0301 |
| 3kyq | noschrodinger | 2.44 | 0.0029 | 0.77 | 0 | 96.86 | 5.81 | 0 | 0.63 | 1.48 | 0.1812 | 0.2266 | 0.0454 |
| 3l0a | opls | 2.19 | 0.0917 | 2.29 | 0 | 98.06 | 2.82 | 2 | 1.85 | 1.29 | 0.2212 | 0.2499 | 0.0287 |
| 3l0a | noschrodinger | 2.19 | 0.003 | 0.83 | 0 | 98.11 | 3.54 | 0 | 1.4 | 1.29 | 0.2018 | 0.239 | 0.0372 |
| 3l2b | opls | 2.27 | 0.0919 | 2.26 | 0 | 98.67 | 1.73 | 1 | 0.82 | 0.94 | 0.2179 | 0.2509 | 0.033 |
| 3l2b | noschrodinger | 2.27 | 0.0029 | 0.75 | 0.22 | 98.04 | 2.46 | 0 | 1.92 | 1.25 | 0.2162 | 0.244 | 0.0278 |
| 3l5h | opls | 3.6 | 0.0864 | 3 | 5.25 | 82.58 | 19.22 | 15 | 5.01 | 2.94 | 0.2697 | 0.314 | 0.0443 |
| 3l5h | noschrodinger | 3.6 | 0.0058 | 1.18 | 16 | 65.74 | 30.77 | 0 | 39.12 | 4.08 | 0.2519 | 0.2857 | 0.0338 |
| 3l5z | opls | 2.9 | 0.0913 | 2.29 | 0 | 98.44 | 4.84 | 0 | 0.43 | 1.17 | 0.2061 | 0.2246 | 0.0185 |
| 3l5z | noschrodinger | 2.9 | 0.0028 | 0.76 | 0 | 98.23 | 4.65 | 0 | 1.72 | 1.43 | 0.2023 | 0.2222 | 0.0199 |
| 3l6g | opls | 1.9 | 0.1019 | 2.16 | 0.4 | 98.81 | 0.46 | 1 | 0.73 | 0.73 | 0.1646 | 0.2063 | 0.0417 |
| 3l6g | noschrodinger | 1.9 | 0.0038 | 0.84 | 0.39 | 98.82 | 0.91 | 0 | 1.71 | 0.93 | 0.1621 | 0.2005 | 0.0384 |
| 3l86 | opls | 2.06 | 0.0937 | 2.34 | 0.94 | 97.18 | 2.13 | 0 | 0.53 | 1.08 | 0.2088 | 0.2416 | 0.0328 |
| 3l86 | noschrodinger | 2.06 | 0.0043 | 1.02 | 0.82 | 97.53 | 3.17 | 0 | 2.69 | 1.53 | 0.2012 | 0.2451 | 0.0439 |
| 3l8g | opls | 2.18 | 0.0937 | 2.31 | 0 | 97.48 | 1.38 | 0 | 0.7 | 0.94 | 0.21 | 0.2421 | 0.0321 |
| 3l8g | noschrodinger | 2.18 | 0.0034 | 0.8 | 0 | 96.13 | 2.61 | 0 | 2.8 | 1.65 | 0.1837 | 0.2237 | 0.04 |
| 3lac | opls | 2 | 0.0965 | 2.26 | 0 | 97.03 | 0.3 | 3 | 1.22 | 1.01 | 0.1917 | 0.2208 | 0.0291 |
| 3lac | noschrodinger | 2 | 0.0031 | 0.86 | 0 | 96.98 | 1.72 | 0 | 3.21 | 1.47 | 0.1911 | 0.2179 | 0.0268 |
| 3lat | opls | 1.7 | 0.1039 | 2.41 | 0 | 97.8 | 0.55 | 4 | 2.54 | 1.08 | 0.147 | 0.1755 | 0.0285 |
| 3lat | noschrodinger | 1.7 | 0.0141 | 1.36 | 0 | 97.8 | 0.83 | 0 | 0.9 | 0.82 | 0.1495 | 0.1746 | 0.0251 |
| 3lbg | opls | 1.5 | 0.102 | 2.83 | 0.34 | 98.29 | 1.86 | 3 | 1.86 | 1.15 | 0.1551 | 0.174 | 0.0189 |
| 3lbg | noschrodinger | 1.5 | 0.007 | 2.04 | 0.34 | 97.95 | 1.12 | 0 | 0.82 | 0.8 | 0.1666 | 0.182 | 0.0154 |
| 3lea | opls | 2 | 0.0985 | 2.13 | 0.43 | 97.41 | 0.47 | 0 | 0.87 | 0.88 | 0.1816 | 0.206 | 0.0244 |
| 3lea | noschrodinger | 2 | 0.003 | 0.85 | 0.39 | 97.48 | 0 | 0 | 1.38 | 0.97 | 0.1717 | 0.1993 | 0.0276 |
| 3leg | opls | 2.01 | 0.099 | 2.16 | 0 | 97.12 | 0.85 | 0 | 1.31 | 1.01 | 0.1926 | 0.213 | 0.0204 |
| 3leg | noschrodinger | 2.01 | 0.0024 | 0.75 | 0 | 96.43 | 2.54 | 0 | 0 | 1.04 | 0.1848 | 0.2086 | 0.0238 |
| 3lev | opls | 2.5 | 0.0921 | 2.3 | 0.32 | 93.69 | 2.62 | 1 | 1.43 | 1.61 | 0.2352 | 0.2739 | 0.0387 |
| 3lev | noschrodinger | 2.5 | 0.0029 | 0.96 | 1.13 | 90.03 | 2.98 | 0 | 3.67 | 2.07 | 0.2261 | 0.2615 | 0.0354 |
| 3lff | opls | 1.5 | 0.0976 | 2.53 | 0 | 97.44 | 2.35 | 7 | 4.78 | 1.64 | 0.1673 | 0.1993 | 0.032 |
| 3lff | noschrodinger | 1.5 | 0.0038 | 0.97 | 0 | 98.19 | 1.67 | 0 | 2.84 | 1.24 | 0.1768 | 0.1962 | 0.0194 |
| 3lfr | opls | 1.53 | 0.0963 | 2.35 | 0 | 98.49 | 1.44 | 1 | 1.71 | 1.05 | 0.1763 | 0.2077 | 0.0314 |
| 3lfr | noschrodinger | 1.53 | 0.0064 | 1.02 | 0 | 99.18 | 1.39 | 0 | 1.97 | 1.07 | 0.1777 | 0.2032 | 0.0255 |
| 3lhh | opls | 2.1 | 0.0912 | 2.28 | 0 | 96.12 | 8.93 | 0 | 0.48 | 1.65 | 0.2239 | 0.2632 | 0.0393 |
| 3lhh | noschrodinger | 2.1 | 0.0031 | 0.79 | 0 | 97.56 | 10.74 | 0 | 1.44 | 1.75 | 0.2107 | 0.2573 | 0.0466 |
| 3lho | opls | 1.8 | 0.0979 | 2.28 | 0.42 | 97.48 | 0 | 0 | 3.13 | 1.21 | 0.1657 | 0.2033 | 0.0376 |
| 3lho | noschrodinger | 1.8 | 0.0137 | 1.1 | 0.4 | 96.83 | 0.46 | 0 | 1.45 | 1.08 | 0.1669 | 0.2072 | 0.0403 |

| | | | | | | | | | | | | | |
|---|---|---|---|---|---|---|---|---|---|---|---|---|---|
| 3lhq | opls | 1.56 | 0.0982 | 2.49 | 0 | 100 | 2.18 | 2 | 5.99 | 1.59 | 0.1688 | 0.2082 | 0.0394 |
| 3lhq | noschrodinger | 1.56 | 0.0122 | 1.22 | 0 | 99.76 | 2.42 | 0 | 5.7 | 1.6 | 0.1695 | 0.2075 | 0.038 |
| 3li2 | opls | 2.2 | 0.095 | 2.3 | 1.4 | 95.8 | 4.33 | 6 | 1.25 | 1.62 | 0.2215 | 0.2602 | 0.0387 |
| 3li2 | noschrodinger | 2.2 | 0.0031 | 0.82 | 1.4 | 95.64 | 8.82 | 0 | 3.53 | 2.17 | 0.2173 | 0.2535 | 0.0362 |
| 3ljy | opls | 2.4 | 0.0906 | 2.19 | 0 | 96.84 | 1.74 | 0 | 0.64 | 1.08 | 0.2204 | 0.2589 | 0.0385 |
| 3ljy | noschrodinger | 2.4 | 0.004 | 0.81 | 0.42 | 95.49 | 2.57 | 0 | 1.93 | 1.58 | 0.2114 | 0.2532 | 0.0418 |
| 3lkk | opls | 2 | 0.0952 | 2.28 | 0 | 98.47 | 1.01 | 2 | 0.8 | 0.75 | 0.1866 | 0.2048 | 0.0182 |
| 3lkk | noschrodinger | 2 | 0.0031 | 0.87 | 0.22 | 98.7 | 2.24 | 0 | 2.55 | 1.31 | 0.1769 | 0.1982 | 0.0213 |
| 3ll4 | opls | 2.49 | 0.0922 | 2.32 | 0.39 | 98.45 | 2.19 | 0 | 1.42 | 1.13 | 0.21 | 0.2551 | 0.0451 |
| 3ll4 | noschrodinger | 2.49 | 0.0043 | 0.85 | 0.39 | 97.68 | 4.15 | 0 | 1.65 | 1.45 | 0.2135 | 0.2531 | 0.0396 |
| 3llk | opls | 2 | 0.0972 | 2.31 | 0 | 97.85 | 2.98 | 5 | 1.68 | 1.31 | 0.2254 | 0.2576 | 0.0322 |
| 3llk | noschrodinger | 2 | 0.0028 | 0.8 | 0.28 | 95.23 | 2.79 | 0 | 3.7 | 1.83 | 0.2229 | 0.2518 | 0.0289 |
| 3llm | opls | 2.8 | 0.0889 | 2.26 | 0 | 97.3 | 0 | 2 | 0.4 | 0.78 | 0.2046 | 0.2252 | 0.0206 |
| 3llm | noschrodinger | 2.8 | 0.0047 | 0.91 | 0 | 95.71 | 2.92 | 0 | 1.08 | 1.46 | 0.2106 | 0.2313 | 0.0207 |
| 3loq | opls | 2.32 | 0.0904 | 2.35 | 0.41 | 97.72 | 4.54 | 3 | 0.35 | 1.19 | 0.2134 | 0.23 | 0.0166 |
| 3loq | noschrodinger | 2.32 | 0.0029 | 0.82 | 0.38 | 96.42 | 6.97 | 0 | 1.52 | 1.77 | 0.1955 | 0.2169 | 0.0214 |
| 3lp3 | opls | 2.8 | 0.0904 | 2.24 | 0 | 98.05 | 8.8 | 0 | 1.67 | 1.64 | 0.2075 | 0.2538 | 0.0463 |
| 3lp3 | noschrodinger | 2.8 | 0.0046 | 0.87 | 0.39 | 98.06 | 11.01 | 0 | 3.34 | 1.92 | 0.2093 | 0.2441 | 0.0348 |
| 3lpw | opls | 1.65 | 0.108 | 2.22 | 0 | 97.93 | 0.29 | 2 | 1.93 | 0.97 | 0.1799 | 0.2083 | 0.0284 |
| 3lpw | noschrodinger | 1.65 | 0.0068 | 0.91 | 0 | 97.42 | 0.29 | 0 | 2.89 | 1.19 | 0.178 | 0.2003 | 0.0223 |
| 3lqq | opls | 3.53 | 0.0904 | 2.47 | 1.93 | 91.26 | 11.04 | 14 | 2.58 | 2.35 | 0.2838 | 0.3199 | 0.0361 |
| 3lqq | noschrodinger | 3.53 | 0.0033 | 0.88 | 3.02 | 82.16 | 17.83 | 0 | 6.52 | 3.02 | 0.2682 | 0.3063 | 0.0381 |
| 3lqu | opls | 2.52 | 0.0945 | 2.35 | 0 | 98.73 | 3.19 | 0 | 1.33 | 1.24 | 0.2168 | 0.2587 | 0.0419 |
| 3lqu | noschrodinger | 2.52 | 0.0029 | 0.78 | 0.25 | 95.72 | 5.38 | 0 | 2.83 | 1.92 | 0.1926 | 0.2436 | 0.051 |
| 3lrt | opls | 1.53 | 0.0992 | 2.39 | 0.18 | 95.91 | 0.99 | 8 | 1.94 | 1.24 | 0.1681 | 0.2077 | 0.0396 |
| 3lrt | noschrodinger | 1.53 | 0.0075 | 1.06 | 0.35 | 95.76 | 1.38 | 0 | 2.38 | 1.42 | 0.169 | 0.2031 | 0.0341 |
| 3lsj | opls | 2.3 | 0.0948 | 2.46 | 0 | 98.72 | 3.81 | 2 | 1.67 | 1.36 | 0.2065 | 0.2533 | 0.0468 |
| 3lsj | noschrodinger | 2.3 | 0.0045 | 0.96 | 0 | 98.5 | 5.02 | 0 | 3.35 | 1.66 | 0.2197 | 0.2491 | 0.0294 |
| 3ltm | opls | 2.15 | 0.1001 | 2.37 | 0 | 99.17 | 5.49 | 9 | 1.02 | 1.36 | 0.2096 | 0.2376 | 0.028 |
| 3ltm | noschrodinger | 2.15 | 0.0029 | 0.75 | 0 | 98.62 | 5.82 | 0 | 2.56 | 1.62 | 0.183 | 0.2267 | 0.0437 |
| 3luh | opls | 2 | 0.1004 | 2.31 | 0 | 100 | 1.16 | 2 | 0.78 | 0.8 | 0.1699 | 0.2114 | 0.0415 |
| 3luh | noschrodinger | 2 | 0.0027 | 0.78 | 0 | 98.98 | 1.74 | 0 | 0.94 | 0.97 | 0.1719 | 0.2113 | 0.0394 |
| 3lur | opls | 1.82 | 0.0952 | 2.22 | 0 | 99.21 | 0.76 | 3 | 1.89 | 0.95 | 0.1646 | 0.1932 | 0.0286 |
| 3lur | noschrodinger | 1.82 | 0.0087 | 1 | 0 | 98.92 | 0.48 | 0 | 3.15 | 1.11 | 0.1634 | 0.1928 | 0.0294 |
| 3lw7 | opls | 2.3 | 0.0919 | 2.3 | 0 | 97.35 | 4.03 | 1 | 0.55 | 1.27 | 0.2127 | 0.2388 | 0.0261 |
| 3lw7 | noschrodinger | 2.3 | 0.0028 | 0.77 | 0 | 97.58 | 6.5 | 0 | 1.49 | 1.59 | 0.2039 | 0.2426 | 0.0387 |
| 3lx3 | opls | 1.55 | 0.1017 | 2.44 | 0.62 | 95.65 | 1.81 | 2 | 2.48 | 1.53 | 0.1568 | 0.1754 | 0.0186 |
| 3lx3 | noschrodinger | 1.55 | 0.0094 | 1.14 | 0.62 | 95.65 | 1.81 | 0 | 2.12 | 1.48 | 0.1579 | 0.1742 | 0.0163 |
| 3lxr | opls | 1.68 | 0.1045 | 2.33 | 0 | 98.03 | 1.18 | 1 | 1.5 | 0.95 | 0.1663 | 0.211 | 0.0447 |
| 3lxr | noschrodinger | 1.68 | 0.0041 | 0.87 | 0 | 97.48 | 2.07 | 0 | 3.32 | 1.47 | 0.1711 | 0.203 | 0.0319 |
| 3lxs | opls | 1.5 | 0.1009 | 2.26 | 0 | 97.41 | 0 | 0 | 3.21 | 1.23 | 0.1161 | 0.1287 | 0.0126 |
| 3lxs | noschrodinger | 1.5 | 0.0061 | 1.15 | 0 | 97.42 | 0 | 0 | 0.92 | 0.89 | 0.1112 | 0.1424 | 0.0312 |
| 3lzz | opls | 2.5 | 0.0963 | 2.37 | 0 | 99.03 | 0.36 | 0 | 1.73 | 0.93 | 0.1983 | 0.2112 | 0.0129 |
| 3lzz | noschrodinger | 2.5 | 0.0044 | 1 | 0 | 98.11 | 0.36 | 0 | 4.45 | 1.22 | 0.1841 | 0.2053 | 0.0212 |
| 3m1i | opls | 2 | 0.0973 | 2.26 | 0 | 97.72 | 2.77 | 10 | 0.78 | 1.14 | 0.2043 | 0.2173 | 0.013 |
| 3m1i | noschrodinger | 2 | 0.0029 | 0.8 | 0.37 | 97.1 | 4.95 | 0 | 1.66 | 1.6 | 0.1935 | 0.2121 | 0.0186 |
| 3m2k | opls | 3.5 | 0.0914 | 2.43 | 1.03 | 95.88 | 10.17 | 7 | 1.02 | 1.85 | 0.1974 | 0.2284 | 0.031 |
| 3m2k | noschrodinger | 3.5 | 0.0037 | 0.98 | 2.25 | 89.75 | 19.23 | 0 | 5.72 | 2.84 | 0.1907 | 0.2308 | 0.0401 |
| 3m3c | opls | 2.01 | 0.0997 | 2.3 | 0.32 | 96.47 | 2.29 | 1 | 2.24 | 1.51 | 0.1494 | 0.1883 | 0.0389 |

| | | | | | | | | | | | | | |
|---|---|---|---|---|---|---|---|---|---|---|---|---|---|
| 3m3c | noschrodinger | 2.01 | 0.0031 | 0.89 | 0 | 96.5 | 2.65 | 0 | 3.67 | 1.71 | 0.1653 | 0.1951 | 0.0298 |
| 3m93 | opls | 2.9 | 0.0865 | 2.31 | 0 | 97.9 | 5.92 | 1 | 1.3 | 1.46 | 0.2317 | 0.261 | 0.0293 |
| 3m93 | noschrodinger | 2.9 | 0.0028 | 0.79 | 0.53 | 96.26 | 8.97 | 0 | 2.98 | 2.06 | 0.2246 | 0.2627 | 0.0381 |
| 3maw | opls | 3.51 | 0.0877 | 2.29 | 0 | 95.32 | 1.06 | 4 | 0.89 | 1.12 | 0.2655 | 0.2953 | 0.0298 |
| 3maw | noschrodinger | 3.51 | 0.0028 | 0.77 | 0.12 | 90.22 | 0 | 0 | 1.64 | 1.46 | 0.272 | 0.3051 | 0.0331 |
| 3mb3 | opls | 2.25 | 0.0979 | 2.47 | 0 | 100 | 2.78 | 1 | 0 | 0.84 | 0.2263 | 0.2468 | 0.0205 |
| 3mb3 | noschrodinger | 2.25 | 0.003 | 0.76 | 0 | 99.12 | 3.7 | 0 | 0.54 | 1.11 | 0.2172 | 0.2462 | 0.029 |
| 3mbz | opls | 2.6 | 0.0955 | 2.25 | 0 | 97.48 | 4.78 | 0 | 1.25 | 1.47 | 0.1851 | 0.2154 | 0.0303 |
| 3mbz | noschrodinger | 2.6 | 0.0028 | 0.95 | 0.42 | 95.82 | 7.14 | 0 | 2.25 | 1.94 | 0.1684 | 0.2023 | 0.0339 |
| 3mct | opls | 1.87 | 0.0959 | 2.31 | 0 | 99.65 | 4.53 | 3 | 1.02 | 1.3 | 0.197 | 0.2425 | 0.0455 |
| 3mct | noschrodinger | 1.87 | 0.0034 | 0.91 | 0 | 98.95 | 5.99 | 0 | 3.48 | 1.73 | 0.2069 | 0.2386 | 0.0317 |
| 3mdp | opls | 1.9 | 0.0994 | 2.34 | 0 | 98.11 | 0 | 0 | 4.55 | 1.23 | 0.1732 | 0.2119 | 0.0387 |
| 3mdp | noschrodinger | 1.9 | 0.0031 | 0.8 | 0 | 96.88 | 1.74 | 0 | 2.29 | 1.38 | 0.1878 | 0.2095 | 0.0217 |
| 3mgd | opls | 1.9 | 0.0991 | 2.48 | 0.36 | 97.49 | 2.64 | 0 | 1.51 | 1.32 | 0.1819 | 0.2227 | 0.0408 |
| 3mgd | noschrodinger | 1.9 | 0.003 | 0.81 | 0 | 97.69 | 4.33 | 0 | 2.64 | 1.6 | 0.1786 | 0.2137 | 0.0351 |
| 3mh6 | opls | 3.6 | 0.0908 | 2.35 | 0.78 | 94.82 | 2.89 | 1 | 1.02 | 1.51 | 0.2605 | 0.2944 | 0.0339 |
| 3mh6 | noschrodinger | 3.6 | 0.0029 | 0.77 | 2.32 | 83.51 | 7.99 | 0 | 6.33 | 2.72 | 0.2279 | 0.2783 | 0.0504 |
| 3mhp | opls | 1.7 | 0.0991 | 2.28 | 0 | 98.83 | 1.33 | 0 | 2.73 | 1.15 | 0.1454 | 0.1777 | 0.0323 |
| 3mhp | noschrodinger | 1.7 | 0.0061 | 0.96 | 0 | 99.34 | 1.69 | 0 | 2.12 | 1.16 | 0.1483 | 0.1747 | 0.0264 |
| 3mj6 | opls | 2.19 | 0.0963 | 2.3 | 0 | 95.54 | 2.79 | 2 | 1.04 | 1.45 | 0.1926 | 0.2212 | 0.0286 |
| 3mj6 | noschrodinger | 2.19 | 0.0043 | 1.68 | 0.44 | 96.89 | 6.02 | 0 | 1.83 | 1.72 | 0.1699 | 0.2154 | 0.0455 |
| 3mj7 | opls | 2.8 | 0.089 | 2.3 | 0.5 | 94.29 | 2.11 | 1 | 0.44 | 1.29 | 0.2674 | 0.3048 | 0.0374 |
| 3mj7 | noschrodinger | 2.8 | 0.0028 | 0.76 | 0.25 | 91.18 | 4.42 | 0 | 3.22 | 2.12 | 0.2946 | 0.3179 | 0.0233 |
| 3ml1 | opls | 1.5 | 0.0982 | 2.33 | 0.11 | 96.18 | 0.78 | 2 | 2.38 | 1.28 | 0.17 | 0.1881 | 0.0181 |
| 3ml1 | noschrodinger | 1.5 | 0.0144 | 1.32 | 0.11 | 96.56 | 1.3 | 0 | 1.29 | 1.16 | 0.1698 | 0.1886 | 0.0188 |
| 3mql | opls | 3 | 0.0888 | 2.76 | 0.49 | 92.2 | 1.69 | 8 | 4.39 | 1.87 | 0.2108 | 0.2572 | 0.0464 |
| 3mql | noschrodinger | 3 | 0.0027 | 0.73 | 0.48 | 94.2 | 1.68 | 0 | 3.46 | 1.7 | 0.2353 | 0.2521 | 0.0168 |
| 3msw | opls | 1.9 | 0.0978 | 2.18 | 1.72 | 97.41 | 3.2 | 3 | 1.69 | 1.42 | 0.2043 | 0.2499 | 0.0456 |
| 3msw | noschrodinger | 1.9 | 0.0039 | 0.78 | 0.73 | 94.89 | 3.05 | 0 | 0.85 | 1.48 | 0.2112 | 0.2515 | 0.0403 |
| 3mtx | opls | 2 | 0.0927 | 2.31 | 0 | 97.05 | 0.43 | 3 | 0.87 | 0.93 | 0.2136 | 0.2577 | 0.0441 |
| 3mtx | noschrodinger | 2 | 0.0037 | 0.85 | 0 | 96.35 | 1.72 | 0 | 2.21 | 1.42 | 0.2192 | 0.2574 | 0.0382 |
| 3mwn | opls | 2.6 | 0.0896 | 2.74 | 2.14 | 88.03 | 11.34 | 8 | 3.5 | 2.54 | 0.3201 | 0.3547 | 0.0346 |
| 3mwn | noschrodinger | 2.6 | 0.004 | 0.96 | 2.67 | 86.26 | 19.9 | 0 | 10.81 | 3.17 | 0.3067 | 0.342 | 0.0353 |
| 3mwy | opls | 3.1 | 0.0906 | 2.44 | 1.19 | 92.59 | 12.98 | 13 | 2.36 | 2.33 | 0.294 | 0.3389 | 0.0449 |
| 3mwy | noschrodinger | 3.1 | 0.0039 | 0.92 | 5.87 | 75.81 | 25.83 | 0 | 16.13 | 3.57 | 0.2888 | 0.3255 | 0.0367 |
| 3mxn | opls | 1.55 | 0.1004 | 2.32 | 0 | 97.83 | 1.15 | 2 | 3.02 | 1.18 | 0.1678 | 0.1967 | 0.0289 |
| 3mxn | noschrodinger | 1.55 | 0.0046 | 1 | 0 | 97.48 | 1.52 | 0 | 3.66 | 1.4 | 0.1719 | 0.1944 | 0.0225 |
| 3mxo | opls | 1.7 | 0.0981 | 2.33 | 0 | 98.86 | 1.48 | 0 | 3.04 | 1.23 | 0.1649 | 0.1932 | 0.0283 |
| 3mxo | noschrodinger | 1.7 | 0.0041 | 0.87 | 0 | 99.19 | 1.76 | 0 | 2.74 | 1.25 | 0.168 | 0.1923 | 0.0243 |
| 3my1 | opls | 2.8 | 0.0915 | 2.28 | 0.72 | 95.5 | 1.19 | 1 | 0.21 | 0.95 | 0.2103 | 0.2249 | 0.0146 |
| 3my1 | noschrodinger | 2.8 | 0.0054 | 0.95 | 0.9 | 90.14 | 2.95 | 0 | 7.69 | 2.32 | 0.1992 | 0.2175 | 0.0183 |
| 3mzo | opls | 1.98 | 0.0955 | 2.28 | 0.19 | 97.96 | 0.93 | 9 | 1.38 | 0.88 | 0.1678 | 0.2065 | 0.0387 |
| 3mzo | noschrodinger | 1.98 | 0.0049 | 0.83 | 0.16 | 97.7 | 1.63 | 0 | 0.79 | 0.98 | 0.1705 | 0.2007 | 0.0302 |
| 3n0n | opls | 1.5 | 0.1023 | 2.21 | 0 | 97.27 | 0.43 | 0 | 2.32 | 1.15 | 0.1373 | 0.1559 | 0.0186 |
| 3n0n | noschrodinger | 1.5 | 0.0062 | 1.04 | 0 | 97.66 | 0.43 | 0 | 1.16 | 0.9 | 0.1374 | 0.1534 | 0.016 |
| 3n10 | opls | 1.6 | 0.0964 | 2.37 | 0 | 99.71 | 2.98 | 2 | 1.23 | 1.2 | 0.1944 | 0.2317 | 0.0373 |
| 3n10 | noschrodinger | 1.6 | 0.01 | 1.27 | 0 | 99.43 | 3.64 | 0 | 2.65 | 1.48 | 0.1961 | 0.2342 | 0.0381 |
| 3n2v | opls | 1.55 | 0.107 | 2.21 | 0 | 97.42 | 1.57 | 2 | 2.04 | 1.24 | 0.1657 | 0.1994 | 0.0337 |
| 3n2v | noschrodinger | 1.55 | 0.004 | 0.92 | 0 | 97.44 | 0.79 | 0 | 2.04 | 1.09 | 0.1652 | 0.1955 | 0.0303 |

| | | | | | | | | | | | | | |
|---|---|---|---|---|---|---|---|---|---|---|---|---|---|
| 3n35 | opls | 2 | 0.0975 | 2.18 | 0 | 99.16 | 0.97 | 0 | 1.08 | 0.81 | 0.1388 | 0.1644 | 0.0256 |
| 3n35 | noschrodinger | 2 | 0.0034 | 0.91 | 0 | 99.17 | 0.48 | 0 | 0.81 | 0.75 | 0.1418 | 0.164 | 0.0222 |
| 3n5b | opls | 1.9 | 0.098 | 2.35 | 0 | 97.91 | 0.58 | 3 | 2.13 | 1.01 | 0.1994 | 0.2317 | 0.0323 |
| 3n5b | noschrodinger | 1.9 | 0.0114 | 1.1 | 0 | 97.94 | 0.57 | 0 | 3.65 | 1.17 | 0.1986 | 0.2306 | 0.032 |
| 3n9t | opls | 2 | 0.0936 | 2.23 | 0 | 97.17 | 0.83 | 0 | 1.54 | 1.05 | 0.1797 | 0.2102 | 0.0305 |
| 3n9t | noschrodinger | 2 | 0.006 | 1.3 | 0 | 96.13 | 2.49 | 0 | 7.73 | 1.99 | 0.1773 | 0.2087 | 0.0314 |
| 3nbe | opls | 1.86 | 0.1036 | 2.34 | 0.35 | 96.5 | 0.77 | 1 | 1.29 | 1.08 | 0.1723 | 0.197 | 0.0247 |
| 3nbe | noschrodinger | 1.86 | 0.0062 | 1.06 | 0.69 | 97.24 | 1.53 | 0 | 3.66 | 1.44 | 0.166 | 0.1961 | 0.0301 |
| 3nhf | opls | 2 | 0.0983 | 2.28 | 0 | 96.7 | 1.02 | 6 | 2.13 | 1.2 | 0.167 | 0.199 | 0.032 |
| 3nhf | noschrodinger | 2 | 0.0035 | 0.91 | 0.22 | 94.96 | 2.8 | 0 | 2.26 | 1.69 | 0.1729 | 0.1935 | 0.0206 |
| 3nkh | opls | 2.5 | 0.0937 | 2.27 | 0 | 98.19 | 4 | 2 | 0.52 | 1.14 | 0.1858 | 0.2033 | 0.0175 |
| 3nkh | noschrodinger | 2.5 | 0.003 | 0.77 | 0 | 99.09 | 6.21 | 0 | 2.22 | 1.6 | 0.1712 | 0.1953 | 0.0241 |
| 3nku | opls | 2.1 | 0.0931 | 2.3 | 0 | 98.06 | 2.89 | 0 | 0.33 | 0.97 | 0.2029 | 0.2305 | 0.0276 |
| 3nku | noschrodinger | 2.1 | 0.0029 | 0.75 | 0.54 | 97.55 | 4.27 | 0 | 1.51 | 1.46 | 0.1884 | 0.2198 | 0.0314 |
| 3nlx | opls | 1.87 | 0.0962 | 2.27 | 0 | 98.51 | 1.09 | 8 | 2.35 | 1.04 | 0.1882 | 0.2245 | 0.0363 |
| 3nlx | noschrodinger | 1.87 | 0.003 | 0.87 | 0 | 98.77 | 1.63 | 0 | 1.4 | 1.03 | 0.1907 | 0.2188 | 0.0281 |
| 3nlz | opls | 1.92 | 0.0953 | 2.3 | 0 | 97.78 | 0.82 | 8 | 2.49 | 1.08 | 0.1829 | 0.2126 | 0.0297 |
| 3nlz | noschrodinger | 1.92 | 0.003 | 0.86 | 0 | 98.28 | 1.62 | 0 | 1.9 | 1.11 | 0.1918 | 0.2079 | 0.0161 |
| 3nm0 | opls | 1.81 | 0.0959 | 2.24 | 0 | 98.76 | 0.54 | 6 | 1.83 | 0.94 | 0.1895 | 0.2227 | 0.0332 |
| 3nm0 | noschrodinger | 1.81 | 0.0033 | 0.89 | 0 | 98.52 | 1.49 | 0 | 1.97 | 1.1 | 0.1921 | 0.2227 | 0.0306 |
| 3noj | opls | 1.82 | 0.0971 | 2.5 | 0.43 | 97.41 | 1.06 | 3 | 2.46 | 1.16 | 0.1583 | 0.1843 | 0.026 |
| 3noj | noschrodinger | 1.82 | 0.0141 | 1.33 | 0.43 | 97 | 1.05 | 0 | 1.91 | 1.15 | 0.1591 | 0.1846 | 0.0255 |
| 3nok | opls | 1.65 | 0.0974 | 2.28 | 0 | 96.75 | 1.04 | 0 | 2.44 | 1.24 | 0.1825 | 0.2117 | 0.0292 |
| 3nok | noschrodinger | 1.65 | 0.0041 | 0.89 | 0 | 96.54 | 1.56 | 0 | 2.17 | 1.36 | 0.1816 | 0.2099 | 0.0283 |
| 3npd | opls | 1.6 | 0.098 | 2.24 | 0 | 100 | 1.03 | 0 | 3.07 | 1.11 | 0.1623 | 0.1986 | 0.0363 |
| 3npd | noschrodinger | 1.6 | 0.0052 | 0.89 | 0 | 100 | 0.99 | 0 | 1.03 | 0.8 | 0.1605 | 0.1962 | 0.0357 |
| 3npk | opls | 1.5 | 0.1004 | 2.26 | 0 | 99.16 | 2.24 | 1 | 2.23 | 1.27 | 0.1713 | 0.1876 | 0.0163 |
| 3npk | noschrodinger | 1.5 | 0.0082 | 1.02 | 0.19 | 97.88 | 3.69 | 0 | 2.7 | 1.52 | 0.1679 | 0.189 | 0.0211 |
| 3nri | opls | 2.85 | 0.0901 | 2.43 | 0 | 98.53 | 4.38 | 0 | 0.42 | 1.14 | 0.2063 | 0.2599 | 0.0536 |
| 3nri | noschrodinger | 2.85 | 0.003 | 0.87 | 0.36 | 96.73 | 7.51 | 0 | 6.57 | 2.23 | 0.1877 | 0.2429 | 0.0552 |
| 3ns7 | opls | 2.6 | 0.0901 | 2.44 | 0.9 | 90.58 | 8.8 | 5 | 0.74 | 1.99 | 0.2544 | 0.2837 | 0.0293 |
| 3ns7 | noschrodinger | 2.6 | 0.0035 | 0.94 | 1.22 | 89.84 | 15.62 | 0 | 4.21 | 2.66 | 0.2478 | 0.2822 | 0.0344 |
| 3nuq | opls | 1.7 | 0.0961 | 2.25 | 0 | 96.89 | 0.4 | 0 | 3.11 | 1.29 | 0.1734 | 0.1976 | 0.0242 |
| 3nuq | noschrodinger | 1.7 | 0.0039 | 0.89 | 0.38 | 96.56 | 1.99 | 0 | 0.67 | 1.17 | 0.1741 | 0.1929 | 0.0188 |
| 3nv0 | opls | 1.84 | 0.0997 | 2.72 | 0 | 98.16 | 2.25 | 4 | 7.74 | 1.69 | 0.1584 | 0.2076 | 0.0492 |
| 3nv0 | noschrodinger | 1.84 | 0.0061 | 0.99 | 0.3 | 97.87 | 4.15 | 0 | 7.96 | 1.94 | 0.1601 | 0.2066 | 0.0465 |
| 3nwe | opls | 1.5 | 0.0995 | 2.3 | 0 | 96.19 | 1.04 | 0 | 2.78 | 1.34 | 0.1283 | 0.1601 | 0.0318 |
| 3nwe | noschrodinger | 1.5 | 0.0043 | 0.97 | 0 | 96.21 | 0.52 | 0 | 2.23 | 1.26 | 0.1324 | 0.1598 | 0.0274 |
| 3nx2 | opls | 2.01 | 0.0991 | 2.29 | 0 | 96.88 | 2.8 | 1 | 1.37 | 1.4 | 0.2079 | 0.232 | 0.0241 |
| 3nx2 | noschrodinger | 2.01 | 0.0029 | 0.84 | 0 | 93.48 | 6.6 | 0 | 2.55 | 2.09 | 0.2003 | 0.2336 | 0.0333 |
| 3ny7 | opls | 1.92 | 0.1023 | 2.52 | 0 | 96.83 | 0.61 | 2 | 1.57 | 1.1 | 0.1598 | 0.2002 | 0.0404 |
| 3ny7 | noschrodinger | 1.92 | 0.0046 | 0.99 | 0 | 97.91 | 1.22 | 0 | 4.11 | 1.28 | 0.1642 | 0.2016 | 0.0374 |
| 3nym | opls | 1.9 | 0.0921 | 2.4 | 0 | 98.92 | 1.95 | 0 | 1.46 | 1.1 | 0.1704 | 0.2062 | 0.0358 |
| 3nym | noschrodinger | 1.9 | 0.0135 | 0.82 | 0 | 98.55 | 1.85 | 0 | 1.46 | 1.09 | 0.1718 | 0.2071 | 0.0353 |
| 3nzm | opls | 1.55 | 0.1044 | 2.26 | 0 | 98.76 | 0 | 0 | 0.78 | 0.74 | 0.1644 | 0.2072 | 0.0428 |
| 3nzm | noschrodinger | 1.55 | 0.0075 | 0.98 | 0 | 98.76 | 0 | 0 | 0.77 | 0.74 | 0.1679 | 0.2089 | 0.041 |
| 3o1h | opls | 3.1 | 0.0926 | 3.05 | 0.9 | 91.04 | 8.4 | 49 | 4.75 | 2.47 | 0.1975 | 0.2348 | 0.0373 |
| 3o1h | noschrodinger | 3.1 | 0.0029 | 0.75 | 0.54 | 91.07 | 8.76 | 0 | 3.43 | 2.37 | 0.2038 | 0.2333 | 0.0295 |
| 3o2i | opls | 2.2 | 0.0969 | 2.24 | 0 | 99.25 | 1.22 | 0 | 1.5 | 0.96 | 0.2177 | 0.2583 | 0.0406 |

| | | | | | | | | | | | | | |
|---|---|---|---|---|---|---|---|---|---|---|---|---|---|
| 3o2i | noschrodinger | 2.2 | 0.0024 | 0.7 | 0 | 98.19 | 2.4 | 0 | 1.08 | 1.1 | 0.2107 | 0.2468 | 0.0361 |
| 3o3j | opls | 3 | 0.0937 | 2.37 | 0.72 | 95.68 | 7.59 | 1 | 1.09 | 1.78 | 0.3134 | 0.3596 | 0.0462 |
| 3o3j | noschrodinger | 3 | 0.0044 | 0.93 | 0 | 92.13 | 22.07 | 0 | 6.27 | 2.85 | 0.2686 | 0.3186 | 0.05 |
| 3o57 | opls | 2 | 0.0997 | 2.27 | 0 | 98.2 | 0.66 | 1 | 2.01 | 0.97 | 0.2013 | 0.2241 | 0.0228 |
| 3o57 | noschrodinger | 2 | 0.0036 | 0.88 | 0 | 99.1 | 1.97 | 0 | 4.03 | 1.41 | 0.1912 | 0.2187 | 0.0275 |
| 3o5t | opls | 2.09 | 0.1021 | 2.28 | 0 | 96.88 | 1.95 | 2 | 0.49 | 1.08 | 0.1605 | 0.1889 | 0.0284 |
| 3o5t | noschrodinger | 2.09 | 0.0033 | 0.85 | 0 | 97.43 | 4.5 | 0 | 1.8 | 1.55 | 0.1486 | 0.187 | 0.0384 |
| 3o69 | opls | 2.1 | 0.102 | 2.25 | 0.85 | 95.44 | 2.19 | 0 | 2.19 | 1.57 | 0.1985 | 0.2358 | 0.0373 |
| 3o69 | noschrodinger | 2.1 | 0.0028 | 0.76 | 0.85 | 95.77 | 3.7 | 0 | 2.36 | 1.74 | 0.1914 | 0.227 | 0.0356 |
| 3o7j | opls | 2 | 0.0974 | 2.32 | 0 | 97.52 | 2.38 | 0 | 0 | 0.89 | 0.1841 | 0.2074 | 0.0233 |
| 3o7j | noschrodinger | 2 | 0.0028 | 0.77 | 0 | 98.17 | 3.97 | 0 | 0.39 | 1.1 | 0.1821 | 0.2096 | 0.0275 |
| 3ocm | opls | 1.8 | 0.1012 | 2.33 | 0 | 99.63 | 2.02 | 0 | 2.29 | 1.24 | 0.1675 | 0.1954 | 0.0279 |
| 3ocm | noschrodinger | 1.8 | 0.0056 | 1.01 | 0 | 100 | 1.56 | 0 | 1.66 | 1.06 | 0.1714 | 0.1963 | 0.0249 |
| 3ocp | opls | 2.49 | 0.0941 | 2.47 | 0 | 98.8 | 3.06 | 2 | 2.42 | 1.39 | 0.1849 | 0.2232 | 0.0383 |
| 3ocp | noschrodinger | 2.49 | 0.0063 | 0.96 | 0 | 97.21 | 3.9 | 0 | 2.42 | 1.62 | 0.1878 | 0.2238 | 0.036 |
| 3ocv | opls | 1.55 | 0.1016 | 2.11 | 0 | 97.72 | 0 | 1 | 0.26 | 0.66 | 0.1544 | 0.1678 | 0.0134 |
| 3ocv | noschrodinger | 1.55 | 0.0053 | 0.97 | 0 | 97.55 | 0 | 0 | 0.26 | 0.69 | 0.1521 | 0.168 | 0.0159 |
| 3od2 | opls | 2.6 | 0.0912 | 2.35 | 0.45 | 98.65 | 3.81 | 0 | 0.74 | 1.18 | 0.2496 | 0.2843 | 0.0347 |
| 3od2 | noschrodinger | 2.6 | 0.0032 | 0.86 | 1.18 | 92.94 | 4.21 | 0 | 2.51 | 1.96 | 0.2323 | 0.2731 | 0.0408 |
| 3og9 | opls | 1.88 | 0.1026 | 2.22 | 0 | 97.78 | 2.45 | 1 | 1.93 | 1.3 | 0.1622 | 0.2033 | 0.0411 |
| 3og9 | noschrodinger | 1.88 | 0.0028 | 0.8 | 0 | 97.55 | 2.43 | 0 | 1.48 | 1.27 | 0.1626 | 0.1989 | 0.0363 |
| 3oi8 | opls | 1.99 | 0.0947 | 2.3 | 0 | 98.92 | 0.75 | 1 | 0.79 | 0.75 | 0.1972 | 0.2312 | 0.034 |
| 3oi8 | noschrodinger | 1.99 | 0.0041 | 0.83 | 0 | 97.39 | 1.44 | 0 | 2.56 | 1.28 | 0.188 | 0.2276 | 0.0396 |
| 3oou | opls | 1.57 | 0.1016 | 2.43 | 0 | 100 | 1.02 | 3 | 1.6 | 0.91 | 0.1823 | 0.2258 | 0.0435 |
| 3oou | noschrodinger | 1.57 | 0.0105 | 1.02 | 0 | 100 | 0 | 0 | 2.13 | 0.99 | 0.1807 | 0.2193 | 0.0386 |
| 3oqi | opls | 1.7 | 0.1004 | 2.42 | 0 | 98.28 | 0 | 0 | 3.86 | 1.17 | 0.1428 | 0.1718 | 0.029 |
| 3oqi | noschrodinger | 1.7 | 0.0084 | 1.05 | 0 | 98.17 | 0.82 | 0 | 3.04 | 1.1 | 0.1472 | 0.1703 | 0.0231 |
| 3oqj | opls | 2.4 | 0.095 | 2.31 | 0 | 98.47 | 0.86 | 2 | 1.16 | 0.83 | 0.1882 | 0.2248 | 0.0366 |
| 3oqj | noschrodinger | 2.4 | 0.0031 | 0.79 | 0.24 | 97.18 | 1.43 | 0 | 2.93 | 1.35 | 0.1749 | 0.2157 | 0.0408 |
| 3oqv | opls | 1.9 | 0.0959 | 2.41 | 0 | 98.36 | 0 | 2 | 1.84 | 0.95 | 0.1858 | 0.2345 | 0.0487 |
| 3oqv | noschrodinger | 1.9 | 0.003 | 0.79 | 0 | 96.62 | 0.61 | 0 | 4.04 | 1.41 | 0.1877 | 0.2268 | 0.0391 |
| 3ose | opls | 1.65 | 0.0999 | 2.37 | 0 | 98.91 | 3.09 | 0 | 6.23 | 1.72 | 0.1483 | 0.1683 | 0.02 |
| 3ose | noschrodinger | 1.65 | 0.0055 | 0.98 | 0 | 98.98 | 2.06 | 0 | 9.08 | 1.72 | 0.1509 | 0.1667 | 0.0158 |
| 3ot2 | opls | 1.96 | 0.0982 | 2.3 | 0 | 97.25 | 0 | 0 | 1.53 | 1.03 | 0.2466 | 0.2793 | 0.0327 |
| 3ot2 | noschrodinger | 1.96 | 0.0086 | 0.93 | 0 | 98.31 | 0.3 | 0 | 1.36 | 0.87 | 0.252 | 0.2784 | 0.0264 |
| 3ot6 | opls | 2.51 | 0.0937 | 2.3 | 0 | 96.08 | 3.55 | 2 | 0.57 | 1.38 | 0.2028 | 0.2257 | 0.0229 |
| 3ot6 | noschrodinger | 2.51 | 0.007 | 1.01 | 0 | 96.49 | 4.49 | 0 | 0.85 | 1.49 | 0.1778 | 0.2209 | 0.0431 |
| 3ouh | opls | 2.51 | 0.0923 | 2.35 | 0 | 96.6 | 5.14 | 1 | 1.19 | 1.59 | 0.2202 | 0.2443 | 0.0241 |
| 3ouh | noschrodinger | 2.51 | 0.0029 | 0.79 | 1.44 | 93.75 | 5.65 | 0 | 1.79 | 1.92 | 0.2007 | 0.2266 | 0.0259 |
| 3oum | opls | 2 | 0.0954 | 2.26 | 0 | 96.67 | 2.79 | 3 | 0.58 | 1.24 | 0.189 | 0.2203 | 0.0313 |
| 3oum | noschrodinger | 2 | 0.0059 | 0.93 | 0 | 96.36 | 4.37 | 0 | 2.31 | 1.74 | 0.193 | 0.2229 | 0.0299 |
| 3owc | opls | 1.9 | 0.096 | 2.61 | 0 | 99.01 | 2.51 | 1 | 3.26 | 1.42 | 0.1895 | 0.2183 | 0.0288 |
| 3owc | noschrodinger | 1.9 | 0.0083 | 1.51 | 0 | 98.82 | 3.51 | 0 | 5.22 | 1.69 | 0.195 | 0.2195 | 0.0245 |
| 3oyw | opls | 2.5 | 0.092 | 2.24 | 0 | 95.47 | 0 | 2 | 0.48 | 0.98 | 0.1819 | 0.2238 | 0.0419 |
| 3oyw | noschrodinger | 2.5 | 0.0033 | 0.88 | 0 | 96.34 | 0.93 | 0 | 1.43 | 1.12 | 0.161 | 0.2122 | 0.0512 |
| 3p0f | opls | 1.54 | 0.1023 | 2.47 | 0 | 97.59 | 0.78 | 1 | 0.64 | 0.8 | 0.1618 | 0.1881 | 0.0263 |
| 3p0f | noschrodinger | 1.54 | 0.004 | 0.9 | 0 | 97.59 | 0.39 | 0 | 0.43 | 0.74 | 0.1681 | 0.1843 | 0.0162 |
| 3p0m | opls | 2 | 0.0964 | 2.19 | 0 | 96.91 | 0.34 | 0 | 0.35 | 0.81 | 0.1893 | 0.215 | 0.0257 |
| 3p0m | noschrodinger | 2 | 0.0105 | 1.04 | 0 | 97.11 | 0.68 | 0 | 1.05 | 0.97 | 0.1724 | 0.2064 | 0.034 |

| | | | | | | | | | | | | | |
|---|---|---|---|---|---|---|---|---|---|---|---|---|---|
| 3p0y | opls | 1.8 | 0.1013 | 2.18 | 0 | 98.03 | 0.37 | 4 | 1.94 | 0.96 | 0.175 | 0.2119 | 0.0369 |
| 3p0y | noschrodinger | 1.8 | 0.0034 | 0.84 | 0.16 | 98.06 | 0.18 | 0 | 1.84 | 0.94 | 0.1768 | 0.2084 | 0.0316 |
| 3p2h | opls | 2 | 0.0958 | 2.3 | 0 | 98.31 | 1.39 | 0 | 2.04 | 1.08 | 0.1782 | 0.2042 | 0.026 |
| 3p2h | noschrodinger | 2 | 0.0036 | 0.89 | 0 | 98.33 | 1.36 | 0 | 1.36 | 0.97 | 0.1802 | 0.2065 | 0.0263 |
| 3p2r | opls | 2.46 | 0.094 | 2.32 | 0 | 97.69 | 4.59 | 0 | 0.96 | 1.36 | 0.2172 | 0.257 | 0.0398 |
| 3p2r | noschrodinger | 2.46 | 0.003 | 0.81 | 0 | 96.95 | 9.09 | 0 | 5.07 | 2.18 | 0.1883 | 0.247 | 0.0587 |
| 3p3v | opls | 1.65 | 0.0993 | 2.34 | 0 | 98.58 | 0.7 | 0 | 1.33 | 0.86 | 0.1621 | 0.1961 | 0.034 |
| 3p3v | noschrodinger | 1.65 | 0.0068 | 0.99 | 0 | 98.44 | 1.03 | 0 | 0.57 | 0.7 | 0.162 | 0.1988 | 0.0368 |
| 3p7n | opls | 2.1 | 0.0931 | 2.32 | 0 | 97.3 | 1.89 | 3 | 0.65 | 1.06 | 0.2688 | 0.3105 | 0.0417 |
| 3p7n | noschrodinger | 2.1 | 0.0028 | 0.76 | 0 | 96.26 | 2.48 | 0 | 3.77 | 1.72 | 0.2523 | 0.2886 | 0.0363 |
| 3p8a | opls | 1.95 | 0.095 | 2.52 | 0 | 96.09 | 3.07 | 1 | 1.62 | 1.55 | 0.2269 | 0.2619 | 0.035 |
| 3p8a | noschrodinger | 1.95 | 0.0057 | 0.89 | 0 | 97.18 | 4.25 | 0 | 2.32 | 1.64 | 0.219 | 0.2556 | 0.0366 |
| 3p8k | opls | 1.7 | 0.1008 | 2.43 | 0 | 96.57 | 0.85 | 5 | 3.78 | 1.39 | 0.182 | 0.2199 | 0.0379 |
| 3p8k | noschrodinger | 1.7 | 0.0032 | 0.88 | 0.19 | 96.05 | 1.04 | 0 | 2.4 | 1.3 | 0.1908 | 0.2153 | 0.0245 |
| 3pa8 | opls | 2 | 0.0966 | 2.26 | 0 | 98.93 | 1.6 | 0 | 0.88 | 0.92 | 0.1924 | 0.2284 | 0.036 |
| 3pa8 | noschrodinger | 2 | 0.0029 | 1.53 | 0 | 98.55 | 2.5 | 0 | 1.14 | 1.13 | 0.1829 | 0.2288 | 0.0459 |
| 3pd0 | opls | 2 | 0.1007 | 2.22 | 0 | 99.57 | 0.47 | 2 | 0.77 | 0.74 | 0.1628 | 0.1976 | 0.0348 |
| 3pd0 | noschrodinger | 2 | 0.0025 | 0.74 | 0 | 97.03 | 0.46 | 0 | 1.81 | 1.11 | 0.1525 | 0.1928 | 0.0403 |
| 3pds | opls | 3.5 | 0.0906 | 2.27 | 0.23 | 98.17 | 2.62 | 1 | 0.41 | 0.96 | 0.221 | 0.2682 | 0.0472 |
| 3pds | noschrodinger | 3.5 | 0.003 | 0.85 | 0 | 97.27 | 3.66 | 0 | 0.55 | 1.25 | 0.2344 | 0.2645 | 0.0301 |
| 3pdt | opls | 1.8 | 0.1029 | 2.27 | 0 | 97.11 | 0 | 6 | 1.47 | 1.05 | 0.1676 | 0.2005 | 0.0329 |
| 3pdt | noschrodinger | 1.8 | 0.005 | 1.06 | 0.41 | 96.73 | 0.45 | 0 | 4.93 | 1.46 | 0.1702 | 0.2022 | 0.032 |
| 3peg | opls | 2.52 | 0.0902 | 2.25 | 0 | 97.48 | 3.29 | 3 | 1.47 | 1.38 | 0.2427 | 0.2671 | 0.0244 |
| 3peg | noschrodinger | 2.52 | 0.0029 | 0.81 | 0.42 | 94.17 | 3.27 | 0 | 2.94 | 1.87 | 0.2237 | 0.2557 | 0.032 |
| 3pev | opls | 2.5 | 0.092 | 2.36 | 0 | 97.69 | 0.98 | 2 | 1.03 | 0.87 | 0.2226 | 0.2589 | 0.0363 |
| 3pev | noschrodinger | 2.5 | 0.0087 | 1.25 | 0 | 97.22 | 1.2 | 0 | 3.09 | 1.3 | 0.2212 | 0.2576 | 0.0364 |
| 3pfq | opls | 4 | 0.0902 | 2.35 | 1.97 | 90.75 | 5.86 | 2 | 1.07 | 1.92 | 0.2013 | 0.2448 | 0.0435 |
| 3pfq | noschrodinger | 4 | 0.0026 | 0.72 | 4.31 | 81.76 | 6.48 | 0 | 4.26 | 2.54 | 0.1938 | 0.2415 | 0.0477 |
| 3pft | opls | 1.6 | 0.1054 | 2.29 | 0 | 99.02 | 1.54 | 1 | 2.04 | 1.12 | 0.1651 | 0.2017 | 0.0366 |
| 3pft | noschrodinger | 1.6 | 0.0083 | 1.1 | 0 | 99.03 | 1.91 | 0 | 2.24 | 1.21 | 0.1639 | 0.1963 | 0.0324 |
| 3pg0 | opls | 1.62 | 0.0971 | 2.53 | 0 | 98.52 | 2.5 | 0 | 7.15 | 1.7 | 0.1434 | 0.1789 | 0.0355 |
| 3pg0 | noschrodinger | 1.62 | 0.0043 | 0.92 | 0 | 98.55 | 4.17 | 0 | 7.96 | 1.9 | 0.1465 | 0.1744 | 0.0279 |
| 3pg7 | opls | 2.19 | 0.0941 | 2.33 | 0.23 | 95.57 | 3.11 | 0 | 0.61 | 1.38 | 0.2455 | 0.2764 | 0.0309 |
| 3pg7 | noschrodinger | 2.19 | 0.0026 | 0.78 | 0.79 | 95.67 | 6.22 | 0 | 3.82 | 2.07 | 0.229 | 0.265 | 0.036 |
| 3pjt | opls | 2.52 | 0.0939 | 2.39 | 0 | 98.31 | 9.2 | 1 | 1.52 | 1.63 | 0.1902 | 0.2302 | 0.04 |
| 3pjt | noschrodinger | 2.52 | 0.0028 | 0.83 | 0 | 97.69 | 13.18 | 0 | 3.67 | 2.08 | 0.1931 | 0.225 | 0.0319 |
| 3pju | opls | 2.5 | 0.0966 | 2.41 | 0 | 98.78 | 9.13 | 0 | 1.47 | 1.62 | 0.2145 | 0.2304 | 0.0159 |
| 3pju | noschrodinger | 2.5 | 0.0027 | 0.82 | 0 | 98.38 | 11.96 | 0 | 3.68 | 1.98 | 0.2018 | 0.2252 | 0.0234 |
| 3pmd | opls | 1.76 | 0.097 | 2.47 | 0 | 99.33 | 2.65 | 2 | 2.2 | 1.32 | 0.1807 | 0.2125 | 0.0318 |
| 3pmd | noschrodinger | 1.76 | 0.004 | 0.78 | 0 | 98.68 | 1.32 | 0 | 1.46 | 0.98 | 0.1889 | 0.2061 | 0.0172 |
| 3pn4 | opls | 1.9 | 0.1031 | 2.22 | 0 | 99.33 | 1.33 | 0 | 2.46 | 1.12 | 0.1882 | 0.2277 | 0.0395 |
| 3pn4 | noschrodinger | 1.9 | 0.0066 | 0.97 | 0 | 98.88 | 1.33 | 0 | 4.61 | 1.33 | 0.1908 | 0.2282 | 0.0374 |
| 3pne | opls | 1.97 | 0.0942 | 2.26 | 0 | 97.66 | 1.22 | 4 | 1.47 | 1.02 | 0.1974 | 0.2197 | 0.0223 |
| 3pne | noschrodinger | 1.97 | 0.0035 | 0.87 | 0 | 97.79 | 1.9 | 0 | 2.49 | 1.29 | 0.2046 | 0.2245 | 0.0199 |
| 3pnl | opls | 2.2 | 0.0931 | 2.37 | 0.19 | 96.36 | 4.75 | 1 | 1.06 | 1.56 | 0.2123 | 0.2442 | 0.0319 |
| 3pnl | noschrodinger | 2.2 | 0.003 | 0.86 | 0.18 | 96.63 | 8.55 | 0 | 3.05 | 2.02 | 0.203 | 0.23 | 0.027 |
| 3pp9 | opls | 1.6 | 0.0958 | 2.33 | 0.22 | 96.73 | 0.82 | 3 | 2.49 | 1.24 | 0.1953 | 0.2249 | 0.0296 |
| 3pp9 | noschrodinger | 1.6 | 0.0135 | 1.3 | 0.2 | 96.06 | 1.76 | 0 | 7.14 | 1.85 | 0.1919 | 0.226 | 0.0341 |
| 3ppb | opls | 2.1 | 0.092 | 2.22 | 0 | 99.09 | 0 | 0 | 1.92 | 0.96 | 0.1783 | 0.2223 | 0.044 |

| | | | | | | | | | | | | | |
|---|---|---|---|---|---|---|---|---|---|---|---|---|---|
| 3ppb | noschrodinger | 2.1 | 0.0026 | 0.7 | 0 | 98.66 | 0 | 0 | 0.8 | 0.75 | 0.1848 | 0.2163 | 0.0315 |
| 3pqc | opls | 1.9 | 0.1012 | 2.3 | 0.26 | 95.54 | 1.14 | 2 | 2.91 | 1.43 | 0.1591 | 0.2089 | 0.0498 |
| 3pqc | noschrodinger | 1.9 | 0.0041 | 0.92 | 0.26 | 95.81 | 0.85 | 0 | 2.3 | 1.3 | 0.1655 | 0.2054 | 0.0399 |
| 3prn | opls | 1.9 | 0.0941 | 2.48 | 0.35 | 97.2 | 2.63 | 2 | 0 | 0.97 | 0.1553 | 0.1728 | 0.0175 |
| 3prn | noschrodinger | 1.9 | 0.0051 | 0.99 | 0.35 | 95.47 | 3.93 | 0 | 1.59 | 1.67 | 0.1551 | 0.1727 | 0.0176 |
| 3ps8 | opls | 2.55 | 0.0984 | 2.27 | 0 | 99.07 | 2.15 | 0 | 1.15 | 1.08 | 0.2205 | 0.2532 | 0.0327 |
| 3ps8 | noschrodinger | 2.55 | 0.0028 | 0.74 | 0 | 98.13 | 1.08 | 0 | 0 | 0.52 | 0.2089 | 0.2504 | 0.0415 |
| 3put | opls | 1.83 | 0.0972 | 2.27 | 0 | 96.83 | 0.4 | 0 | 0.82 | 0.95 | 0.1696 | 0.2108 | 0.0412 |
| 3put | noschrodinger | 1.83 | 0.0028 | 0.82 | 0.33 | 96.32 | 1.93 | 0 | 1.64 | 1.38 | 0.1734 | 0.2067 | 0.0333 |
| 3pvt | opls | 2.03 | 0.0958 | 2.27 | 0.13 | 98.71 | 0.47 | 1 | 0.16 | 0.56 | 0.1893 | 0.2136 | 0.0243 |
| 3pvt | noschrodinger | 2.03 | 0.0026 | 0.74 | 0.26 | 98.21 | 0.94 | 0 | 1.2 | 0.84 | 0.1865 | 0.209 | 0.0225 |
| 3pwy | opls | 3.5 | 0.09 | 2.26 | 0 | 96.55 | 0.41 | 1 | 1.09 | 1.04 | 0.2146 | 0.2454 | 0.0308 |
| 3pwy | noschrodinger | 3.5 | 0.0026 | 0.74 | 0.72 | 93.86 | 1.64 | 0 | 1.76 | 1.51 | 0.191 | 0.2378 | 0.0468 |
| 3px2 | opls | 1.65 | 0.0995 | 2.39 | 0 | 98.95 | 1.87 | 3 | 2.67 | 1.26 | 0.1837 | 0.217 | 0.0333 |
| 3px2 | noschrodinger | 1.65 | 0.0081 | 1.02 | 0 | 98.95 | 2.67 | 0 | 2.67 | 1.38 | 0.1856 | 0.2143 | 0.0287 |
| 3q18 | opls | 1.7 | 0.0987 | 2.35 | 0.22 | 98.49 | 1.11 | 5 | 3.69 | 1.19 | 0.1421 | 0.1739 | 0.0318 |
| 3q18 | noschrodinger | 1.7 | 0.0053 | 0.94 | 0.22 | 98.28 | 0.44 | 1 | 2.34 | 1.01 | 0.1496 | 0.1722 | 0.0226 |
| 3q20 | opls | 1.71 | 0.101 | 2.9 | 0 | 98.12 | 2.96 | 5 | 5.62 | 1.66 | 0.19 | 0.2304 | 0.0404 |
| 3q20 | noschrodinger | 1.71 | 0.0081 | 0.98 | 0 | 98.15 | 3.4 | 0 | 2.7 | 1.46 | 0.1956 | 0.229 | 0.0334 |
| 3q33 | opls | 2.8 | 0.0907 | 2.31 | 0.85 | 94.07 | 4.58 | 2 | 1.09 | 1.71 | 0.2681 | 0.2924 | 0.0243 |
| 3q33 | noschrodinger | 2.8 | 0.0027 | 0.76 | 2.18 | 90.27 | 7.44 | 0 | 3.17 | 2.31 | 0.2495 | 0.2775 | 0.028 |
| 3q4r | opls | 1.6 | 0.1004 | 2.28 | 0 | 100 | 1.28 | 1 | 2.11 | 1.07 | 0.1748 | 0.2117 | 0.0369 |
| 3q4r | noschrodinger | 1.6 | 0.0067 | 0.85 | 0 | 99.4 | 1.91 | 0 | 3.53 | 1.36 | 0.1789 | 0.2099 | 0.031 |
| 3q7s | opls | 2.09 | 0.0943 | 2.22 | 0.52 | 98.43 | 0.53 | 0 | 1.16 | 0.83 | 0.1943 | 0.227 | 0.0327 |
| 3q7s | noschrodinger | 2.09 | 0.0025 | 0.76 | 0.51 | 98.97 | 1.03 | 0 | 0.58 | 0.71 | 0.1923 | 0.2219 | 0.0296 |
| 3q80 | opls | 2 | 0.0998 | 2.36 | 0 | 98.55 | 0.88 | 4 | 2.23 | 1 | 0.162 | 0.2036 | 0.0416 |
| 3q80 | noschrodinger | 2 | 0.005 | 1.03 | 0.24 | 98.11 | 1.17 | 0 | 4.03 | 1.24 | 0.1657 | 0.2069 | 0.0412 |
| 3qbm | opls | 1.8 | 0.098 | 2.38 | 0 | 99.43 | 1.27 | 6 | 2.72 | 1.14 | 0.1651 | 0.205 | 0.0399 |
| 3qbm | noschrodinger | 1.8 | 0.0032 | 0.8 | 0 | 99.49 | 0.91 | 0 | 1.96 | 0.96 | 0.1721 | 0.1997 | 0.0276 |
| 3qfo | opls | 2.2 | 0.097 | 2.25 | 0.2 | 96.56 | 0.69 | 1 | 0.73 | 0.96 | 0.1826 | 0.2147 | 0.0321 |
| 3qfo | noschrodinger | 2.2 | 0.0026 | 0.79 | 0.4 | 96.37 | 0.68 | 0 | 1.1 | 1.06 | 0.1767 | 0.2119 | 0.0352 |
| 3qh8 | opls | 1.6 | 0.102 | 2.3 | 0.4 | 97.19 | 0 | 1 | 3.19 | 1.26 | 0.1369 | 0.1664 | 0.0295 |
| 3qh8 | noschrodinger | 1.6 | 0.0084 | 1.18 | 0.37 | 97.78 | 0.84 | 0 | 0.92 | 0.83 | 0.136 | 0.1636 | 0.0276 |
| 3qme | opls | 2 | 0.0953 | 2.3 | 0 | 98.23 | 0.94 | 0 | 2.09 | 0.98 | 0.1854 | 0.2355 | 0.0501 |
| 3qme | noschrodinger | 2 | 0.0026 | 0.71 | 0 | 99.13 | 0.93 | 0 | 1.04 | 0.8 | 0.1679 | 0.215 | 0.0471 |
| 3qp4 | opls | 1.55 | 0.0997 | 2.59 | 0 | 98.29 | 1.37 | 3 | 1.76 | 1.04 | 0.1929 | 0.2211 | 0.0282 |
| 3qp4 | noschrodinger | 1.55 | 0.0128 | 1.31 | 0 | 98.86 | 0.68 | 0 | 1.06 | 0.81 | 0.1918 | 0.2176 | 0.0258 |
| 3qp6 | opls | 2 | 0.1014 | 2.23 | 0 | 99.22 | 1.38 | 0 | 1.47 | 0.99 | 0.1729 | 0.2094 | 0.0365 |
| 3qp6 | noschrodinger | 2 | 0.003 | 0.79 | 0 | 99.61 | 1.38 | 0 | 0.98 | 0.9 | 0.1711 | 0.2034 | 0.0323 |
| 3qq7 | opls | 2.65 | 0.0921 | 2.43 | 1.2 | 96.39 | 2.6 | 4 | 1.45 | 1.44 | 0.2189 | 0.2379 | 0.019 |
| 3qq7 | noschrodinger | 2.65 | 0.0029 | 0.81 | 1.81 | 93.98 | 5.19 | 0 | 6.15 | 2.29 | 0.2295 | 0.2481 | 0.0186 |
| 3qqa | opls | 2.2 | 0.0933 | 2.49 | 0.51 | 97.46 | 2.79 | 2 | 0.89 | 1.22 | 0.2338 | 0.2588 | 0.025 |
| 3qqa | noschrodinger | 2.2 | 0.0065 | 1.28 | 0 | 96.97 | 3.89 | 0 | 13.07 | 2.25 | 0.2188 | 0.2482 | 0.0294 |
| 3qra | opls | 1.8 | 0.0898 | 2.18 | 0 | 98.32 | 0.9 | 0 | 1.59 | 0.91 | 0.1889 | 0.2143 | 0.0254 |
| 3qra | noschrodinger | 1.8 | 0.0075 | 1.11 | 0 | 98.63 | 0.87 | 0 | 4.04 | 1.19 | 0.192 | 0.2174 | 0.0254 |
| 3qtb | opls | 2.08 | 0.0921 | 2.28 | 0 | 98.28 | 0.98 | 0 | 0.74 | 0.74 | 0.2209 | 0.246 | 0.0251 |
| 3qtb | noschrodinger | 2.08 | 0.0032 | 0.78 | 0 | 99.21 | 1.45 | 0 | 0.25 | 0.72 | 0.1951 | 0.2392 | 0.0441 |
| 3qtt | opls | 2.6 | 0.0924 | 2.36 | 0.2 | 96.73 | 5.66 | 4 | 0.57 | 1.47 | 0.2214 | 0.2496 | 0.0282 |
| 3qtt | noschrodinger | 2.6 | 0.0049 | 0.99 | 0 | 97.67 | 9.78 | 2 | 6.72 | 2.19 | 0.2115 | 0.24 | 0.0285 |

| | | | | | | | | | | | | | |
|---|---|---|---|---|---|---|---|---|---|---|---|---|---|
| 3qut | opls | 1.5 | 0.1023 | 2.53 | 0 | 98.61 | 0.51 | 0 | 1.93 | 0.96 | 0.1325 | 0.1519 | 0.0194 |
| 3qut | noschrodinger | 1.5 | 0.0066 | 1.11 | 0 | 98.65 | 1.03 | 0 | 2.48 | 1.04 | 0.1359 | 0.1497 | 0.0138 |
| 3qvk | opls | 2 | 0.0975 | 2.29 | 0 | 97.58 | 1.69 | 1 | 0.55 | 0.95 | 0.17 | 0.2004 | 0.0304 |
| 3qvk | noschrodinger | 2 | 0.007 | 1.11 | 0 | 97.53 | 2.23 | 0 | 0.97 | 1.15 | 0.1698 | 0.2024 | 0.0326 |
| 3qvu | opls | 2.5 | 0.0919 | 2.26 | 0 | 97.2 | 3.44 | 0 | 1.36 | 1.42 | 0.2037 | 0.2435 | 0.0398 |
| 3qvu | noschrodinger | 2.5 | 0.0073 | 1.17 | 0.17 | 95.98 | 2.87 | 2 | 3.25 | 1.74 | 0.1937 | 0.2314 | 0.0377 |
| 3qw9 | opls | 2 | 0.0939 | 2.22 | 0.95 | 96.84 | 2.61 | 0 | 1.97 | 1.47 | 0.2206 | 0.257 | 0.0364 |
| 3qw9 | noschrodinger | 2 | 0.0041 | 0.83 | 0.91 | 96.06 | 2.57 | 0 | 5.74 | 1.89 | 0.207 | 0.2489 | 0.0419 |
| 3qwq | opls | 2.75 | 0.0868 | 2.34 | 0.48 | 95.66 | 5.14 | 5 | 1.37 | 1.71 | 0.2412 | 0.2579 | 0.0167 |
| 3qwq | noschrodinger | 2.75 | 0.0093 | 1.29 | 0.85 | 92.24 | 9.91 | 0 | 7.01 | 2.62 | 0.2332 | 0.2569 | 0.0237 |
| 3r03 | opls | 2.49 | 0.092 | 2.38 | 0 | 97.91 | 3.48 | 0 | 0.97 | 1.22 | 0.2196 | 0.2562 | 0.0366 |
| 3r03 | noschrodinger | 2.49 | 0.003 | 0.85 | 0 | 95.67 | 5.31 | 0 | 2.44 | 1.88 | 0.2211 | 0.2528 | 0.0317 |
| 3r1v | opls | 2.19 | 0.0921 | 2.49 | 0.42 | 95.78 | 8.84 | 10 | 1.27 | 1.86 | 0.2081 | 0.2552 | 0.0471 |
| 3r1v | noschrodinger | 2.19 | 0.0036 | 0.9 | 0.41 | 93.83 | 13.95 | 0 | 3.58 | 2.43 | 0.2197 | 0.2578 | 0.0381 |
| 3r20 | opls | 2 | 0.1023 | 2.27 | 0 | 99.05 | 0 | 0 | 0 | 0.5 | 0.1851 | 0.2107 | 0.0256 |
| 3r20 | noschrodinger | 2 | 0.0039 | 0.83 | 0 | 99.07 | 0 | 0 | 0.31 | 0.62 | 0.1704 | 0.2133 | 0.0429 |
| 3r5z | opls | 1.5 | 0.0982 | 2.36 | 0 | 99.64 | 0.84 | 2 | 3.01 | 1.09 | 0.1647 | 0.1851 | 0.0204 |
| 3r5z | noschrodinger | 1.5 | 0.0086 | 1.19 | 0 | 99.64 | 0.83 | 0 | 2.8 | 1.07 | 0.165 | 0.185 | 0.02 |
| 3r95 | opls | 1.6 | 0.1025 | 2.33 | 0 | 99.71 | 0.66 | 1 | 1.2 | 0.84 | 0.1625 | 0.1941 | 0.0316 |
| 3r95 | noschrodinger | 1.6 | 0.0041 | 1.04 | 0 | 99.15 | 0.33 | 0 | 2.74 | 1.06 | 0.1682 | 0.1924 | 0.0242 |
| 3r9v | opls | 1.9 | 0.0967 | 2.35 | 0.86 | 97.13 | 0 | 1 | 1.04 | 0.96 | 0.212 | 0.2302 | 0.0182 |
| 3r9v | noschrodinger | 1.9 | 0.0105 | 1.32 | 1.13 | 96.6 | 1.27 | 0 | 6.41 | 1.65 | 0.2098 | 0.2336 | 0.0238 |
| 3ret | opls | 1.79 | 0.0919 | 2.25 | 0 | 98.87 | 0 | 2 | 0.33 | 0.62 | 0.2058 | 0.2347 | 0.0289 |
| 3ret | noschrodinger | 1.79 | 0.0032 | 0.8 | 0 | 98.9 | 0.66 | 0 | 1.96 | 0.96 | 0.2034 | 0.2312 | 0.0278 |
| 3rfv | opls | 2.1 | 0.1012 | 2.38 | 0 | 97.84 | 0 | 1 | 0.89 | 0.81 | 0.1603 | 0.1789 | 0.0186 |
| 3rfv | noschrodinger | 2.1 | 0.0028 | 0.84 | 0 | 97.59 | 0.31 | 0 | 1.05 | 0.89 | 0.1525 | 0.1683 | 0.0158 |
| 3rg9 | opls | 2 | 0.1032 | 2.3 | 0 | 98.15 | 0.27 | 0 | 1 | 0.8 | 0.1699 | 0.1959 | 0.026 |
| 3rg9 | noschrodinger | 2 | 0.0031 | 0.83 | 0 | 98.61 | 0.82 | 0 | 1.86 | 0.95 | 0.1621 | 0.1901 | 0.028 |
| 3rga | opls | 1.59 | 0.0961 | 2.44 | 0 | 97.47 | 0.44 | 0 | 5.65 | 1.41 | 0.162 | 0.1956 | 0.0336 |
| 3rga | noschrodinger | 1.59 | 0.0043 | 0.99 | 0 | 97.85 | 0.44 | 0 | 3.49 | 1.17 | 0.1666 | 0.1912 | 0.0246 |
| 3rgq | opls | 2.05 | 0.1004 | 2.45 | 0 | 98.03 | 3.01 | 1 | 1.55 | 1.26 | 0.1676 | 0.2031 | 0.0355 |
| 3rgq | noschrodinger | 2.05 | 0.0091 | 1.2 | 0 | 97.4 | 3.73 | 0 | 3.89 | 1.73 | 0.1567 | 0.1968 | 0.0401 |
| 3rh3 | opls | 2.1 | 0.0897 | 2.12 | 0 | 99.31 | 0.29 | 0 | 0.13 | 0.55 | 0.2316 | 0.2599 | 0.0283 |
| 3rh3 | noschrodinger | 2.1 | 0.0027 | 0.71 | 0 | 98.29 | 0.27 | 0 | 0.95 | 0.79 | 0.23 | 0.2626 | 0.0326 |
| 3rh8 | opls | 2.71 | 0.0876 | 2.32 | 0.85 | 94.07 | 4.13 | 0 | 2.06 | 1.84 | 0.2205 | 0.2484 | 0.0279 |
| 3rh8 | noschrodinger | 2.71 | 0.004 | 0.85 | 1.71 | 92.47 | 5.36 | 0 | 4.33 | 2.24 | 0.2031 | 0.2262 | 0.0231 |
| 3rhn | opls | 2.1 | 0.0969 | 2.33 | 0 | 100 | 1.08 | 0 | 1.67 | 0.94 | 0.1706 | 0.2106 | 0.04 |
| 3rhn | noschrodinger | 2.1 | 0.003 | 0.92 | 0 | 99.12 | 2.15 | 0 | 2.24 | 1.25 | 0.1655 | 0.2112 | 0.0457 |
| 3rjd | opls | 2.65 | 0.0893 | 2.45 | 0 | 96.81 | 4.89 | 3 | 1.58 | 1.62 | 0.2722 | 0.3013 | 0.0291 |
| 3rjd | noschrodinger | 2.65 | 0.0036 | 0.84 | 1.19 | 96.84 | 11.01 | 0 | 5.43 | 2.28 | 0.2602 | 0.2978 | 0.0376 |
| 3rk1 | opls | 2.29 | 0.0938 | 2.3 | 0.25 | 98.26 | 2.49 | 0 | 1.11 | 1.12 | 0.2211 | 0.2464 | 0.0253 |
| 3rk1 | noschrodinger | 2.29 | 0.0026 | 0.76 | 0.23 | 96.79 | 2.66 | 0 | 2.22 | 1.52 | 0.2086 | 0.2477 | 0.0391 |
| 3rl6 | opls | 2 | 0.0973 | 2.28 | 0 | 97.72 | 0.4 | 3 | 1.25 | 0.91 | 0.1667 | 0.2019 | 0.0352 |
| 3rl6 | noschrodinger | 2 | 0.0027 | 0.79 | 0.17 | 97.38 | 0.6 | 0 | 1.46 | 1 | 0.1756 | 0.2014 | 0.0258 |
| 3rmi | opls | 2.4 | 0.0911 | 2.27 | 0 | 100 | 3.53 | 2 | 0 | 0.92 | 0.2281 | 0.2489 | 0.0208 |
| 3rmi | noschrodinger | 2.4 | 0.003 | 0.79 | 0 | 100 | 3.53 | 0 | 0.59 | 1.11 | 0.216 | 0.2499 | 0.0339 |
| 3rno | opls | 2.48 | 0.0888 | 2.26 | 0.47 | 95.79 | 0.52 | 2 | 1.36 | 1.16 | 0.1778 | 0.2102 | 0.0324 |
| 3rno | noschrodinger | 2.48 | 0.0033 | 0.84 | 0.43 | 96.1 | 3.98 | 0 | 1.36 | 1.59 | 0.1797 | 0.2142 | 0.0345 |
| 3rom | opls | 2.04 | 0.0958 | 2.33 | 0 | 96.13 | 1.08 | 3 | 0.99 | 1.08 | 0.1765 | 0.2244 | 0.0479 |

| | | | | | | | | | | | | | |
|---|---|---|---|---|---|---|---|---|---|---|---|---|---|
| 3rom | noschrodinger | 2.04 | 0.0026 | 0.76 | 0 | 96.61 | 0.86 | 0 | 1.12 | 1.04 | 0.1927 | 0.2197 | 0.027 |
| 3rpq | opls | 2.61 | 0.0909 | 2.26 | 0 | 97.73 | 1.45 | 0 | 2.12 | 1.17 | 0.2007 | 0.2368 | 0.0361 |
| 3rpq | noschrodinger | 2.61 | 0.0029 | 0.79 | 0.25 | 97.24 | 1.15 | 0 | 1.97 | 1.15 | 0.2105 | 0.2336 | 0.0231 |
| 3rpz | opls | 1.51 | 0.0998 | 2.6 | 0 | 98.18 | 0.44 | 1 | 2.5 | 1.03 | 0.1348 | 0.1521 | 0.0173 |
| 3rpz | noschrodinger | 1.51 | 0.0145 | 1.52 | 0 | 98.18 | 0.44 | 0 | 1.36 | 0.87 | 0.1364 | 0.1533 | 0.0169 |
| 3rq0 | opls | 2.02 | 0.0966 | 2.36 | 0 | 95.95 | 0 | 0 | 1.86 | 1.23 | 0.1538 | 0.1809 | 0.0271 |
| 3rq0 | noschrodinger | 2.02 | 0.0026 | 0.8 | 0 | 95.18 | 0.57 | 0 | 0.8 | 1.09 | 0.1628 | 0.1824 | 0.0196 |
| 3rt3 | opls | 2.01 | 0.0948 | 2.26 | 0.83 | 98.76 | 5.61 | 0 | 1.01 | 1.37 | 0.2135 | 0.2454 | 0.0319 |
| 3rt3 | noschrodinger | 2.01 | 0.0027 | 0.76 | 0 | 97.94 | 6.94 | 0 | 2.27 | 1.66 | 0.1997 | 0.2394 | 0.0397 |
| 3ryk | opls | 1.63 | 0.0993 | 2.51 | 0 | 97.38 | 0.95 | 9 | 2.07 | 1.1 | 0.2464 | 0.2901 | 0.0437 |
| 3ryk | noschrodinger | 1.63 | 0.014 | 1.45 | 0 | 96.82 | 0.63 | 0 | 4.81 | 1.44 | 0.247 | 0.2904 | 0.0434 |
| 3ryt | opls | 3.58 | 0.085 | 2.28 | 0.9 | 96.26 | 2.92 | 7 | 0.88 | 1.37 | 0.288 | 0.3318 | 0.0438 |
| 3ryt | noschrodinger | 3.58 | 0.0029 | 0.78 | 1.25 | 89.57 | 3.14 | 0 | 2.48 | 1.97 | 0.2632 | 0.3105 | 0.0473 |
| 3rzf | opls | 3.9 | 0.0902 | 2.64 | 6.68 | 75.25 | 10.62 | 21 | 5.63 | 2.88 | 0.2676 | 0.3158 | 0.0482 |
| 3rzf | noschrodinger | 3.9 | 0.0039 | 1 | 8.13 | 66.92 | 13.14 | 1 | 14.35 | 3.38 | 0.27 | 0.3132 | 0.0432 |
| 3s0e | opls | 1.6 | 0.1024 | 2.33 | 0 | 100 | 1.77 | 1 | 1.54 | 1.09 | 0.1639 | 0.2104 | 0.0465 |
| 3s0e | noschrodinger | 1.6 | 0.0036 | 0.82 | 0 | 100 | 0.88 | 0 | 3.08 | 1.1 | 0.1725 | 0.2069 | 0.0344 |
| 3s4o | opls | 2.16 | 0.0935 | 2.31 | 0.96 | 96.82 | 2.17 | 0 | 1.34 | 1.31 | 0.2163 | 0.2541 | 0.0378 |
| 3s4o | noschrodinger | 2.16 | 0.0033 | 0.87 | 1.24 | 96.27 | 5.65 | 0 | 3.06 | 1.92 | 0.1998 | 0.2389 | 0.0391 |
| 3s5r | opls | 2.6 | 0.0865 | 2.28 | 0.38 | 97.73 | 0.69 | 3 | 0.82 | 0.82 | 0.2756 | 0.302 | 0.0264 |
| 3s5r | noschrodinger | 2.6 | 0.003 | 0.76 | 0.26 | 97.69 | 0 | 0 | 0.5 | 0.74 | 0.2796 | 0.2974 | 0.0178 |
| 3s7a | opls | 1.8 | 0.0901 | 2.35 | 0 | 97.81 | 4.76 | 2 | 1.58 | 1.46 | 0.2185 | 0.2575 | 0.039 |
| 3s7a | noschrodinger | 1.8 | 0.0039 | 0.96 | 0 | 97.83 | 8.28 | 0 | 1.9 | 1.69 | 0.2233 | 0.2553 | 0.032 |
| 3s7z | opls | 2 | 0.0958 | 2.31 | 0 | 97.35 | 0.52 | 0 | 1.29 | 0.98 | 0.1466 | 0.1864 | 0.0398 |
| 3s7z | noschrodinger | 2 | 0.003 | 0.87 | 0 | 98.92 | 0.5 | 0 | 2.32 | 1.01 | 0.1463 | 0.1835 | 0.0372 |
| 3s8x | opls | 1.3 | 0.1003 | 2.58 | 0 | 97.25 | 1.72 | 0 | 3.94 | 1.5 | 0.1334 | 0.1449 | 0.0115 |
| 3s8x | noschrodinger | 1.3 | 0.0088 | 1.28 | 0 | 97.25 | 1.29 | 0 | 2.77 | 1.29 | 0.1344 | 0.1451 | 0.0107 |
| 3sbu | opls | 2.16 | 0.0891 | 2.26 | 0 | 97.71 | 0.26 | 1 | 1.14 | 0.89 | 0.2122 | 0.2314 | 0.0192 |
| 3sbu | noschrodinger | 2.16 | 0.0128 | 0.78 | 0 | 97.87 | 1.52 | 0 | 1.67 | 1.08 | 0.2063 | 0.2288 | 0.0225 |
| 3sch | opls | 2.1 | 0.0937 | 2.36 | 0 | 97.94 | 2.73 | 4 | 1.56 | 1.25 | 0.1884 | 0.2241 | 0.0357 |
| 3sch | noschrodinger | 2.1 | 0.0041 | 0.86 | 0 | 96.34 | 2.99 | 0 | 3.14 | 1.71 | 0.1968 | 0.2314 | 0.0346 |
| 3sgi | opls | 3.5 | 0.0909 | 2.64 | 5.38 | 83.85 | 12.23 | 17 | 2.83 | 2.58 | 0.329 | 0.3819 | 0.0529 |
| 3sgi | noschrodinger | 3.5 | 0.0039 | 1.09 | 8.62 | 76.85 | 22.26 | 0 | 18.18 | 3.56 | 0.3164 | 0.3445 | 0.0281 |
| 3sgj | opls | 2.15 | 0.0943 | 2.31 | 0 | 99.33 | 3.79 | 3 | 0.59 | 1.14 | 0.2159 | 0.2501 | 0.0342 |
| 3sgj | noschrodinger | 2.15 | 0.0042 | 0.87 | 0 | 98.33 | 5.22 | 0 | 5.34 | 1.83 | 0.2084 | 0.2383 | 0.0299 |
| 3sgv | opls | 1.61 | 0.0989 | 2.31 | 0.47 | 98.12 | 1.67 | 0 | 2.08 | 1.15 | 0.1769 | 0.2047 | 0.0278 |
| 3sgv | noschrodinger | 1.61 | 0.0037 | 0.89 | 0.92 | 97.23 | 1.93 | 0 | 3.75 | 1.52 | 0.1809 | 0.2029 | 0.022 |
| 3sh7 | opls | 2.5 | 0.0998 | 2.47 | 1.58 | 94.86 | 3.49 | 8 | 1.36 | 1.63 | 0.1861 | 0.2259 | 0.0398 |
| 3sh7 | noschrodinger | 2.5 | 0.0033 | 0.92 | 2.36 | 94.49 | 5.79 | 0 | 4.57 | 2.19 | 0.1723 | 0.217 | 0.0447 |
| 3shf | opls | 3.55 | 0.0914 | 2.41 | 2.67 | 86.49 | 5.98 | 13 | 1.45 | 2.1 | 0.2373 | 0.2639 | 0.0266 |
| 3shf | noschrodinger | 3.55 | 0.0029 | 0.84 | 6.92 | 73.38 | 8.76 | 0 | 9.58 | 3.03 | 0.2023 | 0.2375 | 0.0352 |
| 3shr | opls | 2.44 | 0.088 | 2.25 | 0.45 | 96.44 | 3.7 | 2 | 0.7 | 1.39 | 0.2423 | 0.2719 | 0.0296 |
| 3shr | noschrodinger | 2.44 | 0.003 | 0.96 | 2.54 | 90.22 | 4.86 | 0 | 3.58 | 2.21 | 0.2387 | 0.2683 | 0.0296 |
| 3smj | opls | 1.5 | 0.0989 | 2.53 | 0 | 98.07 | 0 | 2 | 1.51 | 0.89 | 0.1642 | 0.1896 | 0.0254 |
| 3smj | noschrodinger | 1.5 | 0.0071 | 1.05 | 0 | 97.3 | 0.61 | 1 | 1.01 | 0.93 | 0.1712 | 0.1913 | 0.0201 |
| 3soa | opls | 3.55 | 0.0906 | 2.32 | 0.7 | 93.49 | 1.34 | 2 | 1.98 | 1.49 | 0.2644 | 0.3077 | 0.0433 |
| 3soa | noschrodinger | 3.55 | 0.0032 | 0.83 | 0.93 | 89.58 | 3.74 | 0 | 9.65 | 2.5 | 0.2674 | 0.3149 | 0.0475 |
| 3sp8 | opls | 1.86 | 0.0971 | 2.22 | 0.2 | 96.58 | 1.02 | 3 | 2.96 | 1.31 | 0.1953 | 0.2248 | 0.0295 |
| 3sp8 | noschrodinger | 1.86 | 0.006 | 0.9 | 0.2 | 95.46 | 2.44 | 0 | 4.89 | 1.87 | 0.1943 | 0.2264 | 0.0321 |

| | | | | | | | | | | | | | |
|---|---|---|---|---|---|---|---|---|---|---|---|---|---|
| 3st4 | opls | 2 | 0.0982 | 2.26 | 0 | 97.94 | 1.75 | 1 | 1.39 | 1.07 | 0.1881 | 0.2203 | 0.0322 |
| 3st4 | noschrodinger | 2 | 0.0035 | 0.85 | 0 | 97.95 | 2.69 | 0 | 4.08 | 1.53 | 0.1773 | 0.2088 | 0.0315 |
| 3svl | opls | 2.19 | 0.0932 | 2.28 | 0.28 | 98.04 | 2.36 | 0 | 0 | 0.78 | 0.2239 | 0.252 | 0.0281 |
| 3svl | noschrodinger | 2.19 | 0.0027 | 0.81 | 0.56 | 96.1 | 2.69 | 0 | 2.49 | 1.63 | 0.2178 | 0.2467 | 0.0289 |
| 3sy6 | opls | 1.9 | 0.0973 | 2.2 | 0 | 96.49 | 0.92 | 0 | 1.34 | 1.09 | 0.1802 | 0.22 | 0.0398 |
| 3sy6 | noschrodinger | 1.9 | 0.0031 | 0.79 | 0 | 97.22 | 0.68 | 0 | 1.59 | 1.05 | 0.1905 | 0.2155 | 0.025 |
| 3sz4 | opls | 2.59 | 0.0905 | 2.29 | 0 | 99.46 | 2.52 | 0 | 0.32 | 0.92 | 0.2344 | 0.2592 | 0.0248 |
| 3sz4 | noschrodinger | 2.59 | 0.0032 | 0.8 | 0 | 96.83 | 6.83 | 0 | 2.93 | 1.91 | 0.2343 | 0.2627 | 0.0284 |
| 3t0w | opls | 1.5 | 0.1043 | 2.39 | 0 | 95.33 | 0.49 | 1 | 5.91 | 1.65 | 0.1549 | 0.1853 | 0.0304 |
| 3t0w | noschrodinger | 1.5 | 0.006 | 1.07 | 0 | 95.83 | 0.49 | 0 | 4.79 | 1.54 | 0.1562 | 0.1798 | 0.0236 |
| 3t1k | opls | 1.5 | 0.0998 | 2.35 | 0.48 | 96.2 | 3.27 | 0 | 4.09 | 1.84 | 0.1515 | 0.1654 | 0.0139 |
| 3t1k | noschrodinger | 1.5 | 0.0054 | 1.04 | 0.24 | 95.27 | 4.07 | 0 | 2.49 | 1.82 | 0.1509 | 0.1773 | 0.0264 |
| 3t1q | opls | 2.7 | 0.092 | 2.29 | 0 | 97.68 | 4.08 | 3 | 0.72 | 1.26 | 0.2043 | 0.2437 | 0.0394 |
| 3t1q | noschrodinger | 2.7 | 0.004 | 1 | 0 | 96.55 | 7.49 | 0 | 1.86 | 1.84 | 0.1894 | 0.2362 | 0.0468 |
| 3t38 | opls | 2.2 | 0.092 | 2.35 | 0.56 | 97.49 | 2.51 | 0 | 0.99 | 1.2 | 0.1925 | 0.2314 | 0.0389 |
| 3t38 | noschrodinger | 2.2 | 0.0034 | 0.84 | 0.79 | 95.51 | 4.57 | 0 | 1.48 | 1.7 | 0.1788 | 0.2218 | 0.043 |
| 3t4l | opls | 1.53 | 0.1017 | 2.38 | 0 | 98.43 | 0.81 | 5 | 4.06 | 1.19 | 0.1836 | 0.2189 | 0.0353 |
| 3t4l | noschrodinger | 1.53 | 0.0034 | 0.82 | 0 | 98.27 | 0.81 | 0 | 1.92 | 0.96 | 0.1929 | 0.2125 | 0.0196 |
| 3t4n | opls | 2.3 | 0.0965 | 2.29 | 0 | 98.73 | 2.07 | 4 | 1.47 | 1.13 | 0.1848 | 0.2224 | 0.0376 |
| 3t4n | noschrodinger | 2.3 | 0.0027 | 0.8 | 0.37 | 97.22 | 2.04 | 0 | 1.83 | 1.32 | 0.1908 | 0.2222 | 0.0314 |
| 3t4q | opls | 2.3 | 0.0994 | 2.22 | 0 | 98.43 | 1.05 | 0 | 1.04 | 0.82 | 0.1935 | 0.2203 | 0.0268 |
| 3t4q | noschrodinger | 2.3 | 0.0025 | 0.72 | 0 | 97.89 | 0.62 | 0 | 1.04 | 0.83 | 0.1783 | 0.21 | 0.0317 |
| 3t50 | opls | 1.64 | 0.097 | 2.26 | 0 | 99.01 | 0 | 0 | 3.09 | 1.1 | 0.1813 | 0.2107 | 0.0294 |
| 3t50 | noschrodinger | 1.64 | 0.0038 | 0.92 | 0 | 99.54 | 0 | 0 | 1.69 | 0.92 | 0.1861 | 0.2106 | 0.0245 |
| 3t91 | opls | 2.6 | 0.099 | 2.75 | 2.7 | 93.38 | 10.7 | 4 | 8.24 | 2.66 | 0.2084 | 0.2597 | 0.0513 |
| 3t91 | noschrodinger | 2.6 | 0.0047 | 1.05 | 2.17 | 92.77 | 12.93 | 0 | 12.88 | 2.92 | 0.2312 | 0.2621 | 0.0309 |
| 3tb0 | opls | 2 | 0.1033 | 2.22 | 0.63 | 96.84 | 3.33 | 1 | 3.38 | 1.72 | 0.1759 | 0.2019 | 0.026 |
| 3tb0 | noschrodinger | 2 | 0.0036 | 0.87 | 0.63 | 96.23 | 2.65 | 0 | 2.27 | 1.58 | 0.159 | 0.1938 | 0.0348 |
| 3tb6 | opls | 2.21 | 0.0956 | 2.3 | 0.36 | 97.14 | 6.05 | 5 | 1.09 | 1.56 | 0.2079 | 0.2258 | 0.0179 |
| 3tb6 | noschrodinger | 2.21 | 0.0032 | 1.21 | 0.71 | 96.44 | 8.75 | 0 | 3.04 | 2.05 | 0.1948 | 0.2265 | 0.0317 |
| 3td7 | opls | 2.21 | 0.096 | 2.41 | 0 | 97.56 | 1.77 | 2 | 0.7 | 1.01 | 0.1971 | 0.2223 | 0.0252 |
| 3td7 | noschrodinger | 2.21 | 0.0026 | 0.71 | 0 | 97.59 | 1.76 | 0 | 0.71 | 1 | 0.2122 | 0.2271 | 0.0149 |
| 3tdg | opls | 2.1 | 0.0963 | 2.22 | 0 | 98.25 | 1.45 | 0 | 1.32 | 0.98 | 0.2098 | 0.2276 | 0.0178 |
| 3tdg | noschrodinger | 2.1 | 0.0041 | 0.84 | 0 | 99.13 | 1.44 | 0 | 3.17 | 1.23 | 0.2023 | 0.2188 | 0.0165 |
| 3te5 | opls | 2.5 | 0.0943 | 2.26 | 0 | 98.57 | 2.48 | 4 | 0.45 | 0.96 | 0.2331 | 0.2569 | 0.0238 |
| 3te5 | noschrodinger | 2.5 | 0.0028 | 0.79 | 0.37 | 96.85 | 4.69 | 0 | 2.15 | 1.69 | 0.2114 | 0.2432 | 0.0318 |
| 3tfp | opls | 1.93 | 0.0976 | 2.22 | 0 | 98.93 | 0.39 | 1 | 1.26 | 0.85 | 0.1895 | 0.2247 | 0.0352 |
| 3tfp | noschrodinger | 1.93 | 0.0052 | 0.95 | 0 | 97.87 | 1.55 | 0 | 4.83 | 1.43 | 0.1807 | 0.2219 | 0.0412 |
| 3thm | opls | 2.1 | 0.0976 | 2.19 | 0 | 97.8 | 0.68 | 0 | 0.89 | 0.82 | 0.1984 | 0.228 | 0.0296 |
| 3thm | noschrodinger | 2.1 | 0.0027 | 0.79 | 0 | 97.64 | 0.22 | 0 | 1.53 | 0.97 | 0.1903 | 0.2175 | 0.0272 |
| 3tkl | opls | 2.18 | 0.094 | 2.28 | 0.28 | 98.04 | 5 | 2 | 1.36 | 1.4 | 0.1826 | 0.2324 | 0.0498 |
| 3tkl | noschrodinger | 2.18 | 0.0027 | 0.73 | 0.28 | 98.33 | 4.97 | 0 | 1.71 | 1.45 | 0.1893 | 0.2261 | 0.0368 |
| 3tm0 | opls | 2.1 | 0.0968 | 2.29 | 0.38 | 97.31 | 2.49 | 1 | 1.37 | 1.3 | 0.1874 | 0.2229 | 0.0355 |
| 3tm0 | noschrodinger | 2.1 | 0.0032 | 0.97 | 0 | 98.08 | 2.49 | 0 | 2.07 | 1.28 | 0.1854 | 0.2176 | 0.0322 |
| 3tnj | opls | 2 | 0.0919 | 2.34 | 0 | 100 | 1.89 | 0 | 0 | 0.71 | 0.1901 | 0.2372 | 0.0471 |
| 3tnj | noschrodinger | 2 | 0.0027 | 0.75 | 0 | 100 | 0.93 | 0 | 1 | 0.79 | 0.1981 | 0.2371 | 0.039 |
| 3to6 | opls | 2.1 | 0.0995 | 2.33 | 0 | 97.77 | 2.33 | 3 | 0.42 | 0.98 | 0.1839 | 0.2033 | 0.0194 |
| 3to6 | noschrodinger | 2.1 | 0.0032 | 0.85 | 0 | 98.15 | 3.46 | 0 | 1.46 | 1.29 | 0.1779 | 0.1947 | 0.0168 |
| 3tri | opls | 2.4 | 0.0921 | 2.38 | 0 | 97.6 | 1.15 | 1 | 1.29 | 0.98 | 0.201 | 0.231 | 0.03 |

| | | | | | | | | | | | | | |
|---|---|---|---|---|---|---|---|---|---|---|---|---|---|
| 3tri | noschrodinger | 2.4 | 0.0029 | 0.76 | 0.37 | 96.84 | 1.34 | 0 | 4.23 | 1.49 | 0.1896 | 0.2197 | 0.0301 |
| 3tta | opls | 2 | 0.0976 | 2.31 | 0 | 100 | 2.25 | 1 | 1.15 | 1.09 | 0.1782 | 0.217 | 0.0388 |
| 3tta | noschrodinger | 2 | 0.0028 | 0.88 | 0 | 99.1 | 4.44 | 0 | 1.76 | 1.42 | 0.1867 | 0.2096 | 0.0229 |
| 3tu5 | opls | 3 | 0.092 | 2.3 | 0 | 97.78 | 4.28 | 3 | 0.5 | 1.2 | 0.1886 | 0.2252 | 0.0366 |
| 3tu5 | noschrodinger | 3 | 0.003 | 0.84 | 0.2 | 96.79 | 10.17 | 0 | 1.51 | 1.86 | 0.2103 | 0.2388 | 0.0285 |
| 3tvk | opls | 1.8 | 0.0973 | 2.33 | 0 | 98.81 | 0 | 0 | 1.02 | 0.8 | 0.1596 | 0.197 | 0.0374 |
| 3tvk | noschrodinger | 1.8 | 0.0048 | 1.11 | 0 | 99.41 | 1.32 | 0 | 2.39 | 1.11 | 0.1634 | 0.197 | 0.0336 |
| 3tvq | opls | 1.67 | 0.101 | 2.24 | 0.68 | 97.97 | 0.79 | 0 | 1.22 | 0.85 | 0.1773 | 0.2095 | 0.0322 |
| 3tvq | noschrodinger | 1.67 | 0.0098 | 1.07 | 0.67 | 98.67 | 0.78 | 0 | 0 | 0.5 | 0.1766 | 0.2135 | 0.0369 |
| 3tx2 | opls | 1.5 | 0.1055 | 2.4 | 0 | 97.84 | 1.58 | 2 | 4.79 | 1.44 | 0.1577 | 0.1975 | 0.0398 |
| 3tx2 | noschrodinger | 1.5 | 0.0029 | 0.87 | 0 | 97.55 | 0.53 | 0 | 1.07 | 0.9 | 0.1516 | 0.1912 | 0.0396 |
| 3tx3 | opls | 2.3 | 0.0918 | 2.31 | 0 | 98.19 | 0.77 | 0 | 1.23 | 0.84 | 0.2035 | 0.226 | 0.0225 |
| 3tx3 | noschrodinger | 2.3 | 0.0027 | 0.79 | 0.45 | 97.98 | 1.26 | 0 | 1.99 | 1.05 | 0.1999 | 0.2252 | 0.0253 |
| 3u11 | opls | 2.5 | 0.0947 | 2.3 | 0.26 | 97.42 | 4.14 | 0 | 1.61 | 1.49 | 0.217 | 0.2455 | 0.0285 |
| 3u11 | noschrodinger | 2.5 | 0.0033 | 0.82 | 0.51 | 96.41 | 6.87 | 0 | 6.02 | 2.2 | 0.1974 | 0.227 | 0.0296 |
| 3u2v | opls | 1.5 | 0.1007 | 2.45 | 0 | 98.22 | 0.63 | 0 | 3.16 | 1.11 | 0.1791 | 0.2151 | 0.036 |
| 3u2v | noschrodinger | 1.5 | 0.0035 | 0.91 | 0 | 97.51 | 0.63 | 0 | 3.06 | 1.2 | 0.186 | 0.2154 | 0.0294 |
| 3u3u | opls | 2.5 | 0.0947 | 2.29 | 0.44 | 97.82 | 0 | 1 | 1.62 | 0.95 | 0.2229 | 0.2645 | 0.0416 |
| 3u3u | noschrodinger | 2.5 | 0.0034 | 0.86 | 0.43 | 96.96 | 1.02 | 0 | 3.51 | 1.33 | 0.2022 | 0.2597 | 0.0575 |
| 3u5s | opls | 1.5 | 0.0991 | 2.58 | 0 | 97.92 | 1.96 | 2 | 5.67 | 1.55 | 0.1745 | 0.2072 | 0.0327 |
| 3u5s | noschrodinger | 1.5 | 0.0125 | 1.31 | 1.83 | 94.5 | 2.83 | 0 | 4.83 | 1.97 | 0.1653 | 0.1938 | 0.0285 |
| 3u9e | opls | 2.04 | 0.0925 | 2.29 | 0 | 97.61 | 1.53 | 0 | 0.65 | 0.94 | 0.1968 | 0.2253 | 0.0285 |
| 3u9e | noschrodinger | 2.04 | 0.0028 | 0.77 | 0.18 | 96.47 | 3.42 | 0 | 1.42 | 1.52 | 0.191 | 0.2196 | 0.0286 |
| 3ubp | opls | 2 | 0.1017 | 2.29 | 0.27 | 96.14 | 1.58 | 4 | 1.58 | 1.32 | 0.1583 | 0.1898 | 0.0315 |
| 3ubp | noschrodinger | 2 | 0.003 | 0.84 | 0.38 | 95.66 | 2.63 | 0 | 1.83 | 1.56 | 0.1511 | 0.173 | 0.0219 |
| 3ucj | opls | 1.85 | 0.0998 | 2.31 | 0 | 97.5 | 0.82 | 0 | 1.76 | 1.03 | 0.1764 | 0.206 | 0.0296 |
| 3ucj | noschrodinger | 1.85 | 0.0067 | 0.99 | 0 | 97.95 | 1.65 | 0 | 1.61 | 1.08 | 0.1751 | 0.2041 | 0.029 |
| 3uda | opls | 2.51 | 0.0914 | 2.47 | 1.07 | 95.71 | 6.95 | 3 | 2.12 | 1.92 | 0.2042 | 0.2556 | 0.0514 |
| 3uda | noschrodinger | 2.51 | 0.003 | 0.83 | 1.86 | 91.76 | 10.18 | 0 | 5.07 | 2.53 | 0.2086 | 0.2472 | 0.0386 |
| 3ueh | opls | 2.59 | 0.0874 | 2.23 | 0 | 97.93 | 0.48 | 2 | 0.23 | 0.61 | 0.2218 | 0.2505 | 0.0287 |
| 3ueh | noschrodinger | 2.59 | 0.0029 | 0.8 | 0 | 96.67 | 0.96 | 0 | 0.24 | 0.8 | 0.2088 | 0.2394 | 0.0306 |
| 3uf6 | opls | 1.8 | 0.098 | 2.27 | 0.18 | 97.62 | 1.49 | 0 | 1.23 | 1.05 | 0.1604 | 0.1855 | 0.0251 |
| 3uf6 | noschrodinger | 1.8 | 0.0042 | 0.87 | 0 | 96.65 | 1.05 | 0 | 1.78 | 1.17 | 0.1625 | 0.1848 | 0.0223 |
| 3uf8 | opls | 1.5 | 0.1028 | 2.17 | 0 | 99.41 | 0.62 | 0 | 2.25 | 1 | 0.1494 | 0.1818 | 0.0324 |
| 3uf8 | noschrodinger | 1.5 | 0.0105 | 1.19 | 0 | 99.47 | 0.62 | 0 | 2.26 | 1 | 0.1466 | 0.1821 | 0.0355 |
| 3ufo | opls | 2.17 | 0.0925 | 2.28 | 0 | 97.4 | 1.37 | 5 | 0.52 | 0.9 | 0.2306 | 0.2508 | 0.0202 |
| 3ufo | noschrodinger | 2.17 | 0.0042 | 0.9 | 0 | 96.05 | 3.68 | 0 | 2.14 | 1.69 | 0.2155 | 0.2436 | 0.0281 |
| 3uj0 | opls | 3.6 | 0.0884 | 2.35 | 0.26 | 92.86 | 6.6 | 10 | 2.18 | 2.07 | 0.2568 | 0.3026 | 0.0458 |
| 3uj0 | noschrodinger | 3.6 | 0.0135 | 0.94 | 0.72 | 86.29 | 11.22 | 0 | 3.27 | 2.55 | 0.2504 | 0.2851 | 0.0347 |
| 3ul6 | opls | 2.63 | 0.0879 | 2.23 | 0 | 98.18 | 2.68 | 1 | 0.81 | 1.08 | 0.2475 | 0.277 | 0.0295 |
| 3ul6 | noschrodinger | 2.63 | 0.0031 | 0.73 | 0 | 94.19 | 5.31 | 0 | 1.25 | 1.79 | 0.2585 | 0.2736 | 0.0151 |
| 3up1 | opls | 2.15 | 0.0926 | 2.37 | 0 | 94.86 | 1.65 | 2 | 0.74 | 1.26 | 0.2263 | 0.2459 | 0.0196 |
| 3up1 | noschrodinger | 2.15 | 0.0033 | 0.84 | 0.51 | 93.89 | 5.48 | 0 | 4.63 | 2.2 | 0.2143 | 0.2399 | 0.0256 |
| 3urg | opls | 2 | 0.0966 | 2.23 | 0.9 | 98.2 | 2.86 | 1 | 1.52 | 1.24 | 0.233 | 0.2516 | 0.0186 |
| 3urg | noschrodinger | 2 | 0.0032 | 0.76 | 0.84 | 97.48 | 5.66 | 0 | 1.01 | 1.48 | 0.2178 | 0.2445 | 0.0267 |
| 3ust | opls | 2.1 | 0.0943 | 2.21 | 0 | 98.48 | 0.93 | 1 | 1.17 | 0.83 | 0.1981 | 0.216 | 0.0179 |
| 3ust | noschrodinger | 2.1 | 0.0038 | 0.75 | 0 | 97.94 | 1.28 | 0 | 0.47 | 0.76 | 0.1789 | 0.2134 | 0.0345 |
| 3ux4 | opls | 3.26 | 0.0883 | 2.64 | 1.72 | 90.61 | 2.7 | 17 | 2.63 | 1.91 | 0.363 | 0.384 | 0.021 |
| 3ux4 | noschrodinger | 3.26 | 0.0032 | 0.84 | 0.57 | 87.12 | 3.56 | 0 | 9.53 | 2.54 | 0.3491 | 0.3749 | 0.0258 |

| | | | | | | | | | | | | | |
|---|---|---|---|---|---|---|---|---|---|---|---|---|---|
| 3v4e | opls | 1.95 | 0.0932 | 2.34 | 0 | 97.57 | 0.42 | 0 | 1.68 | 1.01 | 0.1863 | 0.2198 | 0.0335 |
| 3v4e | noschrodinger | 1.95 | 0.0029 | 0.79 | 0 | 97.67 | 0.85 | 0 | 2.25 | 1.07 | 0.1892 | 0.2139 | 0.0247 |
| 3vaa | opls | 1.7 | 0.1002 | 2.34 | 0 | 98.44 | 0.43 | 3 | 1.91 | 0.96 | 0.1701 | 0.1982 | 0.0281 |
| 3vaa | noschrodinger | 1.7 | 0.0107 | 1.08 | 0.19 | 98.25 | 0.21 | 1 | 2.02 | 0.97 | 0.1735 | 0.1975 | 0.024 |
| 3vae | opls | 2.8 | 0.0934 | 2.25 | 0.2 | 96.85 | 2.31 | 0 | 0.75 | 1.21 | 0.1945 | 0.2265 | 0.032 |
| 3vae | noschrodinger | 2.8 | 0.0028 | 0.81 | 0.2 | 94.92 | 5.06 | 0 | 1.25 | 1.73 | 0.1929 | 0.2206 | 0.0277 |
| 3vbj | opls | 1.8 | 0.0982 | 2.39 | 0 | 97.64 | 1.86 | 6 | 3.11 | 1.38 | 0.1771 | 0.2068 | 0.0297 |
| 3vbj | noschrodinger | 1.8 | 0.006 | 1.03 | 0.18 | 96.21 | 3.3 | 0 | 3.64 | 1.8 | 0.1744 | 0.2063 | 0.0319 |
| 3vbk | opls | 2.2 | 0.0924 | 2.37 | 0 | 97.81 | 2.78 | 2 | 1.41 | 1.25 | 0.1965 | 0.2139 | 0.0174 |
| 3vbk | noschrodinger | 2.2 | 0.0027 | 0.78 | 0 | 95.83 | 5.11 | 0 | 2.06 | 1.8 | 0.1953 | 0.2174 | 0.0221 |
| 3vbn | opls | 2.5 | 0.0909 | 2.31 | 0 | 97.1 | 1.28 | 0 | 0.66 | 0.96 | 0.204 | 0.2429 | 0.0389 |
| 3vbn | noschrodinger | 2.5 | 0.0029 | 0.78 | 0 | 91.86 | 5.11 | 0 | 2.73 | 2.09 | 0.1982 | 0.232 | 0.0338 |
| 3vka | opls | 1.57 | 0.1007 | 2.34 | 0.4 | 97.6 | 1.72 | 0 | 3.16 | 1.37 | 0.1426 | 0.1764 | 0.0338 |
| 3vka | noschrodinger | 1.57 | 0.0071 | 1.01 | 0.4 | 96.61 | 2.45 | 0 | 2.65 | 1.57 | 0.1491 | 0.1748 | 0.0257 |
| 3vkh | opls | 3.79 | 0.0886 | 2.37 | 0.68 | 94.73 | 5.11 | 31 | 2.24 | 1.9 | 0.2071 | 0.2569 | 0.0498 |
| 3vkh | noschrodinger | 3.79 | 0.0028 | 0.74 | 2.43 | 84.81 | 6.78 | 0 | 6.48 | 2.65 | 0.1999 | 0.2411 | 0.0412 |
| 3vkl | opls | 2.54 | 0.094 | 2.29 | 0.35 | 97.74 | 3.12 | 0 | 0.93 | 1.22 | 0.1975 | 0.2496 | 0.0521 |
| 3vkl | noschrodinger | 2.54 | 0.0028 | 0.82 | 1.05 | 95.82 | 5.08 | 0 | 4.87 | 2.08 | 0.1964 | 0.2472 | 0.0508 |
| 3vmr | opls | 3.69 | 0.0894 | 2.74 | 1.36 | 90.95 | 16.1 | 10 | 4.17 | 2.64 | 0.3729 | 0.4291 | 0.0562 |
| 3vmr | noschrodinger | 3.69 | 0.0034 | 1.1 | 2.69 | 84.3 | 23.19 | 0 | 22.46 | 3.55 | 0.3687 | 0.4041 | 0.0354 |
| 3vmt | opls | 2.3 | 0.0922 | 2.36 | 0.47 | 96.5 | 7.05 | 8 | 1.38 | 1.74 | 0.2319 | 0.2565 | 0.0246 |
| 3vmt | noschrodinger | 2.3 | 0.003 | 0.79 | 0 | 94.21 | 8.23 | 0 | 3.32 | 2.21 | 0.2172 | 0.2457 | 0.0285 |
| 3vpd | opls | 1.95 | 0.0992 | 2.4 | 0.19 | 97.96 | 5.79 | 5 | 1.26 | 1.44 | 0.1677 | 0.2037 | 0.036 |
| 3vpd | noschrodinger | 1.95 | 0.0033 | 0.9 | 0.18 | 98.16 | 6.9 | 0 | 2.17 | 1.63 | 0.1749 | 0.1959 | 0.021 |
| 3vqh | opls | 1.95 | 0.0958 | 2.24 | 0 | 97.73 | 1.27 | 3 | 1.63 | 1.05 | 0.173 | 0.1969 | 0.0239 |
| 3vqh | noschrodinger | 1.95 | 0.0031 | 0.8 | 0 | 96.89 | 2.53 | 0 | 0.98 | 1.28 | 0.1768 | 0.1955 | 0.0187 |
| 3vv4 | opls | 1.95 | 0.0935 | 2.51 | 0 | 98.37 | 8.18 | 8 | 1.54 | 1.59 | 0.1847 | 0.2145 | 0.0298 |
| 3vv4 | noschrodinger | 1.95 | 0.0094 | 1.2 | 0 | 98.06 | 10.7 | 0 | 4.25 | 1.99 | 0.188 | 0.2163 | 0.0283 |
| 3vx3 | opls | 2.1 | 0.0943 | 2.3 | 0.22 | 98.01 | 1.77 | 2 | 1.17 | 1.02 | 0.2304 | 0.2537 | 0.0233 |
| 3vx3 | noschrodinger | 2.1 | 0.0028 | 0.85 | 0.22 | 97.6 | 3.48 | 0 | 1.7 | 1.42 | 0.2176 | 0.2452 | 0.0276 |
| 3vya | opls | 2.4 | 0.0941 | 2.32 | 0 | 98.31 | 2.15 | 0 | 0 | 0.75 | 0.2043 | 0.2535 | 0.0492 |
| 3vya | noschrodinger | 2.4 | 0.0026 | 0.74 | 0 | 98.31 | 4.3 | 0 | 1.07 | 1.29 | 0.1844 | 0.2389 | 0.0545 |
| 3vyf | opls | 2.8 | 0.0905 | 2.23 | 0 | 98.33 | 1.59 | 3 | 0.28 | 0.76 | 0.2103 | 0.2355 | 0.0252 |
| 3vyf | noschrodinger | 2.8 | 0.0029 | 0.83 | 0.15 | 97.74 | 3.33 | 0 | 1.32 | 1.31 | 0.2045 | 0.2373 | 0.0328 |
| 3w34 | opls | 1.91 | 0.0971 | 2.62 | 0 | 99.01 | 2.36 | 2 | 2.37 | 1.3 | 0.1982 | 0.2273 | 0.0291 |
| 3w34 | noschrodinger | 1.91 | 0.0035 | 0.89 | 0 | 98.53 | 1.76 | 0 | 4.9 | 1.44 | 0.1973 | 0.2261 | 0.0288 |
| 3w44 | opls | 2.3 | 0.0976 | 2.27 | 0 | 98.13 | 3.2 | 0 | 1.28 | 1.23 | 0.2081 | 0.2452 | 0.0371 |
| 3w44 | noschrodinger | 2.3 | 0.0028 | 0.78 | 0.51 | 97.72 | 5.68 | 0 | 2.23 | 1.64 | 0.1814 | 0.234 | 0.0526 |
| 3w5j | opls | 1.93 | 0.0966 | 2.24 | 0 | 98.78 | 1.28 | 0 | 0.83 | 0.84 | 0.1793 | 0.2151 | 0.0358 |
| 3w5j | noschrodinger | 1.93 | 0.0029 | 0.81 | 0 | 98.94 | 0.95 | 0 | 0.84 | 0.76 | 0.1841 | 0.2137 | 0.0296 |
| 3w7r | opls | 1.68 | 0.0959 | 2.46 | 0.27 | 96.44 | 2.24 | 0 | 5.36 | 1.79 | 0.1439 | 0.1583 | 0.0144 |
| 3w7r | noschrodinger | 1.68 | 0.0134 | 1.4 | 0.27 | 96.16 | 1.92 | 1 | 8.92 | 1.95 | 0.1403 | 0.1574 | 0.0171 |
| 3w8r | opls | 2.49 | 0.0923 | 2.71 | 0.25 | 97.3 | 2.35 | 2 | 1.48 | 1.3 | 0.2245 | 0.2575 | 0.033 |
| 3w8r | noschrodinger | 2.49 | 0.003 | 0.81 | 0 | 96.09 | 2.35 | 0 | 4.44 | 1.77 | 0.2182 | 0.255 | 0.0368 |
| 3wal | opls | 2 | 0.0992 | 2.22 | 0 | 100 | 0 | 0 | 2.05 | 0.97 | 0.2085 | 0.2326 | 0.0241 |
| 3wal | noschrodinger | 2 | 0.005 | 0.78 | 0 | 99.12 | 3.67 | 0 | 1.54 | 1.33 | 0.196 | 0.2228 | 0.0268 |
| 3wcx | opls | 2 | 0.1007 | 2.36 | 0 | 98.95 | 1.32 | 1 | 2.5 | 1.13 | 0.1589 | 0.1993 | 0.0404 |
| 3wcx | noschrodinger | 2 | 0.0026 | 0.78 | 0 | 99.3 | 1.32 | 0 | 0.46 | 0.75 | 0.1574 | 0.1944 | 0.037 |
| 3wgh | opls | 2.05 | 0.0994 | 2.28 | 0 | 99.25 | 1.91 | 1 | 0.43 | 0.87 | 0.2027 | 0.2264 | 0.0237 |

| | | | | | | | | | | | |
|---|---|---|---|---|---|---|---|---|---|---|---|
| 3wgh | noschrodinger | 2.05 | 0.0031 | 0.8 | 0 | 99.01 | 4.07 | 0 | 2.17 | 1.45 | 0.1915 | 0.2179 | 0.0264 |
| 3wgp | opls | 2 | 0.1039 | 2.24 | 0 | 97.54 | 0.88 | 0 | 0.24 | 0.69 | 0.1684 | 0.2116 | 0.0432 |
| 3wgp | noschrodinger | 2 | 0.0047 | 0.97 | 0 | 97.97 | 1.32 | 0 | 1.46 | 0.98 | 0.1625 | 0.203 | 0.0405 |
| 3wgw | opls | 2.8 | 0.0929 | 2.3 | 0.21 | 96.69 | 3.04 | 3 | 0.77 | 1.32 | 0.2119 | 0.2412 | 0.0293 |
| 3wgw | noschrodinger | 2.8 | 0.0031 | 0.88 | 2.04 | 96.52 | 9.47 | 0 | 3.08 | 2.07 | 0.1969 | 0.2272 | 0.0303 |
| 3wnr | opls | 2.01 | 0.0999 | 2.21 | 0 | 98.51 | 0.34 | 0 | 1.85 | 0.95 | 0.1659 | 0.1899 | 0.024 |
| 3wnr | noschrodinger | 2.01 | 0.0029 | 0.79 | 0 | 97.92 | 0.34 | 0 | 0.56 | 0.71 | 0.1586 | 0.1806 | 0.022 |
| 3wo4 | opls | 3.09 | 0.0859 | 2.34 | 0.52 | 94.29 | 10.37 | 3 | 0.68 | 1.88 | 0.2329 | 0.2655 | 0.0326 |
| 3wo4 | noschrodinger | 3.09 | 0.0035 | 0.85 | 0.28 | 91.64 | 15.65 | 0 | 3 | 2.5 | 0.231 | 0.2626 | 0.0316 |
| 3wo6 | opls | 2.4 | 0.0852 | 2.23 | 0 | 99.33 | 1.89 | 1 | 0.58 | 0.91 | 0.2353 | 0.254 | 0.0187 |
| 3wo6 | noschrodinger | 2.4 | 0.0028 | 0.81 | 0 | 98.56 | 1.89 | 0 | 1.53 | 1.1 | 0.2285 | 0.2574 | 0.0289 |
| 3wqn | opls | 2.68 | 0.095 | 2.35 | 0.35 | 97.2 | 2.37 | 4 | 1.06 | 1.24 | 0.2435 | 0.2785 | 0.035 |
| 3wqn | noschrodinger | 2.68 | 0.0031 | 0.81 | 0 | 93.75 | 5.51 | 0 | 5.28 | 2.26 | 0.2286 | 0.2798 | 0.0512 |
| 3wv7 | opls | 1.6 | 0.0981 | 2.33 | 0.17 | 97.64 | 1.23 | 1 | 1.08 | 0.96 | 0.1676 | 0.2012 | 0.0336 |
| 3wv7 | noschrodinger | 1.6 | 0.0036 | 0.89 | 0.17 | 97.15 | 1.85 | 0 | 2.27 | 1.36 | 0.1738 | 0.1983 | 0.0245 |
| 3wvc | opls | 2 | 0.0977 | 2.3 | 0 | 97.9 | 0.34 | 0 | 0.71 | 0.75 | 0.1867 | 0.2148 | 0.0281 |
| 3wvc | noschrodinger | 2 | 0.0027 | 0.78 | 0.3 | 98.2 | 2.37 | 0 | 2.11 | 1.27 | 0.1684 | 0.1988 | 0.0304 |
| 3ww2 | opls | 2 | 0.0923 | 2.33 | 0.21 | 96.6 | 1.79 | 4 | 0.95 | 1.2 | 0.2057 | 0.2448 | 0.0391 |
| 3ww2 | noschrodinger | 2 | 0.0029 | 0.93 | 0.63 | 95.77 | 2.04 | 0 | 3.12 | 1.63 | 0.1987 | 0.2335 | 0.0348 |
| 3wwm | opls | 2.8 | 0.0918 | 2.31 | 0.75 | 97.74 | 3.33 | 1 | 1.19 | 1.29 | 0.2119 | 0.2345 | 0.0226 |
| 3wwm | noschrodinger | 2.8 | 0.003 | 0.89 | 0.75 | 96.63 | 5.19 | 0 | 0.95 | 1.54 | 0.2009 | 0.2283 | 0.0274 |
| 3wzs | opls | 1.7 | 0.0978 | 2.32 | 0 | 99.26 | 2.36 | 1 | 0.82 | 1.04 | 0.1977 | 0.2217 | 0.024 |
| 3wzs | noschrodinger | 1.7 | 0.0042 | 0.87 | 0 | 98.6 | 2.29 | 0 | 0 | 0.77 | 0.2013 | 0.2211 | 0.0198 |
| 3zc7 | opls | 2.1 | 0.1002 | 2.28 | 0.78 | 99.22 | 1.32 | 0 | 1.64 | 1 | 0.2034 | 0.2257 | 0.0223 |
| 3zc7 | noschrodinger | 2.1 | 0.0027 | 0.78 | 0.39 | 98.45 | 1.3 | 0 | 1.45 | 0.97 | 0.1733 | 0.2174 | 0.0441 |
| 3zet | opls | 2.31 | 0.0951 | 2.25 | 0.18 | 97.64 | 1.65 | 1 | 0.7 | 0.97 | 0.1833 | 0.2148 | 0.0315 |
| 3zet | noschrodinger | 2.31 | 0.0029 | 0.8 | 0 | 97.67 | 3.52 | 0 | 0.82 | 1.24 | 0.179 | 0.2118 | 0.0328 |
| 3zj0 | opls | 1.8 | 0.0973 | 2.34 | 0 | 98.49 | 0.64 | 0 | 1.28 | 0.85 | 0.1795 | 0.2069 | 0.0274 |
| 3zj0 | noschrodinger | 1.8 | 0.0099 | 1.1 | 0 | 98 | 0.64 | 0 | 1.92 | 0.96 | 0.1775 | 0.2033 | 0.0258 |
| 3zlr | opls | 2.03 | 0.0995 | 2.22 | 0 | 99.29 | 1.24 | 0 | 1.06 | 0.88 | 0.1901 | 0.2076 | 0.0175 |
| 3zlr | noschrodinger | 2.03 | 0.0027 | 0.88 | 0.35 | 98.95 | 0.82 | 0 | 0.85 | 0.76 | 0.1703 | 0.2044 | 0.0341 |
| 3zoc | opls | 2.11 | 0.0914 | 2.27 | 0 | 97.7 | 1.96 | 1 | 1.38 | 1.16 | 0.2338 | 0.2472 | 0.0134 |
| 3zoc | noschrodinger | 2.11 | 0.0034 | 0.85 | 0 | 96.55 | 5.23 | 0 | 2.06 | 1.75 | 0.2213 | 0.2412 | 0.0199 |
| 3zod | opls | 1.68 | 0.096 | 2.23 | 0.56 | 98.31 | 1.85 | 0 | 1.63 | 1.12 | 0.1992 | 0.2316 | 0.0324 |
| 3zod | noschrodinger | 1.68 | 0.0047 | 0.95 | 0 | 97.19 | 2.47 | 0 | 2.28 | 1.45 | 0.196 | 0.2272 | 0.0312 |
| 3zoe | opls | 1.85 | 0.0952 | 2.23 | 0 | 98.52 | 1.08 | 0 | 1.78 | 0.96 | 0.2263 | 0.2584 | 0.0321 |
| 3zoe | noschrodinger | 1.85 | 0.0028 | 0.82 | 0 | 98.83 | 2.83 | 0 | 2.31 | 1.35 | 0.2181 | 0.2532 | 0.0351 |
| 3zof | opls | 2.15 | 0.0934 | 2.27 | 0 | 98.29 | 1.4 | 0 | 0.88 | 0.88 | 0.1988 | 0.2313 | 0.0325 |
| 3zof | noschrodinger | 2.15 | 0.003 | 0.86 | 0 | 97.16 | 1.74 | 0 | 2.82 | 1.41 | 0.203 | 0.2269 | 0.0239 |
| 3zt9 | opls | 1.75 | 0.1005 | 2.41 | 0 | 98.42 | 1.33 | 2 | 3.66 | 1.25 | 0.152 | 0.1878 | 0.0358 |
| 3zt9 | noschrodinger | 1.75 | 0.0081 | 1.12 | 0 | 98.42 | 1.33 | 1 | 8.66 | 1.56 | 0.1569 | 0.1884 | 0.0315 |
| 3zui | opls | 1.71 | 0.1006 | 2.26 | 0 | 97.22 | 0 | 0 | 2.64 | 1.19 | 0.1677 | 0.2083 | 0.0406 |
| 3zui | noschrodinger | 1.71 | 0.0036 | 0.88 | 0.69 | 97.22 | 1.57 | 0 | 2.64 | 1.34 | 0.1718 | 0.2015 | 0.0297 |
| 3zyh | opls | 1.5 | 0.1152 | 2.39 | 0 | 97.48 | 0.53 | 0 | 8.87 | 1.58 | 0.1843 | 0.2058 | 0.0215 |
| 3zyh | noschrodinger | 1.5 | 0.0035 | 0.87 | 0 | 97.06 | 0 | 0 | 5.27 | 1.45 | 0.1849 | 0.2018 | 0.0169 |
| 3zyp | opls | 1.5 | 0.0992 | 2.49 | 0 | 95.79 | 0.52 | 2 | 3.76 | 1.46 | 0.1677 | 0.1954 | 0.0277 |
| 3zyp | noschrodinger | 1.5 | 0.0069 | 1.09 | 0 | 95.81 | 0.52 | 0 | 3.74 | 1.45 | 0.1695 | 0.1951 | 0.0256 |
| 4a0t | opls | 1.9 | 0.1056 | 2.21 | 0.18 | 97.05 | 0 | 1 | 1.64 | 1.08 | 0.145 | 0.1853 | 0.0403 |
| 4a0t | noschrodinger | 1.9 | 0.0033 | 0.82 | 0.18 | 97.05 | 0 | 0 | 0.82 | 0.92 | 0.1472 | 0.1759 | 0.0287 |

| | | | | | | | | | | | | | |
|---|---|---|---|---|---|---|---|---|---|---|---|---|---|
| 4a0z | opls | 1.9 | 0.0937 | 2.21 | 0 | 97.76 | 0.74 | 0 | 0.19 | 0.63 | 0.2085 | 0.2362 | 0.0277 |
| 4a0z | noschrodinger | 1.9 | 0.0027 | 0.78 | 0 | 98.11 | 1.1 | 0 | 1.34 | 0.9 | 0.2063 | 0.2355 | 0.0292 |
| 4a55 | opls | 3.5 | 0.0907 | 2.35 | 1.34 | 91.35 | 3.42 | 4 | 0.95 | 1.7 | 0.2112 | 0.2564 | 0.0452 |
| 4a55 | noschrodinger | 3.5 | 0.0028 | 0.81 | 2.57 | 87.59 | 6.23 | 0 | 3.26 | 2.33 | 0.2036 | 0.2404 | 0.0368 |
| 4a5w | opls | 3.5 | 0.09 | 2.46 | 2.68 | 87.18 | 5.38 | 57 | 2.12 | 2.16 | 0.2794 | 0.3296 | 0.0502 |
| 4a5w | noschrodinger | 3.5 | 0.0034 | 0.88 | 2.06 | 88.15 | 6.99 | 0 | 7.65 | 2.66 | 0.2763 | 0.3259 | 0.0496 |
| 4a6o | opls | 2.8 | 0.0937 | 2.26 | 0.71 | 94.68 | 1.25 | 0 | 1.37 | 1.31 | 0.2557 | 0.3144 | 0.0587 |
| 4a6o | noschrodinger | 2.8 | 0.003 | 0.85 | 0.35 | 91.9 | 2.07 | 0 | 3.87 | 1.9 | 0.2504 | 0.3017 | 0.0513 |
| 4a7w | opls | 1.8 | 0.0978 | 2.43 | 0 | 98.03 | 2.22 | 6 | 2.25 | 1.27 | 0.1675 | 0.1917 | 0.0242 |
| 4a7w | noschrodinger | 1.8 | 0.0097 | 1.2 | 0 | 97.97 | 2.75 | 0 | 2.27 | 1.35 | 0.1694 | 0.1907 | 0.0213 |
| 4a9h | opls | 2.04 | 0.0998 | 2.21 | 0 | 99.68 | 0.97 | 2 | 0.87 | 0.77 | 0.1664 | 0.2081 | 0.0417 |
| 4a9h | noschrodinger | 2.04 | 0.0027 | 0.75 | 0 | 100 | 0.96 | 0 | 1.04 | 0.8 | 0.1798 | 0.2194 | 0.0396 |
| 4aa7 | opls | 2 | 0.1001 | 2.27 | 0 | 98.39 | 3.49 | 1 | 1.04 | 1.22 | 0.1656 | 0.2076 | 0.042 |
| 4aa7 | noschrodinger | 2 | 0.0028 | 0.81 | 0 | 97.95 | 4.07 | 0 | 1.19 | 1.31 | 0.1695 | 0.2067 | 0.0372 |
| 4adn | opls | 1.6 | 0.0993 | 2.18 | 0 | 98.54 | 0.24 | 2 | 1.87 | 0.95 | 0.1784 | 0.2013 | 0.0229 |
| 4adn | noschrodinger | 1.6 | 0.0087 | 1 | 0 | 98.07 | 1.18 | 0 | 4.26 | 1.26 | 0.1774 | 0.2036 | 0.0262 |
| 4ag7 | opls | 1.55 | 0.1016 | 2.37 | 0.31 | 96.91 | 1.03 | 4 | 3.18 | 1.3 | 0.173 | 0.2046 | 0.0316 |
| 4ag7 | noschrodinger | 1.55 | 0.0066 | 0.98 | 0 | 96.31 | 3.07 | 0 | 2.62 | 1.67 | 0.1762 | 0.2043 | 0.0281 |
| 4aiw | opls | 1.5 | 0.1026 | 2.31 | 0 | 100 | 0.75 | 0 | 0.81 | 0.75 | 0.1553 | 0.1876 | 0.0323 |
| 4aiw | noschrodinger | 1.5 | 0.0058 | 1.56 | 0 | 99.32 | 0.75 | 0 | 1.62 | 0.91 | 0.1541 | 0.1851 | 0.031 |
| 4alb | opls | 3.03 | 0.0887 | 2.24 | 0.24 | 96.35 | 2.96 | 0 | 1.59 | 1.51 | 0.2192 | 0.2573 | 0.0381 |
| 4alb | noschrodinger | 3.03 | 0.0029 | 0.78 | 1.49 | 93.82 | 4.42 | 0 | 3.22 | 2.02 | 0.2128 | 0.2517 | 0.0389 |
| 4am9 | opls | 2.5 | 0.0903 | 2.24 | 0.75 | 94.78 | 2.52 | 0 | 0.86 | 1.43 | 0.221 | 0.2366 | 0.0156 |
| 4am9 | noschrodinger | 2.5 | 0.0026 | 0.73 | 0 | 93.38 | 4.13 | 0 | 0.86 | 1.66 | 0.2101 | 0.2307 | 0.0206 |
| 4anw | opls | 2.31 | 0.0916 | 2.4 | 1.97 | 93.34 | 5.92 | 15 | 1.32 | 1.88 | 0.2476 | 0.2671 | 0.0195 |
| 4anw | noschrodinger | 2.31 | 0.0033 | 0.9 | 2.93 | 89.87 | 11.75 | 0 | 6.09 | 2.7 | 0.2374 | 0.2592 | 0.0218 |
| 4arr | opls | 3 | 0.0902 | 2.3 | 0.19 | 93.66 | 1.21 | 3 | 0.45 | 1.14 | 0.2526 | 0.2746 | 0.022 |
| 4arr | noschrodinger | 3 | 0.0034 | 0.81 | 0 | 92.96 | 3.21 | 0 | 2.04 | 1.81 | 0.2435 | 0.2714 | 0.0279 |
| 4atg | opls | 1.89 | 0.1006 | 2.33 | 0 | 97.91 | 0.54 | 2 | 0.9 | 0.8 | 0.1656 | 0.2087 | 0.0431 |
| 4atg | noschrodinger | 1.89 | 0.0082 | 1.43 | 0 | 97.93 | 1.07 | 2 | 3.89 | 1.22 | 0.1745 | 0.2076 | 0.0331 |
| 4auy | opls | 2.1 | 0.102 | 2.17 | 0 | 96.45 | 0.37 | 1 | 2.07 | 1.21 | 0.1667 | 0.1959 | 0.0292 |
| 4auy | noschrodinger | 2.1 | 0.0046 | 0.93 | 0 | 96.15 | 1.48 | 0 | 3.7 | 1.55 | 0.1506 | 0.1835 | 0.0329 |
| 4avx | opls | 1.68 | 0.0964 | 2.27 | 0 | 99.49 | 0.52 | 1 | 0.82 | 0.76 | 0.174 | 0.2113 | 0.0373 |
| 4avx | noschrodinger | 1.68 | 0.0058 | 0.94 | 0 | 99.54 | 0.52 | 0 | 2.2 | 1 | 0.1767 | 0.2061 | 0.0294 |
| 4ayc | opls | 1.9 | 0.0914 | 2.22 | 0 | 97.63 | 0.78 | 0 | 1.5 | 0.97 | 0.2067 | 0.2561 | 0.0494 |
| 4ayc | noschrodinger | 1.9 | 0.0052 | 0.83 | 0 | 97.66 | 0.78 | 0 | 2.58 | 1.12 | 0.2113 | 0.2489 | 0.0376 |
| 4az2 | opls | 2.6 | 0.0929 | 2.21 | 0 | 96.89 | 1.53 | 0 | 0.61 | 1.03 | 0.153 | 0.1776 | 0.0246 |
| 4az2 | noschrodinger | 2.6 | 0.0033 | 0.88 | 0 | 96.58 | 3.8 | 0 | 1.22 | 1.5 | 0.1406 | 0.1717 | 0.0311 |
| 4az3 | opls | 2.04 | 0.1 | 2.2 | 0.25 | 96.5 | 0.84 | 1 | 0.75 | 0.97 | 0.1699 | 0.2045 | 0.0346 |
| 4az3 | noschrodinger | 2.04 | 0.003 | 0.83 | 0.49 | 96.59 | 1.39 | 0 | 1.94 | 1.29 | 0.1719 | 0.2058 | 0.0339 |
| 4b1l | opls | 1.65 | 0.0943 | 2.13 | 0 | 97.48 | 2.27 | 0 | 2.01 | 1.34 | 0.2012 | 0.2277 | 0.0265 |
| 4b1l | noschrodinger | 1.65 | 0.0066 | 1.02 | 0 | 97.5 | 3.01 | 0 | 0.4 | 1.11 | 0.1985 | 0.2219 | 0.0234 |
| 4b4q | opls | 2 | 0.0907 | 2.23 | 0.38 | 96.98 | 2.11 | 2 | 1.57 | 1.32 | 0.1932 | 0.2455 | 0.0523 |
| 4b4q | noschrodinger | 2 | 0.0034 | 1.39 | 0.37 | 97.03 | 1.25 | 0 | 7.04 | 1.63 | 0.2136 | 0.2425 | 0.0289 |
| 4b53 | opls | 1.8 | 0.0972 | 2.2 | 0 | 100 | 2.08 | 0 | 2.9 | 1.32 | 0.1898 | 0.228 | 0.0382 |
| 4b53 | noschrodinger | 1.8 | 0.0082 | 1.05 | 0 | 100 | 1.55 | 0 | 3.21 | 1.26 | 0.1872 | 0.224 | 0.0368 |
| 4b8v | opls | 1.59 | 0.0957 | 2.6 | 0 | 97.31 | 0 | 1 | 1.34 | 0.99 | 0.1935 | 0.2254 | 0.0319 |
| 4b8v | noschrodinger | 1.59 | 0.0138 | 1.45 | 0 | 97.33 | 1.21 | 0 | 6.02 | 1.52 | 0.1959 | 0.2254 | 0.0295 |
| 4b9h | opls | 2.1 | 0.0909 | 2.27 | 0 | 96.24 | 1.22 | 0 | 1 | 1.12 | 0.223 | 0.2577 | 0.0347 |

| | | | | | | | | | | | | | |
|---|---|---|---|---|---|---|---|---|---|---|---|---|---|
| 4b9h | noschrodinger | 2.1 | 0.0037 | 0.85 | 0 | 95.72 | 2.42 | 0 | 4 | 1.77 | 0.2293 | 0.2564 | 0.0271 |
| 4bcp | opls | 2.26 | 0.0943 | 2.27 | 0.28 | 97.82 | 1.36 | 7 | 0.95 | 0.93 | 0.1956 | 0.2139 | 0.0183 |
| 4bcp | noschrodinger | 2.26 | 0.0027 | 0.77 | 0.19 | 98.21 | 1.98 | 0 | 2.58 | 1.27 | 0.185 | 0.2045 | 0.0195 |
| 4bcr | opls | 2.5 | 0.0898 | 2.29 | 0 | 98.43 | 0.47 | 1 | 0.36 | 0.63 | 0.288 | 0.302 | 0.014 |
| 4bcr | noschrodinger | 2.5 | 0.0027 | 0.76 | 0 | 97.13 | 1.16 | 0 | 2.05 | 1.18 | 0.2739 | 0.2999 | 0.026 |
| 4bcx | opls | 2 | 0.1011 | 2.23 | 0 | 98.31 | 0 | 1 | 2.03 | 0.97 | 0.2387 | 0.28 | 0.0413 |
| 4bcx | noschrodinger | 2 | 0.0031 | 0.8 | 0 | 98.31 | 0 | 0 | 2.03 | 0.97 | 0.2288 | 0.2714 | 0.0426 |
| 4bgd | opls | 3.1 | 0.0908 | 2.28 | 0.41 | 96.71 | 1.71 | 10 | 0.99 | 1.18 | 0.2371 | 0.2511 | 0.014 |
| 4bgd | noschrodinger | 3.1 | 0.0031 | 0.82 | 0.26 | 94 | 3.24 | 0 | 3.73 | 1.95 | 0.2249 | 0.2416 | 0.0167 |
| 4bix | opls | 2 | 0.0986 | 2.25 | 0.23 | 98.19 | 0 | 0 | 0.83 | 0.76 | 0.223 | 0.2399 | 0.0169 |
| 4bix | noschrodinger | 2 | 0.0029 | 0.81 | 0.22 | 97.53 | 0.77 | 0 | 0.97 | 0.88 | 0.2114 | 0.2333 | 0.0219 |
| 4bjx | opls | 1.59 | 0.1101 | 2.14 | 0 | 100 | 0 | 0 | 3.13 | 1.1 | 0.1302 | 0.17 | 0.0398 |
| 4bjx | noschrodinger | 1.59 | 0.0043 | 0.95 | 0 | 99.2 | 0 | 0 | 2.24 | 1 | 0.1286 | 0.1658 | 0.0372 |
| 4bk0 | opls | 1.9 | 0.0967 | 2.36 | 0 | 100 | 0 | 0 | 0.97 | 0.79 | 0.207 | 0.2389 | 0.0319 |
| 4bk0 | noschrodinger | 1.9 | 0.0035 | 0.78 | 0 | 98.92 | 0.61 | 0 | 1.63 | 0.91 | 0.2072 | 0.2364 | 0.0292 |
| 4blu | opls | 1.85 | 0.1009 | 2.38 | 0 | 96.52 | 0.4 | 1 | 3.26 | 1.34 | 0.1569 | 0.2037 | 0.0468 |
| 4blu | noschrodinger | 1.85 | 0.0032 | 0.92 | 0.18 | 95.81 | 0.2 | 0 | 3.89 | 1.47 | 0.164 | 0.1968 | 0.0328 |
| 4bol | opls | 1.7 | 0.1011 | 2.29 | 0 | 98.34 | 1.72 | 2 | 1.54 | 1.08 | 0.1606 | 0.1965 | 0.0359 |
| 4bol | noschrodinger | 1.7 | 0.0039 | 0.96 | 0.41 | 97.51 | 1.97 | 0 | 1.41 | 1.2 | 0.1646 | 0.1914 | 0.0268 |
| 4bqc | opls | 3.2 | 0.0893 | 2.29 | 0.56 | 95.52 | 1.53 | 0 | 0.82 | 1.21 | 0.2849 | 0.3123 | 0.0274 |
| 4bqc | noschrodinger | 3.2 | 0.0034 | 0.85 | 0.55 | 95.87 | 4.23 | 0 | 1.16 | 1.59 | 0.2721 | 0.3013 | 0.0292 |
| 4bum | opls | 2.8 | 0.0902 | 2.2 | 0 | 97.42 | 5.36 | 1 | 0.47 | 1.33 | 0.2831 | 0.2976 | 0.0145 |
| 4bum | noschrodinger | 2.8 | 0.004 | 0.8 | 0.36 | 93.59 | 7.11 | 0 | 1.18 | 1.9 | 0.2656 | 0.2917 | 0.0261 |
| 4bv3 | opls | 2 | 0.1103 | 3.51 | 0 | 98.13 | 0.83 | 3 | 28.46 | 1.94 | 0.2012 | 0.2395 | 0.0383 |
| 4bv3 | noschrodinger | 2 | 0.0245 | 2.18 | 0 | 97.01 | 1.23 | 0 | 35.18 | 2.27 | 0.2025 | 0.2361 | 0.0336 |
| 4bvx | opls | 1.6 | 0.096 | 2.33 | 0.55 | 98.08 | 1.2 | 3 | 1.47 | 0.95 | 0.1947 | 0.2273 | 0.0326 |
| 4bvx | noschrodinger | 1.6 | 0.0037 | 0.88 | 0.54 | 97.56 | 2.99 | 0 | 4.55 | 1.68 | 0.1982 | 0.2245 | 0.0263 |
| 4bwc | opls | 1.89 | 0.0936 | 2.35 | 0 | 96.07 | 1.62 | 2 | 2.55 | 1.47 | 0.18 | 0.2109 | 0.0309 |
| 4bwc | noschrodinger | 1.89 | 0.0056 | 0.97 | 0 | 96.3 | 2.08 | 0 | 2.8 | 1.56 | 0.1836 | 0.2092 | 0.0256 |
| 4bwp | opls | 3.6 | 0.0907 | 2.37 | 0.13 | 95.56 | 1.78 | 3 | 0.82 | 1.25 | 0.1969 | 0.2463 | 0.0494 |
| 4bwp | noschrodinger | 3.6 | 0.003 | 0.77 | 0 | 95.59 | 1.62 | 0 | 3.27 | 1.58 | 0.198 | 0.2392 | 0.0412 |
| 4c0x | opls | 1.5 | 0.102 | 2.49 | 0 | 98.47 | 1.29 | 3 | 3.7 | 1.24 | 0.1603 | 0.1883 | 0.028 |
| 4c0x | noschrodinger | 1.5 | 0.018 | 2.02 | 0 | 98.48 | 1.27 | 2 | 5.23 | 1.36 | 0.1616 | 0.184 | 0.0224 |
| 4c1w | opls | 1.84 | 0.0995 | 2.37 | 0 | 97.3 | 1.19 | 0 | 1.9 | 1.14 | 0.1328 | 0.1822 | 0.0494 |
| 4c1w | noschrodinger | 1.84 | 0.0042 | 1.09 | 0 | 97.31 | 3.57 | 0 | 1.58 | 1.46 | 0.1349 | 0.1754 | 0.0405 |
| 4c2a | opls | 2.08 | 0.096 | 2.2 | 0.22 | 96.53 | 0.72 | 1 | 1.19 | 1.06 | 0.1982 | 0.221 | 0.0228 |
| 4c2a | noschrodinger | 2.08 | 0.0025 | 0.77 | 0.22 | 96.34 | 0.71 | 0 | 1.45 | 1.13 | 0.188 | 0.2196 | 0.0316 |
| 4c4p | opls | 2 | 0.0995 | 2.28 | 0 | 97.69 | 4.39 | 1 | 2.11 | 1.54 | 0.1993 | 0.2346 | 0.0353 |
| 4c4p | noschrodinger | 2 | 0.0028 | 0.91 | 0 | 98.17 | 6.28 | 0 | 1.32 | 1.47 | 0.1889 | 0.2299 | 0.041 |
| 4c69 | opls | 2.28 | 0.0905 | 2.3 | 0 | 97.15 | 4.68 | 1 | 0.63 | 1.37 | 0.2414 | 0.2672 | 0.0258 |
| 4c69 | noschrodinger | 2.28 | 0.0027 | 0.82 | 0.71 | 95.73 | 8.94 | 0 | 2.77 | 2.08 | 0.2249 | 0.2571 | 0.0322 |
| 4c76 | opls | 1.96 | 0.0977 | 2.37 | 0 | 97.44 | 3.07 | 2 | 3.5 | 1.62 | 0.1419 | 0.1826 | 0.0407 |
| 4c76 | noschrodinger | 1.96 | 0.0043 | 0.95 | 0.28 | 97.21 | 3.93 | 0 | 2.21 | 1.59 | 0.1403 | 0.1763 | 0.036 |
| 4ca4 | opls | 2.84 | 0.0916 | 2.25 | 0.32 | 94.84 | 3.37 | 0 | 1.43 | 1.64 | 0.2052 | 0.2403 | 0.0351 |
| 4ca4 | noschrodinger | 2.84 | 0.003 | 0.85 | 0 | 95.83 | 5.9 | 0 | 3.03 | 1.97 | 0.1978 | 0.2352 | 0.0374 |
| 4cb5 | opls | 1.5 | 0.0966 | 2.47 | 0 | 98.75 | 2.76 | 1 | 2.57 | 1.38 | 0.131 | 0.1427 | 0.0117 |
| 4cb5 | noschrodinger | 1.5 | 0.0129 | 1.37 | 0 | 99.38 | 2.07 | 0 | 5.14 | 1.51 | 0.1316 | 0.1431 | 0.0115 |
| 4cf8 | opls | 1.65 | 0.0942 | 2.51 | 0 | 99.3 | 2.25 | 4 | 5.16 | 1.54 | 0.1494 | 0.1833 | 0.0339 |
| 4cf8 | noschrodinger | 1.65 | 0.0089 | 1.4 | 0 | 99.31 | 2.95 | 0 | 4.96 | 1.62 | 0.1541 | 0.1823 | 0.0282 |

| | | | | | | | | | | | | | |
|---|---|---|---|---|---|---|---|---|---|---|---|---|---|
| 4ci3 | opls | 3.5 | 0.09 | 2.27 | 0.23 | 95.86 | 1.45 | 4 | 0.48 | 1.08 | 0.2282 | 0.2671 | 0.0389 |
| 4ci3 | noschrodinger | 3.5 | 0.0027 | 0.78 | 0.14 | 93.97 | 3.28 | 0 | 0.97 | 1.59 | 0.2168 | 0.257 | 0.0402 |
| 4cii | opls | 2.15 | 0.0937 | 2.24 | 0 | 98.31 | 1.22 | 0 | 0.33 | 0.69 | 0.2509 | 0.2704 | 0.0195 |
| 4cii | noschrodinger | 2.15 | 0.0025 | 0.65 | 0.56 | 96.63 | 2.42 | 0 | 0 | 1.01 | 0.2275 | 0.2521 | 0.0246 |
| 4ciu | opls | 3.5 | 0.0895 | 2.4 | 0.51 | 97.29 | 4.83 | 4 | 1.51 | 1.55 | 0.2179 | 0.2466 | 0.0287 |
| 4ciu | noschrodinger | 3.5 | 0.0031 | 0.78 | 0.47 | 97.36 | 7.53 | 0 | 4.16 | 1.99 | 0.2144 | 0.2544 | 0.04 |
| 4cjp | opls | 2 | 0.0926 | 2.19 | 0 | 99.3 | 0.38 | 0 | 2.55 | 1.04 | 0.1769 | 0.1919 | 0.015 |
| 4cjp | noschrodinger | 2 | 0.0045 | 0.83 | 0 | 99.31 | 1.86 | 0 | 3.72 | 1.37 | 0.1596 | 0.1837 | 0.0241 |
| 4cmw | opls | 2.21 | 0.0968 | 2.27 | 0 | 96.24 | 2.51 | 4 | 1.58 | 1.46 | 0.1975 | 0.2377 | 0.0402 |
| 4cmw | noschrodinger | 2.21 | 0.0028 | 0.77 | 0.18 | 96.81 | 3.27 | 0 | 2.21 | 1.58 | 0.1951 | 0.2304 | 0.0353 |
| 4cnd | opls | 1.5 | 0.0982 | 2.26 | 0 | 98.14 | 0.76 | 1 | 1.39 | 0.87 | 0.1928 | 0.2109 | 0.0181 |
| 4cnd | noschrodinger | 1.5 | 0.0047 | 0.94 | 0 | 97.85 | 0.38 | 0 | 2.77 | 1.1 | 0.1914 | 0.2057 | 0.0143 |
| 4cnl | opls | 1.7 | 0.1031 | 2.21 | 0 | 98.88 | 0.68 | 0 | 2.25 | 1 | 0.1477 | 0.1822 | 0.0345 |
| 4cnl | noschrodinger | 1.7 | 0.0034 | 0.79 | 0 | 99.44 | 0 | 0 | 2.89 | 1.08 | 0.1435 | 0.1769 | 0.0334 |
| 4co1 | opls | 1.5 | 0.099 | 2.21 | 0 | 99.33 | 1.89 | 0 | 1.73 | 1.14 | 0.2638 | 0.2829 | 0.0191 |
| 4co1 | noschrodinger | 1.5 | 0.0033 | 0.91 | 0 | 97.73 | 0.62 | 0 | 6.47 | 1.42 | 0.2594 | 0.2851 | 0.0257 |
| 4cp1 | opls | 2.5 | 0.0973 | 2.22 | 0 | 96.23 | 1.55 | 0 | 3.42 | 1.53 | 0.1766 | 0.2049 | 0.0283 |
| 4cp1 | noschrodinger | 2.5 | 0.0034 | 0.87 | 0 | 94.39 | 5.15 | 0 | 2.86 | 2 | 0.1776 | 0.2051 | 0.0275 |
| 4cry | opls | 1.61 | 0.0959 | 2.51 | 1.65 | 97.52 | 5.53 | 4 | 4.66 | 1.9 | 0.1342 | 0.1555 | 0.0213 |
| 4cry | noschrodinger | 1.61 | 0.01 | 1.4 | 1.63 | 97.14 | 5.43 | 0 | 3.95 | 1.89 | 0.1361 | 0.1548 | 0.0187 |
| 4cs0 | opls | 2.1 | 0.0933 | 2.29 | 0.41 | 97.55 | 1.92 | 1 | 0.97 | 1.1 | 0.1478 | 0.1736 | 0.0258 |
| 4cs0 | noschrodinger | 2.1 | 0.003 | 0.8 | 0.4 | 96.36 | 4.29 | 0 | 3.63 | 1.88 | 0.1436 | 0.1699 | 0.0263 |
| 4ct0 | opls | 2.45 | 0.0925 | 2.27 | 0 | 97.62 | 2.58 | 2 | 0.33 | 1.01 | 0.252 | 0.2748 | 0.0228 |
| 4ct0 | noschrodinger | 2.45 | 0.0027 | 0.78 | 0 | 93.92 | 5.15 | 0 | 2.34 | 1.96 | 0.2318 | 0.2637 | 0.0319 |
| 4cub | opls | 2.1 | 0.0988 | 2.2 | 0 | 97.1 | 0.32 | 0 | 0.87 | 0.93 | 0.1988 | 0.2334 | 0.0346 |
| 4cub | noschrodinger | 2.1 | 0.0032 | 0.85 | 0 | 98.31 | 0.32 | 0 | 2.62 | 1.05 | 0.1826 | 0.2236 | 0.041 |
| 4d05 | opls | 1.65 | 0.1028 | 2.43 | 0 | 96.7 | 1.38 | 5 | 3.72 | 1.47 | 0.1691 | 0.1997 | 0.0306 |
| 4d05 | noschrodinger | 1.65 | 0.0143 | 1.29 | 0 | 96.41 | 1.98 | 0 | 3.84 | 1.64 | 0.1677 | 0.1986 | 0.0309 |
| 4d1y | opls | 2.6 | 0.0923 | 2.22 | 0 | 96.24 | 0.69 | 1 | 0.37 | 0.89 | 0.228 | 0.2715 | 0.0435 |
| 4d1y | noschrodinger | 2.6 | 0.0028 | 0.75 | 0 | 95.38 | 0.69 | 0 | 1.47 | 1.21 | 0.213 | 0.2693 | 0.0563 |
| 4d3h | opls | 2 | 0.0959 | 2.32 | 0 | 98.98 | 0 | 0 | 0 | 0.5 | 0.1954 | 0.2351 | 0.0397 |
| 4d3h | noschrodinger | 2 | 0.0026 | 0.81 | 0 | 100 | 0.77 | 0 | 1.64 | 0.91 | 0.1868 | 0.2213 | 0.0345 |
| 4d62 | opls | 2.5 | 0.0928 | 2.25 | 0 | 97.79 | 0.8 | 0 | 2.2 | 1.04 | 0.2305 | 0.2695 | 0.039 |
| 4d62 | noschrodinger | 2.5 | 0.0027 | 0.84 | 0.74 | 97.79 | 4.8 | 0 | 2.64 | 1.61 | 0.215 | 0.2585 | 0.0435 |
| 4d63 | opls | 2.2 | 0.0963 | 2.25 | 0 | 98.53 | 2.4 | 0 | 1.76 | 1.22 | 0.2082 | 0.2348 | 0.0266 |
| 4d63 | noschrodinger | 2.2 | 0.0027 | 0.82 | 0.74 | 97.79 | 0.8 | 0 | 0.88 | 0.82 | 0.1945 | 0.2238 | 0.0293 |
| 4d6w | opls | 3.6 | 0.0897 | 2.25 | 0.41 | 96.04 | 2.07 | 2 | 0.99 | 1.3 | 0.2172 | 0.2617 | 0.0445 |
| 4d6w | noschrodinger | 3.6 | 0.0034 | 0.83 | 0.74 | 93.6 | 2.8 | 0 | 5.47 | 2.06 | 0.1973 | 0.2503 | 0.053 |
| 4d87 | opls | 3.5 | 0.0904 | 2.44 | 0.71 | 93.1 | 8.89 | 14 | 2.72 | 2.22 | 0.26 | 0.3074 | 0.0474 |
| 4d87 | noschrodinger | 3.5 | 0.0035 | 0.88 | 1.59 | 86.6 | 16.9 | 0 | 17.22 | 3.3 | 0.2322 | 0.2677 | 0.0355 |
| 4d8e | opls | 1.5 | 0.1061 | 2.33 | 0 | 99.2 | 0 | 3 | 1.53 | 0.9 | 0.1391 | 0.162 | 0.0229 |
| 4d8e | noschrodinger | 1.5 | 0.0046 | 0.96 | 0 | 98.81 | 0 | 0 | 0.26 | 0.6 | 0.1424 | 0.1588 | 0.0164 |
| 4d90 | opls | 2.6 | 0.0885 | 2.33 | 0.83 | 96.68 | 2.82 | 3 | 1.1 | 1.37 | 0.2052 | 0.2348 | 0.0296 |
| 4d90 | noschrodinger | 2.6 | 0.0032 | 0.87 | 0.82 | 94.69 | 5.99 | 0 | 3.85 | 2.13 | 0.2015 | 0.2242 | 0.0227 |
| 4dfw | opls | 1.55 | 0.1025 | 2.37 | 0 | 98.13 | 1.92 | 1 | 3.7 | 1.38 | 0.1448 | 0.1811 | 0.0363 |
| 4dfw | noschrodinger | 1.55 | 0.0039 | 1.02 | 0 | 98.15 | 2.38 | 0 | 2.65 | 1.34 | 0.1497 | 0.1773 | 0.0276 |
| 4did | opls | 2.35 | 0.0933 | 2.3 | 0 | 99 | 3.47 | 1 | 1.64 | 1.33 | 0.2186 | 0.2424 | 0.0238 |
| 4did | noschrodinger | 2.35 | 0.003 | 0.83 | 0 | 97.02 | 5.36 | 0 | 0.41 | 1.37 | 0.1995 | 0.229 | 0.0295 |
| 4dl8 | opls | 1.7 | 0.0982 | 2.37 | 0 | 99.51 | 1.6 | 1 | 1.41 | 1.03 | 0.1623 | 0.1876 | 0.0253 |

| | | | | | | | | | | | | | |
|---|---|---|---|---|---|---|---|---|---|---|---|---|---|
| 4dl8 | noschrodinger | 1.7 | 0.0071 | 1.15 | 0 | 98.64 | 1.07 | 0 | 1.42 | 0.9 | 0.1652 | 0.1843 | 0.0191 |
| 4dmo | opls | 2.14 | 0.0931 | 2.3 | 0 | 98.18 | 0.9 | 0 | 0.61 | 0.7 | 0.224 | 0.2575 | 0.0335 |
| 4dmo | noschrodinger | 2.14 | 0.0029 | 0.89 | 0 | 97.39 | 1.56 | 0 | 0.61 | 0.97 | 0.2092 | 0.248 | 0.0388 |
| 4dn9 | opls | 2.05 | 0.0947 | 2.26 | 0 | 98.84 | 0 | 0 | 0.69 | 0.72 | 0.2098 | 0.2204 | 0.0106 |
| 4dn9 | noschrodinger | 2.05 | 0.0032 | 0.83 | 0 | 99.48 | 1.27 | 0 | 0.34 | 0.71 | 0.1893 | 0.2174 | 0.0281 |
| 4doo | opls | 1.9 | 0.1028 | 2.25 | 0 | 97.75 | 0 | 1 | 1.69 | 0.98 | 0.184 | 0.221 | 0.037 |
| 4doo | noschrodinger | 1.9 | 0.0059 | 0.92 | 0 | 97.76 | 0.57 | 0 | 3.69 | 1.21 | 0.1741 | 0.2234 | 0.0493 |
| 4dpn | opls | 2.56 | 0.0922 | 2.63 | 0.67 | 93.78 | 4.62 | 10 | 4.59 | 2.15 | 0.1955 | 0.2513 | 0.0558 |
| 4dpn | noschrodinger | 2.56 | 0.0032 | 0.85 | 0.44 | 92.73 | 7.14 | 0 | 5.46 | 2.4 | 0.2141 | 0.2362 | 0.0221 |
| 4dqo | opls | 2.44 | 0.0915 | 2.19 | 0.38 | 96.17 | 1.33 | 0 | 0.59 | 1.05 | 0.2173 | 0.2532 | 0.0359 |
| 4dqo | noschrodinger | 2.44 | 0.004 | 1.54 | 0.38 | 94.7 | 1.31 | 0 | 5.18 | 1.73 | 0.222 | 0.2553 | 0.0333 |
| 4drk | opls | 1.5 | 0.1039 | 2.26 | 0 | 97.5 | 2.18 | 1 | 5.14 | 1.63 | 0.1471 | 0.1807 | 0.0336 |
| 4drk | noschrodinger | 1.5 | 0.0039 | 0.95 | 0 | 98.01 | 2.18 | 0 | 2.23 | 1.26 | 0.151 | 0.1764 | 0.0254 |
| 4dt6 | opls | 2.6 | 0.0919 | 2.45 | 0 | 97.79 | 2.37 | 3 | 0.94 | 1.11 | 0.188 | 0.2246 | 0.0366 |
| 4dt6 | noschrodinger | 2.6 | 0.003 | 0.8 | 0 | 97.81 | 4.09 | 0 | 2.5 | 1.54 | 0.1845 | 0.2104 | 0.0259 |
| 4dvl | opls | 2.75 | 0.0865 | 2.65 | 0 | 95.89 | 4.18 | 0 | 1.73 | 1.68 | 0.2452 | 0.2675 | 0.0223 |
| 4dvl | noschrodinger | 2.75 | 0.0052 | 0.97 | 0 | 94.97 | 4.11 | 0 | 10.17 | 2.34 | 0.2209 | 0.2554 | 0.0345 |
| 4dw6 | opls | 2 | 0.0926 | 2.32 | 0 | 98.9 | 2 | 1 | 1.33 | 1.09 | 0.1715 | 0.2094 | 0.0379 |
| 4dw6 | noschrodinger | 2 | 0.0075 | 1.07 | 0 | 98.37 | 2.63 | 0 | 2.66 | 1.37 | 0.1793 | 0.214 | 0.0347 |
| 4dxg | opls | 2.5 | 0.0933 | 2.25 | 0 | 97.38 | 0.57 | 0 | 0.3 | 0.73 | 0.1968 | 0.2264 | 0.0296 |
| 4dxg | noschrodinger | 2.5 | 0.0033 | 0.85 | 0 | 95.83 | 2.27 | 0 | 4.46 | 1.78 | 0.1874 | 0.2329 | 0.0455 |
| 4dxs | opls | 2.71 | 0.0923 | 2.29 | 0.47 | 93.49 | 2.14 | 3 | 0.86 | 1.44 | 0.1976 | 0.2403 | 0.0427 |
| 4dxs | noschrodinger | 2.71 | 0.0052 | 1.01 | 1.39 | 91.67 | 5.85 | 0 | 3.75 | 2.25 | 0.2143 | 0.2499 | 0.0356 |
| 4dzz | opls | 1.79 | 0.0973 | 2.28 | 0 | 98.27 | 3.29 | 1 | 1.1 | 1.21 | 0.1821 | 0.2129 | 0.0308 |
| 4dzz | noschrodinger | 1.79 | 0.0067 | 0.95 | 0 | 97.3 | 2.96 | 0 | 2.36 | 1.51 | 0.1834 | 0.2158 | 0.0324 |
| 4.00E+03 | opls | 2.4 | 0.0945 | 2.31 | 0 | 96.75 | 5.12 | 1 | 0.95 | 1.53 | 0.2038 | 0.2348 | 0.031 |
| 4.00E+03 | noschrodinger | 2.4 | 0.0027 | 0.77 | 0.5 | 96.29 | 6.55 | 0 | 1.26 | 1.72 | 0.1968 | 0.233 | 0.0362 |
| 4.00E+07 | opls | 2.8 | 0.0906 | 2.4 | 0.75 | 95.27 | 6.27 | 2 | 1.26 | 1.78 | 0.2482 | 0.2786 | 0.0304 |
| 4.00E+07 | noschrodinger | 2.8 | 0.0028 | 0.75 | 1.23 | 92.84 | 9.2 | 0 | 3.62 | 2.34 | 0.2468 | 0.2745 | 0.0277 |
| 4e0f | opls | 2.85 | 0.0908 | 2.28 | 0 | 96.49 | 4.05 | 5 | 1.08 | 1.5 | 0.2152 | 0.2453 | 0.0301 |
| 4e0f | noschrodinger | 2.85 | 0.0028 | 0.79 | 0 | 95.16 | 5.71 | 1 | 1.2 | 1.75 | 0.1926 | 0.233 | 0.0404 |
| 4e0h | opls | 2 | 0.0903 | 2.31 | 0 | 97.03 | 0 | 0 | 0 | 0.67 | 0.185 | 0.2312 | 0.0462 |
| 4e0h | noschrodinger | 2 | 0.0027 | 0.77 | 0 | 98.06 | 0 | 0 | 2.76 | 1.06 | 0.187 | 0.2372 | 0.0502 |
| 4e0i | opls | 3 | 0.0888 | 2.33 | 1.06 | 94.16 | 1.15 | 1 | 1.21 | 1.28 | 0.2337 | 0.2797 | 0.046 |
| 4e0i | noschrodinger | 3 | 0.0032 | 0.77 | 0.79 | 95.8 | 2.56 | 0 | 1.96 | 1.56 | 0.2148 | 0.263 | 0.0482 |
| 4e1t | opls | 2.26 | 0.0894 | 2.43 | 0 | 99.15 | 4.95 | 2 | 0.87 | 1.3 | 0.2012 | 0.2362 | 0.035 |
| 4e1t | noschrodinger | 2.26 | 0.0029 | 0.88 | 0.41 | 96.71 | 7.35 | 0 | 1.98 | 1.83 | 0.1918 | 0.2335 | 0.0417 |
| 4eaj | opls | 2.61 | 0.0919 | 2.35 | 0 | 97.63 | 5.81 | 1 | 1.77 | 1.59 | 0.2111 | 0.2406 | 0.0295 |
| 4eaj | noschrodinger | 2.61 | 0.0029 | 0.83 | 0.23 | 96.03 | 8 | 0 | 2.46 | 1.99 | 0.2092 | 0.2372 | 0.028 |
| 4ebk | opls | 2.15 | 0.1001 | 2.28 | 0 | 98.33 | 0.94 | 1 | 1.54 | 0.9 | 0.1866 | 0.2184 | 0.0318 |
| 4ebk | noschrodinger | 2.15 | 0.0072 | 1.39 | 0 | 99.01 | 2.07 | 0 | 4.39 | 1.46 | 0.1692 | 0.2038 | 0.0346 |
| 4egw | opls | 2.5 | 0.091 | 2.45 | 2.43 | 93.12 | 8.32 | 10 | 4.82 | 2.39 | 0.277 | 0.3123 | 0.0353 |
| 4egw | noschrodinger | 2.5 | 0.0035 | 0.86 | 2.82 | 90.93 | 12.1 | 0 | 14 | 3 | 0.2593 | 0.2974 | 0.0381 |
| 4eh1 | opls | 2.19 | 0.0947 | 2.31 | 0.21 | 97 | 2.43 | 1 | 3.12 | 1.57 | 0.1947 | 0.2261 | 0.0314 |
| 4eh1 | noschrodinger | 2.19 | 0.0033 | 0.84 | 0.21 | 97.23 | 3.61 | 1 | 2.5 | 1.6 | 0.1792 | 0.2139 | 0.0347 |
| 4ehg | opls | 3.5 | 0.0904 | 2.4 | 0.78 | 92.16 | 7.03 | 15 | 1.17 | 1.95 | 0.2071 | 0.253 | 0.0459 |
| 4ehg | noschrodinger | 3.5 | 0.003 | 0.89 | 2.72 | 86.96 | 11.76 | 0 | 4.46 | 2.66 | 0.1833 | 0.228 | 0.0447 |
| 4ehu | opls | 1.6 | 0.1018 | 2.48 | 0 | 97.69 | 0.23 | 0 | 3.02 | 1.16 | 0.179 | 0.2153 | 0.0363 |
| 4ehu | noschrodinger | 1.6 | 0.012 | 1.17 | 0 | 98.09 | 0.45 | 0 | 2.66 | 1.05 | 0.18 | 0.2116 | 0.0316 |

| | | | | | | | | | | | | | |
|---|---|---|---|---|---|---|---|---|---|---|---|---|---|
| 4eiu | opls | 1.9 | 0.0943 | 2.21 | 0 | 98.6 | 0 | 0 | 2.73 | 1.06 | 0.1991 | 0.2132 | 0.0141 |
| 4eiu | noschrodinger | 1.9 | 0.0038 | 0.83 | 0.42 | 97.92 | 0.47 | 0 | 1.99 | 0.99 | 0.198 | 0.2124 | 0.0144 |
| 4ek6 | opls | 1.52 | 0.0996 | 2.29 | 0 | 98.88 | 0.77 | 2 | 2.9 | 1.08 | 0.1762 | 0.2056 | 0.0294 |
| 4ek6 | noschrodinger | 1.52 | 0.0048 | 0.92 | 0 | 98.53 | 0.38 | 0 | 2.69 | 1.06 | 0.1784 | 0.2035 | 0.0251 |
| 4ekd | opls | 2.71 | 0.0909 | 2.28 | 0 | 96.85 | 2.47 | 2 | 0.54 | 1.17 | 0.2556 | 0.2806 | 0.025 |
| 4ekd | noschrodinger | 2.71 | 0.0028 | 0.76 | 0.45 | 94.17 | 3.44 | 0 | 1.76 | 1.73 | 0.2001 | 0.2478 | 0.0477 |
| 4ekv | opls | 1.95 | 0.0969 | 2.38 | 0 | 96.58 | 2.27 | 0 | 3.36 | 1.62 | 0.172 | 0.2162 | 0.0442 |
| 4ekv | noschrodinger | 1.95 | 0.003 | 0.93 | 0 | 95.8 | 0 | 0 | 1.69 | 1.21 | 0.183 | 0.2114 | 0.0284 |
| 4em0 | opls | 2.74 | 0.0936 | 2.69 | 0 | 98.61 | 5.84 | 0 | 3.46 | 1.72 | 0.2786 | 0.3134 | 0.0348 |
| 4em0 | noschrodinger | 2.74 | 0.0031 | 0.84 | 1.37 | 91.78 | 5.8 | 0 | 8.49 | 2.53 | 0.2861 | 0.3226 | 0.0365 |
| 4em2 | opls | 2.08 | 0.1036 | 2.28 | 0 | 100 | 1.46 | 1 | 0.39 | 0.76 | 0.2255 | 0.2514 | 0.0259 |
| 4em2 | noschrodinger | 2.08 | 0.0026 | 0.7 | 0 | 99.32 | 4.35 | 0 | 1.16 | 1.31 | 0.2056 | 0.2432 | 0.0376 |
| 4emt | opls | 1.5 | 0.0987 | 2.42 | 0 | 97.22 | 1.68 | 7 | 2.29 | 1.32 | 0.1711 | 0.1849 | 0.0138 |
| 4emt | noschrodinger | 1.5 | 0.0062 | 1.13 | 0.29 | 97.95 | 2.92 | 0 | 4.42 | 1.59 | 0.1551 | 0.1835 | 0.0284 |
| 4en6 | opls | 2.56 | 0.0981 | 2.28 | 0.34 | 95.72 | 2.43 | 3 | 2.62 | 1.64 | 0.2026 | 0.2173 | 0.0147 |
| 4en6 | noschrodinger | 2.56 | 0.0029 | 0.82 | 0.85 | 93.52 | 1.86 | 0 | 5.25 | 1.91 | 0.1948 | 0.2071 | 0.0123 |
| 4eoc | opls | 1.49 | 0.1027 | 2.52 | 0.43 | 97.41 | 0 | 6 | 7.59 | 1.53 | 0.1405 | 0.1767 | 0.0362 |
| 4eoc | noschrodinger | 1.49 | 0.0086 | 1.16 | 0.43 | 97.44 | 0 | 0 | 3.42 | 1.24 | 0.1443 | 0.1717 | 0.0274 |
| 4er9 | opls | 1.9 | 0.0924 | 2.38 | 0 | 100 | 2.41 | 0 | 4.19 | 1.49 | 0.2487 | 0.2859 | 0.0372 |
| 4er9 | noschrodinger | 1.9 | 0.0117 | 1.1 | 0 | 99.03 | 5.95 | 0 | 7.8 | 2.02 | 0.2404 | 0.2827 | 0.0423 |
| 4eru | opls | 2.1 | 0.0936 | 2.23 | 0 | 99.32 | 1.41 | 1 | 0 | 0.61 | 0.1992 | 0.2226 | 0.0234 |
| 4eru | noschrodinger | 2.1 | 0.0063 | 0.89 | 0 | 99.68 | 3.08 | 0 | 1.31 | 1.23 | 0.1787 | 0.219 | 0.0403 |
| 4erx | opls | 2.5 | 0.0963 | 2.27 | 0.26 | 99.21 | 2.01 | 1 | 0.51 | 0.91 | 0.1823 | 0.2101 | 0.0278 |
| 4erx | noschrodinger | 2.5 | 0.0027 | 0.75 | 0 | 99.74 | 3.33 | 0 | 1.69 | 1.32 | 0.1699 | 0.2096 | 0.0397 |
| 4es7 | opls | 2 | 0.0939 | 2.24 | 0 | 96.32 | 2.8 | 0 | 0.74 | 1.32 | 0.2173 | 0.2573 | 0.04 |
| 4es7 | noschrodinger | 2 | 0.0027 | 0.81 | 0 | 95.73 | 5.56 | 0 | 3.72 | 2.02 | 0.2003 | 0.2471 | 0.0468 |
| 4esn | opls | 2.2 | 0.0932 | 2.3 | 0 | 97.12 | 5.3 | 0 | 0 | 1.21 | 0.2196 | 0.2369 | 0.0173 |
| 4esn | noschrodinger | 2.2 | 0.0026 | 0.77 | 0 | 96.03 | 5.22 | 0 | 0 | 1.32 | 0.1955 | 0.2196 | 0.0241 |
| 4esq | opls | 1.7 | 0.1031 | 2.16 | 0 | 96.86 | 1.86 | 0 | 1.71 | 1.32 | 0.1508 | 0.1836 | 0.0328 |
| 4esq | noschrodinger | 1.7 | 0.0028 | 0.78 | 0 | 96.88 | 0.62 | 0 | 1.03 | 0.99 | 0.1528 | 0.1819 | 0.0291 |
| 4eu0 | opls | 1.69 | 0.1003 | 2.51 | 0 | 96.76 | 5.15 | 4 | 4.13 | 1.94 | 0.2036 | 0.2532 | 0.0496 |
| 4eu0 | noschrodinger | 1.69 | 0.0082 | 1.14 | 0.71 | 96.8 | 5.51 | 0 | 6.96 | 2.14 | 0.206 | 0.2493 | 0.0433 |
| 4euv | opls | 2 | 0.0962 | 2.51 | 1.43 | 97.49 | 5.96 | 4 | 3.03 | 1.79 | 0.2154 | 0.2646 | 0.0492 |
| 4euv | noschrodinger | 2 | 0.0092 | 1.09 | 2.84 | 94.33 | 7.59 | 0 | 6.51 | 2.41 | 0.2143 | 0.2591 | 0.0448 |
| 4ev1 | opls | 1.95 | 0.0967 | 2.38 | 0 | 97.72 | 4.52 | 1 | 2.65 | 1.61 | 0.1999 | 0.2389 | 0.039 |
| 4ev1 | noschrodinger | 1.95 | 0.0041 | 0.89 | 0 | 97.32 | 5.47 | 0 | 4.79 | 1.94 | 0.2023 | 0.2327 | 0.0304 |
| 4evy | opls | 1.77 | 0.0983 | 2.22 | 0 | 98.96 | 3.54 | 0 | 2.88 | 1.49 | 0.1588 | 0.1922 | 0.0334 |
| 4evy | noschrodinger | 1.77 | 0.005 | 1.28 | 0 | 98.96 | 3.15 | 0 | 2.26 | 1.38 | 0.1592 | 0.1891 | 0.0299 |
| 4ewl | opls | 1.85 | 0.1005 | 2.44 | 0.87 | 96.37 | 1.39 | 5 | 5.04 | 1.62 | 0.1717 | 0.2083 | 0.0366 |
| 4ewl | noschrodinger | 1.85 | 0.0082 | 1 | 1.03 | 95.88 | 2.76 | 0 | 5.27 | 1.9 | 0.1769 | 0.2068 | 0.0299 |
| 4exo | opls | 1.91 | 0.1047 | 2.25 | 0 | 99.25 | 3.45 | 1 | 1.27 | 1.26 | 0.1459 | 0.1871 | 0.0412 |
| 4exo | noschrodinger | 1.91 | 0.0076 | 1.08 | 0 | 99.31 | 4.1 | 0 | 1.69 | 1.39 | 0.1461 | 0.1801 | 0.034 |
| 4f48 | opls | 3 | 0.0932 | 2.37 | 0 | 96.67 | 7.61 | 3 | 1.88 | 1.83 | 0.2176 | 0.2551 | 0.0375 |
| 4f48 | noschrodinger | 3 | 0.0031 | 0.87 | 1.81 | 91.87 | 10.87 | 0 | 7.52 | 2.69 | 0.1998 | 0.2581 | 0.0583 |
| 4f6s | opls | 2.6 | 0.0923 | 2.24 | 0.17 | 98.27 | 0.75 | 3 | 0.51 | 0.68 | 0.2068 | 0.2401 | 0.0333 |
| 4f6s | noschrodinger | 2.6 | 0.0028 | 0.77 | 0 | 97.95 | 0.75 | 0 | 0.53 | 0.7 | 0.1988 | 0.2337 | 0.0349 |
| 4f6u | opls | 2.1 | 0.096 | 2.22 | 0 | 98.46 | 0.93 | 2 | 0.99 | 0.79 | 0.1876 | 0.2099 | 0.0223 |
| 4f6u | noschrodinger | 2.1 | 0.0028 | 0.78 | 0 | 97.97 | 1.1 | 0 | 0.91 | 0.82 | 0.1816 | 0.2007 | 0.0191 |
| 4f8p | opls | 2.05 | 0.0914 | 2.33 | 0.72 | 98.19 | 5.02 | 1 | 0.45 | 1.19 | 0.2518 | 0.2965 | 0.0447 |

| | | | | | | | | | | | | | |
|---|---|---|---|---|---|---|---|---|---|---|---|---|---|
| 4f8p | noschrodinger | 2.05 | 0.0028 | 0.75 | 0.72 | 95.7 | 10.37 | 0 | 4.1 | 2.26 | 0.2422 | 0.2802 | 0.038 |
| 4f9c | opls | 2.08 | 0.0927 | 2.24 | 0 | 97.05 | 0.87 | 2 | 0.16 | 0.73 | 0.2039 | 0.2194 | 0.0155 |
| 4f9c | noschrodinger | 2.08 | 0.0026 | 0.75 | 0 | 97.38 | 2.29 | 0 | 1.41 | 1.27 | 0.1935 | 0.2163 | 0.0228 |
| 4fbh | opls | 2.3 | 0.0946 | 2.22 | 0.39 | 97.29 | 0.96 | 0 | 0.49 | 0.8 | 0.169 | 0.224 | 0.055 |
| 4fbh | noschrodinger | 2.3 | 0.0032 | 0.82 | 0.38 | 98.09 | 0.48 | 0 | 1.71 | 0.92 | 0.1752 | 0.2141 | 0.0389 |
| 4fda | opls | 2.1 | 0.0918 | 2.4 | 0 | 98.92 | 1.55 | 0 | 1.83 | 1.09 | 0.1947 | 0.2183 | 0.0236 |
| 4fda | noschrodinger | 2.1 | 0.0029 | 0.82 | 0 | 97.87 | 2.05 | 0 | 2.13 | 1.25 | 0.1785 | 0.2055 | 0.027 |
| 4fek | opls | 2 | 0.0946 | 2.29 | 0 | 95.93 | 1.65 | 1 | 1.36 | 1.31 | 0.2442 | 0.2607 | 0.0165 |
| 4fek | noschrodinger | 2 | 0.0029 | 0.83 | 0.2 | 96.12 | 3.92 | 0 | 1.24 | 1.56 | 0.2292 | 0.2533 | 0.0241 |
| 4fer | opls | 2.1 | 0.0964 | 2.16 | 0 | 98.78 | 0.59 | 0 | 1.21 | 0.84 | 0.1808 | 0.207 | 0.0262 |
| 4fer | noschrodinger | 2.1 | 0.0028 | 0.8 | 0 | 98.29 | 0.88 | 0 | 1.06 | 0.81 | 0.1671 | 0.2002 | 0.0331 |
| 4ffb | opls | 2.88 | 0.0896 | 2.34 | 0.69 | 95.45 | 3.75 | 7 | 1.12 | 1.57 | 0.2144 | 0.2618 | 0.0474 |
| 4ffb | noschrodinger | 2.88 | 0.003 | 0.78 | 1.31 | 91.92 | 7.54 | 0 | 3.69 | 2.31 | 0.1988 | 0.2434 | 0.0446 |
| 4fg4 | opls | 2.7 | 0.094 | 2.21 | 0 | 98.29 | 0 | 0 | 0.92 | 0.78 | 0.1971 | 0.2188 | 0.0217 |
| 4fg4 | noschrodinger | 2.7 | 0.0028 | 0.79 | 0.24 | 96.1 | 0.59 | 0 | 1.38 | 1.13 | 0.1785 | 0.2163 | 0.0378 |
| 4fgr | opls | 2.6 | 0.0944 | 2.35 | 0.22 | 94.59 | 9.51 | 5 | 1.69 | 2.04 | 0.261 | 0.3015 | 0.0405 |
| 4fgr | noschrodinger | 2.6 | 0.0033 | 0.89 | 1.08 | 90.73 | 16.99 | 0 | 8.61 | 2.93 | 0.2468 | 0.2833 | 0.0365 |
| 4fhz | opls | 2.01 | 0.0953 | 2.24 | 0 | 97.7 | 1.17 | 1 | 2.32 | 1.13 | 0.2604 | 0.2833 | 0.0229 |
| 4fhz | noschrodinger | 2.01 | 0.0028 | 0.81 | 0 | 97.71 | 2.33 | 0 | 2.89 | 1.42 | 0.2389 | 0.2714 | 0.0325 |
| 4fod | opls | 1.99 | 0.0983 | 2.3 | 0 | 99 | 0.75 | 2 | 2.19 | 0.99 | 0.1653 | 0.2016 | 0.0363 |
| 4fod | noschrodinger | 1.99 | 0.0045 | 0.96 | 0 | 98.01 | 1.48 | 0 | 0.6 | 0.83 | 0.1667 | 0.198 | 0.0313 |
| 4fof | opls | 2.42 | 0.0945 | 2.31 | 0.61 | 98.79 | 1.34 | 2 | 1.05 | 0.9 | 0.1969 | 0.2232 | 0.0263 |
| 4fof | noschrodinger | 2.42 | 0.0028 | 0.78 | 0 | 97.59 | 1.33 | 0 | 2.08 | 1.16 | 0.2037 | 0.2319 | 0.0282 |
| 4fq5 | opls | 2.1 | 0.0982 | 2.38 | 0 | 98.57 | 10.53 | 2 | 1.72 | 1.7 | 0.1985 | 0.2315 | 0.033 |
| 4fq5 | noschrodinger | 2.1 | 0.003 | 0.89 | 0 | 98.17 | 10.5 | 0 | 2.25 | 1.78 | 0.1904 | 0.2221 | 0.0317 |
| 4fqe | opls | 1.93 | 0.0894 | 2.28 | 0 | 98.68 | 4.67 | 0 | 2.19 | 1.5 | 0.1939 | 0.2315 | 0.0376 |
| 4fqe | noschrodinger | 1.93 | 0.0038 | 0.86 | 0 | 97.47 | 3.82 | 0 | 3.13 | 1.65 | 0.1969 | 0.2281 | 0.0312 |
| 4fqp | opls | 3.56 | 0.0859 | 2.47 | 0.66 | 95.7 | 8.43 | 1 | 1.94 | 1.96 | 0.2321 | 0.2555 | 0.0234 |
| 4fqp | noschrodinger | 3.56 | 0.0047 | 0.9 | 0.33 | 92.11 | 15.97 | 0 | 8.35 | 2.85 | 0.2242 | 0.2496 | 0.0254 |
| 4frp | opls | 2 | 0.0976 | 2.29 | 0 | 97.01 | 3.7 | 0 | 0.88 | 1.37 | 0.18 | 0.2146 | 0.0346 |
| 4frp | noschrodinger | 2 | 0.0032 | 0.85 | 0 | 95.93 | 5.71 | 0 | 1.97 | 1.82 | 0.1816 | 0.2094 | 0.0278 |
| 4fry | opls | 2.1 | 0.0873 | 2.34 | 0 | 98.87 | 2.07 | 1 | 0 | 0.74 | 0.2227 | 0.2629 | 0.0402 |
| 4fry | noschrodinger | 2.1 | 0.0053 | 1.03 | 0.37 | 96.3 | 3.11 | 0 | 2.44 | 1.65 | 0.2253 | 0.2676 | 0.0423 |
| 4ftt | opls | 2.29 | 0.0952 | 2.25 | 0 | 97.03 | 0 | 0 | 0.23 | 0.76 | 0.1775 | 0.2053 | 0.0278 |
| 4ftt | noschrodinger | 2.29 | 0.0027 | 0.96 | 0 | 97.69 | 0 | 0 | 0 | 0.57 | 0.1689 | 0.2007 | 0.0318 |
| 4ftx | opls | 2.1 | 0.096 | 2.21 | 0.33 | 96.37 | 1.74 | 1 | 1.42 | 1.3 | 0.2156 | 0.2359 | 0.0203 |
| 4ftx | noschrodinger | 2.1 | 0.0029 | 0.79 | 0 | 95.42 | 2.43 | 0 | 1.42 | 1.49 | 0.1997 | 0.2329 | 0.0332 |
| 4fv3 | opls | 2.2 | 0.095 | 2.23 | 0.31 | 97.19 | 0.33 | 0 | 0.52 | 0.83 | 0.1797 | 0.2211 | 0.0414 |
| 4fv3 | noschrodinger | 2.2 | 0.0026 | 0.77 | 0.29 | 96.78 | 0.65 | 0 | 0.52 | 0.88 | 0.1815 | 0.2184 | 0.0369 |
| 4fyu | opls | 2 | 0.0995 | 2.25 | 0 | 96.94 | 1.9 | 0 | 0.59 | 1.09 | 0.1881 | 0.2119 | 0.0238 |
| 4fyu | noschrodinger | 2 | 0.0027 | 0.8 | 0 | 96.48 | 3.25 | 0 | 1.19 | 1.45 | 0.1771 | 0.2051 | 0.028 |
| 4g01 | opls | 2.2 | 0.0984 | 2.27 | 0.25 | 98.48 | 2.94 | 3 | 0.94 | 1.14 | 0.2098 | 0.2314 | 0.0216 |
| 4g01 | noschrodinger | 2.2 | 0.0032 | 0.82 | 0 | 98.24 | 4.68 | 0 | 2.68 | 1.56 | 0.1878 | 0.2243 | 0.0365 |
| 4g95 | opls | 1.35 | 0.1075 | 2.55 | 0 | 98.91 | 1.16 | 1 | 6.92 | 1.43 | 0.1591 | 0.1827 | 0.0236 |
| 4g95 | noschrodinger | 1.35 | 0.0171 | 1.57 | 0 | 97.83 | 1.73 | 1 | 13.2 | 1.85 | 0.1575 | 0.1776 | 0.0201 |
| 4gah | opls | 2.3 | 0.0953 | 2.38 | 0 | 97.75 | 3.76 | 2 | 1.32 | 1.35 | 0.1979 | 0.2367 | 0.0388 |
| 4gah | noschrodinger | 2.3 | 0.0029 | 0.86 | 0.28 | 98.05 | 5.57 | 0 | 3.31 | 1.69 | 0.1917 | 0.2265 | 0.0348 |
| 4gfq | opls | 2.65 | 0.0923 | 2.34 | 0 | 98.37 | 1.91 | 1 | 0.99 | 1.01 | 0.2307 | 0.2511 | 0.0204 |
| 4gfq | noschrodinger | 2.65 | 0.0038 | 0.78 | 0 | 96.74 | 2.55 | 0 | 4.93 | 1.77 | 0.2249 | 0.2454 | 0.0205 |

| | | | | | | | | | | | | | |
|---|---|---|---|---|---|---|---|---|---|---|---|---|---|
| 4gl7 | opls | 3.9 | 0.0906 | 2.39 | 1.34 | 90.2 | 8.15 | 5 | 2.13 | 2.22 | 0.1821 | 0.2159 | 0.0338 |
| 4gl7 | noschrodinger | 3.9 | 0.0032 | 0.95 | 1.56 | 85.78 | 14.04 | 0 | 4.52 | 2.75 | 0.1801 | 0.2148 | 0.0347 |
| 4gp7 | opls | 2 | 0.0955 | 2.26 | 0 | 98.41 | 1.32 | 4 | 2.11 | 1.07 | 0.1663 | 0.2081 | 0.0418 |
| 4gp7 | noschrodinger | 2 | 0.0057 | 0.93 | 0 | 99.41 | 1.9 | 0 | 3.17 | 1.32 | 0.1641 | 0.2044 | 0.0403 |
| 4gqn | opls | 1.85 | 0.0954 | 2.26 | 0 | 97.53 | 3.35 | 3 | 1.05 | 1.3 | 0.1739 | 0.2031 | 0.0292 |
| 4gqn | noschrodinger | 1.85 | 0.0054 | 0.86 | 0 | 97.16 | 3.73 | 0 | 1.58 | 1.49 | 0.1784 | 0.2014 | 0.023 |
| 4gu4 | opls | 3 | 0.0885 | 2.18 | 0 | 96.9 | 0.39 | 0 | 0.21 | 0.77 | 0.2464 | 0.2638 | 0.0174 |
| 4gu4 | noschrodinger | 3 | 0.0029 | 0.81 | 0.16 | 96.06 | 0.97 | 0 | 1.46 | 1.15 | 0.2262 | 0.2469 | 0.0207 |
| 4gud | opls | 1.91 | 0.0963 | 2.27 | 0.28 | 97.49 | 1.77 | 0 | 1.2 | 1.13 | 0.1583 | 0.201 | 0.0427 |
| 4gud | noschrodinger | 1.91 | 0.0047 | 0.86 | 0.26 | 96.91 | 2.26 | 0 | 1.5 | 1.34 | 0.1605 | 0.2025 | 0.042 |
| 4guj | opls | 1.5 | 0.1005 | 2.45 | 0.2 | 98.19 | 0.7 | 5 | 3.49 | 1.14 | 0.1579 | 0.1972 | 0.0393 |
| 4guj | noschrodinger | 1.5 | 0.0043 | 0.93 | 0 | 98.39 | 1.41 | 0 | 3.74 | 1.28 | 0.1633 | 0.1958 | 0.0325 |
| 4gur | opls | 2.51 | 0.0921 | 2.3 | 0.27 | 96.27 | 4.45 | 3 | 0.99 | 1.54 | 0.1979 | 0.221 | 0.0231 |
| 4gur | noschrodinger | 2.51 | 0.0029 | 0.82 | 0.4 | 95.24 | 9.13 | 0 | 3.12 | 2.16 | 0.1811 | 0.2122 | 0.0311 |
| 4gut | opls | 2 | 0.0959 | 2.4 | 0 | 97.29 | 4.34 | 3 | 1.33 | 1.48 | 0.1861 | 0.2111 | 0.025 |
| 4gut | noschrodinger | 2 | 0.0038 | 0.86 | 0.13 | 97.31 | 5.55 | 0 | 2.84 | 1.77 | 0.1885 | 0.2142 | 0.0257 |
| 4gvt | opls | 2.9 | 0.0902 | 2.9 | 2.45 | 92.65 | 7.87 | 21 | 3.17 | 2.25 | 0.2041 | 0.2621 | 0.058 |
| 4gvt | noschrodinger | 2.9 | 0.0068 | 1.09 | 1.96 | 91.67 | 11.8 | 0 | 2.61 | 2.36 | 0.2173 | 0.2544 | 0.0371 |
| 4gyc | opls | 2.05 | 0.0914 | 2.24 | 0 | 99.27 | 1.97 | 1 | 0.44 | 0.88 | 0.2087 | 0.2321 | 0.0234 |
| 4gyc | noschrodinger | 2.05 | 0.0039 | 0.83 | 0 | 98.92 | 2.46 | 0 | 1.55 | 1.2 | 0.2042 | 0.2321 | 0.0279 |
| 4gyf | opls | 1.65 | 0.0993 | 2.18 | 0.38 | 98.85 | 0 | 0 | 1.8 | 0.94 | 0.1493 | 0.1688 | 0.0195 |
| 4gyf | noschrodinger | 1.65 | 0.005 | 0.97 | 0.38 | 99.23 | 0 | 0 | 1.13 | 0.82 | 0.152 | 0.1685 | 0.0165 |
| 4h2d | opls | 1.8 | 0.096 | 2.37 | 0.33 | 96.39 | 4.6 | 2 | 1 | 1.54 | 0.1941 | 0.2231 | 0.029 |
| 4h2d | noschrodinger | 1.8 | 0.0084 | 1.02 | 0.33 | 96.42 | 5.7 | 0 | 3.2 | 1.92 | 0.189 | 0.2197 | 0.0307 |
| 4h54 | opls | 3.9 | 0.0899 | 2.38 | 0.39 | 97.28 | 1.74 | 1 | 0.68 | 1.04 | 0.2308 | 0.2916 | 0.0608 |
| 4h54 | noschrodinger | 3.9 | 0.0025 | 0.71 | 0.75 | 97.01 | 2.16 | 0 | 2.5 | 1.46 | 0.2299 | 0.2729 | 0.043 |
| 4h6z | opls | 2.7 | 0.0868 | 2.45 | 0 | 98.62 | 2.59 | 0 | 0.42 | 0.96 | 0.2304 | 0.2831 | 0.0527 |
| 4h6z | noschrodinger | 2.7 | 0.0028 | 0.79 | 0.72 | 96.74 | 5.81 | 0 | 2.56 | 1.83 | 0.2241 | 0.2629 | 0.0388 |
| 4h75 | opls | 2.1 | 0.0953 | 2.27 | 0 | 97.92 | 2.78 | 0 | 3.5 | 1.5 | 0.1963 | 0.2277 | 0.0314 |
| 4h75 | noschrodinger | 2.1 | 0.0035 | 0.79 | 0 | 96.46 | 4.95 | 0 | 1.75 | 1.69 | 0.194 | 0.2273 | 0.0333 |
| 4hat | opls | 1.78 | 0.1022 | 2.29 | 0.07 | 98.28 | 0.97 | 8 | 2.14 | 0.99 | 0.17 | 0.1981 | 0.0281 |
| 4hat | noschrodinger | 1.78 | 0.0056 | 0.91 | 0.07 | 98.06 | 0.88 | 0 | 1.2 | 0.84 | 0.1742 | 0.1995 | 0.0253 |
| 4hg0 | opls | 3.1 | 0.0881 | 2.38 | 1.53 | 93.88 | 8.25 | 1 | 0.54 | 1.79 | 0.2612 | 0.2973 | 0.0361 |
| 4hg0 | noschrodinger | 3.1 | 0.0034 | 0.82 | 2.61 | 86.52 | 13.11 | 0 | 5.91 | 2.8 | 0.2435 | 0.2822 | 0.0387 |
| 4hgp | opls | 1.8 | 0.102 | 2.49 | 0 | 98.29 | 3.52 | 0 | 1.8 | 1.35 | 0.2304 | 0.2751 | 0.0447 |
| 4hgp | noschrodinger | 1.8 | 0.0026 | 0.74 | 0 | 96.61 | 3.5 | 0 | 3.6 | 1.78 | 0.2369 | 0.2618 | 0.0249 |
| 4hh1 | opls | 3.5 | 0.0911 | 2.31 | 0.13 | 97.74 | 0.5 | 2 | 0.51 | 0.73 | 0.2088 | 0.2494 | 0.0406 |
| 4hh1 | noschrodinger | 3.5 | 0.0036 | 0.8 | 0.26 | 96.17 | 1 | 0 | 1.78 | 1.2 | 0.1873 | 0.2376 | 0.0503 |
| 4hhu | opls | 2 | 0.0938 | 2.32 | 0 | 98.63 | 2.26 | 0 | 2.56 | 1.31 | 0.1723 | 0.2158 | 0.0435 |
| 4hhu | noschrodinger | 2 | 0.0105 | 0.93 | 0 | 98.8 | 5.38 | 0 | 4.21 | 1.76 | 0.1717 | 0.2107 | 0.039 |
| 4hi5 | opls | 3.6 | 0.0901 | 2.3 | 0.78 | 90.7 | 3.28 | 3 | 1.33 | 1.78 | 0.2253 | 0.2797 | 0.0544 |
| 4hi5 | noschrodinger | 3.6 | 0.0026 | 0.77 | 3.05 | 93.13 | 7.26 | 0 | 0.45 | 1.75 | 0.232 | 0.2787 | 0.0467 |
| 4hj3 | opls | 2.64 | 0.0921 | 2.4 | 0.58 | 94.8 | 6.25 | 3 | 4.33 | 2.18 | 0.2254 | 0.2509 | 0.0255 |
| 4hj3 | noschrodinger | 2.64 | 0.0028 | 0.79 | 1.15 | 93.37 | 7.35 | 0 | 3.79 | 2.26 | 0.2231 | 0.2403 | 0.0172 |
| 4hj6 | opls | 2.2 | 0.093 | 2.3 | 0.29 | 98.56 | 3.31 | 1 | 0.18 | 0.97 | 0.2135 | 0.2358 | 0.0223 |
| 4hj6 | noschrodinger | 2.2 | 0.0027 | 0.76 | 0.57 | 97.13 | 5.15 | 0 | 1.8 | 1.64 | 0.2039 | 0.2274 | 0.0235 |
| 4hjl | opls | 1.5 | 0.1044 | 2.59 | 0.16 | 96.83 | 2.05 | 2 | 3.76 | 1.59 | 0.1339 | 0.157 | 0.0231 |
| 4hjl | noschrodinger | 1.5 | 0.0086 | 1.2 | 0 | 97.32 | 2.04 | 0 | 2.97 | 1.45 | 0.1358 | 0.155 | 0.0192 |
| 4hjm | opls | 1.55 | 0.1049 | 2.34 | 0 | 98.04 | 1.89 | 1 | 2.07 | 1.19 | 0.1299 | 0.1611 | 0.0312 |

| | | | | | | | | | | | | | |
|---|---|---|---|---|---|---|---|---|---|---|---|---|---|
| 4hjm | noschrodinger | 1.55 | 0.0054 | 0.95 | 0 | 99.04 | 1.89 | 0 | 2.57 | 1.25 | 0.1331 | 0.1589 | 0.0258 |
| 4hkf | opls | 1.69 | 0.1037 | 2.34 | 0 | 97.47 | 2.78 | 1 | 1.77 | 1.38 | 0.1513 | 0.1905 | 0.0392 |
| 4hkf | noschrodinger | 1.69 | 0.0066 | 1.06 | 0 | 96.99 | 3.38 | 0 | 1.77 | 1.51 | 0.1524 | 0.1839 | 0.0315 |
| 4hkn | opls | 2.05 | 0.0952 | 2.4 | 0 | 98.04 | 1.68 | 5 | 1.19 | 1 | 0.1716 | 0.2139 | 0.0423 |
| 4hkn | noschrodinger | 2.05 | 0.0037 | 0.87 | 0 | 96.59 | 1.67 | 0 | 0.59 | 1.09 | 0.1868 | 0.216 | 0.0292 |
| 4hnx | opls | 2.34 | 0.0904 | 2.33 | 0 | 98.34 | 0.85 | 10 | 1.25 | 0.85 | 0.2208 | 0.2402 | 0.0194 |
| 4hnx | noschrodinger | 2.34 | 0.0027 | 0.76 | 0 | 96.7 | 0.71 | 0 | 1 | 1 | 0.2291 | 0.2397 | 0.0106 |
| 4hqm | opls | 2.55 | 0.09 | 2.41 | 2.59 | 90.16 | 11.43 | 4 | 2.75 | 2.41 | 0.2873 | 0.3317 | 0.0444 |
| 4hqm | noschrodinger | 2.55 | 0.0031 | 0.86 | 6.67 | 85.13 | 14.12 | 0 | 5.51 | 2.83 | 0.2748 | 0.3142 | 0.0394 |
| 4hsl | opls | 2 | 0.0908 | 2.5 | 0.6 | 96.41 | 3.4 | 5 | 1.81 | 1.58 | 0.2134 | 0.2631 | 0.0497 |
| 4hsl | noschrodinger | 2 | 0.0029 | 0.82 | 0 | 95.35 | 4.73 | 0 | 2.9 | 1.92 | 0.2213 | 0.2562 | 0.0349 |
| 4hun | opls | 3.59 | 0.0906 | 2.67 | 9.39 | 70.72 | 19.63 | 24 | 6.2 | 3.16 | 0.3441 | 0.3816 | 0.0375 |
| 4hun | noschrodinger | 3.59 | 0.0047 | 1.22 | 14.29 | 58.61 | 31.55 | 0 | 37.79 | 4.12 | 0.3221 | 0.3661 | 0.044 |
| 4hur | opls | 2.02 | 0.1001 | 2.38 | 0 | 98.39 | 1.27 | 2 | 0.83 | 0.84 | 0.1841 | 0.219 | 0.0349 |
| 4hur | noschrodinger | 2.02 | 0.0028 | 0.81 | 0.16 | 97.92 | 1.62 | 0 | 1.85 | 1.12 | 0.1734 | 0.2125 | 0.0391 |
| 4hvj | opls | 2.1 | 0.0956 | 2.28 | 0 | 97.85 | 0 | 1 | 0.57 | 0.73 | 0.2013 | 0.2343 | 0.033 |
| 4hvj | noschrodinger | 2.1 | 0.0029 | 0.83 | 0 | 97.44 | 0.72 | 0 | 0.39 | 0.75 | 0.1847 | 0.2196 | 0.0349 |
| 4hvr | opls | 2.69 | 0.0903 | 2.3 | 0 | 96.41 | 1.35 | 0 | 0.36 | 0.97 | 0.2485 | 0.2771 | 0.0286 |
| 4hvr | noschrodinger | 2.69 | 0.003 | 0.83 | 1.74 | 88.95 | 4.03 | 0 | 2.9 | 2.12 | 0.2278 | 0.2576 | 0.0298 |
| 4hxc | opls | 2.15 | 0.0975 | 2.2 | 0 | 97.02 | 0.48 | 1 | 0.74 | 0.91 | 0.21 | 0.2343 | 0.0243 |
| 4hxc | noschrodinger | 2.15 | 0.0081 | 0.92 | 0 | 96.44 | 0.94 | 0 | 1.99 | 1.2 | 0.1911 | 0.231 | 0.0399 |
| 4i09 | opls | 2.05 | 0.0939 | 2.26 | 0 | 98.63 | 2.98 | 1 | 0.62 | 1.07 | 0.2263 | 0.2497 | 0.0234 |
| 4i09 | noschrodinger | 2.05 | 0.0028 | 0.78 | 0.5 | 97.26 | 6.23 | 0 | 2.34 | 1.76 | 0.2118 | 0.2428 | 0.031 |
| 4i0b | opls | 1.5 | 0.0996 | 2.44 | 0 | 99.48 | 3.12 | 6 | 6.5 | 1.73 | 0.1676 | 0.1993 | 0.0317 |
| 4i0b | noschrodinger | 1.5 | 0.0057 | 1.03 | 0 | 99.49 | 3.68 | 0 | 5.75 | 1.74 | 0.1716 | 0.1985 | 0.0269 |
| 4i1v | opls | 2.6 | 0.0901 | 2.3 | 0 | 96.98 | 1.45 | 0 | 0.17 | 0.86 | 0.2602 | 0.2721 | 0.0119 |
| 4i1v | noschrodinger | 2.6 | 0.0029 | 0.83 | 0.25 | 96.3 | 2.9 | 0 | 1.02 | 1.4 | 0.2231 | 0.2494 | 0.0263 |
| 4i5i | opls | 2.5 | 0.0928 | 2.32 | 0 | 97.78 | 2.57 | 2 | 0.46 | 1.02 | 0.1721 | 0.1847 | 0.0126 |
| 4i5i | noschrodinger | 2.5 | 0.0031 | 0.96 | 0 | 98.52 | 3.21 | 0 | 0.69 | 1.11 | 0.1574 | 0.1748 | 0.0174 |
| 4i9a | opls | 2.1 | 0.0924 | 2.32 | 0 | 97.54 | 3.08 | 0 | 1.99 | 1.43 | 0.2204 | 0.2366 | 0.0162 |
| 4i9a | noschrodinger | 2.1 | 0.003 | 0.82 | 0.17 | 97.03 | 3.76 | 0 | 2.7 | 1.66 | 0.2181 | 0.2351 | 0.017 |
| 4i9o | opls | 2 | 0.0894 | 2.27 | 0 | 98.33 | 3.23 | 0 | 1.51 | 1.28 | 0.2068 | 0.2251 | 0.0183 |
| 4i9o | noschrodinger | 2 | 0.0027 | 0.73 | 0 | 100 | 3.08 | 0 | 0.77 | 1.11 | 0.1939 | 0.2255 | 0.0316 |
| 4iae | opls | 2.05 | 0.0938 | 2.24 | 0 | 100 | 2 | 0 | 1.55 | 1.13 | 0.1699 | 0.1859 | 0.016 |
| 4iae | noschrodinger | 2.05 | 0.0029 | 0.77 | 0 | 99.17 | 3.95 | 0 | 1.55 | 1.35 | 0.1617 | 0.1818 | 0.0201 |
| 4ibf | opls | 2.29 | 0.0951 | 2.3 | 0 | 100 | 0.94 | 0 | 0.5 | 0.67 | 0.202 | 0.2467 | 0.0447 |
| 4ibf | noschrodinger | 2.29 | 0.0029 | 0.78 | 0 | 98.76 | 0 | 0 | 1.52 | 0.89 | 0.1916 | 0.2388 | 0.0472 |
| 4ibj | opls | 1.54 | 0.1029 | 2.34 | 0 | 99.59 | 0 | 1 | 2.4 | 1.02 | 0.1778 | 0.207 | 0.0292 |
| 4ibj | noschrodinger | 1.54 | 0.0037 | 0.87 | 0 | 100 | 0.89 | 0 | 2.63 | 1.05 | 0.183 | 0.2036 | 0.0206 |
| 4ihz | opls | 1.5 | 0.1027 | 2.73 | 0 | 97.77 | 2.05 | 6 | 5.04 | 1.56 | 0.173 | 0.2109 | 0.0379 |
| 4ihz | noschrodinger | 1.5 | 0.0098 | 1.11 | 0 | 98.11 | 2.38 | 0 | 2.61 | 1.33 | 0.1781 | 0.2109 | 0.0328 |
| 4ij7 | opls | 2.25 | 0.0942 | 2.27 | 0.3 | 98.21 | 1.04 | 1 | 0.77 | 0.76 | 0.2163 | 0.2526 | 0.0363 |
| 4ij7 | noschrodinger | 2.25 | 0.0032 | 0.82 | 0.29 | 96.47 | 3.1 | 0 | 1.92 | 1.56 | 0.1958 | 0.2395 | 0.0437 |
| 4ijh | opls | 1.5 | 0.0993 | 2.59 | 0 | 95.8 | 0 | 1 | 4.02 | 1.48 | 0.143 | 0.1728 | 0.0298 |
| 4ijh | noschrodinger | 1.5 | 0.0092 | 1.35 | 0 | 95.87 | 0 | 0 | 0.89 | 1.06 | 0.1292 | 0.1685 | 0.0393 |
| 4imq | opls | 1.5 | 0.1054 | 2.57 | 0 | 98.8 | 0.71 | 1 | 4.12 | 1.2 | 0.1551 | 0.1866 | 0.0315 |
| 4imq | noschrodinger | 1.5 | 0.0083 | 1.24 | 0 | 98.82 | 0.7 | 0 | 2.63 | 1.05 | 0.1566 | 0.1827 | 0.0261 |
| 4in4 | opls | 2.59 | 0.0923 | 2.38 | 0.12 | 96.42 | 3.77 | 3 | 0.68 | 1.4 | 0.1977 | 0.2329 | 0.0352 |
| 4in4 | noschrodinger | 2.59 | 0.0028 | 0.8 | 0.35 | 95.56 | 6.93 | 0 | 1.38 | 1.82 | 0.1857 | 0.2204 | 0.0347 |

| | | | | | | | | | | | | | |
|---|---|---|---|---|---|---|---|---|---|---|---|---|---|
| 4inn | opls | 2.6 | 0.0925 | 2.2 | 0.36 | 96.73 | 4.31 | 4 | 1.23 | 1.53 | 0.1985 | 0.2355 | 0.037 |
| 4inn | noschrodinger | 2.6 | 0.0028 | 0.82 | 0.69 | 96.54 | 8.05 | 0 | 2.87 | 1.99 | 0.1817 | 0.2255 | 0.0438 |
| 4ip9 | opls | 2.5 | 0.0885 | 2.29 | 0 | 100 | 5.92 | 0 | 0.6 | 1.29 | 0.2861 | 0.3156 | 0.0295 |
| 4ip9 | noschrodinger | 2.5 | 0.0031 | 0.72 | 0.49 | 96.57 | 3.29 | 0 | 1.5 | 1.5 | 0.264 | 0.3066 | 0.0426 |
| 4ipb | opls | 1.62 | 0.1021 | 2.15 | 0 | 98.76 | 0.42 | 0 | 1.54 | 0.9 | 0.1697 | 0.2033 | 0.0336 |
| 4ipb | noschrodinger | 1.62 | 0.0039 | 0.84 | 0 | 97.48 | 0.4 | 0 | 1.54 | 1 | 0.1652 | 0.2047 | 0.0395 |
| 4iqk | opls | 1.97 | 0.0959 | 2.54 | 0 | 97.16 | 1.32 | 3 | 3.47 | 1.38 | 0.1366 | 0.1642 | 0.0276 |
| 4iqk | noschrodinger | 1.97 | 0.0126 | 1.26 | 0 | 97.17 | 1.75 | 0 | 2.33 | 1.35 | 0.1396 | 0.1638 | 0.0242 |
| 4iqn | opls | 1.75 | 0.0963 | 2.58 | 0.18 | 96.24 | 0.69 | 6 | 3.12 | 1.36 | 0.1659 | 0.1954 | 0.0295 |
| 4iqn | noschrodinger | 1.75 | 0.0134 | 1.19 | 0 | 96.78 | 1.32 | 0 | 2.59 | 1.33 | 0.1681 | 0.1947 | 0.0266 |
| 4it6 | opls | 1.89 | 0.0954 | 2.29 | 0 | 99.51 | 0.52 | 0 | 1.89 | 0.95 | 0.2383 | 0.2847 | 0.0464 |
| 4it6 | noschrodinger | 1.89 | 0.0113 | 1.12 | 0.45 | 99.09 | 0 | 0 | 4.34 | 1.21 | 0.2486 | 0.2985 | 0.0499 |
| 4iup | opls | 1.9 | 0.1063 | 2.38 | 0.42 | 97.92 | 1.76 | 4 | 1.39 | 1.08 | 0.1504 | 0.1965 | 0.0461 |
| 4iup | noschrodinger | 1.9 | 0.0065 | 0.87 | 0.39 | 97.27 | 1.7 | 0 | 3.01 | 1.4 | 0.1568 | 0.1923 | 0.0355 |
| 4iur | opls | 2.5 | 0.0924 | 2.33 | 0.38 | 96.58 | 2.5 | 1 | 1.1 | 1.34 | 0.2175 | 0.2477 | 0.0302 |
| 4iur | noschrodinger | 2.5 | 0.0032 | 0.84 | 0.38 | 95.86 | 4.1 | 0 | 3.54 | 1.9 | 0.2134 | 0.2445 | 0.0311 |
| 4ivn | opls | 1.9 | 0.0961 | 2.3 | 0 | 99.03 | 3.2 | 3 | 0.74 | 1.12 | 0.1897 | 0.2226 | 0.0329 |
| 4ivn | noschrodinger | 1.9 | 0.0033 | 0.83 | 0 | 98.65 | 4.32 | 0 | 1.35 | 1.35 | 0.1924 | 0.222 | 0.0296 |
| 4iwf | opls | 1.93 | 0.0958 | 2.35 | 0 | 99.77 | 0.94 | 3 | 1.63 | 0.91 | 0.1872 | 0.2131 | 0.0259 |
| 4iwf | noschrodinger | 1.93 | 0.003 | 0.78 | 0.21 | 98.72 | 3.24 | 0 | 2.4 | 1.41 | 0.1896 | 0.2164 | 0.0268 |
| 4iwt | opls | 2.6 | 0.0892 | 2.26 | 1.4 | 91.26 | 4.24 | 2 | 1.45 | 1.87 | 0.2237 | 0.2651 | 0.0414 |
| 4iwt | noschrodinger | 2.6 | 0.0027 | 0.78 | 2.78 | 90.28 | 11.34 | 0 | 3.12 | 2.45 | 0.2108 | 0.2565 | 0.0457 |
| 4iwv | opls | 2.1 | 0.0943 | 2.31 | 0.23 | 98.16 | 2.93 | 1 | 1.57 | 1.26 | 0.2249 | 0.2471 | 0.0222 |
| 4iwv | noschrodinger | 2.1 | 0.003 | 0.97 | 0.23 | 96.8 | 5.03 | 0 | 2.86 | 1.81 | 0.2155 | 0.2396 | 0.0241 |
| 4iy0 | opls | 1.9 | 0.0976 | 2.35 | 0 | 99.31 | 5.84 | 2 | 0.8 | 1.33 | 0.1867 | 0.214 | 0.0273 |
| 4iy0 | noschrodinger | 1.9 | 0.0052 | 1.06 | 0 | 98.67 | 7.19 | 0 | 2.83 | 1.72 | 0.1902 | 0.2119 | 0.0217 |
| 4iy5 | opls | 2 | 0.102 | 2.24 | 0 | 99.23 | 1.33 | 0 | 1.99 | 1.06 | 0.1529 | 0.1789 | 0.026 |
| 4iy5 | noschrodinger | 2 | 0.0028 | 0.78 | 0 | 99.43 | 1.99 | 0 | 1.4 | 1.1 | 0.1549 | 0.1803 | 0.0254 |
| 4j30 | opls | 2.3 | 0.093 | 2.33 | 0 | 98.3 | 0 | 1 | 0.94 | 0.78 | 0.2084 | 0.2291 | 0.0207 |
| 4j30 | noschrodinger | 2.3 | 0.0028 | 0.74 | 0 | 99.2 | 1.36 | 0 | 2.12 | 1.09 | 0.1885 | 0.2187 | 0.0302 |
| 4j4b | opls | 1.9 | 0.0939 | 2.32 | 0.2 | 96.64 | 4.72 | 2 | 1.49 | 1.62 | 0.2185 | 0.2493 | 0.0308 |
| 4j4b | noschrodinger | 1.9 | 0.0054 | 1.01 | 0.19 | 95.91 | 5.87 | 0 | 2.74 | 1.93 | 0.2131 | 0.2506 | 0.0375 |
| 4j90 | opls | 3.1 | 0.0903 | 2.32 | 0 | 95.79 | 1.08 | 2 | 0.14 | 0.87 | 0.2283 | 0.2446 | 0.0163 |
| 4j90 | noschrodinger | 3.1 | 0.0044 | 0.94 | 0 | 93.72 | 4.03 | 0 | 4.8 | 2.13 | 0.2386 | 0.2494 | 0.0108 |
| 4jc1 | opls | 1.5 | 0.1002 | 2.42 | 0 | 97.79 | 0 | 1 | 9.78 | 1.56 | 0.1403 | 0.1706 | 0.0303 |
| 4jc1 | noschrodinger | 1.5 | 0.0097 | 1.17 | 0 | 99.26 | 2.14 | 0 | 6.24 | 1.59 | 0.1419 | 0.1617 | 0.0198 |
| 4jcw | opls | 1.75 | 0.1012 | 2.19 | 0 | 97.99 | 0.62 | 0 | 1.1 | 0.82 | 0.1683 | 0.2092 | 0.0409 |
| 4jcw | noschrodinger | 1.75 | 0.0038 | 0.96 | 0 | 97.75 | 0.62 | 0 | 1.73 | 0.98 | 0.1734 | 0.2007 | 0.0273 |
| 4jem | opls | 1.55 | 0.0973 | 2.25 | 0 | 98.98 | 0.42 | 0 | 1.27 | 0.85 | 0.1733 | 0.2002 | 0.0269 |
| 4jem | noschrodinger | 1.55 | 0.0037 | 0.9 | 0 | 98.69 | 0 | 0 | 1.7 | 0.92 | 0.1772 | 0.1998 | 0.0226 |
| 4jg4 | opls | 2.3 | 0.0947 | 2.55 | 0 | 98.15 | 9.9 | 1 | 1.5 | 1.65 | 0.2384 | 0.2813 | 0.0429 |
| 4jg4 | noschrodinger | 2.3 | 0.004 | 0.75 | 0 | 98.21 | 19.61 | 0 | 7.02 | 2.37 | 0.232 | 0.2796 | 0.0476 |
| 4jgp | opls | 2.03 | 0.0983 | 2.2 | 0 | 99.47 | 3.01 | 0 | 1.49 | 1.25 | 0.2111 | 0.2368 | 0.0257 |
| 4jgp | noschrodinger | 2.03 | 0.0061 | 0.92 | 0 | 99.51 | 5.57 | 0 | 1.64 | 1.48 | 0.1957 | 0.2288 | 0.0331 |
| 4jhi | opls | 2.6 | 0.091 | 2.18 | 0 | 96.2 | 2.17 | 0 | 1.18 | 1.35 | 0.2028 | 0.2325 | 0.0297 |
| 4jhi | noschrodinger | 2.6 | 0.003 | 0.79 | 0 | 95.62 | 4.29 | 0 | 1.18 | 1.62 | 0.189 | 0.2196 | 0.0306 |
| 4jin | opls | 2.1 | 0.0939 | 2.19 | 0 | 97.41 | 0.93 | 0 | 1.93 | 1.07 | 0.2001 | 0.2371 | 0.037 |
| 4jin | noschrodinger | 2.1 | 0.0028 | 0.75 | 0 | 98.02 | 1.39 | 0 | 2.92 | 1.19 | 0.1858 | 0.2269 | 0.0411 |
| 4jk3 | opls | 2.5 | 0.0895 | 2.37 | 0.44 | 96.71 | 5.94 | 6 | 0.52 | 1.47 | 0.2311 | 0.2736 | 0.0425 |

| | | | | | | | | | | | | | |
|---|---|---|---|---|---|---|---|---|---|---|---|---|---|
| 4jk3 | noschrodinger | 2.5 | 0.0031 | 0.86 | 0.61 | 96.56 | 6.25 | 0 | 2.59 | 1.87 | 0.237 | 0.2702 | 0.0332 |
| 4jkw | opls | 2.01 | 0.1006 | 2.47 | 0.86 | 97.41 | 2.97 | 1 | 2.13 | 1.46 | 0.1862 | 0.2456 | 0.0594 |
| 4jkw | noschrodinger | 2.01 | 0.0025 | 0.78 | 0.85 | 98.29 | 2.94 | 0 | 1.6 | 1.26 | 0.2039 | 0.2487 | 0.0448 |
| 4jlv | opls | 2.2 | 0.0911 | 2.23 | 0 | 99.09 | 1.51 | 0 | 0.55 | 0.82 | 0.22 | 0.2476 | 0.0276 |
| 4jlv | noschrodinger | 2.2 | 0.0029 | 0.81 | 0 | 98.2 | 2.48 | 0 | 1.1 | 1.11 | 0.2048 | 0.2427 | 0.0379 |
| 4jmu | opls | 2 | 0.0992 | 2.26 | 0 | 100 | 1.09 | 0 | 0.56 | 0.72 | 0.2108 | 0.2417 | 0.0309 |
| 4jmu | noschrodinger | 2 | 0.0029 | 0.8 | 0 | 98.06 | 1.09 | 0 | 2.25 | 1.03 | 0.1955 | 0.2301 | 0.0346 |
| 4jmx | opls | 2.55 | 0.1056 | 2.31 | 0 | 98.59 | 1.11 | 0 | 0.31 | 0.65 | 0.1878 | 0.2353 | 0.0475 |
| 4jmx | noschrodinger | 2.55 | 0.005 | 0.87 | 0 | 97.66 | 1.66 | 0 | 1.23 | 1.08 | 0.1649 | 0.2182 | 0.0533 |
| 4jpq | opls | 2.7 | 0.0898 | 2.27 | 0.5 | 95.04 | 1.65 | 0 | 0.87 | 1.28 | 0.2161 | 0.2412 | 0.0251 |
| 4jpq | noschrodinger | 2.7 | 0.0033 | 0.85 | 0.46 | 96.07 | 0.54 | 0 | 0.73 | 1 | 0.2081 | 0.2366 | 0.0285 |
| 4jps | opls | 2 | 0.0949 | 2.25 | 0 | 97.28 | 0.65 | 2 | 1 | 0.93 | 0.2123 | 0.2445 | 0.0322 |
| 4jps | noschrodinger | 2 | 0.0029 | 0.77 | 0.33 | 97.43 | 1.01 | 0 | 1.06 | 0.92 | 0.2076 | 0.2325 | 0.0249 |
| 4jqs | opls | 2.3 | 0.0936 | 2.18 | 0 | 97.94 | 0.48 | 0 | 1.78 | 0.95 | 0.1774 | 0.2033 | 0.0259 |
| 4jqs | noschrodinger | 2.3 | 0.0029 | 0.8 | 0.41 | 97.39 | 0.95 | 0 | 1.44 | 1 | 0.1704 | 0.2001 | 0.0297 |
| 4jsf | opls | 2.05 | 0.0952 | 2.24 | 0 | 97.52 | 1.22 | 5 | 2.05 | 1.14 | 0.2024 | 0.2228 | 0.0204 |
| 4jsf | noschrodinger | 2.05 | 0.0032 | 0.84 | 0 | 98.15 | 2.44 | 0 | 2.64 | 1.34 | 0.1918 | 0.217 | 0.0252 |
| 4jvc | opls | 2.5 | 0.0905 | 2.39 | 0 | 97.99 | 7.34 | 3 | 1.24 | 1.5 | 0.2848 | 0.3021 | 0.0173 |
| 4jvc | noschrodinger | 2.5 | 0.0028 | 0.83 | 1.99 | 92.04 | 15.17 | 0 | 3.09 | 2.48 | 0.2456 | 0.2729 | 0.0273 |
| 4jwp | opls | 2 | 0.1004 | 2.23 | 0 | 99.65 | 0.4 | 0 | 0.78 | 0.75 | 0.1745 | 0.2103 | 0.0358 |
| 4jwp | noschrodinger | 2 | 0.0031 | 0.82 | 0.31 | 99.07 | 0.8 | 0 | 0.59 | 0.7 | 0.1721 | 0.2073 | 0.0352 |
| 4jyk | opls | 1.49 | 0.0978 | 2.47 | 0 | 99.44 | 0.93 | 4 | 4.75 | 1.25 | 0.1378 | 0.1544 | 0.0166 |
| 4jyk | noschrodinger | 1.49 | 0.0082 | 1.08 | 0 | 100 | 1.53 | 1 | 3.24 | 1.26 | 0.1404 | 0.153 | 0.0126 |
| 4k21 | opls | 1.6 | 0.0993 | 2.3 | 0.43 | 97.86 | 0.46 | 2 | 3.73 | 1.19 | 0.1506 | 0.1877 | 0.0371 |
| 4k21 | noschrodinger | 1.6 | 0.0045 | 0.96 | 0.43 | 98.3 | 0.91 | 0 | 1.85 | 0.95 | 0.1543 | 0.1829 | 0.0286 |
| 4k46 | opls | 2.01 | 0.1004 | 2.29 | 0 | 99.05 | 1.18 | 0 | 1.17 | 0.88 | 0.1791 | 0.2219 | 0.0428 |
| 4k46 | noschrodinger | 2.01 | 0.0108 | 1.18 | 0 | 99.06 | 2.34 | 0 | 1.46 | 1.16 | 0.1548 | 0.2134 | 0.0586 |
| 4k47 | opls | 2.02 | 0.0978 | 2.29 | 0 | 98.89 | 1.37 | 1 | 0.7 | 0.83 | 0.2063 | 0.2367 | 0.0304 |
| 4k47 | noschrodinger | 2.02 | 0.0029 | 0.85 | 0 | 97.79 | 2.72 | 0 | 3.87 | 1.55 | 0.1985 | 0.2325 | 0.034 |
| 4k4d | opls | 2.17 | 0.0914 | 2.66 | 0.38 | 96.99 | 2.65 | 0 | 1.36 | 1.36 | 0.2163 | 0.2596 | 0.0433 |
| 4k4d | noschrodinger | 2.17 | 0.0026 | 0.82 | 0 | 96.3 | 3.95 | 0 | 1.59 | 1.61 | 0.2106 | 0.2478 | 0.0372 |
| 4k7q | opls | 3.41 | 0.091 | 2.33 | 2 | 91.89 | 5.75 | 6 | 1.61 | 1.98 | 0.3005 | 0.3173 | 0.0168 |
| 4k7q | noschrodinger | 3.41 | 0.0029 | 0.76 | 2.69 | 87.34 | 7.79 | 0 | 4.18 | 2.49 | 0.2722 | 0.3051 | 0.0329 |
| 4k80 | opls | 2.05 | 0.102 | 2.24 | 0.52 | 98.97 | 1.67 | 1 | 1.17 | 1 | 0.172 | 0.1927 | 0.0207 |
| 4k80 | noschrodinger | 2.05 | 0.0027 | 0.74 | 0 | 98.55 | 0.56 | 0 | 0.88 | 0.77 | 0.1646 | 0.1922 | 0.0276 |
| 4k84 | opls | 1.9 | 0.1017 | 2.49 | 0 | 97.98 | 3.83 | 3 | 1.58 | 1.35 | 0.1475 | 0.1964 | 0.0489 |
| 4k84 | noschrodinger | 1.9 | 0.0073 | 0.94 | 0 | 97.8 | 4.1 | 0 | 3.02 | 1.6 | 0.1536 | 0.1919 | 0.0383 |
| 4k90 | opls | 1.8 | 0.0995 | 2.29 | 0 | 97.83 | 3.08 | 3 | 2.1 | 1.39 | 0.1532 | 0.1748 | 0.0216 |
| 4k90 | noschrodinger | 1.8 | 0.0078 | 0.99 | 0 | 96.94 | 3.07 | 0 | 1.66 | 1.47 | 0.1539 | 0.1725 | 0.0186 |
| 4kit | opls | 3.6 | 0.0909 | 2.36 | 0.5 | 94.21 | 7.58 | 10 | 1.92 | 2.02 | 0.2328 | 0.2638 | 0.031 |
| 4kit | noschrodinger | 3.6 | 0.003 | 0.81 | 0.35 | 93.91 | 12.85 | 0 | 4.13 | 2.45 | 0.2272 | 0.2568 | 0.0296 |
| 4kmn | opls | 1.52 | 0.1 | 2.57 | 0 | 95.96 | 1.16 | 3 | 4.06 | 1.52 | 0.1648 | 0.187 | 0.0222 |
| 4kmn | noschrodinger | 1.52 | 0.0107 | 1.45 | 0.99 | 96.04 | 1.16 | 0 | 4.65 | 1.56 | 0.1683 | 0.1894 | 0.0211 |
| 4kpu | opls | 1.6 | 0.101 | 2.33 | 0.17 | 97.81 | 2.17 | 1 | 2.3 | 1.31 | 0.158 | 0.1747 | 0.0167 |
| 4kpu | noschrodinger | 1.6 | 0.0056 | 0.99 | 0.34 | 97.99 | 2.54 | 0 | 1.77 | 1.25 | 0.1618 | 0.1752 | 0.0134 |
| 4kqt | opls | 2.83 | 0.0896 | 2.17 | 0 | 98.74 | 1.5 | 0 | 0.37 | 0.77 | 0.2264 | 0.2463 | 0.0199 |
| 4kqt | noschrodinger | 2.83 | 0.0027 | 0.69 | 0 | 96.41 | 1.47 | 0 | 0.74 | 1.1 | 0.2362 | 0.253 | 0.0168 |
| 4kuj | opls | 1.99 | 0.0942 | 2.3 | 0 | 96.46 | 4.09 | 4 | 2.67 | 1.75 | 0.199 | 0.2352 | 0.0362 |
| 4kuj | noschrodinger | 1.99 | 0.007 | 0.93 | 0.19 | 95.54 | 6 | 0 | 3.02 | 1.99 | 0.1932 | 0.2275 | 0.0343 |

| | | | | | | | | | | | | | |
|---|---|---|---|---|---|---|---|---|---|---|---|---|---|
| 4kuo | opls | 2 | 0.0973 | 2.2 | 0 | 100 | 0 | 0 | 3.22 | 1.11 | 0.1491 | 0.1895 | 0.0404 |
| 4kuo | noschrodinger | 2 | 0.0029 | 0.79 | 0 | 98.2 | 0 | 0 | 2.72 | 1.06 | 0.1473 | 0.1863 | 0.039 |
| 4kvx | opls | 2 | 0.0975 | 2.37 | 0 | 98.4 | 6.56 | 0 | 2.37 | 1.64 | 0.1813 | 0.2281 | 0.0468 |
| 4kvx | noschrodinger | 2 | 0.0034 | 0.89 | 0 | 98.33 | 8.4 | 0 | 3.96 | 1.88 | 0.1762 | 0.2211 | 0.0449 |
| 4kwa | opls | 1.8 | 0.096 | 2.39 | 0 | 99.18 | 0.6 | 2 | 3.09 | 1.1 | 0.2047 | 0.2387 | 0.034 |
| 4kwa | noschrodinger | 1.8 | 0.0028 | 0.75 | 0 | 98.43 | 0.59 | 0 | 2.16 | 0.99 | 0.2118 | 0.2322 | 0.0204 |
| 4kyh | opls | 2.5 | 0.0937 | 2.22 | 0 | 97.85 | 0.33 | 1 | 0.36 | 0.67 | 0.1881 | 0.2225 | 0.0344 |
| 4kyh | noschrodinger | 2.5 | 0.0031 | 0.83 | 0 | 95.7 | 1.64 | 0 | 2.17 | 1.45 | 0.1715 | 0.2167 | 0.0452 |
| 4kyr | opls | 2.3 | 0.0904 | 2.31 | 0.49 | 97.07 | 0.55 | 0 | 1.57 | 1.07 | 0.1971 | 0.2282 | 0.0311 |
| 4kyr | noschrodinger | 2.3 | 0.0042 | 1.05 | 0.49 | 97.57 | 1.1 | 0 | 1.58 | 1.03 | 0.1888 | 0.2202 | 0.0314 |
| 4kzn | opls | 1.71 | 0.0926 | 2.26 | 1.09 | 95.65 | 3.33 | 0 | 0.59 | 1.4 | 0.1705 | 0.2067 | 0.0362 |
| 4kzn | noschrodinger | 1.71 | 0.009 | 1.05 | 1.08 | 96.77 | 4.4 | 0 | 6.51 | 2.05 | 0.1762 | 0.2079 | 0.0317 |
| 4l50 | opls | 2.1 | 0.0966 | 2.34 | 0.62 | 97.94 | 1.06 | 0 | 0.4 | 0.68 | 0.1884 | 0.2159 | 0.0275 |
| 4l50 | noschrodinger | 2.1 | 0.0047 | 0.84 | 1.23 | 97.74 | 4.22 | 0 | 2 | 1.5 | 0.1807 | 0.2053 | 0.0246 |
| 4l51 | opls | 1.9 | 0.0994 | 2.34 | 0.21 | 98.97 | 2.66 | 0 | 1.07 | 1.13 | 0.1714 | 0.2007 | 0.0293 |
| 4l51 | noschrodinger | 1.9 | 0.0035 | 0.89 | 0.62 | 99.18 | 4.76 | 0 | 2.54 | 1.55 | 0.175 | 0.1989 | 0.0239 |
| 4lac | opls | 2.82 | 0.0932 | 2.35 | 0 | 96.67 | 1.53 | 4 | 0.91 | 1.13 | 0.1925 | 0.2279 | 0.0354 |
| 4lac | noschrodinger | 2.82 | 0.003 | 0.81 | 0 | 96.44 | 2.5 | 0 | 2.34 | 1.55 | 0.184 | 0.2159 | 0.0319 |
| 4lad | opls | 2.3 | 0.0923 | 2.36 | 0 | 97.64 | 5.64 | 5 | 1.12 | 1.47 | 0.2456 | 0.2803 | 0.0347 |
| 4lad | noschrodinger | 2.3 | 0.0027 | 0.82 | 0.93 | 91.63 | 9.09 | 0 | 1.97 | 2.19 | 0.2352 | 0.2628 | 0.0276 |
| 4lax | opls | 2.01 | 0.0936 | 2.18 | 0 | 99.12 | 0.51 | 0 | 0.51 | 0.68 | 0.1841 | 0.2078 | 0.0237 |
| 4lax | noschrodinger | 2.01 | 0.0088 | 1.04 | 0.43 | 98.72 | 0 | 0 | 1.79 | 0.94 | 0.1831 | 0.2049 | 0.0218 |
| 4lay | opls | 1.7 | 0.1009 | 2.34 | 0 | 98.19 | 0.49 | 1 | 2.28 | 1.01 | 0.1595 | 0.1908 | 0.0313 |
| 4lay | noschrodinger | 1.7 | 0.01 | 1.06 | 0 | 98.28 | 0 | 0 | 1.02 | 0.8 | 0.1644 | 0.1902 | 0.0258 |
| 4lb8 | opls | 2.49 | 0.0928 | 2.17 | 0 | 97.91 | 0.88 | 0 | 0 | 0.52 | 0.2036 | 0.2465 | 0.0429 |
| 4lb8 | noschrodinger | 2.49 | 0.0026 | 0.71 | 0 | 97.79 | 2.14 | 0 | 0.7 | 1.02 | 0.1963 | 0.2461 | 0.0498 |
| 4lbh | opls | 1.75 | 0.0992 | 2.23 | 0 | 100 | 1.32 | 1 | 0.66 | 0.81 | 0.1739 | 0.1979 | 0.024 |
| 4lbh | noschrodinger | 1.75 | 0.0039 | 0.88 | 0 | 100 | 0 | 0 | 1.99 | 0.97 | 0.1789 | 0.1967 | 0.0178 |
| 4lco | opls | 2.7 | 0.092 | 2.35 | 0 | 95.6 | 2.21 | 2 | 0.55 | 1.25 | 0.2014 | 0.2436 | 0.0422 |
| 4lco | noschrodinger | 2.7 | 0.003 | 0.76 | 0.27 | 94.55 | 4.74 | 0 | 1.86 | 1.83 | 0.1784 | 0.2257 | 0.0473 |
| 4lcp | opls | 2 | 0.0962 | 2.33 | 0 | 98.35 | 0.73 | 0 | 1.28 | 0.85 | 0.184 | 0.2149 | 0.0309 |
| 4lcp | noschrodinger | 2 | 0.0029 | 0.78 | 0 | 98.09 | 2.53 | 0 | 2.02 | 1.28 | 0.1661 | 0.1991 | 0.033 |
| 4ld2 | opls | 1.55 | 0.098 | 2.38 | 0.55 | 97.79 | 0.74 | 3 | 1.66 | 0.97 | 0.1597 | 0.1865 | 0.0268 |
| 4ld2 | noschrodinger | 1.55 | 0.0084 | 1.07 | 0.55 | 97.25 | 0.74 | 0 | 3.15 | 1.25 | 0.1599 | 0.1844 | 0.0245 |
| 4lgu | opls | 2 | 0.0978 | 2.32 | 0 | 97.75 | 0 | 0 | 0.65 | 0.77 | 0.1769 | 0.2186 | 0.0417 |
| 4lgu | noschrodinger | 2 | 0.0124 | 1.23 | 0 | 98.33 | 0 | 0 | 1.3 | 0.85 | 0.1786 | 0.2217 | 0.0431 |
| 4ll2 | opls | 2.6 | 0.0968 | 2.24 | 0 | 95.7 | 2.67 | 0 | 2.26 | 1.63 | 0.1895 | 0.2339 | 0.0444 |
| 4ll2 | noschrodinger | 2.6 | 0.0032 | 0.83 | 0.45 | 93.69 | 6.65 | 0 | 4.09 | 2.24 | 0.1885 | 0.2276 | 0.0391 |
| 4llc | opls | 2 | 0.0918 | 2.62 | 0 | 99.56 | 3.09 | 2 | 4.12 | 1.57 | 0.1882 | 0.2366 | 0.0484 |
| 4llc | noschrodinger | 2 | 0.005 | 1.09 | 0 | 99.58 | 4.55 | 0 | 4.9 | 1.76 | 0.1879 | 0.2285 | 0.0406 |
| 4lm9 | opls | 1.6 | 0.1 | 2.4 | 1.75 | 95.61 | 2.88 | 1 | 3.83 | 1.83 | 0.2124 | 0.2527 | 0.0403 |
| 4lm9 | noschrodinger | 1.6 | 0.0125 | 1.27 | 2.34 | 94.53 | 2.75 | 0 | 11.53 | 2.29 | 0.2076 | 0.25 | 0.0424 |
| 4ln0 | opls | 2.9 | 0.0882 | 2.22 | 0.27 | 98.92 | 4.92 | 2 | 0.82 | 1.28 | 0.1977 | 0.2405 | 0.0428 |
| 4ln0 | noschrodinger | 2.9 | 0.0027 | 0.79 | 0 | 95.46 | 6.62 | 0 | 1.66 | 1.86 | 0.1943 | 0.2319 | 0.0376 |
| 4ln7 | opls | 1.7 | 0.0993 | 2.46 | 0 | 97.47 | 1.66 | 2 | 2.38 | 1.29 | 0.184 | 0.2155 | 0.0315 |
| 4ln7 | noschrodinger | 1.7 | 0.0138 | 1.3 | 0 | 96.5 | 1.1 | 0 | 5.37 | 1.55 | 0.186 | 0.2123 | 0.0263 |
| 4lo2 | opls | 2.25 | 0.0946 | 2.21 | 0 | 96.18 | 1.67 | 4 | 0.43 | 1.08 | 0.1896 | 0.2127 | 0.0231 |
| 4lo2 | noschrodinger | 2.25 | 0.0026 | 0.77 | 0 | 96.76 | 3.47 | 0 | 1.82 | 1.55 | 0.1766 | 0.2076 | 0.031 |
| 4lo5 | opls | 2.7 | 0.0929 | 2.21 | 0 | 95.21 | 3.56 | 0 | 0.21 | 1.33 | 0.2374 | 0.2588 | 0.0214 |

| | | | | | | | | | | | | | |
|---|---|---|---|---|---|---|---|---|---|---|---|---|---|
| 4lo5 | noschrodinger | 2.7 | 0.0029 | 0.81 | 0 | 95.23 | 5.77 | 0 | 1.59 | 1.82 | 0.2232 | 0.2485 | 0.0253 |
| 4lo6 | opls | 2.3 | 0.0964 | 2.21 | 0.17 | 95.55 | 3.75 | 0 | 0.85 | 1.51 | 0.2096 | 0.2219 | 0.0123 |
| 4lo6 | noschrodinger | 2.3 | 0.0027 | 0.8 | 0.17 | 96.08 | 6.89 | 0 | 1.27 | 1.75 | 0.2019 | 0.2148 | 0.0129 |
| 4lov | opls | 1.5 | 0.1103 | 2.15 | 0 | 98.03 | 0 | 0 | 2.04 | 0.97 | 0.155 | 0.1766 | 0.0216 |
| 4lov | noschrodinger | 1.5 | 0.0034 | 0.92 | 0 | 97.44 | 0 | 0 | 0.82 | 0.87 | 0.1552 | 0.1728 | 0.0176 |
| 4lps | opls | 2 | 0.0984 | 2.36 | 0.7 | 97.89 | 0.53 | 1 | 2.42 | 1.05 | 0.1604 | 0.2014 | 0.041 |
| 4lps | noschrodinger | 2 | 0.0028 | 0.81 | 0.47 | 97.89 | 0.26 | 0 | 1.71 | 0.95 | 0.1644 | 0.1982 | 0.0338 |
| 4lrj | opls | 1.62 | 0.1001 | 2.45 | 0.33 | 98.34 | 0 | 3 | 1.19 | 0.83 | 0.1949 | 0.2278 | 0.0329 |
| 4lrj | noschrodinger | 1.62 | 0.0102 | 1.19 | 0.64 | 97.75 | 1.08 | 0 | 3.37 | 1.21 | 0.2003 | 0.2258 | 0.0255 |
| 4ltm | opls | 2.5 | 0.0951 | 2.38 | 0 | 97.14 | 0 | 1 | 1.02 | 0.95 | 0.1644 | 0.1932 | 0.0288 |
| 4ltm | noschrodinger | 2.5 | 0.0028 | 0.81 | 0 | 96.17 | 0.66 | 0 | 1.03 | 1.06 | 0.1702 | 0.1905 | 0.0203 |
| 4ltn | opls | 2 | 0.0968 | 2.38 | 0 | 97.71 | 0.68 | 1 | 1.35 | 0.93 | 0.1574 | 0.1737 | 0.0163 |
| 4ltn | noschrodinger | 2 | 0.0136 | 1.33 | 1.64 | 96.72 | 1.32 | 0 | 4.75 | 1.54 | 0.1495 | 0.1696 | 0.0201 |
| 4m4q | opls | 2.5 | 0.0909 | 2.25 | 0 | 96.95 | 0 | 0 | 0.61 | 0.88 | 0.2421 | 0.2612 | 0.0191 |
| 4m4q | noschrodinger | 2.5 | 0.0027 | 0.72 | 0.51 | 95.96 | 0.56 | 0 | 2.42 | 1.3 | 0.2262 | 0.2592 | 0.033 |
| 4m5b | opls | 1.83 | 0.0906 | 2.33 | 0 | 97.14 | 1.21 | 2 | 2.12 | 1.2 | 0.1883 | 0.1999 | 0.0116 |
| 4m5b | noschrodinger | 1.83 | 0.0107 | 1.07 | 0.47 | 95.75 | 2.42 | 0 | 2.39 | 1.61 | 0.1873 | 0.1989 | 0.0116 |
| 4m5c | opls | 2.5 | 0.0923 | 2.27 | 0 | 97.14 | 3.64 | 1 | 0.29 | 1.19 | 0.2148 | 0.2253 | 0.0105 |
| 4m5c | noschrodinger | 2.5 | 0.0028 | 0.79 | 0.47 | 95.75 | 3.64 | 0 | 0.89 | 1.49 | 0.2012 | 0.2195 | 0.0183 |
| 4m5n | opls | 2 | 0.0997 | 2.32 | 0 | 98.41 | 0.36 | 2 | 0.19 | 0.57 | 0.1436 | 0.1697 | 0.0261 |
| 4m5n | noschrodinger | 2 | 0.0042 | 0.9 | 0 | 98.42 | 1.79 | 0 | 1.32 | 1.05 | 0.1513 | 0.1729 | 0.0216 |
| 4m5o | opls | 1.9 | 0.0978 | 2.35 | 0 | 97.97 | 0.56 | 1 | 1.49 | 0.9 | 0.2131 | 0.2447 | 0.0316 |
| 4m5o | noschrodinger | 1.9 | 0.0026 | 0.72 | 0 | 97.47 | 0 | 0 | 1.79 | 1.04 | 0.2185 | 0.2403 | 0.0218 |
| 4m8y | opls | 2.22 | 0.121 | 5.8 | 0 | 97.38 | 2.4 | 4 | 63.16 | 2.68 | 0.206 | 0.263 | 0.057 |
| 4m8y | noschrodinger | 2.22 | 0.0147 | 2.92 | 0 | 97.45 | 1.78 | 0 | 50.67 | 2.48 | 0.2278 | 0.2862 | 0.0584 |
| 4m99 | opls | 2.08 | 0.0923 | 2.34 | 0 | 96.86 | 0.67 | 0 | 0.56 | 0.88 | 0.233 | 0.2435 | 0.0105 |
| 4m99 | noschrodinger | 2.08 | 0.0179 | 1.04 | 0.49 | 95.91 | 1.56 | 0 | 1.34 | 1.29 | 0.2241 | 0.2413 | 0.0172 |
| 4m9q | opls | 2.5 | 0.0913 | 2.48 | 0.18 | 97.31 | 6.41 | 15 | 1.03 | 1.55 | 0.202 | 0.25 | 0.048 |
| 4m9q | noschrodinger | 2.5 | 0.0035 | 1.05 | 0.35 | 97.04 | 9.98 | 1 | 3.72 | 2.09 | 0.2111 | 0.2459 | 0.0348 |
| 4mda | opls | 1.7 | 0.0986 | 2.38 | 0 | 98.45 | 2.19 | 1 | 2.28 | 1.26 | 0.1885 | 0.2193 | 0.0308 |
| 4mda | noschrodinger | 1.7 | 0.0041 | 0.86 | 0 | 98.47 | 3.24 | 0 | 3.7 | 1.55 | 0.1925 | 0.2203 | 0.0278 |
| 4mdy | opls | 1.78 | 0.0984 | 2.22 | 0 | 98.18 | 0.88 | 2 | 1.82 | 0.94 | 0.1623 | 0.2021 | 0.0398 |
| 4mdy | noschrodinger | 1.78 | 0.0032 | 0.83 | 0 | 98.23 | 0.43 | 0 | 1.59 | 0.91 | 0.1674 | 0.1955 | 0.0281 |
| 4mik | opls | 1.95 | 0.0973 | 2.38 | 0 | 98.44 | 0 | 0 | 3.26 | 1.12 | 0.1826 | 0.2209 | 0.0383 |
| 4mik | noschrodinger | 1.95 | 0.0076 | 1.17 | 0 | 98.45 | 0.46 | 0 | 5.56 | 1.3 | 0.1851 | 0.2187 | 0.0336 |
| 4ml9 | opls | 1.84 | 0.0988 | 2.41 | 0.2 | 98.4 | 0.66 | 4 | 5.41 | 1.29 | 0.1489 | 0.1764 | 0.0275 |
| 4ml9 | noschrodinger | 1.84 | 0.0136 | 1.32 | 0.18 | 98.35 | 1.27 | 0 | 4.84 | 1.33 | 0.1482 | 0.1771 | 0.0289 |
| 4mln | opls | 2.1 | 0.0926 | 2.35 | 0 | 97.4 | 1.08 | 1 | 1.8 | 1.08 | 0.2111 | 0.2397 | 0.0286 |
| 4mln | noschrodinger | 2.1 | 0.0027 | 0.73 | 0 | 96.88 | 1.42 | 0 | 3.49 | 1.44 | 0.2005 | 0.2299 | 0.0294 |
| 4mlo | opls | 1.65 | 0.0968 | 2.21 | 0 | 99.11 | 0 | 1 | 0.45 | 0.66 | 0.1778 | 0.2084 | 0.0306 |
| 4mlo | noschrodinger | 1.65 | 0.0044 | 0.82 | 0 | 99.62 | 0.4 | 0 | 0.91 | 0.78 | 0.1806 | 0.2079 | 0.0273 |
| 4mph | opls | 2.03 | 0.104 | 2.17 | 0.28 | 97.46 | 0.63 | 0 | 1.86 | 1.06 | 0.1724 | 0.192 | 0.0196 |
| 4mph | noschrodinger | 2.03 | 0.0026 | 0.74 | 0 | 96.64 | 0.63 | 0 | 2.02 | 1.19 | 0.1596 | 0.185 | 0.0254 |
| 4mrd | opls | 2.55 | 0.0887 | 2.19 | 0.69 | 95.17 | 1.56 | 0 | 0.42 | 1.13 | 0.2028 | 0.2432 | 0.0404 |
| 4mrd | noschrodinger | 2.55 | 0.0034 | 0.84 | 0 | 93.88 | 1.54 | 0 | 4.22 | 1.75 | 0.1992 | 0.2365 | 0.0373 |
| 4mrf | opls | 1.55 | 0.0938 | 2.27 | 0 | 96.58 | 0 | 0 | 2.82 | 1.29 | 0.1585 | 0.1872 | 0.0287 |
| 4mrf | noschrodinger | 1.55 | 0.0082 | 1.07 | 0 | 95.27 | 0.71 | 0 | 4.02 | 1.52 | 0.1594 | 0.184 | 0.0246 |
| 4mrt | opls | 2 | 0.0959 | 2.33 | 0 | 98.94 | 0.77 | 1 | 0.2 | 0.58 | 0.2077 | 0.2301 | 0.0224 |
| 4mrt | noschrodinger | 2 | 0.003 | 0.81 | 0.34 | 97.65 | 3.42 | 0 | 2.44 | 1.51 | 0.1872 | 0.2248 | 0.0376 |

| | | | | | | | | | | | | | |
|---|---|---|---|---|---|---|---|---|---|---|---|---|---|
| 4mvc | opls | 3 | 0.087 | 2.3 | 0.34 | 96.27 | 5.52 | 0 | 0.81 | 1.57 | 0.2338 | 0.2878 | 0.054 |
| 4mvc | noschrodinger | 3 | 0.0028 | 0.76 | 0 | 96.26 | 10.32 | 0 | 6.94 | 2.41 | 0.2187 | 0.2766 | 0.0579 |
| 4mwz | opls | 1.5 | 0.1041 | 2.55 | 0 | 97.03 | 0.86 | 5 | 4.86 | 1.42 | 0.1377 | 0.1643 | 0.0266 |
| 4mwz | noschrodinger | 1.5 | 0.0063 | 1.02 | 0 | 97.48 | 1.07 | 0 | 3.29 | 1.25 | 0.143 | 0.1618 | 0.0188 |
| 4mxe | opls | 2.6 | 0.0901 | 2.44 | 0.63 | 95.9 | 2.27 | 6 | 2.19 | 1.55 | 0.2248 | 0.2672 | 0.0424 |
| 4mxe | noschrodinger | 2.6 | 0.0031 | 0.82 | 1.15 | 93.1 | 4.84 | 0 | 4.75 | 2.21 | 0.2202 | 0.2589 | 0.0387 |
| 4mym | opls | 1.9 | 0.093 | 2.18 | 0 | 99.28 | 2.46 | 0 | 0.43 | 0.95 | 0.1737 | 0.1951 | 0.0214 |
| 4mym | noschrodinger | 1.9 | 0.003 | 0.82 | 0 | 98.66 | 0.8 | 0 | 0.43 | 0.65 | 0.176 | 0.1979 | 0.0219 |
| 4myt | opls | 3.5 | 0.0902 | 2.36 | 1.27 | 91.26 | 8.09 | 7 | 1.16 | 2.03 | 0.2393 | 0.2815 | 0.0422 |
| 4myt | noschrodinger | 3.5 | 0.0032 | 0.83 | 2.76 | 81.44 | 17.22 | 0 | 5.25 | 2.94 | 0.2147 | 0.26 | 0.0453 |
| 4n04 | opls | 2.49 | 0.0911 | 2.27 | 0 | 97.13 | 1.21 | 0 | 0.59 | 0.92 | 0.21 | 0.2617 | 0.0517 |
| 4n04 | noschrodinger | 2.49 | 0.0027 | 0.78 | 0 | 96.31 | 5.95 | 0 | 0.88 | 1.61 | 0.2058 | 0.2556 | 0.0498 |
| 4n1f | opls | 2.09 | 0.0997 | 2.24 | 0 | 98.9 | 1.88 | 3 | 0.67 | 0.92 | 0.2107 | 0.2293 | 0.0186 |
| 4n1f | noschrodinger | 2.09 | 0.0027 | 0.76 | 0 | 97.8 | 1.86 | 0 | 1.33 | 1.11 | 0.1895 | 0.2247 | 0.0352 |
| 4n1g | opls | 1.5 | 0.1049 | 2.23 | 0 | 97.4 | 1.49 | 1 | 3.03 | 1.34 | 0.1428 | 0.1832 | 0.0404 |
| 4n1g | noschrodinger | 1.5 | 0.0037 | 0.9 | 0 | 98.96 | 1.79 | 0 | 3.67 | 1.35 | 0.1431 | 0.1816 | 0.0385 |
| 4n31 | opls | 2.2 | 0.0925 | 2.24 | 0 | 97.56 | 0.44 | 0 | 0 | 0.59 | 0.2116 | 0.2357 | 0.0241 |
| 4n31 | noschrodinger | 2.2 | 0.0091 | 1.13 | 0 | 95.2 | 2.2 | 0 | 1.61 | 1.5 | 0.1949 | 0.2279 | 0.033 |
| 4n5x | opls | 2.1 | 0.0993 | 2.27 | 0 | 98.68 | 5.11 | 1 | 1.6 | 1.45 | 0.1977 | 0.2251 | 0.0274 |
| 4n5x | noschrodinger | 2.1 | 0.003 | 0.82 | 0 | 98.04 | 7.97 | 0 | 2 | 1.65 | 0.1821 | 0.2222 | 0.0401 |
| 4n6o | opls | 1.8 | 0.0977 | 2.3 | 0.27 | 98.64 | 0.92 | 2 | 2.48 | 1.03 | 0.1637 | 0.205 | 0.0413 |
| 4n6o | noschrodinger | 1.8 | 0.0095 | 1.06 | 0.27 | 98.11 | 0.3 | 0 | 5.64 | 1.31 | 0.168 | 0.2039 | 0.0359 |
| 4nfd | opls | 1.71 | 0.0997 | 2.31 | 0 | 98.28 | 0.91 | 2 | 4.38 | 1.22 | 0.1507 | 0.2014 | 0.0507 |
| 4nfd | noschrodinger | 1.71 | 0.0064 | 1.02 | 0 | 95.76 | 2.68 | 0 | 1.46 | 1.5 | 0.1596 | 0.1991 | 0.0395 |
| 4nl0 | opls | 2.41 | 0.0914 | 2.29 | 0 | 97.25 | 2.15 | 4 | 0.69 | 1.12 | 0.2171 | 0.2402 | 0.0231 |
| 4nl0 | noschrodinger | 2.41 | 0.0035 | 0.89 | 0 | 95.54 | 7.92 | 0 | 2.32 | 2 | 0.2101 | 0.2348 | 0.0247 |
| 4nml | opls | 2.6 | 0.0931 | 2.28 | 0.44 | 98.25 | 2.05 | 1 | 1.29 | 1.09 | 0.222 | 0.2344 | 0.0124 |
| 4nml | noschrodinger | 2.6 | 0.0029 | 0.81 | 0.41 | 97.94 | 2.49 | 0 | 3.11 | 1.42 | 0.2018 | 0.2214 | 0.0196 |
| 4nmw | opls | 1.5 | 0.1012 | 2.53 | 0 | 97.84 | 1.75 | 2 | 4.99 | 1.49 | 0.136 | 0.1535 | 0.0175 |
| 4nmw | noschrodinger | 1.5 | 0.005 | 1.03 | 0 | 97.62 | 2.94 | 0 | 4.29 | 1.65 | 0.133 | 0.165 | 0.032 |
| 4nni | opls | 2.64 | 0.0912 | 2.24 | 0.32 | 97.45 | 4.92 | 2 | 1.36 | 1.5 | 0.2571 | 0.2694 | 0.0123 |
| 4nni | noschrodinger | 2.64 | 0.0035 | 0.76 | 0.31 | 98.11 | 7.89 | 0 | 4.65 | 1.92 | 0.2359 | 0.2636 | 0.0277 |
| 4nnr | opls | 1.98 | 0.0946 | 2.29 | 0 | 99.5 | 6.43 | 0 | 0.29 | 1.22 | 0.1687 | 0.2176 | 0.0489 |
| 4nnr | noschrodinger | 1.98 | 0.0028 | 0.97 | 0 | 98.49 | 8.77 | 0 | 1.76 | 1.65 | 0.1704 | 0.2034 | 0.033 |
| 4non | opls | 2.5 | 0.0921 | 2.26 | 0 | 98.1 | 1.13 | 1 | 0.37 | 0.67 | 0.208 | 0.2373 | 0.0293 |
| 4non | noschrodinger | 2.5 | 0.0026 | 0.76 | 0 | 96.48 | 1.34 | 0 | 0.61 | 1.03 | 0.1933 | 0.2316 | 0.0383 |
| 4nqj | opls | 2.15 | 0.0933 | 2.34 | 0 | 99.61 | 5.17 | 1 | 0.76 | 1.28 | 0.2598 | 0.2707 | 0.0109 |
| 4nqj | noschrodinger | 2.15 | 0.0029 | 0.74 | 0 | 99.05 | 7.39 | 0 | 2.07 | 1.64 | 0.2381 | 0.257 | 0.0189 |
| 4nrq | opls | 2.5 | 0.0871 | 2.28 | 0 | 94.23 | 1.82 | 2 | 1.54 | 1.49 | 0.2248 | 0.2681 | 0.0433 |
| 4nrq | noschrodinger | 2.5 | 0.0027 | 0.79 | 0.97 | 95.63 | 4.79 | 0 | 0.95 | 1.6 | 0.2339 | 0.2639 | 0.03 |
| 4ntg | opls | 2.55 | 0.0918 | 2.3 | 0 | 98.7 | 2.1 | 5 | 0.31 | 0.86 | 0.2062 | 0.2366 | 0.0304 |
| 4ntg | noschrodinger | 2.55 | 0.0028 | 0.74 | 0 | 97.22 | 2.7 | 0 | 0.47 | 1.13 | 0.1937 | 0.2233 | 0.0296 |
| 4nto | opls | 2.15 | 0.0957 | 2.3 | 0 | 98.62 | 1.8 | 4 | 0.1 | 0.73 | 0.2087 | 0.2449 | 0.0362 |
| 4nto | noschrodinger | 2.15 | 0.0033 | 0.8 | 0 | 98.15 | 2.4 | 0 | 1.23 | 1.13 | 0.2002 | 0.2366 | 0.0364 |
| 4nv4 | opls | 1.8 | 0.0988 | 2.37 | 0 | 99.1 | 2.82 | 2 | 2.94 | 1.43 | 0.1827 | 0.2123 | 0.0296 |
| 4nv4 | noschrodinger | 1.8 | 0.0057 | 0.93 | 0 | 98.65 | 1.4 | 0 | 1.22 | 0.95 | 0.1876 | 0.2084 | 0.0208 |
| 4nv5 | opls | 2.79 | 0.0903 | 2.9 | 1.21 | 90.71 | 7.5 | 36 | 4.33 | 2.41 | 0.2718 | 0.3056 | 0.0338 |
| 4nv5 | noschrodinger | 2.79 | 0.0049 | 0.92 | 0.79 | 96.27 | 4.71 | 0 | 6.34 | 2.11 | 0.2744 | 0.2963 | 0.0219 |
| 4nwk | opls | 1.62 | 0.1008 | 3.1 | 0 | 97.3 | 3.07 | 0 | 6.56 | 1.86 | 0.1421 | 0.1732 | 0.0311 |

| | | | | | | | | | | | |
|---|---|---|---|---|---|---|---|---|---|---|---|
| 4nwk | noschrodinger | 1.62 | 0.012 | 1.32 | 0 | 97.31 | 1.22 | 0 | 10.43 | 1.73 | 0.1416 | 0.1706 | 0.029 |
| 4nxg | opls | 2.09 | 0.093 | 2.22 | 0 | 97.95 | 1.64 | 0 | 0.57 | 0.87 | 0.2151 | 0.2422 | 0.0271 |
| 4nxg | noschrodinger | 2.09 | 0.0028 | 0.81 | 0 | 98.48 | 3.24 | 0 | 2.29 | 1.39 | 0.2007 | 0.2267 | 0.026 |
| 4nzl | opls | 1.8 | 0.1005 | 2.52 | 0.33 | 97.7 | 1.56 | 3 | 4.39 | 1.43 | 0.1532 | 0.1905 | 0.0373 |
| 4nzl | noschrodinger | 1.8 | 0.0045 | 0.98 | 0 | 97.08 | 0.38 | 0 | 4.33 | 1.38 | 0.1591 | 0.1883 | 0.0292 |
| 4nzw | opls | 3.58 | 0.0905 | 2.33 | 1.64 | 94.73 | 3.13 | 3 | 0.94 | 1.52 | 0.2254 | 0.2702 | 0.0448 |
| 4nzw | noschrodinger | 3.58 | 0.0027 | 0.76 | 3.03 | 90.2 | 5.23 | 0 | 3.79 | 2.26 | 0.2124 | 0.2627 | 0.0503 |
| 4o0p | opls | 3.8 | 0.09 | 2.32 | 0.11 | 96.79 | 1.58 | 0 | 0.54 | 1.03 | 0.1979 | 0.2455 | 0.0476 |
| 4o0p | noschrodinger | 3.8 | 0.0026 | 0.84 | 0.63 | 92.71 | 2.08 | 0 | 1.27 | 1.55 | 0.1956 | 0.2334 | 0.0378 |
| 4o1s | opls | 2.7 | 0.0947 | 2.25 | 0.6 | 96.71 | 4.58 | 1 | 0.74 | 1.45 | 0.2028 | 0.2328 | 0.03 |
| 4o1s | noschrodinger | 2.7 | 0.0028 | 0.8 | 0.89 | 93.45 | 8.74 | 0 | 5.36 | 2.43 | 0.1863 | 0.2204 | 0.0341 |
| 4o77 | opls | 2 | 0.0992 | 2.16 | 0 | 98.79 | 2.06 | 2 | 0.45 | 0.9 | 0.1816 | 0.2129 | 0.0313 |
| 4o77 | noschrodinger | 2 | 0.0032 | 0.81 | 0 | 98.8 | 2.04 | 0 | 1.14 | 1.06 | 0.1695 | 0.206 | 0.0365 |
| 4o79 | opls | 2 | 0.0922 | 2.22 | 0 | 97.83 | 1.43 | 2 | 1.13 | 0.98 | 0.202 | 0.2278 | 0.0258 |
| 4o79 | noschrodinger | 2 | 0.0033 | 0.72 | 0 | 98.56 | 1.43 | 0 | 3.27 | 1.24 | 0.2039 | 0.2263 | 0.0224 |
| 4o9k | opls | 1.85 | 0.0973 | 2.41 | 0 | 99.52 | 0 | 1 | 1.7 | 0.92 | 0.1551 | 0.1863 | 0.0312 |
| 4o9k | noschrodinger | 1.85 | 0.0038 | 0.93 | 0 | 98.82 | 0.49 | 0 | 3.17 | 1.11 | 0.1552 | 0.1817 | 0.0265 |
| 4o9s | opls | 2.3 | 0.0959 | 2.17 | 0 | 97.96 | 0.66 | 1 | 1.19 | 0.84 | 0.1808 | 0.2121 | 0.0313 |
| 4o9s | noschrodinger | 2.3 | 0.003 | 0.92 | 0 | 96.23 | 0.99 | 0 | 1.53 | 1.15 | 0.1726 | 0.2079 | 0.0353 |
| 4oaa | opls | 2.98 | 0.0914 | 2.32 | 0.27 | 95.89 | 3.4 | 5 | 0.16 | 1.25 | 0.295 | 0.32 | 0.025 |
| 4oaa | noschrodinger | 2.98 | 0.0031 | 0.77 | 0.9 | 91.09 | 4.77 | 0 | 2.55 | 2.07 | 0.3043 | 0.3186 | 0.0143 |
| 4ob1 | opls | 1.63 | 0.1071 | 2.54 | 0.48 | 97.38 | 1.05 | 7 | 5.39 | 1.43 | 0.1356 | 0.1693 | 0.0337 |
| 4ob1 | noschrodinger | 1.63 | 0.006 | 1.06 | 1.42 | 96.7 | 1.3 | 1 | 5.38 | 1.59 | 0.1393 | 0.1616 | 0.0223 |
| 4oc6 | opls | 2.64 | 0.0888 | 2.24 | 0 | 97.01 | 1.75 | 2 | 1.28 | 1.21 | 0.2742 | 0.292 | 0.0178 |
| 4oc6 | noschrodinger | 2.64 | 0.0044 | 0.85 | 0 | 96.44 | 4.51 | 0 | 1.65 | 1.65 | 0.2609 | 0.2797 | 0.0188 |
| 4ogt | opls | 1.54 | 0.0991 | 2.4 | 0 | 97.8 | 1.06 | 1 | 2.81 | 1.14 | 0.2164 | 0.2456 | 0.0292 |
| 4ogt | noschrodinger | 1.54 | 0.0073 | 1.23 | 0 | 98.97 | 2.13 | 0 | 2.25 | 1.25 | 0.22 | 0.2451 | 0.0251 |
| 4oh6 | opls | 3.56 | 0.0908 | 2.43 | 0.42 | 96.23 | 3.57 | 0 | 0.24 | 1.27 | 0.1878 | 0.2528 | 0.065 |
| 4oh6 | noschrodinger | 3.56 | 0.003 | 0.83 | 0.41 | 93.06 | 9.29 | 0 | 5.06 | 2.45 | 0.179 | 0.2271 | 0.0481 |
| 4ohb | opls | 2.4 | 0.0932 | 2.31 | 0 | 98.08 | 1.6 | 1 | 0.82 | 0.91 | 0.2007 | 0.2387 | 0.038 |
| 4ohb | noschrodinger | 2.4 | 0.0032 | 0.78 | 0.64 | 96.82 | 4.8 | 0 | 2.86 | 1.79 | 0.1899 | 0.231 | 0.0411 |
| 4oke | opls | 1.7 | 0.1014 | 2.53 | 0 | 96.77 | 1.43 | 1 | 3.57 | 1.47 | 0.1773 | 0.2167 | 0.0394 |
| 4oke | noschrodinger | 1.7 | 0.0035 | 0.92 | 0 | 97.13 | 2.83 | 0 | 3.2 | 1.61 | 0.1827 | 0.2078 | 0.0251 |
| 4olk | opls | 2.69 | 0.0959 | 2.25 | 0 | 96.28 | 3.3 | 1 | 0.96 | 1.43 | 0.185 | 0.2106 | 0.0256 |
| 4olk | noschrodinger | 2.69 | 0.0068 | 1.02 | 0.62 | 97.54 | 5.47 | 0 | 1.53 | 1.55 | 0.1602 | 0.2028 | 0.0426 |
| 4onc | opls | 1.83 | 0.0995 | 2.37 | 0 | 98.72 | 1.28 | 2 | 1.55 | 0.98 | 0.1562 | 0.1856 | 0.0294 |
| 4onc | noschrodinger | 1.83 | 0.0046 | 0.98 | 0.36 | 97.83 | 1.9 | 0 | 2.22 | 1.25 | 0.1628 | 0.1823 | 0.0195 |
| 4onw | opls | 1.65 | 0.1012 | 2.6 | 0.2 | 97.43 | 1.13 | 5 | 5.15 | 1.43 | 0.126 | 0.1506 | 0.0246 |
| 4onw | noschrodinger | 1.65 | 0.0136 | 1.49 | 0.19 | 97.53 | 1.13 | 0 | 3.83 | 1.31 | 0.127 | 0.1483 | 0.0213 |
| 4or4 | opls | 2.21 | 0.0963 | 2.31 | 0 | 98.12 | 3.75 | 1 | 1.99 | 1.4 | 0.1912 | 0.223 | 0.0318 |
| 4or4 | noschrodinger | 2.21 | 0.0028 | 0.78 | 0 | 96.27 | 5.41 | 0 | 3.25 | 1.92 | 0.1728 | 0.2214 | 0.0486 |
| 4ozn | opls | 2.6 | 0.089 | 2.39 | 0 | 98.26 | 4.53 | 0 | 0 | 1 | 0.2161 | 0.2516 | 0.0355 |
| 4ozn | noschrodinger | 2.6 | 0.0028 | 0.85 | 0 | 97.51 | 2.45 | 0 | 0.62 | 1.1 | 0.2019 | 0.2405 | 0.0386 |
| 4p1g | opls | 2.6 | 0.0891 | 2.23 | 0 | 98.59 | 2.22 | 0 | 0 | 0.76 | 0.2172 | 0.2428 | 0.0256 |
| 4p1g | noschrodinger | 2.6 | 0.0027 | 0.77 | 0 | 98.67 | 5.11 | 0 | 1.63 | 1.45 | 0.2363 | 0.2512 | 0.0149 |
| 4p1q | opls | 1.5 | 0.1029 | 2.28 | 0 | 99.1 | 0.48 | 4 | 2.66 | 1.05 | 0.1344 | 0.1587 | 0.0243 |
| 4p1q | noschrodinger | 1.5 | 0.0092 | 1.15 | 0 | 99.1 | 1.9 | 0 | 3.45 | 1.35 | 0.1346 | 0.1553 | 0.0207 |
| 4p3f | opls | 1.7 | 0.101 | 2.32 | 0 | 99.21 | 1.42 | 1 | 2.5 | 1.15 | 0.177 | 0.215 | 0.038 |
| 4p3f | noschrodinger | 1.7 | 0.0081 | 0.89 | 0 | 99.22 | 1.42 | 0 | 4.12 | 1.31 | 0.18 | 0.2151 | 0.0351 |

| | | | | | | | | | | | | |
|---|---|---|---|---|---|---|---|---|---|---|---|---|
| 4p3y | opls | 2.15 | 0.0962 | 2.23 | 0.36 | 97.68 | 0.84 | 1 | 0.67 | 0.79 | 0.2032 | 0.2304 | 0.0272 |
| 4p3y | noschrodinger | 2.15 | 0.0029 | 0.8 | 0.36 | 96.27 | 1.47 | 0 | 1.45 | 1.26 | 0.1933 | 0.2226 | 0.0293 |
| 4p4l | opls | 2.01 | 0.092 | 2.34 | 0 | 98.69 | 1.12 | 4 | 0.97 | 0.83 | 0.206 | 0.2518 | 0.0458 |
| 4p4l | noschrodinger | 2.01 | 0.0033 | 0.87 | 0 | 98.32 | 1.67 | 0 | 1.06 | 0.98 | 0.2144 | 0.2518 | 0.0374 |
| 4p58 | opls | 2.06 | 0.0985 | 2.25 | 0 | 96.97 | 1.71 | 0 | 1.22 | 1.19 | 0.1839 | 0.2209 | 0.037 |
| 4p58 | noschrodinger | 2.06 | 0.0029 | 0.8 | 0 | 97.01 | 2.25 | 0 | 0.61 | 1.14 | 0.1839 | 0.2091 | 0.0252 |
| 4p5d | opls | 2.11 | 0.0964 | 2.35 | 1.43 | 94.29 | 3.86 | 3 | 1.29 | 1.69 | 0.194 | 0.2179 | 0.0239 |
| 4p5d | noschrodinger | 2.11 | 0.0028 | 0.88 | 1.42 | 93.62 | 6.81 | 0 | 4.74 | 2.3 | 0.1913 | 0.2177 | 0.0264 |
| 4p5y | opls | 2.5 | 0.0929 | 2.15 | 0 | 96.4 | 2.4 | 0 | 0.41 | 1.17 | 0.2096 | 0.2381 | 0.0285 |
| 4p5y | noschrodinger | 2.5 | 0.0026 | 0.76 | 0 | 94.84 | 4 | 0 | 3.31 | 1.94 | 0.195 | 0.2299 | 0.0349 |
| 4p6t | opls | 2.5 | 0.0936 | 2.36 | 0.35 | 96.81 | 4.44 | 3 | 1.84 | 1.63 | 0.2097 | 0.2588 | 0.0491 |
| 4p6t | noschrodinger | 2.5 | 0.0031 | 0.85 | 0.7 | 95.43 | 6.61 | 0 | 7.82 | 2.37 | 0.1894 | 0.2353 | 0.0459 |
| 4p77 | opls | 2.04 | 0.0985 | 2.3 | 0 | 97.55 | 2.34 | 0 | 0.46 | 1.04 | 0.1853 | 0.2119 | 0.0266 |
| 4p77 | noschrodinger | 2.04 | 0.0028 | 0.78 | 0 | 97.59 | 2.02 | 0 | 1.08 | 1.13 | 0.1761 | 0.2067 | 0.0306 |
| 4p79 | opls | 2.4 | 0.0874 | 2.18 | 0 | 97.81 | 1.6 | 0 | 0.36 | 0.83 | 0.2191 | 0.2531 | 0.034 |
| 4p79 | noschrodinger | 2.4 | 0.0026 | 0.68 | 0 | 97.74 | 0.75 | 0 | 1.83 | 1 | 0.2084 | 0.2474 | 0.039 |
| 4p7u | opls | 1.5 | 0.0976 | 2.32 | 0 | 97.98 | 0.93 | 2 | 6.22 | 1.35 | 0.1592 | 0.1898 | 0.0306 |
| 4p7u | noschrodinger | 1.5 | 0.0106 | 1.22 | 0 | 98 | 2.78 | 0 | 1.69 | 1.26 | 0.1588 | 0.1875 | 0.0287 |
| 4p8i | opls | 1.85 | 0.0956 | 2.28 | 0.43 | 95.11 | 0.48 | 5 | 1.24 | 1.18 | 0.1831 | 0.2124 | 0.0293 |
| 4p8i | noschrodinger | 1.85 | 0.0036 | 0.8 | 0.21 | 93.91 | 0.71 | 0 | 1.24 | 1.25 | 0.1882 | 0.2105 | 0.0223 |
| 4pc7 | opls | 3.6 | 0.0899 | 2.34 | 1.95 | 90.1 | 3.75 | 5 | 1.12 | 1.8 | 0.2261 | 0.2722 | 0.0461 |
| 4pc7 | noschrodinger | 3.6 | 0.0031 | 0.84 | 2.74 | 84.03 | 3.94 | 0 | 4.17 | 2.33 | 0.2263 | 0.2726 | 0.0463 |
| 4pde | opls | 2.8 | 0.0911 | 2.44 | 1.26 | 93.7 | 7.84 | 1 | 0.78 | 1.84 | 0.2253 | 0.2616 | 0.0363 |
| 4pde | noschrodinger | 2.8 | 0.0032 | 0.87 | 2.92 | 90.83 | 9.22 | 0 | 6.03 | 2.59 | 0.214 | 0.2421 | 0.0281 |
| 4phs | opls | 1.54 | 0.0993 | 2.35 | 0 | 96.55 | 3.72 | 3 | 4.19 | 1.86 | 0.2034 | 0.2443 | 0.0409 |
| 4phs | noschrodinger | 1.54 | 0.0132 | 1.28 | 0.36 | 96 | 4.45 | 0 | 5.07 | 2.04 | 0.1976 | 0.2402 | 0.0426 |
| 4pi7 | opls | 1.6 | 0.1015 | 2.39 | 0 | 97.71 | 1.55 | 3 | 2.16 | 1.2 | 0.145 | 0.1701 | 0.0251 |
| 4pi7 | noschrodinger | 1.6 | 0.0101 | 1.15 | 0 | 96.8 | 1.55 | 0 | 4.59 | 1.57 | 0.1483 | 0.1678 | 0.0195 |
| 4pis | opls | 2.1 | 0.0925 | 2.31 | 0 | 99.04 | 2.2 | 0 | 0.57 | 0.95 | 0.2566 | 0.2788 | 0.0222 |
| 4pis | noschrodinger | 2.1 | 0.0029 | 0.8 | 0 | 97.63 | 4.37 | 0 | 0.57 | 1.26 | 0.2362 | 0.2665 | 0.0303 |
| 4pqn | opls | 1.71 | 0.0991 | 2.27 | 0.43 | 97.84 | 2.27 | 3 | 2.68 | 1.36 | 0.1657 | 0.1968 | 0.0311 |
| 4pqn | noschrodinger | 1.71 | 0.0033 | 0.82 | 0.43 | 97.01 | 2.73 | 0 | 2.68 | 1.56 | 0.1682 | 0.1996 | 0.0314 |
| 4pr3 | opls | 2.61 | 0.0904 | 2.41 | 0.89 | 97.92 | 3.19 | 4 | 0.84 | 1.16 | 0.2176 | 0.2596 | 0.042 |
| 4pr3 | noschrodinger | 2.61 | 0.0043 | 0.9 | 1.26 | 95.2 | 4 | 0 | 3.53 | 1.93 | 0.2008 | 0.2542 | 0.0534 |
| 4pr7 | opls | 2.1 | 0.0883 | 2.22 | 0 | 98.03 | 2.74 | 0 | 0.99 | 1.12 | 0.2184 | 0.246 | 0.0276 |
| 4pr7 | noschrodinger | 2.1 | 0.0024 | 0.78 | 0 | 98.74 | 4.58 | 0 | 0.33 | 1.12 | 0.2054 | 0.2383 | 0.0329 |
| 4pse | opls | 1.65 | 0.094 | 2.3 | 0.48 | 96.38 | 2.92 | 5 | 2.31 | 1.6 | 0.2226 | 0.249 | 0.0264 |
| 4pse | noschrodinger | 1.65 | 0.0081 | 1.14 | 1.84 | 94.25 | 5.23 | 1 | 6.8 | 2.31 | 0.2212 | 0.2494 | 0.0282 |
| 4psu | opls | 2.2 | 0.0916 | 2.26 | 0 | 98.35 | 1.95 | 0 | 0.23 | 0.81 | 0.202 | 0.2388 | 0.0368 |
| 4psu | noschrodinger | 2.2 | 0.0028 | 0.82 | 0 | 96.81 | 3.29 | 0 | 0.46 | 1.25 | 0.1925 | 0.2213 | 0.0288 |
| 4pt1 | opls | 1.65 | 0.093 | 2.33 | 0 | 96.37 | 1.91 | 2 | 2.01 | 1.43 | 0.2017 | 0.2468 | 0.0451 |
| 4pt1 | noschrodinger | 1.65 | 0.0039 | 0.92 | 1.58 | 94.47 | 3.74 | 1 | 6.16 | 2.15 | 0.2085 | 0.2406 | 0.0321 |
| 4pur | opls | 2.95 | 0.091 | 2.3 | 0 | 97.78 | 6.42 | 0 | 0.75 | 1.4 | 0.2192 | 0.2392 | 0.02 |
| 4pur | noschrodinger | 2.95 | 0.003 | 0.78 | 0 | 98.21 | 13.48 | 0 | 3.43 | 1.99 | 0.1985 | 0.2254 | 0.0269 |
| 4q12 | opls | 2.85 | 0.0947 | 2.32 | 0 | 98.64 | 1.78 | 0 | 1.61 | 1.1 | 0.1757 | 0.1945 | 0.0188 |
| 4q12 | noschrodinger | 2.85 | 0.0032 | 0.8 | 0 | 97.83 | 3.55 | 0 | 1.61 | 1.37 | 0.1787 | 0.1947 | 0.016 |
| 4q18 | opls | 2 | 0.0904 | 2.48 | 0 | 97.3 | 2.93 | 1 | 2.28 | 1.49 | 0.22 | 0.2411 | 0.0211 |
| 4q18 | noschrodinger | 2 | 0.0039 | 0.84 | 0.22 | 95.36 | 3.88 | 0 | 2.56 | 1.81 | 0.2282 | 0.2497 | 0.0215 |
| 4q24 | opls | 2.9 | 0.091 | 2.35 | 0 | 98.64 | 7.03 | 1 | 0.84 | 1.4 | 0.2509 | 0.2879 | 0.037 |

| | | | | | | | | | | | | |
|---|---|---|---|---|---|---|---|---|---|---|---|---|
| 4q24 | noschrodinger | 2.9 | 0.0027 | 0.74 | 0 | 96.85 | 14.52 | 0 | 1.12 | 1.9 | 0.2494 | 0.2772 | 0.0278 |
| 4q2g | opls | 3.3 | 0.091 | 2.45 | 0.76 | 90.11 | 5.7 | 6 | 2.44 | 2.15 | 0.3112 | 0.3315 | 0.0203 |
| 4q2g | noschrodinger | 3.3 | 0.003 | 0.84 | 1.33 | 86.5 | 3.95 | 0 | 3.36 | 2.21 | 0.3059 | 0.3219 | 0.016 |
| 4q35 | opls | 2.39 | 0.0958 | 2.22 | 0.22 | 96.98 | 1.39 | 3 | 0.73 | 1.02 | 0.2015 | 0.2401 | 0.0386 |
| 4q35 | noschrodinger | 2.39 | 0.0027 | 0.8 | 0.33 | 96.34 | 1.88 | 0 | 2.81 | 1.52 | 0.1981 | 0.2319 | 0.0338 |
| 4q39 | opls | 2.2 | 0.0941 | 2.27 | 0 | 97.28 | 3.97 | 1 | 0.49 | 1.26 | 0.1709 | 0.2043 | 0.0334 |
| 4q39 | noschrodinger | 2.2 | 0.0033 | 0.89 | 0.19 | 96.3 | 4.44 | 0 | 0.49 | 1.41 | 0.1646 | 0.1967 | 0.0321 |
| 4q5h | opls | 2 | 0.0965 | 2.28 | 0 | 99.2 | 0.83 | 4 | 2.27 | 1 | 0.1917 | 0.2237 | 0.032 |
| 4q5h | noschrodinger | 2 | 0.0036 | 0.86 | 0 | 98.22 | 1.09 | 0 | 3.64 | 1.18 | 0.2013 | 0.2212 | 0.0199 |
| 4q95 | opls | 2.2 | 0.0931 | 2.22 | 0 | 96.53 | 0.84 | 0 | 0 | 0.73 | 0.2148 | 0.244 | 0.0292 |
| 4q95 | noschrodinger | 2.2 | 0.0033 | 0.79 | 0 | 96.18 | 2.92 | 0 | 0.91 | 1.39 | 0.2107 | 0.2466 | 0.0359 |
| 4qag | opls | 1.71 | 0.098 | 2.24 | 0 | 99.62 | 1.82 | 5 | 1.18 | 1.03 | 0.2051 | 0.2465 | 0.0414 |
| 4qag | noschrodinger | 1.71 | 0.0025 | 0.7 | 0 | 97.71 | 2.25 | 0 | 2.12 | 1.32 | 0.2149 | 0.2368 | 0.0219 |
| 4qak | opls | 2.02 | 0.0986 | 2.23 | 0 | 99.37 | 1.05 | 0 | 1.38 | 0.89 | 0.1889 | 0.2198 | 0.0309 |
| 4qak | noschrodinger | 2.02 | 0.0031 | 0.82 | 0 | 98.23 | 2.05 | 0 | 2.93 | 1.32 | 0.1794 | 0.2105 | 0.0311 |
| 4qb2 | opls | 1.5 | 0.0998 | 2.38 | 0 | 98.43 | 1.69 | 0 | 2.84 | 1.25 | 0.1454 | 0.1616 | 0.0162 |
| 4qb2 | noschrodinger | 1.5 | 0.009 | 1.19 | 0 | 98.43 | 2.54 | 0 | 1.89 | 1.26 | 0.1491 | 0.1617 | 0.0126 |
| 4qdy | opls | 2.74 | 0.0875 | 2.18 | 0 | 98.65 | 0 | 1 | 0 | 0.5 | 0.2447 | 0.2987 | 0.054 |
| 4qdy | noschrodinger | 2.74 | 0.003 | 0.83 | 0 | 97.96 | 0 | 0 | 1.03 | 0.81 | 0.2304 | 0.2855 | 0.0551 |
| 4qj0 | opls | 1.55 | 0.1005 | 2.37 | 0.1 | 97.96 | 1.68 | 11 | 3.39 | 1.31 | 0.1723 | 0.2039 | 0.0316 |
| 4qj0 | noschrodinger | 1.55 | 0.0072 | 1.11 | 0.1 | 97.58 | 2.09 | 0 | 3.97 | 1.51 | 0.1728 | 0.202 | 0.0292 |
| 4qji | opls | 2.65 | 0.0886 | 2.35 | 0 | 97.39 | 1.08 | 1 | 0.49 | 0.81 | 0.2141 | 0.232 | 0.0179 |
| 4qji | noschrodinger | 2.65 | 0.0026 | 0.78 | 0 | 97.13 | 0.71 | 0 | 1.66 | 1.07 | 0.1999 | 0.2338 | 0.0339 |
| 4qnd | opls | 1.7 | 0.0912 | 2.32 | 0 | 97.87 | 0 | 0 | 1.6 | 0.94 | 0.1777 | 0.1945 | 0.0168 |
| 4qnd | noschrodinger | 1.7 | 0.0052 | 0.91 | 0 | 97.89 | 0 | 0 | 2.14 | 1.01 | 0.1795 | 0.1964 | 0.0169 |
| 4qnz | opls | 2.55 | 0.0884 | 2.23 | 0.61 | 97.55 | 0 | 1 | 0.66 | 0.81 | 0.2156 | 0.2367 | 0.0211 |
| 4qnz | noschrodinger | 2.55 | 0.0027 | 0.78 | 0 | 98.35 | 2.03 | 0 | 2.66 | 1.29 | 0.2077 | 0.2243 | 0.0166 |
| 4qo6 | opls | 2.26 | 0.0953 | 2.3 | 0.97 | 97.58 | 1.06 | 2 | 0.29 | 0.72 | 0.2065 | 0.2434 | 0.0369 |
| 4qo6 | noschrodinger | 2.26 | 0.0028 | 0.77 | 0.97 | 97.1 | 2.65 | 0 | 0.59 | 1.18 | 0.2051 | 0.2393 | 0.0342 |
| 4qsj | opls | 1.7 | 0.101 | 2.4 | 0 | 97.67 | 2.32 | 1 | 4.64 | 1.59 | 0.155 | 0.1861 | 0.0311 |
| 4qsj | noschrodinger | 1.7 | 0.0034 | 0.9 | 0 | 96.89 | 2.53 | 0 | 3.48 | 1.63 | 0.1543 | 0.1844 | 0.0301 |
| 4qtn | opls | 2.8 | 0.0891 | 2.34 | 0.16 | 95.73 | 6.45 | 6 | 0.85 | 1.67 | 0.2726 | 0.2963 | 0.0237 |
| 4qtn | noschrodinger | 2.8 | 0.003 | 0.81 | 0.29 | 92.44 | 13.31 | 0 | 4.08 | 2.52 | 0.2738 | 0.2927 | 0.0189 |
| 4qwo | opls | 1.52 | 0.1057 | 2.26 | 0 | 95.87 | 0 | 5 | 3.13 | 1.39 | 0.1489 | 0.1748 | 0.0259 |
| 4qwo | noschrodinger | 1.52 | 0.0037 | 0.93 | 0 | 97.33 | 0 | 0 | 1.87 | 1.08 | 0.1488 | 0.1683 | 0.0195 |
| 4qxp | opls | 2.51 | 0.0924 | 3.02 | 1.12 | 92.7 | 8.28 | 24 | 6.45 | 2.51 | 0.206 | 0.2634 | 0.0574 |
| 4qxp | noschrodinger | 2.51 | 0.0033 | 0.84 | 0.56 | 95.28 | 11.95 | 0 | 4.42 | 2.37 | 0.2316 | 0.2504 | 0.0188 |
| 4qxw | opls | 2.04 | 0.093 | 2.2 | 0 | 97.09 | 1.12 | 0 | 0.29 | 0.81 | 0.2069 | 0.2293 | 0.0224 |
| 4qxw | noschrodinger | 2.04 | 0.0082 | 1.04 | 0 | 96.67 | 1.65 | 0 | 0.59 | 1.07 | 0.199 | 0.2265 | 0.0275 |
| 4r1e | opls | 1.98 | 0.0908 | 2.2 | 0 | 100 | 0 | 1 | 0 | 0.5 | 0.1853 | 0.2275 | 0.0422 |
| 4r1e | noschrodinger | 1.98 | 0.0049 | 0.82 | 0 | 100 | 0 | 0 | 0 | 0.5 | 0.1896 | 0.2296 | 0.04 |
| 4r2i | opls | 2.05 | 0.094 | 2.28 | 0 | 95 | 2.7 | 1 | 0 | 1.17 | 0.2059 | 0.2421 | 0.0362 |
| 4r2i | noschrodinger | 2.05 | 0.0037 | 0.86 | 0 | 96.1 | 5.88 | 0 | 2.28 | 1.86 | 0.1822 | 0.222 | 0.0398 |
| 4r3k | opls | 2.13 | 0.0928 | 2.27 | 0 | 98.7 | 0.73 | 0 | 0 | 0.5 | 0.1958 | 0.2267 | 0.0309 |
| 4r3k | noschrodinger | 2.13 | 0.0028 | 0.76 | 0 | 98.06 | 0.72 | 0 | 1.12 | 0.82 | 0.1863 | 0.2237 | 0.0374 |
| 4r3l | opls | 1.84 | 0.0958 | 2.36 | 0 | 96.75 | 0.73 | 1 | 2.22 | 1.2 | 0.1792 | 0.2177 | 0.0385 |
| 4r3l | noschrodinger | 1.84 | 0.0038 | 0.86 | 0 | 96.13 | 0.71 | 0 | 4.07 | 1.46 | 0.1851 | 0.2129 | 0.0278 |
| 4r42 | opls | 1.9 | 0.0967 | 2.34 | 0 | 98.59 | 1.27 | 6 | 2.19 | 1.07 | 0.1351 | 0.1644 | 0.0293 |
| 4r42 | noschrodinger | 1.9 | 0.0048 | 0.91 | 0 | 98.3 | 2.33 | 0 | 1.24 | 1.12 | 0.141 | 0.1631 | 0.0221 |

| | | | | | | | | | | | | | |
|---|---|---|---|---|---|---|---|---|---|---|---|---|---|
| 4r82 | opls | 1.66 | 0.0955 | 2.34 | 0 | 98.91 | 0.81 | 2 | 2.83 | 1.07 | 0.1535 | 0.1812 | 0.0277 |
| 4r82 | noschrodinger | 1.66 | 0.0068 | 1.08 | 0 | 99.38 | 1.17 | 0 | 3.68 | 1.21 | 0.1564 | 0.1793 | 0.0229 |
| 4r9k | opls | 1.5 | 0.0987 | 2.24 | 0 | 99.33 | 0 | 1 | 1.76 | 0.93 | 0.1421 | 0.1667 | 0.0246 |
| 4r9k | noschrodinger | 1.5 | 0.0063 | 1.02 | 0 | 99.34 | 0 | 0 | 0.95 | 0.78 | 0.1448 | 0.1669 | 0.0221 |
| 4raj | opls | 1.84 | 0.0999 | 2.2 | 0 | 98.59 | 1.18 | 0 | 0.88 | 0.82 | 0.1742 | 0.1991 | 0.0249 |
| 4raj | noschrodinger | 1.84 | 0.0036 | 0.8 | 0 | 98.6 | 1.17 | 0 | 1.76 | 0.98 | 0.174 | 0.1958 | 0.0218 |
| 4rda | opls | 2.5 | 0.0922 | 2.36 | 0 | 99.75 | 2.89 | 0 | 0.15 | 0.91 | 0.2391 | 0.2661 | 0.027 |
| 4rda | noschrodinger | 2.5 | 0.0026 | 0.78 | 0.25 | 99.24 | 4.02 | 0 | 2.94 | 1.54 | 0.242 | 0.267 | 0.025 |
| 4rdn | opls | 2.1 | 0.0988 | 2.28 | 0 | 97.52 | 1.91 | 2 | 1.43 | 1.19 | 0.2096 | 0.243 | 0.0334 |
| 4rdn | noschrodinger | 2.1 | 0.0024 | 0.74 | 0 | 96.59 | 2.66 | 0 | 1.23 | 1.38 | 0.2089 | 0.2414 | 0.0325 |
| 4rfr | opls | 1.5 | 0.104 | 2.5 | 0.51 | 98.98 | 0 | 0 | 1.86 | 0.95 | 0.1387 | 0.1671 | 0.0284 |
| 4rfr | noschrodinger | 1.5 | 0.0117 | 1.22 | 0.5 | 98.99 | 0 | 0 | 3.41 | 1.13 | 0.1401 | 0.1646 | 0.0245 |
| 4rgt | opls | 2 | 0.0967 | 2.17 | 0 | 95.66 | 1.64 | 4 | 0.29 | 1.07 | 0.2386 | 0.2637 | 0.0251 |
| 4rgt | noschrodinger | 2 | 0.0026 | 0.75 | 0 | 96.77 | 1.35 | 0 | 3.77 | 1.47 | 0.2305 | 0.2612 | 0.0307 |
| 4rgu | opls | 1.89 | 0.0962 | 2.39 | 0 | 100 | 4.25 | 3 | 1.22 | 1.32 | 0.1768 | 0.2168 | 0.04 |
| 4rgu | noschrodinger | 1.89 | 0.0102 | 1.02 | 0 | 100 | 3.36 | 0 | 3.04 | 1.49 | 0.1784 | 0.2155 | 0.0371 |
| 4rgx | opls | 2.1 | 0.0974 | 2.24 | 0 | 99.19 | 1.65 | 1 | 0.43 | 0.82 | 0.1947 | 0.2231 | 0.0284 |
| 4rgx | noschrodinger | 2.1 | 0.0026 | 0.72 | 0 | 99.63 | 1.59 | 0 | 1.52 | 1.05 | 0.1818 | 0.2142 | 0.0324 |
| 4rhp | opls | 2.39 | 0.0908 | 2.28 | 0 | 99.1 | 5.06 | 4 | 1.11 | 1.35 | 0.2001 | 0.2276 | 0.0275 |
| 4rhp | noschrodinger | 2.39 | 0.0031 | 0.8 | 0 | 97.68 | 7.08 | 0 | 2.37 | 1.73 | 0.1847 | 0.2187 | 0.034 |
| 4ri2 | opls | 2.35 | 0.0901 | 2.4 | 0 | 98.78 | 0.77 | 0 | 0.72 | 0.73 | 0.2174 | 0.2504 | 0.033 |
| 4ri2 | noschrodinger | 2.35 | 0.0028 | 0.76 | 0 | 99.1 | 1.52 | 0 | 2.35 | 1.15 | 0.2219 | 0.2514 | 0.0295 |
| 4rk6 | opls | 1.76 | 0.0979 | 2.21 | 0.42 | 97.03 | 1.07 | 3 | 1.63 | 1.1 | 0.1737 | 0.2027 | 0.029 |
| 4rk6 | noschrodinger | 1.76 | 0.0039 | 0.95 | 0.38 | 97.36 | 0.61 | 0 | 1.87 | 1.07 | 0.1752 | 0.2002 | 0.025 |
| 4rlc | opls | 1.6 | 0.0876 | 2.3 | 0 | 100 | 1.96 | 1 | 0.44 | 0.88 | 0.1905 | 0.2226 | 0.0321 |
| 4rlc | noschrodinger | 1.6 | 0.0099 | 1.12 | 0 | 99.21 | 2.86 | 0 | 4 | 1.53 | 0.1922 | 0.2157 | 0.0235 |
| 4rlt | opls | 2.05 | 0.0994 | 2.19 | 0 | 97.59 | 0.85 | 1 | 0.87 | 0.85 | 0.1758 | 0.1988 | 0.023 |
| 4rlt | noschrodinger | 2.05 | 0.0027 | 0.75 | 0 | 97.64 | 0.42 | 0 | 0.43 | 0.73 | 0.1753 | 0.1972 | 0.0219 |
| 4rmp | opls | 2.51 | 0.0965 | 2.32 | 0 | 97.44 | 0.39 | 0 | 0.99 | 0.9 | 0.1872 | 0.2168 | 0.0296 |
| 4rmp | noschrodinger | 2.51 | 0.0029 | 0.83 | 0 | 98.73 | 1.17 | 0 | 2.97 | 1.14 | 0.18 | 0.2158 | 0.0358 |
| 4rs2 | opls | 1.55 | 0.1021 | 2.44 | 0 | 98.33 | 0.96 | 5 | 4.97 | 1.26 | 0.16 | 0.1998 | 0.0398 |
| 4rs2 | noschrodinger | 1.55 | 0.0045 | 0.99 | 0 | 98.61 | 1.29 | 0 | 4.95 | 1.34 | 0.1628 | 0.1976 | 0.0348 |
| 4rso | opls | 3.5 | 0.0905 | 2.26 | 0.19 | 95.56 | 3.79 | 1 | 1.12 | 1.57 | 0.5285 | 0.5369 | 0.0084 |
| 4rso | noschrodinger | 3.5 | 0.0029 | 0.78 | 0.19 | 95.17 | 3.34 | 0 | 4.35 | 1.95 | 0.5295 | 0.5411 | 0.0116 |
| 4rww | opls | 1.6 | 0.0978 | 2.32 | 0 | 100 | 0.37 | 1 | 1.51 | 0.89 | 0.1531 | 0.1831 | 0.03 |
| 4rww | noschrodinger | 1.6 | 0.014 | 1.49 | 0 | 99.68 | 0.73 | 0 | 0.95 | 0.78 | 0.1534 | 0.1818 | 0.0284 |
| 4ryn | opls | 2.01 | 0.0862 | 2.18 | 0 | 100 | 0 | 0 | 0.68 | 0.72 | 0.2134 | 0.234 | 0.0206 |
| 4ryn | noschrodinger | 2.01 | 0.0027 | 1.08 | 0 | 98.68 | 0 | 0 | 0.71 | 0.73 | 0.2009 | 0.2259 | 0.025 |
| 4rz3 | opls | 1.9 | 0.0969 | 2.27 | 0.62 | 98.13 | 0 | 3 | 0.8 | 0.75 | 0.2068 | 0.2554 | 0.0486 |
| 4rz3 | noschrodinger | 1.9 | 0.0036 | 0.89 | 0.21 | 97.31 | 1.03 | 0 | 3.06 | 1.24 | 0.2092 | 0.2514 | 0.0422 |
| 4s0k | opls | 2.1 | 0.0933 | 2.19 | 0 | 98.5 | 3.23 | 0 | 0 | 0.89 | 0.1747 | 0.1968 | 0.0221 |
| 4s0k | noschrodinger | 2.1 | 0.007 | 1.01 | 0 | 98.52 | 3.17 | 0 | 2.95 | 1.47 | 0.1678 | 0.1923 | 0.0245 |
| 4s12 | opls | 1.55 | 0.1036 | 2.57 | 0.23 | 98.95 | 0.79 | 6 | 5.05 | 1.27 | 0.145 | 0.1718 | 0.0268 |
| 4s12 | noschrodinger | 1.55 | 0.0085 | 1.1 | 0.23 | 99.07 | 0.66 | 0 | 4 | 1.19 | 0.1466 | 0.1688 | 0.0222 |
| 4s14 | opls | 3.54 | 0.0902 | 2.33 | 0 | 97.98 | 2.68 | 0 | 0 | 0.83 | 0.2014 | 0.2346 | 0.0332 |
| 4s14 | noschrodinger | 3.54 | 0.0028 | 0.87 | 1.2 | 97.6 | 2.64 | 0 | 2.6 | 1.45 | 0.1946 | 0.2241 | 0.0295 |
| 4s1b | opls | 2.1 | 0.0956 | 2.3 | 0.24 | 99.28 | 0.81 | 0 | 0.72 | 0.73 | 0.1584 | 0.2011 | 0.0427 |
| 4s1b | noschrodinger | 2.1 | 0.0028 | 0.8 | 0.24 | 99.29 | 0.8 | 0 | 1.29 | 0.85 | 0.1663 | 0.194 | 0.0277 |
| 4s1h | opls | 1.6 | 0.0968 | 2.36 | 0 | 98.39 | 0.98 | 1 | 2.11 | 0.98 | 0.1784 | 0.2043 | 0.0259 |

| | | | | | | | | | | | | | |
|---|---|---|---|---|---|---|---|---|---|---|---|---|---|
| 4s1h | noschrodinger | 1.6 | 0.0106 | 1.2 | 0 | 98.75 | 1.57 | 0 | 1.33 | 1.01 | 0.1787 | 0.2024 | 0.0237 |
| 4s1k | opls | 2.2 | 0.0976 | 2.29 | 0.43 | 97.01 | 1.45 | 2 | 1.05 | 1.1 | 0.1694 | 0.2201 | 0.0507 |
| 4s1k | noschrodinger | 2.2 | 0.0031 | 0.89 | 0.43 | 97.44 | 1.93 | 0 | 0.79 | 1.08 | 0.1576 | 0.1989 | 0.0413 |
| 4s24 | opls | 1.7 | 0.0976 | 2.37 | 0 | 98.2 | 0.61 | 4 | 5.31 | 1.28 | 0.1595 | 0.1936 | 0.0341 |
| 4s24 | noschrodinger | 1.7 | 0.0074 | 0.99 | 0 | 97.34 | 0.57 | 0 | 4.13 | 1.32 | 0.1614 | 0.1929 | 0.0315 |
| 4tl6 | opls | 1.76 | 0.0962 | 2.39 | 0 | 97.98 | 0.52 | 7 | 2.72 | 1.06 | 0.1714 | 0.1969 | 0.0255 |
| 4tl6 | noschrodinger | 1.76 | 0.0042 | 0.9 | 0 | 98.46 | 0.51 | 0 | 3.18 | 1.11 | 0.1754 | 0.1958 | 0.0204 |
| 4tqf | opls | 2.71 | 0.0926 | 2.28 | 0 | 97.2 | 1.72 | 0 | 0.23 | 0.91 | 0.1913 | 0.2151 | 0.0238 |
| 4tqf | noschrodinger | 2.71 | 0.0037 | 0.84 | 0.4 | 97.62 | 3.42 | 0 | 1.14 | 1.31 | 0.1806 | 0.2079 | 0.0273 |
| 4ts5 | opls | 2.39 | 0.095 | 2.3 | 0.97 | 97.34 | 1.58 | 2 | 0.43 | 0.93 | 0.1841 | 0.2219 | 0.0378 |
| 4ts5 | noschrodinger | 2.39 | 0.003 | 0.78 | 1.45 | 96.38 | 1.84 | 0 | 2.71 | 1.5 | 0.1768 | 0.2189 | 0.0421 |
| 4ts7 | opls | 2.79 | 0.0928 | 2.32 | 0.72 | 98.31 | 2.37 | 6 | 0.57 | 0.98 | 0.1742 | 0.223 | 0.0488 |
| 4ts7 | noschrodinger | 2.79 | 0.0032 | 0.8 | 0.72 | 97.1 | 4.21 | 0 | 3.14 | 1.74 | 0.1771 | 0.218 | 0.0409 |
| 4tv6 | opls | 2.6 | 0.0927 | 2.33 | 0 | 97.69 | 0.53 | 1 | 0.57 | 0.76 | 0.2442 | 0.2896 | 0.0454 |
| 4tv6 | noschrodinger | 2.6 | 0.0025 | 0.71 | 0.46 | 95.89 | 2.11 | 0 | 1.99 | 1.5 | 0.2374 | 0.27 | 0.0326 |
| 4tw3 | opls | 1.6 | 0.1033 | 2.29 | 0 | 99.52 | 0 | 1 | 2.43 | 1.02 | 0.162 | 0.1839 | 0.0219 |
| 4tw3 | noschrodinger | 1.6 | 0.0044 | 0.92 | 0 | 99.55 | 0 | 0 | 1.89 | 0.95 | 0.1662 | 0.1856 | 0.0194 |
| 4twk | opls | 2.6 | 0.0876 | 2.21 | 0.24 | 98.05 | 0.26 | 0 | 0.38 | 0.64 | 0.2863 | 0.3063 | 0.02 |
| 4twk | noschrodinger | 2.6 | 0.0031 | 0.77 | 0.21 | 98.35 | 0.52 | 0 | 1.28 | 0.85 | 0.2639 | 0.2925 | 0.0286 |
| 4u9n | opls | 2.19 | 0.0918 | 2.27 | 0 | 99.38 | 0.7 | 0 | 1.2 | 0.84 | 0.2314 | 0.2472 | 0.0158 |
| 4u9n | noschrodinger | 2.19 | 0.0027 | 0.74 | 0 | 99.43 | 1.4 | 0 | 1.22 | 0.95 | 0.2214 | 0.2441 | 0.0227 |
| 4u9r | opls | 2.17 | 0.0917 | 2.4 | 0 | 97.22 | 1.68 | 1 | 0 | 0.82 | 0.2317 | 0.2458 | 0.0141 |
| 4u9r | noschrodinger | 2.17 | 0.0028 | 0.8 | 0 | 96.55 | 3.36 | 0 | 1.76 | 1.56 | 0.2313 | 0.2432 | 0.0119 |
| 4u9s | opls | 1.7 | 0.099 | 2.45 | 0 | 96.74 | 1.71 | 2 | 1.17 | 1.21 | 0.1822 | 0.225 | 0.0428 |
| 4u9s | noschrodinger | 1.7 | 0.01 | 1.1 | 0 | 97.24 | 1.69 | 0 | 3.5 | 1.46 | 0.1874 | 0.2231 | 0.0357 |
| 4u9u | opls | 1.55 | 0.0995 | 2.34 | 0 | 97.64 | 0.41 | 0 | 4.18 | 1.28 | 0.1681 | 0.2049 | 0.0368 |
| 4u9u | noschrodinger | 1.55 | 0.0068 | 1 | 0 | 98.19 | 1.24 | 0 | 2.31 | 1.08 | 0.1709 | 0.1991 | 0.0282 |
| 4u9v | opls | 1.77 | 0.0985 | 2.47 | 0.58 | 98.26 | 1.2 | 1 | 0.62 | 0.77 | 0.1554 | 0.198 | 0.0426 |
| 4u9v | noschrodinger | 1.77 | 0.0145 | 1.34 | 0.51 | 97.95 | 1.2 | 0 | 4.72 | 1.32 | 0.1558 | 0.1978 | 0.042 |
| 4u9y | opls | 2.2 | 0.0961 | 2.23 | 0 | 96.61 | 1.23 | 0 | 0.32 | 0.91 | 0.2211 | 0.2407 | 0.0196 |
| 4u9y | noschrodinger | 2.2 | 0.0027 | 0.77 | 0 | 96.77 | 3.07 | 0 | 0.32 | 1.19 | 0.1927 | 0.2296 | 0.0369 |
| 4ua3 | opls | 1.85 | 0.0945 | 2.28 | 0 | 99.67 | 0.62 | 1 | 1.25 | 0.85 | 0.2334 | 0.2834 | 0.05 |
| 4ua3 | noschrodinger | 1.85 | 0.0037 | 0.81 | 0 | 99.2 | 1.82 | 0 | 2.21 | 1.19 | 0.2249 | 0.2714 | 0.0465 |
| 4uaj | opls | 2.7 | 0.0916 | 2.23 | 0 | 97.12 | 0 | 0 | 0.89 | 0.93 | 0.2389 | 0.2527 | 0.0138 |
| 4uaj | noschrodinger | 2.7 | 0.0027 | 0.77 | 0 | 95.68 | 1.27 | 0 | 1.57 | 1.28 | 0.2069 | 0.237 | 0.0301 |
| 4uc3 | opls | 2.3 | 0.0884 | 2.38 | 0.77 | 97.7 | 2.46 | 4 | 1.74 | 1.29 | 0.2335 | 0.2711 | 0.0376 |
| 4uc3 | noschrodinger | 2.3 | 0.0034 | 0.83 | 0.37 | 94.38 | 1.92 | 0 | 4.17 | 1.8 | 0.2149 | 0.2605 | 0.0456 |
| 4unn | opls | 2.5 | 0.0906 | 2.24 | 0.28 | 97.15 | 1.21 | 0 | 0.9 | 0.99 | 0.1592 | 0.1908 | 0.0316 |
| 4unn | noschrodinger | 2.5 | 0.0034 | 0.78 | 0.55 | 95.04 | 3.15 | 0 | 1.26 | 1.57 | 0.1481 | 0.1727 | 0.0246 |
| 4urs | opls | 2.27 | 0.0925 | 2.4 | 0.62 | 97.19 | 1.72 | 0 | 2.09 | 1.31 | 0.2482 | 0.2863 | 0.0381 |
| 4urs | noschrodinger | 2.27 | 0.0026 | 0.84 | 0.31 | 96.88 | 4.83 | 0 | 3.14 | 1.81 | 0.227 | 0.2709 | 0.0439 |
| 4uvl | opls | 2 | 0.0989 | 2.19 | 0 | 98.76 | 0.28 | 0 | 1.35 | 0.86 | 0.1605 | 0.1895 | 0.029 |
| 4uvl | noschrodinger | 2 | 0.0083 | 0.93 | 0 | 98.28 | 0.84 | 0 | 1.8 | 0.94 | 0.1591 | 0.1934 | 0.0343 |
| 4ux7 | opls | 2.55 | 0.092 | 2.17 | 0 | 96.03 | 1.05 | 1 | 0.73 | 1.02 | 0.2127 | 0.2536 | 0.0409 |
| 4ux7 | noschrodinger | 2.55 | 0.0026 | 0.72 | 0.25 | 95.54 | 2.62 | 0 | 1.76 | 1.56 | 0.2059 | 0.2354 | 0.0295 |
| 4v08 | opls | 2.03 | 0.0978 | 2.3 | 0.25 | 97.52 | 0.3 | 2 | 0.6 | 0.8 | 0.2069 | 0.2476 | 0.0407 |
| 4v08 | noschrodinger | 2.03 | 0.0028 | 0.8 | 0.23 | 98.14 | 1.49 | 0 | 1.65 | 1.05 | 0.1968 | 0.2392 | 0.0424 |
| 4w4s | opls | 2 | 0.0983 | 2.4 | 0.19 | 98.48 | 1.16 | 4 | 2.21 | 1.04 | 0.1852 | 0.2235 | 0.0383 |
| 4w4s | noschrodinger | 2 | 0.0029 | 0.78 | 0.75 | 96.63 | 2.28 | 0 | 3.84 | 1.66 | 0.1874 | 0.2193 | 0.0319 |

| | | | | | | | | | | | | | |
|---|---|---|---|---|---|---|---|---|---|---|---|---|---|
| 4w69 | opls | 3.98 | 0.0893 | 2.37 | 0.25 | 96.18 | 0.82 | 4 | 0.73 | 0.99 | 0.3097 | 0.3496 | 0.0399 |
| 4w69 | noschrodinger | 3.98 | 0.0027 | 0.74 | 0 | 96.43 | 0 | 0 | 3.98 | 1.42 | 0.3077 | 0.3469 | 0.0392 |
| 4wab | opls | 2.7 | 0.0916 | 2.27 | 0 | 99.37 | 0.76 | 0 | 0.37 | 0.63 | 0.2146 | 0.236 | 0.0214 |
| 4wab | noschrodinger | 2.7 | 0.0094 | 1.09 | 0 | 98.11 | 2.26 | 0 | 0.74 | 1 | 0.191 | 0.2369 | 0.0459 |
| 4wei | opls | 2.3 | 0.0954 | 2.19 | 0.4 | 98.81 | 1 | 0 | 0.26 | 0.6 | 0.1984 | 0.2247 | 0.0263 |
| 4wei | noschrodinger | 2.3 | 0.003 | 1 | 0.39 | 99.22 | 1.49 | 0 | 1.82 | 1.07 | 0.1995 | 0.2201 | 0.0206 |
| 4wh5 | opls | 1.82 | 0.1039 | 2.37 | 0 | 96.94 | 1.38 | 4 | 4.12 | 1.48 | 0.1875 | 0.2339 | 0.0464 |
| 4wh5 | noschrodinger | 1.82 | 0.0053 | 0.91 | 0 | 96.79 | 0.68 | 0 | 4.5 | 1.42 | 0.1907 | 0.231 | 0.0403 |
| 4wh9 | opls | 1.5 | 0.106 | 2.46 | 0 | 97.13 | 1.18 | 3 | 4.5 | 1.44 | 0.1431 | 0.1701 | 0.027 |
| 4wh9 | noschrodinger | 1.5 | 0.0142 | 1.37 | 0 | 97.14 | 1.18 | 0 | 5.13 | 1.48 | 0.1445 | 0.1699 | 0.0254 |
| 4wlk | opls | 2.03 | 0.0968 | 2.13 | 0 | 97.52 | 0.41 | 0 | 0.22 | 0.68 | 0.1882 | 0.2092 | 0.021 |
| 4wlk | noschrodinger | 2.03 | 0.003 | 0.8 | 0 | 98.25 | 0 | 0 | 0.22 | 0.58 | 0.179 | 0.2101 | 0.0311 |
| 4wnb | opls | 1.69 | 0.0995 | 2.47 | 0 | 97.12 | 0 | 3 | 3.66 | 1.31 | 0.2457 | 0.2928 | 0.0471 |
| 4wnb | noschrodinger | 1.69 | 0.005 | 0.85 | 0.34 | 97.24 | 0 | 0 | 3.45 | 1.28 | 0.2553 | 0.2916 | 0.0363 |
| 4wqb | opls | 1.5 | 0.1005 | 2.47 | 0 | 98.19 | 0.58 | 0 | 2.21 | 1 | 0.1504 | 0.1785 | 0.0281 |
| 4wqb | noschrodinger | 1.5 | 0.0064 | 1 | 0 | 98.21 | 0.57 | 0 | 0.56 | 0.69 | 0.1541 | 0.1789 | 0.0248 |
| 4wtr | opls | 2.27 | 0.0971 | 2.25 | 0 | 98.11 | 0.45 | 2 | 0 | 0.5 | 0.171 | 0.2155 | 0.0445 |
| 4wtr | noschrodinger | 2.27 | 0.0033 | 0.84 | 0 | 97.35 | 0.89 | 0 | 0.96 | 0.91 | 0.1768 | 0.221 | 0.0442 |
| 4wvy | opls | 3.64 | 0.0908 | 2.35 | 0.25 | 96.06 | 7.7 | 3 | 1.45 | 1.82 | 0.22 | 0.2638 | 0.0438 |
| 4wvy | noschrodinger | 3.64 | 0.003 | 0.77 | 0.61 | 95.5 | 12.12 | 0 | 1.53 | 2.03 | 0.2066 | 0.2552 | 0.0486 |
| 4ww5 | opls | 2 | 0.0917 | 2.34 | 0.32 | 98.09 | 0.61 | 4 | 1.22 | 0.84 | 0.1675 | 0.2002 | 0.0327 |
| 4ww5 | noschrodinger | 2 | 0.0065 | 0.99 | 0 | 97.02 | 1.19 | 0 | 1.09 | 1.04 | 0.1727 | 0.1986 | 0.0259 |
| 4ww7 | opls | 1.67 | 0.0925 | 2.19 | 0 | 97.87 | 0.64 | 0 | 1.25 | 0.87 | 0.1769 | 0.1854 | 0.0085 |
| 4ww7 | noschrodinger | 1.67 | 0.004 | 0.91 | 0 | 97.73 | 0.94 | 0 | 1.28 | 0.91 | 0.1823 | 0.1897 | 0.0074 |
| 4wx0 | opls | 1.7 | 0.1024 | 2.44 | 0 | 98.42 | 0.46 | 5 | 2.63 | 1.05 | 0.1661 | 0.1927 | 0.0266 |
| 4wx0 | noschrodinger | 1.7 | 0.0036 | 0.82 | 0 | 98.82 | 0.23 | 0 | 3.94 | 1.18 | 0.1719 | 0.1888 | 0.0169 |
| 4wxs | opls | 1.9 | 0.1024 | 2.35 | 0 | 96.88 | 0.45 | 1 | 8.13 | 1.63 | 0.1582 | 0.2078 | 0.0496 |
| 4wxs | noschrodinger | 1.9 | 0.0121 | 1.17 | 0 | 96.89 | 1.78 | 0 | 11.01 | 1.94 | 0.1542 | 0.203 | 0.0488 |
| 4wz6 | opls | 2.05 | 0.092 | 2.25 | 0 | 99.15 | 2.29 | 0 | 0 | 0.77 | 0.2202 | 0.2425 | 0.0223 |
| 4wz6 | noschrodinger | 2.05 | 0.0082 | 1.52 | 0 | 98.8 | 5 | 2 | 4.65 | 1.77 | 0.1981 | 0.2351 | 0.037 |
| 4x24 | opls | 1.5 | 0.0984 | 2.29 | 0 | 96.83 | 1.33 | 3 | 2.49 | 1.32 | 0.158 | 0.1847 | 0.0267 |
| 4x24 | noschrodinger | 1.5 | 0.0055 | 1 | 0 | 96.81 | 1.06 | 0 | 1.25 | 1.06 | 0.161 | 0.1803 | 0.0193 |
| 4x3l | opls | 1.7 | 0.101 | 2.3 | 0 | 99.42 | 1 | 2 | 1.57 | 0.9 | 0.1889 | 0.2121 | 0.0232 |
| 4x3l | noschrodinger | 1.7 | 0.0036 | 0.85 | 0 | 99.03 | 0.75 | 0 | 1.68 | 0.92 | 0.1941 | 0.2104 | 0.0163 |
| 4x81 | opls | 1.59 | 0.0998 | 2.29 | 0 | 96.56 | 0.48 | 0 | 0.86 | 0.99 | 0.1615 | 0.1969 | 0.0354 |
| 4x81 | noschrodinger | 1.59 | 0.007 | 1.13 | 0 | 96.17 | 0.71 | 0 | 1.22 | 1.1 | 0.1643 | 0.198 | 0.0337 |
| 4xa7 | opls | 1.56 | 0.0953 | 2.38 | 0.47 | 95.26 | 0.57 | 3 | 1.42 | 1.21 | 0.1967 | 0.2387 | 0.042 |
| 4xa7 | noschrodinger | 1.56 | 0.008 | 0.98 | 0.45 | 94.64 | 1.12 | 0 | 3.42 | 1.54 | 0.2065 | 0.2352 | 0.0287 |
| 4xb4 | opls | 1.54 | 0.1028 | 2.28 | 0 | 99.26 | 0.45 | 0 | 1.09 | 0.81 | 0.1456 | 0.1649 | 0.0193 |
| 4xb4 | noschrodinger | 1.54 | 0.0127 | 1.23 | 0 | 99.28 | 0.88 | 0 | 1.53 | 0.89 | 0.1447 | 0.1654 | 0.0207 |
| 4xba | opls | 1.5 | 0.1 | 2.51 | 0 | 97.59 | 0 | 1 | 3.17 | 1.19 | 0.1586 | 0.1855 | 0.0269 |
| 4xba | noschrodinger | 1.5 | 0.0057 | 1 | 0 | 97.93 | 0.52 | 0 | 3.47 | 1.16 | 0.1647 | 0.183 | 0.0183 |
| 4xdg | opls | 1.5 | 0.1015 | 2.23 | 0 | 96.64 | 0 | 5 | 2.5 | 1.25 | 0.1392 | 0.1779 | 0.0387 |
| 4xdg | noschrodinger | 1.5 | 0.0041 | 0.96 | 0.22 | 97.14 | 0.5 | 0 | 1.84 | 1.1 | 0.1569 | 0.1764 | 0.0195 |
| 4xf6 | opls | 2.08 | 0.0974 | 2.83 | 0 | 96.81 | 1.13 | 2 | 7.62 | 1.65 | 0.2444 | 0.2854 | 0.041 |
| 4xf6 | noschrodinger | 2.08 | 0.0049 | 1.28 | 0 | 96.86 | 2.03 | 0 | 16.18 | 2.14 | 0.2269 | 0.2686 | 0.0417 |
| 4xiv | opls | 2.83 | 0.091 | 2.38 | 0.8 | 94.57 | 3.31 | 5 | 1.74 | 1.7 | 0.261 | 0.2766 | 0.0156 |
| 4xiv | noschrodinger | 2.83 | 0.0028 | 0.77 | 1 | 92.17 | 4.41 | 0 | 2.8 | 2.04 | 0.244 | 0.2625 | 0.0185 |
| 4xj2 | opls | 1.8 | 0.1039 | 2.33 | 0.64 | 97.44 | 0.81 | 0 | 2.03 | 1.08 | 0.2075 | 0.2535 | 0.046 |

| | | | | | | | | | | | | | |
|---|---|---|---|---|---|---|---|---|---|---|---|---|---|
| 4xj2 | noschrodinger | 1.8 | 0.0031 | 0.85 | 0.63 | 97.47 | 1.61 | 0 | 3.66 | 1.42 | 0.2052 | 0.2494 | 0.0442 |
| 4xli | opls | 2.5 | 0.0904 | 2.25 | 0.19 | 96.89 | 1.55 | 0 | 0.93 | 1.11 | 0.199 | 0.218 | 0.019 |
| 4xli | noschrodinger | 2.5 | 0.0033 | 0.83 | 0.77 | 95 | 2.4 | 0 | 2.34 | 1.65 | 0.1805 | 0.2039 | 0.0234 |
| 4xly | opls | 1.82 | 0.0987 | 2.49 | 0 | 99.4 | 2.67 | 3 | 2.41 | 1.35 | 0.1889 | 0.2233 | 0.0344 |
| 4xly | noschrodinger | 1.82 | 0.0038 | 0.88 | 0.39 | 97.85 | 2.39 | 0 | 4.1 | 1.52 | 0.1941 | 0.2205 | 0.0264 |
| 4xmb | opls | 2.43 | 0.0965 | 2.29 | 0 | 97.18 | 0.43 | 0 | 1.83 | 1.09 | 0.195 | 0.228 | 0.033 |
| 4xmb | noschrodinger | 2.43 | 0.0026 | 0.74 | 0 | 96.83 | 2.15 | 0 | 2.52 | 1.48 | 0.175 | 0.2184 | 0.0434 |
| 4xoe | opls | 2.4 | 0.1032 | 2.15 | 0 | 97.55 | 0.85 | 0 | 0.69 | 0.82 | 0.1882 | 0.2076 | 0.0194 |
| 4xoe | noschrodinger | 2.4 | 0.0026 | 0.77 | 0 | 97.92 | 0.42 | 0 | 0.69 | 0.74 | 0.1706 | 0.2018 | 0.0312 |
| 4xpl | opls | 1.95 | 0.0944 | 2.29 | 0 | 100 | 0 | 0 | 1.62 | 0.91 | 0.1702 | 0.2032 | 0.033 |
| 4xpl | noschrodinger | 1.95 | 0.0035 | 0.85 | 0 | 98.54 | 0 | 0 | 1.64 | 0.91 | 0.1764 | 0.2025 | 0.0261 |
| 4xrb | opls | 1.89 | 0.0938 | 2.43 | 0 | 98.3 | 1.53 | 1 | 1.58 | 1.04 | 0.1926 | 0.2334 | 0.0408 |
| 4xrb | noschrodinger | 1.89 | 0.0045 | 0.94 | 0 | 98.76 | 2.03 | 0 | 4.49 | 1.46 | 0.1912 | 0.2273 | 0.0361 |
| 4xsh | opls | 2.5 | 0.0916 | 2.3 | 0 | 96.81 | 3.61 | 0 | 0.95 | 1.4 | 0.2033 | 0.2425 | 0.0392 |
| 4xsh | noschrodinger | 2.5 | 0.0028 | 0.78 | 0 | 96.83 | 7.78 | 0 | 2.86 | 1.95 | 0.1844 | 0.2318 | 0.0474 |
| 4xt5 | opls | 2.13 | 0.0935 | 2.39 | 0 | 96.31 | 1.01 | 0 | 1.05 | 1.05 | 0.2094 | 0.2468 | 0.0374 |
| 4xt5 | noschrodinger | 2.13 | 0.0032 | 0.8 | 0 | 95.08 | 3.52 | 0 | 4.19 | 1.96 | 0.2039 | 0.2398 | 0.0359 |
| 4xu0 | opls | 1.6 | 0.1041 | 2.29 | 0 | 97.88 | 0.5 | 0 | 1.72 | 0.95 | 0.1665 | 0.1897 | 0.0232 |
| 4xu0 | noschrodinger | 1.6 | 0.0049 | 1.15 | 0.19 | 97.31 | 0.25 | 0 | 4.06 | 1.32 | 0.1664 | 0.1877 | 0.0213 |
| 4xu3 | opls | 2.24 | 0.0949 | 2.35 | 0.19 | 97.08 | 1.01 | 0 | 0.74 | 0.9 | 0.1862 | 0.229 | 0.0428 |
| 4xu3 | noschrodinger | 2.24 | 0.0031 | 0.81 | 0.19 | 96.71 | 0.5 | 0 | 1.86 | 1.16 | 0.1797 | 0.2246 | 0.0449 |
| 4xu5 | opls | 2.1 | 0.0892 | 2.41 | 0 | 98.35 | 0.75 | 0 | 3.9 | 1.18 | 0.1834 | 0.2017 | 0.0183 |
| 4xu5 | noschrodinger | 2.1 | 0.0029 | 0.8 | 0 | 98.93 | 1.47 | 0 | 1.63 | 1.04 | 0.1869 | 0.1997 | 0.0128 |
| 4xui | opls | 2.51 | 0.0904 | 2.35 | 0 | 96.23 | 0.6 | 1 | 1.05 | 1.06 | 0.2782 | 0.2902 | 0.012 |
| 4xui | noschrodinger | 2.51 | 0.0028 | 0.86 | 0.47 | 94.37 | 3.78 | 0 | 2.62 | 1.87 | 0.2567 | 0.2744 | 0.0177 |
| 4xur | opls | 1.67 | 0.0916 | 2.2 | 0 | 98.52 | 1.96 | 2 | 0.65 | 0.94 | 0.2886 | 0.3141 | 0.0255 |
| 4xur | noschrodinger | 1.67 | 0.0027 | 0.87 | 0.21 | 97.48 | 0.24 | 0 | 4.45 | 1.33 | 0.288 | 0.311 | 0.023 |
| 4xw2 | opls | 2 | 0.0947 | 2.25 | 0 | 96.13 | 0.61 | 0 | 0.33 | 0.88 | 0.2066 | 0.2183 | 0.0117 |
| 4xw2 | noschrodinger | 2 | 0.0032 | 0.84 | 0 | 96.15 | 0 | 0 | 2.28 | 1.27 | 0.1937 | 0.2116 | 0.0179 |
| 4xxh | opls | 2.4 | 0.0901 | 2.25 | 0 | 97.58 | 2.07 | 3 | 1.42 | 1.2 | 0.1524 | 0.1935 | 0.0411 |
| 4xxh | noschrodinger | 2.4 | 0.0029 | 0.81 | 0 | 97.23 | 2.32 | 0 | 0.66 | 1.14 | 0.1586 | 0.193 | 0.0344 |
| 4xz0 | opls | 2 | 0.0945 | 2.6 | 1.57 | 96.08 | 2.33 | 8 | 1.95 | 1.51 | 0.1949 | 0.236 | 0.0411 |
| 4xz0 | noschrodinger | 2 | 0.0026 | 0.75 | 0.39 | 94.92 | 0.93 | 0 | 1.46 | 1.23 | 0.2111 | 0.2347 | 0.0236 |
| 4y6d | opls | 1.55 | 0.1012 | 2.5 | 0 | 98.52 | 1.64 | 4 | 1.98 | 1.13 | 0.1777 | 0.2123 | 0.0346 |
| 4y6d | noschrodinger | 1.55 | 0.0073 | 1.09 | 0 | 97.83 | 0.4 | 0 | 1.98 | 1.01 | 0.1829 | 0.2138 | 0.0309 |
| 4yag | opls | 1.5 | 0.1037 | 2.26 | 0 | 98.5 | 0.24 | 3 | 1.81 | 0.94 | 0.1417 | 0.1675 | 0.0258 |
| 4yag | noschrodinger | 1.5 | 0.0048 | 0.99 | 0 | 98.14 | 0.47 | 0 | 1.32 | 0.86 | 0.1455 | 0.167 | 0.0215 |
| 4yat | opls | 2.18 | 0.0999 | 2.26 | 0 | 98.78 | 0.94 | 3 | 1.42 | 0.88 | 0.1384 | 0.1696 | 0.0312 |
| 4yat | noschrodinger | 2.18 | 0.0029 | 0.79 | 0 | 99.4 | 0.93 | 0 | 1.42 | 0.88 | 0.1272 | 0.1608 | 0.0336 |
| 4ybn | opls | 1.9 | 0.0975 | 2.32 | 0 | 98.79 | 1.79 | 3 | 3.97 | 1.38 | 0.18 | 0.2141 | 0.0341 |
| 4ybn | noschrodinger | 1.9 | 0.0049 | 1.15 | 0 | 98.31 | 2.66 | 1 | 5.85 | 1.64 | 0.1834 | 0.2097 | 0.0263 |
| 4yc0 | opls | 1.5 | 0.1058 | 2.48 | 0 | 98.63 | 0 | 0 | 1.71 | 0.92 | 0.1639 | 0.1951 | 0.0312 |
| 4yc0 | noschrodinger | 1.5 | 0.0099 | 1.07 | 0 | 99.09 | 0 | 0 | 1.99 | 0.97 | 0.1668 | 0.1921 | 0.0253 |
| 4yca | opls | 1.8 | 0.0985 | 2.33 | 0 | 96.97 | 0.39 | 1 | 2.12 | 1.16 | 0.1763 | 0.202 | 0.0257 |
| 4yca | noschrodinger | 1.8 | 0.0034 | 0.83 | 0 | 96.28 | 0.58 | 0 | 2.11 | 1.23 | 0.1791 | 0.1996 | 0.0205 |
| 4ycn | opls | 3.5 | 0.0906 | 2.33 | 0.2 | 95.26 | 3.1 | 6 | 1.22 | 1.54 | 0.2227 | 0.2682 | 0.0455 |
| 4ycn | noschrodinger | 3.5 | 0.0029 | 0.74 | 0.4 | 91.42 | 2.98 | 0 | 2.5 | 1.9 | 0.2172 | 0.2635 | 0.0463 |
| 4ye2 | opls | 3.1 | 0.0909 | 2.2 | 0.72 | 94.95 | 0.76 | 1 | 0.21 | 0.93 | 0.2598 | 0.3006 | 0.0408 |
| 4ye2 | noschrodinger | 3.1 | 0.0029 | 0.75 | 0 | 96.77 | 3.01 | 0 | 1.27 | 1.41 | 0.2483 | 0.2913 | 0.043 |

| | | | | | | | | | | | | | |
|---|---|---|---|---|---|---|---|---|---|---|---|---|---|
| 4yfz | opls | 1.5 | 0.1102 | 2.5 | 0 | 94.97 | 0 | 2 | 3.79 | 1.52 | 0.1607 | 0.2046 | 0.0439 |
| 4yfz | noschrodinger | 1.5 | 0.0138 | 1.45 | 0 | 94.38 | 0 | 0 | 6.07 | 1.72 | 0.1631 | 0.205 | 0.0419 |
| 4ygq | opls | 2 | 0.0972 | 2.24 | 0 | 97.63 | 0.52 | 2 | 1.66 | 1 | 0.1642 | 0.1964 | 0.0322 |
| 4ygq | noschrodinger | 2 | 0.0048 | 0.96 | 0 | 98.59 | 1.02 | 0 | 1.38 | 0.88 | 0.1608 | 0.1982 | 0.0374 |
| 4ymi | opls | 1.7 | 0.0984 | 2.52 | 0 | 98.21 | 0 | 7 | 1.67 | 0.92 | 0.1597 | 0.1992 | 0.0395 |
| 4ymi | noschrodinger | 1.7 | 0.0038 | 0.87 | 0.27 | 98.91 | 0.33 | 0 | 1.86 | 0.95 | 0.1731 | 0.1972 | 0.0241 |
| 4yp6 | opls | 1.9 | 0.1005 | 2.45 | 0 | 99.39 | 0.45 | 6 | 1.44 | 0.88 | 0.1628 | 0.2006 | 0.0378 |
| 4yp6 | noschrodinger | 1.9 | 0.0087 | 1.06 | 0 | 99.2 | 0.67 | 0 | 0.36 | 0.63 | 0.1682 | 0.1977 | 0.0295 |
| 4yqa | opls | 1.5 | 0.0987 | 2.56 | 0 | 99.51 | 0.5 | 1 | 1.53 | 0.9 | 0.1703 | 0.1968 | 0.0265 |
| 4yqa | noschrodinger | 1.5 | 0.0081 | 1.26 | 0 | 99.57 | 0 | 0 | 3.07 | 1.1 | 0.1752 | 0.1964 | 0.0212 |
| 4yqf | opls | 2.73 | 0.0904 | 2.28 | 0 | 97.83 | 1.63 | 1 | 0.49 | 0.87 | 0.2222 | 0.2482 | 0.026 |
| 4yqf | noschrodinger | 2.73 | 0.0039 | 0.83 | 0.4 | 96.15 | 2.07 | 0 | 2.88 | 1.58 | 0.1964 | 0.2399 | 0.0435 |
| 4z07 | opls | 2.5 | 0.0957 | 2.3 | 0.17 | 97.47 | 0.18 | 4 | 1.36 | 0.97 | 0.1855 | 0.2245 | 0.039 |
| 4z07 | noschrodinger | 2.5 | 0.008 | 1.16 | 0 | 97.49 | 1.08 | 0 | 3.22 | 1.24 | 0.1807 | 0.2185 | 0.0378 |
| 4z1o | opls | 2.15 | 0.1011 | 2.25 | 0 | 97.63 | 0.63 | 0 | 1.2 | 0.91 | 0.2199 | 0.2543 | 0.0344 |
| 4z1o | noschrodinger | 2.15 | 0.0027 | 0.75 | 0 | 97.07 | 1.89 | 0 | 5.16 | 1.65 | 0.1936 | 0.2333 | 0.0397 |
| 4z28 | opls | 1.66 | 0.0925 | 2.3 | 0 | 99.19 | 1.96 | 3 | 3.16 | 1.33 | 0.2008 | 0.2259 | 0.0251 |
| 4z28 | noschrodinger | 1.66 | 0.0061 | 1.11 | 0 | 99.2 | 1.46 | 0 | 2.64 | 1.17 | 0.2015 | 0.2285 | 0.027 |
| 4z3h | opls | 1.5 | 0.1055 | 2.11 | 0 | 98.96 | 0.58 | 0 | 0.61 | 0.7 | 0.1401 | 0.1727 | 0.0326 |
| 4z3h | noschrodinger | 1.5 | 0.0077 | 1.1 | 0 | 98.46 | 0.57 | 0 | 3.66 | 1.16 | 0.1422 | 0.1749 | 0.0327 |
| 4z4k | opls | 2.79 | 0.0911 | 2.28 | 0.18 | 97.62 | 5.77 | 3 | 0.67 | 1.38 | 0.2237 | 0.265 | 0.0413 |
| 4z4k | noschrodinger | 2.79 | 0.0031 | 0.85 | 0.36 | 96.19 | 9.11 | 0 | 2.46 | 2.02 | 0.2014 | 0.2479 | 0.0465 |
| 4z4m | opls | 2.15 | 0.0941 | 2.24 | 0 | 98.37 | 4.73 | 1 | 0.75 | 1.25 | 0.2634 | 0.2853 | 0.0219 |
| 4z4m | noschrodinger | 2.15 | 0.0028 | 0.83 | 0 | 97.98 | 5.36 | 0 | 3.49 | 1.7 | 0.2454 | 0.2714 | 0.026 |
| 4z7e | opls | 1.5 | 0.0996 | 2.26 | 0.37 | 99.07 | 1.85 | 2 | 3.01 | 1.3 | 0.1579 | 0.1748 | 0.0169 |
| 4z7e | noschrodinger | 1.5 | 0.0035 | 0.87 | 0.37 | 98.69 | 2.47 | 0 | 2.68 | 1.35 | 0.1604 | 0.177 | 0.0166 |
| 4z7j | opls | 1.5 | 0.1011 | 2.7 | 0 | 99.36 | 0.75 | 2 | 6.44 | 1.35 | 0.1284 | 0.1465 | 0.0181 |
| 4z7j | noschrodinger | 1.5 | 0.0084 | 1.14 | 0 | 98.73 | 1.12 | 0 | 4.9 | 1.29 | 0.1325 | 0.1463 | 0.0138 |
| 4z7x | opls | 1.55 | 0.1007 | 2.44 | 0 | 98.97 | 1.34 | 9 | 3.46 | 1.23 | 0.1666 | 0.1883 | 0.0217 |
| 4z7x | noschrodinger | 1.55 | 0.0063 | 1.02 | 0 | 99.51 | 1.56 | 0 | 2.29 | 1.15 | 0.1675 | 0.189 | 0.0215 |
| 4z8n | opls | 2.12 | 0.0946 | 2.25 | 0 | 97.73 | 1.32 | 3 | 0.84 | 0.91 | 0.216 | 0.2512 | 0.0352 |
| 4z8n | noschrodinger | 2.12 | 0.0027 | 0.71 | 0.17 | 97.57 | 1.5 | 0 | 1.58 | 1.13 | 0.2194 | 0.2461 | 0.0267 |
| 4zay | opls | 1.54 | 0.097 | 2.53 | 0 | 98.03 | 3.26 | 3 | 6.47 | 1.75 | 0.1483 | 0.1739 | 0.0256 |
| 4zay | noschrodinger | 1.54 | 0.0056 | 2.01 | 0 | 99.02 | 2.7 | 0 | 6.73 | 1.7 | 0.1684 | 0.1909 | 0.0225 |
| 4zby | opls | 1.7 | 0.1003 | 2.28 | 0 | 97.91 | 0 | 0 | 2.36 | 1.04 | 0.1765 | 0.2167 | 0.0402 |
| 4zby | noschrodinger | 1.7 | 0.0111 | 2.31 | 0 | 96.35 | 0.54 | 0 | 4.4 | 1.46 | 0.2097 | 0.2592 | 0.0495 |
| 4zcr | opls | 1.8 | 0.0976 | 2.3 | 0 | 99.14 | 0.88 | 2 | 1.39 | 0.87 | 0.1925 | 0.2127 | 0.0202 |
| 4zcr | noschrodinger | 1.8 | 0.0036 | 0.78 | 0 | 97.58 | 1.72 | 0 | 2.8 | 1.34 | 0.1974 | 0.2131 | 0.0157 |
| 4zdx | opls | 2 | 0.097 | 2.16 | 0 | 96.94 | 0.49 | 1 | 0.78 | 0.93 | 0.1866 | 0.2052 | 0.0186 |
| 4zdx | noschrodinger | 2 | 0.0028 | 0.76 | 0 | 97.03 | 0.49 | 0 | 0.52 | 0.85 | 0.1745 | 0.202 | 0.0275 |
| 4zev | opls | 1.8 | 0.0982 | 2.27 | 0 | 97.65 | 0.39 | 2 | 1.76 | 1.01 | 0.1895 | 0.2169 | 0.0274 |
| 4zev | noschrodinger | 1.8 | 0.0025 | 0.74 | 0 | 97.84 | 0.96 | 0 | 0.55 | 0.72 | 0.1892 | 0.2093 | 0.0201 |
| 4zex | opls | 2 | 0.0971 | 2.26 | 0 | 97.29 | 0.58 | 1 | 2.11 | 1.12 | 0.187 | 0.2291 | 0.0421 |
| 4zex | noschrodinger | 2 | 0.0024 | 0.73 | 0 | 96.94 | 0.19 | 0 | 0.44 | 0.84 | 0.1833 | 0.2254 | 0.0421 |
| 4zf7 | opls | 1.89 | 0.1003 | 2.45 | 0 | 99.17 | 1.25 | 4 | 4.19 | 1.27 | 0.1705 | 0.2071 | 0.0366 |
| 4zf7 | noschrodinger | 1.89 | 0.0095 | 1.07 | 0 | 99.6 | 2.07 | 0 | 5.38 | 1.53 | 0.1729 | 0.206 | 0.0331 |
| 4zfn | opls | 1.9 | 0.1026 | 2.25 | 0 | 96.47 | 0.95 | 3 | 2.04 | 1.21 | 0.1822 | 0.2199 | 0.0377 |
| 4zfn | noschrodinger | 1.9 | 0.0032 | 0.81 | 0 | 97.38 | 2.18 | 0 | 3.4 | 1.51 | 0.178 | 0.2141 | 0.0361 |
| 4zfw | opls | 2.54 | 0.0994 | 2.49 | 0 | 96.75 | 0.68 | 2 | 1.49 | 1.09 | 0.1894 | 0.2223 | 0.0329 |

| | | | | | | | | | | | | | |
|---|---|---|---|---|---|---|---|---|---|---|---|---|---|
| 4zfw | noschrodinger | 2.54 | 0.0035 | 0.83 | 0 | 96.03 | 1.58 | 0 | 4.59 | 1.66 | 0.1872 | 0.2281 | 0.0409 |
| 4zgm | opls | 1.8 | 0.0952 | 2.14 | 0 | 97.56 | 0.88 | 0 | 0 | 0.59 | 0.1634 | 0.1977 | 0.0343 |
| 4zgm | noschrodinger | 1.8 | 0.0044 | 0.93 | 0 | 98.39 | 0.88 | 0 | 1.86 | 0.95 | 0.1685 | 0.1968 | 0.0283 |
| 4zgn | opls | 2.9 | 0.0902 | 2.37 | 0.56 | 95 | 4.75 | 6 | 2.23 | 1.86 | 0.2374 | 0.2784 | 0.041 |
| 4zgn | noschrodinger | 2.9 | 0.0029 | 0.81 | 0.27 | 93.55 | 8.53 | 0 | 1.76 | 2.06 | 0.2543 | 0.2828 | 0.0285 |
| 4zgp | opls | 1.85 | 0.0975 | 2.36 | 0 | 98.09 | 1.51 | 7 | 3.96 | 1.32 | 0.1798 | 0.2139 | 0.0341 |
| 4zgp | noschrodinger | 1.85 | 0.0089 | 1.11 | 0 | 98.55 | 2.34 | 0 | 3.85 | 1.45 | 0.1792 | 0.2081 | 0.0289 |
| 4zib | opls | 2.05 | 0.096 | 2.22 | 0 | 97.99 | 0 | 0 | 0.45 | 0.66 | 0.1789 | 0.1972 | 0.0183 |
| 4zib | noschrodinger | 2.05 | 0.003 | 0.78 | 0 | 98.01 | 0.42 | 0 | 0.9 | 0.77 | 0.1718 | 0.1912 | 0.0194 |
| 4zir | opls | 2.88 | 0.0905 | 2.36 | 0.21 | 94.86 | 0.69 | 5 | 1.35 | 1.22 | 0.2495 | 0.2924 | 0.0429 |
| 4zir | noschrodinger | 2.88 | 0.0034 | 0.82 | 1.79 | 87.85 | 3.43 | 0 | 5.65 | 2.32 | 0.2347 | 0.2686 | 0.0339 |
| 4zjb | opls | 2.55 | 0.0926 | 2.28 | 0 | 95.71 | 3.7 | 1 | 0.65 | 1.44 | 0.2215 | 0.2531 | 0.0316 |
| 4zjb | noschrodinger | 2.55 | 0.0029 | 0.8 | 0 | 93.88 | 4.28 | 0 | 2.75 | 1.95 | 0.2181 | 0.2404 | 0.0223 |
| 4zjp | opls | 1.63 | 0.1045 | 2.28 | 0.38 | 98.85 | 0.95 | 1 | 2.4 | 1.02 | 0.1324 | 0.1768 | 0.0444 |
| 4zjp | noschrodinger | 1.63 | 0.0034 | 0.82 | 0.37 | 98.88 | 1.4 | 0 | 1.2 | 0.95 | 0.1374 | 0.1736 | 0.0362 |
| 4zk6 | opls | 1.89 | 0.0962 | 2.3 | 0 | 98.98 | 1.32 | 2 | 1.4 | 0.97 | 0.1874 | 0.2328 | 0.0454 |
| 4zk6 | noschrodinger | 1.89 | 0.0031 | 0.78 | 0 | 98.99 | 2.44 | 0 | 1.3 | 1.15 | 0.1804 | 0.2276 | 0.0472 |
| 4zmm | opls | 2.5 | 0.0938 | 2.67 | 0 | 98.13 | 8.95 | 3 | 0.58 | 1.42 | 0.2048 | 0.244 | 0.0392 |
| 4zmm | noschrodinger | 2.5 | 0.003 | 0.82 | 0 | 96.26 | 11.28 | 0 | 2.13 | 2.04 | 0.2046 | 0.2409 | 0.0363 |
| 4zo3 | opls | 1.67 | 0.1016 | 2.21 | 0.52 | 96.74 | 0.19 | 2 | 1.26 | 1.05 | 0.2113 | 0.2412 | 0.0299 |
| 4zo3 | noschrodinger | 1.67 | 0.0068 | 0.97 | 0.17 | 96.92 | 1.36 | 0 | 2.52 | 1.32 | 0.2068 | 0.2433 | 0.0365 |
| 4zoh | opls | 2.2 | 0.0967 | 2.3 | 0.36 | 96.87 | 1.82 | 3 | 0.83 | 1.14 | 0.2066 | 0.2371 | 0.0305 |
| 4zoh | noschrodinger | 2.2 | 0.0056 | 1.01 | 0.27 | 96.27 | 4.24 | 0 | 2.05 | 1.7 | 0.1793 | 0.2252 | 0.0459 |
| 4zr1 | opls | 2.6 | 0.0875 | 2.26 | 0.37 | 95.39 | 1.45 | 2 | 1.43 | 1.32 | 0.2512 | 0.268 | 0.0168 |
| 4zr1 | noschrodinger | 2.6 | 0.0028 | 0.78 | 0.55 | 94.68 | 3.09 | 0 | 3.27 | 1.86 | 0.2303 | 0.2488 | 0.0185 |
| 4zra | opls | 1.83 | 0.0975 | 2.26 | 0.78 | 96.61 | 2.29 | 3 | 1.53 | 1.39 | 0.2052 | 0.2493 | 0.0441 |
| 4zra | noschrodinger | 1.83 | 0.0031 | 0.91 | 0.78 | 96.37 | 5.52 | 0 | 4.25 | 2.01 | 0.207 | 0.2417 | 0.0347 |
| 4zrw | opls | 2.6 | 0.0931 | 2.19 | 0.72 | 94.96 | 1.5 | 0 | 1.27 | 1.33 | 0.1749 | 0.2052 | 0.0303 |
| 4zrw | noschrodinger | 2.6 | 0.0045 | 0.92 | 0.7 | 95.77 | 2.22 | 0 | 2.13 | 1.54 | 0.1584 | 0.1964 | 0.038 |
| 4zsi | opls | 1.65 | 0.1001 | 2.25 | 0.62 | 98.13 | 0.72 | 1 | 3.36 | 1.13 | 0.1819 | 0.2096 | 0.0277 |
| 4zsi | noschrodinger | 1.65 | 0.0033 | 0.84 | 0 | 98.47 | 0.36 | 0 | 2.62 | 1.05 | 0.1857 | 0.2101 | 0.0244 |
| 4zvc | opls | 1.5 | 0.1028 | 2.41 | 0 | 98.67 | 1.36 | 0 | 2.29 | 1.11 | 0.1531 | 0.1829 | 0.0298 |
| 4zvc | noschrodinger | 1.5 | 0.0099 | 1.14 | 0 | 98.24 | 0 | 0 | 3.56 | 1.15 | 0.1523 | 0.1815 | 0.0292 |
| 4zy7 | opls | 1.7 | 0.102 | 2.3 | 0.87 | 98.27 | 0 | 2 | 4.63 | 1.24 | 0.1833 | 0.2108 | 0.0275 |
| 4zy7 | noschrodinger | 1.7 | 0.004 | 0.95 | 0.83 | 97.93 | 0 | 0 | 5.66 | 1.33 | 0.1862 | 0.2093 | 0.0231 |
| 4zyw | opls | 2.05 | 0.0961 | 2.28 | 0 | 97.34 | 0 | 0 | 0.62 | 0.83 | 0.2007 | 0.2135 | 0.0128 |
| 4zyw | noschrodinger | 2.05 | 0.0027 | 0.82 | 0 | 97.98 | 0 | 0 | 1.25 | 0.85 | 0.1869 | 0.204 | 0.0171 |
| 4zz3 | opls | 2.5 | 0.0901 | 2.32 | 0 | 97.7 | 0 | 1 | 1.95 | 1.03 | 0.2155 | 0.2438 | 0.0283 |
| 4zz3 | noschrodinger | 2.5 | 0.0035 | 0.81 | 0 | 95.45 | 0 | 0 | 2.63 | 1.37 | 0.2043 | 0.2352 | 0.0309 |
| 4zz8 | opls | 1.65 | 0.1023 | 2.21 | 0 | 98.05 | 0.95 | 0 | 3.17 | 1.11 | 0.1712 | 0.2021 | 0.0309 |
| 4zz8 | noschrodinger | 1.65 | 0.0035 | 1.02 | 0 | 97.3 | 1.42 | 0 | 2.68 | 1.3 | 0.177 | 0.1991 | 0.0221 |
| 5a64 | opls | 2.1 | 0.0925 | 2.29 | 0 | 98.53 | 0.32 | 0 | 0.8 | 0.75 | 0.2391 | 0.2592 | 0.0201 |
| 5a64 | noschrodinger | 2.1 | 0.0026 | 0.77 | 0 | 98.18 | 1.59 | 0 | 1.94 | 1.11 | 0.2323 | 0.2554 | 0.0231 |
| 5a66 | opls | 2.1 | 0.0968 | 2.21 | 0 | 98.8 | 0 | 0 | 0.99 | 0.79 | 0.1847 | 0.2083 | 0.0236 |
| 5a66 | noschrodinger | 2.1 | 0.0027 | 0.75 | 0 | 99.52 | 0.26 | 0 | 1.7 | 0.92 | 0.183 | 0.2065 | 0.0235 |
| 5a7t | opls | 2.4 | 0.092 | 2.26 | 0 | 97.48 | 0.45 | 0 | 0.97 | 0.89 | 0.1988 | 0.2255 | 0.0267 |
| 5a7t | noschrodinger | 2.4 | 0.0027 | 0.74 | 0 | 97.21 | 1.33 | 0 | 0.48 | 0.91 | 0.1875 | 0.2147 | 0.0272 |
| 5a89 | opls | 1.65 | 0.1024 | 2.35 | 0 | 100 | 0 | 2 | 2.67 | 1.05 | 0.1375 | 0.1683 | 0.0308 |
| 5a89 | noschrodinger | 1.65 | 0.0047 | 1.01 | 0 | 99.33 | 0.8 | 0 | 2.05 | 0.98 | 0.1454 | 0.1654 | 0.02 |

| | | | | | | | | | | | | | |
|---|---|---|---|---|---|---|---|---|---|---|---|---|---|
| 5a96 | opls | 1.91 | 0.0976 | 2.29 | 0.41 | 96.34 | 0 | 1 | 1.49 | 1.13 | 0.1627 | 0.206 | 0.0433 |
| 5a96 | noschrodinger | 1.91 | 0.0033 | 0.92 | 0 | 96.36 | 0 | 0 | 3.49 | 1.38 | 0.1405 | 0.182 | 0.0415 |
| 5a9a | opls | 1.82 | 0.0982 | 2.37 | 0 | 97.49 | 0.47 | 1 | 1.53 | 1 | 0.1445 | 0.1852 | 0.0407 |
| 5a9a | noschrodinger | 1.82 | 0.0043 | 0.9 | 0 | 97.08 | 0.93 | 0 | 2.82 | 1.23 | 0.1397 | 0.1834 | 0.0437 |
| 5a9s | opls | 2.06 | 0.0922 | 2.3 | 0.2 | 96.16 | 3.49 | 2 | 0.97 | 1.46 | 0.1936 | 0.2306 | 0.037 |
| 5a9s | noschrodinger | 2.06 | 0.0032 | 0.87 | 0.36 | 96.18 | 6.16 | 0 | 1.35 | 1.72 | 0.1792 | 0.2185 | 0.0393 |
| 5a9t | opls | 1.43 | 0.0943 | 2.4 | 0 | 96.88 | 0.47 | 0 | 1.4 | 1.06 | 0.1792 | 0.21 | 0.0308 |
| 5a9t | noschrodinger | 1.43 | 0.0034 | 0.91 | 0 | 97.54 | 0 | 0 | 0.47 | 0.76 | 0.1856 | 0.2089 | 0.0233 |
| 5a9w | opls | 3.7 | 0.0885 | 2.58 | 4.23 | 84.57 | 7.47 | 12 | 2.41 | 2.35 | 0.2976 | 0.3381 | 0.0405 |
| 5a9w | noschrodinger | 3.7 | 0.0038 | 0.92 | 6.87 | 75.77 | 8.82 | 0 | 13.37 | 3.14 | 0.2695 | 0.3134 | 0.0439 |
| 5ads | opls | 1.8 | 0.0986 | 2.22 | 0 | 98.03 | 0.56 | 0 | 1.46 | 0.88 | 0.1663 | 0.192 | 0.0257 |
| 5ads | noschrodinger | 1.8 | 0.0055 | 0.94 | 0 | 97.55 | 1.11 | 0 | 2.33 | 1.14 | 0.1693 | 0.1941 | 0.0248 |
| 5afe | opls | 2.6 | 0.0896 | 2.15 | 0 | 97.52 | 1.94 | 0 | 0.48 | 0.99 | 0.2663 | 0.2837 | 0.0174 |
| 5afe | noschrodinger | 2.6 | 0.0063 | 1.45 | 0 | 90.98 | 1.94 | 0 | 8.71 | 2.21 | 0.2731 | 0.2912 | 0.0181 |
| 5agj | opls | 2 | 0.0992 | 2.21 | 0 | 97.98 | 0 | 0 | 0.24 | 0.6 | 0.1746 | 0.2203 | 0.0457 |
| 5agj | noschrodinger | 2 | 0.0028 | 0.81 | 0 | 97.98 | 0.45 | 0 | 0.48 | 0.67 | 0.1735 | 0.2104 | 0.0369 |
| 5aip | opls | 2.3 | 0.0924 | 2.29 | 0 | 100 | 1.75 | 0 | 1.11 | 1 | 0.209 | 0.2338 | 0.0248 |
| 5aip | noschrodinger | 2.3 | 0.0032 | 0.73 | 0 | 100 | 1.69 | 0 | 2.44 | 1.2 | 0.1935 | 0.2313 | 0.0378 |
| 5ame | opls | 1.58 | 0.1023 | 2.34 | 0 | 99.51 | 0 | 3 | 1.69 | 0.92 | 0.1801 | 0.2132 | 0.0331 |
| 5ame | noschrodinger | 1.58 | 0.0046 | 0.85 | 0 | 99.59 | 0.92 | 0 | 0.73 | 0.73 | 0.1836 | 0.2087 | 0.0251 |
| 5ao9 | opls | 1.58 | 0.0966 | 2.48 | 0 | 97.07 | 4.65 | 4 | 7.74 | 2.09 | 0.1462 | 0.1715 | 0.0253 |
| 5ao9 | noschrodinger | 1.58 | 0.0119 | 1.24 | 0 | 96.74 | 5.38 | 0 | 8.45 | 2.22 | 0.147 | 0.1675 | 0.0205 |
| 5aol | opls | 1.5 | 0.1051 | 2.45 | 0 | 99.48 | 0.8 | 1 | 7.48 | 1.41 | 0.1497 | 0.174 | 0.0243 |
| 5aol | noschrodinger | 1.5 | 0.0087 | 1.15 | 0 | 99.74 | 1.06 | 0 | 2.65 | 1.07 | 0.1527 | 0.1723 | 0.0196 |
| 5apa | opls | 2.05 | 0.0977 | 2.23 | 0 | 98.88 | 2.87 | 0 | 1.24 | 1.19 | 0.1804 | 0.1928 | 0.0124 |
| 5apa | noschrodinger | 2.05 | 0.0026 | 0.78 | 0 | 98.45 | 3.45 | 0 | 1.56 | 1.31 | 0.1746 | 0.1964 | 0.0218 |
| 5aqh | opls | 2 | 0.0937 | 2.21 | 0 | 98.26 | 0.52 | 2 | 0.52 | 0.68 | 0.2338 | 0.2526 | 0.0188 |
| 5aqh | noschrodinger | 2 | 0.0044 | 0.78 | 0 | 98.15 | 0.26 | 0 | 0.27 | 0.6 | 0.2155 | 0.2458 | 0.0303 |
| 5aqv | opls | 1.75 | 0.0954 | 2.19 | 0 | 98.14 | 0.5 | 1 | 0.89 | 0.77 | 0.1979 | 0.2319 | 0.034 |
| 5aqv | noschrodinger | 1.75 | 0.0071 | 0.95 | 0 | 99.18 | 0.5 | 0 | 0.64 | 0.71 | 0.1949 | 0.2289 | 0.034 |
| 5aun | opls | 1.63 | 0.0999 | 2.36 | 0 | 97.95 | 1.61 | 2 | 3.27 | 1.29 | 0.1675 | 0.1968 | 0.0293 |
| 5aun | noschrodinger | 1.63 | 0.0049 | 0.99 | 0 | 97.57 | 1.6 | 0 | 2.8 | 1.31 | 0.1722 | 0.199 | 0.0268 |
| 5axl | opls | 3 | 0.091 | 2.3 | 0 | 98.37 | 2.11 | 0 | 0.95 | 1.03 | 0.2198 | 0.2477 | 0.0279 |
| 5axl | noschrodinger | 3 | 0.0027 | 0.77 | 0 | 98.16 | 6.02 | 0 | 1.76 | 1.53 | 0.228 | 0.2524 | 0.0244 |
| 5azb | opls | 1.59 | 0.0909 | 2.28 | 0 | 97.51 | 0 | 2 | 1.83 | 1.04 | 0.2065 | 0.2212 | 0.0147 |
| 5azb | noschrodinger | 1.59 | 0.0069 | 0.99 | 0.35 | 96.81 | 0.4 | 0 | 3.52 | 1.34 | 0.1971 | 0.2197 | 0.0226 |
| 5b00 | opls | 2.94 | 0.0938 | 2.48 | 0.52 | 93.93 | 5.88 | 2 | 1.33 | 1.85 | 0.2882 | 0.3122 | 0.024 |
| 5b00 | noschrodinger | 2.94 | 0.0038 | 0.88 | 3.6 | 85.33 | 14.8 | 0 | 11.22 | 3.11 | 0.266 | 0.3069 | 0.0409 |
| 5b1u | opls | 1.57 | 0.1057 | 2.29 | 0 | 97.27 | 0.96 | 1 | 3.02 | 1.23 | 0.1176 | 0.1489 | 0.0313 |
| 5b1u | noschrodinger | 1.57 | 0.0057 | 1 | 0 | 97.28 | 0.48 | 0 | 1.51 | 1.03 | 0.1197 | 0.1422 | 0.0225 |
| 5b4b | opls | 1.6 | 0.101 | 2.34 | 0 | 98.95 | 0 | 1 | 3.77 | 1.17 | 0.1584 | 0.1832 | 0.0248 |
| 5b4b | noschrodinger | 1.6 | 0.0072 | 1.08 | 0 | 98.74 | 0 | 0 | 2.43 | 1.03 | 0.1596 | 0.1865 | 0.0269 |
| 5b6v | opls | 1.77 | 0.0846 | 2.15 | 0.44 | 98.68 | 0.54 | 1 | 0.23 | 0.59 | 0.1836 | 0.2155 | 0.0319 |
| 5b6v | noschrodinger | 1.77 | 0.0052 | 0.9 | 0.44 | 98.69 | 0 | 0 | 0.69 | 0.72 | 0.1825 | 0.2097 | 0.0272 |
| 5b7i | opls | 2.6 | 0.0909 | 2.35 | 1.29 | 95.07 | 2.38 | 10 | 1.06 | 1.44 | 0.2135 | 0.2445 | 0.031 |
| 5b7i | noschrodinger | 2.6 | 0.0029 | 0.82 | 1.83 | 93.28 | 3.91 | 0 | 2.94 | 1.97 | 0.1885 | 0.2341 | 0.0456 |
| 5b7z | opls | 1.5 | 0.1033 | 2.57 | 0.75 | 98.49 | 1.32 | 4 | 4.4 | 1.31 | 0.1525 | 0.1775 | 0.025 |
| 5b7z | noschrodinger | 1.5 | 0.0106 | 1.17 | 0.75 | 98.31 | 1.09 | 2 | 4.4 | 1.25 | 0.1542 | 0.1788 | 0.0246 |
| 5b8h | opls | 2.2 | 0.095 | 2.41 | 0 | 99.14 | 0.57 | 0 | 1.31 | 0.86 | 0.1681 | 0.2269 | 0.0588 |

| | | | | | | | | | | | | | |
|---|---|---|---|---|---|---|---|---|---|---|---|---|---|
| 5b8h | noschrodinger | 2.2 | 0.0027 | 0.77 | 0 | 98.21 | 0.56 | 0 | 0.93 | 0.78 | 0.1888 | 0.2164 | 0.0276 |
| 5bni | opls | 2.5 | 0.094 | 2.32 | 0 | 98.25 | 5.31 | 6 | 0.68 | 1.27 | 0.2308 | 0.2824 | 0.0516 |
| 5bni | noschrodinger | 2.5 | 0.0029 | 0.77 | 0.22 | 96.51 | 11.51 | 0 | 6.5 | 2.39 | 0.2178 | 0.2727 | 0.0549 |
| 5bp9 | opls | 1.5 | 0.1011 | 2.24 | 0 | 98.88 | 0 | 1 | 4.78 | 1.25 | 0.1343 | 0.1595 | 0.0252 |
| 5bp9 | noschrodinger | 1.5 | 0.0058 | 1.09 | 0 | 98.7 | 1.47 | 0 | 4.3 | 1.34 | 0.1352 | 0.1542 | 0.019 |
| 5bpe | opls | 2.69 | 0.0906 | 2.3 | 0 | 95.38 | 5.41 | 2 | 1.39 | 1.75 | 0.2137 | 0.2458 | 0.0321 |
| 5bpe | noschrodinger | 2.69 | 0.0029 | 0.88 | 0.56 | 93.26 | 7.38 | 0 | 3.48 | 2.24 | 0.1947 | 0.2402 | 0.0455 |
| 5btq | opls | 2.08 | 0.0952 | 2.25 | 0 | 97.87 | 0 | 0 | 0.28 | 0.64 | 0.1977 | 0.2227 | 0.025 |
| 5btq | noschrodinger | 2.08 | 0.0028 | 0.81 | 0 | 98.35 | 0.54 | 0 | 0.84 | 0.76 | 0.1828 | 0.2209 | 0.0381 |
| 5btu | opls | 2.5 | 0.0886 | 2.8 | 0.36 | 96.01 | 8.4 | 14 | 2.66 | 2.03 | 0.1866 | 0.2344 | 0.0478 |
| 5btu | noschrodinger | 2.5 | 0.003 | 0.79 | 0.62 | 95.31 | 11.69 | 0 | 2.27 | 2.14 | 0.2053 | 0.2224 | 0.0171 |
| 5btx | opls | 2.1 | 0.0945 | 2.27 | 0.39 | 96.08 | 1.27 | 3 | 1.4 | 1.22 | 0.2658 | 0.296 | 0.0302 |
| 5btx | noschrodinger | 2.1 | 0.0032 | 0.88 | 1.54 | 93.82 | 3.75 | 0 | 4.64 | 2.08 | 0.2469 | 0.2844 | 0.0375 |
| 5buh | opls | 2.55 | 0.0924 | 2.24 | 0.65 | 96.1 | 2.14 | 1 | 0 | 1.02 | 0.2273 | 0.2609 | 0.0336 |
| 5buh | noschrodinger | 2.55 | 0.0026 | 0.72 | 0 | 97.52 | 6.38 | 0 | 0.38 | 1.35 | 0.2051 | 0.2445 | 0.0394 |
| 5bx1 | opls | 1.84 | 0.0977 | 2.47 | 1.35 | 93.24 | 4.55 | 5 | 3.14 | 2.04 | 0.198 | 0.2465 | 0.0485 |
| 5bx1 | noschrodinger | 1.84 | 0.0035 | 0.85 | 1.32 | 92.72 | 3.03 | 0 | 7.09 | 2.22 | 0.2016 | 0.2321 | 0.0305 |
| 5c1m | opls | 1.8 | 0.0917 | 2.29 | 0.25 | 96.95 | 2.77 | 1 | 0.43 | 1.17 | 0.2217 | 0.2427 | 0.021 |
| 5c1m | noschrodinger | 1.8 | 0.0096 | 1.16 | 0 | 95.31 | 3.57 | 0 | 3.92 | 1.93 | 0.2094 | 0.2438 | 0.0344 |
| 5c2j | opls | 2.49 | 0.0939 | 2.31 | 0 | 95.62 | 2.29 | 2 | 0.64 | 1.29 | 0.1946 | 0.2386 | 0.044 |
| 5c2j | noschrodinger | 2.49 | 0.004 | 0.88 | 0 | 95.1 | 3.43 | 0 | 3.83 | 1.92 | 0.1746 | 0.2277 | 0.0531 |
| 5c2o | opls | 2.35 | 0.0944 | 2.29 | 0 | 97.16 | 2.89 | 1 | 0.88 | 1.27 | 0.1914 | 0.2215 | 0.0301 |
| 5c2o | noschrodinger | 2.35 | 0.0026 | 0.75 | 0 | 97.89 | 2.48 | 0 | 0.22 | 0.91 | 0.1903 | 0.2177 | 0.0274 |
| 5c3c | opls | 2.8 | 0.09 | 2.39 | 0 | 96.87 | 3.35 | 3 | 2.02 | 1.56 | 0.2509 | 0.2793 | 0.0284 |
| 5c3c | noschrodinger | 2.8 | 0.0029 | 0.8 | 0.22 | 93.81 | 4.83 | 0 | 3.37 | 2.06 | 0.2424 | 0.2816 | 0.0392 |
| 5c42 | opls | 3.23 | 0.0902 | 2.22 | 1.18 | 94.33 | 2.33 | 1 | 1.26 | 1.51 | 0.2629 | 0.2997 | 0.0368 |
| 5c42 | noschrodinger | 3.23 | 0.0028 | 0.76 | 0.73 | 93.04 | 3.6 | 0 | 2.65 | 1.92 | 0.2478 | 0.2839 | 0.0361 |
| 5c4n | opls | 1.63 | 0.1023 | 2.65 | 0 | 97.89 | 3.45 | 5 | 5.73 | 1.75 | 0.1723 | 0.2008 | 0.0285 |
| 5c4n | noschrodinger | 1.63 | 0.0092 | 1.38 | 0 | 97.08 | 2.29 | 0 | 6.83 | 1.81 | 0.1735 | 0.1985 | 0.025 |
| 5c50 | opls | 1.63 | 0.0966 | 2.28 | 0 | 98.4 | 0.58 | 2 | 1.43 | 0.88 | 0.1824 | 0.2115 | 0.0291 |
| 5c50 | noschrodinger | 1.63 | 0.0093 | 1.07 | 0.26 | 97.63 | 1.16 | 0 | 0.8 | 0.88 | 0.1817 | 0.2079 | 0.0262 |
| 5c79 | opls | 1.6 | 0.1006 | 2.38 | 0 | 99 | 2.76 | 1 | 1.97 | 1.3 | 0.1745 | 0.2127 | 0.0382 |
| 5c79 | noschrodinger | 1.6 | 0.0074 | 0.99 | 0 | 97.54 | 2.75 | 0 | 1.41 | 1.3 | 0.1789 | 0.2112 | 0.0323 |
| 5c7q | opls | 1.52 | 0.0992 | 2.41 | 0 | 93.8 | 1.56 | 4 | 3.49 | 1.7 | 0.1662 | 0.2048 | 0.0386 |
| 5c7q | noschrodinger | 1.52 | 0.0055 | 0.91 | 0 | 95.25 | 1.85 | 0 | 2.38 | 1.55 | 0.1722 | 0.1996 | 0.0274 |
| 5c7t | opls | 2.06 | 0.0955 | 2.26 | 0.56 | 94.35 | 1.26 | 1 | 1.43 | 1.34 | 0.2026 | 0.2316 | 0.029 |
| 5c7t | noschrodinger | 2.06 | 0.0029 | 0.79 | 0.56 | 94.68 | 1.25 | 0 | 3.35 | 1.57 | 0.1863 | 0.2214 | 0.0351 |
| 5c8c | opls | 2.5 | 0.0981 | 2.28 | 0.39 | 95.83 | 0.6 | 3 | 0.58 | 0.98 | 0.4938 | 0.5254 | 0.0316 |
| 5c8c | noschrodinger | 2.5 | 0.0027 | 0.8 | 0.26 | 95.08 | 1.35 | 0 | 2.72 | 1.5 | 0.4882 | 0.5165 | 0.0283 |
| 5c8z | opls | 1.6 | 0.0966 | 2.57 | 0.95 | 97.34 | 0.63 | 2 | 2.52 | 1.16 | 0.1414 | 0.1565 | 0.0151 |
| 5c8z | noschrodinger | 1.6 | 0.0056 | 1.03 | 0.76 | 97.35 | 0.42 | 0 | 1.48 | 1.01 | 0.1476 | 0.1572 | 0.0096 |
| 5c93 | opls | 2.52 | 0.0904 | 2.48 | 0.45 | 95.69 | 1.97 | 6 | 2.01 | 1.49 | 0.242 | 0.282 | 0.04 |
| 5c93 | noschrodinger | 2.52 | 0.0041 | 0.76 | 0.22 | 95.96 | 2.2 | 0 | 2.69 | 1.59 | 0.2355 | 0.266 | 0.0305 |
| 5c94 | opls | 2.44 | 0.0942 | 2.35 | 0 | 98.08 | 5.26 | 1 | 2.79 | 1.62 | 0.2008 | 0.2319 | 0.0311 |
| 5c94 | noschrodinger | 2.44 | 0.0032 | 0.87 | 0.88 | 94.74 | 4.17 | 0 | 5.05 | 2.1 | 0.2061 | 0.2274 | 0.0213 |
| 5caj | opls | 1.65 | 0.101 | 2.3 | 0 | 98.75 | 0.22 | 1 | 2.67 | 1.05 | 0.1575 | 0.1895 | 0.032 |
| 5caj | noschrodinger | 1.65 | 0.0065 | 0.96 | 0 | 98.84 | 0.22 | 0 | 2.09 | 0.98 | 0.1616 | 0.1887 | 0.0271 |
| 5cbn | opls | 2.3 | 0.0927 | 2.23 | 0 | 97.93 | 3.77 | 1 | 0.48 | 1.12 | 0.227 | 0.261 | 0.034 |
| 5cbn | noschrodinger | 2.3 | 0.0028 | 0.78 | 0.56 | 95.69 | 6.32 | 0 | 2.17 | 1.9 | 0.2111 | 0.2539 | 0.0428 |

| | | | | | | | | | | | | | |
|---|---|---|---|---|---|---|---|---|---|---|---|---|---|
| 5ced | opls | 2.02 | 0.0947 | 2.2 | 0 | 98.06 | 0.63 | 2 | 0.67 | 0.72 | 0.2129 | 0.2308 | 0.0179 |
| 5ced | noschrodinger | 2.02 | 0.003 | 0.81 | 0 | 97.36 | 1.26 | 0 | 1.35 | 1.06 | 0.1908 | 0.2223 | 0.0315 |
| 5cf9 | opls | 1.52 | 0.1002 | 2.39 | 0.41 | 97.52 | 0.96 | 1 | 1.77 | 1.03 | 0.1573 | 0.1931 | 0.0358 |
| 5cf9 | noschrodinger | 1.52 | 0.0091 | 1.05 | 0.41 | 96.72 | 1.43 | 0 | 0.51 | 1 | 0.1619 | 0.1941 | 0.0322 |
| 5cfp | opls | 2.07 | 0.098 | 2.32 | 0.84 | 95.22 | 1.53 | 1 | 2.14 | 1.46 | 0.1827 | 0.2054 | 0.0227 |
| 5cfp | noschrodinger | 2.07 | 0.0081 | 1.1 | 0.56 | 96.09 | 1.83 | 0 | 4.11 | 1.66 | 0.1827 | 0.2047 | 0.022 |
| 5cft | opls | 1.5 | 0.1007 | 2.46 | 0 | 98.8 | 0 | 1 | 4.53 | 1.23 | 0.1516 | 0.1839 | 0.0323 |
| 5cft | noschrodinger | 1.5 | 0.0083 | 1.37 | 0 | 98.84 | 0.66 | 1 | 8.4 | 1.45 | 0.1556 | 0.1816 | 0.026 |
| 5cjj | opls | 2.42 | 0.0887 | 2.22 | 0 | 96.85 | 1.03 | 1 | 0.68 | 0.92 | 0.302 | 0.3345 | 0.0325 |
| 5cjj | noschrodinger | 2.42 | 0.0026 | 0.74 | 0 | 95.69 | 1.32 | 0 | 1.2 | 1.23 | 0.2819 | 0.3296 | 0.0477 |
| 5cqe | opls | 2.1 | 0.0941 | 2.25 | 0 | 97.74 | 0.73 | 0 | 0.58 | 0.75 | 0.1973 | 0.2155 | 0.0182 |
| 5cqe | noschrodinger | 2.1 | 0.0227 | 1.03 | 0.32 | 97.46 | 0.72 | 0 | 2.12 | 1.09 | 0.1785 | 0.205 | 0.0265 |
| 5ctg | opls | 3.1 | 0.0895 | 2.34 | 0 | 98.94 | 5.34 | 3 | 1.08 | 1.36 | 0.2867 | 0.3214 | 0.0347 |
| 5ctg | noschrodinger | 3.1 | 0.0029 | 0.9 | 0.31 | 97.36 | 12.03 | 0 | 6.23 | 2.29 | 0.3026 | 0.3249 | 0.0223 |
| 5cth | opls | 3.69 | 0.0891 | 2.39 | 0.35 | 96.29 | 4.61 | 5 | 0.68 | 1.48 | 0.3037 | 0.3475 | 0.0438 |
| 5cth | noschrodinger | 3.69 | 0.0028 | 0.76 | 0.32 | 96.21 | 7.2 | 0 | 2.47 | 1.94 | 0.2878 | 0.3369 | 0.0491 |
| 5cuo | opls | 1.54 | 0.1052 | 2.46 | 0 | 99.46 | 0.66 | 2 | 2.72 | 1.06 | 0.1888 | 0.2195 | 0.0307 |
| 5cuo | noschrodinger | 1.54 | 0.007 | 0.98 | 0 | 98.93 | 0.65 | 0 | 1.02 | 0.8 | 0.1925 | 0.2151 | 0.0226 |
| 5cvq | opls | 2.5 | 0.096 | 2.3 | 0 | 98.68 | 2.4 | 1 | 1.59 | 1.19 | 0.209 | 0.2317 | 0.0227 |
| 5cvq | noschrodinger | 2.5 | 0.0033 | 0.88 | 0 | 98.7 | 3.15 | 0 | 3.58 | 1.53 | 0.1954 | 0.2295 | 0.0341 |
| 5cw8 | opls | 2.6 | 0.0886 | 2.26 | 0 | 98.48 | 2.04 | 1 | 0.91 | 1.01 | 0.2054 | 0.2415 | 0.0361 |
| 5cw8 | noschrodinger | 2.6 | 0.0036 | 0.82 | 0.29 | 97.99 | 3.66 | 0 | 7.26 | 1.83 | 0.1959 | 0.2409 | 0.045 |
| 5cxi | opls | 2 | 0.0941 | 2.23 | 0 | 98.42 | 2.45 | 1 | 0.73 | 1.03 | 0.2579 | 0.2725 | 0.0146 |
| 5cxi | noschrodinger | 2 | 0.0035 | 0.78 | 0.29 | 98.57 | 3.6 | 0 | 2.61 | 1.47 | 0.2307 | 0.2606 | 0.0299 |
| 5cxr | opls | 2 | 0.0983 | 2.26 | 0.51 | 97.97 | 0.55 | 1 | 0.3 | 0.62 | 0.1912 | 0.217 | 0.0258 |
| 5cxr | noschrodinger | 2 | 0.0029 | 0.73 | 0 | 95.48 | 0 | 0 | 1.79 | 1.25 | 0.1826 | 0.2123 | 0.0297 |
| 5cxx | opls | 1.55 | 0.1013 | 2.49 | 0 | 98.64 | 0.41 | 12 | 3.58 | 1.15 | 0.1564 | 0.1735 | 0.0171 |
| 5cxx | noschrodinger | 1.55 | 0.014 | 1.32 | 0 | 98.65 | 0.54 | 0 | 2.31 | 1.01 | 0.1578 | 0.1748 | 0.017 |
| 5cyo | opls | 2.04 | 0.0941 | 2.26 | 0 | 98.59 | 1.32 | 0 | 1.41 | 0.97 | 0.186 | 0.2182 | 0.0322 |
| 5cyo | noschrodinger | 2.04 | 0.0036 | 0.82 | 0.2 | 97.84 | 1.3 | 0 | 2.39 | 1.14 | 0.1862 | 0.2143 | 0.0281 |
| 5cyr | opls | 3.5 | 0.0919 | 2.53 | 1.16 | 92.59 | 3.94 | 15 | 2.39 | 1.94 | 0.247 | 0.2966 | 0.0496 |
| 5cyr | noschrodinger | 3.5 | 0.003 | 0.77 | 1.13 | 89.41 | 3.22 | 0 | 5.53 | 2.25 | 0.2432 | 0.2836 | 0.0404 |
| 5d1m | opls | 1.58 | 0.1047 | 2.45 | 0.4 | 97.98 | 2.39 | 3 | 5.79 | 1.61 | 0.1536 | 0.1833 | 0.0297 |
| 5d1m | noschrodinger | 1.58 | 0.0092 | 1.13 | 0 | 98 | 1.98 | 0 | 5.3 | 1.51 | 0.1527 | 0.178 | 0.0253 |
| 5d1r | opls | 2 | 0.0972 | 2.32 | 0 | 98.21 | 3.93 | 3 | 0.29 | 1.06 | 0.2109 | 0.2299 | 0.019 |
| 5d1r | noschrodinger | 2 | 0.0029 | 0.78 | 0 | 98.22 | 6.7 | 0 | 3.43 | 1.76 | 0.1897 | 0.2178 | 0.0281 |
| 5d2k | opls | 1.57 | 0.1037 | 2.54 | 0 | 96.93 | 2.73 | 7 | 3.57 | 1.66 | 0.1327 | 0.1698 | 0.0371 |
| 5d2k | noschrodinger | 1.57 | 0.0046 | 0.95 | 0 | 97.32 | 3.18 | 0 | 2.73 | 1.57 | 0.1394 | 0.169 | 0.0296 |
| 5d4n | opls | 1.6 | 0.0924 | 2.34 | 0 | 99.2 | 0.45 | 4 | 3.24 | 1.11 | 0.1917 | 0.2304 | 0.0387 |
| 5d4n | noschrodinger | 1.6 | 0.0054 | 1.01 | 0 | 99.65 | 0.88 | 0 | 2.18 | 0.99 | 0.1966 | 0.2263 | 0.0297 |
| 5d4w | opls | 3.6 | 0.0897 | 2.54 | 2.88 | 84.77 | 11.49 | 31 | 3.09 | 2.57 | 0.298 | 0.3421 | 0.0441 |
| 5d4w | noschrodinger | 3.6 | 0.0033 | 0.89 | 1.87 | 82.79 | 14.89 | 0 | 13.4 | 3.22 | 0.2807 | 0.323 | 0.0423 |
| 5d61 | opls | 1.6 | 0.0983 | 2.51 | 0 | 97.25 | 0.81 | 0 | 1.86 | 1.09 | 0.1511 | 0.1671 | 0.016 |
| 5d61 | noschrodinger | 1.6 | 0.0076 | 1.15 | 0 | 97.59 | 1.21 | 0 | 5.79 | 1.47 | 0.1583 | 0.1693 | 0.011 |
| 5d6q | opls | 1.5 | 0.1003 | 2.29 | 0 | 97.83 | 1.71 | 1 | 2.16 | 1.21 | 0.1617 | 0.1936 | 0.0319 |
| 5d6q | noschrodinger | 1.5 | 0.0043 | 0.94 | 0 | 97.58 | 1.7 | 0 | 2.62 | 1.31 | 0.1642 | 0.1894 | 0.0252 |
| 5d6t | opls | 1.93 | 0.0993 | 2.21 | 0 | 97.55 | 0.94 | 1 | 0.51 | 0.77 | 0.1629 | 0.1938 | 0.0309 |
| 5d6t | noschrodinger | 1.93 | 0.0031 | 0.82 | 0 | 98.41 | 0.47 | 0 | 0.77 | 0.74 | 0.1642 | 0.1915 | 0.0273 |
| 5d71 | opls | 2.25 | 0.0947 | 2.15 | 0 | 99.01 | 1.37 | 1 | 0.39 | 0.74 | 0.2038 | 0.2192 | 0.0154 |

| | | | | | | | | | | | | | |
|---|---|---|---|---|---|---|---|---|---|---|---|---|---|
| 5d71 | noschrodinger | 2.25 | 0.0027 | 0.79 | 0 | 98.25 | 1.57 | 0 | 0.53 | 0.83 | 0.1882 | 0.2133 | 0.0251 |
| 5d76 | opls | 2.54 | 0.0907 | 2.3 | 0.9 | 93.71 | 0.56 | 3 | 1.3 | 1.27 | 0.2228 | 0.2566 | 0.0338 |
| 5d76 | noschrodinger | 2.54 | 0.0033 | 0.84 | 1.1 | 92.49 | 1.67 | 0 | 3.9 | 1.81 | 0.2121 | 0.2493 | 0.0372 |
| 5dad | opls | 2.61 | 0.0899 | 2.29 | 0 | 97.41 | 0 | 0 | 0.51 | 0.79 | 0.2114 | 0.2395 | 0.0281 |
| 5dad | noschrodinger | 2.61 | 0.0024 | 0.71 | 0 | 94.96 | 0.93 | 0 | 0 | 0.85 | 0.206 | 0.2476 | 0.0416 |
| 5dau | opls | 1.5 | 0.1015 | 2.15 | 0 | 96.03 | 1.61 | 2 | 0.83 | 1.19 | 0.1687 | 0.1973 | 0.0286 |
| 5dau | noschrodinger | 1.5 | 0.0042 | 0.91 | 0 | 95.28 | 3.2 | 0 | 1.66 | 1.63 | 0.1698 | 0.1972 | 0.0274 |
| 5deq | opls | 1.95 | 0.0957 | 2.2 | 0 | 98.94 | 1.39 | 1 | 1.24 | 0.95 | 0.2037 | 0.2387 | 0.035 |
| 5deq | noschrodinger | 1.95 | 0.0034 | 0.77 | 0.23 | 97.91 | 1.32 | 0 | 1.11 | 0.93 | 0.2031 | 0.2346 | 0.0315 |
| 5dfy | opls | 1.6 | 0.1041 | 2.31 | 0 | 99.35 | 0.72 | 0 | 1.5 | 0.89 | 0.1563 | 0.1718 | 0.0155 |
| 5dfy | noschrodinger | 1.6 | 0.0075 | 1.08 | 0 | 99.36 | 0.72 | 0 | 2.25 | 1 | 0.1584 | 0.1733 | 0.0149 |
| 5di3 | opls | 2.5 | 0.0917 | 2.29 | 0 | 98.07 | 0.32 | 3 | 0.68 | 0.72 | 0.2245 | 0.2522 | 0.0277 |
| 5di3 | noschrodinger | 2.5 | 0.0031 | 1.01 | 0 | 96.7 | 2.88 | 0 | 1.86 | 1.51 | 0.2135 | 0.2409 | 0.0274 |
| 5dje | opls | 1.85 | 0.1044 | 2.26 | 0 | 99.6 | 0.84 | 3 | 2.13 | 0.99 | 0.1819 | 0.2219 | 0.04 |
| 5dje | noschrodinger | 1.85 | 0.0052 | 0.81 | 0 | 99.63 | 0.8 | 0 | 0.43 | 0.65 | 0.1863 | 0.2179 | 0.0316 |
| 5djo | opls | 1.74 | 0.0992 | 2.49 | 0 | 98.65 | 1.39 | 6 | 4.24 | 1.31 | 0.1868 | 0.223 | 0.0362 |
| 5djo | noschrodinger | 1.74 | 0.0082 | 0.99 | 0 | 98.09 | 2.08 | 0 | 4.44 | 1.46 | 0.1881 | 0.222 | 0.0339 |
| 5dkk | opls | 2.5 | 0.0923 | 2.24 | 0 | 99.62 | 0.88 | 0 | 1.87 | 0.95 | 0.2118 | 0.2477 | 0.0359 |
| 5dkk | noschrodinger | 2.5 | 0.0028 | 0.78 | 0 | 98.87 | 0.88 | 0 | 4.93 | 1.26 | 0.1998 | 0.2322 | 0.0324 |
| 5dni | opls | 2.23 | 0.0925 | 2.21 | 0.68 | 94.2 | 0.36 | 1 | 0.36 | 1.02 | 0.2078 | 0.234 | 0.0262 |
| 5dni | noschrodinger | 2.23 | 0.0028 | 0.77 | 0.85 | 93.79 | 0.35 | 0 | 1.26 | 1.26 | 0.1919 | 0.2269 | 0.035 |
| 5dnl | opls | 1.53 | 0.1006 | 2.19 | 0 | 96.72 | 0.45 | 0 | 2.45 | 1.23 | 0.1333 | 0.1432 | 0.0099 |
| 5dnl | noschrodinger | 1.53 | 0.0067 | 1.03 | 0 | 97.13 | 0.22 | 0 | 1.52 | 1.05 | 0.1337 | 0.141 | 0.0073 |
| 5dro | opls | 2.01 | 0.0989 | 2.19 | 0 | 97.16 | 0.21 | 1 | 0.56 | 0.84 | 0.1647 | 0.1857 | 0.021 |
| 5dro | noschrodinger | 2.01 | 0.0027 | 0.77 | 0 | 96.04 | 0.63 | 0 | 2.13 | 1.26 | 0.1479 | 0.1814 | 0.0335 |
| 5ds7 | opls | 2 | 0.0914 | 2.22 | 0 | 99.6 | 0.91 | 3 | 0 | 0.5 | 0.2133 | 0.2448 | 0.0315 |
| 5ds7 | noschrodinger | 2 | 0.0057 | 0.95 | 0 | 99.29 | 0.9 | 0 | 1.55 | 0.9 | 0.2113 | 0.2439 | 0.0326 |
| 5.00E+16 | opls | 1.65 | 0.0981 | 2.32 | 0 | 100 | 1.61 | 2 | 3.96 | 1.34 | 0.1885 | 0.2201 | 0.0316 |
| 5.00E+16 | noschrodinger | 1.65 | 0.0115 | 1.42 | 0 | 100 | 0.8 | 0 | 4.42 | 1.22 | 0.1894 | 0.218 | 0.0286 |
| 5e6v | opls | 1.8 | 0.1009 | 2.54 | 0 | 97.18 | 0.52 | 2 | 2.85 | 1.22 | 0.1748 | 0.21 | 0.0352 |
| 5e6v | noschrodinger | 1.8 | 0.0086 | 1.1 | 0.45 | 95.93 | 0 | 0 | 5.14 | 1.55 | 0.1803 | 0.2058 | 0.0255 |
| 5eb3 | opls | 2.4 | 0.0988 | 2.28 | 0 | 97.63 | 0 | 1 | 0.64 | 0.79 | 0.1809 | 0.2124 | 0.0315 |
| 5eb3 | noschrodinger | 2.4 | 0.0041 | 0.84 | 0 | 97.98 | 1.97 | 0 | 2.34 | 1.24 | 0.1774 | 0.2143 | 0.0369 |
| 5ebk | opls | 3.5 | 0.0894 | 2.3 | 0.21 | 95.86 | 2.92 | 2 | 1 | 1.43 | 0.2366 | 0.2629 | 0.0263 |
| 5ebk | noschrodinger | 3.5 | 0.0029 | 0.8 | 0 | 90.82 | 5.31 | 0 | 3.32 | 2.2 | 0.208 | 0.2496 | 0.0416 |
| 5edl | opls | 1.95 | 0.0943 | 2.32 | 0 | 99.48 | 1.32 | 0 | 1.57 | 0.99 | 0.186 | 0.2175 | 0.0315 |
| 5edl | noschrodinger | 1.95 | 0.0096 | 0.97 | 0 | 98.97 | 0.65 | 0 | 0.63 | 0.71 | 0.1897 | 0.211 | 0.0213 |
| 5ee5 | opls | 2.28 | 0.0914 | 2.33 | 0 | 98.59 | 0.64 | 1 | 0.67 | 0.72 | 0.2496 | 0.2659 | 0.0163 |
| 5ee5 | noschrodinger | 2.28 | 0.0029 | 0.77 | 0 | 96.64 | 0.95 | 0 | 2.35 | 1.23 | 0.2299 | 0.26 | 0.0301 |
| 5eg3 | opls | 2.61 | 0.0917 | 2.28 | 0.26 | 97.69 | 1.98 | 1 | 0.9 | 1.07 | 0.1993 | 0.2417 | 0.0424 |
| 5eg3 | noschrodinger | 2.61 | 0.0028 | 0.78 | 0.74 | 97.04 | 1.97 | 0 | 1.5 | 1.28 | 0.1825 | 0.222 | 0.0395 |
| 5egl | opls | 2.1 | 0.0946 | 2.34 | 0 | 98.75 | 0 | 1 | 1.24 | 0.84 | 0.2107 | 0.222 | 0.0113 |
| 5egl | noschrodinger | 2.1 | 0.0054 | 1.02 | 0 | 97.31 | 0 | 0 | 2.48 | 1.16 | 0.2056 | 0.2174 | 0.0118 |
| 5ei3 | opls | 1.71 | 0.1068 | 2.4 | 0 | 97.49 | 0.55 | 1 | 0.87 | 0.87 | 0.1475 | 0.1801 | 0.0326 |
| 5ei3 | noschrodinger | 1.71 | 0.0073 | 0.97 | 0 | 98.01 | 1.09 | 0 | 0.87 | 0.79 | 0.1492 | 0.1773 | 0.0281 |
| 5eik | opls | 2.3 | 0.0902 | 2.55 | 0 | 96.54 | 1.98 | 1 | 1.39 | 1.32 | 0.1998 | 0.2255 | 0.0257 |
| 5eik | noschrodinger | 2.3 | 0.0028 | 0.81 | 0.43 | 96.98 | 2.46 | 0 | 2.56 | 1.51 | 0.2034 | 0.2251 | 0.0217 |
| 5enc | opls | 1.59 | 0.1042 | 2.47 | 0 | 100 | 0 | 3 | 3.88 | 1.18 | 0.2011 | 0.2436 | 0.0425 |
| 5enc | noschrodinger | 1.59 | 0.0078 | 0.97 | 0 | 99.18 | 0.88 | 0 | 2.44 | 1.03 | 0.205 | 0.2425 | 0.0375 |

| | | | | | | | | | | | | | |
|---|---|---|---|---|---|---|---|---|---|---|---|---|---|
| 5ep4 | opls | 1.5 | 0.1005 | 2.52 | 0 | 98.38 | 0.45 | 1 | 2.91 | 1.08 | 0.1448 | 0.1782 | 0.0334 |
| 5ep4 | noschrodinger | 1.5 | 0.0136 | 1.54 | 0 | 98.41 | 1.33 | 0 | 4.84 | 1.35 | 0.1457 | 0.1805 | 0.0348 |
| 5epn | opls | 1.89 | 0.098 | 2.37 | 0 | 98.93 | 0.64 | 0 | 1.32 | 0.86 | 0.1572 | 0.2047 | 0.0475 |
| 5epn | noschrodinger | 1.89 | 0.0037 | 1.05 | 0 | 98.97 | 0.63 | 0 | 1.33 | 0.86 | 0.1664 | 0.2066 | 0.0402 |
| 5eq3 | opls | 2 | 0.1 | 2.22 | 0 | 96.89 | 0.86 | 2 | 0.16 | 0.75 | 0.2015 | 0.2454 | 0.0439 |
| 5eq3 | noschrodinger | 2 | 0.0028 | 0.8 | 0 | 97.16 | 1.14 | 0 | 2.08 | 1.17 | 0.2019 | 0.2345 | 0.0326 |
| 5es9 | opls | 3.46 | 0.0903 | 2.29 | 0.57 | 96.03 | 1.63 | 5 | 0.87 | 1.2 | 0.2962 | 0.3337 | 0.0375 |
| 5es9 | noschrodinger | 3.46 | 0.0028 | 0.76 | 0.56 | 94.69 | 2.02 | 0 | 4.15 | 1.8 | 0.2979 | 0.3292 | 0.0313 |
| 5eux | opls | 2.04 | 0.093 | 2.21 | 0 | 100 | 0 | 0 | 0.48 | 0.67 | 0.2157 | 0.2535 | 0.0378 |
| 5eux | noschrodinger | 2.04 | 0.0035 | 0.81 | 0 | 99.2 | 0 | 0 | 0.48 | 0.67 | 0.1995 | 0.25 | 0.0505 |
| 5exs | opls | 2.5 | 0.0979 | 2.34 | 0 | 97.59 | 1.88 | 0 | 0.74 | 1.03 | 0.2199 | 0.2698 | 0.0499 |
| 5exs | noschrodinger | 2.5 | 0.0027 | 0.76 | 0 | 98 | 8.41 | 0 | 5.16 | 1.98 | 0.1926 | 0.2394 | 0.0468 |
| 5ext | opls | 2.39 | 0.095 | 2.33 | 0 | 96.79 | 5.63 | 1 | 0.74 | 1.5 | 0.2072 | 0.247 | 0.0398 |
| 5ext | noschrodinger | 2.39 | 0.0026 | 0.8 | 0.8 | 96 | 8.41 | 0 | 1.72 | 1.9 | 0.1915 | 0.2376 | 0.0461 |
| 5ey0 | opls | 1.6 | 0.1006 | 2.53 | 0.41 | 97.36 | 1.8 | 5 | 3.76 | 1.48 | 0.1761 | 0.2144 | 0.0383 |
| 5ey0 | noschrodinger | 1.6 | 0.0136 | 1.26 | 0.2 | 96.38 | 2.23 | 0 | 3.77 | 1.67 | 0.1807 | 0.21 | 0.0293 |
| 5ey1 | opls | 1.97 | 0.0996 | 2.37 | 0 | 97.46 | 2.18 | 0 | 2.23 | 1.37 | 0.1955 | 0.2326 | 0.0371 |
| 5ey1 | noschrodinger | 1.97 | 0.0031 | 0.81 | 0 | 97.49 | 2.16 | 0 | 3.73 | 1.52 | 0.2003 | 0.2241 | 0.0238 |
| 5eyi | opls | 2.16 | 0.1003 | 2.23 | 0 | 98.15 | 3.66 | 2 | 1.48 | 1.31 | 0.1748 | 0.2185 | 0.0437 |
| 5eyi | noschrodinger | 2.16 | 0.0028 | 0.83 | 0 | 97.65 | 4.16 | 0 | 1.19 | 1.38 | 0.1838 | 0.2206 | 0.0368 |
| 5ez1 | opls | 2.4 | 0.0942 | 2.2 | 0.22 | 99.11 | 2.24 | 0 | 1.05 | 1.07 | 0.2896 | 0.3053 | 0.0157 |
| 5ez1 | noschrodinger | 2.4 | 0.0027 | 0.73 | 0.22 | 98.24 | 2.22 | 0 | 4.98 | 1.53 | 0.2773 | 0.2986 | 0.0213 |
| 5f12 | opls | 1.5 | 0.0962 | 2.24 | 0.83 | 98.62 | 1.72 | 2 | 1.39 | 1.05 | 0.1769 | 0.2033 | 0.0264 |
| 5f12 | noschrodinger | 1.5 | 0.0072 | 1.12 | 0.54 | 98.91 | 2.05 | 0 | 1.91 | 1.19 | 0.1797 | 0.2018 | 0.0221 |
| 5f47 | opls | 1.5 | 0.111 | 2.25 | 0 | 99.33 | 1.42 | 2 | 1.36 | 0.98 | 0.1496 | 0.1832 | 0.0336 |
| 5f47 | noschrodinger | 1.5 | 0.0068 | 0.97 | 0 | 99.33 | 1.07 | 0 | 2.71 | 1.08 | 0.1491 | 0.174 | 0.0249 |
| 5f5r | opls | 1.85 | 0.1033 | 2.51 | 0.68 | 96.8 | 0.53 | 6 | 3.01 | 1.29 | 0.1844 | 0.2268 | 0.0424 |
| 5f5r | noschrodinger | 1.85 | 0.0115 | 1.1 | 0 | 97.28 | 1.32 | 0 | 2.58 | 1.27 | 0.1921 | 0.2279 | 0.0358 |
| 5f8w | opls | 1.56 | 0.1063 | 2.25 | 0 | 96.92 | 0 | 0 | 1.43 | 1.06 | 0.1217 | 0.1452 | 0.0235 |
| 5f8w | noschrodinger | 1.56 | 0.0064 | 1.15 | 0 | 96.94 | 0 | 0 | 1.44 | 1.06 | 0.1224 | 0.1422 | 0.0198 |
| 5fae | opls | 1.7 | 0.0976 | 2.83 | 0 | 97.96 | 2.06 | 3 | 2.22 | 1.25 | 0.1921 | 0.2407 | 0.0486 |
| 5fae | noschrodinger | 1.7 | 0.0035 | 0.76 | 0 | 95.19 | 1.01 | 0 | 2.22 | 1.34 | 0.2032 | 0.2331 | 0.0299 |
| 5fb9 | opls | 1.5 | 0.1059 | 2.32 | 0 | 97.72 | 2.16 | 2 | 4.08 | 1.51 | 0.1421 | 0.1717 | 0.0296 |
| 5fb9 | noschrodinger | 1.5 | 0.0062 | 1 | 0 | 98.3 | 1.72 | 0 | 1.56 | 1.08 | 0.1421 | 0.1662 | 0.0241 |
| 5fc4 | opls | 1.5 | 0.1044 | 2.49 | 0 | 100 | 1.65 | 0 | 3.31 | 1.29 | 0.151 | 0.1886 | 0.0376 |
| 5fc4 | noschrodinger | 1.5 | 0.0048 | 0.89 | 0 | 99.26 | 0.81 | 0 | 2.9 | 1.08 | 0.1541 | 0.1844 | 0.0303 |
| 5fcy | opls | 2.51 | 0.0906 | 2.26 | 0 | 99.22 | 0 | 1 | 0.88 | 0.77 | 0.2152 | 0.2573 | 0.0421 |
| 5fcy | noschrodinger | 2.51 | 0.0029 | 0.73 | 0 | 99.23 | 0.83 | 0 | 3.51 | 1.14 | 0.2166 | 0.2454 | 0.0288 |
| 5fdd | opls | 2.51 | 0.092 | 2.22 | 0 | 97.74 | 0 | 0 | 0.33 | 0.68 | 0.2223 | 0.2438 | 0.0215 |
| 5fdd | noschrodinger | 2.51 | 0.0026 | 0.7 | 0 | 98.32 | 0.62 | 0 | 1.67 | 0.92 | 0.2362 | 0.2562 | 0.02 |
| 5fea | opls | 2.6 | 0.0872 | 2.22 | 0 | 98.35 | 0.52 | 0 | 1.01 | 0.8 | 0.2395 | 0.2813 | 0.0418 |
| 5fea | noschrodinger | 2.6 | 0.0026 | 0.82 | 0 | 97.93 | 0 | 0 | 0.26 | 0.61 | 0.2308 | 0.2862 | 0.0554 |
| 5ffz | opls | 2.58 | 0.0832 | 2.2 | 0 | 100 | 6.52 | 0 | 0 | 1.12 | 0.2884 | 0.3107 | 0.0223 |
| 5ffz | noschrodinger | 2.58 | 0.0031 | 0.71 | 0 | 97.1 | 6.45 | 0 | 0.51 | 1.45 | 0.2621 | 0.306 | 0.0439 |
| 5flg | opls | 2.04 | 0.0916 | 2.24 | 0 | 98.79 | 0.91 | 0 | 1.1 | 0.82 | 0.2062 | 0.2298 | 0.0236 |
| 5flg | noschrodinger | 2.04 | 0.0032 | 0.83 | 0 | 99.6 | 0.91 | 0 | 2.09 | 0.98 | 0.1937 | 0.2188 | 0.0251 |
| 5fob | opls | 2.6 | 0.0912 | 2.27 | 0.4 | 96.74 | 6.07 | 11 | 1.05 | 1.61 | 0.2593 | 0.2762 | 0.0169 |
| 5fob | noschrodinger | 2.6 | 0.0031 | 0.78 | 0.51 | 96.23 | 11.67 | 0 | 4.93 | 2.32 | 0.2383 | 0.2634 | 0.0251 |
| 5for | opls | 2.5 | 0.089 | 2.42 | 0 | 98.18 | 2.59 | 1 | 0.9 | 1.09 | 0.2314 | 0.2645 | 0.0331 |

| | | | | | | | | | | | | | |
|---|---|---|---|---|---|---|---|---|---|---|---|---|---|
| 5for | noschrodinger | 2.5 | 0.01 | 0.95 | 0.81 | 95.12 | 6.03 | 0 | 3.63 | 2.09 | 0.2085 | 0.2546 | 0.0461 |
| 5fpz | opls | 1.5 | 0.1079 | 2.32 | 0 | 98.13 | 0 | 1 | 2.29 | 1.01 | 0.1438 | 0.1681 | 0.0243 |
| 5fpz | noschrodinger | 1.5 | 0.0061 | 1.04 | 0 | 98.15 | 0 | 0 | 2.29 | 1.01 | 0.1445 | 0.1647 | 0.0202 |
| 5fre | opls | 1.9 | 0.0991 | 2.13 | 0 | 96.41 | 0.4 | 1 | 1.29 | 1.09 | 0.1558 | 0.1948 | 0.039 |
| 5fre | noschrodinger | 1.9 | 0.0046 | 0.91 | 0 | 96.32 | 0.8 | 0 | 1.41 | 1.12 | 0.1566 | 0.1909 | 0.0343 |
| 5frw | opls | 1.96 | 0.093 | 2.34 | 0 | 97.2 | 1.57 | 1 | 1.77 | 1.23 | 0.1882 | 0.229 | 0.0408 |
| 5frw | noschrodinger | 1.96 | 0.0033 | 0.82 | 0.25 | 96.71 | 2.81 | 0 | 3.87 | 1.72 | 0.1928 | 0.2227 | 0.0299 |
| 5fsx | opls | 2 | 0.0927 | 2.35 | 0 | 97.79 | 0.52 | 2 | 0.41 | 0.69 | 0.234 | 0.2564 | 0.0224 |
| 5fsx | noschrodinger | 2 | 0.0028 | 0.82 | 0.22 | 96.72 | 1.04 | 0 | 1.38 | 1.09 | 0.2306 | 0.2554 | 0.0248 |
| 5fsy | opls | 1.7 | 0.0992 | 2.34 | 0 | 98.67 | 0.52 | 0 | 0.55 | 0.69 | 0.1944 | 0.2298 | 0.0354 |
| 5fsy | noschrodinger | 1.7 | 0.0027 | 0.82 | 0 | 98.25 | 0.52 | 0 | 1.09 | 0.82 | 0.2002 | 0.2281 | 0.0279 |
| 5fu3 | opls | 1.61 | 0.0931 | 2.19 | 0 | 97.34 | 1.85 | 0 | 0.66 | 1.05 | 0.2061 | 0.2454 | 0.0393 |
| 5fu3 | noschrodinger | 1.61 | 0.0092 | 1.44 | 0 | 97.88 | 0.61 | 0 | 5.61 | 1.33 | 0.197 | 0.2424 | 0.0454 |
| 5fuq | opls | 2.04 | 0.0974 | 2.24 | 0 | 96.84 | 1.26 | 4 | 1.36 | 1.13 | 0.1744 | 0.2008 | 0.0264 |
| 5fuq | noschrodinger | 2.04 | 0.0027 | 0.78 | 0 | 96.86 | 1.24 | 0 | 1.46 | 1.15 | 0.1791 | 0.1981 | 0.019 |
| 5fus | opls | 1.87 | 0.0973 | 2.26 | 0 | 97.32 | 1.08 | 5 | 1.31 | 1.01 | 0.1732 | 0.1981 | 0.0249 |
| 5fus | noschrodinger | 1.87 | 0.0035 | 0.83 | 0 | 97.14 | 1.23 | 0 | 2.78 | 1.29 | 0.1751 | 0.1942 | 0.0191 |
| 5fvj | opls | 1.7 | 0.1042 | 2.37 | 0 | 99.68 | 1.1 | 5 | 1.68 | 0.95 | 0.1793 | 0.2261 | 0.0468 |
| 5fvj | noschrodinger | 1.7 | 0.0038 | 0.86 | 0 | 99.37 | 0.36 | 0 | 5.22 | 1.28 | 0.1851 | 0.2229 | 0.0378 |
| 5g41 | opls | 1.54 | 0.0994 | 2.48 | 0 | 99.52 | 3.83 | 2 | 0.86 | 1.21 | 0.1877 | 0.2118 | 0.0241 |
| 5g41 | noschrodinger | 1.54 | 0.0095 | 1.27 | 0 | 99.53 | 3.26 | 0 | 0.57 | 1.08 | 0.1922 | 0.2115 | 0.0193 |
| 5g52 | opls | 3.8 | 0.0901 | 2.43 | 2.74 | 86.29 | 5.39 | 11 | 2.81 | 2.26 | 0.4602 | 0.4903 | 0.0301 |
| 5g52 | noschrodinger | 3.8 | 0.0034 | 0.95 | 3.06 | 82.43 | 7.96 | 0 | 13.39 | 3.02 | 0.4643 | 0.4839 | 0.0196 |
| 5g5g | opls | 1.7 | 0.0997 | 2.38 | 0.25 | 98.26 | 0.62 | 4 | 1.85 | 0.95 | 0.1417 | 0.1693 | 0.0276 |
| 5g5g | noschrodinger | 1.7 | 0.0046 | 0.98 | 0.16 | 97.69 | 0.82 | 0 | 1.37 | 0.94 | 0.1492 | 0.1689 | 0.0197 |
| 5g5h | opls | 2.3 | 0.0976 | 2.35 | 0.08 | 97.51 | 1.06 | 5 | 0.65 | 0.83 | 0.1679 | 0.2059 | 0.038 |
| 5g5h | noschrodinger | 2.3 | 0.0031 | 0.83 | 0.25 | 97.44 | 0.95 | 0 | 1.14 | 0.93 | 0.1709 | 0.1991 | 0.0282 |
| 5g5m | opls | 2.07 | 0.0929 | 2.24 | 0.6 | 95.2 | 0.22 | 1 | 0.84 | 1.09 | 0.2068 | 0.2436 | 0.0368 |
| 5g5m | noschrodinger | 2.07 | 0.0047 | 0.81 | 0.6 | 95.62 | 0.89 | 0 | 1.08 | 1.12 | 0.2076 | 0.2405 | 0.0329 |
| 5g5x | opls | 2.8 | 0.0893 | 2.32 | 0 | 96.07 | 2.06 | 1 | 0.54 | 1.19 | 0.2778 | 0.3112 | 0.0334 |
| 5g5x | noschrodinger | 2.8 | 0.0029 | 1.16 | 0 | 96.14 | 3.54 | 0 | 3.26 | 1.8 | 0.2882 | 0.3189 | 0.0307 |
| 5ggo | opls | 1.5 | 0.1022 | 2.21 | 0.35 | 98.26 | 0.41 | 1 | 2.74 | 1.06 | 0.1734 | 0.1998 | 0.0264 |
| 5ggo | noschrodinger | 1.5 | 0.0092 | 1.12 | 0 | 98 | 0.41 | 0 | 4.03 | 1.19 | 0.1714 | 0.2021 | 0.0307 |
| 5gi6 | opls | 1.65 | 0.0975 | 2.3 | 0 | 97.45 | 0.56 | 1 | 1.49 | 1 | 0.1956 | 0.2291 | 0.0335 |
| 5gi6 | noschrodinger | 1.65 | 0.0036 | 0.91 | 0 | 97.98 | 0.56 | 0 | 1.5 | 0.89 | 0.1998 | 0.2284 | 0.0286 |
| 5git | opls | 2.19 | 0.0927 | 2.29 | 0 | 96.85 | 3.6 | 1 | 0.48 | 1.28 | 0.247 | 0.2757 | 0.0287 |
| 5git | noschrodinger | 2.19 | 0.003 | 0.93 | 1.56 | 94.53 | 6.25 | 0 | 9.63 | 2.49 | 0.2349 | 0.2703 | 0.0354 |
| 5gkw | opls | 1.99 | 0.0963 | 2.33 | 0 | 96.33 | 2.37 | 2 | 1 | 1.33 | 0.2656 | 0.2981 | 0.0325 |
| 5gkw | noschrodinger | 1.99 | 0.0033 | 0.87 | 0.4 | 94.05 | 3.74 | 0 | 6.79 | 2.21 | 0.267 | 0.2928 | 0.0258 |
| 5glt | opls | 2 | 0.0981 | 2.64 | 2.9 | 92.75 | 3.96 | 23 | 4.19 | 2.11 | 0.1867 | 0.2316 | 0.0449 |
| 5glt | noschrodinger | 2 | 0.0121 | 1.31 | 3.99 | 92.21 | 2.5 | 1 | 14.67 | 2.45 | 0.195 | 0.2356 | 0.0406 |
| 5gly | opls | 1.58 | 0.1014 | 2.32 | 0 | 98.09 | 1.22 | 0 | 2.24 | 1.07 | 0.1467 | 0.167 | 0.0203 |
| 5gly | noschrodinger | 1.58 | 0.0089 | 1.45 | 0 | 98.58 | 1.21 | 0 | 6.08 | 1.4 | 0.15 | 0.1657 | 0.0157 |
| 5gm0 | opls | 1.7 | 0.1003 | 2.61 | 2.9 | 93.3 | 4.58 | 14 | 6.95 | 2.32 | 0.192 | 0.224 | 0.032 |
| 5gm0 | noschrodinger | 1.7 | 0.0139 | 1.47 | 3.8 | 92.75 | 3.12 | 1 | 13.56 | 2.48 | 0.196 | 0.2275 | 0.0315 |
| 5gro | opls | 2 | 0.0985 | 2.25 | 0 | 99.5 | 1.69 | 0 | 2.98 | 1.26 | 0.1698 | 0.2011 | 0.0313 |
| 5gro | noschrodinger | 2 | 0.0033 | 0.82 | 0 | 100 | 1.68 | 0 | 1.19 | 1 | 0.1759 | 0.2015 | 0.0256 |
| 5guf | opls | 2.4 | 0.0964 | 2.36 | 0 | 98.06 | 1.85 | 0 | 0 | 0.7 | 0.2054 | 0.24 | 0.0346 |
| 5guf | noschrodinger | 2.4 | 0.0036 | 0.92 | 0 | 98.72 | 0.91 | 0 | 3.77 | 1.17 | 0.2091 | 0.2444 | 0.0353 |

| | | | | | | | | | | | | | |
|---|---|---|---|---|---|---|---|---|---|---|---|---|---|
| 5gvr | opls | 1.5 | 0.0969 | 2.33 | 0 | 99.13 | 0.5 | 3 | 2.66 | 1.05 | 0.1798 | 0.2141 | 0.0343 |
| 5gvr | noschrodinger | 1.5 | 0.0072 | 1.02 | 0 | 99.57 | 1.49 | 0 | 2.66 | 1.18 | 0.1809 | 0.2105 | 0.0296 |
| 5gzf | opls | 2 | 0.0963 | 2.23 | 0.73 | 97.81 | 0.77 | 1 | 0.83 | 0.8 | 0.2032 | 0.2279 | 0.0247 |
| 5gzf | noschrodinger | 2 | 0.0028 | 0.85 | 0.68 | 97.3 | 1.53 | 0 | 1.67 | 1.19 | 0.2022 | 0.2268 | 0.0246 |
| 5h23 | opls | 2.2 | 0.093 | 2.26 | 0.34 | 97.99 | 2.45 | 2 | 1.45 | 1.18 | 0.2226 | 0.2406 | 0.018 |
| 5h23 | noschrodinger | 2.2 | 0.0031 | 0.8 | 0 | 96.64 | 2.45 | 0 | 2.28 | 1.52 | 0.1961 | 0.2388 | 0.0427 |
| 5h4s | opls | 1.8 | 0.1057 | 2.33 | 0 | 96.8 | 2.45 | 3 | 2.85 | 1.57 | 0.1525 | 0.1929 | 0.0404 |
| 5h4s | noschrodinger | 1.8 | 0.0041 | 0.9 | 0 | 96.45 | 3.27 | 0 | 2.14 | 1.61 | 0.1572 | 0.1856 | 0.0284 |
| 5h5o | opls | 2.12 | 0.0953 | 2.57 | 0.95 | 95.26 | 2.73 | 9 | 1.55 | 1.56 | 0.1898 | 0.2392 | 0.0494 |
| 5h5o | noschrodinger | 2.12 | 0.0054 | 1.39 | 2.05 | 94.26 | 5.13 | 0 | 6.97 | 2.31 | 0.1999 | 0.2285 | 0.0286 |
| 5h60 | opls | 3.64 | 0.0901 | 2.36 | 1 | 94.68 | 4.46 | 2 | 1.22 | 1.7 | 0.2577 | 0.3085 | 0.0508 |
| 5h60 | noschrodinger | 3.64 | 0.0028 | 0.76 | 0.66 | 93.38 | 5.19 | 0 | 8.38 | 2.43 | 0.2556 | 0.2935 | 0.0379 |
| 5h67 | opls | 2.07 | 0.0957 | 2.26 | 0 | 98.17 | 0.5 | 0 | 0.82 | 0.75 | 0.1928 | 0.2147 | 0.0219 |
| 5h67 | noschrodinger | 2.07 | 0.0029 | 0.77 | 0 | 97.73 | 0.99 | 0 | 2.04 | 1.03 | 0.1834 | 0.2144 | 0.031 |
| 5h6l | opls | 2.1 | 0.0942 | 2.26 | 0 | 99.08 | 0.26 | 1 | 0.27 | 0.6 | 0.1963 | 0.2228 | 0.0265 |
| 5h6l | noschrodinger | 2.1 | 0.0028 | 0.8 | 0 | 98.86 | 0 | 0 | 0.41 | 0.65 | 0.1914 | 0.2199 | 0.0285 |
| 5h6q | opls | 2.53 | 0.0906 | 2.25 | 0 | 97.47 | 0.45 | 0 | 0.39 | 0.75 | 0.2258 | 0.2324 | 0.0066 |
| 5h6q | noschrodinger | 2.53 | 0.0093 | 1.17 | 0.5 | 95.13 | 2.24 | 0 | 3.24 | 1.72 | 0.2154 | 0.2312 | 0.0158 |
| 5h6z | opls | 2 | 0.0992 | 2.31 | 0.42 | 96.19 | 0.92 | 1 | 1.74 | 1.19 | 0.1741 | 0.2163 | 0.0422 |
| 5h6z | noschrodinger | 2 | 0.0078 | 1 | 0 | 95.4 | 1.36 | 0 | 1.74 | 1.35 | 0.1745 | 0.213 | 0.0385 |
| 5h84 | opls | 2 | 0.0963 | 2.28 | 0.66 | 98.68 | 0.7 | 1 | 1.78 | 0.94 | 0.1602 | 0.2056 | 0.0454 |
| 5h84 | noschrodinger | 2 | 0.003 | 0.81 | 0.62 | 98.77 | 1.41 | 0 | 2.15 | 1.1 | 0.1497 | 0.1993 | 0.0496 |
| 5h9i | opls | 1.5 | 0.099 | 2.54 | 0 | 97.38 | 0.54 | 2 | 3.73 | 1.28 | 0.1656 | 0.1856 | 0.02 |
| 5h9i | noschrodinger | 1.5 | 0.01 | 1.1 | 0 | 97.48 | 0.54 | 0 | 2.4 | 1.13 | 0.1672 | 0.1827 | 0.0155 |
| 5hae | opls | 2 | 0.102 | 2.12 | 0 | 95.39 | 0.75 | 0 | 1.66 | 1.24 | 0.1773 | 0.2186 | 0.0413 |
| 5hae | noschrodinger | 2 | 0.0025 | 0.76 | 0 | 95.42 | 0 | 0 | 1.66 | 1.24 | 0.1492 | 0.1955 | 0.0463 |
| 5hc7 | opls | 2.05 | 0.0955 | 2.38 | 0.88 | 96.92 | 4.08 | 5 | 1.88 | 1.6 | 0.185 | 0.2119 | 0.0269 |
| 5hc7 | noschrodinger | 2.05 | 0.0028 | 0.78 | 0 | 96.49 | 4.57 | 0 | 1.88 | 1.68 | 0.1912 | 0.2181 | 0.0269 |
| 5hcn | opls | 2.2 | 0.0894 | 2.31 | 0.81 | 97.15 | 3.12 | 3 | 1.22 | 1.37 | 0.2482 | 0.2631 | 0.0149 |
| 5hcn | noschrodinger | 2.2 | 0.0029 | 0.77 | 0.8 | 96.79 | 6.17 | 0 | 2.2 | 1.8 | 0.2395 | 0.2556 | 0.0161 |
| 5hh0 | opls | 1.6 | 0.1028 | 2.46 | 0 | 99.44 | 0.59 | 1 | 3.99 | 1.19 | 0.1659 | 0.1969 | 0.031 |
| 5hh0 | noschrodinger | 1.6 | 0.0058 | 0.96 | 0 | 98.45 | 1.76 | 0 | 2.16 | 1.18 | 0.1672 | 0.1963 | 0.0291 |
| 5hif | opls | 1.6 | 0.0998 | 2.25 | 0.17 | 96.44 | 0.43 | 0 | 1.88 | 1.18 | 0.164 | 0.1923 | 0.0283 |
| 5hif | noschrodinger | 1.6 | 0.0068 | 1 | 0 | 95.96 | 0.43 | 0 | 1.32 | 1.14 | 0.1695 | 0.1923 | 0.0228 |
| 5hj1 | opls | 1.5 | 0.0979 | 2.65 | 0 | 98.9 | 2.38 | 3 | 2.98 | 1.37 | 0.2014 | 0.2333 | 0.0319 |
| 5hj1 | noschrodinger | 1.5 | 0.0137 | 1.36 | 1.05 | 98.95 | 2.33 | 0 | 1.78 | 1.21 | 0.1994 | 0.2251 | 0.0257 |
| 5hkf | opls | 2.25 | 0.0935 | 2.33 | 0 | 99.02 | 0 | 0 | 0.4 | 0.64 | 0.2076 | 0.2499 | 0.0423 |
| 5hkf | noschrodinger | 2.25 | 0.0027 | 0.77 | 0 | 97.13 | 3.59 | 0 | 3.01 | 1.67 | 0.2032 | 0.2475 | 0.0443 |
| 5hmc | opls | 1.95 | 0.0919 | 2.24 | 0 | 98.21 | 1.09 | 1 | 0 | 0.53 | 0.2046 | 0.2211 | 0.0165 |
| 5hmc | noschrodinger | 1.95 | 0.0027 | 0.75 | 0 | 98.37 | 0 | 0 | 1.07 | 0.81 | 0.1983 | 0.218 | 0.0197 |
| 5hmk | opls | 2.17 | 0.0923 | 2.3 | 0 | 95.32 | 1.83 | 0 | 0.63 | 1.23 | 0.2656 | 0.2924 | 0.0268 |
| 5hmk | noschrodinger | 2.17 | 0.0038 | 0.94 | 0 | 95.95 | 3.61 | 0 | 2.22 | 1.7 | 0.2475 | 0.2846 | 0.0371 |
| 5hol | opls | 1.59 | 0.102 | 2.24 | 0 | 99.39 | 0 | 0 | 1.52 | 0.89 | 0.1485 | 0.171 | 0.0225 |
| 5hol | noschrodinger | 1.59 | 0.0067 | 1.06 | 0 | 98.79 | 0 | 0 | 1.52 | 0.89 | 0.1469 | 0.1652 | 0.0183 |
| 5hqt | opls | 1.6 | 0.1004 | 2.34 | 0 | 98.7 | 2.14 | 0 | 1.9 | 1.2 | 0.177 | 0.1965 | 0.0195 |
| 5hqt | noschrodinger | 1.6 | 0.0066 | 0.91 | 0 | 98.28 | 1.06 | 0 | 1.36 | 0.88 | 0.1763 | 0.1955 | 0.0192 |
| 5hwc | opls | 2.05 | 0.1025 | 2.2 | 0 | 97.27 | 1.08 | 0 | 1.63 | 1.07 | 0.1629 | 0.2162 | 0.0533 |
| 5hwc | noschrodinger | 2.05 | 0.0073 | 0.98 | 0 | 97.3 | 3.23 | 0 | 2.72 | 1.58 | 0.1522 | 0.2091 | 0.0569 |
| 5hwh | opls | 1.79 | 0.1051 | 2.23 | 0 | 98.22 | 0.73 | 1 | 0.75 | 0.74 | 0.1627 | 0.1908 | 0.0281 |

| | | | | | | | | | | | | | |
|---|---|---|---|---|---|---|---|---|---|---|---|---|---|
| 5hwh | noschrodinger | 1.79 | 0.0065 | 0.96 | 0 | 98.27 | 1.45 | 0 | 1.89 | 1.07 | 0.1557 | 0.1895 | 0.0338 |
| 5hwi | opls | 1.75 | 0.0942 | 2.28 | 0 | 100 | 0 | 0 | 2.18 | 0.99 | 0.1739 | 0.2122 | 0.0383 |
| 5hwi | noschrodinger | 1.75 | 0.0039 | 0.84 | 0 | 99.11 | 0.53 | 0 | 1.94 | 0.96 | 0.1809 | 0.2114 | 0.0305 |
| 5hwu | opls | 2.1 | 0.0967 | 2.34 | 0 | 97.24 | 0.5 | 0 | 0.72 | 0.87 | 0.2275 | 0.2534 | 0.0259 |
| 5hwu | noschrodinger | 2.1 | 0.0029 | 0.77 | 0 | 97.31 | 0.49 | 0 | 2.66 | 1.18 | 0.208 | 0.2402 | 0.0322 |
| 5hwv | opls | 1.65 | 0.1034 | 2.32 | 0 | 100 | 1.01 | 0 | 2.27 | 1.01 | 0.1826 | 0.2198 | 0.0372 |
| 5hwv | noschrodinger | 1.65 | 0.0042 | 0.9 | 0 | 99.59 | 1.95 | 0 | 2.53 | 1.26 | 0.1834 | 0.2205 | 0.0371 |
| 5hww | opls | 2 | 0.0979 | 2.25 | 0 | 99.15 | 1.02 | 0 | 0.78 | 0.75 | 0.2196 | 0.2546 | 0.035 |
| 5hww | noschrodinger | 2 | 0.0026 | 0.73 | 0.42 | 98.32 | 3.54 | 0 | 2.34 | 1.43 | 0.2023 | 0.2446 | 0.0423 |
| 5hwy | opls | 2.1 | 0.0888 | 2.26 | 0 | 97.64 | 1.23 | 0 | 0.77 | 0.89 | 0.1836 | 0.2055 | 0.0219 |
| 5hwy | noschrodinger | 2.1 | 0.0037 | 0.79 | 0 | 97.32 | 1.22 | 0 | 1.18 | 1.03 | 0.1804 | 0.207 | 0.0266 |
| 5hxh | opls | 2.8 | 0.0909 | 2.3 | 0.18 | 96.49 | 1.93 | 2 | 0.63 | 1.16 | 0.2144 | 0.2591 | 0.0447 |
| 5hxh | noschrodinger | 2.8 | 0.0027 | 0.72 | 0.35 | 93.23 | 2.55 | 0 | 1.7 | 1.67 | 0.2059 | 0.2581 | 0.0522 |
| 5i0a | opls | 1.5 | 0.1052 | 2.4 | 0 | 97.84 | 0 | 0 | 2.2 | 1.03 | 0.1645 | 0.1937 | 0.0292 |
| 5i0a | noschrodinger | 1.5 | 0.003 | 0.82 | 0 | 98.57 | 0 | 0 | 1.32 | 0.86 | 0.1721 | 0.1898 | 0.0177 |
| 5i1q | opls | 1.49 | 0.1028 | 2.16 | 0 | 99.24 | 0 | 0 | 0.44 | 0.65 | 0.1615 | 0.1864 | 0.0249 |
| 5i1q | noschrodinger | 1.49 | 0.0041 | 0.88 | 0 | 100 | 2.31 | 0 | 0.44 | 0.93 | 0.1727 | 0.1835 | 0.0108 |
| 5i6c | opls | 3.7 | 0.0902 | 2.39 | 1.81 | 94.46 | 0.67 | 5 | 1.17 | 1.21 | 0.2757 | 0.3243 | 0.0486 |
| 5i6c | noschrodinger | 3.7 | 0.0033 | 0.8 | 1.7 | 91.08 | 0.4 | 0 | 6.5 | 1.88 | 0.2623 | 0.3126 | 0.0503 |
| 5i8b | opls | 1.52 | 0.0983 | 2.26 | 0.45 | 97.29 | 0.46 | 1 | 3.55 | 1.28 | 0.1834 | 0.2063 | 0.0229 |
| 5i8b | noschrodinger | 1.52 | 0.0066 | 1.15 | 0.45 | 97.3 | 0.92 | 0 | 6.33 | 1.48 | 0.1805 | 0.205 | 0.0245 |
| 5i8t | opls | 1.75 | 0.1057 | 2.28 | 0 | 97.22 | 1.23 | 0 | 2.93 | 1.3 | 0.1594 | 0.1829 | 0.0235 |
| 5i8t | noschrodinger | 1.75 | 0.0072 | 0.99 | 0 | 97.18 | 1.85 | 0 | 2.96 | 1.44 | 0.1597 | 0.1824 | 0.0227 |
| 5ian | opls | 2.7 | 0.0915 | 2.27 | 0 | 97.87 | 0.44 | 3 | 1.21 | 0.87 | 0.1907 | 0.2245 | 0.0338 |
| 5ian | noschrodinger | 2.7 | 0.0028 | 0.78 | 0 | 97.07 | 2.59 | 0 | 2.43 | 1.5 | 0.1801 | 0.218 | 0.0379 |
| 5iax | opls | 2.1 | 0.0927 | 2.18 | 0 | 97.77 | 0.63 | 2 | 0.49 | 0.72 | 0.2201 | 0.2487 | 0.0286 |
| 5iax | noschrodinger | 2.1 | 0.0035 | 0.81 | 0 | 97.65 | 0 | 0 | 0.83 | 0.83 | 0.1943 | 0.2416 | 0.0473 |
| 5ibo | opls | 1.95 | 0.091 | 2.29 | 0 | 97.47 | 1.87 | 0 | 2.68 | 1.37 | 0.2144 | 0.2483 | 0.0339 |
| 5ibo | noschrodinger | 1.95 | 0.0026 | 0.77 | 0 | 97.52 | 1.85 | 0 | 1.74 | 1.23 | 0.2209 | 0.2433 | 0.0224 |
| 5ie5 | opls | 2.38 | 0.0947 | 2.31 | 0.77 | 95.86 | 5.89 | 5 | 0.47 | 1.54 | 0.25 | 0.2728 | 0.0228 |
| 5ie5 | noschrodinger | 2.38 | 0.0031 | 0.87 | 0.75 | 95.65 | 9.76 | 0 | 2.45 | 2.08 | 0.2386 | 0.269 | 0.0304 |
| 5ie7 | opls | 2.5 | 0.0963 | 2.33 | 0.8 | 95.66 | 7.45 | 5 | 0.79 | 1.71 | 0.2471 | 0.2737 | 0.0266 |
| 5ie7 | noschrodinger | 2.5 | 0.0034 | 0.89 | 0.94 | 94.02 | 10.02 | 0 | 4.07 | 2.35 | 0.2468 | 0.2754 | 0.0286 |
| 5ien | opls | 2.09 | 0.0893 | 2.17 | 0 | 98.32 | 0 | 0 | 0.26 | 0.6 | 0.2135 | 0.2478 | 0.0343 |
| 5ien | noschrodinger | 2.09 | 0.0052 | 1.06 | 0.82 | 97.55 | 0.59 | 0 | 7.78 | 1.52 | 0.2034 | 0.2431 | 0.0397 |
| 5ieo | opls | 1.85 | 0.0919 | 2.11 | 0 | 100 | 0 | 0 | 1.59 | 0.9 | 0.1665 | 0.1899 | 0.0234 |
| 5ieo | noschrodinger | 1.85 | 0.0044 | 1.21 | 0 | 99.11 | 0 | 0 | 12.95 | 1.62 | 0.1667 | 0.1896 | 0.0229 |
| 5ijj | opls | 1.95 | 0.0935 | 2.43 | 0 | 99.07 | 0 | 3 | 1.43 | 0.88 | 0.2168 | 0.2467 | 0.0299 |
| 5ijj | noschrodinger | 1.95 | 0.006 | 1.69 | 0 | 98.78 | 1.39 | 0 | 2.87 | 1.19 | 0.2095 | 0.2431 | 0.0336 |
| 5ijp | opls | 2.75 | 0.0882 | 2.4 | 0 | 99.3 | 0.69 | 1 | 1.04 | 0.8 | 0.2502 | 0.2723 | 0.0221 |
| 5ijp | noschrodinger | 2.75 | 0.0039 | 1.07 | 0 | 97.67 | 2.03 | 0 | 0.7 | 1.03 | 0.2461 | 0.28 | 0.0339 |
| 5in9 | opls | 2.6 | 0.0909 | 2.41 | 1.19 | 96.66 | 0.78 | 3 | 1.24 | 1.06 | 0.2081 | 0.2496 | 0.0415 |
| 5in9 | noschrodinger | 2.6 | 0.0027 | 0.77 | 1.18 | 93.62 | 2.57 | 0 | 5.94 | 2.06 | 0.1928 | 0.2504 | 0.0576 |
| 5iqj | opls | 1.9 | 0.1007 | 2.31 | 0 | 100 | 1.39 | 6 | 2.79 | 1.18 | 0.1887 | 0.2308 | 0.0421 |
| 5iqj | noschrodinger | 1.9 | 0.0037 | 0.83 | 0 | 99.47 | 1.11 | 0 | 3.25 | 1.15 | 0.1889 | 0.2315 | 0.0426 |
| 5ir2 | opls | 2.08 | 0.0985 | 2.21 | 0 | 98.98 | 0 | 1 | 0.81 | 0.75 | 0.1768 | 0.1914 | 0.0146 |
| 5ir2 | noschrodinger | 2.08 | 0.0028 | 0.75 | 0 | 98.18 | 0 | 0 | 0.81 | 0.75 | 0.1713 | 0.19 | 0.0187 |
| 5iu1 | opls | 1.7 | 0.0996 | 2.25 | 0 | 98.59 | 0.51 | 0 | 1.97 | 0.96 | 0.157 | 0.1837 | 0.0267 |
| 5iu1 | noschrodinger | 1.7 | 0.0065 | 1.03 | 0 | 98.6 | 0.5 | 0 | 1.97 | 0.96 | 0.1556 | 0.1827 | 0.0271 |

| | | | | | | | | | | | | | |
|---|---|---|---|---|---|---|---|---|---|---|---|---|---|
| 5iu6 | opls | 2.51 | 0.0927 | 2.32 | 0.14 | 96.42 | 3.55 | 3 | 1.1 | 1.47 | 0.1835 | 0.2224 | 0.0389 |
| 5iu6 | noschrodinger | 2.51 | 0.003 | 0.81 | 0.28 | 95.89 | 5.83 | 0 | 1.65 | 1.78 | 0.1863 | 0.2136 | 0.0273 |
| 5iva | opls | 2.99 | 0.0894 | 2.3 | 1.1 | 93.52 | 3.29 | 6 | 0.92 | 1.6 | 0.2514 | 0.2793 | 0.0279 |
| 5iva | noschrodinger | 2.99 | 0.0027 | 0.75 | 0.83 | 91.33 | 3.12 | 0 | 3.17 | 1.99 | 0.2417 | 0.2717 | 0.03 |
| 5iw4 | opls | 2.6 | 0.0935 | 2.31 | 0.2 | 96.84 | 2.05 | 4 | 0.72 | 1.16 | 0.2305 | 0.249 | 0.0185 |
| 5iw4 | noschrodinger | 2.6 | 0.0028 | 0.76 | 0.39 | 95.69 | 4.06 | 0 | 2.98 | 1.85 | 0.215 | 0.2454 | 0.0304 |
| 5iyy | opls | 1.6 | 0.1057 | 2.14 | 0 | 96.75 | 0.71 | 1 | 0.78 | 0.95 | 0.1529 | 0.1812 | 0.0283 |
| 5iyy | noschrodinger | 1.6 | 0.0036 | 0.95 | 0 | 96.44 | 1.07 | 0 | 0.98 | 1.05 | 0.1502 | 0.1756 | 0.0254 |
| 5j0a | opls | 2.74 | 0.0927 | 2.32 | 1.34 | 92.81 | 4.76 | 4 | 1.26 | 1.82 | 0.2214 | 0.2576 | 0.0362 |
| 5j0a | noschrodinger | 2.74 | 0.003 | 0.8 | 1.84 | 88.13 | 8 | 0 | 5.69 | 2.59 | 0.2174 | 0.2474 | 0.03 |
| 5j1j | opls | 1.55 | 0.1031 | 2.47 | 0.39 | 98.06 | 1.16 | 1 | 4.49 | 1.28 | 0.1579 | 0.184 | 0.0261 |
| 5j1j | noschrodinger | 1.55 | 0.0111 | 1.2 | 0.39 | 97.87 | 1.39 | 0 | 4.01 | 1.33 | 0.1585 | 0.1812 | 0.0227 |
| 5j62 | opls | 2.15 | 0.0942 | 2.25 | 0.24 | 98.78 | 2.18 | 2 | 0.6 | 0.96 | 0.1954 | 0.2171 | 0.0217 |
| 5j62 | noschrodinger | 2.15 | 0.0029 | 0.8 | 0.24 | 97.82 | 3.79 | 0 | 0.45 | 1.14 | 0.1741 | 0.2005 | 0.0264 |
| 5j7m | opls | 2.06 | 0.0912 | 2.16 | 0 | 97.2 | 0 | 0 | 0.53 | 0.83 | 0.2001 | 0.2389 | 0.0388 |
| 5j7m | noschrodinger | 2.06 | 0.0027 | 0.81 | 0 | 97.08 | 0 | 0 | 0.54 | 0.85 | 0.1893 | 0.2273 | 0.038 |
| 5jf1 | opls | 2 | 0.1029 | 2.21 | 0 | 98.32 | 1.2 | 0 | 0.93 | 0.84 | 0.1749 | 0.2006 | 0.0257 |
| 5jf1 | noschrodinger | 2 | 0.0029 | 0.84 | 0 | 98.48 | 1.8 | 0 | 0 | 0.69 | 0.165 | 0.1888 | 0.0238 |
| 5jfe | opls | 2.03 | 0.0942 | 2.29 | 0 | 96.95 | 1.02 | 0 | 2.26 | 1.19 | 0.1627 | 0.1874 | 0.0247 |
| 5jfe | noschrodinger | 2.03 | 0.0028 | 0.8 | 0 | 97.61 | 1 | 0 | 1.01 | 0.88 | 0.1571 | 0.1834 | 0.0263 |
| 5jje | opls | 1.9 | 0.0891 | 2.22 | 0 | 99.64 | 2.46 | 0 | 2.71 | 1.36 | 0.1939 | 0.2153 | 0.0214 |
| 5jje | noschrodinger | 1.9 | 0.0039 | 0.89 | 0 | 99.29 | 2.93 | 0 | 3.98 | 1.54 | 0.1952 | 0.2134 | 0.0182 |
| 5jjj | opls | 2.5 | 0.0896 | 2.3 | 0 | 99.26 | 0.51 | 0 | 0.44 | 0.66 | 0.2151 | 0.2356 | 0.0205 |
| 5jjj | noschrodinger | 2.5 | 0.0028 | 0.73 | 0 | 97.79 | 1 | 0 | 1.56 | 0.95 | 0.1968 | 0.2238 | 0.027 |
| 5jlj | opls | 2.5 | 0.0925 | 2.26 | 0.15 | 97.52 | 0.66 | 2 | 0.68 | 0.82 | 0.2155 | 0.2412 | 0.0257 |
| 5jlj | noschrodinger | 2.5 | 0.0027 | 0.78 | 0.07 | 97.23 | 0.98 | 0 | 0.9 | 0.92 | 0.2047 | 0.2315 | 0.0268 |
| 5joh | opls | 1.99 | 0.0908 | 2.48 | 0 | 93.4 | 4.4 | 3 | 2.25 | 1.92 | 0.2202 | 0.2517 | 0.0315 |
| 5joh | noschrodinger | 1.99 | 0.0039 | 0.87 | 0 | 94.42 | 5.41 | 0 | 2.54 | 1.98 | 0.2315 | 0.2463 | 0.0148 |
| 5jqb | opls | 2.68 | 0.0899 | 2.24 | 0.55 | 95.57 | 0.32 | 1 | 0.96 | 1.09 | 0.2298 | 0.2458 | 0.016 |
| 5jqb | noschrodinger | 2.68 | 0.0045 | 0.82 | 0.54 | 94.61 | 1.57 | 0 | 2.42 | 1.54 | 0.2117 | 0.2293 | 0.0176 |
| 5jqc | opls | 2.15 | 0.0893 | 2.19 | 0 | 97.99 | 1.06 | 1 | 0.76 | 0.76 | 0.2234 | 0.259 | 0.0356 |
| 5jqc | noschrodinger | 2.15 | 0.0025 | 0.71 | 0.41 | 96.75 | 0 | 0 | 1.02 | 1 | 0.2351 | 0.2616 | 0.0265 |
| 5jwy | opls | 3.2 | 0.0897 | 2.31 | 0.82 | 97.53 | 3.83 | 1 | 0.46 | 1.2 | 0.2846 | 0.3077 | 0.0231 |
| 5jwy | noschrodinger | 3.2 | 0.0028 | 0.75 | 0.39 | 95.33 | 3.26 | 0 | 2.06 | 1.69 | 0.2694 | 0.2985 | 0.0291 |
| 5jxd | opls | 2.03 | 0.091 | 2.38 | 0 | 96.17 | 0.58 | 0 | 0.94 | 1.04 | 0.2416 | 0.2711 | 0.0295 |
| 5jxd | noschrodinger | 2.03 | 0.0027 | 0.71 | 0.54 | 97.28 | 1.16 | 0 | 4.07 | 1.38 | 0.2329 | 0.2623 | 0.0294 |
| 5jzv | opls | 2.07 | 0.101 | 2.29 | 0 | 98.26 | 2.53 | 0 | 0.35 | 0.93 | 0.1883 | 0.2034 | 0.0151 |
| 5jzv | noschrodinger | 2.07 | 0.003 | 0.83 | 0 | 95.95 | 5.03 | 0 | 0.35 | 1.44 | 0.1857 | 0.2006 | 0.0149 |
| 5k21 | opls | 1.8 | 0.0947 | 2.29 | 0.35 | 97.56 | 0 | 0 | 0.95 | 0.88 | 0.1713 | 0.207 | 0.0357 |
| 5k21 | noschrodinger | 1.8 | 0.0066 | 1.02 | 0.28 | 97.78 | 0 | 0 | 1.16 | 0.88 | 0.1799 | 0.2067 | 0.0268 |
| 5k25 | opls | 3.05 | 0.0893 | 2.38 | 0 | 95.31 | 10.69 | 1 | 0.82 | 1.86 | 0.2631 | 0.2798 | 0.0167 |
| 5k25 | noschrodinger | 3.05 | 0.0031 | 0.9 | 0.66 | 92.08 | 16.29 | 0 | 3.32 | 2.53 | 0.229 | 0.2609 | 0.0319 |
| 5k6w | opls | 3.5 | 0.0864 | 2.25 | 0.29 | 98.4 | 0.86 | 1 | 0.99 | 0.79 | 0.2052 | 0.2556 | 0.0504 |
| 5k6w | noschrodinger | 3.5 | 0.0028 | 0.77 | 0.32 | 95.88 | 0.29 | 0 | 2.89 | 1.36 | 0.2053 | 0.248 | 0.0427 |
| 5k7h | opls | 2.35 | 0.0895 | 2.34 | 0 | 98.69 | 0 | 0 | 0.33 | 0.62 | 0.2066 | 0.2243 | 0.0177 |
| 5k7h | noschrodinger | 2.35 | 0.0072 | 0.88 | 0 | 97.53 | 0 | 0 | 1.69 | 1.02 | 0.1906 | 0.217 | 0.0264 |
| 5k8k | opls | 2.55 | 0.0865 | 2.24 | 0 | 98.77 | 0.53 | 3 | 0.75 | 0.74 | 0.244 | 0.2616 | 0.0176 |
| 5k8k | noschrodinger | 2.55 | 0.003 | 0.77 | 0 | 97.02 | 0 | 0 | 0.76 | 0.91 | 0.2393 | 0.2639 | 0.0246 |
| 5k95 | opls | 2.77 | 0.0936 | 2.31 | 0.42 | 97.08 | 2.99 | 2 | 1.14 | 1.35 | 0.2409 | 0.2652 | 0.0243 |

| | | | | | | | | | | | | | |
|---|---|---|---|---|---|---|---|---|---|---|---|---|---|
| 5k95 | noschrodinger | 2.77 | 0.0028 | 0.78 | 0.62 | 94.58 | 5.75 | 0 | 2.53 | 1.99 | 0.2192 | 0.2509 | 0.0317 |
| 5kax | opls | 2 | 0.0967 | 2.15 | 0 | 98.65 | 1.62 | 0 | 0.83 | 0.92 | 0.2206 | 0.2469 | 0.0263 |
| 5kax | noschrodinger | 2 | 0.0026 | 0.75 | 0 | 98.07 | 0.81 | 0 | 3.53 | 1.14 | 0.2111 | 0.2426 | 0.0315 |
| 5kbe | opls | 2.5 | 0.0901 | 2.26 | 0.28 | 97.77 | 3.65 | 1 | 0.63 | 1.19 | 0.2037 | 0.2383 | 0.0346 |
| 5kbe | noschrodinger | 2.5 | 0.0031 | 0.79 | 0.25 | 96.22 | 7.23 | 0 | 2.39 | 1.93 | 0.1889 | 0.2296 | 0.0407 |
| 5kgo | opls | 2.9 | 0.0912 | 2.5 | 0.7 | 94.85 | 6.73 | 6 | 1.04 | 1.79 | 0.2488 | 0.2963 | 0.0475 |
| 5kgo | noschrodinger | 2.9 | 0.0037 | 0.88 | 1.38 | 89.45 | 9.65 | 0 | 7.4 | 2.72 | 0.2488 | 0.2815 | 0.0327 |
| 5khh | opls | 1.77 | 0.0988 | 2.32 | 0 | 98.82 | 2.96 | 0 | 3.39 | 1.49 | 0.1994 | 0.2371 | 0.0377 |
| 5khh | noschrodinger | 1.77 | 0.0059 | 0.86 | 0.53 | 96.84 | 2.96 | 0 | 5.29 | 1.83 | 0.1977 | 0.2306 | 0.0329 |
| 5kiq | opls | 1.64 | 0.1042 | 2.38 | 0 | 98.17 | 0.3 | 3 | 1.46 | 0.88 | 0.1872 | 0.2233 | 0.0361 |
| 5kiq | noschrodinger | 1.64 | 0.0033 | 0.87 | 0.26 | 96.63 | 0.88 | 0 | 3.58 | 1.36 | 0.1941 | 0.2178 | 0.0237 |
| 5kjy | opls | 2 | 0.0953 | 2.25 | 0 | 97.52 | 0 | 1 | 0.44 | 0.75 | 0.1976 | 0.2245 | 0.0269 |
| 5kjy | noschrodinger | 2 | 0.0058 | 0.91 | 0 | 96.32 | 0 | 0 | 0.88 | 1.02 | 0.1815 | 0.222 | 0.0405 |
| 5km8 | opls | 2 | 0.095 | 2.26 | 0 | 98.95 | 0 | 0 | 0 | 0.5 | 0.1837 | 0.2039 | 0.0202 |
| 5km8 | noschrodinger | 2 | 0.0029 | 0.86 | 0 | 99.05 | 1.17 | 0 | 0.59 | 0.75 | 0.161 | 0.1926 | 0.0316 |
| 5knt | opls | 2.55 | 0.0963 | 2.41 | 0.32 | 95.9 | 1.65 | 3 | 1.23 | 1.29 | 0.1743 | 0.2138 | 0.0395 |
| 5knt | noschrodinger | 2.55 | 0.0027 | 0.76 | 0.3 | 96.69 | 2.28 | 0 | 2.48 | 1.51 | 0.1937 | 0.2175 | 0.0238 |
| 5kqg | opls | 1.5 | 0.1056 | 2.57 | 0 | 98.68 | 1.38 | 0 | 4.64 | 1.34 | 0.1399 | 0.1664 | 0.0265 |
| 5kqg | noschrodinger | 1.5 | 0.006 | 1.09 | 0 | 98.68 | 1.38 | 0 | 2.7 | 1.16 | 0.1448 | 0.1618 | 0.017 |
| 5kqs | opls | 1.5 | 0.0987 | 2.32 | 0 | 98.23 | 0.97 | 2 | 4.46 | 1.22 | 0.1294 | 0.1548 | 0.0254 |
| 5kqs | noschrodinger | 1.5 | 0.0047 | 1.04 | 0 | 98.44 | 0 | 0 | 5.7 | 1.31 | 0.1344 | 0.1581 | 0.0237 |
| 5ks7 | opls | 2.9 | 0.0896 | 2.34 | 0.44 | 96 | 3.69 | 0 | 0 | 1.21 | 0.3153 | 0.348 | 0.0327 |
| 5ks7 | noschrodinger | 2.9 | 0.0029 | 0.81 | 0.41 | 93.78 | 11.36 | 0 | 2.75 | 2.28 | 0.2798 | 0.3272 | 0.0474 |
| 5ksd | opls | 3.5 | 0.0908 | 2.33 | 0.42 | 94.15 | 2.12 | 9 | 0.84 | 1.4 | 0.305 | 0.3461 | 0.0411 |
| 5ksd | noschrodinger | 3.5 | 0.0026 | 0.75 | 0.18 | 94.28 | 1.75 | 0 | 1.52 | 1.47 | 0.3005 | 0.339 | 0.0385 |
| 5ksp | opls | 2.16 | 0.1004 | 2.28 | 0 | 98.68 | 1.07 | 0 | 0.78 | 0.77 | 0.1967 | 0.2232 | 0.0265 |
| 5ksp | noschrodinger | 2.16 | 0.0029 | 0.79 | 0 | 98.05 | 0.35 | 0 | 1.95 | 0.96 | 0.176 | 0.2193 | 0.0433 |
| 5ktc | opls | 1.8 | 0.0976 | 2.27 | 0 | 96.76 | 1.75 | 3 | 0.92 | 1.17 | 0.1974 | 0.2311 | 0.0337 |
| 5ktc | noschrodinger | 1.8 | 0.0038 | 0.88 | 0 | 95.14 | 1.75 | 0 | 1.85 | 1.47 | 0.1976 | 0.2261 | 0.0285 |
| 5kym | opls | 2.8 | 0.0874 | 2.27 | 0.23 | 96.61 | 2.72 | 5 | 1.28 | 1.4 | 0.2629 | 0.2786 | 0.0157 |
| 5kym | noschrodinger | 2.8 | 0.0086 | 1.52 | 0.86 | 93.95 | 4.87 | 0 | 4.23 | 2.13 | 0.2433 | 0.266 | 0.0227 |
| 5kzv | opls | 1.62 | 0.1085 | 2.25 | 0 | 97.37 | 0 | 0 | 1.6 | 1.03 | 0.1703 | 0.2193 | 0.049 |
| 5kzv | noschrodinger | 1.62 | 0.0039 | 0.95 | 0 | 97.39 | 0 | 0 | 9.1 | 1.6 | 0.1695 | 0.2124 | 0.0429 |
| 5kzy | opls | 2.48 | 0.0908 | 2.26 | 0 | 96.43 | 2.49 | 0 | 0.54 | 1.22 | 0.2343 | 0.2618 | 0.0275 |
| 5kzy | noschrodinger | 2.48 | 0.0036 | 1.18 | 0.44 | 94.69 | 2.96 | 0 | 13.03 | 2.35 | 0.2183 | 0.2644 | 0.0461 |
| 5l09 | opls | 1.99 | 0.0931 | 2.33 | 0 | 99.32 | 2.35 | 1 | 1.24 | 1.13 | 0.1715 | 0.2156 | 0.0441 |
| 5l09 | noschrodinger | 1.99 | 0.0126 | 1.06 | 0 | 98.75 | 3.56 | 0 | 3.38 | 1.55 | 0.1705 | 0.2166 | 0.0461 |
| 5l1a | opls | 2.4 | 0.0955 | 2.25 | 0 | 100 | 0 | 0 | 0.55 | 0.69 | 0.1909 | 0.244 | 0.0531 |
| 5l1a | noschrodinger | 2.4 | 0.0025 | 0.69 | 0 | 98.08 | 0 | 0 | 4.94 | 1.26 | 0.1748 | 0.2282 | 0.0534 |
| 5l1n | opls | 3.6 | 0.09 | 2.32 | 0.11 | 97.15 | 1.84 | 1 | 0.64 | 1.07 | 0.2432 | 0.2828 | 0.0396 |
| 5l1n | noschrodinger | 3.6 | 0.0031 | 0.84 | 0.46 | 96.25 | 6.63 | 0 | 4.18 | 2.08 | 0.2318 | 0.2699 | 0.0381 |
| 5l3e | opls | 2.8 | 0.0893 | 2.37 | 0.13 | 97.23 | 0.29 | 7 | 0.38 | 0.78 | 0.2246 | 0.2478 | 0.0232 |
| 5l3e | noschrodinger | 2.8 | 0.003 | 0.81 | 0 | 97.11 | 0.15 | 0 | 1.83 | 1.1 | 0.2362 | 0.2509 | 0.0147 |
| 5l6w | opls | 2.53 | 0.0829 | 2.3 | 0.79 | 95.24 | 2.6 | 6 | 1.33 | 1.51 | 0.2454 | 0.2674 | 0.022 |
| 5l6w | noschrodinger | 2.53 | 0.0032 | 0.88 | 2 | 90.87 | 3.55 | 0 | 2.62 | 1.99 | 0.239 | 0.2617 | 0.0227 |
| 5l73 | opls | 2.24 | 0.0938 | 2.2 | 0.29 | 96.18 | 1.32 | 1 | 1.08 | 1.16 | 0.1804 | 0.2091 | 0.0287 |
| 5l73 | noschrodinger | 2.24 | 0.0071 | 0.97 | 1.45 | 93.9 | 1.98 | 0 | 1.62 | 1.54 | 0.159 | 0.2048 | 0.0458 |
| 5l8x | opls | 1.85 | 0.0938 | 2.32 | 0 | 96.73 | 0.74 | 2 | 0.19 | 0.78 | 0.1998 | 0.2349 | 0.0351 |
| 5l8x | noschrodinger | 1.85 | 0.003 | 0.8 | 0.29 | 97.35 | 1.82 | 0 | 0.59 | 1.02 | 0.2083 | 0.2303 | 0.022 |

| | | | | | | | | | | | | | |
|---|---|---|---|---|---|---|---|---|---|---|---|---|---|
| 5lb1 | opls | 1.55 | 0.1042 | 2.26 | 0 | 98 | 1.15 | 0 | 2.11 | 1.03 | 0.1642 | 0.1867 | 0.0225 |
| 5lb1 | noschrodinger | 1.55 | 0.0059 | 1.04 | 0 | 97.43 | 0.91 | 0 | 2.36 | 1.13 | 0.164 | 0.1882 | 0.0242 |
| 5lb6 | opls | 1.7 | 0.0975 | 2.34 | 0 | 97.93 | 0.55 | 1 | 2.8 | 1.09 | 0.1684 | 0.1982 | 0.0298 |
| 5lb6 | noschrodinger | 1.7 | 0.0041 | 0.86 | 0 | 98.16 | 1.09 | 0 | 1.41 | 0.9 | 0.1774 | 0.2006 | 0.0232 |
| 5lc0 | opls | 2.7 | 0.0861 | 2.34 | 0 | 94.86 | 8.33 | 4 | 1.36 | 1.92 | 0.3533 | 0.3787 | 0.0254 |
| 5lc0 | noschrodinger | 2.7 | 0.0037 | 0.88 | 0.8 | 89.07 | 9.8 | 0 | 6.99 | 2.71 | 0.3541 | 0.3829 | 0.0288 |
| 5lf5 | opls | 3.5 | 0.0884 | 2.32 | 0.42 | 94.74 | 2.16 | 1 | 1.17 | 1.45 | 0.255 | 0.2911 | 0.0361 |
| 5lf5 | noschrodinger | 3.5 | 0.0031 | 0.8 | 0.42 | 93.96 | 2.15 | 0 | 5.36 | 1.94 | 0.2459 | 0.2792 | 0.0333 |
| 5lir | opls | 1.75 | 0.0949 | 2.34 | 0 | 98.58 | 0.56 | 1 | 1.18 | 0.83 | 0.1726 | 0.1908 | 0.0182 |
| 5lir | noschrodinger | 1.75 | 0.0095 | 1.11 | 0 | 99.53 | 0 | 0 | 0.88 | 0.77 | 0.1676 | 0.1914 | 0.0238 |
| 5lkm | opls | 3.19 | 0.0903 | 2.36 | 0.47 | 95.77 | 3.01 | 7 | 1.14 | 1.48 | 0.2231 | 0.2719 | 0.0488 |
| 5lkm | noschrodinger | 3.19 | 0.0032 | 0.79 | 0.37 | 93.62 | 4.56 | 0 | 2.01 | 1.89 | 0.2279 | 0.2578 | 0.0299 |
| 5lkw | opls | 2 | 0.0983 | 2.21 | 0 | 97.41 | 1 | 0 | 0.26 | 0.72 | 0.2089 | 0.2354 | 0.0265 |
| 5lkw | noschrodinger | 2 | 0.0026 | 0.77 | 0 | 98.72 | 4.95 | 0 | 1.33 | 1.39 | 0.1955 | 0.2254 | 0.0299 |
| 5llt | opls | 2.2 | 0.0941 | 2.26 | 0 | 99.26 | 0.51 | 0 | 0.56 | 0.69 | 0.18 | 0.2224 | 0.0424 |
| 5llt | noschrodinger | 2.2 | 0.0031 | 0.78 | 0 | 98.28 | 1.53 | 0 | 2.38 | 1.16 | 0.1672 | 0.212 | 0.0448 |
| 5lm3 | opls | 2.5 | 0.092 | 2.25 | 0 | 98.97 | 0.53 | 0 | 0.29 | 0.61 | 0.2154 | 0.2587 | 0.0433 |
| 5lm3 | noschrodinger | 2.5 | 0.0025 | 0.72 | 0 | 98.48 | 0 | 0 | 1.17 | 0.83 | 0.2096 | 0.2574 | 0.0478 |
| 5lo8 | opls | 2.5 | 0.0891 | 2.41 | 1.48 | 95.56 | 3.69 | 1 | 3.68 | 1.9 | 0.273 | 0.3285 | 0.0555 |
| 5lo8 | noschrodinger | 2.5 | 0.0026 | 0.71 | 1.08 | 89.93 | 1.63 | 0 | 3.49 | 1.85 | 0.2786 | 0.3365 | 0.0579 |
| 5lpg | opls | 1.7 | 0.0958 | 2.32 | 0 | 98.31 | 0.75 | 0 | 1.21 | 0.84 | 0.2342 | 0.2626 | 0.0284 |
| 5lpg | noschrodinger | 1.7 | 0.0043 | 0.86 | 0 | 98.05 | 0.75 | 0 | 2.03 | 0.97 | 0.2292 | 0.2608 | 0.0316 |
| 5lq7 | opls | 1.6 | 0.1011 | 2.37 | 0.49 | 97.54 | 0.5 | 3 | 2.6 | 1.14 | 0.1517 | 0.1942 | 0.0425 |
| 5lq7 | noschrodinger | 1.6 | 0.0091 | 1.16 | 0.49 | 98.04 | 1 | 0 | 3.17 | 1.11 | 0.1543 | 0.1969 | 0.0426 |
| 5lq8 | opls | 1.52 | 0.0953 | 2.2 | 0 | 99.26 | 1.25 | 0 | 2.02 | 1.04 | 0.1881 | 0.2092 | 0.0211 |
| 5lq8 | noschrodinger | 1.52 | 0.0049 | 0.89 | 0 | 97.41 | 0 | 0 | 1.35 | 0.98 | 0.1886 | 0.2091 | 0.0205 |
| 5lsx | opls | 2.69 | 0.0893 | 2.3 | 0 | 94.37 | 4.02 | 1 | 0.27 | 1.44 | 0.2703 | 0.29 | 0.0197 |
| 5lsx | noschrodinger | 2.69 | 0.0027 | 0.75 | 0.89 | 89.33 | 8 | 0 | 4.36 | 2.47 | 0.2457 | 0.2817 | 0.036 |
| 5lua | opls | 2 | 0.0948 | 2.26 | 0 | 97.25 | 0.44 | 3 | 2.48 | 1.17 | 0.2018 | 0.2311 | 0.0293 |
| 5lua | noschrodinger | 2 | 0.0028 | 0.79 | 0.19 | 97.67 | 2.19 | 0 | 2.49 | 1.36 | 0.202 | 0.2275 | 0.0255 |
| 5luh | opls | 1.73 | 0.0998 | 2.27 | 0 | 99.2 | 0.87 | 0 | 1.44 | 0.88 | 0.1827 | 0.2122 | 0.0295 |
| 5luh | noschrodinger | 1.73 | 0.0055 | 1.54 | 0 | 99.01 | 1.51 | 0 | 1.99 | 1.1 | 0.1815 | 0.2156 | 0.0341 |
| 5lui | opls | 1.5 | 0.1033 | 2.31 | 0 | 97.7 | 0.92 | 2 | 1.98 | 1.03 | 0.1424 | 0.1768 | 0.0344 |
| 5lui | noschrodinger | 1.5 | 0.0054 | 1.01 | 0 | 97.33 | 0.91 | 0 | 1.98 | 1.09 | 0.1438 | 0.1739 | 0.0301 |
| 5lva | opls | 2.53 | 0.0944 | 2.22 | 0.59 | 95.28 | 0.35 | 0 | 0.54 | 1.01 | 0.2047 | 0.2364 | 0.0317 |
| 5lva | noschrodinger | 2.53 | 0.0036 | 0.84 | 1.17 | 93.57 | 1.4 | 0 | 3.25 | 1.65 | 0.1927 | 0.2329 | 0.0402 |
| 5lw0 | opls | 1.65 | 0.1004 | 2.55 | 0 | 98.25 | 0.28 | 5 | 3.47 | 1.14 | 0.1494 | 0.1799 | 0.0305 |
| 5lw0 | noschrodinger | 1.65 | 0.0104 | 1.25 | 0 | 98.51 | 0.55 | 0 | 5.13 | 1.27 | 0.1496 | 0.1767 | 0.0271 |
| 5lwy | opls | 2.4 | 0.0952 | 2.26 | 0 | 98 | 4.43 | 1 | 1.16 | 1.32 | 0.2448 | 0.2743 | 0.0295 |
| 5lwy | noschrodinger | 2.4 | 0.0026 | 0.76 | 0.2 | 97.62 | 7.85 | 0 | 1.97 | 1.72 | 0.2322 | 0.2689 | 0.0367 |
| 5lz6 | opls | 2.6 | 0.0884 | 2.17 | 0 | 96.3 | 1.5 | 0 | 0 | 0.88 | 0.2176 | 0.2388 | 0.0212 |
| 5lz6 | noschrodinger | 2.6 | 0.0025 | 0.73 | 0 | 96.03 | 0.74 | 0 | 1.21 | 1.11 | 0.2326 | 0.254 | 0.0214 |
| 5m0y | opls | 1.5 | 0.0961 | 2.43 | 0 | 97.54 | 0.7 | 1 | 2.61 | 1.14 | 0.1726 | 0.1999 | 0.0273 |
| 5m0y | noschrodinger | 1.5 | 0.0088 | 1.1 | 0 | 97.86 | 0.69 | 0 | 1.12 | 0.85 | 0.176 | 0.1951 | 0.0191 |
| 5m3e | opls | 2.5 | 0.0917 | 2.28 | 0 | 95.97 | 0.79 | 0 | 0 | 0.78 | 0.2102 | 0.265 | 0.0548 |
| 5m3e | noschrodinger | 2.5 | 0.0029 | 0.79 | 0 | 95.33 | 3.12 | 0 | 0.39 | 1.34 | 0.1981 | 0.2494 | 0.0513 |
| 5m42 | opls | 2.2 | 0.0943 | 2.31 | 0 | 98.33 | 2.96 | 0 | 0.25 | 0.95 | 0.1906 | 0.2251 | 0.0345 |
| 5m42 | noschrodinger | 2.2 | 0.0029 | 0.82 | 0 | 98.33 | 6.9 | 0 | 1.48 | 1.52 | 0.1861 | 0.2157 | 0.0296 |
| 5m47 | opls | 2.59 | 0.0932 | 2.29 | 0 | 97.1 | 0.92 | 1 | 1.22 | 1 | 0.2214 | 0.2441 | 0.0227 |

| | | | | | | | | | | | | | |
|---|---|---|---|---|---|---|---|---|---|---|---|---|---|
| 5m47 | noschrodinger | 2.59 | 0.0043 | 0.96 | 0 | 97.12 | 5.45 | 0 | 3.17 | 1.83 | 0.1987 | 0.2359 | 0.0372 |
| 5m5e | opls | 2.3 | 0.0928 | 2.25 | 0 | 96.19 | 0.63 | 2 | 0.68 | 0.98 | 0.1981 | 0.221 | 0.0229 |
| 5m5e | noschrodinger | 2.3 | 0.0029 | 0.78 | 0 | 95.27 | 1.45 | 0 | 2.05 | 1.43 | 0.1953 | 0.2164 | 0.0211 |
| 5m5q | opls | 2.2 | 0.0914 | 2.2 | 0 | 96.67 | 1.67 | 1 | 0.84 | 1.14 | 0.2078 | 0.219 | 0.0112 |
| 5m5q | noschrodinger | 2.2 | 0.0035 | 0.83 | 0.47 | 95.31 | 3.28 | 0 | 1.13 | 1.54 | 0.1976 | 0.2271 | 0.0295 |
| 5mfn | opls | 2.8 | 0.0906 | 2.33 | 0 | 99.09 | 3.67 | 0 | 0.6 | 1.13 | 0.2873 | 0.3273 | 0.04 |
| 5mfn | noschrodinger | 2.8 | 0.003 | 0.76 | 0 | 98.74 | 6.62 | 0 | 1.08 | 1.44 | 0.2923 | 0.3213 | 0.029 |
| 5mfu | opls | 2.15 | 0.0962 | 2.29 | 0 | 98.8 | 0.46 | 0 | 0 | 0.5 | 0.2161 | 0.2457 | 0.0296 |
| 5mfu | noschrodinger | 2.15 | 0.0034 | 0.82 | 0.4 | 97.62 | 0.91 | 0 | 1.44 | 0.96 | 0.1959 | 0.2439 | 0.048 |
| 5miu | opls | 3.5 | 0.0909 | 2.31 | 0.11 | 96.27 | 3.81 | 2 | 0.21 | 1.27 | 0.244 | 0.2659 | 0.0219 |
| 5miu | noschrodinger | 3.5 | 0.0028 | 0.76 | 0.34 | 93.03 | 7.17 | 0 | 1.77 | 2.03 | 0.2215 | 0.2604 | 0.0389 |
| 5mj1 | opls | 1.8 | 0.0966 | 2.36 | 0 | 95.77 | 1.3 | 1 | 8.28 | 1.83 | 0.176 | 0.2202 | 0.0442 |
| 5mj1 | noschrodinger | 1.8 | 0.0053 | 0.97 | 0 | 94.81 | 0 | 0 | 4.54 | 1.59 | 0.1813 | 0.2198 | 0.0385 |
| 5mje | opls | 2.6 | 0.088 | 2.26 | 1.04 | 95.83 | 3.31 | 1 | 3.89 | 1.86 | 0.2902 | 0.32 | 0.0298 |
| 5mje | noschrodinger | 2.6 | 0.0027 | 0.77 | 0.44 | 93.89 | 6.38 | 0 | 5.35 | 2.31 | 0.2495 | 0.302 | 0.0525 |
| 5mji | opls | 2 | 0.0981 | 2.48 | 0.4 | 97.59 | 0.95 | 3 | 3.23 | 1.2 | 0.173 | 0.1965 | 0.0235 |
| 5mji | noschrodinger | 2 | 0.0091 | 1.19 | 0.8 | 98 | 1.42 | 0 | 3.73 | 1.28 | 0.1802 | 0.1971 | 0.0169 |
| 5mju | opls | 3.4 | 0.0906 | 2.36 | 0 | 95.51 | 6.5 | 1 | 1.1 | 1.75 | 0.2629 | 0.3005 | 0.0376 |
| 5mju | noschrodinger | 3.4 | 0.0028 | 0.75 | 0 | 93.02 | 9.79 | 0 | 1.58 | 2.1 | 0.2545 | 0.2869 | 0.0324 |
| 5ml2 | opls | 1.6 | 0.0994 | 2.22 | 0 | 98.47 | 0 | 0 | 1.24 | 0.84 | 0.1906 | 0.2179 | 0.0273 |
| 5ml2 | noschrodinger | 1.6 | 0.0051 | 1.04 | 0 | 98.57 | 0 | 0 | 0.42 | 0.65 | 0.1949 | 0.2188 | 0.0239 |
| 5mn8 | opls | 3.5 | 0.0902 | 2.34 | 0.34 | 96.76 | 3.56 | 0 | 0.8 | 1.37 | 0.2548 | 0.3031 | 0.0483 |
| 5mn8 | noschrodinger | 3.5 | 0.0026 | 0.74 | 0.85 | 91.19 | 0.66 | 0 | 0.8 | 1.26 | 0.2531 | 0.2923 | 0.0392 |
| 5mob | opls | 1.67 | 0.1005 | 2.24 | 0 | 98.42 | 1.74 | 1 | 0.98 | 0.97 | 0.1891 | 0.2091 | 0.02 |
| 5mob | noschrodinger | 1.67 | 0.003 | 0.95 | 0 | 97.91 | 4.65 | 0 | 0.66 | 1.24 | 0.1921 | 0.2084 | 0.0163 |
| 5mwa | opls | 1.46 | 0.0963 | 2.46 | 0 | 99 | 0.57 | 0 | 1.77 | 0.93 | 0.1781 | 0.2093 | 0.0312 |
| 5mwa | noschrodinger | 1.46 | 0.0131 | 1.33 | 0 | 99.01 | 1.69 | 0 | 2.67 | 1.23 | 0.1797 | 0.2062 | 0.0265 |
| 5myi | opls | 1.9 | 0.0946 | 2.27 | 0 | 97.04 | 0.68 | 3 | 2.17 | 1.16 | 0.1682 | 0.1998 | 0.0316 |
| 5myi | noschrodinger | 1.9 | 0.0034 | 0.82 | 0 | 98.16 | 1.37 | 0 | 1.1 | 0.92 | 0.1747 | 0.2025 | 0.0278 |
| 5mza | opls | 2.78 | 0.0898 | 2.29 | 0.33 | 97.69 | 0.75 | 3 | 0.8 | 0.82 | 0.2169 | 0.2451 | 0.0282 |
| 5mza | noschrodinger | 2.78 | 0.0086 | 1.23 | 0.16 | 96.75 | 1.11 | 0 | 1.6 | 1.15 | 0.208 | 0.2395 | 0.0315 |
| 5n0g | opls | 1.57 | 0.1 | 2.44 | 0 | 98.05 | 0 | 0 | 0.61 | 0.7 | 0.184 | 0.2076 | 0.0236 |
| 5n0g | noschrodinger | 1.57 | 0.0106 | 1.36 | 0 | 98.54 | 0.65 | 0 | 1.82 | 0.94 | 0.1837 | 0.2073 | 0.0236 |
| 5n1u | opls | 2.98 | 0.0882 | 2.41 | 0.36 | 97.08 | 5.73 | 2 | 1.1 | 1.56 | 0.2623 | 0.2881 | 0.0258 |
| 5n1u | noschrodinger | 2.98 | 0.0028 | 0.78 | 0.61 | 92.73 | 6.38 | 0 | 3.16 | 2.18 | 0.2503 | 0.2686 | 0.0183 |
| 5n21 | opls | 1.58 | 0.1002 | 2.31 | 0 | 95.76 | 0.89 | 1 | 2.17 | 1.29 | 0.1499 | 0.1895 | 0.0396 |
| 5n21 | noschrodinger | 1.58 | 0.0037 | 1.02 | 0 | 97.08 | 1.31 | 0 | 1.45 | 1.13 | 0.157 | 0.1869 | 0.0299 |
| 5n92 | opls | 2.3 | 0.0947 | 2.33 | 0 | 98.64 | 0.94 | 2 | 1.09 | 0.81 | 0.2062 | 0.2229 | 0.0167 |
| 5n92 | noschrodinger | 2.3 | 0.0028 | 0.83 | 0.45 | 97.31 | 1.4 | 0 | 1.09 | 1.06 | 0.2175 | 0.2327 | 0.0152 |
| 5nc8 | opls | 3.09 | 0.0903 | 2.27 | 0.33 | 98.69 | 1.62 | 1 | 0.6 | 0.86 | 0.2438 | 0.2635 | 0.0197 |
| 5nc8 | noschrodinger | 3.09 | 0.0044 | 0.88 | 0.97 | 93.85 | 5.6 | 0 | 2.41 | 2 | 0.233 | 0.2521 | 0.0191 |
| 5ndz | opls | 3.6 | 0.0909 | 2.25 | 0 | 96.71 | 0.21 | 5 | 0.44 | 0.86 | 0.2544 | 0.2764 | 0.022 |
| 5ndz | noschrodinger | 3.6 | 0.0027 | 0.72 | 0 | 96.55 | 0.42 | 0 | 1.33 | 1.08 | 0.2459 | 0.2758 | 0.0299 |
| 5ngz | opls | 2.4 | 0.094 | 2.24 | 0 | 99.32 | 0.75 | 0 | 0.41 | 0.65 | 0.2201 | 0.2296 | 0.0095 |
| 5ngz | noschrodinger | 2.4 | 0.0029 | 0.86 | 0 | 98 | 0 | 0 | 1.63 | 0.91 | 0.2116 | 0.2263 | 0.0147 |
| 5nh0 | opls | 2.35 | 0.0968 | 2.28 | 0 | 98.19 | 1.74 | 1 | 0.58 | 0.88 | 0.2084 | 0.2366 | 0.0282 |
| 5nh0 | noschrodinger | 2.35 | 0.0026 | 0.78 | 0 | 98.43 | 3.46 | 0 | 1.17 | 1.24 | 0.2018 | 0.2344 | 0.0326 |
| 5nim | opls | 1.5 | 0.0982 | 2.58 | 0 | 99.44 | 0.6 | 2 | 5.59 | 1.3 | 0.2105 | 0.2264 | 0.0159 |
| 5nim | noschrodinger | 1.5 | 0.0178 | 1.57 | 0 | 97.94 | 1.2 | 1 | 2.47 | 1.11 | 0.1738 | 0.2049 | 0.0311 |

| | | | | | | | | | | | | | |
|---|---|---|---|---|---|---|---|---|---|---|---|---|---|
| 5nmm | opls | 2.02 | 0.0918 | 2.26 | 0 | 97.92 | 0.53 | 0 | 0.87 | 0.79 | 0.2028 | 0.2333 | 0.0305 |
| 5nmm | noschrodinger | 2.02 | 0.0029 | 0.79 | 0 | 95.26 | 1.06 | 0 | 1.46 | 1.23 | 0.1878 | 0.2316 | 0.0438 |
| 5nmo | opls | 1.9 | 0.0914 | 2.47 | 0.38 | 99.62 | 0.4 | 6 | 1.98 | 0.97 | 0.2051 | 0.2399 | 0.0348 |
| 5nmo | noschrodinger | 1.9 | 0.0143 | 1.2 | 0.34 | 99.32 | 0.39 | 0 | 3.2 | 1.11 | 0.2074 | 0.242 | 0.0346 |
| 5non | opls | 1.85 | 0.0994 | 2.4 | 0 | 98.68 | 0.98 | 4 | 6.26 | 1.34 | 0.1897 | 0.2363 | 0.0466 |
| 5non | noschrodinger | 1.85 | 0.004 | 0.92 | 0 | 98.71 | 1.46 | 0 | 2.35 | 1.14 | 0.1961 | 0.2319 | 0.0358 |
| 5npl | opls | 2.79 | 0.0905 | 2.35 | 0 | 99.39 | 0.93 | 0 | 0.59 | 0.7 | 0.1825 | 0.2373 | 0.0548 |
| 5npl | noschrodinger | 2.79 | 0.0028 | 0.82 | 0 | 98.12 | 2.31 | 0 | 1.54 | 1.17 | 0.1738 | 0.2288 | 0.055 |
| 5nro | opls | 3.25 | 0.0908 | 2.24 | 0.15 | 98.18 | 0.73 | 1 | 0.49 | 0.67 | 0.2441 | 0.2661 | 0.022 |
| 5nro | noschrodinger | 3.25 | 0.0029 | 0.77 | 0.15 | 97.58 | 0.37 | 0 | 1.85 | 1.03 | 0.2182 | 0.2583 | 0.0401 |
| 5nrr | opls | 1.7 | 0.1031 | 2.31 | 1.15 | 96.56 | 0 | 1 | 1.99 | 1.19 | 0.1851 | 0.2031 | 0.018 |
| 5nrr | noschrodinger | 1.7 | 0.0133 | 1.2 | 0.75 | 96.99 | 0 | 0 | 3.98 | 1.36 | 0.1856 | 0.2083 | 0.0227 |
| 5nv7 | opls | 2.57 | 0.0922 | 2.22 | 0.8 | 96.8 | 0.96 | 0 | 0.94 | 0.98 | 0.2071 | 0.2431 | 0.036 |
| 5nv7 | noschrodinger | 2.57 | 0.0028 | 0.74 | 0.39 | 94.09 | 4.25 | 0 | 2.11 | 1.86 | 0.2264 | 0.2572 | 0.0308 |
| 5nwh | opls | 2.6 | 0.091 | 2.3 | 0 | 97.36 | 1.81 | 0 | 1.13 | 1.14 | 0.2216 | 0.2475 | 0.0259 |
| 5nwh | noschrodinger | 2.6 | 0.0033 | 0.83 | 0.52 | 95.29 | 2.1 | 0 | 4.54 | 1.8 | 0.2209 | 0.2515 | 0.0306 |
| 5nx2 | opls | 3.7 | 0.089 | 2.33 | 0.26 | 96.9 | 1.73 | 3 | 1.47 | 1.25 | 0.3068 | 0.3484 | 0.0416 |
| 5nx2 | noschrodinger | 3.7 | 0.0036 | 0.86 | 0.26 | 96.91 | 0.86 | 0 | 7.65 | 1.6 | 0.3365 | 0.3657 | 0.0292 |
| 5nyc | opls | 1.51 | 0.0993 | 2.56 | 0 | 99.6 | 0.49 | 0 | 4.45 | 1.22 | 0.1444 | 0.167 | 0.0226 |
| 5nyc | noschrodinger | 1.51 | 0.0054 | 1.08 | 0 | 99.22 | 0.48 | 0 | 4.2 | 1.2 | 0.1477 | 0.1623 | 0.0146 |
| 5o00 | opls | 2.03 | 0.0956 | 2.23 | 0.93 | 97.2 | 0.53 | 1 | 1.86 | 1.09 | 0.1726 | 0.1932 | 0.0206 |
| 5o00 | noschrodinger | 2.03 | 0.003 | 0.79 | 0.93 | 96.76 | 1.04 | 0 | 1.6 | 1.12 | 0.1669 | 0.1931 | 0.0262 |
| 5o0t | opls | 2.05 | 0.0897 | 2.13 | 0 | 97.96 | 0.6 | 0 | 0 | 0.51 | 0.2052 | 0.2216 | 0.0164 |
| 5o0t | noschrodinger | 2.05 | 0.01 | 1.06 | 0 | 98.04 | 1.2 | 0 | 1.32 | 0.92 | 0.184 | 0.2139 | 0.0299 |
| 5o0x | opls | 2.2 | 0.0897 | 2.22 | 0 | 99.12 | 0.93 | 0 | 1.75 | 0.93 | 0.1886 | 0.2087 | 0.0201 |
| 5o0x | noschrodinger | 2.2 | 0.0025 | 0.82 | 0.39 | 98.05 | 0.46 | 0 | 1.12 | 0.82 | 0.1874 | 0.2013 | 0.0139 |
| 5o2d | opls | 1.6 | 0.0975 | 2.58 | 0 | 96 | 0 | 4 | 6.85 | 1.65 | 0.1554 | 0.194 | 0.0386 |
| 5o2d | noschrodinger | 1.6 | 0.004 | 0.92 | 0 | 96.79 | 0 | 0 | 1.63 | 1.11 | 0.1638 | 0.1879 | 0.0241 |
| 5o2f | opls | 2.01 | 0.1008 | 2.21 | 0 | 97.93 | 0.81 | 0 | 1.22 | 0.86 | 0.1667 | 0.193 | 0.0263 |
| 5o2f | noschrodinger | 2.01 | 0.003 | 0.82 | 0 | 98.34 | 0.81 | 0 | 1.77 | 0.93 | 0.1438 | 0.1805 | 0.0367 |
| 5o33 | opls | 1.64 | 0.0978 | 2.34 | 0.32 | 98.05 | 0.67 | 4 | 0.52 | 0.68 | 0.2243 | 0.2582 | 0.0339 |
| 5o33 | noschrodinger | 1.64 | 0.012 | 1.2 | 0.28 | 97.73 | 1.32 | 0 | 2.47 | 1.18 | 0.2244 | 0.2509 | 0.0265 |
| 5o3p | opls | 1.75 | 0.096 | 2.23 | 0 | 98.49 | 0.42 | 2 | 1.89 | 0.95 | 0.1695 | 0.1849 | 0.0154 |
| 5o3p | noschrodinger | 1.75 | 0.0039 | 0.86 | 0.35 | 97.89 | 1.25 | 0 | 1.48 | 0.99 | 0.1709 | 0.1826 | 0.0117 |
| 5o9j | opls | 2 | 0.0952 | 2.24 | 0 | 96.9 | 1.84 | 1 | 2.54 | 1.43 | 0.2236 | 0.2576 | 0.034 |
| 5o9j | noschrodinger | 2 | 0.0025 | 0.75 | 0.28 | 94.74 | 2.76 | 0 | 3.82 | 1.87 | 0.2098 | 0.2503 | 0.0405 |
| 5oec | opls | 2.3 | 0.0907 | 2.32 | 0.31 | 96.64 | 1.32 | 1 | 1.06 | 1.11 | 0.2382 | 0.2843 | 0.0461 |
| 5oec | noschrodinger | 2.3 | 0.0082 | 0.75 | 0.61 | 97.27 | 1.62 | 0 | 0.89 | 1.07 | 0.2298 | 0.2801 | 0.0503 |
| 5oed | opls | 2.9 | 0.0903 | 3.06 | 0.89 | 91.42 | 5.08 | 32 | 3.28 | 2.16 | 0.2393 | 0.2876 | 0.0483 |
| 5oed | noschrodinger | 2.9 | 0.0075 | 0.73 | 0.29 | 96.19 | 2.52 | 0 | 0.69 | 1.29 | 0.2567 | 0.2759 | 0.0192 |
| 5ofw | opls | 1.5 | 0.0969 | 2.42 | 0 | 98.18 | 3.3 | 3 | 4 | 1.58 | 0.1741 | 0.1893 | 0.0152 |
| 5ofw | noschrodinger | 1.5 | 0.0066 | 1.09 | 0 | 98.45 | 2.99 | 0 | 4.95 | 1.62 | 0.1751 | 0.1882 | 0.0131 |
| 5oh3 | opls | 2.1 | 0.0951 | 2.22 | 0 | 98.02 | 0.93 | 0 | 1.63 | 0.91 | 0.1981 | 0.2252 | 0.0271 |
| 5oh3 | noschrodinger | 2.1 | 0.0026 | 0.76 | 0 | 96.97 | 0.46 | 0 | 1.17 | 1.01 | 0.1854 | 0.2115 | 0.0261 |
| 5oha | opls | 1.55 | 0.0971 | 2.44 | 0 | 99.29 | 0.4 | 3 | 3.67 | 1.16 | 0.1688 | 0.1966 | 0.0278 |
| 5oha | noschrodinger | 1.55 | 0.0062 | 0.96 | 0 | 98.35 | 0 | 0 | 3.07 | 1.1 | 0.1722 | 0.1949 | 0.0227 |
| 5oly | opls | 2 | 0.1003 | 2.24 | 0 | 96.91 | 1.98 | 0 | 0 | 0.91 | 0.237 | 0.2725 | 0.0355 |
| 5oly | noschrodinger | 2 | 0.0033 | 0.82 | 0.24 | 96.94 | 1.96 | 0 | 1.74 | 1.33 | 0.2091 | 0.26 | 0.0509 |
| 5ovo | opls | 1.55 | 0.1042 | 2.55 | 0 | 97.94 | 3.15 | 3 | 4.17 | 1.59 | 0.1229 | 0.1422 | 0.0193 |

| | | | | | | | | | | | |
|---|---|---|---|---|---|---|---|---|---|---|---|
| 5ovo | noschrodinger | 1.55 | 0.0143 | 1.59 | 0 | 97.95 | 4.09 | 1 | 5.45 | 1.77 | 0.1233 | 0.1423 | 0.019 |
| 5ow0 | opls | 1.7 | 0.1044 | 2.3 | 0 | 98.63 | 0.23 | 4 | 2.23 | 1 | 0.1428 | 0.1798 | 0.037 |
| 5ow0 | noschrodinger | 1.7 | 0.0048 | 0.95 | 0 | 98.89 | 0 | 0 | 1.18 | 0.83 | 0.1452 | 0.1761 | 0.0309 |
| 5prn | opls | 2 | 0.095 | 2.5 | 0 | 97.2 | 2.59 | 5 | 0.89 | 1.23 | 0.1426 | 0.174 | 0.0314 |
| 5prn | noschrodinger | 2 | 0.0078 | 1.1 | 0 | 96.86 | 4.7 | 0 | 2 | 1.67 | 0.1461 | 0.1741 | 0.028 |
| 5qc9 | opls | 2 | 0.1034 | 2.13 | 0 | 98.55 | 2.56 | 0 | 1.03 | 1.11 | 0.132 | 0.1711 | 0.0391 |
| 5qc9 | noschrodinger | 2 | 0.003 | 0.82 | 0 | 97.94 | 3.4 | 0 | 0.3 | 1.03 | 0.1261 | 0.1713 | 0.0452 |
| 5rnt | opls | 3.19 | 0.0982 | 2.51 | 0 | 94.06 | 5.88 | 2 | 5.33 | 2.27 | 0.2334 | 0.2843 | 0.0509 |
| 5rnt | noschrodinger | 3.19 | 0.0035 | 0.94 | 2.94 | 87.25 | 20 | 0 | 11.35 | 3.18 | 0.194 | 0.2561 | 0.0621 |
| 5sux | opls | 2 | 0.0941 | 2.24 | 0 | 98.48 | 1.2 | 0 | 0.89 | 0.83 | 0.1758 | 0.2293 | 0.0535 |
| 5sux | noschrodinger | 2 | 0.0024 | 0.75 | 0 | 98.11 | 0.79 | 0 | 0.89 | 0.77 | 0.1842 | 0.2313 | 0.0471 |
| 5szh | opls | 2.3 | 0.0904 | 2.26 | 0 | 98.21 | 0.74 | 1 | 0.19 | 0.58 | 0.2142 | 0.2388 | 0.0246 |
| 5szh | noschrodinger | 2.3 | 0.0026 | 0.86 | 0 | 97.44 | 0.74 | 0 | 1.96 | 1.07 | 0.2021 | 0.2316 | 0.0295 |
| 5szj | opls | 2.66 | 0.0905 | 2.28 | 0 | 98.51 | 2.26 | 1 | 0.58 | 0.96 | 0.24 | 0.2649 | 0.0249 |
| 5szj | noschrodinger | 2.66 | 0.0026 | 0.83 | 0 | 96.82 | 2.99 | 0 | 3.12 | 1.66 | 0.255 | 0.2724 | 0.0174 |
| 5szm | opls | 3.6 | 0.0881 | 2.26 | 0 | 96.31 | 1.97 | 2 | 0.61 | 1.18 | 0.2291 | 0.2591 | 0.03 |
| 5szm | noschrodinger | 3.6 | 0.0031 | 0.76 | 0.48 | 94.44 | 0.84 | 0 | 3.69 | 1.54 | 0.2244 | 0.2554 | 0.031 |
| 5t3u | opls | 2.15 | 0.1018 | 2.27 | 0 | 99.23 | 1.71 | 4 | 1.67 | 1.09 | 0.1912 | 0.2271 | 0.0359 |
| 5t3u | noschrodinger | 2.15 | 0.0107 | 1.15 | 0 | 99.62 | 3.39 | 0 | 1.2 | 1.24 | 0.1815 | 0.2188 | 0.0373 |
| 5t53 | opls | 2.7 | 0.0843 | 2.5 | 0 | 98.06 | 4.58 | 2 | 0.35 | 1.13 | 0.2745 | 0.2916 | 0.0171 |
| 5t53 | noschrodinger | 2.7 | 0.0034 | 0.83 | 0 | 93.85 | 0.75 | 0 | 8.49 | 1.87 | 0.2705 | 0.2853 | 0.0148 |
| 5t54 | opls | 1.65 | 0.0989 | 2.41 | 0 | 98 | 0.25 | 2 | 1.09 | 0.81 | 0.1506 | 0.1849 | 0.0343 |
| 5t54 | noschrodinger | 1.65 | 0.0087 | 1.16 | 0 | 97.8 | 0.74 | 0 | 2.87 | 1.12 | 0.1515 | 0.1815 | 0.03 |
| 5t80 | opls | 2.62 | 0.0908 | 2.22 | 0.36 | 97.3 | 2.82 | 2 | 0.56 | 1.16 | 0.2397 | 0.2562 | 0.0165 |
| 5t80 | noschrodinger | 2.62 | 0.0029 | 0.79 | 0.51 | 95.41 | 5.2 | 0 | 2.9 | 1.94 | 0.2346 | 0.2491 | 0.0145 |
| 5t9e | opls | 2.03 | 0.0942 | 2.29 | 0 | 96.54 | 1.16 | 1 | 0.73 | 1.01 | 0.2086 | 0.2322 | 0.0236 |
| 5t9e | noschrodinger | 2.03 | 0.0043 | 0.84 | 0.2 | 97.18 | 2.85 | 0 | 3.05 | 1.59 | 0.1905 | 0.2206 | 0.0301 |
| 5t9t | opls | 3.5 | 0.0879 | 2.27 | 0.33 | 97.92 | 1.59 | 5 | 0.25 | 0.77 | 0.2256 | 0.2678 | 0.0422 |
| 5t9t | noschrodinger | 3.5 | 0.003 | 0.77 | 0.19 | 95.13 | 0.68 | 0 | 1.94 | 1.3 | 0.2214 | 0.2598 | 0.0384 |
| 5tcx | opls | 2.95 | 0.089 | 2.26 | 0 | 98.43 | 3.01 | 1 | 0.6 | 1.07 | 0.2575 | 0.3109 | 0.0534 |
| 5tcx | noschrodinger | 2.95 | 0.0029 | 0.96 | 0 | 94.92 | 2.37 | 0 | 2.13 | 1.62 | 0.2575 | 0.3168 | 0.0593 |
| 5tfz | opls | 2.2 | 0.0945 | 2.19 | 0 | 98.17 | 0 | 0 | 0.75 | 0.74 | 0.2003 | 0.2242 | 0.0239 |
| 5tfz | noschrodinger | 2.2 | 0.0027 | 0.77 | 0 | 98.43 | 0 | 0 | 1.26 | 0.85 | 0.1835 | 0.2228 | 0.0393 |
| 5tgs | opls | 2.55 | 0.0924 | 2.23 | 0 | 96.44 | 1.53 | 0 | 0.55 | 1.06 | 0.2115 | 0.2376 | 0.0261 |
| 5tgs | noschrodinger | 2.55 | 0.0026 | 0.73 | 0.22 | 95.14 | 1.26 | 0 | 1.1 | 1.23 | 0.2089 | 0.2386 | 0.0297 |
| 5tiz | opls | 2.87 | 0.0906 | 2.31 | 0.4 | 96.4 | 2.97 | 1 | 0.48 | 1.26 | 0.2439 | 0.2704 | 0.0265 |
| 5tiz | noschrodinger | 2.87 | 0.0047 | 0.85 | 0.4 | 96.41 | 5.06 | 0 | 1.92 | 1.73 | 0.2165 | 0.2615 | 0.045 |
| 5tk8 | opls | 1.64 | 0.0969 | 2.31 | 0 | 96.81 | 0.55 | 1 | 3.42 | 1.33 | 0.1673 | 0.2017 | 0.0344 |
| 5tk8 | noschrodinger | 1.64 | 0.0069 | 1 | 0 | 97.35 | 0.54 | 0 | 2.18 | 1.12 | 0.1688 | 0.1989 | 0.0301 |
| 5tph | opls | 2.47 | 0.0844 | 2.12 | 0 | 97.58 | 0 | 0 | 0.27 | 0.69 | 0.2364 | 0.2757 | 0.0393 |
| 5tph | noschrodinger | 2.47 | 0.0025 | 0.72 | 0 | 97.05 | 0 | 0 | 1.13 | 0.99 | 0.2193 | 0.2644 | 0.0451 |
| 5trv | opls | 2.91 | 0.0858 | 2.21 | 0 | 97.06 | 0 | 0 | 0.54 | 0.85 | 0.323 | 0.3357 | 0.0127 |
| 5trv | noschrodinger | 2.91 | 0.0024 | 0.69 | 0 | 99.14 | 0 | 0 | 1.67 | 0.92 | 0.2751 | 0.3246 | 0.0495 |
| 5tty | opls | 1.8 | 0.1008 | 2.36 | 0.68 | 96.59 | 2.97 | 2 | 2.65 | 1.63 | 0.1604 | 0.2033 | 0.0429 |
| 5tty | noschrodinger | 1.8 | 0.0044 | 0.87 | 0.34 | 96.95 | 2.22 | 0 | 1.22 | 1.28 | 0.1687 | 0.2059 | 0.0372 |
| 5tu4 | opls | 2.1 | 0.098 | 2.3 | 1.37 | 95.22 | 2.23 | 1 | 1.41 | 1.47 | 0.1925 | 0.2227 | 0.0302 |
| 5tu4 | noschrodinger | 2.1 | 0.0026 | 0.77 | 0.68 | 96.61 | 2.21 | 0 | 2.22 | 1.48 | 0.1808 | 0.2153 | 0.0345 |
| 5tuy | opls | 2.6 | 0.0919 | 2.3 | 0.19 | 95.38 | 1.06 | 2 | 0.7 | 1.07 | 0.2159 | 0.2433 | 0.0274 |
| 5tuy | noschrodinger | 2.6 | 0.0027 | 0.83 | 0.19 | 95.23 | 2.1 | 0 | 2.11 | 1.56 | 0.1912 | 0.229 | 0.0378 |

| | | | | | | | | | | | | | |
|---|---|---|---|---|---|---|---|---|---|---|---|---|---|
| 5tv8 | opls | 2.55 | 0.0937 | 2.28 | 0 | 98.09 | 3.35 | 1 | 0.5 | 1.07 | 0.2292 | 0.2497 | 0.0205 |
| 5tv8 | noschrodinger | 2.55 | 0.0028 | 0.76 | 0 | 97.68 | 4.52 | 0 | 1.89 | 1.52 | 0.224 | 0.2398 | 0.0158 |
| 5tvu | opls | 2.3 | 0.0905 | 2.29 | 0.52 | 95.49 | 0.77 | 2 | 0.57 | 1.01 | 0.2914 | 0.3396 | 0.0482 |
| 5tvu | noschrodinger | 2.3 | 0.0025 | 0.73 | 0.69 | 92.96 | 0.76 | 0 | 1.97 | 1.41 | 0.2928 | 0.3352 | 0.0424 |
| 5tya | opls | 1.5 | 0.1026 | 2.24 | 0 | 96.88 | 0.41 | 0 | 3.42 | 1.32 | 0.1404 | 0.1646 | 0.0242 |
| 5tya | noschrodinger | 1.5 | 0.0046 | 1.01 | 0 | 97.27 | 0.41 | 0 | 3.63 | 1.29 | 0.1427 | 0.1623 | 0.0196 |
| 5u1u | opls | 3.6 | 0.084 | 2.3 | 0 | 97.52 | 3.11 | 0 | 1.07 | 1.28 | 0.2218 | 0.2684 | 0.0466 |
| 5u1u | noschrodinger | 3.6 | 0.0027 | 0.8 | 0 | 95.83 | 1.33 | 0 | 4.89 | 1.64 | 0.2321 | 0.267 | 0.0349 |
| 5u32 | opls | 2.19 | 0.096 | 2.26 | 0 | 98.55 | 0.5 | 1 | 0.57 | 0.69 | 0.2088 | 0.2349 | 0.0261 |
| 5u32 | noschrodinger | 2.19 | 0.0039 | 0.83 | 0 | 98.09 | 1.48 | 0 | 3.14 | 1.23 | 0.1914 | 0.2292 | 0.0378 |
| 5u5i | opls | 2.2 | 0.0922 | 2.25 | 0.22 | 97.56 | 0.72 | 0 | 0.47 | 0.76 | 0.225 | 0.2401 | 0.0151 |
| 5u5i | noschrodinger | 2.2 | 0.0027 | 0.79 | 0.37 | 96.32 | 0.68 | 0 | 0.36 | 0.88 | 0.2163 | 0.2398 | 0.0235 |
| 5u75 | opls | 1.66 | 0.0992 | 2.39 | 0 | 98.36 | 0 | 0 | 2.42 | 1.02 | 0.177 | 0.1962 | 0.0192 |
| 5u75 | noschrodinger | 1.66 | 0.0111 | 1.23 | 0 | 98.59 | 0 | 0 | 4.88 | 1.25 | 0.1785 | 0.1961 | 0.0176 |
| 5u77 | opls | 2.16 | 0.0927 | 2.39 | 0 | 96.49 | 0 | 0 | 0 | 0.73 | 0.2559 | 0.2687 | 0.0128 |
| 5u77 | noschrodinger | 2.16 | 0.0028 | 0.8 | 0.87 | 98.26 | 3 | 0 | 3.15 | 1.47 | 0.2466 | 0.2695 | 0.0229 |
| 5u86 | opls | 1.62 | 0.1047 | 2.49 | 0 | 96.41 | 0.4 | 4 | 8.63 | 1.7 | 0.1254 | 0.1567 | 0.0313 |
| 5u86 | noschrodinger | 1.62 | 0.0049 | 1.21 | 0 | 97.38 | 0.4 | 0 | 4.11 | 1.32 | 0.1295 | 0.1528 | 0.0233 |
| 5u93 | opls | 2 | 0.0964 | 2.47 | 0 | 97.95 | 3.42 | 2 | 1.55 | 1.32 | 0.1787 | 0.2164 | 0.0377 |
| 5u93 | noschrodinger | 2 | 0.003 | 0.82 | 0 | 99.05 | 2.56 | 0 | 1.55 | 1.21 | 0.1898 | 0.2126 | 0.0228 |
| 5u9j | opls | 2.1 | 0.0943 | 2.27 | 0 | 97.4 | 1.17 | 2 | 2.45 | 1.2 | 0.2596 | 0.2876 | 0.028 |
| 5u9j | noschrodinger | 2.1 | 0.0027 | 0.79 | 0 | 95.67 | 1.54 | 0 | 3.3 | 1.57 | 0.2517 | 0.2846 | 0.0329 |
| 5u9n | opls | 2.4 | 0.089 | 2.2 | 0 | 100 | 0 | 0 | 0 | 0.5 | 0.3005 | 0.3153 | 0.0148 |
| 5u9n | noschrodinger | 2.4 | 0.0049 | 0.84 | 0 | 98.28 | 0 | 0 | 0.77 | 0.74 | 0.2698 | 0.2917 | 0.0219 |
| 5ubh | opls | 2 | 0.0931 | 2.27 | 0 | 98.02 | 1.15 | 0 | 0.15 | 0.6 | 0.2297 | 0.2523 | 0.0226 |
| 5ubh | noschrodinger | 2 | 0.0026 | 0.85 | 0 | 97.7 | 1.71 | 0 | 0.59 | 0.94 | 0.215 | 0.2407 | 0.0257 |
| 5uc1 | opls | 2.35 | 0.0858 | 2.19 | 0 | 97.77 | 0 | 0 | 0.14 | 0.61 | 0.213 | 0.2407 | 0.0277 |
| 5uc1 | noschrodinger | 2.35 | 0.0101 | 1.7 | 0.25 | 97.26 | 0.3 | 0 | 7.44 | 1.55 | 0.2025 | 0.2258 | 0.0233 |
| 5ucv | opls | 1.8 | 0.0988 | 2.27 | 0 | 99.04 | 1.28 | 2 | 2.31 | 1.09 | 0.166 | 0.2136 | 0.0476 |
| 5ucv | noschrodinger | 1.8 | 0.0045 | 0.82 | 0 | 98.63 | 1.27 | 0 | 1 | 0.87 | 0.1767 | 0.2071 | 0.0304 |
| 5ueh | opls | 2 | 0.0996 | 2.27 | 0 | 98.68 | 0.47 | 1 | 1.27 | 0.85 | 0.1685 | 0.2113 | 0.0428 |
| 5ueh | noschrodinger | 2 | 0.0084 | 0.97 | 0 | 99.16 | 0.47 | 0 | 1.52 | 0.89 | 0.1649 | 0.2017 | 0.0368 |
| 5ujv | opls | 2.46 | 0.0913 | 2.3 | 0 | 96.64 | 0 | 1 | 0.57 | 0.91 | 0.2673 | 0.3046 | 0.0373 |
| 5ujv | noschrodinger | 2.46 | 0.0027 | 0.91 | 0 | 97.18 | 0 | 0 | 1.14 | 0.97 | 0.2681 | 0.2998 | 0.0317 |
| 5uk5 | opls | 2.51 | 0.0897 | 2.29 | 0 | 95.37 | 0.25 | 2 | 1.08 | 1.13 | 0.2568 | 0.296 | 0.0392 |
| 5uk5 | noschrodinger | 2.51 | 0.0089 | 0.83 | 0.21 | 95.97 | 0.74 | 0 | 4.35 | 1.49 | 0.2512 | 0.2873 | 0.0361 |
| 5uk8 | opls | 2.5 | 0.0916 | 2.27 | 0 | 97.3 | 1.64 | 1 | 0.84 | 1.05 | 0.24 | 0.2669 | 0.0269 |
| 5uk8 | noschrodinger | 2.5 | 0.0029 | 0.79 | 0.16 | 95.62 | 2.54 | 0 | 1.04 | 1.42 | 0.2244 | 0.2494 | 0.025 |
| 5ula | opls | 1.5 | 0.1063 | 2.36 | 0 | 100 | 0.44 | 2 | 0.71 | 0.73 | 0.1626 | 0.1931 | 0.0305 |
| 5ula | noschrodinger | 1.5 | 0.0063 | 1.04 | 0 | 99.16 | 0.43 | 0 | 1.67 | 0.92 | 0.1664 | 0.1919 | 0.0255 |
| 5umx | opls | 1.59 | 0.1007 | 2.27 | 0 | 97.46 | 0.99 | 0 | 1.32 | 0.97 | 0.1534 | 0.1814 | 0.028 |
| 5umx | noschrodinger | 1.59 | 0.0069 | 1.01 | 0 | 97.9 | 0.98 | 0 | 1.06 | 0.83 | 0.1569 | 0.1783 | 0.0214 |
| 5uni | opls | 2.2 | 0.0899 | 2.23 | 0 | 98.53 | 1.56 | 1 | 0.52 | 0.82 | 0.2021 | 0.2323 | 0.0302 |
| 5uni | noschrodinger | 2.2 | 0.003 | 0.75 | 0 | 96.87 | 1.53 | 0 | 0.87 | 1.1 | 0.1939 | 0.2295 | 0.0356 |
| 5uos | opls | 2.51 | 0.0917 | 2.68 | 0 | 99.57 | 0.5 | 0 | 1.71 | 0.92 | 0.1878 | 0.2254 | 0.0376 |
| 5uos | noschrodinger | 2.51 | 0.0041 | 1.26 | 0 | 97.05 | 1.47 | 0 | 4.4 | 1.51 | 0.1842 | 0.2163 | 0.0321 |
| 5uqi | opls | 1.62 | 0.0972 | 2.45 | 0 | 98.91 | 0 | 1 | 2.09 | 0.98 | 0.155 | 0.1804 | 0.0254 |
| 5uqi | noschrodinger | 1.62 | 0.0111 | 1.22 | 0 | 99.46 | 0.63 | 0 | 0.7 | 0.73 | 0.1579 | 0.1797 | 0.0218 |
| 5uqw | opls | 1.5 | 0.0993 | 2.43 | 0 | 98.68 | 0.35 | 2 | 4.99 | 1.26 | 0.1069 | 0.1321 | 0.0252 |

| | | | | | | | | | | | | | |
|---|---|---|---|---|---|---|---|---|---|---|---|---|---|
| 5uqw | noschrodinger | 1.5 | 0.0108 | 1.28 | 0 | 98.11 | 0.7 | 0 | 4.43 | 1.22 | 0.1068 | 0.126 | 0.0192 |
| 5ur3 | opls | 1.76 | 0.0953 | 2.65 | 0 | 98.28 | 2.68 | 8 | 4.07 | 1.52 | 0.1782 | 0.2229 | 0.0447 |
| 5ur3 | noschrodinger | 1.76 | 0.0053 | 1.06 | 0 | 97.33 | 3.53 | 0 | 4.43 | 1.77 | 0.1888 | 0.2185 | 0.0297 |
| 5ut3 | opls | 1.5 | 0.1021 | 2.5 | 0 | 98.64 | 0 | 1 | 1.41 | 0.87 | 0.1492 | 0.1779 | 0.0287 |
| 5ut3 | noschrodinger | 1.5 | 0.0132 | 1.3 | 0 | 98.89 | 0 | 0 | 0.95 | 0.78 | 0.1521 | 0.1774 | 0.0253 |
| 5ute | opls | 2.05 | 0.099 | 2.33 | 0 | 98.45 | 4.5 | 1 | 1.89 | 1.45 | 0.203 | 0.2345 | 0.0315 |
| 5ute | noschrodinger | 2.05 | 0.0075 | 1.07 | 0 | 97.82 | 7.03 | 0 | 5.56 | 1.99 | 0.1785 | 0.2192 | 0.0407 |
| 5uwa | opls | 1.5 | 0.0984 | 2.42 | 0 | 98.63 | 0.3 | 2 | 3.97 | 1.18 | 0.1818 | 0.2022 | 0.0204 |
| 5uwa | noschrodinger | 1.5 | 0.0071 | 1.01 | 0 | 98.91 | 0.3 | 0 | 3.06 | 1.1 | 0.1894 | 0.2024 | 0.013 |
| 5uwb | opls | 2.6 | 0.0867 | 2.19 | 0 | 97.93 | 1 | 0 | 0.8 | 0.77 | 0.2314 | 0.2725 | 0.0411 |
| 5uwb | noschrodinger | 2.6 | 0.0025 | 0.66 | 0 | 97.03 | 1.28 | 0 | 0.48 | 0.92 | 0.2212 | 0.2629 | 0.0417 |
| 5uwc | opls | 2.4 | 0.0915 | 2.33 | 1.14 | 94.32 | 2.48 | 5 | 1.32 | 1.55 | 0.2205 | 0.2429 | 0.0224 |
| 5uwc | noschrodinger | 2.4 | 0.0068 | 0.91 | 0.84 | 93.56 | 4.29 | 0 | 5.46 | 2.2 | 0.2208 | 0.2399 | 0.0191 |
| 5uxc | opls | 1.72 | 0.1041 | 2.38 | 0 | 98.57 | 0.45 | 1 | 1.85 | 0.95 | 0.1601 | 0.1985 | 0.0384 |
| 5uxc | noschrodinger | 1.72 | 0.0036 | 1.51 | 0 | 98.64 | 0.88 | 0 | 1.85 | 0.95 | 0.1651 | 0.1943 | 0.0292 |
| 5uxi | opls | 2 | 0.0957 | 2.27 | 0 | 99.08 | 0 | 1 | 0 | 0.5 | 0.2122 | 0.2462 | 0.034 |
| 5uxi | noschrodinger | 2 | 0.0029 | 0.74 | 0 | 99.08 | 0.54 | 0 | 1.66 | 0.92 | 0.1993 | 0.2378 | 0.0385 |
| 5uxy | opls | 1.8 | 0.1035 | 2.36 | 0 | 97.98 | 0.91 | 4 | 2.05 | 0.98 | 0.1482 | 0.1864 | 0.0382 |
| 5uxy | noschrodinger | 1.8 | 0.0083 | 0.93 | 0 | 98.18 | 0.43 | 0 | 2.5 | 1.03 | 0.1492 | 0.1815 | 0.0323 |
| 5v35 | opls | 2.5 | 0.0873 | 2.26 | 0 | 93.86 | 0.82 | 0 | 0 | 0.91 | 0.262 | 0.2961 | 0.0341 |
| 5v35 | noschrodinger | 2.5 | 0.0053 | 0.93 | 0 | 93.33 | 1.63 | 0 | 1.73 | 1.52 | 0.2294 | 0.2885 | 0.0591 |
| 5v3w | opls | 1.72 | 0.0998 | 2.37 | 0 | 98.6 | 0 | 2 | 1.9 | 0.95 | 0.1625 | 0.1975 | 0.035 |
| 5v3w | noschrodinger | 1.72 | 0.0045 | 0.94 | 0 | 98.34 | 1.18 | 0 | 1.56 | 0.96 | 0.1686 | 0.1945 | 0.0259 |
| 5v6i | opls | 1.7 | 0.0983 | 2.32 | 0 | 95.41 | 1.58 | 2 | 1.45 | 1.35 | 0.1523 | 0.1859 | 0.0336 |
| 5v6i | noschrodinger | 1.7 | 0.0047 | 0.99 | 0 | 94.5 | 1.05 | 0 | 2.9 | 1.47 | 0.1574 | 0.183 | 0.0256 |
| 5v87 | opls | 1.69 | 0.0975 | 2.24 | 0 | 98.86 | 0 | 0 | 1.56 | 0.9 | 0.1872 | 0.2235 | 0.0363 |
| 5v87 | noschrodinger | 1.69 | 0.0064 | 0.92 | 0 | 98.95 | 0 | 0 | 1.18 | 0.83 | 0.1899 | 0.2253 | 0.0354 |
| 5v8c | opls | 2.51 | 0.0891 | 2.35 | 0 | 96.98 | 5.8 | 1 | 0.75 | 1.49 | 0.2568 | 0.3047 | 0.0479 |
| 5v8c | noschrodinger | 2.51 | 0.003 | 0.8 | 1.49 | 95.15 | 11.11 | 0 | 1.51 | 2.02 | 0.2407 | 0.2908 | 0.0501 |
| 5v8t | opls | 2.1 | 0.097 | 2.18 | 0.32 | 99.05 | 0 | 1 | 0.68 | 0.72 | 0.1921 | 0.2218 | 0.0297 |
| 5v8t | noschrodinger | 2.1 | 0.0027 | 0.77 | 0 | 99.21 | 0.7 | 0 | 1.39 | 0.87 | 0.1731 | 0.216 | 0.0429 |
| 5veg | opls | 1.99 | 0.094 | 2.26 | 0 | 97.48 | 0.82 | 0 | 0.73 | 0.84 | 0.2023 | 0.236 | 0.0337 |
| 5veg | noschrodinger | 1.99 | 0.0036 | 0.8 | 0 | 96.13 | 0.27 | 0 | 3.67 | 1.42 | 0.2018 | 0.231 | 0.0292 |
| 5vj7 | opls | 2.55 | 0.0932 | 2.31 | 0.14 | 95.08 | 0.65 | 4 | 1 | 1.14 | 0.2243 | 0.267 | 0.0427 |
| 5vj7 | noschrodinger | 2.55 | 0.0034 | 0.82 | 0.14 | 94.7 | 0.65 | 0 | 2.09 | 1.35 | 0.2079 | 0.2528 | 0.0449 |
| 5vt7 | opls | 2.62 | 0.0923 | 2.3 | 0.21 | 97.44 | 1.22 | 1 | 0.92 | 0.95 | 0.1995 | 0.2386 | 0.0391 |
| 5vt7 | noschrodinger | 2.62 | 0.0028 | 0.78 | 0 | 97.46 | 2.65 | 0 | 1.45 | 1.31 | 0.1858 | 0.2268 | 0.041 |
| 5vx6 | opls | 3.2 | 0.0864 | 2.38 | 0.33 | 95.44 | 3.05 | 3 | 0.89 | 1.46 | 0.2567 | 0.2953 | 0.0386 |
| 5vx6 | noschrodinger | 3.2 | 0.0043 | 0.96 | 1.22 | 88.54 | 3.03 | 0 | 4.69 | 2.19 | 0.2209 | 0.2762 | 0.0553 |
| 5vyr | opls | 1.7 | 0.099 | 2.4 | 0 | 97.92 | 1.92 | 3 | 3.91 | 1.41 | 0.1842 | 0.217 | 0.0328 |
| 5vyr | noschrodinger | 1.7 | 0.0066 | 0.97 | 0 | 97.93 | 2.39 | 0 | 3.28 | 1.42 | 0.1861 | 0.2124 | 0.0263 |
| 5vys | opls | 2.2 | 0.0963 | 2.27 | 0 | 97.94 | 1.46 | 0 | 0.38 | 0.78 | 0.2037 | 0.2391 | 0.0354 |
| 5vys | noschrodinger | 2.2 | 0.0029 | 0.85 | 0 | 97.75 | 1.93 | 0 | 1.76 | 1.21 | 0.1994 | 0.2262 | 0.0268 |
| 5vyv | opls | 2.48 | 0.0946 | 2.29 | 0 | 98.8 | 0.66 | 1 | 0.9 | 0.77 | 0.2323 | 0.272 | 0.0397 |
| 5vyv | noschrodinger | 2.48 | 0.0027 | 0.72 | 0 | 96.14 | 2.29 | 0 | 2.33 | 1.55 | 0.2084 | 0.265 | 0.0566 |
| 5w37 | opls | 2.52 | 0.0901 | 2.19 | 0 | 96.71 | 2.82 | 0 | 0.74 | 1.28 | 0.2516 | 0.2739 | 0.0223 |
| 5w37 | noschrodinger | 2.52 | 0.0038 | 0.77 | 0 | 92.31 | 5.59 | 0 | 1.12 | 1.86 | 0.26 | 0.2719 | 0.0119 |
| 5w5v | opls | 3.65 | 0.0906 | 2.28 | 0.35 | 97.4 | 1.31 | 1 | 0.51 | 0.88 | 0.2604 | 0.2973 | 0.0369 |
| 5w5v | noschrodinger | 3.65 | 0.0025 | 0.7 | 0 | 97.95 | 2.03 | 0 | 1.64 | 1.16 | 0.2714 | 0.2996 | 0.0282 |

| | | | | | | | | | | | |
|---|---|---|---|---|---|---|---|---|---|---|---|
| 5w78 | opls | 2.27 | 0.093 | 2.24 | 0.19 | 97.09 | 0.21 | 1 | 0.69 | 0.89 | 0.1753 | 0.2126 | 0.0373 |
| 5w78 | noschrodinger | 2.27 | 0.0032 | 1.22 | 0.39 | 96.72 | 0.21 | 0 | 2.54 | 1.24 | 0.1778 | 0.2078 | 0.03 |
| 5w7l | opls | 2.74 | 0.0907 | 2.28 | 0 | 96.43 | 0 | 0 | 0.31 | 0.85 | 0.2676 | 0.3055 | 0.0379 |
| 5w7l | noschrodinger | 2.74 | 0.0023 | 0.71 | 0 | 96.99 | 0 | 0 | 1.88 | 1.12 | 0.2997 | 0.3175 | 0.0178 |
| 5w8a | opls | 2 | 0.1021 | 2.31 | 0 | 97.58 | 2.21 | 0 | 2.26 | 1.35 | 0.1781 | 0.1928 | 0.0147 |
| 5w8a | noschrodinger | 2 | 0.0026 | 0.84 | 0 | 98.09 | 1.65 | 0 | 1.13 | 0.99 | 0.1673 | 0.1842 | 0.0169 |
| 5w8e | opls | 1.8 | 0.1008 | 2.49 | 0 | 98.09 | 1.08 | 6 | 3.08 | 1.12 | 0.1698 | 0.1888 | 0.019 |
| 5w8e | noschrodinger | 1.8 | 0.01 | 1.13 | 0 | 98.58 | 1.6 | 0 | 1.4 | 1.03 | 0.1715 | 0.1913 | 0.0198 |
| 5wb5 | opls | 2.7 | 0.0904 | 2.21 | 0 | 98.9 | 0 | 0 | 0.6 | 0.7 | 0.2858 | 0.3152 | 0.0294 |
| 5wb5 | noschrodinger | 2.7 | 0.0028 | 0.71 | 0 | 96.95 | 0 | 0 | 2.42 | 1.2 | 0.2578 | 0.2987 | 0.0409 |
| 5wep | opls | 3.51 | 0.0857 | 2.26 | 0 | 92.86 | 3.28 | 1 | 0.5 | 1.52 | 0.2451 | 0.2761 | 0.031 |
| 5wep | noschrodinger | 3.51 | 0.0027 | 0.85 | 1.13 | 88.68 | 2.15 | 0 | 2.82 | 1.91 | 0.242 | 0.2739 | 0.0319 |
| 5whx | opls | 1.69 | 0.1011 | 2.33 | 0 | 97.38 | 1.29 | 3 | 1.67 | 1.12 | 0.1484 | 0.1754 | 0.027 |
| 5whx | noschrodinger | 1.69 | 0.0066 | 0.92 | 0 | 97.19 | 2.14 | 0 | 1.55 | 1.3 | 0.1521 | 0.1761 | 0.024 |
| 5wif | opls | 2.51 | 0.0915 | 2.24 | 0 | 98.37 | 0.67 | 1 | 0.23 | 0.59 | 0.2169 | 0.2437 | 0.0268 |
| 5wif | noschrodinger | 2.51 | 0.0028 | 0.76 | 0 | 96.39 | 2.18 | 0 | 1.63 | 1.41 | 0.2134 | 0.2414 | 0.028 |
| 5wje | opls | 1.77 | 0.1 | 2.4 | 0 | 97.9 | 1.41 | 1 | 3 | 1.23 | 0.147 | 0.1799 | 0.0329 |
| 5wje | noschrodinger | 1.77 | 0.0062 | 1.07 | 0 | 97.48 | 1.41 | 0 | 3.02 | 1.31 | 0.1503 | 0.1803 | 0.03 |
| 5wlq | opls | 3.1 | 0.0905 | 2.27 | 0 | 100 | 0 | 1 | 0 | 0.5 | 0.2543 | 0.2722 | 0.0179 |
| 5wlq | noschrodinger | 3.1 | 0.0027 | 0.69 | 0 | 99.02 | 2.06 | 0 | 0 | 0.74 | 0.2329 | 0.2615 | 0.0286 |
| 5wqk | opls | 1.7 | 0.0978 | 2.21 | 0 | 98.24 | 0.4 | 0 | 0.63 | 0.71 | 0.1768 | 0.2067 | 0.0299 |
| 5wqk | noschrodinger | 1.7 | 0.0035 | 0.87 | 0 | 98.25 | 1.19 | 0 | 1.69 | 0.98 | 0.1764 | 0.2057 | 0.0293 |
| 5wsh | opls | 2 | 0.1033 | 2.22 | 0 | 98.66 | 1.81 | 0 | 2.1 | 1.18 | 0.1784 | 0.2132 | 0.0348 |
| 5wsh | noschrodinger | 2 | 0.0026 | 0.75 | 0 | 98.15 | 1.79 | 0 | 1.94 | 1.15 | 0.164 | 0.2036 | 0.0396 |
| 5wt9 | opls | 2.4 | 0.0961 | 2.22 | 0 | 98.06 | 2.43 | 4 | 0.37 | 0.93 | 0.2036 | 0.2233 | 0.0197 |
| 5wt9 | noschrodinger | 2.4 | 0.0039 | 0.89 | 0.19 | 96.94 | 4.78 | 0 | 2.09 | 1.68 | 0.1915 | 0.2252 | 0.0337 |
| 5wts | opls | 3 | 0.0905 | 2.27 | 0 | 97.71 | 2.06 | 0 | 0.84 | 1.06 | 0.2308 | 0.2756 | 0.0448 |
| 5wts | noschrodinger | 3 | 0.0031 | 0.75 | 0 | 93.85 | 2.56 | 0 | 3.79 | 1.89 | 0.2278 | 0.2732 | 0.0454 |
| 5wy6 | opls | 1.78 | 0.097 | 2.27 | 0 | 97.79 | 0 | 1 | 1.57 | 0.95 | 0.1884 | 0.2218 | 0.0334 |
| 5wy6 | noschrodinger | 1.78 | 0.0035 | 0.84 | 0 | 97.8 | 0.83 | 0 | 1.12 | 0.87 | 0.1948 | 0.2232 | 0.0284 |
| 5wz2 | opls | 2.6 | 0.0913 | 2.31 | 0.13 | 98.31 | 1.71 | 1 | 1.36 | 1.04 | 0.209 | 0.2517 | 0.0427 |
| 5wz2 | noschrodinger | 2.6 | 0.0026 | 0.78 | 0.52 | 95.85 | 2.47 | 0 | 2.08 | 1.56 | 0.1958 | 0.2433 | 0.0475 |
| 5x0e | opls | 2 | 0.0952 | 2.18 | 0 | 97.01 | 0 | 0 | 1.51 | 1.06 | 0.1656 | 0.2041 | 0.0385 |
| 5x0e | noschrodinger | 2 | 0.0032 | 0.82 | 0 | 97.49 | 0 | 0 | 0.25 | 0.7 | 0.1729 | 0.2028 | 0.0299 |
| 5x0k | opls | 1.65 | 0.089 | 2.17 | 0 | 97.1 | 0 | 1 | 1.36 | 1.03 | 0.2358 | 0.2746 | 0.0388 |
| 5x0k | noschrodinger | 1.65 | 0.003 | 0.82 | 0 | 97.48 | 0 | 0 | 1.13 | 0.93 | 0.2459 | 0.2763 | 0.0304 |
| 5x13 | opls | 1.7 | 0.0998 | 2.21 | 0 | 98.12 | 0.62 | 1 | 0.99 | 0.79 | 0.1626 | 0.1864 | 0.0238 |
| 5x13 | noschrodinger | 1.7 | 0.0032 | 0.81 | 0 | 98.27 | 1.84 | 0 | 1.32 | 1.06 | 0.1666 | 0.1827 | 0.0161 |
| 5x33 | opls | 3.7 | 0.0898 | 2.36 | 0 | 96.88 | 9.06 | 2 | 1.46 | 1.8 | 0.2647 | 0.3126 | 0.0479 |
| 5x33 | noschrodinger | 3.7 | 0.0028 | 0.8 | 0 | 95.86 | 18.42 | 0 | 5.6 | 2.55 | 0.2717 | 0.313 | 0.0413 |
| 5x3d | opls | 1.93 | 0.0967 | 2.28 | 0 | 99.19 | 1.89 | 2 | 1.47 | 1.09 | 0.1905 | 0.2124 | 0.0219 |
| 5x3d | noschrodinger | 1.93 | 0.0029 | 0.86 | 0 | 99.19 | 3.74 | 0 | 1.47 | 1.32 | 0.195 | 0.2137 | 0.0187 |
| 5x4b | opls | 1.5 | 0.0977 | 2.25 | 0 | 99.17 | 0.45 | 0 | 0.44 | 0.66 | 0.1536 | 0.1753 | 0.0217 |
| 5x4b | noschrodinger | 1.5 | 0.0049 | 1.04 | 0.35 | 97.92 | 1.79 | 0 | 1.34 | 1.07 | 0.1568 | 0.1738 | 0.017 |
| 5x73 | opls | 2.5 | 0.0911 | 2.18 | 0 | 97.87 | 0.75 | 0 | 1.19 | 0.86 | 0.2326 | 0.262 | 0.0294 |
| 5x73 | noschrodinger | 2.5 | 0.0088 | 1 | 0 | 97.22 | 1.48 | 0 | 1.98 | 1.24 | 0.214 | 0.2541 | 0.0401 |
| 5x7z | opls | 2.2 | 0.0947 | 2.38 | 0 | 99.19 | 5.1 | 2 | 1.48 | 1.42 | 0.2194 | 0.2484 | 0.029 |
| 5x7z | noschrodinger | 2.2 | 0.0032 | 0.82 | 0 | 97.62 | 3 | 0 | 2.97 | 1.53 | 0.2277 | 0.2493 | 0.0216 |
| 5x9h | opls | 3.6 | 0.0888 | 2.37 | 0.14 | 97.16 | 5.36 | 5 | 0.68 | 1.43 | 0.255 | 0.2989 | 0.0439 |

| | | | | | | | | | | | | | |
|---|---|---|---|---|---|---|---|---|---|---|---|---|---|
| 5x9h | noschrodinger | 3.6 | 0.003 | 0.82 | 0.35 | 94.35 | 13.22 | 0 | 4.84 | 2.49 | 0.247 | 0.2907 | 0.0437 |
| 5x9r | opls | 3.98 | 0.0913 | 2.33 | 0 | 95.69 | 0.31 | 0 | 0.55 | 0.98 | 0.2476 | 0.2732 | 0.0256 |
| 5x9r | noschrodinger | 3.98 | 0.0031 | 0.75 | 0.49 | 92.04 | 0.16 | 0 | 3.83 | 1.66 | 0.24 | 0.2722 | 0.0322 |
| 5xaw | opls | 2.1 | 0.0895 | 2.3 | 0 | 97.06 | 0.51 | 0 | 1.6 | 1.07 | 0.2248 | 0.2456 | 0.0208 |
| 5xaw | noschrodinger | 2.1 | 0.0028 | 0.72 | 0 | 95.71 | 0 | 0 | 4.3 | 1.51 | 0.204 | 0.2398 | 0.0358 |
| 5xbf | opls | 1.8 | 0.0941 | 2.25 | 0 | 98.36 | 1.03 | 0 | 1.83 | 0.95 | 0.202 | 0.2159 | 0.0139 |
| 5xbf | noschrodinger | 1.8 | 0.0029 | 0.82 | 0 | 98.08 | 1.43 | 0 | 1.64 | 1.03 | 0.201 | 0.2142 | 0.0132 |
| 5xbt | opls | 2.49 | 0.093 | 2.41 | 0 | 97.76 | 3.07 | 2 | 1.09 | 1.24 | 0.2313 | 0.2537 | 0.0224 |
| 5xbt | noschrodinger | 2.49 | 0.0031 | 0.84 | 0.37 | 96.1 | 6.55 | 0 | 5.44 | 2.18 | 0.2159 | 0.242 | 0.0261 |
| 5xe8 | opls | 2.8 | 0.0907 | 2.33 | 1.61 | 93.55 | 3.2 | 2 | 0.96 | 1.59 | 0.2774 | 0.3012 | 0.0238 |
| 5xe8 | noschrodinger | 2.8 | 0.0054 | 0.96 | 1.2 | 89.24 | 8.56 | 0 | 12.06 | 2.87 | 0.2806 | 0.3065 | 0.0259 |
| 5xfd | opls | 1.5 | 0.0972 | 2.42 | 0 | 97.16 | 1.5 | 3 | 2.58 | 1.33 | 0.1409 | 0.1595 | 0.0186 |
| 5xfd | noschrodinger | 1.5 | 0.0131 | 1.44 | 0 | 97.69 | 1.1 | 0 | 6.17 | 1.44 | 0.1435 | 0.1589 | 0.0154 |
| 5xg5 | opls | 1.54 | 0.0987 | 2.26 | 0 | 100 | 0 | 4 | 0.84 | 0.76 | 0.146 | 0.1859 | 0.0399 |
| 5xg5 | noschrodinger | 1.54 | 0.0031 | 0.91 | 0 | 100 | 0 | 0 | 0.84 | 0.76 | 0.1512 | 0.1806 | 0.0294 |
| 5xga | opls | 1.95 | 0.0932 | 2.4 | 0 | 96.52 | 3.81 | 0 | 3.28 | 1.79 | 0.1899 | 0.2329 | 0.043 |
| 5xga | noschrodinger | 1.95 | 0.0135 | 1.94 | 0 | 96.55 | 2.83 | 0 | 10.34 | 2.1 | 0.19 | 0.2304 | 0.0404 |
| 5xj6 | opls | 2.37 | 0.0897 | 2.25 | 0 | 97.98 | 1.23 | 1 | 0.57 | 0.77 | 0.231 | 0.2735 | 0.0425 |
| 5xj6 | noschrodinger | 2.37 | 0.0029 | 0.75 | 0 | 97.47 | 0.61 | 0 | 2.29 | 1.11 | 0.2139 | 0.2614 | 0.0475 |
| 5xj7 | opls | 1.77 | 0.0889 | 2.31 | 0 | 99.47 | 0.64 | 1 | 4.4 | 1.22 | 0.1806 | 0.2117 | 0.0311 |
| 5xj7 | noschrodinger | 1.77 | 0.0056 | 0.9 | 0 | 99.47 | 0 | 0 | 3.02 | 1.09 | 0.1828 | 0.2079 | 0.0251 |
| 5xkt | opls | 1.8 | 0.104 | 2.4 | 0.25 | 97.97 | 0 | 3 | 2.47 | 1.04 | 0.1513 | 0.1856 | 0.0343 |
| 5xkt | noschrodinger | 1.8 | 0.0033 | 1.07 | 0 | 98.22 | 0 | 0 | 1.32 | 0.86 | 0.1601 | 0.1833 | 0.0232 |
| 5xpd | opls | 2.79 | 0.0911 | 2.3 | 0 | 98.46 | 1.32 | 2 | 0.46 | 0.75 | 0.2419 | 0.2688 | 0.0269 |
| 5xpd | noschrodinger | 2.79 | 0.0027 | 0.76 | 0 | 97.38 | 1.31 | 0 | 2.09 | 1.19 | 0.2348 | 0.2565 | 0.0217 |
| 5xpi | opls | 2.2 | 0.0985 | 2.19 | 0 | 96.99 | 3.25 | 0 | 0.85 | 1.32 | 0.197 | 0.2342 | 0.0372 |
| 5xpi | noschrodinger | 2.2 | 0.0028 | 0.76 | 0 | 94.85 | 7.14 | 0 | 0.43 | 1.66 | 0.1913 | 0.2345 | 0.0432 |
| 5xsj | opls | 2.2 | 0.0985 | 2.27 | 0 | 96.89 | 1.76 | 6 | 0.81 | 1.13 | 0.1819 | 0.2131 | 0.0312 |
| 5xsj | noschrodinger | 2.2 | 0.0045 | 0.77 | 0.26 | 96.4 | 1.75 | 0 | 1.62 | 1.33 | 0.1893 | 0.2141 | 0.0248 |
| 5xtk | opls | 1.74 | 0.097 | 2.29 | 0 | 98.57 | 1.57 | 1 | 2.85 | 1.22 | 0.1639 | 0.1851 | 0.0212 |
| 5xtk | noschrodinger | 1.74 | 0.0047 | 0.99 | 0 | 98.65 | 1.97 | 0 | 2.04 | 1.2 | 0.1694 | 0.1849 | 0.0155 |
| 5xuh | opls | 2.02 | 0.1024 | 2.29 | 0 | 99.16 | 0.35 | 0 | 0.66 | 0.72 | 0.1873 | 0.2142 | 0.0269 |
| 5xuh | noschrodinger | 2.02 | 0.0028 | 0.79 | 0 | 99.17 | 1.39 | 0 | 1.81 | 1.05 | 0.1658 | 0.2039 | 0.0381 |
| 5xun | opls | 2 | 0.096 | 2.28 | 0.63 | 97.8 | 1.44 | 0 | 0.18 | 0.74 | 0.2054 | 0.2452 | 0.0398 |
| 5xun | noschrodinger | 2 | 0.0033 | 0.79 | 0.59 | 97.92 | 1.08 | 0 | 2 | 1.01 | 0.209 | 0.2394 | 0.0304 |
| 5xv2 | opls | 2 | 0.0937 | 2.26 | 0 | 96.36 | 1.68 | 0 | 0.85 | 1.18 | 0.2451 | 0.275 | 0.0299 |
| 5xv2 | noschrodinger | 2 | 0.0027 | 0.77 | 0 | 97.74 | 1.11 | 0 | 1.99 | 1.06 | 0.2311 | 0.2636 | 0.0325 |
| 5xvq | opls | 2.29 | 0.0943 | 2.3 | 0 | 98.77 | 2.88 | 3 | 0.96 | 1.14 | 0.1943 | 0.2342 | 0.0399 |
| 5xvq | noschrodinger | 2.29 | 0.0028 | 0.83 | 0 | 98.79 | 5.05 | 0 | 0.96 | 1.32 | 0.1856 | 0.2274 | 0.0418 |
| 5xw8 | opls | 2 | 0.1031 | 2.29 | 0 | 99.07 | 0.55 | 1 | 1.22 | 0.84 | 0.1531 | 0.178 | 0.0249 |
| 5xw8 | noschrodinger | 2 | 0.0049 | 0.94 | 0 | 96.79 | 0 | 0 | 0.61 | 0.9 | 0.1391 | 0.1705 | 0.0314 |
| 5xxr | opls | 2.65 | 0.0853 | 2.35 | 0 | 97.95 | 1.11 | 0 | 0.52 | 0.73 | 0.2417 | 0.2869 | 0.0452 |
| 5xxr | noschrodinger | 2.65 | 0.0025 | 0.72 | 0 | 95.69 | 2.63 | 0 | 1.6 | 1.53 | 0.2272 | 0.2842 | 0.057 |
| 5xxs | opls | 2.09 | 0.0907 | 2.22 | 0 | 97.86 | 0.51 | 0 | 0.25 | 0.63 | 0.2201 | 0.2554 | 0.0353 |
| 5xxs | noschrodinger | 2.09 | 0.0026 | 0.77 | 0 | 96.19 | 1 | 0 | 1.78 | 1.19 | 0.2034 | 0.2421 | 0.0387 |
| 5y65 | opls | 2.55 | 0.0917 | 2.26 | 0 | 98.06 | 1.5 | 0 | 0.52 | 0.81 | 0.2081 | 0.2407 | 0.0326 |
| 5y65 | noschrodinger | 2.55 | 0.0028 | 0.8 | 0 | 98.28 | 2.99 | 0 | 1.31 | 1.22 | 0.1943 | 0.2256 | 0.0313 |
| 5y6f | opls | 2.2 | 0.0914 | 2.47 | 0 | 96.26 | 7.25 | 0 | 1.56 | 1.81 | 0.2047 | 0.2399 | 0.0352 |
| 5y6f | noschrodinger | 2.2 | 0.0029 | 0.84 | 0.92 | 94.95 | 9.64 | 0 | 3.14 | 2.2 | 0.2172 | 0.2459 | 0.0287 |

| | | | | | | | | | | | | | |
|---|---|---|---|---|---|---|---|---|---|---|---|---|---|
| 5y6g | opls | 2.3 | 0.0931 | 2.39 | 0 | 97.66 | 3.51 | 0 | 1.3 | 1.34 | 0.2208 | 0.2397 | 0.0189 |
| 5y6g | noschrodinger | 2.3 | 0.0025 | 0.75 | 0.78 | 96.12 | 3.48 | 0 | 3.92 | 1.85 | 0.2192 | 0.2418 | 0.0226 |
| 5y6q | opls | 2.5 | 0.0954 | 2.29 | 0.41 | 96.49 | 1.13 | 2 | 0.85 | 1.03 | 0.1792 | 0.2213 | 0.0421 |
| 5y6q | noschrodinger | 2.5 | 0.0032 | 0.84 | 0.16 | 97.4 | 1.44 | 0 | 2.54 | 1.28 | 0.1715 | 0.2133 | 0.0418 |
| 5y8t | opls | 1.99 | 0.0969 | 2.21 | 0 | 99.11 | 0.23 | 2 | 0.61 | 0.7 | 0.1919 | 0.221 | 0.0291 |
| 5y8t | noschrodinger | 1.99 | 0.0045 | 0.8 | 0 | 98.96 | 1.13 | 0 | 0.49 | 0.71 | 0.1971 | 0.2176 | 0.0205 |
| 5y97 | opls | 3.05 | 0.0893 | 2.46 | 0 | 95.8 | 3.2 | 4 | 1.13 | 1.5 | 0.266 | 0.2887 | 0.0227 |
| 5y97 | noschrodinger | 3.05 | 0.0027 | 0.78 | 0 | 94.06 | 3.62 | 0 | 5.3 | 2.11 | 0.2558 | 0.2942 | 0.0384 |
| 5ycc | opls | 2.69 | 0.0913 | 2.44 | 0.54 | 98.64 | 0.97 | 2 | 0.33 | 0.62 | 0.1831 | 0.2316 | 0.0485 |
| 5ycc | noschrodinger | 2.69 | 0.0069 | 1 | 0.27 | 97.28 | 0.97 | 0 | 1.83 | 1.08 | 0.1817 | 0.2255 | 0.0438 |
| 5ydd | opls | 1.5 | 0.0987 | 2.46 | 0 | 100 | 1.65 | 3 | 4.19 | 1.37 | 0.1749 | 0.213 | 0.0381 |
| 5ydd | noschrodinger | 1.5 | 0.0036 | 0.85 | 0 | 100 | 1.09 | 0 | 2.64 | 1.08 | 0.1813 | 0.2127 | 0.0314 |
| 5yge | opls | 2.04 | 0.0962 | 2.36 | 0 | 98.48 | 0.35 | 0 | 2.15 | 0.99 | 0.1959 | 0.2201 | 0.0242 |
| 5yge | noschrodinger | 2.04 | 0.0031 | 0.81 | 0 | 97.89 | 2.08 | 0 | 4.13 | 1.46 | 0.1923 | 0.2187 | 0.0264 |
| 5yk6 | opls | 2.79 | 0.0895 | 2.36 | 0.43 | 96.57 | 2.74 | 1 | 0.97 | 1.34 | 0.2375 | 0.2489 | 0.0114 |
| 5yk6 | noschrodinger | 2.79 | 0.0028 | 0.78 | 0.43 | 95.74 | 4.55 | 0 | 2.67 | 1.85 | 0.227 | 0.245 | 0.018 |
| 5yng | opls | 2.5 | 0.0917 | 2.21 | 0.34 | 98.65 | 2.39 | 0 | 1.07 | 1.1 | 0.196 | 0.2325 | 0.0365 |
| 5yng | noschrodinger | 2.5 | 0.0029 | 0.8 | 0 | 98.65 | 2.39 | 0 | 1.92 | 1.24 | 0.1954 | 0.2227 | 0.0273 |
| 5ynm | opls | 1.68 | 0.0969 | 2.28 | 0 | 97.66 | 0.58 | 1 | 0.95 | 0.86 | 0.1886 | 0.2172 | 0.0286 |
| 5ynm | noschrodinger | 1.68 | 0.0046 | 0.93 | 0.25 | 96.77 | 0.29 | 0 | 1.27 | 1.05 | 0.1895 | 0.2166 | 0.0271 |
| 5yo2 | opls | 3 | 0.0894 | 2.42 | 0 | 98.31 | 1.1 | 0 | 0.75 | 0.77 | 0.2317 | 0.2665 | 0.0348 |
| 5yo2 | noschrodinger | 3 | 0.0027 | 0.8 | 0 | 96.97 | 1.09 | 0 | 2.47 | 1.24 | 0.1958 | 0.252 | 0.0562 |
| 5yoj | opls | 1.5 | 0.1048 | 2.86 | 0 | 99.48 | 3.08 | 0 | 12.33 | 1.97 | 0.1808 | 0.2088 | 0.028 |
| 5yoj | noschrodinger | 1.5 | 0.0052 | 1.12 | 0 | 99.48 | 1.54 | 0 | 10.37 | 1.68 | 0.1632 | 0.1933 | 0.0301 |
| 5yrl | opls | 2.1 | 0.1047 | 2.21 | 0 | 95.68 | 1.72 | 1 | 1.33 | 1.34 | 0.1366 | 0.191 | 0.0544 |
| 5yrl | noschrodinger | 2.1 | 0.0049 | 1.09 | 0 | 95.32 | 3.43 | 0 | 1.55 | 1.63 | 0.1394 | 0.1929 | 0.0535 |
| 5ys0 | opls | 2.6 | 0.0902 | 2.3 | 1.11 | 92.24 | 6.62 | 3 | 0.78 | 1.85 | 0.3056 | 0.3174 | 0.0118 |
| 5ys0 | noschrodinger | 2.6 | 0.0055 | 1.19 | 1.53 | 88.43 | 12.64 | 0 | 6.48 | 2.78 | 0.268 | 0.2866 | 0.0186 |
| 5ys3 | opls | 1.82 | 0.0922 | 2.24 | 0 | 98.9 | 0.25 | 3 | 1.53 | 0.9 | 0.1603 | 0.1883 | 0.028 |
| 5ys3 | noschrodinger | 1.82 | 0.0066 | 0.87 | 0 | 98.35 | 0.49 | 0 | 1.22 | 0.84 | 0.161 | 0.1881 | 0.0271 |
| 5ysi | opls | 1.55 | 0.0921 | 2.48 | 0 | 96.27 | 1.63 | 1 | 4.34 | 1.63 | 0.2046 | 0.2476 | 0.043 |
| 5ysi | noschrodinger | 1.55 | 0.0096 | 1.04 | 0 | 96.53 | 3.15 | 0 | 2.18 | 1.6 | 0.206 | 0.2426 | 0.0366 |
| 5yx4 | opls | 2.1 | 0.1015 | 2.21 | 0 | 98.25 | 2.35 | 0 | 0 | 0.78 | 0.2107 | 0.2358 | 0.0251 |
| 5yx4 | noschrodinger | 2.1 | 0.0035 | 0.87 | 0 | 96.96 | 2.92 | 0 | 2.06 | 1.51 | 0.1951 | 0.2285 | 0.0334 |
| 5z1n | opls | 1.95 | 0.0994 | 3.03 | 0 | 99.39 | 0.67 | 1 | 4.07 | 1.19 | 0.168 | 0.207 | 0.039 |
| 5z1n | noschrodinger | 1.95 | 0.0052 | 0.87 | 0 | 99.39 | 0.67 | 0 | 2.04 | 0.97 | 0.1749 | 0.2073 | 0.0324 |
| 5z39 | opls | 2.74 | 0.0917 | 2.94 | 0 | 97.32 | 2.11 | 0 | 0 | 0.88 | 0.2287 | 0.2729 | 0.0442 |
| 5z39 | noschrodinger | 2.74 | 0.0027 | 0.72 | 0 | 98.17 | 0 | 0 | 2.47 | 1.03 | 0.2147 | 0.2713 | 0.0566 |
| 5z41 | opls | 1.7 | 0.0966 | 2.47 | 1.04 | 94.79 | 2.17 | 1 | 1.74 | 1.54 | 0.2124 | 0.2598 | 0.0474 |
| 5z41 | noschrodinger | 1.7 | 0.0028 | 0.77 | 0 | 95.88 | 2.15 | 0 | 3.48 | 1.68 | 0.2185 | 0.2565 | 0.038 |
| 5z49 | opls | 2.15 | 0.0963 | 2.27 | 0.23 | 97.24 | 1.09 | 2 | 0.43 | 0.82 | 0.1981 | 0.2325 | 0.0344 |
| 5z49 | noschrodinger | 2.15 | 0.0028 | 0.79 | 0 | 98.17 | 0.81 | 0 | 0.87 | 0.77 | 0.1868 | 0.2217 | 0.0349 |
| 5z4h | opls | 2 | 0.0949 | 2.41 | 0.3 | 98.49 | 1.27 | 5 | 2.18 | 1.07 | 0.1735 | 0.2121 | 0.0386 |
| 5z4h | noschrodinger | 2 | 0.0048 | 0.82 | 0.27 | 98.1 | 0.95 | 0 | 0.85 | 0.76 | 0.1826 | 0.2101 | 0.0275 |
| 5z51 | opls | 1.58 | 0.1042 | 2.21 | 0.49 | 99.51 | 0 | 1 | 0.28 | 0.6 | 0.1844 | 0.2148 | 0.0304 |
| 5z51 | noschrodinger | 1.58 | 0.0114 | 1.12 | 0.44 | 98.68 | 0 | 0 | 0.83 | 0.76 | 0.1767 | 0.2163 | 0.0396 |
| 5z7b | opls | 2.1 | 0.095 | 2.21 | 0 | 99.43 | 0.33 | 1 | 0.32 | 0.62 | 0.1846 | 0.2089 | 0.0243 |
| 5z7b | noschrodinger | 2.1 | 0.0028 | 0.74 | 0 | 99.23 | 0.33 | 0 | 0.49 | 0.67 | 0.1749 | 0.2061 | 0.0312 |
| 5z9n | opls | 2.54 | 0.09 | 2.24 | 0.66 | 97.35 | 0 | 0 | 0.71 | 0.85 | 0.2059 | 0.2317 | 0.0258 |

| | | | | | | | | | | | | | |
|---|---|---|---|---|---|---|---|---|---|---|---|---|---|
| 5z9n | noschrodinger | 2.54 | 0.0027 | 0.76 | 0.54 | 97.84 | 0.68 | 0 | 1.08 | 0.85 | 0.1984 | 0.2335 | 0.0351 |
| 5zgq | opls | 1.5 | 0.1037 | 2.39 | 0 | 98.32 | 0.82 | 1 | 3.58 | 1.15 | 0.1321 | 0.1599 | 0.0278 |
| 5zgq | noschrodinger | 1.5 | 0.0061 | 1.05 | 0 | 98.12 | 0.54 | 0 | 1.38 | 0.87 | 0.1357 | 0.1563 | 0.0206 |
| 5zj5 | opls | 1.57 | 0.0969 | 2.26 | 0 | 96.12 | 0 | 0 | 1.95 | 1.23 | 0.1771 | 0.2077 | 0.0306 |
| 5zj5 | noschrodinger | 1.57 | 0.0036 | 0.95 | 0.65 | 96.13 | 0 | 0 | 0.2 | 0.84 | 0.1818 | 0.2112 | 0.0294 |
| 5zmf | opls | 3.56 | 0.0905 | 2.45 | 1.61 | 91.95 | 3.87 | 4 | 1.81 | 1.88 | 0.295 | 0.3324 | 0.0374 |
| 5zmf | noschrodinger | 3.56 | 0.0037 | 0.91 | 1.12 | 90.75 | 6.14 | 0 | 10.3 | 2.66 | 0.2709 | 0.3151 | 0.0442 |
| 5zqa | opls | 1.55 | 0.0982 | 2.2 | 0 | 98.37 | 0.97 | 0 | 1.06 | 0.81 | 0.1509 | 0.1935 | 0.0426 |
| 5zqa | noschrodinger | 1.55 | 0.0064 | 0.95 | 0 | 98.37 | 0.48 | 0 | 0 | 0.5 | 0.1547 | 0.1894 | 0.0347 |
| 5zza | opls | 1.53 | 0.1058 | 2.33 | 0 | 98.55 | 0.46 | 1 | 4.55 | 1.23 | 0.146 | 0.1716 | 0.0256 |
| 5zza | noschrodinger | 1.53 | 0.0069 | 1.04 | 0 | 98.56 | 0.23 | 0 | 2.82 | 1.07 | 0.1469 | 0.1702 | 0.0233 |
| 6a46 | opls | 2 | 0.0934 | 2.29 | 0 | 98.1 | 0.28 | 1 | 0.59 | 0.7 | 0.2149 | 0.2386 | 0.0237 |
| 6a46 | noschrodinger | 2 | 0.0029 | 0.78 | 0.23 | 97.65 | 0.27 | 0 | 1.48 | 0.96 | 0.1908 | 0.2256 | 0.0348 |
| 6afe | opls | 1.5 | 0.1019 | 2.45 | 0.61 | 98.18 | 0.7 | 1 | 7.19 | 1.4 | 0.1517 | 0.1713 | 0.0196 |
| 6afe | noschrodinger | 1.5 | 0.0056 | 0.94 | 0 | 98.38 | 0 | 0 | 2.74 | 1.06 | 0.1574 | 0.1695 | 0.0121 |
| 6alb | opls | 2.05 | 0.0904 | 2.32 | 0 | 96.92 | 0.52 | 1 | 0.28 | 0.79 | 0.2534 | 0.2683 | 0.0149 |
| 6alb | noschrodinger | 2.05 | 0.0026 | 0.83 | 0 | 97.55 | 1.01 | 0 | 0.56 | 0.79 | 0.2306 | 0.2576 | 0.027 |
| 6amx | opls | 2.05 | 0.0911 | 2.27 | 0 | 97.87 | 1.93 | 1 | 0.49 | 0.92 | 0.2401 | 0.2619 | 0.0218 |
| 6amx | noschrodinger | 2.05 | 0.0029 | 0.76 | 0 | 97.47 | 0.96 | 0 | 3.01 | 1.2 | 0.2197 | 0.2512 | 0.0315 |
| 6ao7 | opls | 1.85 | 0.0937 | 2.24 | 0 | 99.17 | 0.8 | 1 | 0.4 | 0.64 | 0.2248 | 0.2441 | 0.0193 |
| 6ao7 | noschrodinger | 1.85 | 0.003 | 0.88 | 0 | 98.68 | 1.6 | 0 | 1.62 | 1.07 | 0.2222 | 0.2527 | 0.0305 |
| 6aoh | opls | 3.44 | 0.0895 | 2.34 | 0.2 | 96.05 | 4.43 | 2 | 1.01 | 1.56 | 0.1793 | 0.234 | 0.0547 |
| 6aoh | noschrodinger | 3.44 | 0.0032 | 0.87 | 0.1 | 94.31 | 6.31 | 1 | 3.58 | 2.14 | 0.1667 | 0.2221 | 0.0554 |
| 6ap7 | opls | 1.51 | 0.1004 | 2.39 | 0 | 98.47 | 1.32 | 2 | 2.46 | 1.12 | 0.1703 | 0.1887 | 0.0184 |
| 6ap7 | noschrodinger | 1.51 | 0.0106 | 1.19 | 0.19 | 98.47 | 1.1 | 0 | 3.28 | 1.15 | 0.1708 | 0.1885 | 0.0177 |
| 6as5 | opls | 2.04 | 0.094 | 2.4 | 0 | 97.35 | 0 | 3 | 0.49 | 0.8 | 0.241 | 0.2679 | 0.0269 |
| 6as5 | noschrodinger | 2.04 | 0.0028 | 0.8 | 0.13 | 97.61 | 0.86 | 0 | 2.29 | 1.09 | 0.2239 | 0.2579 | 0.034 |
| 6ayh | opls | 2.05 | 0.0947 | 2.24 | 0 | 99.31 | 0 | 0 | 1.01 | 0.8 | 0.1922 | 0.2237 | 0.0315 |
| 6ayh | noschrodinger | 2.05 | 0.0027 | 0.72 | 0 | 97.89 | 0 | 0 | 0 | 0.53 | 0.1764 | 0.2149 | 0.0385 |
| 6b37 | opls | 1.5 | 0.0984 | 2.29 | 0 | 97.74 | 0 | 3 | 1.13 | 0.88 | 0.1891 | 0.2099 | 0.0208 |
| 6b37 | noschrodinger | 1.5 | 0.006 | 1.1 | 0 | 98.91 | 0.43 | 0 | 2.49 | 1.03 | 0.1837 | 0.2027 | 0.019 |
| 6b51 | opls | 2.05 | 0.0926 | 2.26 | 0.41 | 96.72 | 0 | 0 | 0 | 0.71 | 0.2089 | 0.2365 | 0.0276 |
| 6b51 | noschrodinger | 2.05 | 0.0029 | 0.85 | 0 | 95.9 | 0 | 0 | 0.49 | 0.95 | 0.1956 | 0.2264 | 0.0308 |
| 6b5k | opls | 1.6 | 0.0971 | 2.3 | 0.39 | 98.04 | 1.4 | 0 | 1.74 | 1.04 | 0.1932 | 0.2165 | 0.0233 |
| 6b5k | noschrodinger | 1.6 | 0.0045 | 0.95 | 0.56 | 97.19 | 3 | 0 | 1.28 | 1.36 | 0.1956 | 0.218 | 0.0224 |
| 6b7f | opls | 2.56 | 0.0957 | 2.28 | 0 | 98.97 | 3.41 | 1 | 1.37 | 1.27 | 0.1909 | 0.2241 | 0.0332 |
| 6b7f | noschrodinger | 2.56 | 0.003 | 0.85 | 0 | 98.72 | 5.28 | 0 | 2.37 | 1.57 | 0.1777 | 0.2097 | 0.032 |
| 6b88 | opls | 2.41 | 0.0902 | 2.25 | 0 | 96.52 | 1.81 | 2 | 1.31 | 1.28 | 0.2305 | 0.2575 | 0.027 |
| 6b88 | noschrodinger | 2.41 | 0.0028 | 0.84 | 0 | 94.9 | 3.33 | 0 | 1.59 | 1.66 | 0.233 | 0.2637 | 0.0307 |
| 6b89 | opls | 2 | 0.0927 | 2.29 | 0 | 97.79 | 0.26 | 0 | 0.93 | 0.83 | 0.2032 | 0.2213 | 0.0181 |
| 6b89 | noschrodinger | 2 | 0.0028 | 1.2 | 0 | 98.68 | 0.26 | 0 | 2 | 0.97 | 0.1832 | 0.212 | 0.0288 |
| 6bbx | opls | 2.2 | 0.0929 | 2.4 | 0 | 97.88 | 0.5 | 0 | 0.52 | 0.71 | 0.2248 | 0.2519 | 0.0271 |
| 6bbx | noschrodinger | 2.2 | 0.0031 | 1.32 | 0 | 96.64 | 2.45 | 0 | 2.36 | 1.53 | 0.2133 | 0.24 | 0.0267 |
| 6bc0 | opls | 2.2 | 0.0923 | 2.37 | 0.33 | 95.65 | 2.22 | 3 | 0.59 | 1.26 | 0.2827 | 0.3055 | 0.0228 |
| 6bc0 | noschrodinger | 2.2 | 0.0031 | 0.84 | 0.33 | 94.37 | 1.84 | 0 | 3.55 | 1.73 | 0.2835 | 0.3099 | 0.0264 |
| 6bds | opls | 1.53 | 0.1033 | 2.33 | 0 | 98.03 | 0.84 | 1 | 2.31 | 1.01 | 0.1693 | 0.2024 | 0.0331 |
| 6bds | noschrodinger | 1.53 | 0.0089 | 1.19 | 0 | 98.44 | 0.83 | 0 | 2.31 | 1.01 | 0.168 | 0.2018 | 0.0338 |
| 6bh8 | opls | 3.8 | 0.0905 | 2.22 | 0 | 98.95 | 1.91 | 0 | 1.22 | 1.05 | 0.3768 | 0.4135 | 0.0367 |
| 6bh8 | noschrodinger | 3.8 | 0.0026 | 0.74 | 0 | 98.44 | 4.35 | 0 | 1.6 | 1.39 | 0.3622 | 0.4111 | 0.0489 |

| | | | | | | | | | | | | | |
|---|---|---|---|---|---|---|---|---|---|---|---|---|---|
| 6bin | opls | 2.5 | 0.0924 | 2.22 | 0 | 97.66 | 0.3 | 2 | 1.14 | 0.9 | 0.216 | 0.2412 | 0.0252 |
| 6bin | noschrodinger | 2.5 | 0.0033 | 0.79 | 0 | 98.12 | 0 | 0 | 2.13 | 0.99 | 0.1923 | 0.2302 | 0.0379 |
| 6bml | opls | 2.95 | 0.0895 | 2.24 | 0.19 | 96.81 | 0.59 | 0 | 0.31 | 0.81 | 0.2631 | 0.2857 | 0.0226 |
| 6bml | noschrodinger | 2.95 | 0.0029 | 0.72 | 0 | 97.77 | 0.59 | 0 | 2.73 | 1.11 | 0.2568 | 0.2789 | 0.0221 |
| 6bmn | opls | 2.25 | 0.0903 | 2.26 | 0.38 | 96.37 | 1.2 | 0 | 0.85 | 1.06 | 0.2608 | 0.2823 | 0.0215 |
| 6bmn | noschrodinger | 2.25 | 0.0025 | 0.73 | 0 | 97.21 | 1.19 | 0 | 2.14 | 1.19 | 0.258 | 0.28 | 0.022 |
| 6bqo | opls | 2.8 | 0.0914 | 2.23 | 0 | 96.75 | 0.79 | 1 | 0.75 | 0.94 | 0.2537 | 0.2695 | 0.0158 |
| 6bqo | noschrodinger | 2.8 | 0.0029 | 0.76 | 0.58 | 95.34 | 0.39 | 0 | 2.06 | 1.3 | 0.2501 | 0.2748 | 0.0247 |
| 6br8 | opls | 2.3 | 0.0858 | 2.41 | 0.21 | 97.46 | 1.5 | 2 | 0.76 | 0.98 | 0.2666 | 0.2988 | 0.0322 |
| 6br8 | noschrodinger | 2.3 | 0.0037 | 0.73 | 0.2 | 94.94 | 0.21 | 0 | 2.1 | 1.33 | 0.2711 | 0.2918 | 0.0207 |
| 6brk | opls | 3.5 | 0.0903 | 2.42 | 0 | 96.47 | 3.39 | 3 | 0.71 | 1.36 | 0.2817 | 0.3139 | 0.0322 |
| 6brk | noschrodinger | 3.5 | 0.0028 | 0.76 | 0 | 94.88 | 6.9 | 0 | 1.79 | 1.93 | 0.2654 | 0.302 | 0.0366 |
| 6bt4 | opls | 2.31 | 0.0911 | 2.19 | 0 | 96.23 | 2.33 | 0 | 0.38 | 1.17 | 0.2132 | 0.2317 | 0.0185 |
| 6bt4 | noschrodinger | 2.31 | 0.0083 | 1.13 | 0.62 | 95.65 | 1.53 | 0 | 0.77 | 1.18 | 0.1955 | 0.2252 | 0.0297 |
| 6bvc | opls | 1.81 | 0.1016 | 2.56 | 0 | 98.03 | 0 | 0 | 3.09 | 1.1 | 0.1525 | 0.1846 | 0.0321 |
| 6bvc | noschrodinger | 1.81 | 0.004 | 0.92 | 0 | 98.68 | 0 | 0 | 2.32 | 1.01 | 0.1572 | 0.1833 | 0.0261 |
| 6bvq | opls | 2.08 | 0.0965 | 2.29 | 0 | 97.56 | 0.68 | 0 | 0.74 | 0.83 | 0.2288 | 0.271 | 0.0422 |
| 6bvq | noschrodinger | 2.08 | 0.0071 | 0.98 | 0.6 | 95.24 | 0.68 | 0 | 2.58 | 1.37 | 0.2232 | 0.2679 | 0.0447 |
| 6bwh | opls | 2.18 | 0.0938 | 2.36 | 0 | 99.15 | 1.65 | 4 | 0.56 | 0.85 | 0.2507 | 0.2784 | 0.0277 |
| 6bwh | noschrodinger | 2.18 | 0.0032 | 0.81 | 0.17 | 95.51 | 3.46 | 0 | 2.93 | 1.8 | 0.2336 | 0.269 | 0.0354 |
| 6bxt | opls | 2.7 | 0.0908 | 2.29 | 0 | 98.95 | 1.92 | 2 | 1.17 | 1.04 | 0.2356 | 0.2838 | 0.0482 |
| 6bxt | noschrodinger | 2.7 | 0.0028 | 0.74 | 0 | 96.47 | 1.11 | 0 | 2 | 1.23 | 0.2233 | 0.2683 | 0.045 |
| 6bym | opls | 2.2 | 0.0939 | 2.19 | 0 | 97.41 | 1.1 | 0 | 0.63 | 0.86 | 0.1807 | 0.2166 | 0.0359 |
| 6bym | noschrodinger | 2.2 | 0.0029 | 0.83 | 0 | 97.19 | 1.1 | 0 | 2.66 | 1.23 | 0.1641 | 0.2029 | 0.0388 |
| 6bzg | opls | 2.13 | 0.0928 | 2.23 | 0 | 98.92 | 0.28 | 0 | 1.08 | 0.81 | 0.2657 | 0.2913 | 0.0256 |
| 6bzg | noschrodinger | 2.13 | 0.0024 | 0.68 | 0 | 98.15 | 0.28 | 0 | 1.39 | 0.87 | 0.258 | 0.2889 | 0.0309 |
| 6c0b | opls | 2.5 | 0.0908 | 2.26 | 0.17 | 97.02 | 2.26 | 3 | 0.8 | 1.19 | 0.2271 | 0.2455 | 0.0184 |
| 6c0b | noschrodinger | 2.5 | 0.0028 | 0.78 | 0.32 | 93.86 | 4.84 | 0 | 3.01 | 2.02 | 0.2324 | 0.2478 | 0.0154 |
| 6c1m | opls | 2.52 | 0.0849 | 2.45 | 0.91 | 96.6 | 3.2 | 2 | 1.65 | 1.52 | 0.2538 | 0.2678 | 0.014 |
| 6c1m | noschrodinger | 2.52 | 0.007 | 1.12 | 0.67 | 94.43 | 3.68 | 0 | 12.36 | 2.41 | 0.2392 | 0.2604 | 0.0212 |
| 6c3n | opls | 2.53 | 0.0942 | 2.33 | 0 | 97.2 | 1.87 | 0 | 1 | 1.15 | 0.195 | 0.234 | 0.039 |
| 6c3n | noschrodinger | 2.53 | 0.0025 | 0.76 | 0 | 96.68 | 1.85 | 0 | 0.76 | 1.15 | 0.2007 | 0.2452 | 0.0445 |
| 6c57 | opls | 3.5 | 0.087 | 2.3 | 0 | 98.5 | 1.93 | 3 | 0.49 | 0.89 | 0.2169 | 0.259 | 0.0421 |
| 6c57 | noschrodinger | 3.5 | 0.0032 | 0.85 | 0.58 | 91.3 | 6.49 | 0 | 2.74 | 2.19 | 0.2219 | 0.2482 | 0.0263 |
| 6c9g | opls | 2.57 | 0.0909 | 2.25 | 0.12 | 97.74 | 0.64 | 1 | 1.05 | 0.86 | 0.2536 | 0.2763 | 0.0227 |
| 6c9g | noschrodinger | 2.57 | 0.0051 | 0.82 | 0 | 97.87 | 0.38 | 0 | 2.59 | 1.07 | 0.2497 | 0.2771 | 0.0274 |
| 6cc0 | opls | 2.5 | 0.0918 | 2.26 | 0 | 97.62 | 0.97 | 0 | 0.64 | 0.79 | 0.2268 | 0.2579 | 0.0311 |
| 6cc0 | noschrodinger | 2.5 | 0.0027 | 0.75 | 0 | 96.98 | 2.67 | 0 | 1.55 | 1.4 | 0.2126 | 0.2461 | 0.0335 |
| 6cd3 | opls | 2.61 | 0.0946 | 2.23 | 0 | 96.34 | 0.68 | 1 | 1.86 | 1.19 | 0.2279 | 0.2567 | 0.0288 |
| 6cd3 | noschrodinger | 2.61 | 0.0039 | 0.9 | 1.19 | 95.83 | 2.05 | 0 | 1.86 | 1.47 | 0.2046 | 0.2484 | 0.0438 |
| 6ci5 | opls | 1.65 | 0.0922 | 2.28 | 0.54 | 98.92 | 0.58 | 1 | 1.53 | 0.9 | 0.2456 | 0.2712 | 0.0256 |
| 6ci5 | noschrodinger | 1.65 | 0.0027 | 0.75 | 0.53 | 98.93 | 0.58 | 0 | 1.53 | 0.9 | 0.2485 | 0.2683 | 0.0198 |
| 6ck0 | opls | 2.25 | 0.0921 | 2.32 | 0.58 | 98.27 | 0.3 | 0 | 0.3 | 0.61 | 0.1902 | 0.2311 | 0.0409 |
| 6ck0 | noschrodinger | 2.25 | 0.006 | 1.02 | 0.48 | 97.59 | 1.19 | 0 | 2.86 | 1.22 | 0.178 | 0.2219 | 0.0439 |
| 6co9 | opls | 1.6 | 0.1026 | 2.24 | 0 | 97.55 | 0.68 | 0 | 2.24 | 1.09 | 0.165 | 0.1888 | 0.0238 |
| 6co9 | noschrodinger | 1.6 | 0.0095 | 1.18 | 0 | 97.77 | 0.68 | 0 | 1.78 | 0.99 | 0.1591 | 0.1884 | 0.0293 |
| 6csn | opls | 2.9 | 0.0892 | 2.37 | 0.4 | 96.05 | 4.37 | 1 | 0.89 | 1.53 | 0.2456 | 0.2956 | 0.05 |
| 6csn | noschrodinger | 2.9 | 0.0026 | 0.75 | 0.36 | 96.73 | 5.22 | 0 | 1.58 | 1.65 | 0.2524 | 0.2976 | 0.0452 |
| 6cvh | opls | 3.5 | 0.0906 | 2.44 | 0.45 | 96.41 | 2.48 | 1 | 1.06 | 1.34 | 0.292 | 0.3184 | 0.0264 |

| | | | | | | | | | | | | | |
|---|---|---|---|---|---|---|---|---|---|---|---|---|---|
| 6cvh | noschrodinger | 3.5 | 0.0034 | 0.93 | 2.22 | 92.44 | 1.49 | 0 | 7.14 | 1.99 | 0.278 | 0.3138 | 0.0358 |
| 6cwi | opls | 2.16 | 0.0942 | 2.16 | 0 | 98.45 | 2.38 | 1 | 0.38 | 0.92 | 0.2427 | 0.2746 | 0.0319 |
| 6cwi | noschrodinger | 2.16 | 0.0026 | 0.73 | 0 | 97.54 | 3.15 | 0 | 1.16 | 1.3 | 0.2282 | 0.2751 | 0.0469 |
| 6cwn | opls | 1.53 | 0.101 | 2.23 | 0 | 99.38 | 1.6 | 1 | 0.76 | 0.9 | 0.1581 | 0.1832 | 0.0251 |
| 6cwn | noschrodinger | 1.53 | 0.0107 | 1.14 | 0 | 99.38 | 1.59 | 0 | 3.83 | 1.32 | 0.1537 | 0.1803 | 0.0266 |
| 6cx0 | opls | 3.5 | 0.0904 | 2.4 | 0.52 | 94.44 | 8.52 | 7 | 0.78 | 1.83 | 0.3026 | 0.3519 | 0.0493 |
| 6cx0 | noschrodinger | 3.5 | 0.0027 | 0.79 | 0.83 | 93.39 | 10.97 | 0 | 4.82 | 2.47 | 0.2968 | 0.3466 | 0.0498 |
| 6cxh | opls | 2.7 | 0.0901 | 2.34 | 1.13 | 93.37 | 2.5 | 8 | 0.86 | 1.5 | 0.2705 | 0.2803 | 0.0098 |
| 6cxh | noschrodinger | 2.7 | 0.0054 | 1.03 | 1.4 | 88.1 | 3.15 | 0 | 10.37 | 2.51 | 0.2594 | 0.2699 | 0.0105 |
| 6cy6 | opls | 1.75 | 0.0958 | 2.41 | 0 | 99.4 | 1.92 | 1 | 0.32 | 0.84 | 0.1588 | 0.186 | 0.0272 |
| 6cy6 | noschrodinger | 1.75 | 0.0076 | 1.02 | 0 | 99.4 | 0.64 | 0 | 1.94 | 0.96 | 0.1676 | 0.1829 | 0.0153 |
| 6czh | opls | 1.8 | 0.0932 | 2.22 | 0 | 98.73 | 1.73 | 0 | 4.22 | 1.39 | 0.1924 | 0.2386 | 0.0462 |
| 6czh | noschrodinger | 1.8 | 0.0129 | 1.45 | 0 | 97.63 | 2.27 | 0 | 12.88 | 1.97 | 0.1962 | 0.2296 | 0.0334 |
| 6d0y | opls | 2.68 | 0.0938 | 2.3 | 0.12 | 97.47 | 2.39 | 4 | 0.41 | 1.04 | 0.1905 | 0.2357 | 0.0452 |
| 6d0y | noschrodinger | 2.68 | 0.0027 | 0.75 | 0 | 98.54 | 3.39 | 0 | 1.09 | 1.22 | 0.1831 | 0.2253 | 0.0422 |
| 6d32 | opls | 3.75 | 0.09 | 2.28 | 0.18 | 96.04 | 1.24 | 2 | 0.55 | 1.03 | 0.2421 | 0.2859 | 0.0438 |
| 6d32 | noschrodinger | 3.75 | 0.0031 | 0.94 | 0.18 | 94.89 | 2.68 | 0 | 7 | 2.06 | 0.239 | 0.2795 | 0.0405 |
| 6d4g | opls | 2.8 | 0.0898 | 2.34 | 0 | 97.44 | 0.28 | 1 | 0.45 | 0.77 | 0.3006 | 0.3291 | 0.0285 |
| 6d4g | noschrodinger | 2.8 | 0.0029 | 0.76 | 0.25 | 96.25 | 0.81 | 0 | 2.1 | 1.23 | 0.2841 | 0.3156 | 0.0315 |
| 6d62 | opls | 1.77 | 0.1071 | 2.3 | 0 | 98.81 | 1.33 | 1 | 2.51 | 1.13 | 0.1662 | 0.2025 | 0.0363 |
| 6d62 | noschrodinger | 1.77 | 0.0057 | 1.04 | 0 | 98.84 | 2 | 0 | 5.38 | 1.52 | 0.1678 | 0.202 | 0.0342 |
| 6d6d | opls | 1.7 | 0.0961 | 2.25 | 0.31 | 99.06 | 2.67 | 4 | 0.97 | 1.11 | 0.1971 | 0.231 | 0.0339 |
| 6d6d | noschrodinger | 1.7 | 0.0034 | 0.84 | 0.31 | 98.45 | 3.41 | 0 | 1.36 | 1.27 | 0.1962 | 0.2269 | 0.0307 |
| 6deb | opls | 1.7 | 0.0996 | 2.37 | 0 | 98.04 | 0.59 | 7 | 2.15 | 0.99 | 0.1567 | 0.1852 | 0.0285 |
| 6deb | noschrodinger | 1.7 | 0.0044 | 0.99 | 0 | 98.23 | 0.99 | 1 | 2.68 | 1.05 | 0.1607 | 0.1815 | 0.0208 |
| 6dex | opls | 2.1 | 0.0937 | 2.29 | 0 | 97.97 | 0.74 | 0 | 0.39 | 0.65 | 0.254 | 0.2746 | 0.0206 |
| 6dex | noschrodinger | 2.1 | 0.0076 | 0.88 | 0 | 98.12 | 1.47 | 0 | 1.75 | 1.06 | 0.2297 | 0.2652 | 0.0355 |
| 6df7 | opls | 2 | 0.0973 | 2.23 | 0 | 98.79 | 1.67 | 2 | 0 | 0.67 | 0.2183 | 0.2365 | 0.0182 |
| 6df7 | noschrodinger | 2 | 0.0031 | 0.88 | 0 | 99.6 | 2.49 | 0 | 0.95 | 1.08 | 0.2047 | 0.2332 | 0.0285 |
| 6dg5 | opls | 2.52 | 0.0869 | 2.28 | 0 | 95.79 | 1.67 | 4 | 1.86 | 1.41 | 0.2219 | 0.2558 | 0.0339 |
| 6dg5 | noschrodinger | 2.52 | 0.0029 | 0.75 | 0.21 | 96.06 | 3.55 | 0 | 2.77 | 1.75 | 0.2114 | 0.2495 | 0.0381 |
| 6dnu | opls | 2.28 | 0.0916 | 2.18 | 0 | 98.24 | 1.04 | 0 | 0.27 | 0.61 | 0.2144 | 0.2397 | 0.0253 |
| 6dnu | noschrodinger | 2.28 | 0.0026 | 0.74 | 0 | 97.38 | 0.51 | 0 | 1.34 | 0.98 | 0.2025 | 0.2389 | 0.0364 |
| 6dxz | opls | 2.7 | 0.0944 | 2.27 | 0 | 95.61 | 1.43 | 0 | 0.54 | 1.11 | 0.1856 | 0.2274 | 0.0418 |
| 6dxz | noschrodinger | 2.7 | 0.003 | 1.81 | 0 | 95.64 | 1.07 | 0 | 1.97 | 1.29 | 0.1834 | 0.2231 | 0.0397 |
| 6.00E+12 | opls | 2.4 | 0.0944 | 2.28 | 0.27 | 98.4 | 4.8 | 3 | 0.88 | 1.29 | 0.1913 | 0.2093 | 0.018 |
| 6.00E+12 | noschrodinger | 2.4 | 0.0028 | 0.8 | 0 | 98.08 | 5.38 | 0 | 1.61 | 1.46 | 0.1837 | 0.2041 | 0.0204 |
| 6e9o | opls | 3.5 | 0.0908 | 2.33 | 0.26 | 95.92 | 3.39 | 5 | 0.88 | 1.45 | 0.3203 | 0.3938 | 0.0735 |
| 6e9o | noschrodinger | 3.5 | 0.0026 | 0.73 | 0.64 | 93.74 | 3.84 | 0 | 2.9 | 1.94 | 0.3119 | 0.3847 | 0.0728 |
| 6efz | opls | 3.5 | 0.0883 | 2.29 | 0.68 | 95.55 | 3.47 | 2 | 0.8 | 1.47 | 0.2538 | 0.2635 | 0.0097 |
| 6efz | noschrodinger | 3.5 | 0.004 | 1.86 | 0.34 | 91.84 | 3.83 | 0 | 4.41 | 2.16 | 0.2477 | 0.2662 | 0.0185 |
| 6eg0 | opls | 2.9 | 0.0891 | 2.28 | 0.2 | 95.44 | 2.23 | 0 | 0.82 | 1.34 | 0.2559 | 0.2757 | 0.0198 |
| 6eg0 | noschrodinger | 2.9 | 0.0036 | 0.78 | 0 | 94.07 | 2.22 | 0 | 7.51 | 2.07 | 0.2479 | 0.2647 | 0.0168 |
| 6ei4 | opls | 1.69 | 0.0974 | 2.34 | 0 | 97.35 | 0.4 | 7 | 2.58 | 1.17 | 0.2369 | 0.275 | 0.0381 |
| 6ei4 | noschrodinger | 1.69 | 0.0035 | 0.85 | 0 | 97.55 | 0.4 | 0 | 3.66 | 1.25 | 0.2329 | 0.2598 | 0.0269 |
| 6el2 | opls | 1.9 | 0.0919 | 2.32 | 0 | 98.61 | 0 | 0 | 0.31 | 0.62 | 0.1992 | 0.2362 | 0.037 |
| 6el2 | noschrodinger | 1.9 | 0.0025 | 0.69 | 0 | 98.62 | 0 | 0 | 1.57 | 0.9 | 0.1965 | 0.2278 | 0.0313 |
| 6eup | opls | 2.65 | 0.0894 | 2.23 | 0 | 99.77 | 2.32 | 0 | 0.29 | 0.88 | 0.2525 | 0.2811 | 0.0286 |
| 6eup | noschrodinger | 2.65 | 0.0027 | 0.67 | 0 | 98.62 | 4.31 | 0 | 0.57 | 1.17 | 0.2333 | 0.2723 | 0.039 |

| | | | | | | | | | | | | | |
|---|---|---|---|---|---|---|---|---|---|---|---|---|---|
| 6euu | opls | 2.6 | 0.0904 | 2.2 | 0 | 97.3 | 1.7 | 0 | 0.53 | 0.99 | 0.2305 | 0.268 | 0.0375 |
| 6euu | noschrodinger | 2.6 | 0.0029 | 0.72 | 0 | 94.18 | 1.94 | 0 | 1.58 | 1.51 | 0.2369 | 0.2716 | 0.0347 |
| 6eux | opls | 2 | 0.0955 | 2.29 | 0.48 | 96.19 | 1 | 0 | 0 | 0.76 | 0.2249 | 0.2617 | 0.0368 |
| 6eux | noschrodinger | 2 | 0.0023 | 0.69 | 0 | 98.14 | 0.98 | 0 | 0 | 0.5 | 0.2199 | 0.2485 | 0.0286 |
| 6ewp | opls | 1.85 | 0.0964 | 2.48 | 0 | 97.58 | 0.63 | 7 | 4.17 | 1.29 | 0.1913 | 0.2204 | 0.0291 |
| 6ewp | noschrodinger | 1.85 | 0.0098 | 1 | 0 | 97.79 | 0 | 0 | 2.26 | 1.05 | 0.1969 | 0.2182 | 0.0213 |
| 6ex0 | opls | 2.78 | 0.0913 | 2.5 | 0.53 | 98.15 | 1.66 | 1 | 1.2 | 1 | 0.2465 | 0.2836 | 0.0371 |
| 6ex0 | noschrodinger | 2.78 | 0.0037 | 0.76 | 0.26 | 95.82 | 3.55 | 0 | 2.86 | 1.78 | 0.2446 | 0.2785 | 0.0339 |
| 6exk | opls | 2.1 | 0.0923 | 2.18 | 0 | 97.78 | 1.92 | 0 | 0.52 | 0.94 | 0.2275 | 0.2549 | 0.0274 |
| 6exk | noschrodinger | 2.1 | 0.0029 | 0.77 | 0.22 | 96.69 | 1.44 | 0 | 1.81 | 1.27 | 0.2231 | 0.256 | 0.0329 |
| 6exm | opls | 1.6 | 0.0985 | 2.38 | 0 | 97.6 | 1.66 | 1 | 2.3 | 1.26 | 0.18 | 0.2064 | 0.0264 |
| 6exm | noschrodinger | 1.6 | 0.0054 | 0.84 | 0 | 98.47 | 0.95 | 0 | 2.43 | 1.02 | 0.1873 | 0.2032 | 0.0159 |
| 6eyi | opls | 2.04 | 0.0924 | 2.24 | 1.09 | 96.38 | 0.41 | 1 | 0.9 | 1.01 | 0.2539 | 0.2883 | 0.0344 |
| 6eyi | noschrodinger | 2.04 | 0.0034 | 0.95 | 1.08 | 96.4 | 0.4 | 0 | 0.45 | 0.9 | 0.2559 | 0.288 | 0.0321 |
| 6eyu | opls | 2.5 | 0.0854 | 2.59 | 0 | 95.71 | 8.04 | 6 | 0.8 | 1.74 | 0.2479 | 0.2569 | 0.009 |
| 6eyu | noschrodinger | 2.5 | 0.0028 | 0.81 | 0.44 | 95.49 | 13.45 | 0 | 3.37 | 2.3 | 0.2348 | 0.2483 | 0.0135 |
| 6f01 | opls | 2.3 | 0.0936 | 2.24 | 0.27 | 98.4 | 0.59 | 1 | 0.3 | 0.61 | 0.1987 | 0.2245 | 0.0258 |
| 6f01 | noschrodinger | 2.3 | 0.003 | 0.86 | 0 | 98.3 | 1.17 | 0 | 2.58 | 1.1 | 0.2017 | 0.2247 | 0.023 |
| 6f20 | opls | 2 | 0.0939 | 2.2 | 0 | 99.34 | 1.1 | 0 | 0.79 | 0.78 | 0.1931 | 0.2415 | 0.0484 |
| 6f20 | noschrodinger | 2 | 0.0029 | 0.78 | 0 | 98.36 | 1.47 | 0 | 0.39 | 0.77 | 0.1972 | 0.2395 | 0.0423 |
| 6f37 | opls | 2.2 | 0.093 | 2.24 | 0 | 97.24 | 0.6 | 1 | 0.87 | 0.91 | 0.2213 | 0.2373 | 0.016 |
| 6f37 | noschrodinger | 2.2 | 0.0033 | 0.75 | 0.45 | 95.93 | 0 | 0 | 1.16 | 1.11 | 0.2289 | 0.243 | 0.0141 |
| 6f5u | opls | 2.07 | 0.0925 | 2.25 | 0 | 96.93 | 0.65 | 0 | 1.28 | 1.03 | 0.2004 | 0.2242 | 0.0238 |
| 6f5u | noschrodinger | 2.07 | 0.003 | 0.79 | 0.27 | 96.98 | 0.32 | 0 | 3.55 | 1.32 | 0.1992 | 0.2207 | 0.0215 |
| 6f6r | opls | 1.8 | 0.1005 | 2.41 | 0 | 96.54 | 2.48 | 2 | 2.73 | 1.59 | 0.1665 | 0.2026 | 0.0361 |
| 6f6r | noschrodinger | 1.8 | 0.0129 | 1.26 | 0 | 96.59 | 2.06 | 0 | 2.74 | 1.52 | 0.1708 | 0.202 | 0.0312 |
| 6fct | opls | 1.89 | 0.0998 | 2.24 | 0 | 99.51 | 2.29 | 0 | 0.3 | 0.88 | 0.1675 | 0.2131 | 0.0456 |
| 6fct | noschrodinger | 1.89 | 0.0029 | 0.88 | 0 | 99.52 | 2.27 | 0 | 0.89 | 1.04 | 0.1687 | 0.2065 | 0.0378 |
| 6fcw | opls | 2 | 0.0983 | 2.32 | 0 | 99.03 | 0.56 | 1 | 0.3 | 0.61 | 0.1947 | 0.2219 | 0.0272 |
| 6fcw | noschrodinger | 2 | 0.0101 | 1.2 | 0 | 98.55 | 1.12 | 0 | 0.59 | 0.74 | 0.1655 | 0.2201 | 0.0546 |
| 6fhw | opls | 3.6 | 0.088 | 2.4 | 1.46 | 91.08 | 7.1 | 17 | 1.86 | 2.11 | 0.2021 | 0.2346 | 0.0325 |
| 6fhw | noschrodinger | 3.6 | 0.0031 | 0.82 | 1.8 | 89.8 | 5.61 | 0 | 2.95 | 2.21 | 0.2112 | 0.2362 | 0.025 |
| 6fjc | opls | 2.6 | 0.0931 | 2.39 | 0.39 | 93.73 | 1.72 | 0 | 2.93 | 1.68 | 0.1999 | 0.2511 | 0.0512 |
| 6fjc | noschrodinger | 2.6 | 0.003 | 0.81 | 0 | 94.9 | 5.15 | 0 | 2.94 | 1.98 | 0.2047 | 0.246 | 0.0413 |
| 6fm2 | opls | 2.8 | 0.0923 | 2.3 | 0.39 | 94.79 | 4.41 | 2 | 0.84 | 1.61 | 0.3118 | 0.3349 | 0.0231 |
| 6fm2 | noschrodinger | 2.8 | 0.0028 | 0.8 | 0.38 | 93.28 | 7.02 | 0 | 2.05 | 2.05 | 0.2873 | 0.3215 | 0.0342 |
| 6fm9 | opls | 3.6 | 0.0875 | 2.27 | 0 | 97.56 | 1.03 | 0 | 0.69 | 0.82 | 0.3319 | 0.3566 | 0.0247 |
| 6fm9 | noschrodinger | 3.6 | 0.0027 | 0.76 | 0 | 97.58 | 2.75 | 0 | 0.71 | 1.15 | 0.3216 | 0.3636 | 0.042 |
| 6fmx | opls | 1.79 | 0.0912 | 2.84 | 0 | 98.41 | 0.6 | 2 | 3.65 | 1.15 | 0.2198 | 0.2462 | 0.0264 |
| 6fmx | noschrodinger | 1.79 | 0.0042 | 0.89 | 0 | 98.99 | 1.19 | 0 | 5.65 | 1.36 | 0.2175 | 0.245 | 0.0275 |
| 6fs2 | opls | 2.55 | 0.0901 | 2.43 | 0.35 | 96.54 | 3.52 | 6 | 0.8 | 1.39 | 0.1846 | 0.2361 | 0.0515 |
| 6fs2 | noschrodinger | 2.55 | 0.0028 | 0.72 | 0.34 | 96.61 | 5.73 | 0 | 1.02 | 1.59 | 0.1912 | 0.2325 | 0.0413 |
| 6fuf | opls | 3.12 | 0.0904 | 2.29 | 0.62 | 93.4 | 0.23 | 2 | 0.88 | 1.2 | 0.263 | 0.2951 | 0.0321 |
| 6fuf | noschrodinger | 3.12 | 0.0036 | 0.8 | 0.61 | 92.06 | 0.23 | 0 | 2.89 | 1.56 | 0.2709 | 0.2994 | 0.0285 |
| 6fy9 | opls | 3.65 | 0.0902 | 2.35 | 0.88 | 95.35 | 1.72 | 2 | 2.02 | 1.47 | 0.2478 | 0.2677 | 0.0199 |
| 6fy9 | noschrodinger | 3.65 | 0.0039 | 0.89 | 0.66 | 94.04 | 3.14 | 0 | 9.26 | 2.27 | 0.2131 | 0.2603 | 0.0472 |
| 6g14 | opls | 1.8 | 0.0986 | 2.3 | 0 | 98.54 | 1.04 | 1 | 1.18 | 0.84 | 0.2156 | 0.2464 | 0.0308 |
| 6g14 | noschrodinger | 1.8 | 0.0119 | 1.14 | 0.18 | 98.92 | 1.65 | 0 | 3.11 | 1.27 | 0.2115 | 0.2442 | 0.0327 |
| 6g5j | opls | 1.8 | 0.0941 | 2.21 | 0 | 99.16 | 2.4 | 1 | 1.28 | 1.14 | 0.1726 | 0.2076 | 0.035 |

| | | | | | | | | | | | | | |
|---|---|---|---|---|---|---|---|---|---|---|---|---|---|
| 6g5j | noschrodinger | 1.8 | 0.0032 | 0.87 | 0 | 99.17 | 2.38 | 0 | 0.77 | 1.03 | 0.1778 | 0.2087 | 0.0309 |
| 6g6w | opls | 2.72 | 0.0906 | 2.27 | 0.1 | 98.09 | 0.73 | 3 | 0.46 | 0.66 | 0.2385 | 0.265 | 0.0265 |
| 6g6w | noschrodinger | 2.72 | 0.0031 | 0.76 | 0.09 | 96.12 | 1.34 | 0 | 1.45 | 1.24 | 0.2484 | 0.2646 | 0.0162 |
| 6g7h | opls | 1.5 | 0.0885 | 2.3 | 0 | 99.08 | 0.55 | 1 | 0.72 | 0.73 | 0.1435 | 0.1664 | 0.0229 |
| 6g7h | noschrodinger | 1.5 | 0.0065 | 1.02 | 0 | 98.68 | 0.54 | 0 | 1.25 | 0.84 | 0.1477 | 0.1648 | 0.0171 |
| 6ga0 | opls | 2 | 0.0969 | 2.24 | 0.66 | 94.7 | 0 | 0 | 0.79 | 1.11 | 0.2071 | 0.2245 | 0.0174 |
| 6ga0 | noschrodinger | 2 | 0.0029 | 0.84 | 0.66 | 96.71 | 0 | 0 | 2.76 | 1.27 | 0.1899 | 0.2162 | 0.0263 |
| 6gaq | opls | 2.5 | 0.089 | 2.31 | 0 | 96.88 | 0.85 | 2 | 0.66 | 0.9 | 0.2495 | 0.2923 | 0.0428 |
| 6gaq | noschrodinger | 2.5 | 0.0031 | 0.79 | 0.35 | 96.88 | 0.85 | 0 | 3.09 | 1.29 | 0.2336 | 0.2895 | 0.0559 |
| 6gb1 | opls | 2.73 | 0.0907 | 2.27 | 1.63 | 94.31 | 4.35 | 0 | 0.46 | 1.53 | 0.2305 | 0.2568 | 0.0263 |
| 6gb1 | noschrodinger | 2.73 | 0.003 | 0.89 | 3.05 | 88.55 | 11.3 | 0 | 1.86 | 2.33 | 0.1993 | 0.2307 | 0.0314 |
| 6ghy | opls | 2.12 | 0.0994 | 2.32 | 0 | 99.46 | 0 | 0 | 0.68 | 0.72 | 0.2015 | 0.2379 | 0.0364 |
| 6ghy | noschrodinger | 2.12 | 0.0027 | 0.78 | 0 | 98.67 | 0.98 | 0 | 1.52 | 0.89 | 0.1846 | 0.232 | 0.0474 |
| 6gie | opls | 2.1 | 0.0905 | 2.17 | 0 | 98.5 | 0.61 | 1 | 0.59 | 0.7 | 0.242 | 0.2719 | 0.0299 |
| 6gie | noschrodinger | 2.1 | 0.0029 | 0.78 | 0 | 98.54 | 2.38 | 0 | 2.69 | 1.34 | 0.2303 | 0.2659 | 0.0356 |
| 6gnk | opls | 2.55 | 0.0907 | 2.33 | 0.15 | 97.85 | 3.96 | 5 | 1.34 | 1.35 | 0.2561 | 0.2715 | 0.0154 |
| 6gnk | noschrodinger | 2.55 | 0.0028 | 0.73 | 0.15 | 97.56 | 6.09 | 0 | 0.67 | 1.41 | 0.2383 | 0.2521 | 0.0138 |
| 6gnn | opls | 3.79 | 0.0904 | 2.34 | 1.69 | 92.03 | 4.89 | 3 | 1.07 | 1.82 | 0.2919 | 0.3385 | 0.0466 |
| 6gnn | noschrodinger | 3.79 | 0.0028 | 0.72 | 3.36 | 85.61 | 3.99 | 0 | 5.19 | 2.38 | 0.2657 | 0.3237 | 0.058 |
| 6gu0 | opls | 2.5 | 0.0999 | 2.17 | 0 | 97.2 | 0 | 0 | 0.93 | 0.93 | 0.193 | 0.2108 | 0.0178 |
| 6gu0 | noschrodinger | 2.5 | 0.0045 | 0.9 | 0 | 96.54 | 0.85 | 0 | 1.85 | 1.17 | 0.1791 | 0.2065 | 0.0274 |
| 6gu2 | opls | 2 | 0.0944 | 2.25 | 0.32 | 97.25 | 3.36 | 3 | 0.77 | 1.28 | 0.2045 | 0.2308 | 0.0263 |
| 6gu2 | noschrodinger | 2 | 0.0056 | 0.98 | 0.48 | 97.11 | 4.93 | 0 | 1.73 | 1.61 | 0.1894 | 0.223 | 0.0336 |
| 6gu3 | opls | 2.65 | 0.0921 | 2.25 | 0.32 | 96.62 | 2.66 | 2 | 0.29 | 1.15 | 0.208 | 0.2412 | 0.0332 |
| 6gu3 | noschrodinger | 2.65 | 0.0027 | 0.8 | 0.64 | 94.87 | 4.94 | 0 | 2.6 | 1.93 | 0.1924 | 0.234 | 0.0416 |
| 6gur | opls | 2.1 | 0.0936 | 2.25 | 0 | 98.72 | 1.09 | 1 | 1.34 | 0.89 | 0.2024 | 0.2274 | 0.025 |
| 6gur | noschrodinger | 2.1 | 0.0027 | 0.75 | 0.18 | 98.55 | 0.86 | 0 | 1.57 | 0.9 | 0.2012 | 0.2309 | 0.0297 |
| 6gzt | opls | 2.1 | 0.0961 | 2.35 | 0 | 97.28 | 0.85 | 0 | 1.58 | 1.04 | 0.185 | 0.2088 | 0.0238 |
| 6gzt | noschrodinger | 2.1 | 0.0027 | 0.77 | 0 | 96.98 | 2.12 | 0 | 2.04 | 1.4 | 0.1849 | 0.2048 | 0.0199 |
| 6h00 | opls | 1.63 | 0.0899 | 2.39 | 0 | 98.19 | 1.17 | 2 | 2.97 | 1.14 | 0.1936 | 0.228 | 0.0344 |
| 6h00 | noschrodinger | 1.63 | 0.0138 | 1.22 | 0.5 | 97 | 2.34 | 0 | 2.71 | 1.51 | 0.1965 | 0.2302 | 0.0337 |
| 6h96 | opls | 1.55 | 0.0986 | 2.45 | 0 | 99.72 | 0.59 | 6 | 4.71 | 1.24 | 0.1972 | 0.2221 | 0.0249 |
| 6h96 | noschrodinger | 1.55 | 0.0045 | 1.06 | 0 | 99.06 | 1.36 | 0 | 4.71 | 1.34 | 0.2015 | 0.2243 | 0.0228 |
| 6hai | opls | 2.2 | 0.0922 | 2.26 | 0 | 99.29 | 1.39 | 2 | 0.98 | 0.9 | 0.2238 | 0.252 | 0.0282 |
| 6hai | noschrodinger | 2.2 | 0.0027 | 0.74 | 0 | 99.06 | 1.65 | 0 | 2.38 | 1.18 | 0.2081 | 0.2475 | 0.0394 |
| 6hdo | opls | 2.61 | 0.1025 | 2.27 | 0 | 100 | 1.67 | 0 | 0.44 | 0.83 | 0.2008 | 0.2283 | 0.0275 |
| 6hdo | noschrodinger | 2.61 | 0.0033 | 0.82 | 0 | 96.88 | 3.31 | 0 | 2.23 | 1.58 | 0.1828 | 0.2197 | 0.0369 |
| 6ho7 | opls | 2.5 | 0.0911 | 2.25 | 0 | 98.94 | 0 | 0 | 0.34 | 0.62 | 0.1814 | 0.2394 | 0.058 |
| 6ho7 | noschrodinger | 2.5 | 0.0029 | 0.74 | 0 | 98.41 | 1.3 | 0 | 0.67 | 0.81 | 0.1954 | 0.2317 | 0.0363 |
| 6hpu | opls | 3.96 | 0.0916 | 2.39 | 0.6 | 93.95 | 6.1 | 2 | 0.61 | 1.7 | 0.2114 | 0.2599 | 0.0485 |
| 6hpu | noschrodinger | 3.96 | 0.0031 | 0.84 | 2.29 | 83.61 | 13.31 | 0 | 3.19 | 2.65 | 0.199 | 0.2357 | 0.0367 |
| 6hpy | opls | 2 | 0.0944 | 2.19 | 0.73 | 96.35 | 0.77 | 0 | 1.23 | 1.09 | 0.2416 | 0.258 | 0.0164 |
| 6hpy | noschrodinger | 2 | 0.003 | 0.81 | 0 | 95.68 | 0 | 0 | 1.65 | 1.21 | 0.2227 | 0.251 | 0.0283 |
| 6hul | opls | 2.5 | 0.0912 | 2.32 | 1.23 | 96.46 | 3.17 | 2 | 1.31 | 1.47 | 0.2262 | 0.246 | 0.0198 |
| 6hul | noschrodinger | 2.5 | 0.0053 | 0.89 | 0.92 | 96.78 | 6.49 | 0 | 4.22 | 2.02 | 0.1959 | 0.2331 | 0.0372 |
| 6ieg | opls | 3.55 | 0.0855 | 2.45 | 1.35 | 94.8 | 11.25 | 28 | 1.88 | 2.11 | 0.2761 | 0.3251 | 0.049 |
| 6ieg | noschrodinger | 3.55 | 0.0043 | 0.97 | 2.5 | 91.64 | 24.66 | 0 | 21.57 | 3.38 | 0.2594 | 0.3078 | 0.0484 |
| 6iq5 | opls | 3.7 | 0.09 | 2.41 | 1.2 | 93.38 | 7.46 | 8 | 0.98 | 1.89 | 0.3498 | 0.3748 | 0.025 |
| 6iq5 | noschrodinger | 3.7 | 0.004 | 0.93 | 1.47 | 86.2 | 11.9 | 0 | 7.74 | 2.88 | 0.3394 | 0.3756 | 0.0362 |

| | | | | | | | | | | | | | |
|---|---|---|---|---|---|---|---|---|---|---|---|---|---|
| 6iq7 | opls | 1.9 | 0.1004 | 2.36 | 0.55 | 98.9 | 0.65 | 3 | 2.03 | 0.97 | 0.1599 | 0.1934 | 0.0335 |
| 6iq7 | noschrodinger | 1.9 | 0.0043 | 0.9 | 0.52 | 98.18 | 2.58 | 0 | 2.21 | 1.31 | 0.165 | 0.1905 | 0.0255 |
| 6mf9 | opls | 2.04 | 0.0915 | 2.31 | 0 | 100 | 0 | 2 | 2.06 | 0.98 | 0.1745 | 0.2142 | 0.0397 |
| 6mf9 | noschrodinger | 2.04 | 0.0058 | 1.2 | 0 | 99.6 | 0 | 0 | 2.79 | 1.07 | 0.179 | 0.2191 | 0.0401 |
| 6mip | opls | 1.79 | 0.1017 | 2.51 | 0 | 97.81 | 0.78 | 2 | 3.06 | 1.14 | 0.1906 | 0.2369 | 0.0463 |
| 6mip | noschrodinger | 1.79 | 0.0048 | 0.94 | 0 | 99.27 | 0.78 | 0 | 1.31 | 0.86 | 0.2 | 0.2332 | 0.0332 |
| 6mn2 | opls | 2.74 | 0.0921 | 2.38 | 0.19 | 97.66 | 1 | 0 | 0.36 | 0.71 | 0.2266 | 0.2576 | 0.031 |
| 6mn2 | noschrodinger | 2.74 | 0.0039 | 1.05 | 0.19 | 96.12 | 2.49 | 0 | 2.92 | 1.65 | 0.2205 | 0.2639 | 0.0434 |
| 6mpt | opls | 1.65 | 0.1008 | 2.28 | 0 | 97.9 | 1.39 | 1 | 1.7 | 1.06 | 0.1926 | 0.2261 | 0.0335 |
| 6mpt | noschrodinger | 1.65 | 0.0045 | 0.87 | 0 | 97.57 | 0.46 | 0 | 2.44 | 1.12 | 0.1914 | 0.2245 | 0.0331 |
| 6mq3 | opls | 3.57 | 0.0901 | 2.22 | 0.05 | 96.43 | 0.48 | 2 | 0.64 | 0.95 | 0.181 | 0.2093 | 0.0283 |
| 6mq3 | noschrodinger | 3.57 | 0.0026 | 0.74 | 0.1 | 94.87 | 0.36 | 0 | 2.66 | 1.41 | 0.161 | 0.1995 | 0.0385 |
| 6msz | opls | 1.97 | 0.0892 | 2.26 | 0 | 95.77 | 1.99 | 1 | 0.88 | 1.29 | 0.2655 | 0.3072 | 0.0417 |
| 6msz | noschrodinger | 1.97 | 0.0026 | 0.76 | 0 | 95.92 | 0.66 | 0 | 1.82 | 1.22 | 0.2658 | 0.2993 | 0.0335 |
| 6mwd | opls | 2.33 | 0.0848 | 2.38 | 0 | 99.43 | 5.08 | 0 | 0.46 | 1.2 | 0.2453 | 0.2613 | 0.016 |
| 6mwd | noschrodinger | 2.33 | 0.0041 | 1.14 | 0 | 97.41 | 8.08 | 0 | 6.64 | 2.17 | 0.233 | 0.2563 | 0.0233 |
| 6mxv | opls | 1.78 | 0.0971 | 2.25 | 0 | 98.08 | 0.71 | 2 | 1.62 | 0.91 | 0.1526 | 0.1931 | 0.0405 |
| 6mxv | noschrodinger | 1.78 | 0.0046 | 0.94 | 0 | 97.99 | 0.92 | 0 | 2.12 | 0.99 | 0.1556 | 0.191 | 0.0354 |
